# The molecular diversity of hippocampal regions and strata at synaptic resolution revealed by integrated transcriptomic and proteomic profiling

**DOI:** 10.1101/2024.08.05.606570

**Authors:** Eva Kaulich, Quinn Waselenchuk, Nicole Fürst, Kristina Desch, Janus Mosbacher, Elena Ciirdaeva, Marcel Juengling, Georgi Tushev, Julian Langer, Erin M. Schuman

**Author notes:** co-first authors (Eva Kaulich, Quinn Waselenchuk).

## Abstract

The molecular diversity of neurons and their synapses underlies the different responses and plasticity profiles that drive all neural circuits and behavior. While the extent of this diversity has been partially revealed by transcriptomic and proteomic profiling, combined studies of neuronal transcripts and proteins are limited. Here, we used microdissection of mouse hippocampal subregions and CA1 strata and fluorescence-activated synaptosome sorting (FASS) to characterize the transcripts and proteins from different hippocampal neurons and their compartments with synaptic resolution. Parallel RNA-seq and LC-MS/MS of microdissections identified over 15,000 mRNA transcripts and 10,000 proteins, revealing thousands with local enrichment such as classes of glutamate receptors and voltage-gated potassium channels, myelin-associated molecules, and adhesion molecules. Synaptosome analysis further identified specific enrichment of molecules from collagen, ribosome, solute carrier, and receptor families at different synapses formed along CA1 neurons. By integrating mRNA and protein data, we defined clusters of co-regulated molecules such as adhesion and neurofilament proteins and transporter mRNAs, and found subsets of mRNA-protein pairs with strong correlation and anti-correlation in their abundance variation. Our findings comprise a rich resource on the molecular landscape of the hippocampus and its synapses that is accessible at <u>syndive.org</u>, and highlight the coordinated organization of transcripts and proteins between regions, neuronal compartments, and synapses.

## MAIN

Transcriptomic and proteomic profiling of brain regions, neuronal populations, and synapse types has revealed the brain’s immense capacity for cellular and subcellular molecular diversity^1–11^. However, comprehensive studies addressing both neuronal transcript and protein diversity in parallel are lacking.

The need to quantify mRNA and protein together is underscored by the consensus that abundance of each molecule is not reliably predicted by the other^12–19^. The complicated relationship between the expression level of an mRNA and the protein it encodes is evident when reconstructing molecular-level networks based on spatial scale –omics data and span from the organismal to the cellular and subcellular levels. Factors that affect the mRNA-protein relationship include mRNA localization to specific sites^20–25^ and mRNA stabilization, such as during fly embryogenesis, where mRNAs in specific regions affect the local pool of protein but do not reflect overall mRNA abundance^13^. Other examples include retinal cone development in *Xenopus*^26–28^ and spatio-temporal regulation of mRNA local dendritic translation during neuronal plasticity, where local mRNAs are translated upon environmental cues and can show high or low correlations with protein abundance depending on the condition^29–31^. A comprehensive multi-omics dataset obtained from the same samples will enable identification of functional units within subcellular compartments, including synapses. Still, such integrated studies remain largely absent.

We set out to systematically characterize the transcripts and proteins that define different neuronal populations, compartments, and synapses to address how both mRNA and proteins give rise to functional diversity in the hippocampus, a brain region where the physiology, cellular composition, connectivity, and function has been extensively studied. Molecular gradients across the hippocampus^6^ and functional differences indicate compartmentalization of these highly organized subregions: the DG (dentate gyrus, with granule cells), and CA1-3 of Ammon’s horn (*Cornu Ammonis*, pyramidal cells). Area CA1 can further be divided into four physiologically distinct strata (*s.oriens, s.pyramidale, s.radiatum and s.lacunosum moleculare*)^32^. Within area CA1, the excitatory pyramidal neurons run orthogonal to the strata with their basal dendrites in *Stratum oriens* (SO), cell bodies in *Stratum pyramidale* (SP), and proximal and distal apical dendrites in *Stratum radiatum* (SR) and *Stratum lacunosum-moleculare* (SLM), respectively^32,33^. The excitatory synaptic inputs and modulatory inputs to each layer are also distinct, including inputs from hippocampal area CA3, entorhinal cortex, the nucleus reunions of the thalamus and other cortical and subcortical areas^34^. Each of the strata also houses a unique combination of interneuron cell bodies and processes^35^. The distinctive nature of the inputs as well as differences in the local excitatory-inhibitory microcircuits gives rise to distinct functional and plasticity properties of these different strata synaptic populations^33,36,37^ (SO^38–43^, SP^33^, SR^44–47^, SLM^48–52^). While the differential distribution of some mRNAs and proteins within the strata has been studied^20,22–24,53–57^, we lack a complete understanding of how their molecular composition gives rise to the well-known differences in synaptic and plasticity properties.

Here, we combined microdissection of mouse hippocampal subregions and four CA1 strata with purification of synapses using fluorescence activated synaptosome sorting (FASS^9^) and downstream transcriptomic and proteomic analyses of tissue and synaptosomes to reveal highly specific and functionally specialized multi-omic profiles, creating a comprehensive map of molecular diversity in the hippocampal formation and the layers of area CA1.

## RESULTS

### A pipeline to analyze subregions, strata and synaptosomes of the mouse hippocampus

We established a dissection pipeline to isolate hippocampal subregions and strata and successfully purified synaptosomes, for the first time, from microdissected strata (Fig. 1a-c, Extended Data Fig. S1). For microdissection of subregions from acute slices, we adapted a previous dissection protocol for DG, CA3 and CA1^20^ to the mouse hippocampus (as they are visually indistinguishable under a dissection microscope, CA2 and CA3 were dissected together). We further microdissected the four CA1 strata, *S. oriens* (SO), *S. pyramidale* (SP), *S. radiatum* (SR) and *S. lacunosum-moleculare* (SLM). For each sample, tissue from two mice was pooled to provide sufficient material for downstream analysis of the transcriptome and proteome from the same source, reducing technical and experimental variability. Next, we conducted Fluorescence-Activated Synaptosome Sorting (FASS), which uses fluorescent labeling of a synaptic protein (Syn-1::tdTomato) from transgenic mice to sort synaptosomes to high purity^2,9,58^. Despite limited starting material, we successfully sorted ∼2 million synaptosomes to high purity and the same number of control particles from each microdissection (Extended Data Fig. S2a, b).

**Fig. 1:**
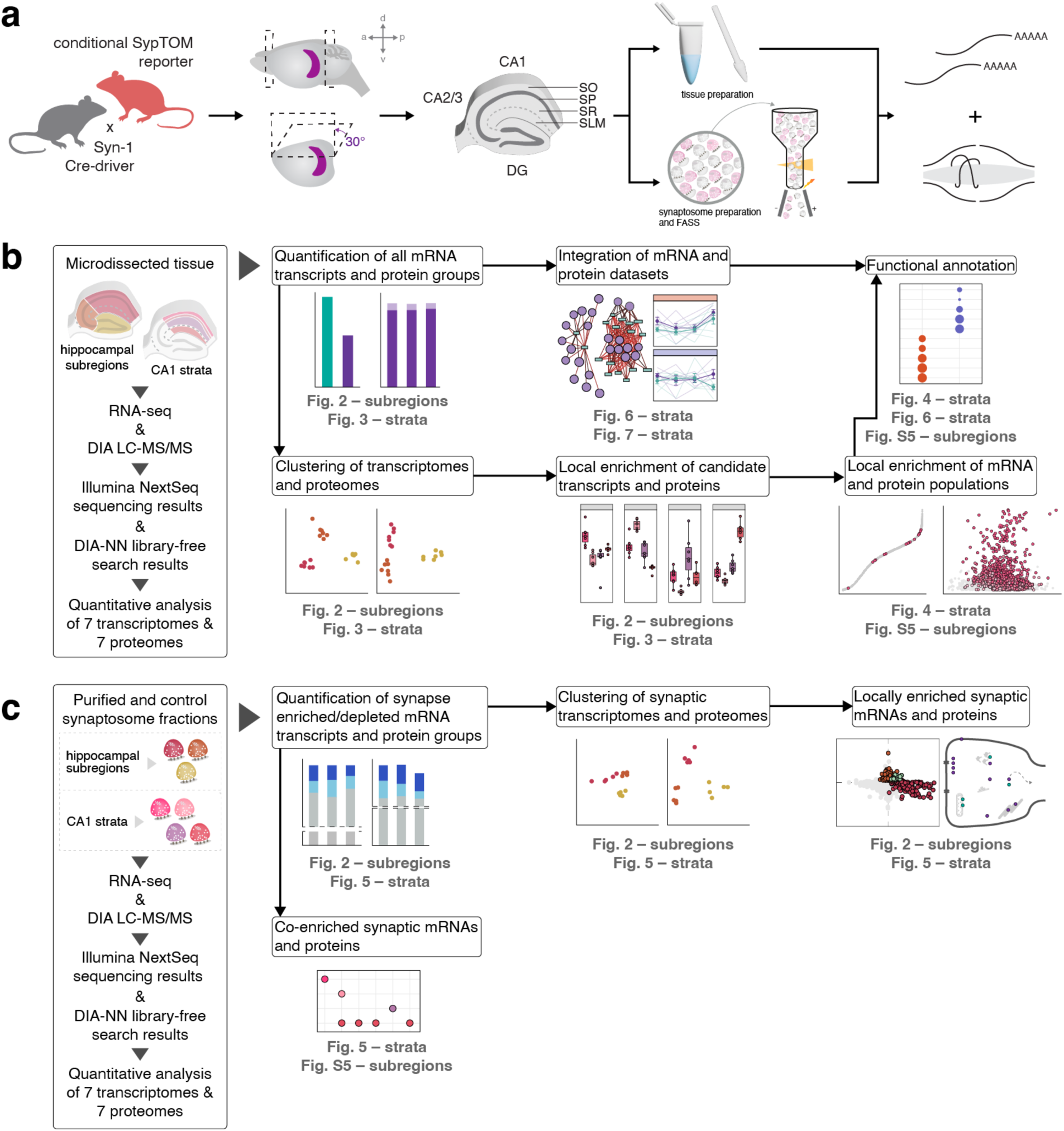
Overall workflow including sample preparation and data analysis. (a) Experimental workflow. A Syn-1 Cre-driver line was crossed with a floxed Synaptophysin-tdTomato (SypTOM) reporter line to label presynaptic terminals. Horizontal brain slices were generated after cutting hemispheres at a 30° angle (see methods) and subregions and CA1 strata were microdissected, and subjected to either tissue preparation or crude synaptosome generation via Percoll-sucrose density gradient followed by fluorescence-activated synaptosome sorting (FASS). Parallel RNA-seq and data independent acquisition (DIA) mass spectrometry were then performed. SO, *Stratum oriens*; SP, *Stratum pyramidale*; SR, *Stratum radiatum*; SLM, *Stratum lacunosum-moleculare*. (b,c) Downstream analysis pipeline of tissue and synaptosome data with relevant figures indicated (full pipeline see Extended data Fig. S1).

### Transcriptomic and proteomic profiling of hippocampal subregions revealed distinct molecular signatures

Using tissue prepared from the microdissected subregions (DG, CA3 and CA1) we split each sample and subjected it to RNA-seq and LC-MS/MS to identify the transcriptome and proteomes, respectively. We achieved deep coverage of subregion transcriptomes and proteomes, quantifying more than 10,000 protein groups across subregions and detecting > 17,000 mRNA transcripts (Fig. 2a, b). While all transcripts and the vast majority of proteins (∼ 97%) were detected in all subregions, we asked if the local populations have different features using an unsupervised Principal Component Analysis (PCA), which revealed clustering of the subregions and indicated that they are molecularly distinct (Fig. 2c). To benchmark our data, we looked for the differential subregion enrichment of previously reported molecular markers (as mRNAs or proteins). For example, we observed the enrichment of *Fibcd1* and *Bok* in areas CA1 and CA2/3, respectively^6,10,59–62^ (Fig. 2d, Extended Data Fig. S3a). At the protein level, we also observed enrichment of Fibcd1 in CA1, while enrichment of Vcan was detected in CA2/3^6,59,63,64^ (Fig. 2e). CA2 markers^65,66^ were also validated in CA2/3 samples (Extended Data Fig. S4b,c). For the DG, we observed enrichment of markers for both inhibitory and excitatory neuronal populations. Within the DG transcriptome, *Prox1*, responsible for subtype-specific development of GABAergic interneurons, was enriched^67,68^ (Fig. 2d, Extended Data Fig. S3a). Calb2, a marker for excitatory cell population of mossy cells at the DG hilus, was enriched at the protein level^6,63,64^ (Fig. 2e). In addition to these candidates, differential expression of thousands of other mRNAs and proteins was detected between subregions (Extended Data Fig. S5a-d and Supplementary Table S1). All of these data (as well as the data described below) can be found on our searchable platform https://syndive.org

**Fig. 2:**
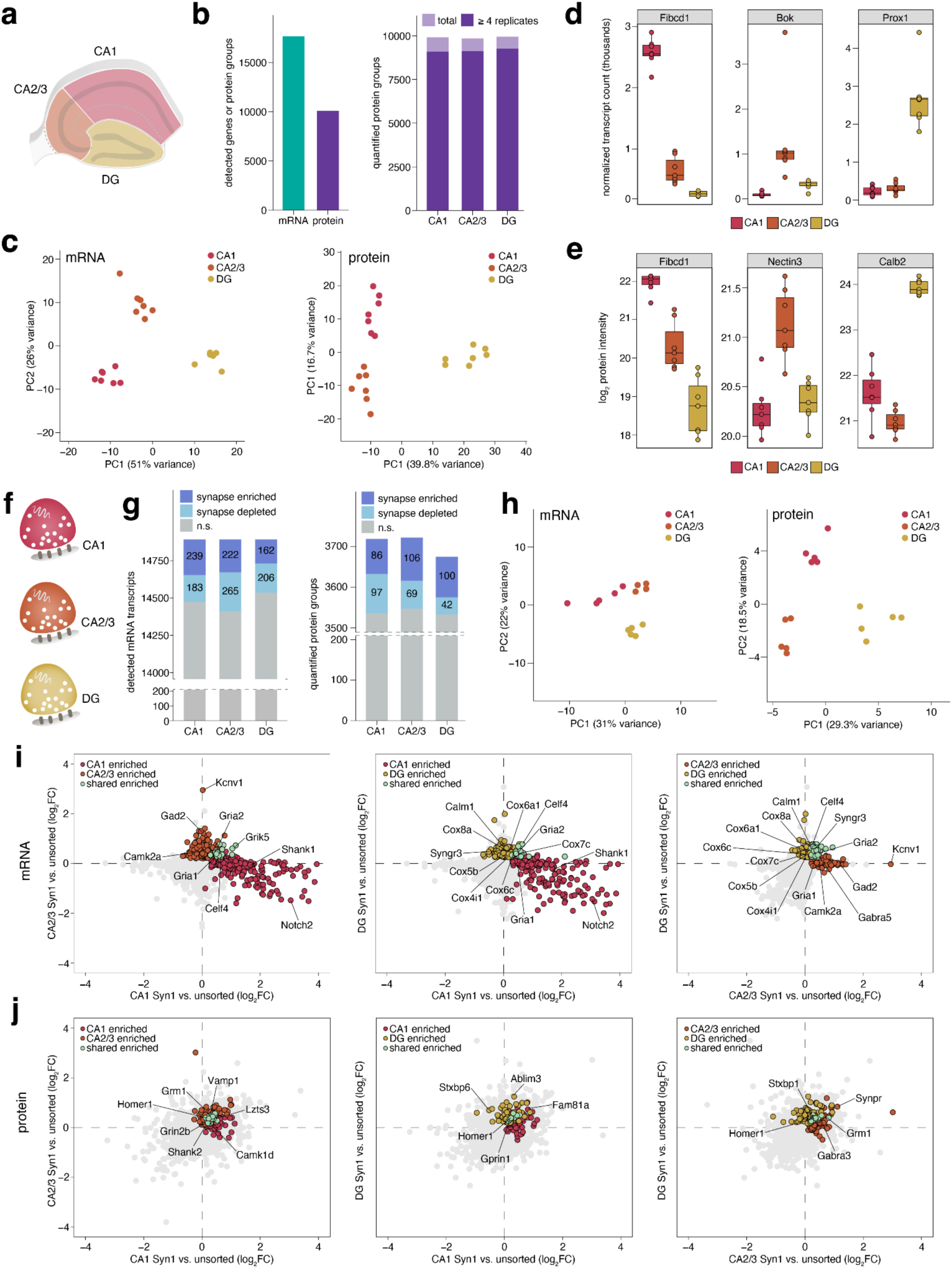
Hippocampal subregion-specific transcriptomic and proteomic differences for tissue and synaptosomes. (a) Schematic indicating subregions CA1, CA2/3 and DG that were microdissected for tissue preparation (b) Left: total number of detected mRNA transcripts and quantified protein groups across all subregions (N = 7 replicates). Right: quantified protein groups per subregion, highlighting proteins detected in ≥ 4 replicates. (c) Principal Component Analysis (PCA) of transcriptome (left) and proteome (right) shows clustering by subregion. Each data point represents one replicate. (d, e) Boxplots showing subregion-specific enrichment of previously described candidates. Plots specify median and 1.5 × IQR. (f) Schematic of synaptosomes from microdissected subregions. (g) Detected transcripts and quantified protein groups in purified synaptosomes for each subregion. Transcripts and proteins that were synapse-enriched, depleted, and non-significant are indicated (N = 5). (h) PCA plot of subregion synapse-enriched and depleted transcripts and proteins. Each data point represents one replicate. (i,j) Correlation plots of Syn-1 enriched fractions vs. controls comparing different subregions at the transcript (i) and protein (j) level. Coloured points represent mRNAs and proteins that were significantly enriched compared to their control fraction (*p* adjusted < 0.1 and < 0.05, respectively), while all other detected mRNAs and proteins are shown in gray.

### Neuronal populations of different subregions shape synaptic diversity in the hippocampus

As indicated above, we detected thousands of transcripts and proteins in individual hippocampal subregions. To understand how molecular differences may underlie differences in physiology and function between the subregions, we zoomed in on the synaptic molecules. To assess potential differences between the synaptic transcriptomes and proteomes, we FASS-purified synapses from microdissected subregions, thus allowing us to examine synaptic diversity. Parallel RNA-seq and LC-MS/MS quantified > 4,000 total protein groups and nearly 15,000 mRNA transcripts across the subregion synapses (Fig. 2f, g). To analyze the molecules enriched in each FASS-purified population, we compared them to the precursor (size-selected control) synaptosome fraction (which also includes contaminants^9^; see methods) and examined which molecules were enriched relative to the controls. Over 550 mRNAs and 250 proteins were synapse-enriched across all regions, while more than 470 mRNAs and 175 proteins were depleted when compared to controls; the enrichment profiles varied by subregion (Fig. 2g). A PCA indicated the distinct clustering of subregion-specific synaptic transcriptomes and proteomes (Fig. 2h).

We next focused on the differentially enriched mRNAs and proteins expressed within the subregion synapses, reasoning that these molecules can give rise to functional specialization. Although nearly all mRNAs and proteins detected in the FASS purified synaptosome fractions were present in all subregions, there were local populations enriched compared to their respective controls. These enriched pre– and postsynaptic mRNAs and proteins generally included receptors, adhesion molecules, extracellular matrix molecules, guidance cues as well as molecules linked to metabolism (Fig. 2i, j, Extended Data Fig. S5a-e). For the synaptic transcriptome, *Gria2* and *Celf4*^69^ transcripts were enriched in the synaptosomes from all subregions. The *Notch2* transcript was uniquely enriched in CA1 synaptosomes,^70,71^, while *Grik5* and *Gria1* transcripts were enriched in both CA1 and CA2/3 synapses. Transcripts for the AMPA receptor auxiliary subunit *Cnih2* and the voltage-gated potassium channel *Kcnv1* (Kv1.1), in contrast, were uniquely enriched at CA2/3 synapses. In DG synaptosomes, we found enrichment of transcripts for several cytochrome c oxidase subunits (*Cox4i1*, *Cox5b*, *Cox6a1*, *Cox6c*, *Cox7c*, *Cox8a*), consistent with previous histochemical evidence of cytochrome oxidase localization in the rat DG, specifically in the dendrites of granule cells molecular layer ^72^.

For the synaptic proteome (Fig. 2j), we detected the enrichment of the glutamate receptors Grin2b in CA1 and CA2/3 pyramidal neurons, and Grm1 uniquely in CA2/3, as previously described^73^. Shared-enriched proteins across all subregions included postsynaptic density scaffold proteins Homer1 and Lzts3 along with Fam81a, and Ablim3. Enriched proteins associated with the presynaptic compartment included Synpr in CA2/3, and Rims2 in both DG and CA2/3. Combined, RNA-seq and LC-MS/MS data of distinct synaptic populations from microdissected hippocampal subregions and identified shared and unique synaptically enriched transcripts and proteins.

### Specialized strata-specific transcriptomes and proteomes reveal distinct molecular signatures and functional roles

We set out next to identify the mRNAs and proteins from the microdissected layers (strata) associated with different synaptic inputs that converge on a single hippocampal cell type, the CA1 pyramidal neuron. We dissected the four strata of CA1, representing the pyramidal cell somata (SP), basal dendrites (SO), apical dendrites (SR), and distal tufts (SLM) and analyzed both the tissue and purified synapses from each strata. Beginning with the analysis of microdissected tissue (Fig. 3, 4), we quantified nearly 10,000 protein groups and over 15,000 mRNA transcripts (Fig. 3a, b) and discovered that these strata have distinct transcriptomes and proteomes, evident as clusters in a PCA (also demonstrating the reproducibility of our dissections) (Fig. 3c). Summing across the CA1 strata, we found thousands of transcripts and proteins were significantly enriched, highlighting previously undescribed expression patterns and reflecting their specific synaptic inputs and functions (Fig. 4a, c; cross referenced with the Allen Mouse Brain Atlas^1^ in Extended Data Fig. S3b). Among the significantly enriched transcripts and proteins we observed a broad distribution of low-to-high abundance mRNAs and proteins, rather than a bias towards lowly– or highly-expressed gene products. Subsequent Gene Ontology (GO) overrepresentation analysis revealed that these enriched molecules contribute to the cellular and functional specialization of each stratum (Fig. 4b, d and Table 1), as indicated in greater detail below.

**Fig. 3:**
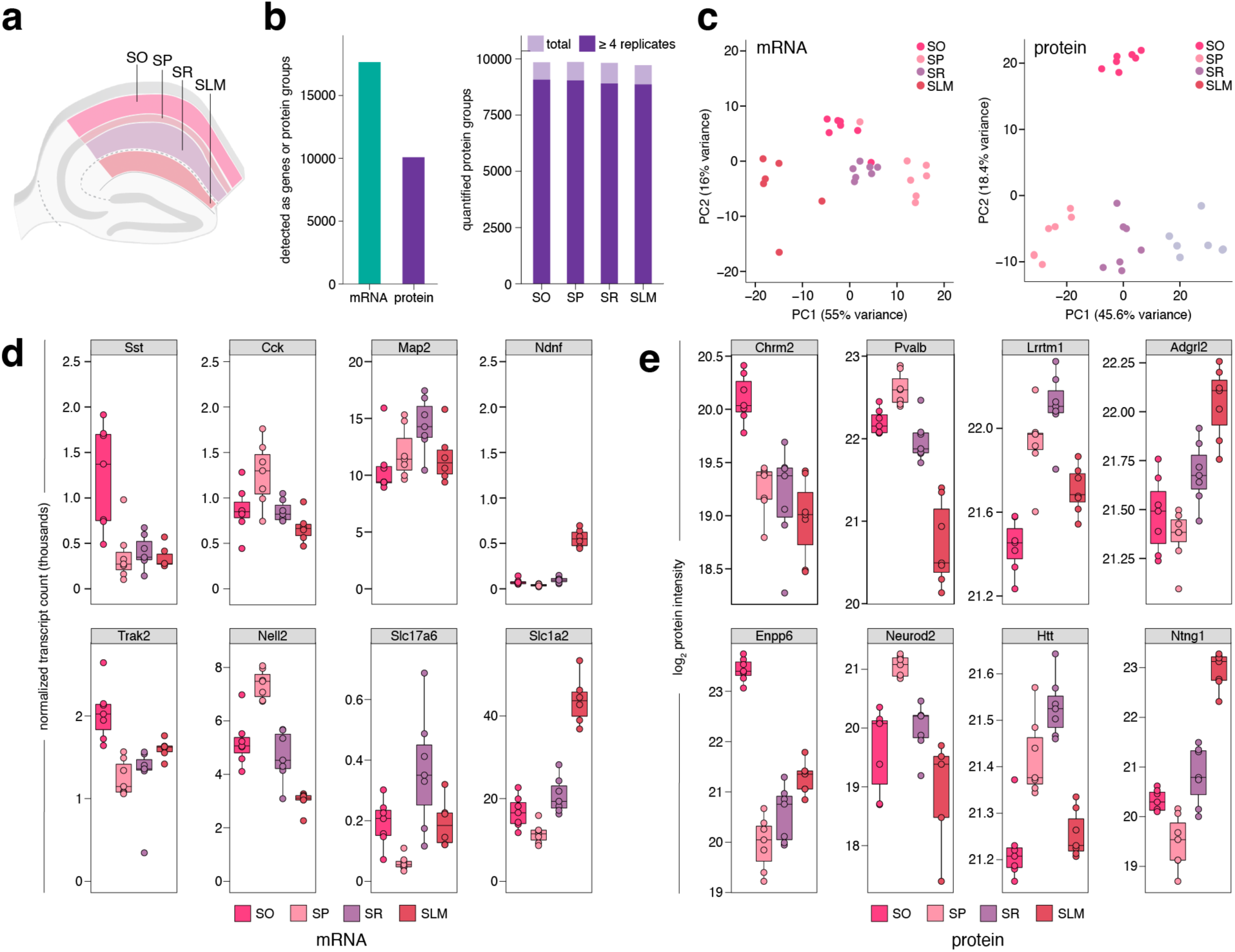
Transcriptomic and proteomic profiles of CA1 strata tissue. (a) Schematic indicating CA1 strata SO, SP, SR and SLM that were microdissected for tissue preparation (b) Left: total detected mRNA transcripts and quantified protein groups across all strata (N = 7 replicates). Right: quantified protein groups per stratum, highlighting proteins detected in ≥ 4 replicates. (c) PCA of transcriptomes (left) and proteomes (right) illustrating separation of strata. Each data point represents one replicate. (d,e) Boxplots showing strata-specific enrichment of previously described (top row) and undescribed (bottom row) candidates. Plots specify median and 1.5 × IQR.

**Fig. 4:**
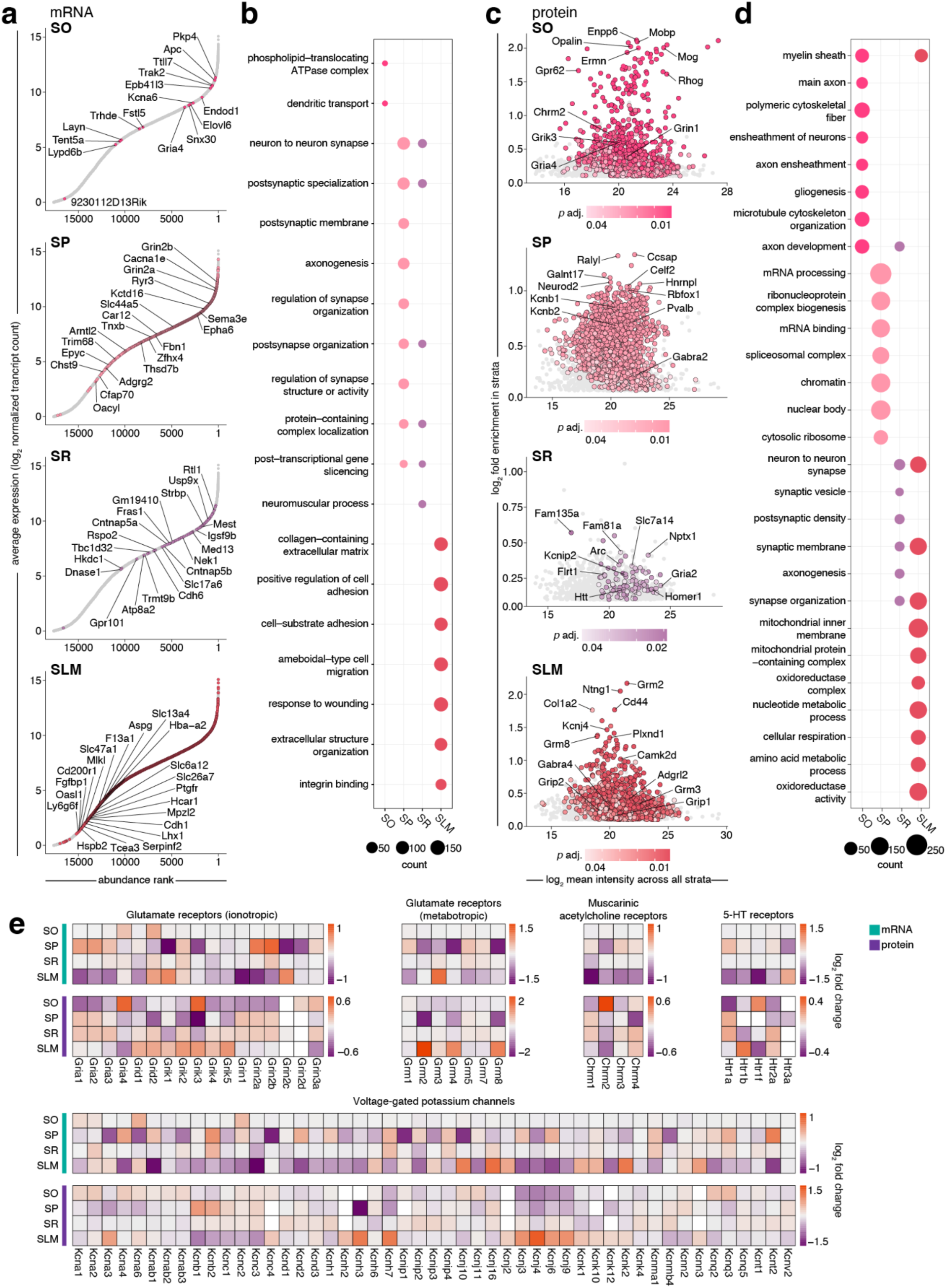
Local transcriptomes and proteomes of CA1 strata (tissue) shape functional specialization. (a) Rank abundance plots of mRNA indicating the relative abundance of transcripts in given strata compared to whole CA1. Transcripts with the highest log_2_ fold-change in a stratum compared to whole CA1 are ranked as 1. Coloured points represent transcripts that showed unique significant enrichment in a stratum (*p* adjusted < 0.1), while all other detected transcripts are shown in gray. The transcripts showing the top 20 highest relative enrichment (uniquely enriched transcripts were ordered by highest log_2_ fold-change) are labeled. (b) Gene ontology (GO) overrepresentation analysis showing top terms (*p* adjusted < 0.01) for significantly enriched transcripts from each strata. (c) Plots showing log_2_ fold enrichment of proteins in a given strata compared to their mean log_2_ intensity across all strata. Significantly enriched proteins are coloured dots (*p* adjusted < 0.05) while all other proteins showing enrichment in the strata are gray. (d) Gene ontology (GO) overrepresentation analysis showing top terms (*p* adjusted < 0.01) for significantly enriched proteins from each strata. (e) Heatmaps of log_2_ fold enrichment or de-enrichment of transcripts and proteins from selected receptor and ion channel classes. Missing values are shown in white.

Zooming in further, the combination of strata microdissection with downstream FASS allowed us to quantify the local transcriptomes and proteomes of synapses formed at different positions along the same neuronal population (CA1 pyramidal neurons) with different presynaptic origins as well as different resident interneurons (Fig.5). In total (considering the union of all synaptosomes across strata), we detected and quantified over 15,000 mRNA transcripts and > 4,000 protein groups, with > 530 mRNAs and > 550 proteins showing synaptic enrichment when compared to controls (Fig. 5a, b). Are the mRNAs and proteins that inhabit the individual layers distinct? Indeed, we found that a PCA revealed the distinct clustering of synaptic transcripts and proteins (Fig. 5c). Thus, despite their physical proximity, we found that subsets of mRNAs and proteins were uniquely enriched at synapses in each stratum, presumably reflecting the distinct functional needs of the synapses in these layers. For example, as indicated below, the differential localization of synaptic mRNAs for many collagens, kinesins, solute carriers and ribosomal proteins between synapses were detected, while many differentially expressed proteins were presynaptic molecules like synaptotagmins as well as pre– and postsynaptic ion channels and receptors.

**Fig. 5:**
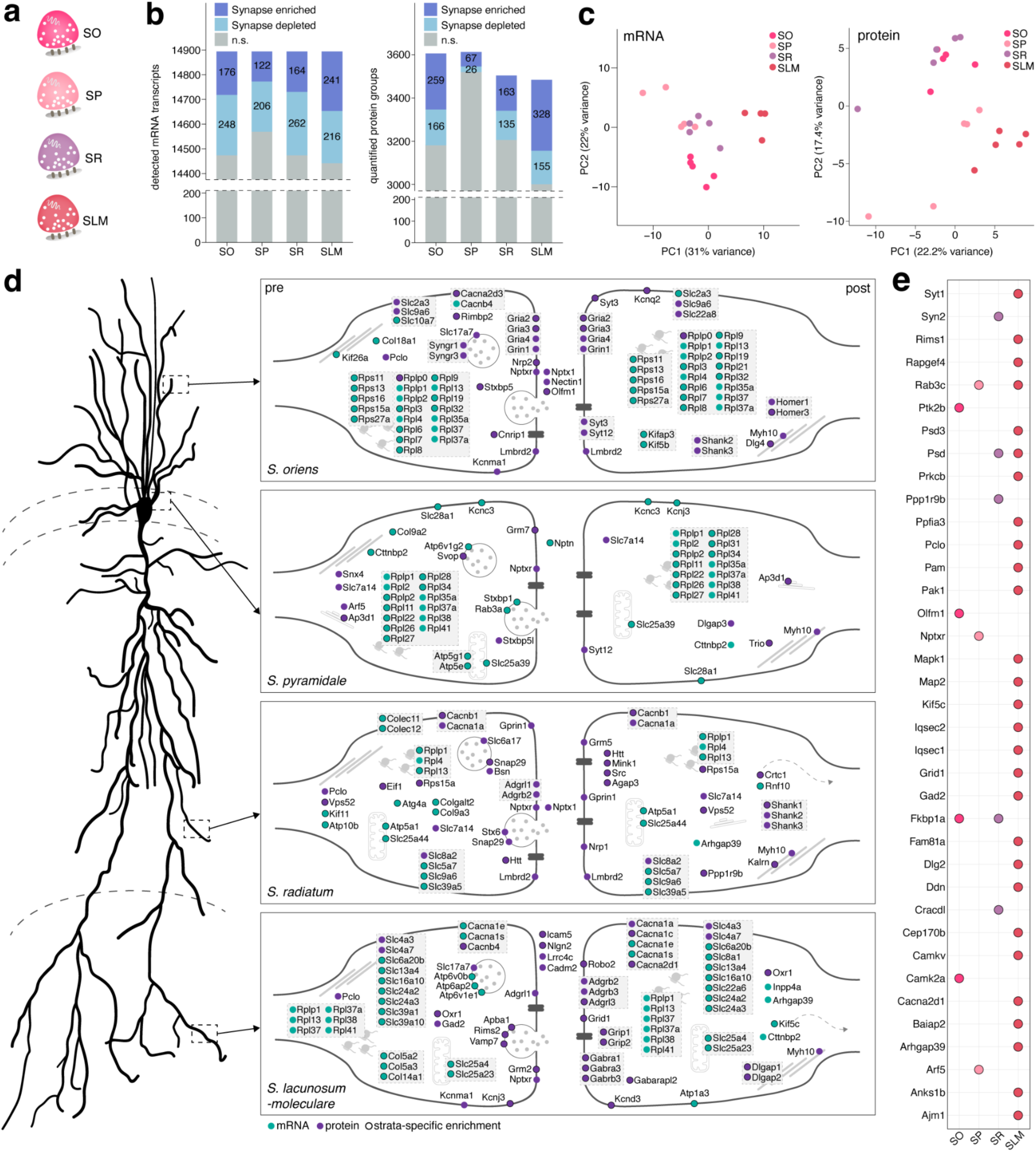
Synaptic transcriptomes and proteomes of CA1 strata show distinct molecular signatures. (a) Schematic of synaptosomes from SO, SP, SR and SLM. (b) Detected transcripts and quantified proteins in purified synaptosomes for each stratum. Transcripts and proteins that were synapse-enriched, depleted, and non-significant are indicated (N = 5). (c) PCA plot of strata synapse-enriched and depleted transcripts and proteins. Each data point represents one replicate. (d) Schematic depicting example synapses (right) along a CA1 pyramidal neuron (left). Shown are a curated selection of synaptic transcripts (cyan) and proteins (purple) that were synapse-enriched in the different CA1 strata. Pre– and/or postsynaptic localization was determined by GO annotation, where applicable. Molecules with similar functions are grouped together. (e) Dot plot of genes that showed co-enrichment of mRNA and the respective protein in each strata.

### Enriched mRNAs and proteins in *S. oriens*

In SO tissue, we found an enrichment of mRNA for *Sst*^74^ and the Chrm2^74^ protein, which are markers for two prominent interneuron populations that reside in this layer (top row, Fig. 3d,e). Additionally, we identified the enrichment for mRNAs of *Trak2*^75,76^, a regulator of mitochondrial mobility in dendrites and axons within the hippocampus, *Pkp4*^77^, a regulator of adhesion and cytoskeletal remodeling, and *Kcna6*^46,78^, a potassium channel that modulates pyramidal cell activity through tonic inhibition (Fig. 3d and Extended Data Fig. S3b and Fig. S6a). Other enriched mRNAs included those encoding proteins involved in cellular transport such as the P-type ATPases *Atp8a2, Atp11b,* and *Atp8a1*, as well as the kinesin motor proteins *Kif5a* and *Kif5b*. Among the previously undescribed but highly enriched proteins in SO, we observed multiple oligodendrocyte-specific proteins such as Enpp6, Plp1, and Mag (Fig. 3e and Extended Data Fig. S6a). Mobp, Cnp, Cldn11, Mog, Ermn, Mbp, and Pllp, which collectively contribute to the formation and maintenance of the myelin sheath^79,80^, were similarly enriched. These findings accordingly overrepresented GO terms associated with axon and myelin (Fig. 4d), consistent with the large population of traversing pyramidal cell myelinated axons that project along the alveus.

Focussing on the FASS-purified SO synaptosomes, we found many transcripts with roles in synaptic targeting, maintenance, and modulation^81^. This included *Col18a1*, *Kif26a*, *Kifap3*, *Kif5b* and *Slc10a7* and *Slc22a8* mRNAs, which were uniquely enriched in SO (Fig. 5d). Other uniquely synapse-enriched mRNAs in SO encoded ribosomal proteins (*Rpl3, Rpl6, Rpl7, Rpl8, Rpl9, Rpl19, Rpl21, Rpl32, Rpl37a*). In contrast, amongst the RPs, only Rplp0 was uniquely synapse-enriched at the protein level, which might reflect a role in onsite and on-demand repair or specialization of the translational machinery^82^. Other enriched proteins included the glutamate receptors Grin1, Gria2, Gria3, and Gria4, as well as the voltage gated channels Kcnq2 and Cacna2d3 (Fig. 5d). Neuronal pentraxin family members Nptxr and Nptx1, expressed at glutamatergic synapses onto interneurons, were also enriched here where they are responsible for Gria4 clustering ^83^.

### Enriched mRNAs and proteins in *S. pyramidale*

In SP tissue, which contains pyramidal cell bodies, we observed enrichment of *Cck* transcripts and Pvalb proteins, identifying the two basket cell populations that form perisomatic connections with pyramidal cells^74,84–88^ (Fig. 3d, e, Extended Data Fig. S3b). Previously undescribed enriched mRNAs included *Nell2*^89,90^, which is involved in neuronal polarization and axon growth, *Ankrd46*, associated with membrane interactions, and the uncharacterized *Vnx*^91,92^ (Fig. 3d, Extended Data Fig. S3b, Extended Data Fig. S6b). GO term analysis revealed an overrepresentation of mRNAs related to neuronal and synaptic membrane organization, including various receptors and ion channels (Fig. 4b). This aligns with previous observations that these proteins are synthesized in cell bodies and then trafficked to distal neuronal compartments^24^. mRNAs associated with axonogenesis were also enriched (Fig. 4b) consistent with presence of the axon initial segment in SP. Among the largely-nuclear proteins enriched in SP, those with undescribed enrichment included Neurod2 and Neurod6, members of the neuronal basic helix-loop-helix (bHLH) transcription factor family, fitting with their roles in brain development and connectivity^93–96^ (Fig. 3e, Extended Data Fig. S6b). Generally, enriched proteins in SP were predominantly associated with nuclear processes and gene regulation, such as chromatin binding and mRNA processing (Fig. 4d).

At SP-purified synapses, a particular collagen mRNA, *Col9a2*, was specifically enriched. Among other transcripts only synapse-enriched in SP were the mRNA mitochondrial transporters *Slc25a39* and *Slc28a1,* involved in the transport of nucleosides, as well as ribosomal protein mRNAs *Rpl2, Rpl11, Rpl22, Rpl26, Rpl27, Rpl28, Rpl31* and *Rpl34* (Fig. 5b). Proteins synapse-enriched in SP included Grm7, involved in glutamatergic and GABAergic transmission in terminals^97^, Dlgap3 involved in anchoring glutamate receptors at the postsynapse, and Trio, involved in receptor endocytosis regulation (Fig. 5d). Many other molecules enriched in SP synaptosomes are involved in vesicle trafficking, such as *Srf5*, Syt12, Stxbp1, Stxbp5l (Fig. 5d).

### Enriched mRNAs and proteins in *S. radiatum*

SR comprises the largest fraction of neuropil in area CA1, consisting of mostly apical dendrites. Consistent with this, from tissue, we detected enrichment of transcripts for the dendritic marker and microtubule-binding protein *Map2*^21,98^ (Fig. 3c, Extended Data Fig. S3b). mRNAs in SR with previously undescribed enrichment included vGlut2 transporter *Slc17a6, Ttc7b*, and the orphan receptor *Gpr101* (Fig. 3d, Extended Data Fig. S3b, Extended Data Fig. S6c). CA1 pyramidal cells also receive Schaffer collateral inputs from CA3 axons in SR, forming excitatory synapses (for a review see^99^). GO terms for enriched transcripts accordingly included postsynaptic specialization and organization. At the protein level, we detected enrichment of postsynaptic cell adhesion molecule Lrrtm1^100,101^ as described, which controls dendritic spine density and excitatory synapse morphology (Fig. 3d). Additional, previously undescribed SR-enriched proteins were similarly linked to synaptic transmission and plasticity in this area, such as the secreted proteins Nptx1 and Olfm1 (Extended Data Fig. S6c). Htt, responsible for Huntington’s disease when mutated, was also uniquely enriched in SR (Fig. 3e). Similar to mRNA, enriched proteins overrepresented GO terms linked to synaptic specialization, such as synaptic vesicle, and postsynaptic density (Fig. 4d, Table 1). Other proteins contributing to this included, for example, the synaptic vesicle proteins Syt12, Rph3a, Stxbp5, Unc13a, and Slc17a7, the excitatory postsynaptic proteins Rph3a, Homer1, Homer2 and Gria2, and ligands and receptors from the guidance cue Ephrin-family (Efnb2, Epha6, Ephb2).

SR-specific transcripts from synaptosomes included another member of the mitochondrial transporters *Slc25a44*, along with diverse solute carriers *Slc9a6*, *Slc39a5*, and *Slc5a7.* Transcripts for the kinesin *Kif11* and collagens *Col9a3*, *Colgalt2*, *Colec11* and *Colec12* were also uniquely enriched at SR synapses (Fig. 5b). Postsynaptic density mRNAs *Psd* and *Psd3* were also enriched (Fig. 5b). Proteins enriched specifically in SR synaptosomes included many involved in regulation of receptor levels (Mink1, Src, Agap3, also Nptxr). The receptor Grm5 was also enriched along with the voltage-gated calcium subunit Cacna1a and calcium-channel regulatory unit Cacnb1, likely modulating neuronal activity directly. We also detected enrichment of RhoGEF Kinase Kalrn, along with the adhesion molecules Adgrl1 and Adgrb1, all of which are implicated in spine development..

### Enriched mRNAs and proteins in *S. lacunosum-moleculare*

The SLM comprises synapses receiving inputs from layer III of the entorhinal cortex, the nucleus reuniens, and the inferotemporal cortex^102–104^. From SLM tissue, we detected an enrichment of the neuronal *Ndnf* transcript, a marker for neurogliaform neurons known to form dense arborizations within this layer^105–107^. Additionally, we identified several new SLM-enriched mRNAs, including the disease-associated guanine nucleotide exchange factor *Dennd5a*^108^. SLM also houses distinct astrocyte subpopulations^109^. Notably, we found enrichment of the astrocytic glutamate transporter *Slc1a2*^110^ and the glia marker *Atp1b2*^111^ (Fig. 3d, Extended Data Fig. S3b, Extended Data Fig. S6d). GO terms for enriched transcripts were accordingly associated with cell adhesion and extracellular organization (Fig. 4b, Table 1). These mRNAs further included semaphorins (*Sema3g, Sema5a, Sema6a, Sema6d, Sema3c*), ephrins (*Ephb4, Ephb1*) and extracellular matrix modifying metallo-proteases such as *Timp3* and members of the *ADAM* family, all mainly enriched in SLM (Fig. 4d, Table 1). At the proteome level, we found enrichment of the protein Adgrl2^112–114^, which is involved in synaptic connectivity between the entorhinal cortex and dendritic spines in SLM (Fig. 3e). Strikingly, we also detected enrichment of many proteins known to be present in thalamocortical and/or entorhino-hippocampal neurons that project to this stratum, as well as connections between the hippocampus and retrosplenial cortex^74,115,116^. These proteins included Ntng1, Flrt3, Slitrk1/3/4, and various LRR proteins such as Lrrtm4, Lrrc24, Lrrc75a, Lrrc73, Lrrc40, and Lrrc4b, some of which^100^ are known to play key roles in organizing synaptic connections (Fig. 4c, Extended Data Fig. S6d). While many other proteins with high fold-enrichment in this stratum were metabotropic glutamate receptors (Figs. 4c,e, Table 1), the overall population of enriched proteins showed significant overrepresentation of terms linked to cellular metabolism (Fig. 4d), likely driven by processes like plasticity^48–51,117^.

Synaptosome-enriched transcripts specific to the distal dendrites in SLM included those for kinesins and collagens (*Kif5c, Col5a2, Col5a3, Col14a1*) and many members of the solute carrier family (*Slc25a23, Slc39a1, Slc13a4, Slc16a10, Slc22a3, Slc22a6, Slc24a2, Slc24a3, Slc25a4, Slc39a10, Slc8a1, Slc6a20b*). For synapse-enriched proteins, adhesion molecules were detected such as Lrrc4c, Cadm2, Adgrl1, Adgrl3, Adgrb2 and Adgrb3, perhaps representative of the specific inputs to this stratum. Other enriched proteins included voltage-gated calcium channel auxiliary proteins Cacna2d1 and Cacnb4 together with the primary subunit Cacna1a, and the transcripts for *Cacna1c*, *Cacna1s*, and *Cacna1e*. Glutamate receptors Grid1 and Grm2 were also enriched, as well as interactors Grip1 and Grip2 and the calcium-activated potassium channel subunit Kcnma1. Our detected enrichment of GABAA receptor subunits, along with receptor-associated Gabrapl2 and Gad2, in SLM fits well with the known strong synaptic inhibition^50,51^ and GABAergic interneuron population^118^.

### Locally enriched functional modules and potential hotspots for local translation

Our above analyses revealed functional specialization of the transcriptome and proteome in different CA1 strata occupied by different neuronal compartments. To further explore how enriched mRNAs and proteins can give rise to the unique functions of the different strata, we examined the reciprocal enrichment of transcripts and proteins across strata for some receptor and voltage-gated channel classes (Fig. 4e). These proteins are often synthesized in pyramidal cell bodies^24^ and most associated transcripts were accordingly enriched in SP. Exceptions, however, included mRNAs for *Grid2*, *Grik1*, *Grm3*, and *Kcnj10*, all of which were de-enriched in SP and enriched in SO and/or SLM (Fig. 4e), which could be locally synthesized. Another transcript, *Htr3a*, was enriched in SLM (Fig. 4e), consistent with the presynaptic input from the entorhinal cortex^119^ or expression in OLM (oriens-lacunosum-moleculare) interneuron axonal projections to the SLM^120^. Differentially localized protein modules included all detected delta and kainate-type glutamate receptors that were enriched in SLM together with metabotropic receptors Grm2, 4, and 8, whereas most detected AMPA and NMDA receptor types were SR-enriched (Fig. 4e). Gria4 and Chrm2 were highly enriched in SO and de-enriched in all other strata, while Htr1b was enriched in SLM and de-enriched in neighboring SR (Fig. 4e). Htr2a was enriched in both SR and SLM, consistent with its interneuron expression^121^. The G protein-activated inwardly rectifying potassium channels Kcnj3, 4, 6, and 9 were enriched in SLM and de-enriched in both SP and SO (Fig. 4e), while the voltage-gated delayed rectifiers Kcnb1 and 2 were opposingly enriched in SP and de-enriched in SLM (Fig. 4e), matching their somatic and proximal dendritic localization^122^. Identification of gradients of transcripts between the different strata and specific ion channels and receptors fits well with previous evidence that glutamate, potassium, and calcium channels exhibit pronounced gradients along CA1 dendrites^54–56,123,124^ as well as general molecular gradients throughout the hippocampus ^6^.

As we found highly organized regulation of expression between the strata in tissue, we sought to investigate a role for local translation, asking which mRNAs and proteins were co-enriched in purified synaptosome fractions. Using stringent thresholds, we found 37 pairs of locally co-enriched candidates across all strata, including presynaptic (Rims1, Pyk2, Syn2, Synt1, Gad2 and Pclo) and postsynaptic (Psd, Psd3, Grid1 and Nptxr) mRNAs and proteins (Fig. 5e). Furthermore, 28 of 37 co-enriched pairs were detected in SLM, suggesting a relatively high abundance of candidates that may be translated in the most distal dendrites.

### Identification of multi-omics signatures of strata tissue

Our previous analysis demonstrated that various synaptic, neuronal and structural molecules govern the unique organization of the hippocampal strata. To define the multi-omic signatures of CA1 strata, where the transcriptome and proteome interact to govern functional dynamics, morphology, and connectivity, we used the integrative DIABLO model^125–127^. This data-driven approach analyzes the transcriptome and proteome simultaneously to identify correlated patterns and clusters of features that distinguish the CA1 strata. From an initial sparse Partial Least Squares (sPLS) analysis, multiple groups of mRNAs and proteins emerged as defining features of the strata when both datasets were combined (Fig. 6a, b, Extended Data Fig. S7a-d). Using the DIABLO model we then determined latent components that maximize the covariance between the datasets, thus identifying the most relevant features that best describe the differences between the strata (Extended Data Fig. S7a, d). This revealed clusters of highly correlated (*r* > 0.9) mRNAs and proteins that may be co-regulated, from which we extracted two separate networks of correlated features between each omic dataset (Fig. 6b,c, Extended Data Fig. S7c, d). Most molecules in the first network (Fig. 6c) were related to cytoskeletal organization and transport, such as adhesion proteins (Nfasc, Jam3), neurofilament proteins (Nefl, Nefm), and several myelin-associated proteins (Car2, Opalin, Pllp, Plp1, Cd9, Cldn11, Cnp, Ermn, Rhog). These proteins support neuronal structure and cell interactions crucial for myelin sheath formation and cytoskeletal maintenance and are mainly enriched in SO (Fig. 6b). Core correlations in this network centered around *Trak2* mRNA (positively connected) and *Pkp4* mRNA (negatively connected), which encode proteins involved in intracellular transport^75,128^. The second network identified proteins and mRNAs related to glutamatergic neurotransmission and neuronal connectivity, such as glutamate receptors (Grm2, Grm4), synaptic proteins (Il1rap1), transporter mRNAs (*Atp1a2, Slc1a3, Slc1a2*), and guidance cues (*Flt1*, *Sema3c*, Ntng1 and its ligand Lrrc4c) all involved in patterning thalamic connections to the hippocampus^129^. This cluster indicates a role in neuronal signaling and connectivity and the molecules are mainly enriched in SLM (Fig. 6b). Integration of both omic datasets thus revealed distinct molecular signatures of different strata and defined two networks with separate functional roles. The polarized structure and compartmentalization of the pyramidal neurons might explain the observed separation into two networks that primarily feature molecules associated with enrichment in basal dendrites (SO) or distal tufts (SLM).

**Fig. 6:**
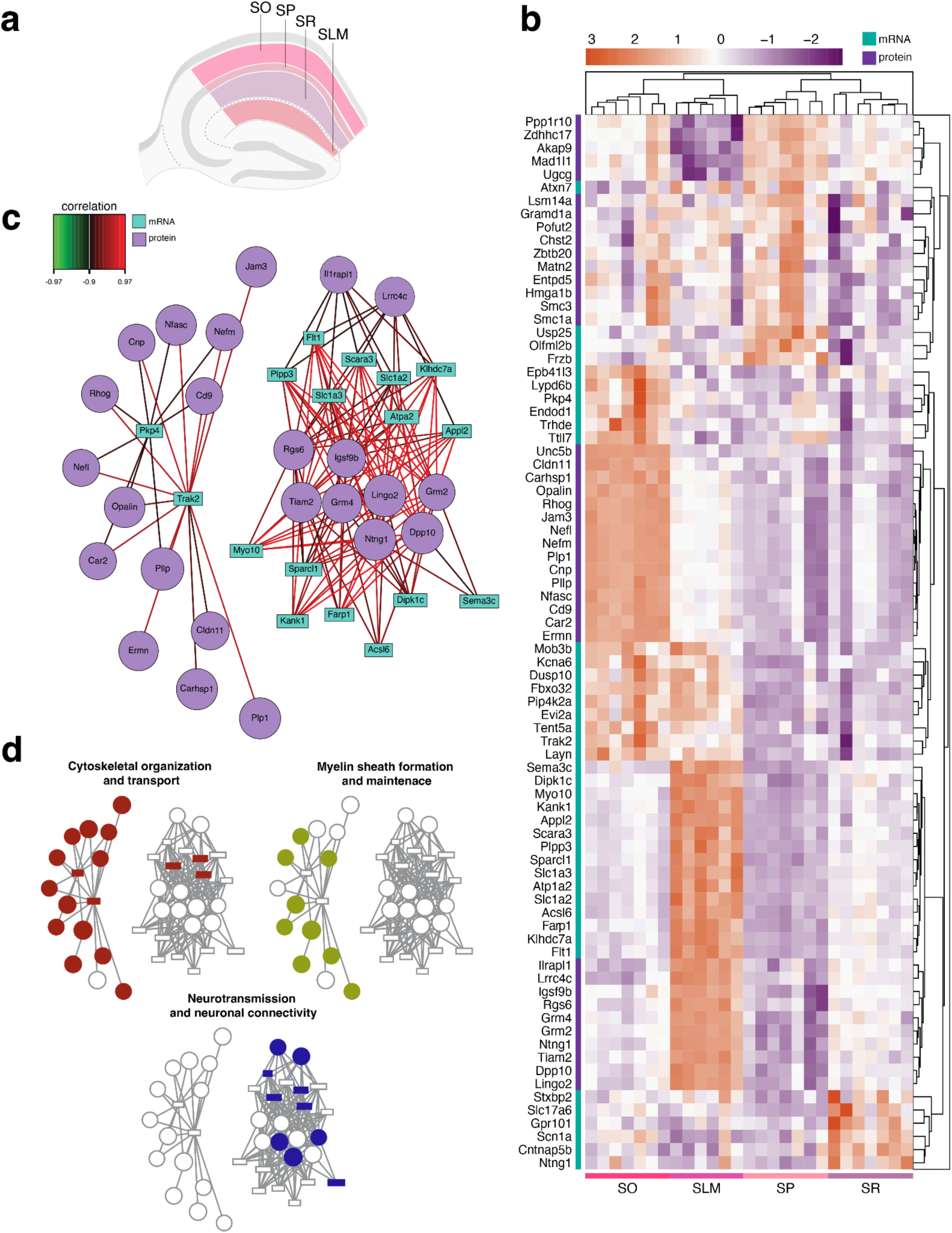
Multi-omics signatures of strata using DIABLO. (a) Clustered Image Map (Euclidean distance, Complete linkage) of highly correlated features from proteomic and transcriptomic Sparse Partial Least Squares (sPLS) analysis. Features show distinct signatures of strata across replicates, with samples grouped based on their similarity in a multi-omics dataset. (b) Relevance associations network plots of highly correlated variables (*r >* 0.9) where edges represent correlations in the data. Nodes represent individual transcripts (cyan squares) and proteins (purple circles). (c) Manually assigned functional annotations of transcripts and proteins.

### Direct comparison of mRNA and protein unveils subcellular relationship patterns and dynamics

By directly comparing mRNA-protein pairs, we asked whether mRNA concentrations are reflected at the protein level, and whether this varied by strata. In line with previous reports^15,53^, we observed a modest correlation of mRNA and protein abundances across all strata (*r* = 0.35; Fig. 7a). We then correlated changes in relative mRNA-protein abundances across strata to deduce the extent to which variation in protein levels between strata might be due to variations in the corresponding mRNA (Fig. 7b). This revealed subsets of hundreds of highly correlated (*r* ≥ 0.9) and anti-correlated (*r* ≤ –0.9) pairs. Highly correlated examples included the synaptic cytoplasmic protein Palm, guidance cue Slit1, and axonal protein Cntn2. Abundance trends for these candidate mRNAs and proteins also correspond to previous findings that they have increased translation efficiency (high footprint-to-mRNA ratio) in either neurite (Cntn2) or soma (Slit1) layers^24^. Anti-correlated pairs included the proteasome subunit Psma1, ribosomal protein Rpl26, and mitochondrial membrane protein Vdac1, none of which are reported to have significant differences in translation between soma and neurite layers^24^ (Fig. 7b). Subsequent GO overrepresentation analysis yielded terms related to neuronal spines, axonogenesis, and adhesion for highly correlated pairs, whereas terms related to the ribosome, proteasome, myelin, and GABAergic synapses were associated with anti-correlated pairs (Fig. 7c, Table 2,3). Previous findings have shown that dendrite-associated proteins have shorter half-lives compared to those associated with mitochondrial or ribosomal GO terms or with the proteasome^130–133^. Short-lived proteins involved in dynamic processes like synaptic plasticity might show high correlation with their mRNA due to rapid turnover and the need for local synthesis, whereas longer-lived mitochondrial and ribosomal proteins may have weaker mRNA-protein correlations due to their stability.

**Fig. 7:**
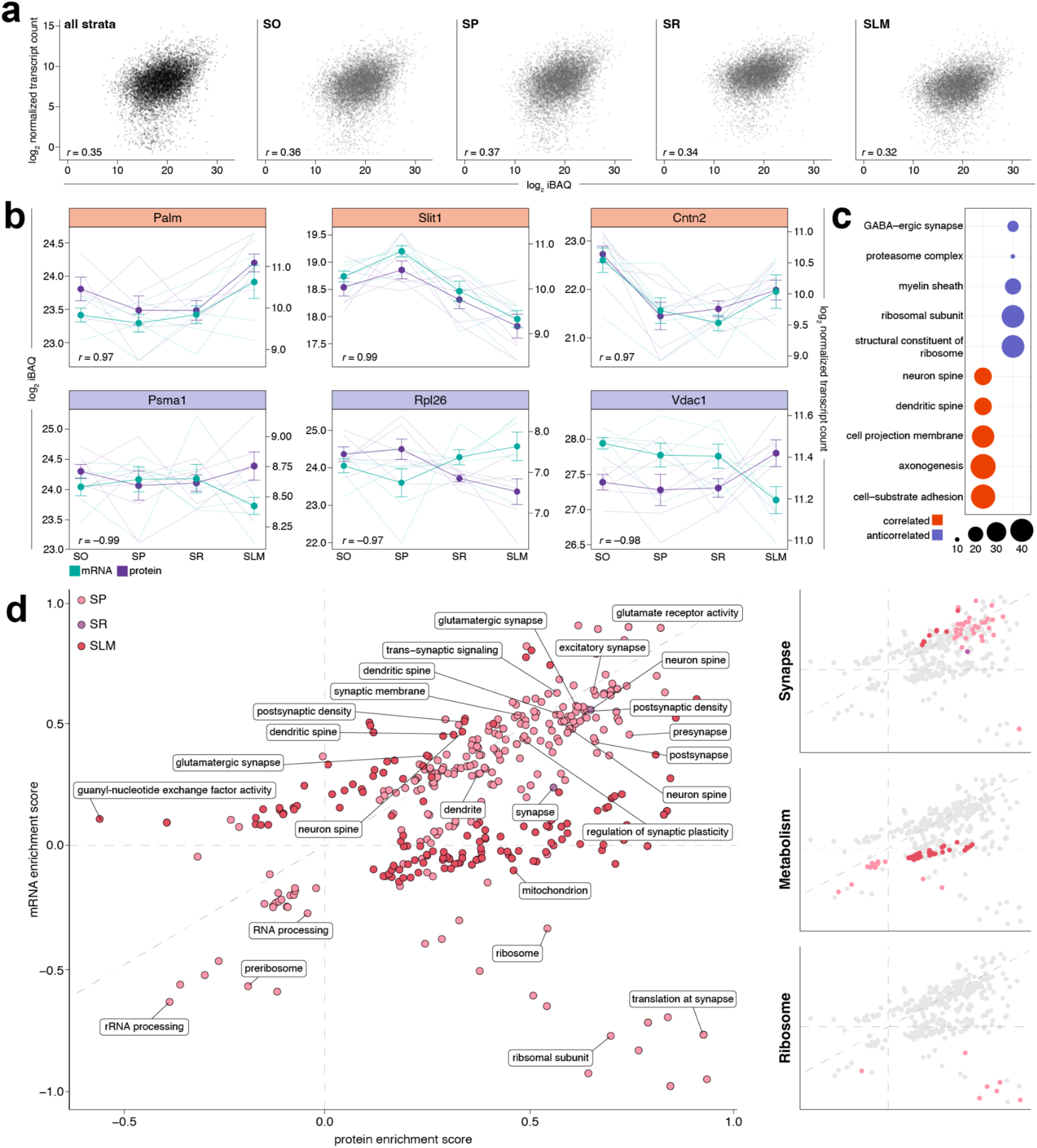
Comparisons of mRNA and protein pairs reveal patterns about their subcellular relationship and local translation. (a) Pearson’s correlation of mean abundances of mRNAs (log_2_ DESeq normalized counts) and their respective protein (log_2_ intensity-based absolute quantification, iBAQ) across CA1 strata and within each stratum. (b) Selected candidates showing significant Pearson’s correlation and anti-correlation (*p* < 0.05) in their mean abundance changes across strata. Abundances for individual replicates are shown as translucent lines, while solid lines represent means ± SD. (c) Selected GO terms overrepresented (*p* adjusted < 0.01) by populations of transcripts and proteins showing a strong positive correlation (≥ 0.9) and anti-correlation (≤ –0.9) in their mean abundance changes across strata. (d) 2D annotation enrichment ^134^ of GO terms for all enriched mRNAs and protein (log_2_ fold-change > 0.1) for each stratum. Common GO terms were found for all strata besides SO. A score of (0,0) means the distribution of annotation terms is similar to the overall distribution. Each point represents one GO annotation for one strata.

Lastly, we tested whether the quantitative relationships between paired mRNAs and proteins could be described in the context of their GO annotations (Table 4). Using 2D annotation enrichment^134^, we evaluated whether members of annotation terms show consistent behavior in either dataset. After filtering for strata-enriched mRNAs and proteins, we found shared GO terms for all strata except SO, indicating either enrichment in both datasets or depletion in one or both dimensions (Fig. 7d, Table 4). Synaptic GO terms were consistently enriched for both mRNAs and proteins across SP, SR, and SLM. In contrast, metabolic terms were predominantly enriched in SLM proteins and depleted in both datasets in SP. Ribosome-related terms were additionally observed in SP, where they were largely enriched for the proteins and depleted for mRNAs.

## DISCUSSION

Understanding the molecular diversity of the brain at multiple layers requires detailed maps of local transcriptomes and proteomes with spatial resolution and high coverage. Efforts to address this challenge have been hindered by the necessity to correlate transcripts from one sample or from one published study with proteins from another. Here, we microdissected hippocampal subregions and strata and performed FASS-based synaptosome purification for RNA-seq and LC-MS/MS analysis of the same samples, in parallel. We found marked differences in the transcriptomic-proteomic landscape of the different hippocampal subregions and CA1 strata defined by their composition of differing cell types, projections, and subcellular compartments.

Our dissection protocol allowed us to collect small areas of the hippocampus from fresh tissue without a requirement for fixation or laser-capture microdissection, providing spatial resolution without use of *in situ* techniques that offer lower coverage, such as *in situ* sequencing or MALDI imaging. The RNA-seq used here provided comprehensive transcriptome profiles and the single-shot LC-MS/MS runs identified ∼10,000 proteins from tissue per subregion or strata, achieving depth comparable to or greater than other studies performing fractionation-based proteomic profiling of the mouse and human hippocampus^4,135^. Thousands of mRNAs and proteins were differentially enriched in subregions or strata from tissue, identifying a previously undescribed spatial organization of molecules. One notable example is the protein Htt which was significantly enriched locally in SR. This fits with deficits in hippocampal plasticity in SR as well as deficiencies in learning and memory that have been reported in Huntington’s disease model mice^136–138^.

At both the mRNA and protein level, we observed an enrichment of cell adhesion molecules and guidance factors that are essential for the formation of strata and their inhibitory and excitatory synaptic connections. Examples include the cell-adhesion protein Nfasc, enriched in SO, which has been shown to regulate the translocation and formation of gephyrin clusters to the axon hillock of inhibitory synapses *in vitro*^139^. Similarly, Latrophilin-2 (encoded by the *Adgrl2* gene) is a synaptic cell-adhesion protein essential for connectivity from layer III of the entorhinal cortex, localized to dendritic spines of excitatory neurons within SLM^85–87^ where we detect its’ enrichment in our dataset. We also detected various LRR-containing proteins and mRNAs in SLM, and to a lesser extent in SR, which can act as organizers for synaptic connections^100,140,141^. Furthermore, we detected several members of guidance cue families including ephrin, Slits, semaphorins and plexins as well as netrin and reelin. Many of these were specifically enriched in SR or SLM tissue including the postsynaptically localized Ephb2 which controls organization and formation of dendritic spines^142^. Other examples include Sema3c, Ntng1 and its ligand Lrrc4c which are essential for correct patterning of thalamic-hippocampal connections^129^, and Reln, crucial for formation and maintenance of entorhinal-hippocampal connections to SLM^143^. These findings closely align with developmental characterization of localized mRNAs of extracellular and membrane-bound guidance molecules^103,125^.

Locally enriched molecules also corresponded to many GO annotations associated with known features of individual strata such as myelination (SO), nuclear processes (SP), and synaptic specialization (SR). SLM enriched proteins, however, uniquely overrepresented multiple mitochondrial and metabolic terms. It was previously reported that increased mitochondrial respiration occurs specifically at the SR/SLM border during fear conditioning^117^ and that mitochondria fuel local protein synthesis during synaptic plasticity^144^. The enrichment of mitochondrial machinery in this layer could therefore serve to enhance local subcellular processes, such as local translation and plasticity.

Our FASS-based purification of synaptosomes from microdissected CA1 strata identified mRNA and protein populations enriched at synapses in different neuronal compartments. These included molecules spanning various functional categories such as ion channels, receptors, exocytosis factors, and ATPases, consistent with previous findings across brain regions^9^. The enriched transcripts encode proteins involved in shaping the synaptic environment, such as collagens which are increasingly recognized for their role in the dynamic cellular processes and neurodegenerative diseases^145^. Specific demands of different synapses along the neuron are also reflected by differential enrichment of diverse transporters (SLC) family members that transport bile acids, ions, protons, neurotransmitters, amino acids, and ADP/ATP. For example, many of these transporters show high enrichment in SLM, a region known for its strong synaptic inhibition^50,51^. High levels of synaptic inhibition often correlated with increased energy demand to maintain pump and transporter activity. We also noted, as observed previously^22,53,146,147^, many mRNAs for ribosomal proteins were enriched.

This fits with recent descriptions of local translation and incorporation of nascent ribosomal proteins into pre-existing ribosomes^82,148^ as well as electrophysiological evidence of protein synthesis required for plasticity of Schaffer collateral-CA1 synapses in SR^149^ and temporoammonic connections to SLM^51^. Our detected enrichment of kinesins in purified synaptosomes was also intriguing, suggesting the possibility of local control of transport and delivery both to the distal dendrites as well as the cell body, given the mixed polarity of dendritic microtubules^150^. Many proteins with spatially localized synaptic enrichment were excitability modulators, such as potassium channels that are known to form gradients along pyramidal neurons^54,55^. Interneurons including the OLM neurons of the SO show both transient and sustained outward potassium currents^38,46^. We found that one subunit forming M-type channels, Kcnq2, was enriched only in SO. As this subunit can form either homo-or heteromeric channels with Kcnq3, this might reflect an abundance of homomeric channels in SO which can alter both the channel surface expression and conductance^151,152^. These channels could contribute to unique synaptic plasticity within this region^40,43^ which is different from plasticity mechanisms observed in other strata.

Our analysis of mRNA and proteins originating from the same sample revealed a modest correlation in their abundances, in line with comparisons made across samples or studies^15,19,53^. When we correlated the extent to which mRNA variation results in protein abundance changes across strata, though, we found strong linear correlations and anti-correlations of certain mRNA-protein pairs with different functional annotations. For certain examples (Palm, Cntn2, Slit1), strong positive correlations can be explained in part by demand-driven localization, as their translation is differentially regulated between somata and dendrites as previously shown by Ribo-seq data^24^. Our anti-correlated examples (Psma1, Rpl26, Vdac1) accordingly are not reported to be differentially translated between compartments^24^. These changes in protein abundances could be explained by other factors including longer half-lives that have been shown for proteasomal, ribosomal, and mitochondrial proteins^130–133^, indicating that these mRNAs may not be needed ‘on-demand’ and therefore might not co-reside with the associated protein. This collectively suggests that combining mRNA concentration variation with protein translation regulation and half-life data provides a more informed prediction of variable protein abundances between neuronal compartments^19^.

While our study comprises a comprehensive map of molecular diversity in the hippocampus, future studies will bring us closer to a complete picture. For example, we purified synapses using Syn-1 as a marker to obtain pan-synaptic information. As FASS can be used to sort different synapse types to high purity^9^, and the hippocampus contains diverse neuronal populations^118,153^ one could define the omics-level differences between specific synapse types (e.g. excitatory or inhibitory) within one subregion or strata. Moreover, additional layers of analyses could be added that describe RNA-protein relationships. This might include investigation of 5’ and 3’ UTRs, miRNAs, specific RNA binding proteins, RNA granules, or post-translational modifications. Lastly, one may explore how our observed molecular landscape is dynamically modulated by different types of activity, or perturbed in disease. Altogether, our dataset provides the foundation for such future work.

## METHODS

### Experimental model and animal details

The procedures involving animal treatment and care were conducted in conformity with the institutional guidelines that are in compliance with national and international laws and policies (DIRECTIVE 2010/63/EU; German animal welfare law; FELASA guidelines). The animals were euthanized according to annex 2 of § 2 Abs. 2 Tierschutz-Versuchstier-Verordnung and animal numbers were reported to the local authorities (Regierungspräsidium Darmstadt).

### Animals used for transcriptomics and proteomics

10/11-week-old *Cre*-positive and *Cre*-negative F1 littermates from offspring of a cross between hemizygous *Syn1-Cre* female mice (B6.Cg-Tg(Syn1-cre)671Jxm/J, RRID:IMSR_JAX:003966) and homozygous *condSypTom* males (B6;129S-Gt(ROSA)26Sortm34.1(CAG-Syp/tdTomato)Hze/J obtained from The Jackson Laboratory, RRID:IMSR_JAX:012570)^154^ were used (deposited by Hongkui Zeng Lab). *Cre*-positive mice were used for the synaptosome generation and Fluorescence-Activated Synaptosome Sorting (FASS), and their *Cre*-negative aged-matched litter mates were used for whole tissue preparations. To allow enough material for downstream transcriptomic and proteomic analysis, hippocampi of two mice, one male one female, were pooled together for one replicate (N=1, hippocampi of two mice) which was then divided following slicing and microdissection for parallel transcriptomic and proteomic analysis.

### Tissue preparation and microdissection

Preparation of acute hippocampal slices from *Cre*-positive mice and their *Cre*-negative siblings was done as previously described^155^. Briefly, mice were decapitated after isoflurane anesthesia, the brain was immediately removed and put in ice-cold, slushed artificial cerebrospinal fluid (ACSF) solution with sucrose (87 mM NaCl, 25 mM NaHCO_3_, 1.25 mM NaH_2_PO_4_, 2.5 mM KCl, 10 mM Glucose, 75 mM Saccharose, 7 mM MgCl and 0.5 mM CaCl_2_) carbogenated with 95% O_2_/5% CO_2_. The brain was cut into halves along the longitudinal fissure, the neocortical side of the hemispheres were cut at an angle of 30° and glued on an ice-cold specimen holder, and sliced with a Vibratome (VT 1200 S, Leica) to 300 µm-thick horizontal/transverse slices in ice-cold, carbogenated ACSF solution with sucrose. Slices from the middle part of the hippocampus were selected for dissection by clear visual identification of subregions and CA1 layers. Before slice preparation, blades were adjusted parallel to the cutting surface using VIBROCHECK and set to a cutting speed of 0.08 mm/s with an amplitude of 1 mm. Generally, the left and right hippocampi taken together from one mouse yielded 8-12 optimal slices that were used for microdissection. Microdissections of CA1, CA2/3, DG and the *Strata oriens*, *pyramidale, radiatum* and *lacunosum-moleculare* were carried out manually under a dissecting microscope at 4 °C using a microdissection scalpel (McKesson,701459/Surgical Specialties, 7503; Stab Knife MSP™ Stainless Steel, Blade / Tip Type 15° Angled Straight Sharp Pointed Tip) and sections were collected in 0.5 mL RNAse-free 1 X PBS supplemented with Millipore Protease inhibitor cocktail III (539134) 1:750 and SUPERase•In™ RNase Inhibitor 1:80 (AM2696) in 1.5 mL LoBind Eppendorf tubes. In brief, the division of the hippocampal fissure serves as a visual separation between Ammon’s horn (*Cornu Ammonis*) and the DG, facilitating the isolation of the upper segment of Ammon’s horn (CA1). Removal of the subiculus from the whole brain slice facilitated access to CA1. Following this, the remaining portion of Ammon’s horn (CA3 and CA2 together as they were visually indistinguishable) was separated from the DG, the boundaries of which are optically identifiable, the hillus was cut from approximately the suprapyramidal blade to the infrapyramidal blade. After dissection, the combined sections from a given region-of-interest were immediately spun down, the supernatant was removed, and the samples were snap-frozen and stored at –79 °C. This dissection technique provided sufficient material for both tissue and synapse purification without laser-associated heat degradation, tissue fixation, or freezing^156^, yielding higher amounts for downstream synaptosomes and tissue preparation.

### Percoll density gradient for crude synaptosome preparation

The same procedure, described above, for preparing hippocampal slices and performing microdissection was followed. Synaptosomes were prepared as previously described^9,157^. After collection in LoBind Eppendorf tubes, the samples were transferred to a clean 1.0 mL glass WHEATON® Dounce Tissue Grinder and homogenized using 20 strokes with the ‘loose’ and 20 strokes with the ‘tight’ pestle in gradient medium (GM; 0.25 M sucrose, 5 mM Tris-HCl, 0.1 mM EDTA supplemented with Millipore Protease inhibitor cocktail III 1:750 (539134) and SUPERase•In™ RNase Inhibitor 1:80 (AM2696)). The homogenate was centrifuged for ten minutes at 4 °C at 1,000 × *g*. The supernatant (S1) was layered onto a Percoll density gradient with 23%, 10% and 3% Percoll in the GM buffer. The gradient was centrifuged for 5 min at 32,500 × *g* at 4 °C with maximum acceleration and minimum deceleration using a Beckman Coulter JA-25.50 rotor in an Avanti J-26S XPI centrifuge (both from Beckman Coulter). The resulting bands were labeled, from top to bottom, F0, F1, F2/3, F4. The F2/3 band was retrieved with a syringe and a blunt cannula and used as an input for subsequent Fluorescence-Activated Synaptosome Sorting (FASS). All procedures were conducted on ice.

### Fluorescence-Activated Synaptosome Sorting (FASS)

Dissections were performed and synaptosomes from each hippocampal region and CA1 strata from the *Cre*-positive mouse strain (see above) were isolated as described above^157^. Subsequent Fluorescence-Activated Synaptosome Sorting (FASS) was then used to sort synaptosomes to high purity^9^ (Extended data Fig. S2a, b) on a FACSAria Fusion (BD Biosciences) running FACSDiva (BD Biosciences). Doublet particles were excluded based on SSC-H and SSC-W and other parameters were as described^9^ including the following settings: 488nm laser (for FM4-64), 561nm laser (for tdTomato signal), sort precision (0-16-0), FSC (317 V), SSC (488/10 nm, 370V), PE ‘‘tdTomato’’ (586/15 nm, 470V), PerCP ‘‘FM4-64’’ (695/40 nm, 470), thresholds (FSC = 200, FM4-64 = 700). Samples were analyzed and sorted at approximately 20,000 events/s and a flow rate of < 5. BD FACSFlow (12756528) was used as a sorting buffer. Synaptosomes were sorted through a 70 µm nozzle and collected on a Whatman GF/F glass microfiber filter (1825-090) supported by a PE drain disk (231100) for stability. Both were punched out together using a Militex biopsy Punch (4 mm, 15110-40-X) which was washed prior to sorting with either 700 µL RNAsecure (1:5; Thermo Fisher Scientific, AM7006) or Millipore Protease inhibitor cocktail III (1:750; 539134) to inhibit RNAses or Proteases in the sorting buffer, respectively. After 2 million synaptosomes were collected on the filter, it was washed with 1 mL RNAse-free 1 X PBS to remove residual FACSFlow and stored at –79 °C until further processing. Purified synaptosome (here referred to as P3 based on the gates used in the sorting layout) fractions were sorted based on the presence of both a membrane dye (FM4-64; Thermo Fisher Scientific, T13320) and the tdTomato signal. Control (here referred to as P2 or synaptosome-sized) fractions were also generated from the F2/3 precursor synaptosome fraction based on only the membrane dye signal intensity but independent of tdTomato signal, resulting in a conservative unsorted fraction which contained both synaptosomes and synaptosome-sized contaminants. Synaptic enrichment was later determined by comparing abundances in P3 relative to P2 fractions. After FASS, filters with synaptosomes were stored at –79 °C until further processing.

### Tissue homogenization, RNA extraction, library preparation and sequencing

Tubes with tissue samples were kept on dry ice until RNA extraction. Frozen tissue samples were resuspended in 1 mL TRIzol, transferred to a clean 1.0 mL glass WHEATON® Dounce Tissue Grinder and homogenized using a ‘loose’ and ‘tight’ pestle for 20 strokes each. The homogenate was then transferred back to the 2 mL RNAse-free Eppendorf tube and triturated 20x using a 23 G needle and 1 mL syringe^24^ and kept on ice. All samples were incubated for 5 min at room temperature and centrifuged for 3 min at 16,100 × *g*. The clean supernatant was collected in a new RNase-free tube and RNA was column purified using ZymoReaserach kit. The RNA from CA1, CA2/3 and DG subregions was extracted using the Direct-Zol RNA Miniprep Kit (R2051) and from the CA1 *Strata* using the Direct-Zol RNA Microprep Kit (R2061). RNA quantity and quality were measured on a Qubit fluorometer (Invitrogen, Q33216) and on a 2100 Bioanalyzer (RNA Pico chip, Agilent Technologies).

mRNA-seq libraries were prepared from 25 ng of total RNA using NEBNext Ultra II Directional RNA Library Prep Kit for Illumina (E7760), Poly(A) mRNA Magnetic Isolation Module (E7490) and Multiplex Oligos for Illumina Set A (E6440). The quantity of the libraries was measured on Qubit fluorometer (Invitrogen, Q33216) and quality assessed using HSDNA 2100 Bioanalyzer assay (Agilent Technologies). Libraries were sequenced on Illumina NextSeq2000 using P2 reagents with 61 bp paired-end reads.

### Synaptosome mRNA library preparation and sequencing

Library preparation and sequencing of synaptosomes was performed using a modified version of an established protocol^25,158^. In brief, to lyse membranes and digest DNA, synaptosomes were resuspended in 7.5 µL of 1 mM DTT, 1 U/µL Recombinant RNAse Inhibitor (2313A, Takara Bio), 1 X SingleShot Lysis Buffer and 1 X DNAse Solution (1725080, Bio-rad), and incubated for 20 min at 20 °C, followed by enzyme inactivation for 5 min at 75 °C. Next, reverse transcription with template switching was carried out. We first hybridized indexed RT primers (final concentration: 1 mM) (see Supplementary Table S2), adding DTT (10 mM), dNTPs (1 mM), dCTPs (1.5 mM) and Recombinant RNAse Inhibitor (1 U/µL) and incubating each sample for 5 min at 65 °C, and then immediately transferred the samples to ice. After adding the reverse transcriptase SuperScript IV (10 U/µL) with SuperScript RT buffer (1 X; Thermo Fisher Scientific, 18090050), MgCl_2_ (6 mM), Betaine (1 M, Merck B0300-5VL), and template switch oligo (1 µM, 5’-/5BiosG/AAGCAGTGGTATCAACGCAGAGTGTCGTGACTGGGAAAACCCTGGCrGrGrG-3’) the sample was incubated for 10 min at 55 °C, followed by enzyme inactivation for 10 min at 80 °C. For PCR, 1 X of the DNA polymerase mix (KAPA HiFi HotStart Ready Mix, 07958935001) and the PCR primer (0.1 µM, 5’-AAGCAGTGGTATCAACGCAGAGT-3’) were added. PCR was then carried out using the following cycling conditions: 3 min incubation at 98 °C for activation of the DNA polymerase, 18 cycles of incubation at 98 °C for 20 s to denature DNA, 67 °C for 10 s for annealing, and 72 °C for 6 min for extension. After the last cycle, the sample was again incubated at 72 °C for a final extension phase. Between incubations, samples were constantly kept on ice. After PCR, the Whatman GF/F glass microfiber filter was removed by transferring the sample to a filter column (Spin-X Centrifuge Tube Filter, 0.45 uM in 2.0 ml Tube; Corning, 8162,) and centrifuged at 3,000 × *g* for 1 min. Next, 28.8 µL AMPure XP beads (Beckman Coulter) were used for DNA purification. Multiplexed samples were subsequently pooled and further prepared for sequencing using the Nextera XT DNA library prep kit (Illumina) according to the manufacturer’s instructions, but using a custom P5 primer (5’-AATGATACGGCGACCACCGAGATCTACACGCCTGTCCGCGGAAGCAGTGGTATCAACGCAGAGTAC-3’). Paired-end sequencing was performed using the NextSeq^TM^ 2000 system (Illumina) with NextSeq^TM^ 1000/2000 P2 (100 cycles) and a custom sequencing primer for Read 1 (5’-GCCTGTCCGCGGAAGCAGTGGTATCAACGCAGAGTAC-3’). The following sequencing parameters were used: Read 1 = 26 bp, Read2 = 82 bp, Read 1 Index = 8 bp.

### Sample processing of tissue for LC-MS/MS

Frozen tissue was briefly thawed and mechanically lysed by addition of 100 μL lysis buffer (5% SDS, 50 mM Tris, pH 7.55 with HCl) and 10 ‘strokes’ with a pestle (VWR International) after which the pestle was washed with 100 μL additional lysis buffer. Further homogenization of lysates was then achieved by pipetting up and down 3 times, 10 min sonication, and 5 min at 75 °C and 400 rpm. After cooling, lysates were incubated with benzonase (1 μL; 250 units/mL stock solution; Sigma) for 10 min at room temperature. To clear debris, lysates were centrifuged for 10 min at 13,000 × *g* and the supernatant was taken. Protein concentration of lysates was determined by a BCA assay (Thermo Fisher Scientific) from a 1:5 dilution to ensure minimal interference from the high detergent concentration.

The samples were prepared for bottom-up proteomics using a suspension trapping protocol as previously reported^159^. Briefly, 30 μg of protein in lysis buffer was reduced using 20 mM DTT for 10 min at RT, and alkylated with 50 mM iodoacetamide for 30 min at RT in the dark. Afterwards, samples were acidified using phosphoric acid to a final concentration of ∼ 1.2%. Binding/wash buffer (BW: 90% methanol, 50 mM Tris, pH 7.1 with H_3_PO_4_) was added in a 1:7 ratio. The protein suspension was loaded onto the S-trap filter (size: “micro”; ProtiFi) in 150 μL steps by centrifugation for 20 s at 4,000 × *g*, and trapped proteins were then washed with 150 μl of BW buffer three times. Digestion buffer was prepared by addition of trypsin (1 μg; Promega) and LysC (0.2 μg; Promega) to 40 μL ammonium bicarbonate buffer buffer (ABC: 50 mM). Digestion was performed overnight (∼18 hours) at RT in a humidified chamber under gentle agitation. Peptides were later eluted by centrifugation at 4,000 × *g* for 60 s via one wash with 40 μL ABC buffer and two additional washes with 0.2% formic acid in MS-grade water.

After digestion, peptides were desalted using a modified C18 StageTip protocol^160^. StageTips were made by addition of two disks of C18 material (Empore 3M) to a 200 μL pipette tip (Eppendorf). Disks were conditioned using 100 μL pure methanol and centrifuged at 2,000 × *g* for 3 min, after which they were washed with 100 μL 50% acetonitrile with 0.5% acetic acid, and equilibrated with two washes of 100 μL 0.5% acetic acid. Peptides were loaded and reloaded on disks by centrifugation at 1,500 × *g* for 5 min, and desalted by washing with 100 μL 0.5% acetic acid twice at 2,000 × *g* for 3 min. Desalted peptides were eluted using 75 μL 50% acetonitrile with 0.5% acetic acid and centrifuged at 2,000 × *g* for 3 min, repeating the elution twice. Eluted peptides were dried *in vacuo* at 45°C.

### Sample processing of synaptosomes for LC-MS/MS

Sorted synaptosomes and control particles were filtered onto Whatman glass microfiber filters (GF/F, Cytiva) as described above, and stored at –79 °C in Eppendorf tubes until further processing as previously described^9^. After thawing, digestion buffer comprising of trypsin (0.1 μg) and LysC (0.1 μg) in 25 μL triethylammonium bicarbonate (TEAB; 50 mM) was added to each sample. Digestion was performed overnight (∼ 18 hours) at 37 °C. 50 μL pure acetonitrile was then added and samples were centrifuged at 16,000 × *g* for 10 min. A ZipTip pipette tip (Millipore) was washed once using 50% acetonitrile in MS-grade water, after which supernatants were filtered through by centrifugation for 1 min at 2,000 × *g*. 50 μL 50% acetonitrile was then added to each sample and the process was repeated, pooling both supernatants during filtering. Eluted peptides were dried *in vacuo* at 45 °C.

### LC-MS/MS analysis

Dried peptides were reconstituted in 5% acetonitrile (ACN) with 0.1% FA. For synaptosome samples, reconstitution buffer was supplemented with iRT peptide standard (Biognosys) at 1:100. Reconstituted peptides were loaded onto a C18-PepMap 100 trapping column (particle size 3 µm, L = 20 mm, Thermo Fisher Scientific) and separated on a C18 analytical column with an integrated emitter (particle size = 1.7 µm, ID = 75 µm, L = 50 cm; CoAnn Technologies) using a nano-HPLC (U3000 RSLCnano, Dionex) coupled to a nanoFlex source (2000 V, Thermo Fisher Scientific). Temperature of the column oven was maintained at 55 °C. Trapping was carried out for 6 min with a flow rate of 6 μL/min using loading buffer (100% H_2_O, 2% ACN with 0.05% TFA). Peptides were separated by a gradient of water (buffer A: 100% H_2_O and 0.1% FA) and acetonitrile (buffer B: 80% ACN, 20% H_2_O and 0.1% FA) with a constant flow rate of 250 nL/min. In 155 min runs, peptides were eluted by a non-linear gradient with 120 min active gradient time, as selected for the respective MS method by Muntel et al.^161^ Analysis was carried out on a Fusion Lumos mass spectrometer (Thermo Fisher Scientific) operated in positive polarity and data independent acquisition (DIA) mode. In brief, the 40-window DIA method had the following settings: Full scan: orbitrap resolution = 120k, AGC target = 125%, mass range = 350-1650 m/z and maximum injection time = 100 ms. DIA scan: activation type: HCD, HCD collision energy = 27%, orbitrap resolution = 30k, AGC target = 2000%, maximum injection time = dynamic.

### Processing of DIA LC-MS/MS data

DIA raw files were processed with the open-source software DIA-NN (version 1.8.2 beta 27) using a library-free approach. The predicted library was generated using the *in silico* FASTA digest (Trypsin/P) option with the UniProtKB database (Proteome_ID: UP000000589) for *mus musculus*. Deep learning-based spectra– and RT-prediction was enabled. The covered peptide length range was set to 7-35 amino acids, missed cleavages to 2 and precursor charge range to 1-5. Methionine oxidation was set as variable, and cysteine carbamidomethylation as a fixed modification. The maximum number of variable modifications per peptide was limited to 3. According to most of DIA-NN’s default settings, MS1 and MS2 accuracies as well as scan-windows were set to 0, isotopologues and match-between-runs were enabled, while shared spectra were disabled. Protein inference was performed using genes with the heuristic protein inference option enabled. The neural network classifier was set to single-pass mode and the quantification strategy was selected as ‘QuantUMS (high precision)’. The cross-run normalization was set to ‘RT-dependent’, library generation to ‘smart profiling’, and speed and Ram usage to ‘optimal results’. The DIA-NN report table and the respective fasta file were imported in the statistical computing software R and analyzed using the MS-DAP R package^162^.

### Preprocessing methods for RNA-Seq tissue datasets

To process the Illumina NextSeq 2000 sequencing data, we implemented a comprehensive bioinformatics pipeline running on a high-performance computing cluster with allocated resources of one node, 32 CPUs, and a runtime limit of 100 hours. The pipeline encompasses steps from raw data demultiplexing to the quantification of gene expression levels. (i) *Demultiplexing*. Initially, raw BCL files generated by the Illumina NextSeq 2000 system were converted to FASTQ format using the Illumina bcl2fastq2(v2.20.0.422) software. The conversion process was tailored to our experimental design by providing a custom sample sheet, enabling the direct demultiplexing of samples without lane splitting. This step utilized 8 threads for loading, processing, and writing, optimizing throughput. (ii) *Quality Control and Adapter Trimming*. Subsequent to demultiplexing, the FASTQ files underwent quality control and adapter trimming using fastp(0.23.1)^163^. This utility was configured to remove adapter sequences, specified both as individual sequences and as a FASTA file, and to trim low-quality bases and poly-X sequences. The quality threshold was set to a minimum average quality score of 20, with a minimum poly-X length of 10 for trimming. This process was parallelized across 16 threads to expedite execution. Each sample’s processing generated a corresponding report in JSON and HTML formats, along with a textual summary. (iii) *Alignment.* The cleaned FASTQ files were aligned to the *Mus musculus* (mm39) reference genome using the STAR(2.7.11b) aligner^164^. The alignment was conducted with the aim of generating sorted BAM files, considering only uniquely mapped reads and filtering out non canonical intron motifs. Alignment was performed in a strand-specific manner to account for RNA-Seq library preparation nuances. Post-alignment, each sample’s BAM file was indexed with samtools for efficient downstream processing. (iv) *Alignment Summary*. An alignment summary was compiled for each sample, detailing the total number of input reads and uniquely mapped reads. This step involved parsing STAR log files and aggregating the relevant metrics into a comprehensive tab-delimited summary file. (v) *Quantification*. For gene expression quantification, featureCounts(v2.0.3)^165^ (part of the Subread package^166^) was employed, counting reads mapping to exon regions and annotated with gene IDs. The software was instructed to count paired-end reads as fragments and to include only reads with a mapping quality of 255, ensuring the consideration of high-confidence alignments only. Quantification was carried out on all samples simultaneously, utilizing 16 threads to improve efficiency, and results were compiled into a single output file.

### Preprocessing methods for RNA-Seq of synaptosome datasets

Preprocessing was performed using a modified pipeline for droplet-based single-cell RNA-seq (Drop-seq)^167^. The two different indices, for replicate and sample used during library preparation, were used to perform demultiplexing during preprocessing. After conversion of base-call files into Fastq files and first demultiplexing on the level of biological replicates with bcl2fastq from Picard, a quality check of the reads was performed by using FastQC^168^. Trimming of Poly-A tails, sequenced Template Switch Oligos and filtering of low-quality reads was carried out using programs of the Drop-seq pipeline in combination with Fastp^169^. STAR aligner^164^ was used to then align quality trimmed reads to the *Mus musculus* (mm39) reference genome (Genome Reference Consortium). GENCODE M33 annotation file was downloaded from UCSC Table browser and used for read annotation. In the last step annotated reads were collapsed UMI-wise, demultiplexed based on the sample-trait indices and finally a Digital Gene Expression – Matrix was created.

### Differential expression analysis of transcriptomes

Differential expression analysis was conducted using the DESeq2 standard workflow^170^. Quality control measures involved assessing read yield versus alignment rate and identifying PCA outliers. Matrices were filtered to include only protein-coding mRNA transcripts from the annotated *Mus musculus* (mm39) reference genome (Genome Reference Consortium). Only replicates passing quality checks were used for the final PCA plots and any replicates failing to meet the quality checks were excluded from downstream analysis (one SLM replicate (replicate 5) was excluded for failing quality control). For the tissue data, DESeq2 objects were created separately for regions and strata, with CA1 as the reference for differential expression analysis. For synaptosome datasets, DESeq2 objects were generated for sorted (FMdye-and tdTomato-positive, P3) fraction over (FMdye-positive only, P2) control fraction as a reference. Effect size shrinkage was applied using the *apeglm* method^171^, and the method of *Independent Hypothesis Weighting*^172^ was implemented for *p*-value adjustment of DESeq2 results. Enrichment lists were derived from the differential expression analysis. Significantly enriched ‘marker’ transcripts were identified as transcripts showing positive log_2_ fold-change > 0 in the selected stratum, with statistical significance using an adjusted *p*-value cut-off of < 0.1. Subregion-or strata-specific ‘uniquely’ enriched transcripts were further refined to exclude overlap with significantly enriched transcripts from other strata. For the rank plots, normalized mRNA counts for the tissue subregions and strata were averaged and log-transformed for ranking of genes. For synaptically enriched transcripts, criteria included a log_2_ fold-change cutoff of ± 0.1 and an adjusted *p*-value cut-off of < 0.1 in the control over the sorted synaptosome fraction. Subregion-or strata-specific ‘uniquely’ enriched synaptic transcripts were determined using the same criteria as for tissue data. For PCA of synaptosomes only the significantly synaptically enriched or depleted transcripts were included. Plots were generated using ggplot2^173^ and heatmaps were generated with pheatmap^174^.

### Data analysis of proteomes

Tissue data were analyzed using MS-DAP in the R environment with the following parameters: filter minimum detect = 1, filter minimum quant = 1, filter minimum peptide per protein = 1, filter by contrast = true, normalization algorithm = rlr. PCA was performed using log_2_ intensities of all quantified protein groups. Differential expression was determined by comparing protein intensity in a given subregion or strata with the mean intensity across all subregions or strata using a two-sided t-test with Benjamini-Hochberg correction. Proteins showing significant enrichment (adjusted *p*-value < 0.05) in a subregion or strata were further filtered for only those detected in ≥ 4 replicates with a log_2_ fold-change > 0.1.

Synaptosome data were analyzed using MS-DAP with the same parameters as tissue. Differential expression analysis was also done using MS-DAP, comparing sorted (FMdye– and tdTomato-positive) fractions to unsorted (FMdye-positive only) fractions for each subregion/strata, as well as between sorted fractions for each subregion/strata. MSqRob was selected as the differential expression algorithm with a q-value threshold of 0.05 and a log_2_ fold-change threshold of 0. Proteins were then classified as synapse-enriched if they met one of the following criteria: (i) significantly enriched in the sorted versus unsorted fraction for a given subregion/strata, or (ii) significantly enriched in the sorted fraction of one subregion/strata compared to that of another subregion/strata and having a positive fold-change when comparing sorted versus unsorted for the given subregion/strata. A fold-change cutoff of 0.1 was then added in every case. Synapse depleted proteins were determined using the same criteria in the opposite direction. All enriched/depleted proteins were further filtered for only those detected with 2 or more unique peptides to ensure data robustness. For PCA, log_2_ intensities of all proteins showing synaptic enrichment or depletion were used. Strata-unique synaptic proteins were defined as those classified as synapse-enriched in one strata and no others.

Bar plots, PCA plots, box plots, enrichment plots, and correlation plots were visualized using ggplot2^173^. Heatmaps were visualized using pheatmap^174^.

### Functional enrichment analysis of individual transcriptomes and proteomes

GO overrepresentation analysis of both transcriptome and proteome tissue data was performed using the ClusterProfiler R package (https://github.com/YuLab-SMU/clusterProfiler). An annotation file was fetched for mus musculus from OrgDB using the access code AH111576. Gene lists of significantly enriched proteins or transcripts in each strata determined by previous differential expression analysis were compared to a custom background consisting of all genes identified in the respective experiment. Analysis was carried out with a Fisher’s exact test using gene symbols, the ontology setting “all” (encompassing biological processes, cellular compartment, and molecular function), Benjamini-Hochberg FDR correction, and an FDR cut-off of 0.01. Redundant GO terms were simplified according to the adjusted *p*-value with a cut-off of 0.7 using the measure Wang. Enrichment results were then merged by count for visualization of the top-7 terms per condition.

### Integration of transcriptomic and proteomic data sets

All comparisons between protein and mRNA data were done using log_2_ scaled DESeq2 normalized counts and intensity Based Absolute Quantification (iBAQ) values. iBAQ values were determined using the DIAgui shiny app^175^, quantifying based on unique and shared peptides and filtering for precursor q-value < 0.01 and protein-group q-value < 0.01. Additional parameters included Use MaxLFQ from diann package, only keep peptide counts all, and peptide length 7-35. DIABLO analysis was done using all quantified protein groups. For all further analyses, mean iBAQ values of protein groups aggregated by gene name were used.

For a holistic analysis of the combined datasets, an explorative approach was adopted for multi-omics data integration using Data Integration Analysis for Biomarker Discovery using Latent components (DIABLO), a method from the mixOmics R package^125,127^. DIABLO employs sparse Partial Least-Squares-Discriminant Analysis (sPLS-DA^126^) to maximize correlations between predefined pairs of datasets. After exploring single omics datasets, transcripts and proteins showing enrichment across four strata (’markers’) from matched samples were selected as model inputs. *S. lacunosum-moleculare* replicate 5 was excluded from both datasets due to earlier failure to pass quality control (see above). A 2×2 matrix served as the design matrix to balance feature correlations and sample classification performance, with values ranging from 0 to 1 (higher values prioritizing feature correlations, lower ones prioritizing classification performance). Based on data-driven exploration indicating high dataset correlations between the datasets (component 1 > 0.91), we chose off-diagonal values to be set to 0.6 for a balance between classification performance and identifying correlations between outcomes. We used an exploratory approach (due to samples > 10) with *N-1* components. For the three components 15,15,5 features were selected for DIABLO from both transcriptomic and proteomic datasets. Multi-omics figures were then generated using mixOmics^127^

For 1:1 analysis of the population level relationship between transcripts and their corresponding protein, Pearson correlations were calculated for complete observations of transcripts and proteins using their mean abundance in each strata and across all strata. To analyze the correlation of variation in individual transcript and protein abundances between strata, row-wise Pearson’s correlation was performed on mean values for each strata. Transcript-protein pairs were classified as highly-correlated (*r* ≥ 0.9) or anti-correlated (*r* ≤ – 0.9) and these subsets were used for GO overrepresentation analysis as described previously. For GO analysis, all detected transcripts were used as the background and results for correlated and anti-correlated gene sets were merged before plotting selected terms.

For comparisons of data in the context of categorical GO annotations (GOMF, GOBP, GOCC terms), a multivariate analysis of variance (MANOVA) test was used as implemented in the 2D-annotation enrichment algorithm^134^ embedded in the open-source software Perseus (version 1.6.2.3; RRID:SCR_015753). Log_2_-transformed normalized counts and iBAQ values of transcripts and proteins with log_2_ fold-enrichment > 0.1 for each strata were imported in Perseus and merged in a protein-centric manner. GO annotations of the mouse proteome were downloaded from UniProt via the Perseus framework. P-values resulting from MANOVA were corrected for multiple testing using the Benjamini-Hochberg method (FDR < 0.05). For each omics-dimension, a significance score was calculated based on the location of the center of the distribution of values for genes/proteins of a certain GO category relative to the overall input distribution of the values. The scores were then compared in a scatterplot to express the degree of categorical enrichment for each term per omics data type.

## DATA AVAILABILITY

The raw RNA-sequencing data for tissue and synaptosomes are available at the NCBI Sequence Read Archive (SRA) under the reference number: PRJNA1142450 (“RNAseq of microdissected mouse hippocampal subregions and CA1 strata”) and PRJNA1142761 (“RNA-Seq of sorted synaptosomes from microdissection of mouse hippocampal subregions and CA1 strata”) and available upon reviewer request. Mass spectrometry data associated with this manuscript are being uploaded to the PRIDE repository and are available upon reviewer request. mRNA and proteins can also be assessed at our lab’s public interface https://syndive.org/.

## CODE AVAILABILITY

All customized R scripts have been deposited on Gitlab: https://gitlab.mpcdf.mpg.de/mpibr/schu/synapticdiversity/

## EXTENDED DATA

**Extended Data Fig. S1:**
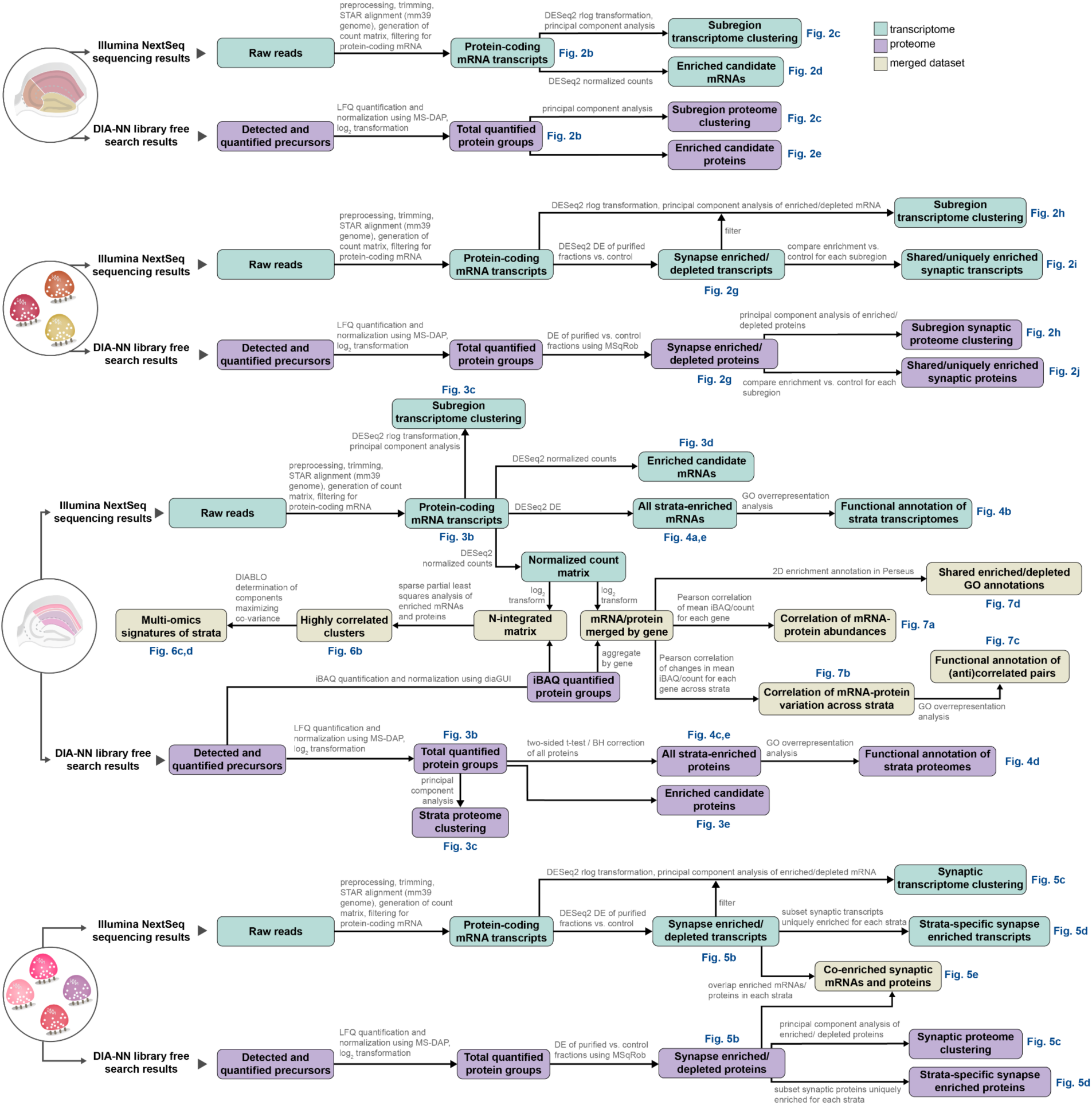
Detailed bioinformatic analysis pipeline of tissue and synaptosome data with relevant steps and final figures indicated.

**Extended Data Fig. S2:**
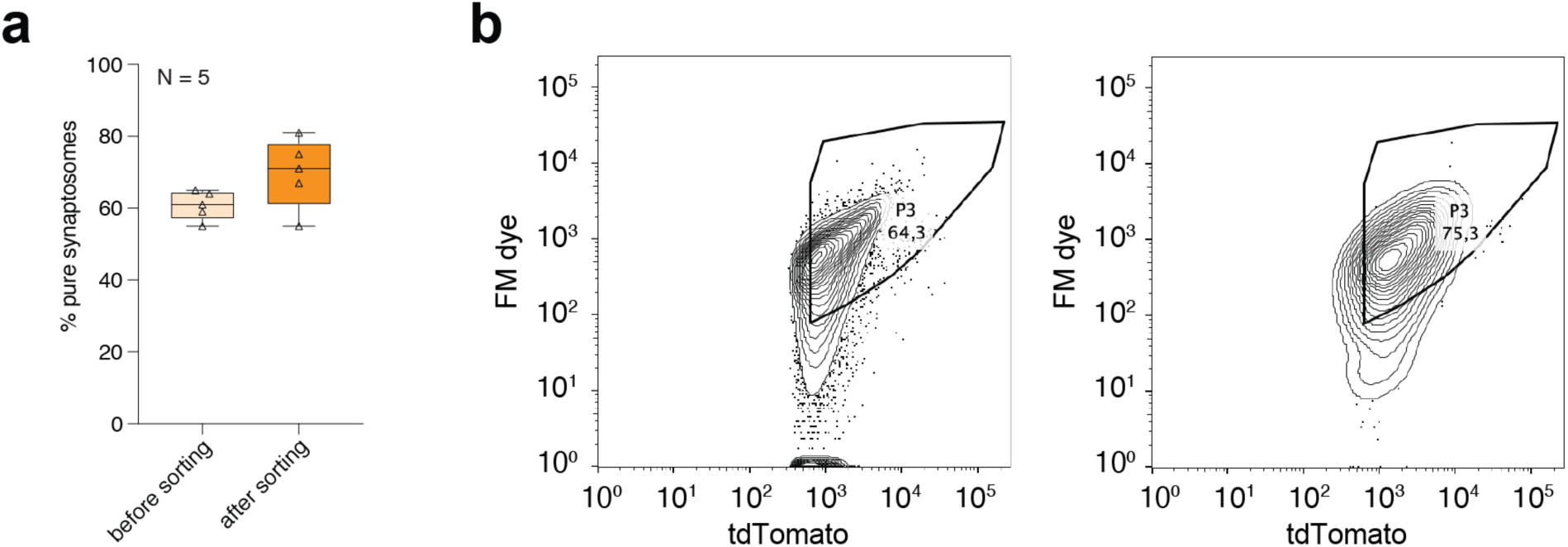
(a) Plot indicating the relative abundance of each fluorescently labeled synaptosome type in the crude synaptosome fraction generated from each sample before and after sorting. Each triangle represents one repeat (2 mice pooled). Shown are 25th to 75th percentiles boxplots, median with error bars representing min to max. (b) Gating strategy and sorting efficiency. FASS contour plots (with outliers) showing the relative density of the targeted tdTomato-positive synaptic population in hippocampal synaptosomes prepared Syn-1::SypTOM mice (64,3%; left). y axes represent fluorescence from a membrane dye (FM4-64) and x axes fluorescence from tdTomato. Following the initial sorting run (left), re-loading of the sorted synaptosomes indicated a high enrichment and purity (75,3%) of the Syn-1::SypTOM sample (right).

**Extended Data Fig. S3:**
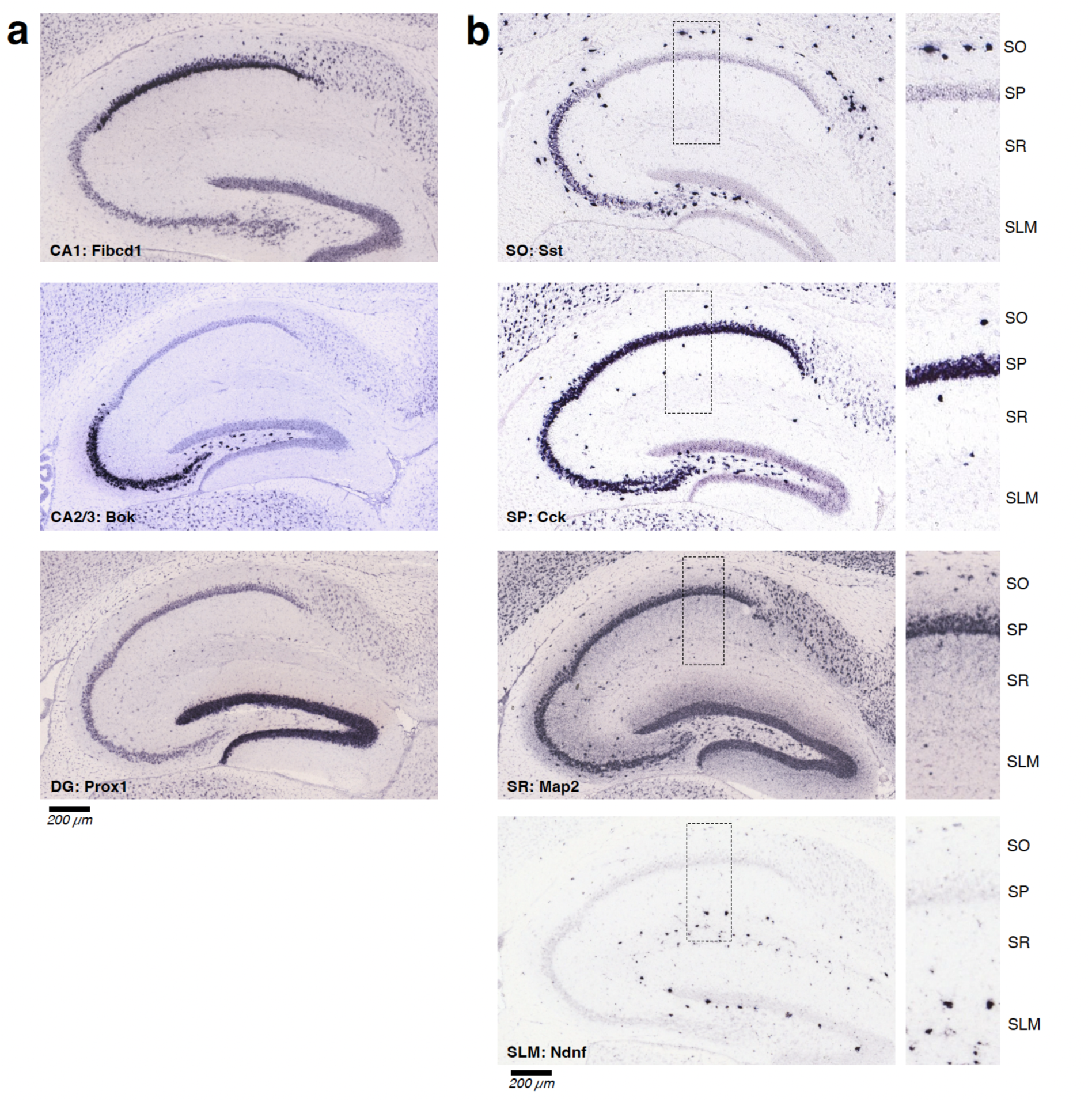
*In situ* hybridization (ISH) assays from the Allen Mouse Brain Atlas sagittal sections (a) Expression of *Fibcd1* in CA1 (mouse.brain-map.org/gene/show/63140), *Bok* in CA2/3 (mouse.brain-map.org/gene/show/31280), *Prox1* in DG (mouse.brain-map.org/gene/show/18893) in the adult mouse brain. (b) Hippocampus expression (left) and zoom in to CA1 strata (right) of *Sst* in SO (mouse.brain-map.org/gene/show/20366), *Cck* in SP (mouse.brain-map.org/gene/show/12209), *Map2* in SR (mouse.brain-map.org/gene/show/17523) and *Ndnf* in SLM (mouse.brain-map.org/gene/show/44012).

**Extended Data Fig. S4:**
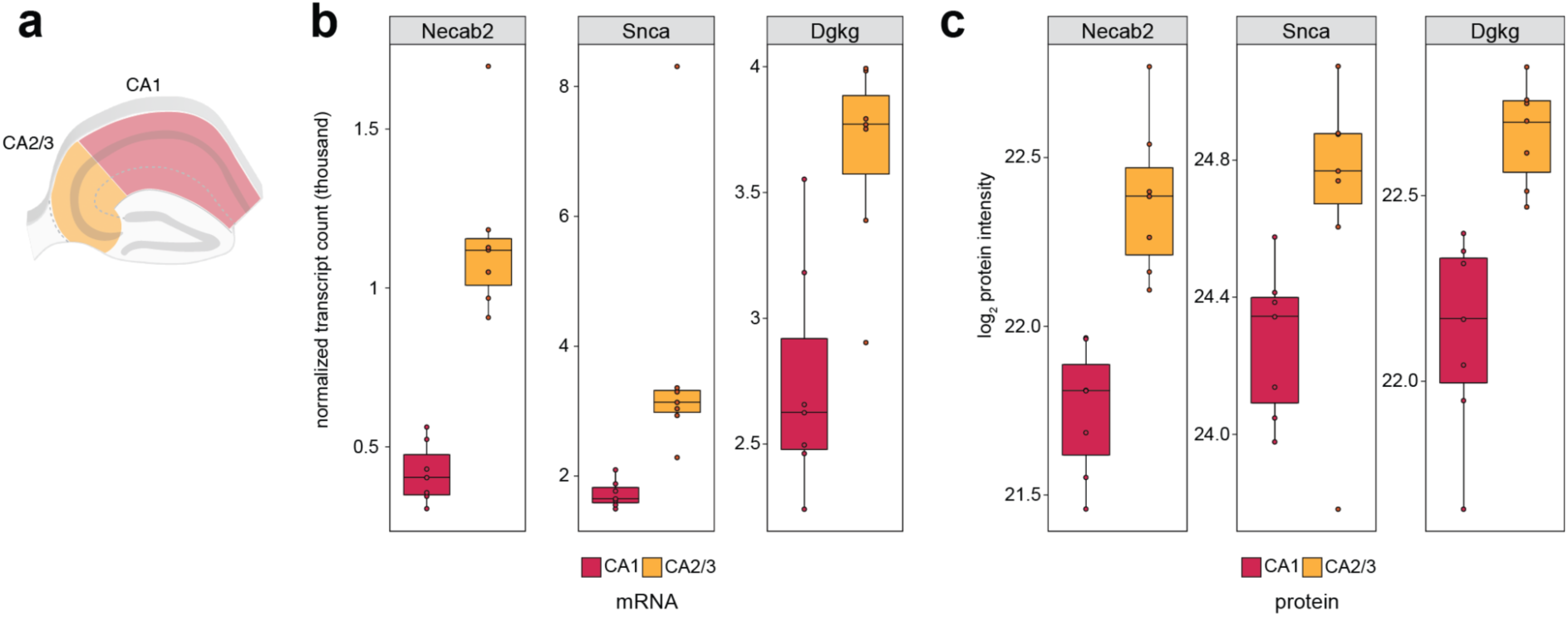
Known CA2 enriched (b) normalized transcripts (thousand) and (c) log_2_ protein intensities are expressed in the CA2/3 dissection of our dataset.

**Extended Data Fig. S5:**
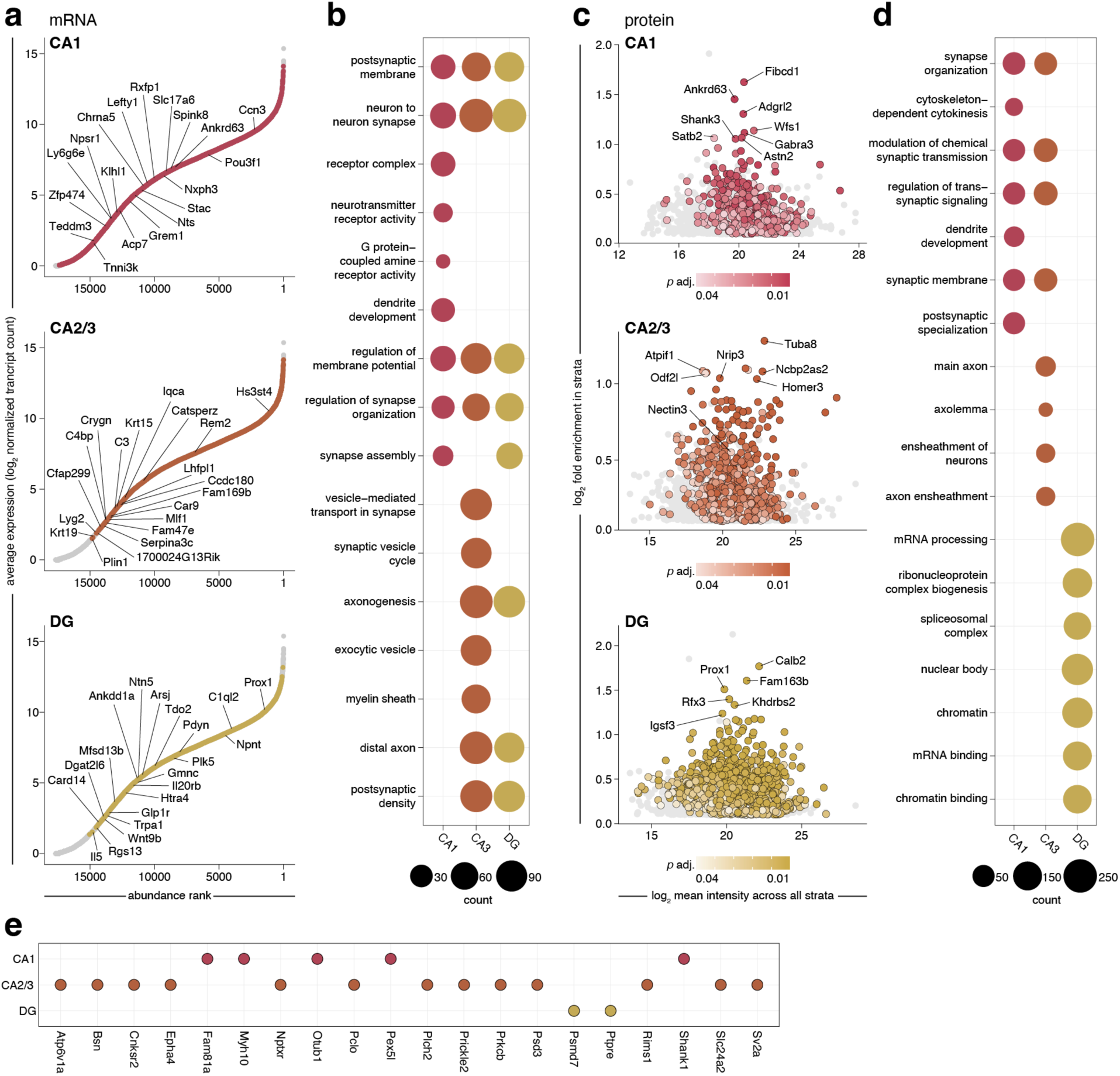
Local transcriptomes and proteomes of hippocampal subregions CA1, CA3 and DG shape functional specialization. (a) Rank abundance plots of mRNA indicating the relative abundance of transcripts in a given subregion. Transcripts with the highest log_2_ fold change are ranked as 1. Coloured points represent transcripts that exhibited unique significant enrichment in each subregion (*p* adjusted < 0.05), while all other detected transcripts are shown in gray. The transcripts that exhibited the top 20 highest relative enrichment (uniquely enriched transcripts were ordered by highest log_2_ fold change) are labeled. (b) Gene ontology (GO) overrepresentation analysis showing top terms (*p* adjusted < 0.01) for significantly enriched transcripts from each subregion. (c) Plots show log_2_ fold enrichment of proteins in a given subregion compared to their mean log_2_ intensity across all subregions. Significantly enriched proteins are shown by coloured dots (*p* adjusted < 0.05) while all other proteins showing enrichment in the subregion are in gray. (d) Gene ontology (GO) overrepresentation analysis showing top terms (*p* adjusted < 0.01) for significantly enriched proteins from each subregion (also see Supplementary Table S1). (e) Dot plot of genes that showed synaptic co-enrichment of mRNA and the respective protein in each subregion.

**Extended Data Fig. S6:**
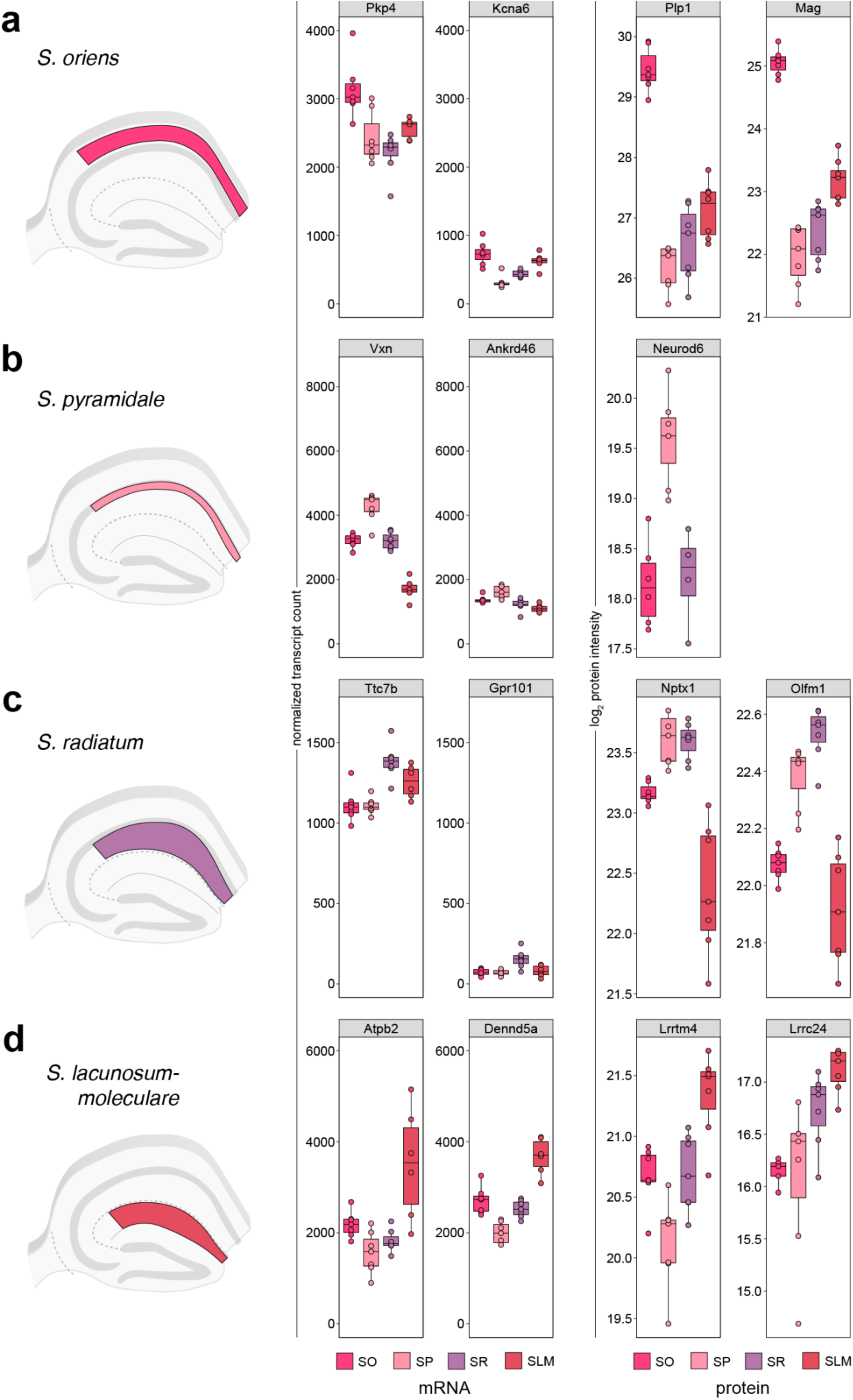
Novel strata-enriched transcripts and proteins. (a) Left: Schematic of CA1 strata. Middle: Boxplots show strata-specific enrichment of previously undescribed mRNA. Right: Boxplots show strata-specific enrichment of previously undescribed proteins. Plots specify median and 1.5 × IQR.

**Extended Data Fig. S7:**
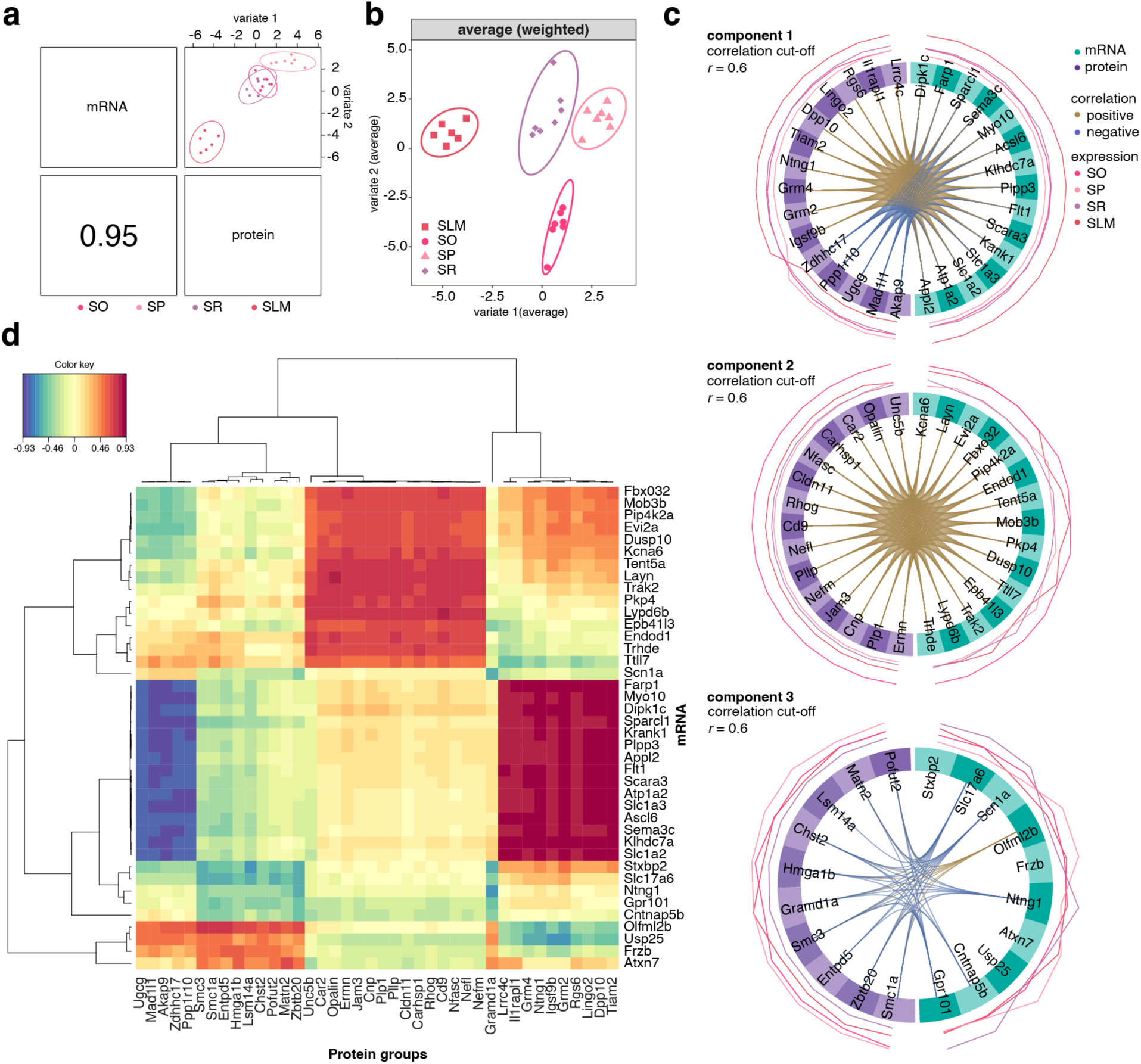
Exploration and integration of transcriptomic and proteomic databases after variable selection using mixOmics. (a) Results of DIABLO analysis to integrate transcriptomic and proteomic data. DIABLO plots depict clustering of samples based on the first component of each dataset. Pearson correlation coefficient (*r*) between the corresponding datasets as indicated and 95% confidence ellipse plots are represented. (b) PCA minimal set of features selected by DIABLO across data types (weighted average based on the first principal component of the data, in order to maximize the variance explained). (c) Circos plot shows the positive (or negative) correlation (*r* > 0.6) between selected features as indicated by the orange (or blue) links in Component 1-3 (left to right), feature names appear in the quadrants. (d) Unsupervised Clustered expression heatmap of the highly correlated features in the DIABLO sPLS-DA model (from 3 components).

## ACKNOWLEDGEMENTS

We thank Belquis Nassim-Assir, Marc van Oostrum, Susanne tom Dieck and the rest of the Schuman Lab and proteomics facility for technical support and assistance. The MPIBR support facilities, in particular Silke Zeissler and her team from Animal facility and Alla Schleif from media and glassware. E.K. acknowledges funding by EMBO (Postdoctoral Fellowship EMBO ALTF 148-2023). The lab of E.M.S. is funded by the Max Planck Society and the European Union (ERC, DiverseSynapse, 101054512). Views and opinions expressed are, however, those of the author(s) only and do not necessarily reflect those of the EU or the ERC. Neither the EU nor the granting authority can be held responsible for them.

