## Supplementary Table S1 for "The molecular diversity of hippocampal regions and strata at synaptic resolution revealed by integrated transcriptomic and proteomic profiling"

**GO_CA1_mRNA**

| ONTOLOGY | ID | Description | GeneRatio | BgRatio | pvalue | p.adjust | qvalue | geneID | Count |
|---|---|---|---|---|---|---|---|---|---|
| CC | GO:0045211 | postsynaptic membrane | 41/703 | 345/17474 | 5,40E+05 | 3,26E+10 | 2,81E+09 | Lin7a/Gabra1/Grin2c/Ntrk2/Hcn1/Adra1a/Slitrk1/Farp1/Arc/Grik1/Slc8a1/Kcnn2/Sorcs3/Chrm4/Lrrc4c/Chrm5/Ptprt/Chrna4/Adgrl2/Grin3a/Grm3/Htr5a/Adra2c/Drd5/Kctd8/Gabrg1/Adgrl3/Met/Cadps2/Chrm2/Grip2/Cacng7/Gabrg3/Efnb2/Ank1/Nrg1/Lzts1/Itgb1/Igsf9b/Chrna5/Gabra3 | 41 |
| CC | GO:0098984 | neuron to neuron synapse | 49/703 | 495/17474 | 5,95E+06 | 1,79E+11 | 1,55E+11 | Bcr/Anks1b/Actr2/Git1/Grin2c/Ntrk2/Homer1/Slitrk1/Ctnnd2/Mal2/Adcy8/Arc/Gap43/Grik1/Grm4/Slc8a1/Nr3c1/Entpd1/Sorcs3/Chrm4/Lrrc4c/Prnp/Plcb4/Ptprt/Ntsr1/Penk/Grin3a/Sh2d5/Grm3/Sema3a/Adra2c/Drd5/Met/Chrm2/Grip2/Itpr1/Cacng7/Homer2/Syt9/Adgra1/Efnb2/Nrg1/Zdhhc2/Lzts1/Itgb1/Igsf9b/Fam81a/Map4/Drp2 | 49 |
| CC | GO:0043235 | receptor complex | 41/703 | 397/17474 | 3,29E+08 | 3,58E+10 | 3,08E+11 | Vwc2l/Ptprb/Gabra1/Skap1/Plxdc1/Grin2c/Ahr/Ntrk2/Alcam/Grik1/Alk/Csf1r/Itga6/Ltk/Tyro3/Mertk/Bmp2/Chrna4/Pex5l/Olfm3/Cntfr/Grin3a/Tek/Cd36/Kctd8/Gabrg1/Flt1/Met/Plxnd1/Cacng7/Gabrg3/Arnt2/Il4ra/Adrb3/Itgb1/Olfm2/Chrna5/Htr1b/Mst1r/Gpr101/Gabra3 | 41 |
| CC | GO:0099572 | postsynaptic specialization | 45/703 | 480/17474 | 1,25E+08 | 4,98E+10 | 4,29E+10 | Bcr/Anks1b/Actr2/Gabra1/Git1/Grin2c/Ntrk2/Homer1/Slitrk1/Ctnnd2/Adcy8/Arc/Gap43/Grik1/Slc8a1/Nr3c1/Entpd1/Sorcs3/Chrm4/Lrrc4c/Prnp/Plcb4/Ptprt/Chrna4/Grin3a/Sh2d5/Grm3/Htr5a/Adra2c/Drd5/Met/Grip2/Itpr1/Cacng7/Homer2/Adgra1/Efnb2/Nrg1/Zdhhc2/Lzts1/Igsf9b/Fam81a/Map4/Gabra3/Drp2 | 45 |
| CC | GO:0005912 | adherens junction | 21/703 | 151/17474 | 7,02E+08 | 1,76E+11 | 1,52E+11 | Cdh20/Cdh7/Lin7a/Vcl/Fermt2/Cdh6/Cdh18/Ctnnd2/Rnd1/Dll1/Vegfa/Efna5/Ptprm/Jcad/Itga6/Cdh22/Cdh4/Plekha7/Kifc3/Itgb1/Myo1e | 21 |
| CC | GO:0034702 | ion channel complex | 31/703 | 295/17474 | 1,09E+10 | 2,52E+12 | 2,17E+12 | Vwc2l/Dpp10/Kcnip1/Gabra1/Kcnab3/Cacna1g/Grin2c/Fkbp1b/Kcns3/Hcn1/Clcn2/Grik1/Clic5/Kcnip3/Chrna4/Pex5l/Trpc4/Kcnab1/Olfm3/Grin3a/Gabrg1/Cacng7/Ttyh1/Slc17a6/Gabrg3/Olfm2/Chrna5/Cacna2d2/Scn5a/Gabra3/Kcne1l | 31 |
| CC | GO:0043679 | axon terminus | 23/703 | 200/17474 | 5,96E+10 | 8,98E+11 | 7,74E+11 | Glul/Nts/Git1/Ntrk2/Crhbp/Hcn1/Grik1/Grm4/Slc8a1/Chrm4/Kcnip3/Prnp/Ntsr1/Prss12/Penk/Adra2c/Chrm2/Prkcb/Pgr/Cplx3/Htr1b/Tpbg/Septin6 | 23 |
| CC | GO:0044306 | neuron projection terminus | 24/703 | 222/17474 | 1,09E+11 | 0.001288949295987809 | 0.00111030432172033 | Glul/Nts/Git1/Ntrk2/Crhbp/Hcn1/Grik1/Grm4/Slc8a1/Flrt1/Chrm4/Kcnip3/Prnp/Ntsr1/Prss12/Penk/Adra2c/Chrm2/Prkcb/Pgr/Cplx3/Htr1b/Tpbg/Septin6 | 24 |
| CC | GO:0005604 | basement membrane | 15/703 | 109/17474 | 3,05E+11 | 0.0026973163816954 | 0.002323475442335609 | Fn1/Timp2/Thbs4/Mmrn2/Matn2/Ccdc80/Vegfa/Efna5/Entpd1/Entpd2/Itga6/Fras1/Ptn/Itgb1/Egfl6 | 15 |
| CC | GO:0060076 | excitatory synapse | 14/703 | 98/17474 | 3,65E+10 | 0.002936014477500941 | 0.0025290906113607 | Git1/Ntrk2/Homer1/Adcy8/Grm4/Plcb4/Sema3a/Elfn1/Met/Cadps2/Cntn6/Slc17a6/Itgb1/Ntm | 14 |
| CC | GO:0019897 | extrinsic component of plasma membrane | 19/703 | 177/17474 | 9,68E+09 | 0.005215157105121305 | 0.004492350079472355 | Serpine2/Cdh20/Cdh7/Esyt2/Gng4/Fermt2/Farp1/Cdh6/Cdh18/Cdh22/Cdh4/Kcnab1/Wnt2/Gng12/Kras/Fes/Sytl2/Olfm2/Stac | 19 |
| CC | GO:1990351 | transporter complex | 32/703 | 397/17474 | 1,58E+12 | 0.0070480817892795465 | 0.006071236234686962 | Vwc2l/Dpp10/Kcnip1/Gabra1/Atp10b/Kcnab3/Cacna1g/Grin2c/Fkbp1b/Kcns3/Hcn1/Clcn2/Grik1/Clic5/Kcnip3/Chrna4/Pex5l/Trpc4/Kcnab1/Olfm3/Grin3a/Gabrg1/Cacng7/Ttyh1/Slc17a6/Gabrg3/Olfm2/Chrna5/Cacna2d2/Scn5a/Gabra3/Kcne1l | 32 |
| CC | GO:0030018 | Z disc | 15/703 | 128/17474 | 1,96E+12 | 0.008050394272120445 | 0.006934630849879809 | Mypn/Ppp1r12a/Fkbp1b/Rtl1/Homer1/Vcl/Myh6/Myh7/Adra1a/Slc8a1/Kcnn2/Jph2/Myo18b/Ank1/Scn5a | 15 |
| MF | GO:0030594 | neurotransmitter receptor activity | 18/703 | 92/17474 | 2,21E+07 | 3,58E+10 | 3,08E+11 | Htr5b/Gabra1/Grin2c/Hrh2/Grik1/Chrm4/Chrm5/Hrh3/Chrna4/Grin3a/Htr5a/Drd5/Gabrg1/Chrm2/Gabrg3/Chrna5/Htr1b/Gabra3 | 18 |
| MF | GO:0008227 | G protein-coupled amine receptor activity | 11/703 | 33/17474 | 3,56E+07 | 3,58E+10 | 3,08E+11 | Htr5b/Hrh2/Adra1a/Chrm4/Chrm5/Hrh3/Htr5a/Adra2c/Chrm2/Adrb3/Htr1b | 11 |
| MF | GO:0050839 | cell adhesion molecule binding | 33/703 | 298/17474 | 1,45E+09 | 5,16E+10 | 4,44E+10 | Fn1/Cdh20/Cdh7/F11r/Utrn/Mypn/Igf1/Ptprb/Timp2/Adam17/Thbs4/Fermt2/Cdh6/Cdh18/Ctnnd2/Ccn3/Ptprm/Itga6/Lrrc4c/Ptprt/Cdh22/Cdh4/Trpc4/Plpp3/Prom1/Ptn/Cntn6/Mfge8/Adam18/Nrg1/Tenm3/Itgb1/Egfl6 | 33 |
| MF | GO:0022839 | monoatomic ion gated channel activity | 31/703 | 302/17474 | 1,79E+09 | 3,73E+11 | 3,21E+12 | Kcnip1/Gabra1/Kcnj12/Kcnab3/Cacna1g/Grin2c/Kcns3/Hcn1/Clcn2/Grik1/Clic5/Kcnn2/Kcnt1/Kcnh7/Chrm5/Kcnip3/Chrna4/Pex5l/Kcnab1/Grin3a/Kcnk3/Gabrg1/Itpr1/Cacng7/Gabrg3/Chrna5/Htr1b/Cacna2d2/Scn5a/Gabra3/Kcne1l | 31 |
| MF | GO:0099589 | serotonin receptor activity | 8/703 | 24/17474 | 2,74E+09 | 5,00E+11 | 4,31E+12 | Htr5b/Hrh2/Chrm4/Chrm5/Hrh3/Htr5a/Chrm2/Htr1b | 8 |
| MF | GO:0005216 | monoatomic ion channel activity | 36/703 | 417/17474 | 1,47E+11 | 0.0016388718338946427 | 0.0014117285184010488 | Ano4/Kcnip1/Gabra1/Kcnj12/Kcnab3/Cacna1g/Grin2c/Kcns3/Hcn1/Apol8/Clcn2/Grik1/Clic5/Kcnn2/Kcnt1/Kcnh7/Chrm5/Kcnip3/Chrna4/Pex5l/Trpc4/Kcnab1/Grin3a/Kcnk3/Gabrg1/Itpr1/Cacng7/Ttyh1/Slc17a6/Gabrg3/Chrna5/Htr1b/Cacna2d2/Scn5a/Gabra3/Kcne1l | 36 |
| MF | GO:0004714 | transmembrane receptor protein tyrosine kinase | 11/703 | 58/17474 | 1,67E+10 | 0.001815950756779007 | 0.0015642647694204608 | Ntrk2/Ephb3/Alk/Csf1r/Ltk/Tyro3/Mertk/Tek/Flt1/Met/Mst1r | 11 |
| MF | GO:0044325 | transmembrane transporter binding | 18/703 | 151/17474 | 3,72E+10 | 0.0029515393197012173 | 0.0025424637513613564 | Dpp10/Kcnip1/Kcnab3/Fkbp1b/Homer1/Rims2/Fgf12/Slc8a1/Kcnip3/Prnp/Rims4/Kcnab1/Rnf207/Itpr1/Ank1/Stac/Scn5a/Kcne1l | 18 |
| MF | GO:0004879 | nuclear receptor activity | 9/703 | 47/17474 | 9,06E+10 | 0.005059118935304204 | 0.004357938388616444 | Rxrg/Ahr/Thrb/Nr4a1/Ppard/Nr3c1/Nr4a2/Pgr/Ar | 9 |
| MF | GO:0098531 | ligand-activated transcription factor activity | 9/703 | 47/17474 | 9,06E+10 | 0.005059118935304204 | 0.004357938388616444 | Rxrg/Ahr/Thrb/Nr4a1/Ppard/Nr3c1/Nr4a2/Pgr/Ar | 9 |
| MF | GO:0005178 | integrin binding | 17/703 | 150/17474 | 1,14E+12 | 0.005838314219485955 | 0.005029139260663223 | Fn1/F11r/Utrn/Igf1/Timp2/Adam17/Thbs4/Fermt2/Ccn3/Itga6/Plpp3/Ptn/Mfge8/Adam18/Nrg1/Itgb1/Egfl6 | 17 |
| MF | GO:0043178 | alcohol binding | 12/703 | 86/17474 | 1,63E+12 | 0.007123107283425625 | 0.006135863393139891 | Syt2/Adap2/Plcl2/Rbp4/Osbpl6/Trpc4/Astn2/Prom1/Osbpl3/Itpr1/Rlbp1/Osbpl10 | 12 |
| MF | GO:0034483 | heparan sulfate sulfotransferase activity | 5/703 | 15/17474 | 2,23E+12 | 0.008844843801715865 | 0.007618971769141865 | Hs6st3/Ndst3/Ndst4/Hs3st2/Hs6st2 | 5 |
| MF | GO:0016247 | channel regulator activity | 16/703 | 144/17474 | 2,27E+12 | 0.008937236947263229 | 0.0076985594682995396 | Bcl2/Dpp10/Stx7/Kcnip1/Kcnab3/Fkbp1b/Tspan13/Akt1/Fgf12/Kcnip3/Kcnab1/Grm3/Itpr1/Cacng7/Prkcb/Kcne1l | 16 |
| BP | GO:0016358 | dendrite development | 36/703 | 320/17474 | 2,62E+08 | 3,58E+10 | 3,08E+11 | Hecw2/Zfp365/Cobl/Actr2/Cdkl3/Git1/Sez6/Dcdc2a/Ntrk2/Plk2/Fezf2/Klhl1/Farp1/Ctnnd2/Matn2/Asap1/Arc/Ephb3/Alk/Nr3c1/Flrt1/Dab2ip/Itpka/Sh3glb1/Wls/Grin3a/Elavl4/Sema3a/Cit/Met/Ptn/Wnt7a/Arhgap33/Lzts1/Itgb1/Tpbg | 36 |
| BP | GO:0042391 | regulation of membrane potential | 46/703 | 480/17474 | 4,71E+08 | 3,97E+10 | 3,42E+10 | Bcl2/Dcn/Kcnip1/Gabra1/Sez6/Cacna1g/Grin2c/Fkbp1b/Akt1/Dcdc2a/Ntrk2/Plk2/Hcn1/Adra1a/Rims2/Fgf12/Grik1/Slc8a1/Kif5b/Kcnn2/Npas4/Kcnh7/Chrm5/Rims4/Ntsr1/Chrna4/Pex5l/Trpc4/Gpr88/Rnf207/Cd36/Kcnk3/Gabrg1/Tac1/Met/Ptn/Wnt7a/Grip2/Foxp1/Tspan9/Gabrg3/Igsf9b/Chrna5/Scn5a/Gabra3/Kcne1l | 46 |
| BP | GO:0007188 | adenylate cyclase-modulating G protein-coupled receptor signaling pathway | 28/703 | 217/17474 | 5,27E+07 | 3,97E+10 | 3,42E+10 | Htr5b/Gpr161/Vip/Adra1a/Npr3/Rims2/Adcy8/Grm4/Adgre1/Chrm4/Chrm5/Sstr4/Hrh3/Rxfp1/S1pr1/Adgrl2/Adgrl4/Htr5a/Adra2c/Drd5/Adgrl3/Grm8/Chrm2/Gipr/Adrb3/Pde4a/Htr1b/Gpr101 | 28 |
| BP | GO:0051932 | synaptic transmission, GABAergic | 15/703 | 68/17474 | 6,15E+06 | 4,12E+11 | 3,55E+10 | Gabra1/Adra1a/Grik1/Plcl2/Kif5b/Npas4/Gabrg1/Tac1/Kras/Gabrg3/Car7/Cntnap4/Htr1b/Tpbg/Gabra3 | 15 |
| BP | GO:0007611 | learning or memory | 35/703 | 320/17474 | 8,29E+07 | 4,29E+09 | 3,70E+11 | Vip/Igf1/Nts/Git1/Ppp1r1b/Dcdc2a/Shc3/Hrh2/Ntrk2/Plk2/Ctnnd2/Adcy8/Arc/Chrd/Grm4/Rcan2/Egr1/Npas4/Sorcs3/Nr4a2/Meis2/Prnp/Hrh3/Ntsr1/Kcnab1/Gpr88/Elavl4/Drd5/Tac1/Ptn/Kras/Nrg1/Itgb1/Aph1b/Tpbg | 35 |
| BP | GO:0048017 | inositol lipid-mediated signaling | 24/703 | 170/17474 | 8,53E+07 | 4,29E+09 | 3,70E+11 | Fn1/Serpine2/Nyap2/Pik3c2b/Igf1/Dcn/Pld2/Adap2/Akt1/Ntrk2/Pik3r1/Npr3/Ppard/Vegfa/Csf1r/Dab2ip/Plcb2/Ltk/Plcb4/Pdgfc/Tek/Flt1/Inpp5f/Dipk2a | 24 |
| BP | GO:0051962 | positive regulation of nervous system development | 38/703 | 369/17474 | 1,14E+09 | 4,98E+10 | 4,29E+10 | Fn1/Serpine2/Zfp365/Igf1/Actr2/Jade2/Cdkl3/Ndel1/Flrt2/Ntrk2/Pik3r1/Slitrk1/Ephb3/Etv5/Sox8/Vegfa/Efna5/Man2a1/Csf1r/Cbln2/Flrt1/Itpka/Bmp2/Cdh4/Sh3glb1/Adgrl2/Wls/Adgrl3/Flt1/Met/Wnt2/Ptn/Wnt7a/Plxnd1/Kras/Nrg1/Itgb1/Tpbg | 38 |
| BP | GO:0050803 | regulation of synapse structure or activity | 34/703 | 310/17474 | 1,19E+09 | 4,98E+10 | 4,29E+10 | Actr2/Arhgef15/Flrt2/Akt1/Ntrk2/Homer1/Pik3r1/Slitrk1/Farp1/Asap1/Arc/Ephb3/Etv5/Efna5/Cbln2/Flrt1/Dab2ip/Itpka/Prnp/Ptprt/Rims4/Adgrl2/Adgrl3/Sparcl1/Ptn/Wnt7a/Arhgap33/Slc17a6/Nrg1/Lzts1/Cntnap4/Itgb1/Il10ra/Tpbg | 34 |
| BP | GO:0050807 | regulation of synapse organization | 33/703 | 302/17474 | 1,99E+09 | 6,66E+09 | 5,74E+10 | Actr2/Arhgef15/Flrt2/Akt1/Ntrk2/Homer1/Pik3r1/Slitrk1/Farp1/Asap1/Arc/Ephb3/Etv5/Efna5/Cbln2/Flrt1/Dab2ip/Itpka/Prnp/Ptprt/Rims4/Adgrl2/Adgrl3/Sparcl1/Ptn/Wnt7a/Arhgap33/Nrg1/Lzts1/Cntnap4/Itgb1/Il10ra/Tpbg | 33 |
| BP | GO:1902074 | response to salt | 34/703 | 327/17474 | 4,23E+09 | 1,26E+12 | 1,09E+12 | Syt2/Pcdh15/Slc13a5/Ppp1r1b/Homer1/Crhbp/Hcn1/Adcy8/Ly6c1/Cpne8/Ly6g6e/Slc8a1/Egr1/Rasgrp2/Chrm4/Chrm5/Itpka/Hrh3/Chrna4/Tacr3/Cd36/Drd5/Adgrl3/Rasal1/Chrm2/Itpr1/Syt5/Gipr/Homer2/Syt9/Prkcb/Pgr/Htr1b/Scn5a | 34 |

| | | | | | | | | | |
|---|---|---|---|---|---|---|---|---|---|
| BP | GO:0034329 | cell junction assembly | 42/703 | 453/17474 | 4,47E+09 | 1,26E+12 | 1,09E+12 | Fn1/Tns1/Cdh20/Bcl2/Cdh7/F11r/Bcr/Gabra1/Arhgef15/Pecam1/Flrt2/Ntrk2/Pik3r1/Vcl/Fermt2/Slitrk1/Farp1/Cdh6/Cdh18/Ctnnd2/Ephb3/Gap43/Vegfa/Efna5/Cbln2/Npas4/Flrt1/Grem1/Cdh22/Cdh4/Adgrl2/Tek/Adgrl3/Micall2/Wnt7a/Plxnd1/Efnb2/Nrg1/Zdhhc2/Lzts1/Itgb1/Tpbg | 42 |
| BP | GO:0034332 | adherens junction organization | 12/703 | 57/17474 | 2,19E+10 | 4,30E+12 | 3,70E+11 | Cdh20/Cdh7/Vcl/Fermt2/Cdh6/Cdh18/Vegfa/Cdh22/Cdh4/Plekha7/Efnb2/Kifc3 | 12 |
| BP | GO:0010959 | regulation of metal ion transport | 39/703 | 445/17474 | 4,55E+09 | 7,60E+11 | 6,54E+11 | Hecw2/Serpine2/Bcl2/Dpp10/Vip/Igf1/Atp2b1/Kcnip1/Kcnab3/Fgf11/Cacna1g/Fkbp1b/Ahr/Tspan13/Akt1/Homer1/Mchr1/Fgf12/Slc8a1/Kif5b/Kcnn2/Grp/Kcnip3/Prnp/Plcb4/Jph2/Ntsr1/Trpc4/Kcnab1/Rnf207/Kcnk3/Wfs1/Itpr1/Homer2/Itgb1/Npsr1/Stac/Scn5a/Kcne1l | 39 |
| BP | GO:0099171 | presynaptic modulation of chemical synaptic | 12/703 | 61/17474 | 4,66E+10 | 7,60E+11 | 6,54E+11 | Git1/Adra1a/Chrd/Grik1/Grm4/Chrna4/Grin3a/Grm8/Chrm2/Prkcb/Chrna5/Htr1b | 12 |
| BP | GO:0051058 | negative regulation of small GTPase mediated | 12/703 | 62/17474 | 5,57E+08 | 8,84E+11 | 7,61E+10 | Git1/Timp2/Ripor2/Adra1a/Cyrib/Arhgap12/Dab2ip/Kctd10/Rasal1/Met/Kctd13/Itgb1 | 12 |
| BP | GO:0048639 | positive regulation of developmental growth | 24/703 | 216/17474 | 6,83E+09 | 9,80E+11 | 8,45E+09 | Fn1/Bcl2/Syt2/Igf1/Ndel1/Stat5b/Dio3/Akt1/Rims2/Dll1/Ppard/Vegfa/Efna5/Nr3c1/Cdh4/Sh3glb1/Pou3f2/Rasal1/Wnt2/Ybx3/Cacng7/Nrg1/Cacna2d2/Arx | 24 |
| BP | GO:0003018 | vascular process in circulatory system | 25/703 | 232/17474 | 7,60E+09 | 9,97E+11 | 8,59E+11 | Bcr/Atp2b1/Cacna1g/Ahr/Akt1/Hrh2/Fermt2/Adra1a/Npr3/Ext1/Ptp4a3/Ppard/Cbs/Vegfa/Ptprm/Slc8a1/Nr3c1/Ddah1/Cd36/Adra2c/Drd5/Grip2/Plekha7/Adrb3/Htr1b | 25 |
| BP | GO:0007626 | locomotory behavior | 27/703 | 262/17474 | 7,66E+09 | 9,97E+11 | 8,59E+11 | Npas2/Pcdh15/Git1/Sez6/Ppp1r1b/Hexb/Fezf2/Klhl1/Adcy8/Etv5/Fgf12/Rcan2/Alk/Egr1/Pbx3/Nr4a2/Ntsr1/Chrna4/Gpr88/Penk/Elavl4/Adgrl3/Lgi4/Inpp5f/Nrg1/Olfm2/Aph1b | 27 |
| BP | GO:1902075 | cellular response to salt | 22/703 | 189/17474 | 7,77E+09 | 9,97E+11 | 8,59E+11 | Syt2/Slc13a5/Ppp1r1b/Crhbp/Adcy8/Ly6c1/Cpne8/Ly6g6e/Egr1/Rasgrp2/Chrm4/Chrm5/Itpka/Hrh3/Chrna4/Rasal1/Chrm2/Itpr1/Syt5/Syt9/Prkcb/Scn5a | 22 |
| BP | GO:0098742 | cell-cell adhesion via plasma-membrane adhesion molecules | 23/703 | 206/17474 | 9,75E+08 | 0.0012002649808762558 | 0.0010339114188779388 | Cdh20/Cdh7/Tnfaip3/Mypn/Pcdh15/Pecam1/Slitrk1/Cdh6/Cdh18/Alcam/Efna5/Ptprm/Lrrc4c/Bmp2/Ptprt/Cdh22/Cdh4/Adgrl3/Sparcl1/Cntn6/Nrg1/Tenm3/Igsf9b | 23 |
| BP | GO:0043542 | endothelial cell migration | 24/703 | 222/17474 | 1,09E+11 | 0.001288949295987809 | 0.00111030432172033 | Glul/Igf1/Dcn/Pecam1/Adam17/Akt1/Plk2/Mmrn2/Ccn3/Ptp4a3/Nr4a1/Vegfa/Ptprm/Jcad/Dab2ip/Grem1/Tek/Plpp3/Met/Wnt7a/Foxp1/Plxnd1/Efnb2/Itgb1 | 24 |
| BP | GO:0034765 | regulation of monoatomic ion transmembrane transport | 40/703 | 480/17474 | 1,15E+10 | 0.0013300799168598622 | 0.0011457343469753443 | Hecw2/Bcl2/Dpp10/Kcnip1/Kcnj12/Kcnab3/Fgf11/Grin2c/Fkbp1b/Kcns3/Ahr/Tspan13/Akt1/Homer1/Hcn1/Arc/Clcn2/Fgf12/Clic5/Slc8a1/Kif5b/Kcnn2/Grp/Tcirg1/Kcnh7/Kcnip3/Prnp/Jph2/Ntsr1/Chrna4/Kcnab1/Rnf207/Kcnk3/Cacng7/Itgb1/Npsr1/Cacna2d2/Stac/Scn5a/Kcne1l | 40 |
| BP | GO:0001666 | response to hypoxia | 23/703 | 213/17474 | 1,69E+10 | 0.001815950756779007 | 0.0015642647694204608 | Bcl2/Cd34/Pld2/Adam17/Akt1/Ppard/Cbs/Vegfa/Slc8a1/Egr1/Nr4a2/Bmp2/Chrna4/Ddah1/Tek/Plk3/Kcnk3/Fosl2/Cpeb2/Ppargc1a/Flt1/Itpr1/Arnt2 | 23 |
| BP | GO:0050769 | positive regulation of neurogenesis | 29/703 | 308/17474 | 2,03E+09 | 0.0020891178166244757 | 0.0017995715949427338 | Fn1/Serpine2/Zfp365/Igf1/Actr2/Jade2/Cdkl3/Ndel1/Ntrk2/Slitrk1/Etv5/Sox8/Vegfa/Efna5/Man2a1/Csf1r/Itpka/Bmp2/Cdh4/Sh3glb1/Wls/Flt1/Met/Wnt2/Ptn/Plxnd1/Kras/Nrg1/Itgb1 | 29 |
| BP | GO:0060537 | muscle tissue development | 38/703 | 461/17474 | 2,41E+10 | 0.0023139089425376685 | 0.001993207263438539 | Col19a1/Bcl2/Igf1/Myl6b/Ahr/Rtl1/Ripor2/Homer1/Myh6/Myh7/Adra1a/Klf5/Nr4a1/Dll1/Sox8/Vegfa/Slc8a1/Egr1/Nr3c1/Rbp4/Grem1/Bmp2/Jph2/S1pr1/Ppargc1a/Myo18b/Met/Wnt2/Foxp1/Ybx3/Kras/Sox6/Efnb2/Nrg1/Acta1/Itgb1/AW551984/Dipk2a | 38 |
| BP | GO:0008217 | regulation of blood pressure | 22/703 | 203/17474 | 2,42E+10 | 0.0023139089425376685 | 0.001993207263438539 | F11r/Cd34/Atp2b1/Tac2/Smtn/Ahr/Adamts16/Pik3r1/Myh6/Adra1a/Npr3/Ext1/Dll1/Hrh3/Ntsr1/Tacr3/Ddah1/Drd5/Tac1/Grip2/Adrb3/Ar | 22 |
| BP | GO:0048754 | branching morphogenesis of an epithelial tube | 21/703 | 190/17474 | 2,74E+11 | 0.002503873907176669 | 0.00215684358479786 | Bcl2/Pbx1/Igf1/Ahr/Adamts16/Rspo2/Ext1/Etv5/Sox8/Vegfa/Grem1/Bmp2/Tek/Flt1/Met/Wnt2/Plxnd1/Kras/Sall1/Pgr/Ar | 21 |
| BP | GO:0031346 | positive regulation of cell projection organization | 37/703 | 449/17474 | 3,10E+10 | 0.0026973163816954 | 0.002323475442335609 | Fn1/Serpine2/Cobl/Actr2/Cdkl3/Ndel1/Fkbp1b/Adam17/Ripor2/Ntrk2/Pik3r1/Slitrk1/Dzip1/Vegfa/Efna5/Alk/Rhoq/Dab2ip/Itga6/Itpka/Ltk/Cdh4/Serpini1/Fnbp1l/Sh3glb1/Wls/Tox/Elavl4/Met/Ptn/Itpr1/Plxnd1/Fes/Nrg1/Tenm3/Itgb1/Rab8b | 37 |
| BP | GO:0001503 | ossification | 34/703 | 398/17474 | 3,13E+11 | 0.0026973163816954 | 0.002323475442335609 | Nab1/Satb2/Bcl2/Pbx1/Igf1/Atp2b1/Git1/Sirt7/Ahr/Dlk1/Akt1/Thrb/Fermt2/Rspo2/Ext1/Chrd/Sox8/Cbs/Vegfa/Slc8a1/Fbn2/Csf1r/Tcirg1/Grem1/Bmp2/Intu/S1pr1/Bmp3/Dmp1/Tac1/Ptn/Clec11a/Sik3/Gdpd2 | 34 |
| BP | GO:0035265 | organ growth | 22/703 | 207/17474 | 3,27E+11 | 0.0027773799955711077 | 0.0023924424504060235 | Bcl2/Igf1/Dusp6/Ahr/Akt1/Myh6/Adra1a/Rspo2/Ext1/Dll1/Nr3c1/Rbp4/S1pr1/Wnt2/Foxp1/Ybx3/Nrg1/Wwc2/Dipk2a/Cacna2d2/Arx/Ar | 22 |
| BP | GO:0043087 | regulation of GTPase activity | 31/703 | 349/17474 | 3,32E+11 | 0.0027802218889665087 | 0.0023948904648691243 | Sh3bp4/F11r/Bcr/Arhgap45/Ndel1/Arhgef15/Adap2/Pecam1/Asap2/Ripor2/Ntrk2/Fermt2/Asap1/Ephb3/Efna5/Arhgap12/Rasgrp2/Vav2/Garnl3/Dab2ip/Itga6/S1pr1/Cpeb2/Tbc1d1/Rasgef1b/Rasal1/Met/Plxnd1/Itgb1/Dock6/Tbc1d2b | 31 |
| BP | GO:0098926 | postsynaptic signal transduction | 11/703 | 63/17474 | 3,77E+10 | 0.002953933457864507 | 0.002544526068287065 | Anks1b/Ly6c1/Ly6g6e/Chrm4/Chrm5/Hrh3/Chrna4/Chrm2/Itpr1/Prkcb/Nrg1 | 11 |
| BP | GO:0060042 | retina morphogenesis in camera-type eye | 12/703 | 75/17474 | 4,17E+11 | 0.0031883030547185612 | 0.0027464126569037906 | Dio3/Ntrk2/Hcn1/Thrb/Dll1/Sox8/Man2a1/Ptprm/Rbp4/Prom1/Tspan12/Ptn | 12 |
| BP | GO:0044331 | cell-cell adhesion mediated by cadherin | 9/703 | 43/17474 | 4,33E+10 | 0.003263463083860242 | 0.002811155704156615 | Cdh20/Cdh7/Cdh6/Cdh18/Vegfa/Cdh22/Cdh4/Ptpru/Plekha7 | 9 |
| BP | GO:0051146 | striated muscle cell differentiation | 29/703 | 323/17474 | 4,87E+10 | 0.0035811161378719782 | 0.003084782882335568 | Bcl2/Mypn/Igf1/Akt1/Ripor2/Homer1/Myh6/Myh7/Adra1a/Klf5/Ccn3/Dll1/Vegfa/Slc8a1/Nr3c1/Csf1r/Grem1/Bmp2/Myo18b/Met/Foxp1/Kras/Sox6/Il4ra/Efnb2/Nrg1/Acta1/Itgb1/AW551984 | 29 |
| BP | GO:0007416 | synapse assembly | 22/703 | 213/17474 | 5,06E+11 | 0.00363178107960871 | 0.003128425796719482 | Gabra1/Arhgef15/Flrt2/Ntrk2/Pik3r1/Slitrk1/Farp1/Ephb3/Gap43/Efna5/Cbln2/Npas4/Flrt1/Adgrl2/Adgrl3/Wnt7a/Plxnd1/Efnb2/Nrg1/Zdhhc2/Lzts1/Tpbg | 22 |
| BP | GO:0099565 | chemical synaptic transmission, postsynaptic | 15/703 | 114/17474 | 5,20E+10 | 0.003639280214993795 | 0.003134885571710743 | Anks1b/Sez6/Grin2c/Plk2/Rims2/Grik1/Npas4/Chrm5/Ntsr1/Chrna4/Met/Wnt7a/Grip2/Igsf9b/Gabra3 | 15 |
| BP | GO:0042220 | response to cocaine | 9/703 | 44/17474 | 5,25E+10 | 0.003639280214993795 | 0.003134885571710743 | Ppp1r1b/Homer1/Crhbp/Tacr3/Drd5/Adgrl3/Homer2/Pgr/Htr1b | 9 |
| BP | GO:0043266 | regulation of potassium ion transport | 15/703 | 115/17474 | 5,76E+10 | 0.0038616730937418037 | 0.003326455383775775 | Dpp10/Vip/Kcnip1/Kcnab3/Kif5b/Kcnn2/Grp/Kcnip3/Prnp/Plcb4/Kcnab1/Rnf207/Kcnk3/Itgb1/Kcne1l | 15 |
| BP | GO:1903522 | regulation of blood circulation | 25/703 | 262/17474 | 5,99E+10 | 0.003872443155282894 | 0.0033357327431812137 | Atp2b1/Smtn/Cacna1g/Fkbp1b/Ahr/Akt1/Hrh2/Pik3r1/Hcn1/Thrb/Myh6/Myh7/Adra1a/Tmem65/Slc8a1/Kcnn2/Tacr3/Rnf207/Sema3a/Adra2c/Tac1/Chrm2/Cacna2d2/Scn5a/Kcne1l | 25 |
| BP | GO:0035296 | regulation of tube diameter | 19/703 | 171/17474 | 6,07E+10 | 0.003872443155282894 | 0.0033357327431812137 | Atp2b1/Cacna1g/Ahr/Akt1/Hrh2/Adra1a/Npr3/Ext1/Ppard/Cbs/Vegfa/Ptprm/Slc8a1/Adra2c/Drd5/Grip2/Plekha7/Adrb3/Htr1b | 19 |
| BP | GO:0097746 | blood vessel diameter maintenance | 19/703 | 171/17474 | 6,07E+10 | 0.003872443155282894 | 0.0033357327431812137 | Atp2b1/Cacna1g/Ahr/Akt1/Hrh2/Adra1a/Npr3/Ext1/Ppard/Cbs/Vegfa/Ptprm/Slc8a1/Adra2c/Drd5/Grip2/Plekha7/Adrb3/Htr1b | 19 |
| BP | GO:0001763 | morphogenesis of a branching structure | 24/703 | 247/17474 | 6,28E+10 | 0.003895889431157599 | 0.0033559294270329096 | Bcl2/Pbx1/Rspo3/Igf1/Ahr/Adamts16/Ctnnd2/Rspo2/Ext1/Etv5/Sox8/Vegfa/Grem1/Bmp2/Tek/Sema3a/Flt1/Met/Wnt2/Plxnd1/Kras/Sall1/Pgr/Ar | 24 |
| BP | GO:0030534 | adult behavior | 20/703 | 188/17474 | 7,27E+10 | 0.0043830937059583875 | 0.003775608473800512 | Pcdh15/Sez6/Ppp1r1b/Homer1/Crhbp/Klhl1/Slitrk1/Fgf12/Grik1/Alk/Pbx3/Nr4a2/Ntsr1/Chrna4/Aldh2/Met/Lgi4/Homer2/Inpp5f/Chrna5 | 20 |
| BP | GO:0042063 | gliogenesis | 31/703 | 366/17474 | 8,12E+10 | 0.004800725545104647 | 0.004135357641076124 | Nab1/Fn1/Serpine2/Zfp365/Igf1/Ptprb/Akt1/Ntrk2/Hexb/Zcchc24/Matn2/Etv5/Gap43/Dll1/Sox8/Nr3c1/Csf1r/Dab2ip/Bmp2/Trpc4/Pou3f2/Plpp3/Pou3f1/Flt1/Ptn/Kras/Mboat7/Lgi4/Idh2/Sox6/Nrg1 | 31 |
| BP | GO:0030336 | negative regulation of cell migration | 27/703 | 300/17474 | 8,43E+10 | 0.004889128342156872 | 0.004211508043520023 | Bcl2/Bcr/Dcn/Srgap1/Evl/Akt1/Ripor2/Mmrn2/Ccn3/Chrd/Dusp1/Ppard/Ptprm/Ldlrad4/Dab2ip/Grem1/Ptprt/Podn/Ptpru/Sema3a/Drd5/Ppargc1a/Ptn/Arhgdib/Idh2/Ing1/Nrg1 | 27 |

| | | | | | | | | |
|---|---|---|---|---|---|---|---|---|
| BP | GO:0090132 | epithelium migration | 28/703 | 319/17474 | 9,67E+10 | 0.005215157105121305 | 0.004492350079472355 | Glul/Igf1/Dcn/Pecam1/Adam17/Evl/Akt1/Plk2/Mmrn2/Ccn3/Ptp4a3/Nr4a1/Ppard/Vegfa/Ptprm/Jcad/Dab2ip/Grem1/Tek/Plpp3/Sema3a/Tac1/Met/Wnt7a/Foxp1/Plxnd1/Efnb2/Itgb1 | 28 |
| BP | GO:0055007 | cardiac muscle cell differentiation | 17/703 | 148/17474 | 9,68E+10 | 0.005215157105121305 | 0.004492350079472355 | Igf1/Myh6/Adra1a/Dll1/Vegfa/Slc8a1/Nr3c1/Grem1/Bmp2/Myo18b/Met/Foxp1/Sox6/Efnb2/Nrg1/Itgb1/AW551984 | 17 |
| BP | GO:0043954 | cellular component maintenance | 12/703 | 82/17474 | 1,02E+12 | 0.00542036325455005 | 0.0046691151975910734 | Homer1/Fermt2/Rims2/Csf1r/Cbln2/Itpka/Prnp/Sema3a/Adgrl3/Plekha7/Kifc3/Itgb1 | 12 |
| BP | GO:0045927 | positive regulation of growth | 27/703 | 304/17474 | 1,05E+12 | 0.005476132409613342 | 0.004717154894791117 | Fn1/Bcl2/Syt2/Igf1/Ndel1/Stat5b/Adam17/Ahr/Dio3/Akt1/Rims2/Dll1/Ppard/Vegfa/Efna5/Nr3c1/Cdh4/Sh3glb1/Pou3f2/Wfs1/Rasal1/Wnt2/Ybx3/Cacng7/Nrg1/Cacna2d2/Arx | 27 |
| BP | GO:0007411 | axon guidance | 23/703 | 241/17474 | 1,16E+12 | 0.005838314219485955 | 0.005029139260663223 | Mypn/Etv1/Flrt2/Evl/Fezf2/Matn2/Ext1/Ephb3/Gap43/Alcam/Vegfa/Efna5/Ptprm/Kif5b/Csf1r/Cdh4/Sema3a/Foxp1/Cntn6/Cntn4/Plxnd1/Efnb2/Arx | 23 |
| BP | GO:1905330 | regulation of morphogenesis of an epithelium | 11/703 | 72/17474 | 1,33E+12 | 0.006370110314346385 | 0.005487229818791054 | Mmrn2/Etv5/Sox8/Vegfa/Grem1/Rnf207/Met/Wnt2/Foxp1/Sall1/Ar | 11 |
| BP | GO:1990778 | protein localization to cell periphery | 31/703 | 377/17474 | 1,39E+12 | 0.006512307810202509 | 0.005609719116607743 | Acsl3/Dpp10/F11r/Stx7/Epb41l2/Lin7a/Git1/Skap1/Grin2c/Rhbdf2/Ttc7b/Akt1/Pik3r1/Rab3c/Tspan14/Rhoq/Kif5b/Kcnip3/Prnp/Rab3b/Grip2/Cacng7/Rab38/Sytl2/Ank1/Zdhhc2/Itgb1/Rab8b/Tpbg/Stac/Ar | 31 |
| BP | GO:1905144 | response to acetylcholine | 9/703 | 50/17474 | 1,50E+11 | 0.006758522266536169 | 0.005821808898407718 | Ly6c1/Ly6g6e/Chrm4/Chrm5/Hrh3/Chrna4/Chrm2/Itpr1/Prkcb | 9 |
| BP | GO:0015012 | heparan sulfate proteoglycan biosynthetic process | 7/703 | 30/17474 | 1,50E+12 | 0.006758522266536169 | 0.005821808898407718 | Hs6st3/Ext1/Ndst3/Ndst4/Hs3st2/Lipc/Hs6st2 | 7 |
| BP | GO:0044706 | multi-multicellular organism process | 18/703 | 169/17474 | 1,62E+12 | 0.007123107283425625 | 0.006135863393139891 | Serpine2/Havcr2/Stat5b/Akt1/Crhbp/Hexb/Ppard/Cbs/Vegfa/Fbn2/Polr1b/Thbd/Rxfp1/Adra2c/Ptn/Arhgdib/Pgr/Ar | 18 |
| BP | GO:1901890 | positive regulation of cell junction assembly | 15/703 | 126/17474 | 1,64E+12 | 0.007123107283425625 | 0.006135863393139891 | Flrt2/Ntrk2/Pik3r1/Fermt2/Slitrk1/Ephb3/Vegfa/Efna5/Cbln2/Flrt1/Adgrl2/Tek/Adgrl3/Wnt7a/Tpbg | 15 |
| BP | GO:0019226 | transmission of nerve impulse | 13/703 | 99/17474 | 1,65E+12 | 0.007123107283425625 | 0.006135863393139891 | Cacna1g/Fkbp1b/Dcdc2a/Ntrk2/Hcn1/Fgf12/Chrm5/Chrna4/Gpr88/Drd5/Cacng7/Chrna5/Scn5a | 13 |
| BP | GO:1900746 | regulation of vascular endothelial growth factor | 6/703 | 22/17474 | 1,78E+12 | 0.007464402308286624 | 0.006429855827337549 | Dcn/Ptp4a3/Dll1/Vegfa/Jcad/Dab2ip | 6 |
| BP | GO:0060047 | heart contraction | 22/703 | 232/17474 | 1,78E+12 | 0.007464402308286624 | 0.006429855827337549 | Smtn/Cacna1g/Fkbp1b/Pik3r1/Hcn1/Thrb/Myh6/Myh7/Adra1a/Ext1/Tmem65/Slc8a1/Kcnn2/Tacr3/Rnf207/Sema3a/Tac1/Met/Chrm2/Cacna2d2/Scn5a/Kcne1l | 22 |
| BP | GO:0001764 | neuron migration | 20/703 | 201/17474 | 1,82E+10 | 0.007582151266330742 | 0.006531285090227412 | Satb2/Ndel1/Adam17/Flrt2/Dcdc2a/Ntrk2/Fezf2/Matn2/Nde1/Vegfa/Dab2ip/Nr4a2/Tyro3/Astn2/Sema3a/Adgrl3/Met/Nrg1/Sh3rf1/Arx | 20 |
| BP | GO:0044703 | multi-organism reproductive process | 17/703 | 156/17474 | 1,85E+11 | 0.007643196792908622 | 0.006583869867760628 | Serpine2/Havcr2/Stat5b/Akt1/Crhbp/Ppard/Cbs/Vegfa/Fbn2/Polr1b/Thbd/Rxfp1/Adra2c/Ptn/Arhgdib/Pgr/Ar | 17 |

**GO_CA3_mRNA**

| ONTOLOGY | ID | Description | GeneRatio | BgRatio | pvalue | p.adjust | qvalue | geneID | Count |
|---|---|---|---|---|---|---|---|---|---|
| BP | GO:0099003 | vesicle-mediated transport in synapse | 94/1983 | 269/17474 | 8,4E-10 | 5,3379E-06 | 4,69445E-06 | Atp6v1h/Rims1/Kcnh1/Cd24a/Ap3d1/Pip5k1c/Slc17a8/Ppfia2/Syt1/Ppp3r1/Canx/Snap47/Vamp2/Atp6v0a1/Itgb3/Prkca/Actg1/Psen1/Amph/Cplx2/Cadps/Ppp3cb/Erc2/Fgf14/Atp6v1c1/Septin5/Ap2m1/Fbxo45/Atp6v1a/Synj1/Pacsin1/Atp6v1g2/Nrxn1/Cdh2/Bin1/Myo5b/Syt7/Slc18a2/Dnm1/Stxbp1/Rapgef4/Snap25/Syndig1/Dnajc5/Nlgn1/Syt11/Sv2a/Sh3gl2/Plaa/Dnajc6/Eps15/Hpca/Pclo/Nsg1/Cplx1/Rph3a/P2rx7/Rimbp2/Stx1a/Atp6v0e2/Scrn1/Snca/Aak1/Syn2/Erc1/Atp6v1e1/Necap1/Vamp1/Napa/Ap2s1/Grik5/Slc17a7/Ppfia3/Cyfip1/Sv2b/Ap3b2/Sh3gl3/Prrt2/Ctbp2/Ap2a2/Ap3m2/Atp6v1b2/Rab3a/Prkaca/Cacna1a/Vps35/Atp6v0d1/Vac14/Scamp5/Snap91/Bsn/Syp/Atp6ap2/Syn1 | 94 |
| BP | GO:0099504 | synaptic vesicle cycle | 85/1983 | 229/17474 | 1,4E-08 | 5,3379E-06 | 4,69445E-06 | Atp6v1h/Rims1/Kcnh1/Cd24a/Ap3d1/Pip5k1c/Slc17a8/Ppfia2/Syt1/Ppp3r1/Canx/Snap47/Vamp2/Atp6v0a1/Prkca/Actg1/Psen1/Amph/Cplx2/Cadps/Ppp3cb/Erc2/Fgf14/Atp6v1c1/Septin5/Ap2m1/Fbxo45/Atp6v1a/Synj1/Pacsin1/Atp6v1g2/Nrxn1/Cdh2/Bin1/Myo5b/Syt7/Slc18a2/Dnm1/Stxbp1/Rapgef4/Snap25/Syndig1/Dnajc5/Nlgn1/Syt11/Sv2a/Sh3gl2/Plaa/Dnajc6/Pclo/Cplx1/Rph3a/P2rx7/Rimbp2/Stx1a/Atp6v0e2/Scrn1/Snca/Syn2/Erc1/Atp6v1e1/Vamp1/Napa/Ap2s1/Grik5/Slc17a7/Ppfia3/Cyfip1/Sv2b/Ap3b2/Sh3gl3/Prrt2/Ctbp2/Ap3m2/Atp6v1b2/Rab3a/Prkaca/Cacna1a/Atp6v0d1/Scamp5/Snap91/Bsn/Syp/Atp6ap2/Syn1 | 85 |
| BP | GO:0007409 | axonogenesis | 106/1983 | 482/17474 | 978,537 | 5416902,528 | 4763931,255 | Trak2/Nrp2/Klf7/Epha4/Cxcr4/Tgfb2/Ust/Kifbp/Adarb1/Map2k2/Pip5k1c/Rab21/Arhgef25/Nefh/Rtn4/Fstl4/Pafah1b1/Rtn4rl1/Pitpna/Cntnap1/Mapt/Prkca/Nptx1/Rab10/Crppa/Psen1/Hsp90aa1/Nrn1/Map1b/Emb/Nefl/Nefm/Sema5a/Cntn1/Rtn4r/Fbxo45/Sema5b/Epha3/Robo1/Hsp90ab1/Sema6b/Cdh2/B4galt6/Wdr36/Dcc/Myo5b/Rnf165/B4gat1/Prkg1/Lgi1/Slit1/Zfyve27/Ablim1/Vim/Olfm1/Stxbp1/Kif5c/Tbr1/Lrp2/Chn1/Bdnf/Ttl/Flrt3/Ythdf1/Bhlhe22/Unc5c/Lmo4/Epha7/Nr4a3/Nfib/Ephb2/Sema3e/Slit2/Apbb2/Uchl1/Epha5/Rufy3/Prdm8/Auts2/Rnf6/Sema4f/Mgll/Atg7/Lrtm2/Ntf3/Mag/Scn1b/Cyfip1/Igf1r/Pak1/Apbb1/Unc5d/Rab3a/Cacna1a/Nrp1/Aplp2/Nectin1/Thy1/Islr2/Nptn/Map2k1/Snap91/Ephb1/Slc9a6/Fgf13/Gdi1 | 106 |
| BP | GO:0006887 | exocytosis | 88/1983 | 376/17474 | 18812,1 | 9112087,441 | 8013686,409 | Rims1/Vps4b/Kcnh1/Pip5k1c/Ppfia2/Syt1/Rab21/Rab3ip/Snap47/Vamp2/Srcin1/Nsf/Prkca/Hgs/Rab10/Rab15/Psen1/Chga/Vps41/Cplx2/Nkd2/Scamp1/Cadps/Ppp3cb/Erc2/Lgi3/Ywhaz/Pla2g6/Syngr1/Septin5/Fbxo45/Synj1/Washc1/Myo5b/Rab3il1/Syt7/Anxa1/Pi4k2a/Slc18a2/Stam/Stxbp1/Rapgef4/Syt13/Bloc1s6/Snap25/Atp9a/Nlgn1/Syt11/Sv2a/Sdcbp/Nr4a3/Pclo/Kit/Cplx1/Rph3a/P2rx7/Rimbp2/Stx1a/Scrn1/Snca/Rab7/Arl8b/Syn2/Cacna1c/Erc1/Vamp1/D6Wsu163e/Napa/Grik5/Syt3/Ppfia3/Sphk2/Sv2b/Pak1/Prrt2/Ctbp2/Rab3a/Prkaca/Cacna1a/Gnao1/Smpd3/Exoc8/Exph5/Scamp5/Syp/Cdk16/Syn1/Sytl4 | 88 |
| BP | GO:0016079 | synaptic vesicle exocytosis | 40/1983 | 115/17474 | 288024 | 131304856,6 | 115476936,7 | Rims1/Kcnh1/Ppfia2/Syt1/Snap47/Vamp2/Prkca/Psen1/Cplx2/Cadps/Erc2/Septin5/Fbxo45/Synj1/Syt7/Stxbp1/Snap25/Nlgn1/Syt11/Sv2a/Pclo/Cplx1/Rph3a/P2rx7/Rimbp2/Stx1a/Snca/Erc1/Vamp1/Napa/Grik5/Ppfia3/Sv2b/Prrt2/Ctbp2/Rab3a/Prkaca/Cacna1a/Syp/Syn1 | 40 |
| BP | GO:0006836 | neurotransmitter transport | 61/1983 | 225/17474 | 46776,8 | 201400035,7 | 177122612 | Rims1/Kcnh1/Slc17a8/Slc6a15/Ppfia2/Syt1/Snap47/Vamp2/Itgb3/Prkca/Psen1/Nrxn3/Cplx2/Mctp1/Cadps/Erc2/Ptger4/Septin5/Fbxo45/Synj1/Slc29a1/Nrxn1/Slc6a7/Nrxn2/Syt7/Slc18a2/Stxbp1/Snap25/Nlgn1/Syt11/Sv2a/Slc6a17/Htr6/Pclo/Cplx1/Rph3a/P2rx7/Rimbp2/Stx1a/Slc29a4/Snca/Arl6ip5/Syn2/Erc1/Vamp1/Napa/Grik5/Kcnc3/Slc17a7/Ppfia3/Sv2b/Pak1/Prrt2/Ctbp2/Rab3a/Prkaca/Cacna1a/Snap91/Syp/Syn1/Gpm6b | 61 |
| BP | GO:0099643 | signal release from synapse | 50/1983 | 169/17474 | 90621,3 | 33265409,27 | 292554872,7 | Rims1/Kcnh1/Ppfia2/Syt1/Snap47/Vamp2/Prkca/Psen1/Nrxn3/Cplx2/Mctp1/Cadps/Erc2/Ptger4/Septin5/Fbxo45/Synj1/Nrxn1/Nrxn2/Syt7/Stxbp1/Snap25/Nlgn1/Syt11/Sv2a/Htr6/Pclo/Cplx1/Rph3a/P2rx7/Rimbp2/Stx1a/Snca/Syn2/Erc1/Vamp1/Napa/Grik5/Kcnc3/Ppfia3/Sv2b/Pak1/Prrt2/Ctbp2/Rab3a/Prkaca/Cacna1a/Snap91/Syp/Syn1 | 50 |
| BP | GO:0001505 | regulation of neurotransmitter levels | 62/1983 | 241/17474 | 339921 | 1053754014 | 92673103,43 | Rims1/Kcnh1/Slc17a8/Ppfia2/Syt1/Snap47/Vamp2/Itgb3/Prkca/Psen1/Nrxn3/Cplx2/Agtpbp1/Mctp1/Cadps/Erc2/Ptger4/Septin5/Fbxo45/Synj1/Slc44a4/Slc29a1/Nrxn1/Nrxn2/Dagla/Syt7/Slc18a2/Stxbp1/Snap25/Nlgn1/Syt11/Sv2a/Htr6/Pclo/Cplx1/Pebp1/Rph3a/P2rx7/Rimbp2/Stx1a/Slc29a4/Snca/Arl6ip5/Syn2/Erc1/Vamp1/Napa/Grik5/Kcnc3/Slc17a7/Ppfia3/Sv2b/Pak1/Prrt2/Ctbp2/Rab3a/Prkaca/Cacna1a/Snap91/Syp/Syn1/Gpm6b | 62 |
| BP | GO:0016050 | vesicle organization | 79/1983 | 349/17474 | 1080786 | 3221572511 | 283323355 | Atp6v1h/Rims1/Vps4b/Klhl12/Tmem9/Sec16b/Sar1a/Ap3d1/Washc4/Syt1/Pmel/Ap1b1/Rab1a/Sqstm1/Septin8/Snap47/Vamp2/Pafah1b1/Atp6v0a1/Vps41/Cplx2/Tmed9/Nkd2/Cadps/Erc2/Chmp7/Atp6v1c1/Trappc9/Mapk15/Ap2m1/Atp6v1a/Synj1/Bace2/Atp6v1g2/Washc1/Pik3c3/Syt7/Anxa1/Pi4k2a/Hps6/Stam/Dnm1/Stxbp1/Rab14/Bloc1s6/Snap25/Syt11/Sdcbp/Tyrp1/AU040320/Insig1/Cplx1/P2rx7/Rimbp2/Stx1a/Atp6v0e2/Snx10/Snca/Rab7/Arl8b/Tmcc1/Erc1/Atp6v1e1/Vamp1/Grik5/Ankrd27/Syt3/Slc17a7/Ap3b2/Aqp11/Prrt2/Ap3m2/Atp6v1b2/Rab3a/Atp6v0d1/Exoc8/Rnf26/Snap91/Atp6ap2 | 79 |
| BP | GO:0048499 | synaptic vesicle membrane organization | 18/1983 | 35/17474 | 6099228 | 15248069135 | 13410016658 | Rims1/Ap3d1/Syt1/Snap47/Cplx2/Erc2/Syngr1/Syt7/Stxbp1/Snap25/Cplx1/Rimbp2/Stx1a/Erc1/Vamp1/Grik5/Prrt2/Rab3a | 18 |

| | | | | | | | | | |
|---|---|---|---|---|---|---|---|---|---|
| BP | GO:0048167 | regulation of synaptic plasticity | 59/1983 | 243/17474 | 945832 | 22581317151 | 19859290806 | Stau2/Rims1/Epha4/Adora1/Ptgs2/Psen2/Grik2/Camk2b/Sqstm1/Snap47/Shisa6/Vamp2/Neurod2/Mapt/Psen1/Scgn/Cplx2/Rgs14/Rasgrf2/Abhd6/Erc2/Ptk2b/Fgf14/Syngr1/Mapk1/Neto1/Cfl1/Syt7/Htr7/Cpeb3/Neurl1a/Nsmf/Stxbp1/Bdnf/Snap25/Ythdf1/Nlgn1/Rapgef2/Musk/Slc24a2/Ncdn/Ephb2/Htr6/Nsg1/Kit/Ywhag/Snca/Mgll/Erc1/Slc8a2/Ppfia3/Cpeb1/Pak1/Prrt2/Rab3a/Nrgn/Nptn/Syp/Syn1 | 59 |
| BP | GO:0008088 | axo-dendritic transport | 30/1983 | 87/17474 | 1,1E+07 | 24102870062 | 21197430716 | Stau2/Trak2/Arl8a/Wasf1/Ap3d1/Rab21/Fbxw11/Pafah1b1/Flot2/Mapt/Klc1/Agtpbp1/Nefl/Nefm/Sybu/Neto1/Cnih2/Kif5c/Madd/Bloc1s6/Kif3b/Uchl1/Arl8b/Borcs5/Ap3b2/Ap3m2/Hspa8/Tmem108/Bsn/Armcx3 | 30 |
| BP | GO:0032409 | regulation of transporter activity | 70/1983 | 311/17474 | 1,2E+08 | 27130066454 | 23859718885 | Atp1b1/Hk1/Jsrp1/Kcnc2/Ppp3r1/Shisa6/Vamp2/Ywhae/Asic2/Vmp1/Stac2/Taco1/Cacng5/Prkca/Osr1/Twist1/Hspa2/Chrm3/Actn2/Ubqln1/Glrx/Rasgrf2/Ppp3cb/Prkcd/Ptk2b/Fgf14/Kcns2/Cacng2/Pla2g6/Mapk8ip2/Cacnb3/Ifngr2/Ehd3/Nrxn1/Ndfip1/Neto1/Cnih2/Cfl1/Gsto1/Lrrc55/Plcb1/Oprl1/Nlgn1/Asph/Ptpn3/Hpca/Ephb2/Ywhah/Slc5a1/Stim2/Cabp1/Tesc/Rph3a/Kctd7/Tcaf1/Snca/Kcna1/Calm3/Fxyd7/Scn1b/Sphk2/Abcc8/Kcnc1/P2ry6/Cacna1a/Neto2/Scn2b/Htr3a/Stoml1/Fgf13 | 70 |
| BP | GO:0031338 | regulation of vesicle fusion | 17/1983 | 34/17474 | 2,8E+08 | 5905240248 | 5193403139 | Rims1/Syt1/Cplx2/Erc2/Syt7/Anxa1/Stxbp1/Cplx1/Rimbp2/Snca/Erc1/Vamp1/Grik5/Ankrd27/Syt3/Prrt2/Rab3a | 17 |
| BP | GO:0007611 | learning or memory | 70/1983 | 320/17474 | 4,1E+07 | 8019818129 | 70530828365 | Chst10/Ptgs2/Abl2/Psen2/Sgk1/Tafa2/Vdac1/Pafah1b1/Serpinf1/Neurod2/Mapt/Taco1/Prkca/Lamb1/Psen1/Nrxn3/Amph/Rgs14/Gmfb/Pla2g6/Mapk8ip2/Tuba1a/Synj1/Glp1r/Pja2/Nrxn1/Camk4/Neto1/Nrxn2/Fen1/Htr7/Cpeb3/Tanc1/Tbr1/Bdnf/Lrrn4/Plcb1/Snap25/Ythdf1/Oprl1/Cyp7b1/Syt11/Musk/Brinp1/Slc24a2/Jun/Ephb2/Camk2n1/Htr6/Atp8a1/Kit/Arl6ip5/Grm7/Oxtr/Cacna1c/Ntf3/Ccnd2/Slc8a2/Klk8/Slc17a7/Abcc8/Gabra5/Pak1/Ric8a/Pomk/Ndrg4/Ldlr/Stra6/Nptn/Fgf13 | 70 |
| BP | GO:0022898 | regulation of transmembrane transporter activity | 66/1983 | 297/17474 | 5342278 | 1,00982E+11 | 8880937674 | Atp1b1/Hk1/Jsrp1/Kcnc2/Ppp3r1/Shisa6/Vamp2/Ywhae/Asic2/Vmp1/Stac2/Taco1/Cacng5/Prkca/Osr1/Twist1/Hspa2/Chrm3/Actn2/Ubqln1/Glrx/Rasgrf2/Ppp3cb/Ptk2b/Fgf14/Kcns2/Cacng2/Pla2g6/Mapk8ip2/Cacnb3/Ifngr2/Ehd3/Nrxn1/Neto1/Cnih2/Cfl1/Gsto1/Lrrc55/Plcb1/Oprl1/Nlgn1/Asph/Ptpn3/Hpca/Ephb2/Ywhah/Stim2/Cabp1/Tesc/Rph3a/Tcaf1/Snca/Kcna1/Calm3/Fxyd7/Scn1b/Sphk2/Abcc8/Kcnc1/P2ry6/Cacna1a/Neto2/Scn2b/Htr3a/Stoml1/Fgf13 | 66 |
| BP | GO:0070050 | neuron cellular homeostasis | 25/1983 | 69/17474 | 5,7E+07 | 1,03344E+11 | 9088634842 | Atp6v1h/Adora1/Psen2/Kcnh1/Atp6v0a1/Psen1/Scgn/Erc2/Atp6v1c1/Nell2/Atp6v1a/Atp6v1g2/Chrna1/Slc24a2/Atp6v0e2/Immt/Erc1/Atp6v1e1/Slc8a2/Slc17a7/Sv2b/Atp6v1b2/Cacna1a/Atp6v0d1/Atp6ap2 | 25 |
| BP | GO:0046939 | nucleotide phosphorylation | 34/1983 | 114/17474 | 7,2E+07 | 1,21458E+11 | 1,06817E+11 | Hdac4/Ak9/Hk1/Pfkl/Zbtb7a/Ogdh/Ppp2ca/Stat3/Cmpk2/Aldoart2/Psen1/Ak7/Pfkp/Pfkm/Zbtb20/Eno1b/Slc2a6/Ak1/Mtch2/Eif6/Ak5/Aldoart1/Ak4/Eno1/Prkag2/Dck/P2rx7/Tigar/Gpi1/Aldoa/Prkaca/Pkm/Nme9/Pgk1 | 34 |
| BP | GO:0006165 | nucleoside diphosphate phosphorylation | 33/1983 | 110/17474 | 9,5E+07 | 1,57349E+11 | 1,38381E+11 | Hdac4/Ak9/Hk1/Pfkl/Zbtb7a/Ogdh/Ppp2ca/Stat3/Cmpk2/Aldoart2/Psen1/Ak7/Pfkp/Pfkm/Zbtb20/Eno1b/Slc2a6/Ak1/Mtch2/Eif6/Ak5/Aldoart1/Ak4/Eno1/Prkag2/P2rx7/Tigar/Gpi1/Aldoa/Prkaca/Pkm/Nme9/Pgk1 | 33 |
| BP | GO:0010959 | regulation of metal ion transport | 88/1983 | 445/17474 | 1,2E+09 | 1,79843E+11 | 1,58164E+11 | Cxcr4/Adora1/Ptgs2/Atp1b1/Rgs7/Psen2/Tgfb2/Sgk1/Fyn/Atg5/Jsrp1/Kcnc2/Ppp3r1/Stc2/Kcnmb1/Vdac1/Gjc2/Vamp2/Ywhae/Vmp1/Stac2/Wnk4/Itgb3/Osr1/Hspa2/Actn2/Agtr1a/Ubqln1/Glrx/Ppp3cb/Ptk2b/Stc1/Fgf14/Kcns2/Pla2g6/Cntn1/Ano6/Cacnb3/Hes1/Mylk/Glp1r/Ehd3/Prkce/Bin1/Neto1/Gnaq/Kcnip2/Gsto1/Lrrc55/Plcb1/Snta1/Gnas/Oprl1/Chd7/Asph/Nkain3/Ptpn3/Hpca/Nkain1/Kcnab2/Ywhah/Stim2/Cabp1/Tesc/P2rx7/Snca/Cacna1c/Kcna1/Calm3/Fxyd7/Scn1b/Sphk2/Abcc8/Kcnc1/Il16/P2ry6/Saraf/Homer3/Tmem38a/Cacna1a/Neto2/Gnao1/Ubash3b/Thy1/Scn2b/Stoml1/Trf/Fgf13 | 88 |
| BP | GO:0097401 | synaptic vesicle lumen acidification | 11/1983 | 18/17474 | 5,8E+08 | 8352538577 | 73456960660 | Atp6v1h/Atp6v0a1/Atp6v1c1/Atp6v1a/Atp6v1g2/Atp6v0e2/Atp6v1e1/Slc17a7/Atp6v1b2/Atp6v0d1/Atp6ap2 | 11 |
| BP | GO:0051648 | vesicle localization | 49/1983 | 212/17474 | 8E+08 | 1,07544E+12 | 94580448504 | Trak2/Mreg/Vps4b/Klhl12/Sec16b/Sar1a/Ap3d1/Ppfia2/Dctn2/Rab1a/Fbxw11/Pafah1b1/Hgs/Psen1/Tmed9/Sybu/Trappc9/Mapk15/Synj1/Nrxn1/Cdh2/Myo5b/Cnih2/Stam/Dnm1/Kif5c/Madd/Bloc1s6/Syndig1/Atp9a/Nlgn1/Syt11/Sdcbp/Pclo/Dync1i1/Snca/Rab7/Syn2/Borcs5/Ap3b2/Ap3m2/Rab3a/Smpd3/Exoc8/Map2k1/Snap91/Bsn/Myrip/Syn1 | 49 |
| BP | GO:0099072 | regulation of postsynaptic membrane neurotransmitter receptor levels | 35/1983 | 132/17474 | 1,1E+10 | 13612726716 | 1,19718E+12 | Pip5k1c/Ppp3r1/Snap47/Shisa6/Vamp2/Ywhae/Flot2/Itgb3/Cacng5/Nptx1/Gphn/Nrxn3/Hsp90aa1/Cacng2/Ap2m1/Synj1/Pacsin1/Neto1/Lgi1/Rapgef4/Snap25/Frrs1l/Eps15/Hpca/Nsg1/Cplx1/Ap2s1/Cpt1c/Ap2a2/Vps35/Neto2/Vac14/Map2k1/Porcn/Slc9a6 | 35 |
| BP | GO:0055074 | calcium ion homeostasis | 68/1983 | 335/17474 | 1,2E+10 | 1,47367E+12 | 1,29603E+12 | Efhc1/Bok/Adora1/Atp1b1/Psen2/Tgfb2/Kcnh1/Atg5/Grik2/Mcu/Micu1/Trpm2/Jsrp1/Stc2/Anxa6/Ywhae/Wnk4/Itgb3/Prkca/Psen1/Chga/Scgn/Mcur1/Anxa7/Erc2/Ptk2b/Stc1/Ank/Grina/Kctd17/Gp5/Glp1r/Tmem178/Prkce/Gsto1/Plcb1/Gnas/Sv2a/Rap1gds1/Chd7/Asph/Slc24a2/Hcrtr1/Camk2n1/Plch2/Stim2/P2rx7/Gpr12/Snx10/Snca/Immt/Cacna1c/Erc1/Slc8a2/Calm3/Sv2b/P2ry6/Bnip3/Fgfr1/Micu3/Tmem38a/Cacna1a/Ubash3b/Thy1/Slc37a4/Cib2/Nptn/Nt5e | 68 |

| | | | | | | | | | |
|---|---|---|---|---|---|---|---|---|---|
| BP | GO:0006090 | pyruvate metabolic process | 33/1983 | 124/17474 | 2E+10 | 2,21368E+12 | 1,94684E+12 | Hdac4/Hk1/Pfkl/Zbtb7a/Ogdh/Ppp2ca/Vdac1/Pdk2/Stat3/Aldoart2/Psen1/Pfkp/Pck2/Pfkm/Zbtb20/Eno1b/Slc2a6/Mtch2/Pdhx/Eif6/Nr4a3/Aldoart1/Eno1/Prkag2/P2rx7/Tigar/Gpi1/Aldoa/Prkaca/Dlat/Pkm/Me1/Pgk1 | 33 |
| BP | GO:0021537 | telencephalon development | 55/1983 | 258/17474 | 2,6E+09 | 2,80106E+11 | 2,46342E+11 | Efhc1/Cxcr4/Uchl5/Zbtb18/Ogdh/Pex13/Rtn4/Pafah1b1/Rtn4rl1/Ywhae/Socs7/Lamb1/Psen1/Inhba/Tubb2a/Agtpbp1/Rarb/Nrg3/Dnah5/Trappc9/Tuba1a/Bmerb1/Rtn4r/Hes1/Fbxo45/Robo1/Sema6b/Cdh2/Cep120/Nars/Slit1/Crtac1/Tbr1/Neurod1/Plcb1/Mkks/Btbd3/Bhlhe22/Chd7/Nr4a3/Nfib/Ephb2/Htr6/Trp73/Slit2/Epha5/Prdm8/Neurod6/Atg7/Kcna1/Aldh1a3/Fgfr1/Dmxl2/Tmem108/Fgf13 | 55 |
| BP | GO:0060560 | developmental growth involved in morphogenesis | 59/1983 | 284/17474 | 2,8E+09 | 28723362184 | 2,5261E+11 | Rdh10/Rims1/Nrp2/Cxcr4/Kif26b/Wasf1/Syt1/Rab21/Llph/Rtn4/Fstl4/Pafah1b1/Mapt/Hsp90aa1/Nrn1/Map1b/Fgf10/Cpne6/Bin3/Sema5a/Spag6l/Rtn4r/Sema5b/Robo1/Ddr1/Hsp90ab1/Sema6b/Wdr36/Myo5b/Cfl1/Prkg1/Slit1/Zfyve27/Olfm1/Bdnf/Ttl/Flrt3/Epha7/Sh3gl2/Plaa/Sema3e/Slit2/Rufy3/Auts2/Rnf6/Sema4f/Mgll/Cpne9/Mag/Syt3/Cyfip1/Pak1/Nrp1/Islr2/Ppp2r3a/Tmem108/Slc9a6/Fgf13/Gdi1 | 59 |
| BP | GO:0006735 | NADH regeneration | 10/1983 | 17/17474 | 3,2E+10 | 3,10205E+11 | 2,72811E+11 | Hk1/Pfkl/Pfkp/Pfkm/Eno1b/Eno1/Gpi1/Aldoa/Pkm/Pgk1 | 10 |
| BP | GO:0061621 | canonical glycolysis | 10/1983 | 17/17474 | 3,2E+10 | 3,10205E+11 | 2,72811E+11 | Hk1/Pfkl/Pfkp/Pfkm/Eno1b/Eno1/Gpi1/Aldoa/Pkm/Pgk1 | 10 |
| BP | GO:0061718 | glucose catabolic process to pyruvate | 10/1983 | 17/17474 | 3,2E+10 | 3,10205E+11 | 2,72811E+11 | Hk1/Pfkl/Pfkp/Pfkm/Eno1b/Eno1/Gpi1/Aldoa/Pkm/Pgk1 | 10 |
| BP | GO:0021954 | central nervous system neuron development | 28/1983 | 99/17474 | 3,2E+10 | 3,10205E+11 | 2,72811E+11 | Nrp2/Epha4/Kifbp/Adarb1/Ogdh/Pafah1b1/Prkca/Hsp90aa1/Agtpbp1/Fbxo45/Hsp90ab1/B4galt6/Dcc/Gnaq/Bhlhe22/Rapgef2/Nhlh2/Nfib/Ephb2/Sema3e/Slit2/Prdm8/Atg7/Scn1b/Fgfr1/Nrp1/Lingo1/Ephb1 | 28 |
| BP | GO:0046034 | ATP metabolic process | 48/1983 | 216/17474 | 3,4E+10 | 3,15422E+11 | 2,774E+12 | Hdac4/Ndufa10/Atp1b1/Ak9/Hk1/Pfkl/Zbtb7a/Atp5b/Ogdh/Ppp2ca/Atp5g1/Stat3/Aldoart2/Psen1/Pfkp/Pfkm/Zbtb20/Atp6v1a/Hspa1b/Hspa1a/Eno1b/Atp5a1/Slc2a6/Ak1/Kif5c/Ola1/Mtch2/Eif6/Ak5/Vcp/Aldoart1/Ak4/Eno1/Prkag2/P2rx7/Tigar/Gpi1/Sphk2/Aldoa/Atp6v1b2/Prkaca/Ldhd/Hspa8/Pkm/Map2k1/Clpx/Nt5e/Pgk1 | 48 |
| BP | GO:0031346 | positive regulation of cell projection organization | 83/1983 | 449/17474 | 4,5E+08 | 3,78575E+11 | 3,32941E+12 | Stau2/Epha4/Hdac4/Abl2/Sgk1/Fyn/Fig4/Cd24a/Map2k2/Rab21/Camk2b/Efemp1/Pfn1/Pafah1b1/Serpinf1/Bcas3/Mapt/Hsp90aa1/Arsb/Enc1/Map1b/Ptk2b/Nefl/Sema5a/Kctd17/Cntn1/Epha3/Robo1/Adamts1/Pcp4/Pacsin1/Neu1/Ddr1/Nrxn1/Dpysl3/Cep120/Dcc/Myo5b/Cfl1/Ehd1/Htr7/Zfyve27/Neurl1a/Camk1d/Hspa5/Bdnf/Src/Stmn2/Nlgn1/Clrn1/Rapgef2/Musk/Akirin1/Ephb2/Wrap73/Hgf/Tapt1/Kit/Rufy3/P2rx7/Clip1/Auts2/Scn1b/Ankrd27/Cyfip1/Igf1r/Cpeb1/Pak1/Apbb1/Fgfr1/Ndrg4/Carmil2/Nrp1/Islr2/Nptn/Parp6/Map2k1/Mns1/Tmem30a/Snap91/Nckipsd/Frmd7/Gdi1 | 83 |
| BP | GO:0050803 | regulation of synapse structure or activity | 62/1983 | 310/17474 | 5,8E+09 | 4,79684E+12 | 4,21862E+11 | Stau2/Nrp2/Epha4/Psen2/Fyn/Slc17a8/Ppfia2/Camk2b/Pafah1b1/Asic2/Srcin1/Neurod2/Cntnap1/Prkca/Rhob/Dgkb/Psen1/Rps6ka5/Arhgap22/Ptk2b/Nefl/Pcdh8/Ywhaz/Sybu/Amigo2/Tuba1a/Eif4g1/Mapk14/Tubb5/Nrxn1/Cdh2/Myo5b/Cfl1/Abhd17b/Neurl1a/Tanc1/Rapgef4/Bdnf/Flrt3/Syndig1/Nlgn1/Pdlim5/Epha7/Vcp/Musk/Elavl2/Ephb2/Snca/Oxtr/Vhl/Lrtm2/Chd4/Klk8/Slc17a7/Cyfip1/Vps35/Cdh8/Hspa8/Nectin1/Snap91/Clstn2/Ephb1 | 62 |
| BP | GO:0042391 | regulation of membrane potential | 87/1983 | 480/17474 | 6E+09 | 4,94395E+11 | 4,34799E+12 | Rims1/Bok/Adora1/Atp1b1/Rgs7/Kcnh1/Hebp2/Popdc3/Bves/Grik2/Kcnc2/Kcnmb4/Ywhae/Asic2/Jup/Cntnap1/Mapt/Eif4a3/Ppp2r3c/Psen1/Actn2/Dsp/Glrx/Ptk2b/Fgf14/Cacng2/Mapk8ip2/Ano6/Cacnb3/Pkp2/Sod2/Slc29a1/Ehd3/Prkce/Nrxn1/Dsg2/Celf4/Bin1/Neto1/Cnih2/Kcnk4/Gnaq/Got1/Kcnip2/Scn7a/Rapgef4/Chrna1/Mtch2/Bdnf/Snta1/Src/Nlgn1/Npy2r/Mllt11/Vcp/Ptpn3/Jun/Slc25a33/Kcnab2/Pclo/Ywhah/Wdr1/P2rx7/Rimbp2/Kctd7/Stx1a/Snca/Arl6ip5/Cacna1c/Bid/Kcna1/Slc8a2/Calm3/Grik5/Scn1b/Myh14/Slc17a7/Gabra5/Tmem135/Bnip3/Cacna1a/Kcnk1/Tmem25/Scn2b/Htr3a/Tmem108/Fgf13 | 87 |
| BP | GO:0051650 | establishment of vesicle localization | 43/1983 | 191/17474 | 7,6E+10 | 6,11789E+11 | 5,38042E+11 | Trak2/Mreg/Vps4b/Klhl12/Sec16b/Sar1a/Ap3d1/Ppfia2/Dctn2/Rab1a/Fbxw11/Pafah1b1/Hgs/Psen1/Tmed9/Sybu/Trappc9/Mapk15/Synj1/Myo5b/Cnih2/Stam/Dnm1/Kif5c/Madd/Bloc1s6/Atp9a/Nlgn1/Syt11/Sdcbp/Pclo/Dync1i1/Snca/Rab7/Borcs5/Ap3b2/Ap3m2/Rab3a/Smpd3/Exoc8/Map2k1/Snap91/Myrip | 43 |
| BP | GO:0007018 | microtubule-based movement | 75/1983 | 405/17474 | 1,2E+10 | 84004102468 | 7,3878E+11 | Stau2/Trak2/Spag16/Mreg/Daw1/Arl8a/Kifap3/Kif26b/Wasf1/Ap3d1/Rab21/Nefh/Rab1a/Fbxw11/Cfap52/Tekt1/Pafah1b1/Flot2/Ttll6/Mapt/Ccdc40/Ak7/Klc1/Agtpbp1/Map1b/Cfap70/Dnah12/Nefl/Nefm/Spef2/Dnah5/Ropn1l/Sybu/Tuba1a/Spag6l/Iqcg/Neto1/Cnih2/Catsperz/Borcs7/Neurl1a/Meig1/Enkur/Kif5c/Madd/Sord/Bloc1s6/Stard7/Mkks/Kif3b/Cenpe/Ssx2ip/Cfap206/Dnaja1/Dnali1/AU040320/Uchl1/Dnah10/Dync1i1/Arl8b/Borcs5/Ap3b2/Dnah3/Ap3m2/Ttc29/Hydin/Dynlrb2/Hspa8/Map2k1/Mns1/Tmem108/Bsn/Ttc21a/Dynlt3/Armcx3 | 75 |
| BP | GO:0042026 | protein refolding | 10/1983 | 19/17474 | 1,2E+10 | 84004102468 | 7,3878E+11 | Hspd1/Hspa2/Hsp90aa1/Hspa1b/Hspa1a/Hspa5/Dnaja1/Dnaja2/Hspa8/Dnaja4 | 10 |
| BP | GO:0006874 | intracellular calcium ion homeostasis | 60/1983 | 306/17474 | 1,5E+10 | 98384081905 | 8,65245E+11 | Efhc1/Bok/Adora1/Atp1b1/Psen2/Tgfb2/Kcnh1/Atg5/Grik2/Mcu/Micu1/Trpm2/Jsrp1/Stc2/Anxa6/Ywhae/Itgb3/Prkca/Psen1/Chga/Scgn/Mcur1/Anxa7/Erc2/Ptk2b/Stc1/Grina/Kctd17/Gp5/Glp1r/Tmem178/Prkce/Gsto1/Plcb1/Sv2a/Rap1gds1/Chd7/Asph/Slc24a2/Hcrtr1/Plch2/Stim2/P2rx7/Gpr12/Snca/Immt/Cacna1c/Erc1/Slc8a2/Calm3/Sv2b/P2ry6/Bnip3/Micu3/Tmem38a/Cacna1a/Ubash3b/Thy1/Slc37a4/Nptn | 60 |

| | | | | | | | | |
|---|---|---|---|---|---|---|---|---|
| BP | GO:0048588 | developmental cell growth | 56/1983 | 280/17474 | 1,6E+11 | 0.0010157091756753174 | 89327224618 | Rims1/Nrp2/Cxcr4/Wasf1/Syt1/Rab21/Llph/Rtn4/Fstl4/Map2k4/Pafah1b1/Mapt/Hsp90aa1/Nrn1/Map1b/Cpne6/Sema5a/Spag6l/Rtn4r/Sema5b/Ddr1/Hsp90ab1/Sema6b/Wdr36/Mex3c/Myo5b/Prkg1/Slit1/Zfyve27/Olfm1/Bdnf/Ttl/Flrt3/Pdlim5/Epha7/Sh3gl2/Plaa/Sema3e/Slit2/Rufy3/Auts2/Rnf6/Sema4f/Mgll/Cpne9/Mag/Syt3/Cyfip1/Akap13/Pak1/Nrp1/Islr2/Tmem108/Slc9a6/Fgf13/Gdi1 | 56 |
| BP | GO:0097120 | receptor localization to synapse | 24/1983 | 85/17474 | 1,6E+11 | 0.0010190116412071614 | 8,96177E+11 | Snap47/Shisa6/Vamp2/Itgb3/Cacng5/Nptx1/Gphn/Nrxn3/Cacng2/Nrxn1/Neto1/Cnih2/Lgi1/Kif5c/Rapgef4/Snap25/Kif3b/Nlgn1/Nsg1/Cplx1/Vps35/Neto2/Nptn/Map2k1 | 24 |
| BP | GO:1904951 | positive regulation of establishment of protein localization | 66/1983 | 349/17474 | 2E+11 | 0.001209473991272272 | 0.0010636800126978554 | Ptgs2/Tgfb2/Fyn/Mcu/Sar1a/Trpm2/Vamp2/Ywhae/Bcas3/Jup/Psen1/Gpr68/Hsp90aa1/Chrm3/Anxa7/Ppp3cb/Prkcd/Ang/Pck2/Oxct1/Ptger4/Cct5/Sybu/Pla2g6/Pfkm/Cacnb3/Mapk1/Cct8/Tcp1/Mapk14/Glp1r/Hsp90ab1/Prkce/Camk4/Cep120/B3gat3/Gnaq/Lrp2/Rapgef4/Plcb1/Src/Gnas/Asph/Hpca/Camk2n1/Rufy3/P2rx7/Aacs/Cct6a/Tcaf1/Edem1/Atg7/Cacna1c/Abcc8/Pak1/Apbb1/Irs2/Prkaca/Vps35/Exph5/Coro2b/Oaz2/Tmem30a/Myrip/Sytl4/Acsl4 | 66 |
| BP | GO:0050807 | regulation of synapse organization | 59/1983 | 302/17474 | 2E+09 | 0.0012116438533221565 | 0.001065588312429333 | Stau2/Nrp2/Epha4/Psen2/Fyn/Ppfia2/Camk2b/Pafah1b1/Asic2/Srcin1/Neurod2/Cntnap1/Prkca/Rhob/Dgkb/Psen1/Rps6ka5/Arhgap22/Ptk2b/Nefl/Pcdh8/Ywhaz/Amigo2/Tuba1a/Eif4g1/Mapk14/Tubb5/Nrxn1/Cdh2/Myo5b/Cfl1/Abhd17b/Neurl1a/Tanc1/Rapgef4/Bdnf/Flrt3/Syndig1/Nlgn1/Pdlim5/Epha7/Vcp/Musk/Elavl2/Ephb2/Snca/Oxtr/Vhl/Lrtm2/Chd4/Klk8/Cyfip1/Vps35/Cdh8/Hspa8/Nectin1/Snap91/Clstn2/Ephb1 | 59 |
| BP | GO:0042886 | amide transport | 74/1983 | 407/17474 | 2,6E+11 | 0.0014990688569363943 | 0.0013183661594109546 | Klf7/Adora1/Grm1/Mcu/Psap/Gnaz/Trpm2/Pfkl/Slc17a8/Rab1a/Sgpp1/Psen1/Gpr68/Chga/Chrm3/Anxa7/Ppp3cb/Pck2/Oxct1/Ptger4/Sybu/Mafa/Pla2g6/Pfkm/Glp1r/Tap2/Prkce/Syt7/Gnaq/Anxa1/Slc18a2/Stxbp1/Acvr1c/Lrp2/Rapgef4/Neurod1/Madd/Bdnf/Plcb1/Snap25/Gnas/Myt1/Foxo1/Npy2r/Ensa/Chd7/Camk2n1/Htr6/Cptp/Pclo/Epha5/Cplx1/P2rx7/Aacs/Stx1a/Snca/Arl6ip5/Grm7/Atg7/Cacna1c/Slc17a7/Abcc8/Pak1/Slc25a22/Irs2/Rab3a/Prkaca/Gnao1/Smpd3/Cyb5r4/Myrip/Cdk16/Sytl4/Acsl4 | 74 |
| BP | GO:0061387 | regulation of extent of cell growth | 31/1983 | 128/17474 | 3,1E+11 | 0.0017493308521434523 | 0.0015384607494232738 | Rab21/Rtn4/Fstl4/Pafah1b1/Mapt/Map1b/Sema5a/Rtn4r/Sema5b/Sema6b/Wdr36/Myo5b/Slit1/Zfyve27/Olfm1/Bdnf/Ttl/Epha7/Sema3e/Slit2/Rufy3/Rnf6/Sema4f/Mgll/Mag/Cyfip1/Pak1/Nrp1/Islr2/Fgf13/Gdi1 | 31 |
| BP | GO:0017157 | regulation of exocytosis | 46/1983 | 222/17474 | 3,6E+10 | 0.0019512159192476488 | 0.0017160099255862176 | Rims1/Vps4b/Kcnh1/Ppfia2/Syt1/Rab21/Vamp2/Nsf/Prkca/Hgs/Rab15/Cadps/Lgi3/Pla2g6/Septin5/Fbxo45/Myo5b/Syt7/Anxa1/Stam/Stxbp1/Rapgef4/Syt13/Atp9a/Nlgn1/Syt11/Sdcbp/Pclo/Cplx1/Rph3a/P2rx7/Stx1a/Snca/Rab7/D6Wsu163e/Syt3/Sphk2/Sv2b/Rab3a/Cacna1a/Smpd3/Exph5/Scamp5/Syp/Syn1/Sytl4 | 46 |
| BP | GO:0030073 | insulin secretion | 50/1983 | 248/17474 | 3,6E+10 | 0.0019512159192476488 | 0.0017160099255862176 | Klf7/Mcu/Gnaz/Trpm2/Pfkl/Gpr68/Chga/Chrm3/Anxa7/Ppp3cb/Pck2/Oxct1/Sybu/Mafa/Pla2g6/Pfkm/Glp1r/Prkce/Syt7/Gnaq/Anxa1/Slc18a2/Acvr1c/Rapgef4/Neurod1/Plcb1/Snap25/Gnas/Myt1/Foxo1/Ensa/Camk2n1/Pclo/Epha5/Cplx1/Aacs/Stx1a/Atg7/Cacna1c/Abcc8/Slc25a22/Irs2/Rab3a/Prkaca/Gnao1/Cyb5r4/Myrip/Cdk16/Sytl4/Acsl4 | 50 |
| BP | GO:0019693 | ribose phosphate metabolic process | 77/1983 | 433/17474 | 4E+10 | 0.0021221369055126975 | 0.0018663275332013198 | Hdac4/Ndufa10/Atp1b1/Ak9/Hk1/Pfkl/Zbtb7a/Nudt4/Atp5b/Ogdh/Ppp2ca/Vdac1/Acaca/Pdk2/Atp5g1/Stat3/Aldoart2/Pygl/Psen1/Acot5/Acot6/Pfkp/Hmgcs1/Tkt/Pfkm/Zbtb20/Atp6v1a/Hspa1b/Hspa1a/Eno1b/Atp5a1/Rfk/Entpd7/Acsl5/Slc2a6/Ak1/Kif5c/Ola1/Mtch2/Pdhx/Acss2/Eif6/Tdo2/Slc27a3/Npr1/Dpyd/Ak5/Vcp/Aldoart1/Ak4/Ctps/Eno1/Acot7/Prkag2/P2rx7/Snca/Tigar/Gpi1/Sphk2/Dgat2/Aldoa/Acsl1/Atp6v1b2/Prkaca/Ldhd/Mvd/Aprt/Hspa8/Dlat/Pkm/Map2k1/Clpx/Nt5e/Nme9/Pgk1/Prps1/Acsl4 | 77 |
| BP | GO:0032535 | regulation of cellular component size | 71/1983 | 392/17474 | 4,3E+09 | 0.0022575505626675177 | 0.001985417982108275 | Nck2/Rab21/Rtn4/Fstl4/Pfn1/Rap1gap2/Pafah1b1/Mapt/Grb2/Scin/Cfl2/Hsp90aa1/Inf2/Actn2/Map1b/Anxa7/Prkcd/Ptk2b/Nefl/Nefm/Sema5a/Ano6/Rtn4r/Ap2m1/Sema5b/Hsp90ab1/Sema6b/Washc1/Prkce/Bin1/Wdr36/Myo5b/Cfl1/Capn1/Slit1/Zfyve27/Borcs7/Add3/Olfm1/Bdnf/Ttl/Mkks/Dstn/Epha7/Tmod1/Capzb/Trp73/Sema3e/Pclo/Wdr1/Slit2/Rufy3/P2rx7/Rnf6/Capza2/Sema4f/Mgll/Arpc4/Borcs5/Mag/Cyfip1/Pex11a/Aqp11/Pak1/Carmil2/Cotl1/Nrp1/Islr2/Tpm1/Fgf13/Gdi1 | 71 |
| BP | GO:1901617 | organic hydroxy compound biosynthetic process | 46/1983 | 225/17474 | 5,1E+11 | 0.002542381099775197 | 0.0022359141081214804 | Avpr1b/Cyb5r1/Hsd17b7/Ephx1/Tgfb2/Pcbd1/Mfsd12/Tph2/Pmel/Sec14l2/Alox8/Agtr1a/Bmp6/Ippk/Sptlc1/Hmgcs1/Pck2/Fdft1/Ptk2b/Sqle/Cyb5r3/Mas1/Pgp/Stard4/Got1/Slc27a2/Hdc/Cyp7b1/Rapgef2/Tyrp1/Dhcr24/Dhdds/H6pd/Insig1/Gpr37/Snca/Sphk2/P2ry6/Dhcr7/Msmo1/Vps35/Mvd/Sc5d/Ip6k2/Nsdhl/Cited1 | 46 |
| BP | GO:0007274 | neuromuscular synaptic transmission | 12/1983 | 30/17474 | 5,4E+10 | 0.0026591202177991156 | 0.002338581108353042 | Cd24a/Adarb1/Erc2/Nrxn1/Stxbp1/Chrna1/Nlgn1/Rimbp2/Erc1/Ntf3/Rab3a/Cacna1a | 12 |
| BP | GO:0032984 | protein-containing complex disassembly | 47/1983 | 233/17474 | 6E+10 | 0.0028435420318660017 | 0.0025007721095188264 | Vps4b/Sgk1/Atg5/Gas2l1/Eif5a/Vmp1/Becn1/Nsf/Smarcd2/Scin/Cfl2/Klc1/Actn2/Ubqln1/Map1b/Chmp7/Sema5a/Bmerb1/Map6d1/Synj1/Pik3c3/Etf1/Atg12/Cfl1/Frat1/Add3/Dstn/Stmn2/Asph/Vcp/Tmod1/Sh3gl2/Dnajc6/Capzb/Wdr1/Tecpr1/Capza2/Napa/Igf1r/Sh3gl3/Bnip3/Abce1/Carmil2/Smarca4/Hspa8/Tpm1/Fgf13 | 47 |

| | | | | | | | | |
|---|---|---|---|---|---|---|---|---|
| BP | GO:0009914 | hormone transport | 70/1983 | 389/17474 | 6E+11 | 0.0028435420318660017 | 0.0025007721095188264 | Klf7/Adora1/Myb/Mcu/Gnaz/Trpm2/Pfkl/Igfbp3/Rab1a/Vamp2/Wnk4/Gpr68/Chga/Chrm3/Inhba/Agtr1a/Bmp6/Anxa7/Ppp3cb/Slc7a8/Pck2/Nkx3-1/Oxct1/Ptger4/Ywhaz/Sybu/Mafa/Pla2g6/Nell2/Pfkm/Glp1r/Prkce/Syt7/Gnaq/Anxa1/Slc18a2/Acvr1c/Lrp2/Rapgef4/Neurod1/Madd/Plcb1/Snap25/Gnas/Myt1/Foxo1/Ensa/Chd7/Camk2n1/Pclo/Epha5/Cplx1/Aacs/Stx1a/Atg7/Cacna1c/Abcc8/Nmb/Slc25a22/Irs2/Fgfr1/Rab3a/Prkaca/Gnao1/Smpd3/Cyb5r4/Myrip/Cdk16/Sytl4/Acsl4 | 70 |
| BP | GO:0032411 | positive regulation of transporter activity | 32/1983 | 139/17474 | 6,7E+10 | 0.003119941103928707 | 0.0027438531270544486 | Atp1b1/Kcnc2/Ppp3r1/Asic2/Vmp1/Stac2/Hspa2/Actn2/Glrx/Ppp3cb/Prkcd/Fgf14/Cacng2/Cacnb3/Ifngr2/Ehd3/Cfl1/Gsto1/Lrrc55/Plcb1/Asph/Ephb2/Stim2/Tesc/Kctd7/Tcaf1/Kcna1/Calm3/Abcc8/Kcnc1/P2ry6/Htr3a | 32 |
| BP | GO:0006066 | alcohol metabolic process | 60/1983 | 322/17474 | 7,2E+09 | 0.003278031429253957 | 0.0028828867238600165 | Rdh10/Idh1/Sgpp2/Avpr1b/Hsd17b7/Ephx1/Pcbd1/Nudt4/Sec14l2/Aldh3a2/Gdpd1/Nfe2l1/Dhrs7/Sgpp1/Itpk1/Bmp6/Ippk/Sptlc1/Hmgcs1/Pck2/Fdft1/Ptk2b/Sqle/Cyb5r3/Mas1/Pgp/Lmf1/Stard4/Aldh1a1/Myof/Got1/Lrp2/Cat/Sord/Plcb1/Impa1/Cyp7b1/Dhcr24/Dhdds/Disp3/H6pd/Napepld/Insig1/Plb1/Coq2/Snca/Cebpa/Sphk2/Dgat2/P2ry6/Dhcr7/Fgfr1/Msmo1/Mvd/Ldlr/Aplp2/Sc5d/Idh3a/Ip6k2/Nsdhl | 60 |
| BP | GO:0071692 | protein localization to extracellular region | 74/1983 | 420/17474 | 7,6E+10 | 0.003434504202370928 | 0.0030204977535282524 | Klf7/Tgfb2/Mcu/Psap/Gnaz/Trpm2/Pfkl/Ap1b1/Pafah1b1/Srcin1/Gpr68/Chga/Chrm3/Agtr1a/Anxa7/Ppp3cb/Ang/Pck2/Oxct1/Ptger4/Sybu/Mafa/Pla2g6/Pfkm/Tango2/Nrros/Cd200/Lmf1/Glp1r/Prkce/Pik3c3/Syt7/Gnaq/Anxa1/Slc18a2/Acvr1c/Rapgef4/Neurod1/Madd/Nr1h3/Bloc1s6/Plcb1/Snap25/Gnas/Myt1/Foxo1/Ensa/Rap1gds1/Camk2n1/Nbl1/Pclo/Epha5/Cplx1/P2rx7/Aacs/Stx1a/Atg7/Cacna1c/M6pr/Abcc8/Apbb1/Slc25a22/Irs2/Rab3a/Prkaca/Vps35/Gnao1/Exph5/Cyb5r4/Myrip/Porcn/Cdk16/Sytl4/Acsl4 | 74 |
| BP | GO:0021700 | developmental maturation | 65/1983 | 359/17474 | 9E+10 | 0.0039695804394365465 | 0.0034910741385876594 | Atp6v1h/Mreg/Ptgs2/Tdrd5/Abl2/Psen2/Tgfb2/Ap3d1/Zbtb7a/Gdf11/Camk2b/Pfn1/Phospho1/Neurod2/Atp6v0a1/Cntnap1/Foxj1/Psen1/Nsun2/Ptk2b/Nefl/Ropn1l/Ywhaz/Atp6v1c1/Sybu/Ano6/Hes1/Atp6v1a/Ppp2r1a/Atp6v1g2/C3/Washc1/Nrxn1/Cabyr/B4galt6/Catsperz/Gnaq/Kcnip2/Neurl1a/Stxbp1/Grb14/Bloc1s6/Fermt1/Plcb1/Trpc4ap/Nlgn1/Wdr77/Ythdf2/Pebp1/Atp6v0e2/Snx10/Atp6v1e1/Cebpa/Ankrd27/Slc17a7/Fgfr1/Atp6v1b2/Rab3a/Prkaca/Cacna1a/Vps35/Atp6v0d1/Gldn/Atp6ap2/Mbtps2 | 65 |
| BP | GO:0010035 | response to inorganic substance | 82/1983 | 480/17474 | 9,5E+10 | 0.004123295426021259 | 0.0036262598143956062 | Adora1/Kcnh1/Lrp11/Map3k5/Fyn/Psap/Trpm2/Syt1/Kcnc2/Kcnmb4/Nefh/Camk2b/Kcnmb1/Rnf112/Nfe2l1/Neurod2/Mapt/Nptx1/Rhob/Gphn/Bmp6/Anxa7/Anxa11/Prkcd/Cpne6/Gja3/Ptk2b/Nefl/Ank/Tuba1a/Mapk1/Txnrd2/Tfrc/Mylk/Sod2/Nrxn1/Cfl1/Sipa1/Capn1/Syt7/Anxa1/Syt13/Cat/Sord/Plcb1/Gss/Src/Nlgn1/Foxo1/Kpna4/Syt11/Asph/Vcp/Alg2/Nr4a3/Jun/Hpca/Hgf/Slc5a1/P2rx7/Gpr37/Snca/Cpne9/Kcna1/Tigar/Cdkn1b/Selenow/Calm3/Cebpa/Syt3/Abcc8/Nsmce3/Mef2a/Plekha1/Bnip3/Fgfr1/Smpd3/Bcar1/Nt5e/Trf/Cpne4/Gdi1 | 82 |
| BP | GO:0010976 | positive regulation of neuron projection development | 46/1983 | 231/17474 | 9,8E+09 | 0.004154198831536309 | 0.003653438021622763 | Stau2/Abl2/Fyn/Fig4/Cd24a/Camk2b/Pafah1b1/Serpinf1/Mapt/Arsb/Enc1/Ptk2b/Cntn1/Epha3/Adamts1/Pcp4/Pacsin1/Neu1/Ddr1/Nrxn1/Dpysl3/Dcc/Ehd1/Htr7/Camk1d/Hspa5/Bdnf/Stmn2/Nlgn1/Rapgef2/Musk/Ephb2/Hgf/Clip1/Scn1b/Ankrd27/Cyfip1/Igf1r/Cpeb1/Apbb1/Fgfr1/Ndrg4/Nrp1/Nptn/Tmem30a/Nckipsd | 46 |
| BP | GO:0031345 | negative regulation of cell projection organization | 43/1983 | 213/17474 | 1,2E+12 | 0.004818079878548371 | 0.004237292660590927 | Trak2/Epha4/Itm2c/Fyn/Rab3ip/Efemp1/Rtn4/Fstl4/Rap1gap2/Pafah1b1/Rtn4rl1/Psen1/Spock1/Prkcd/Sema5a/Rtn4r/Hes1/Sema5b/Epha3/Sema6b/Nrxn1/Dpysl3/Cfl1/Slit1/Vim/Stmn2/Nlgn1/Rapgef2/Epha7/Ephb2/Capzb/Sema3e/Ywhah/Slit2/Rufy3/Rnf6/Sema4f/Mag/Klk8/Nrp1/Thy1/Fgf13/Gdi1 | 43 |
| BP | GO:0002790 | peptide secretion | 57/1983 | 308/17474 | 1,3E+12 | 0.005210957607762797 | 0.0045828116142973335 | Klf7/Adora1/Mcu/Gnaz/Trpm2/Pfkl/Rab1a/Gpr68/Chga/Chrm3/Anxa7/Ppp3cb/Pck2/Oxct1/Ptger4/Sybu/Mafa/Pla2g6/Pfkm/Glp1r/Prkce/Syt7/Gnaq/Anxa1/Slc18a2/Acvr1c/Rapgef4/Neurod1/Madd/Plcb1/Snap25/Gnas/Myt1/Foxo1/Npy2r/Ensa/Chd7/Camk2n1/Pclo/Epha5/Cplx1/Aacs/Stx1a/Atg7/Cacna1c/Abcc8/Slc25a22/Irs2/Rab3a/Prkaca/Gnao1/Smpd3/Cyb5r4/Myrip/Cdk16/Sytl4/Acsl4 | 57 |
| BP | GO:1900242 | regulation of synaptic vesicle endocytosis | 11/1983 | 28/17474 | 1,3E+12 | 0.005210957607762797 | 0.0045828116142973335 | Pip5k1c/Actg1/Ppp3cb/Ap2m1/Syt7/Dnm1/Nlgn1/Syt11/Slc17a7/Scamp5/Snap91 | 11 |
| BP | GO:0030433 | ubiquitin-dependent ERAD pathway | 21/1983 | 79/17474 | 1,4E+12 | 0.005463184762233951 | 0.004804634476803371 | Ube2g2/Erlec1/Canx/Tmub2/Ubqln1/Derl1/Nrros/Usp14/Hspa5/Dnajc10/Ube2j1/Vcp/Ubxn10/Fbxo2/Ube4b/Rnf103/Edem1/Ubxn8/Calr/Amfr/Ubqln2 | 21 |
| BP | GO:0035592 | establishment of protein localization to extracellular region | 72/1983 | 414/17474 | 1,4E+12 | 0.005518803981853901 | 0.0048535491724963 | Klf7/Tgfb2/Mcu/Psap/Gnaz/Trpm2/Pfkl/Ap1b1/Pafah1b1/Srcin1/Gpr68/Chga/Chrm3/Agtr1a/Anxa7/Ppp3cb/Ang/Pck2/Oxct1/Ptger4/Sybu/Mafa/Pla2g6/Pfkm/Tango2/Cd200/Lmf1/Glp1r/Prkce/Pik3c3/Syt7/Gnaq/Anxa1/Slc18a2/Acvr1c/Rapgef4/Neurod1/Madd/Nr1h3/Bloc1s6/Plcb1/Snap25/Gnas/Myt1/Foxo1/Ensa/Rap1gds1/Camk2n1/Pclo/Epha5/Cplx1/P2rx7/Aacs/Stx1a/Atg7/Cacna1c/M6pr/Abcc8/Apbb1/Slc25a22/Irs2/Rab3a/Prkaca/Vps35/Gnao1/Exph5/Cyb5r4/Myrip/Porcn/Cdk16/Sytl4/Acsl4 | 72 |

| | | | | | | | | | |
|---|---|---|---|---|---|---|---|---|---|
| BP | GO:1903532 | positive regulation of secretion by cell | 66/1983 | 373/17474 | 1,6E+12 | 0.00599503951275564 | 0.005272377703917523 | Vps4b/Tgfb2/Myb/Mcu/Trpm2/Syt1/Rtn4/Prkca/Hgs/Rab15/Gpr68/Chrm3/Bmp6/Cadps/Anxa7/Ppp3cb/Ang/Pck2/Nkx3-1/Oxct1/Ptger4/Sybu/Pla2g6/Nell2/Pfkm/Septin5/Pcp4/Glp1r/Prkce/Myo5b/Syt7/Gnaq/Stam/Stxbp1/Rapgef4/Plcb1/Snap25/Gnas/Nlgn1/Npy2r/Sdcbp/Camk2n1/Htr6/P2rx7/Aacs/Stx1a/Snca/Rab7/Oxtr/Atg7/Cacna1c/Sphk2/Abcc8/Nmb/Apbb1/Irs2/Fgfr1/Rab3a/Prkaca/Vps35/Smpd3/Exph5/Scamp5/Myrip/Sytl4/Acsl4 | 66 |
| BP | GO:0048284 | organelle fusion | 35/1983 | 165/17474 | 1,8E+12 | 0.006657143274441635 | 0.005854669297386701 | Gdap1/Rims1/Syt1/Septin8/Snap47/Vamp2/Vat1/Vps41/Cplx2/Nkd2/Erc2/Chmp7/Tfrc/Syt7/Anxa1/Stxbp1/Mtch2/Bloc1s6/Snap25/Syt11/AU040320/Cplx1/Rimbp2/Stx1a/Snca/Rab7/Arl8b/Erc1/Vamp1/Grik5/Ankrd27/Syt3/Prrt2/Bnip3/Rab3a | 35 |
| BP | GO:0051222 | positive regulation of protein transport | 60/1983 | 333/17474 | 1,9E+11 | 0.006811648449065929 | 0.005990549909365282 | Ptgs2/Tgfb2/Fyn/Mcu/Sar1a/Trpm2/Vamp2/Ywhae/Bcas3/Jup/Psen1/Gpr68/Hsp90aa1/Chrm3/Anxa7/Ppp3cb/Prkcd/Ang/Pck2/Oxct1/Ptger4/Sybu/Pla2g6/Pfkm/Cacnb3/Mapk1/Mapk14/Glp1r/Hsp90ab1/Prkce/Camk4/B3gat3/Gnaq/Lrp2/Rapgef4/Plcb1/Src/Gnas/Asph/Hpca/Camk2n1/Rufy3/P2rx7/Aacs/Tcaf1/Edem1/Atg7/Cacna1c/Abcc8/Pak1/Apbb1/Irs2/Prkaca/Vps35/Exph5/Oaz2/Tmem30a/Myrip/Sytl4/Acsl4 | 60 |
| BP | GO:0032024 | positive regulation of insulin secretion | 26/1983 | 110/17474 | 2E+11 | 0.007181311959436442 | 0.0063156529626283145 | Mcu/Trpm2/Gpr68/Chrm3/Anxa7/Ppp3cb/Pck2/Oxct1/Sybu/Pla2g6/Pfkm/Glp1r/Prkce/Gnaq/Rapgef4/Plcb1/Gnas/Camk2n1/Aacs/Atg7/Cacna1c/Abcc8/Irs2/Prkaca/Myrip/Acsl4 | 26 |
| BP | GO:0030836 | positive regulation of actin filament depolymerization | 7/1983 | 13/17474 | 2,2E+11 | 0.007753322671942588 | 0.006818711619806893 | Cfl2/Actn2/Sema5a/Cfl1/Dstn/Wdr1/Carmil2 | 7 |
| BP | GO:0043270 | positive regulation of monoatomic ion transport | 50/1983 | 266/17474 | 2,2E+12 | 0.007778911508418684 | 0.006841215893651747 | Cxcr4/Adora1/Atp1b1/Rgs7/Sgk1/Kcnc2/Ppp3r1/Kcnmb1/Gjc2/Asic2/Vmp1/Stac2/Wnk4/Hspa2/Actn2/Agtr1a/Glrx/Ppp3cb/Stc1/Fgf14/Cacng2/Cntn1/Ano6/Cacnb3/Mylk/Ifngr2/Ehd3/Cfl1/Kcnip2/Gsto1/Lrrc55/Plcb1/Gnas/Asph/Ephb2/Stim2/Tesc/P2rx7/Tcaf1/Snca/Cacna1c/Kcna1/Calm3/Scn1b/Sphk2/Abcc8/Kcnc1/P2ry6/Thy1/Htr3a | 50 |
| BP | GO:0051495 | positive regulation of cytoskeleton organization | 40/1983 | 200/17474 | 2,5E+12 | 0.008290297513073985 | 0.0072909577449445226 | Nck2/Vps4b/Wasf1/Drg1/Pfn1/Serpinf2/Bcas3/Mapt/Grb2/Scin/Cfl2/Togaram1/Actn2/Map1b/Ptk2b/Sema5a/Mapk15/Cd47/Hspa1b/Hspa1a/Prkce/Bin1/Cep120/Cfl1/Add3/Dstn/Stmn2/Stil/Wdr1/Pxn/P2rx7/Clip1/Ntf3/Cdkn1b/Cyfip1/Pak1/Mylk3/Carmil2/Nrp1/Tpm1 | 40 |
| BP | GO:0001921 | positive regulation of receptor recycling | 8/1983 | 17/17474 | 2,6E+12 | 0.008441653807926074 | 0.007424069053490163 | Psen2/Bves/Nsf/Psen1/Agtr1a/Eps15/Nsg1/Snca | 8 |
| BP | GO:0002934 | desmosome organization | 6/1983 | 10/17474 | 3E+11 | 0.009500033920880896 | 0.008354868541623605 | Jup/Prkca/Dsp/Pkp2/Dsg2/Nectin1 | 6 |
| BP | GO:0090084 | negative regulation of inclusion body assembly | 6/1983 | 10/17474 | 3E+11 | 0.009500033920880896 | 0.008354868541623605 | Hspa2/Sacs/Hspa1b/Hspa1a/Dnajb6/Dnaja4 | 6 |
| BP | GO:0099173 | postsynapse organization | 47/1983 | 249/17474 | 3,1E+11 | 0.00974588243375411 | 0.00857108166499937 | Stau2/Nrp2/Epha4/Abl2/Psen2/Fyn/Ppfia2/Camk2b/Shisa6/Pafah1b1/Srcin1/Cntnap1/Itgb3/Nptx1/Dgkb/Gphn/Psen1/Nrxn3/Rps6ka5/Arhgap22/Cpne6/Ptk2b/Dnaja3/Nrxn1/Cdh2/Myo5b/Cfl1/Nrxn2/Abhd17b/Tanc1/Rapgef4/Nlgn1/Pdlim5/Epha7/Frrs1l/Musk/Sh3gl2/Ephb2/Epha5/Rph3a/Vhl/Igf1r/Vps35/Nrp1/Hspa8/Ephb1/Tmem108 | 47 |
| BP | GO:0099590 | neurotransmitter receptor internalization | 14/1983 | 45/17474 | 3,1E+11 | 0.009750575674464414 | 0.008575209167016241 | Pip5k1c/Ppp3r1/Itgb3/Cacng5/Cacng2/Ap2m1/Synj1/Pacsin1/Snap25/Eps15/Hpca/Ap2s1/Ap2a2/Vac14 | 14 |
| BP | GO:0099601 | regulation of neurotransmitter receptor activity | 19/1983 | 72/17474 | 3,2E+12 | 0.009915817193642032 | 0.008720531929213198 | Shisa6/Cacng5/Nptx1/Rasgrf2/Ptk2b/Cacng2/Mapk8ip2/Ifngr2/Nrxn1/Neto1/Cnih2/Cfl1/Nrxn2/Src/Nlgn1/Ephb2/Rph3a/Homer3/Neto2 | 19 |
| CC | GO:0070382 | exocytic vesicle | 86/1983 | 272/17474 | 0,00149 | 0,384759235 | 3,383790892 | Atp6v1h/Atg9a/Psen2/Slc17a8/Ppfia2/Syt1/Rab3ip/Vdac1/Snap47/Rnf112/Vamp2/Atp6v0a1/Rab10/Psen1/Amph/Cltb/Tmed9/Nkd2/Mctp1/Scamp1/Synpr/Lgi3/Atp6v1c1/Syngr1/Septin5/Atp6v1a/Atp6v0c/Rab40c/Atp6v1g2/Bin1/Dpysl3/Slc6a7/Wdr7/Rab3il1/Syt7/Htr7/Pi4k2a/Slc18a2/Dnm1/Rab14/Madd/Bdnf/Snap25/Syndig1/Dnajc5/Syt11/Sv2a/Slc6a17/Sh3gl2/Atp8a1/Svop/Rab35/Pebp1/Rph3a/Stx1a/Atp6v0e2/Snca/Rab7/Syn2/Erc1/Atp6v1e1/Slc2a3/Ntf3/Borcs5/Clcn4/Calm3/Slc17a7/Abcc8/Apba2/Sv2b/Rab6a/Prrt2/Ap2a2/Atp6v1b2/Rab3a/Atp6v0d1/Hspa8/Dmxl2/Scamp5/Snap91/Bsn/Syp/Atp6ap2/Cdk16/Syn1/Sytl4 | 86 |
| CC | GO:0043209 | myelin sheath | 71/1983 | 216/17474 | 0,21866 | 28,24398522 | 248,3936222 | Hspd1/Ndufa10/Atp1b1/Atp5b/Nefh/Cnrip1/Rtn4/Canx/Vdac1/Septin8/Gjc2/Pitpna/Cntnap1/Nsf/Actg1/Hspa2/Hsp90aa1/Tkt/Nefl/Nefm/Cct5/Cntn1/Tuba1b/Tuba1a/Atp6v1a/Tagln3/Tcp1/Sod2/Pacsin1/Tubb4a/Ehd3/Napg/Atp5a1/Ehd1/Stip1/Actr1a/Ina/Gsto1/Tubb4b/Dnm1/Stxbp1/Hspa5/Pdia3/Plcb1/Snap25/Vcp/Eno1/Wdr1/Uchl1/Pebp1/Ubc/Ywhag/Immt/Syn2/Napa/Calm3/Mag/Gpi1/Myh14/Aldoa/Atp6v1b2/Gnao1/Got2/Hspa8/Thy1/Dlat/Idh3a/Pkm/Syn1/Gdi1/Gpm6b | 71 |
| CC | GO:0150034 | distal axon | 104/1983 | 394/17474 | 0,05031 | 557,0512884 | 48,99024913 | Kcnq5/Trak2/Epha4/Cxcr4/Adora1/Psen2/Grik2/Ap3d1/Slc17a8/Kcnc2/Dctn2/Ppp2ca/Fxr2/Pafah1b1/Ywhae/Atp5g1/Mapt/Prkca/Actg1/Psen1/Hsp90aa1/Klc1/Chrm3/Amph/Scgn/Nrsn1/Cplx2/Map1b/Snx18/Erc2/Ang/Ptk2b/Nefl/Pcdh9/Rtn4r/Septin5/Pacsin1/Hsp90ab1/Nrxn1/Bin1/Dpysl3/Eno1b/Dcc/Cfl1/Syt7/Htr7/Zfyve27/Crtac1/Got1/Slc18a2/Olfm1/Kif5c/Tanc1/Lrp2/Rapgef4/Mapk8ip1/Bdnf/Lamp5/Snap25/Flrt3/Src/Stmn2/Syt11/Unc5c/Eno1/Kcnab2/Pclo/Rufy3/Cplx1/Pebp1/P2rx7/Rimbp2/Auts2/Orai2/Snca/Aak1/Grm7/Kcna1/Kcna6/Slc8a2/Calm3/Grik5/Kcnc3/Myh14/Kcnc1/Cyfip1/Cpeb1/Pak1/Apbb1/Prrt2/Rab3a/Amfr/Cdh8/Atp6v0d1/Nrp1/Hspa8/Grik4/Nectin1/Thy1/Snap91/Syp/Frmd7/Slc9a6/Fgf13 | 104 |

| | | | | | | | | | |
|---|---|---|---|---|---|---|---|---|---|
| CC | GO:0098984 | neuron to neuron synapse | 113/1983 | 495/17474 | 146,352 | 11342,29812 | 99750,60147 | Rims1/Il1r1/Nck2/Epha4/Adora1/Grm1/Fyn/Grik2/Pip5k1c/Syt1/Nefh/Camk2b/Ppp3r1/Rtn4/Ppp2ca/Vdac1/Snap47/Rnf112/Shisa6/Fxr2/Dynll2/Srcin1/Stat3/Nsf/Mapt/Cacng5/Dtnb/Dgkb/Rtn1/Gphn/Chrm3/Actn2/Rgs14/Map1b/Synpr/Erc2/Ptk2b/Nefm/Cdh9/Ywhaz/Cacng2/Mapk8ip2/Mapk1/Fbxo45/Nectin3/Pacsin1/Nrxn1/Cdh2/Dcc/Neto1/Cnih2/Rtn3/Dagla/Syt7/Abhd17b/Ina/Neurl1a/Add3/Ablim1/Nsmf/Pkp4/Tanc1/Rapgef4/Bdnf/Flrt3/Syndig1/Src/Bcas1/Nlgn1/Syt11/Pdlim5/Sdcbp/Epha7/Sh3gl2/Elavl2/Dnajc6/Ephb2/Camk2n1/Capzb/Kcnab2/Pclo/Ywhah/Cabp1/Stx1a/Cald1/Sema4f/Grm7/Vhl/Syn2/Cacna1c/Erc1/Chd4/Slc8a2/Grik5/Cpeb1/Sh3gl3/Pak1/Bnip3/Homer3/Vps35/Neto2/Nrgn/Hspa8/Grik4/Nectin1/Nptn/Map2k1/Snap91/Clstn2/Tmem108/Bsn/Syp/Syn1 | 113 |
| CC | GO:0014069 | postsynaptic density | 98/1983 | 437/17474 | 186606 | 9112087,441 | 8013686,409 | Rims1/Il1r1/Nck2/Epha4/Adora1/Grm1/Fyn/Grik2/Pip5k1c/Nefh/Camk2b/Rtn4/Ppp2ca/Vdac1/Snap47/Rnf112/Shisa6/Fxr2/Dynll2/Srcin1/Stat3/Nsf/Mapt/Cacng5/Dtnb/Dgkb/Rtn1/Gphn/Chrm3/Actn2/Rgs14/Map1b/Erc2/Ptk2b/Nefm/Ywhaz/Cacng2/Mapk8ip2/Mapk1/Fbxo45/Nectin3/Pacsin1/Cdh2/Dcc/Neto1/Cnih2/Rtn3/Dagla/Abhd17b/Ina/Neurl1a/Add3/Ablim1/Nsmf/Pkp4/Tanc1/Rapgef4/Flrt3/Syndig1/Src/Bcas1/Syt11/Pdlim5/Sdcbp/Epha7/Dnajc6/Camk2n1/Capzb/Kcnab2/Pclo/Cabp1/Stx1a/Cald1/Sema4f/Vhl/Syn2/Cacna1c/Erc1/Slc8a2/Grik5/Cpeb1/Sh3gl3/Pak1/Bnip3/Homer3/Vps35/Neto2/Nrgn/Hspa8/Grik4/Nptn/Map2k1/Snap91/Clstn2/Tmem108/Bsn/Syp/Syn1 | 98 |
| CC | GO:0030427 | site of polarized growth | 60/1983 | 220/17474 | 517157 | 210945517,4 | 18551744,99 | Trak2/Epha4/Cxcr4/Psen2/Dctn2/Fxr2/Pafah1b1/Ywhae/Mapt/Psen1/Hsp90aa1/Klc1/Nrsn1/Map1b/Snx18/Erc2/Ang/Ptk2b/Nefl/Pcdh9/Rtn4r/Hsp90ab1/Nrxn1/Dpysl3/Eno1b/Dcc/Cfl1/Pi4k2a/Zfyve27/Crtac1/Olfm1/Kif5c/Lrp2/Rapgef4/Mapk8ip1/Lamp5/Snap25/Flrt3/Src/Stmn2/Unc5c/Eno1/Pclo/Rufy3/Auts2/Orai2/Snca/Calm3/Myh14/Cyfip1/Cpeb1/Pak1/Apbb1/Amfr/Cotl1/Nrp1/Nectin1/Thy1/Frmd7/Fgf13 | 60 |
| CC | GO:0044306 | neuron projection terminus | 57/1983 | 222/17474 | 1,9E+07 | 520898521,7 | 458107698,2 | Kcnq5/Epha4/Adora1/Grik2/Ap3d1/Slc17a8/Syt1/Kcnc2/Ppp2ca/Vamp2/Prkca/Actg1/Chrm3/Amph/Scgn/Cplx2/Erc2/Ptger4/Septin5/Pacsin1/Bin1/Syt7/Htr7/Got1/Slc18a2/Tanc1/Rapgef4/Bdnf/Flrt3/Syt11/Kcnab2/Pclo/Uchl1/Cplx1/Pebp1/P2rx7/Rimbp2/Snca/Aak1/Grm7/Kcna1/Kcna6/Slc8a2/Grik5/Kcnc3/Kcnc1/Cyfip1/Prrt2/Rab3a/Cdh8/Atp6v0d1/Hspa8/Grik4/Snap91/Bsn/Syp/Slc9a6 | 57 |
| CC | GO:0043679 | axon terminus | 52/1983 | 200/17474 | 6015077 | 15248069135 | 13410016658 | Kcnq5/Epha4/Adora1/Grik2/Ap3d1/Slc17a8/Kcnc2/Ppp2ca/Prkca/Actg1/Chrm3/Amph/Scgn/Cplx2/Erc2/Septin5/Pacsin1/Bin1/Syt7/Htr7/Got1/Slc18a2/Tanc1/Rapgef4/Bdnf/Flrt3/Syt11/Kcnab2/Pclo/Cplx1/Pebp1/P2rx7/Rimbp2/Snca/Aak1/Grm7/Kcna1/Kcna6/Slc8a2/Grik5/Kcnc3/Kcnc1/Cyfip1/Prrt2/Rab3a/Cdh8/Atp6v0d1/Hspa8/Grik4/Snap91/Syp/Slc9a6 | 52 |
| CC | GO:0042734 | presynaptic membrane | 57/1983 | 238/17474 | 2,8E+08 | 5905240248 | 5193403139 | Kcnq5/Rims1/Epha4/Adora1/Cadm3/Psen2/Kcnh1/Grm1/Grik2/Pip5k1c/Ppfia2/Syt1/Kcnc2/Canx/Vdac1/Flot2/Psen1/Nrxn3/Chrm3/Cltb/Erc2/Cdh9/Cntn1/Ap2m1/Nrxn1/Cdh2/Slc6a7/Nrxn2/Syt7/Htr7/Pi4k2a/Dnm1/Stxbp1/Snap25/Syt11/Gabbr2/Dnajc6/Ephb2/Stx1a/Grm7/Cacna1c/Kcna1/Napa/Ap2s1/Grik5/Kcnc3/Kcnc1/Gabra5/Apbb1/Prrt2/Cacna1a/Grik4/Nectin1/Htr3a/Nptn/Snap91/Syp | 57 |
| CC | GO:0019898 | extrinsic component of membrane | 67/1983 | 313/17474 | 1,9E+09 | 28338952896 | 2,49229E+11 | Atp6v1h/Rims1/Zap70/Rgs8/Coq8a/Fyn/Gnaz/Socs2/Arhgef25/Rnf112/Alox8/Socs7/Stac2/Jup/Becn1/Hid1/Osr1/Dgkb/Gphn/Amph/Snx18/Cadps/Gng2/Cdh24/Ptk2b/Cdh9/Cdh12/Atp6v1c1/Dnaja3/Ppl/Ap2m1/Atp6v1a/Atp6v1g2/Cdh2/Pik3c3/Bin1/Gnaq/Anxa1/Olfm1/Stxbp1/Bloc1s6/Snap25/Src/Gnas/Dnajc6/Hpca/Fbxo2/Errfi1/Kcnab2/Cabp1/Rph3a/Ubc/Snx10/Mgll/Syn2/Atp6v1e1/Ap2s1/Atp6v1b2/Gnao1/Cdh8/Vac14/Snap91/Trf/Bsn/Cdk16/Syn1/Sytl4 | 67 |
| CC | GO:0098685 | Schaffer collateral - CA1 synapse | 38/1983 | 143/17474 | 3,5E+09 | 5,18297E+11 | 45581992544 | Epha4/Grm1/Fyn/Nefh/Ppp3r1/Pfn1/Pafah1b1/Stat3/Actg1/Rhob/Dgkb/Ptk2b/Nefl/Pcdh8/Cacng2/Synj1/Nrxn1/Slc6a7/Dcc/Myo5b/Neto1/Ina/Ptpra/Syt11/Epha7/Sh3gl2/Capzb/Cplx1/Stx1a/Syn2/Apba2/Apbb1/Nptn/Snap91/Bsn/Syp/Syn1/Slc9a6 | 38 |
| CC | GO:0031594 | neuromuscular junction | 31/1983 | 108/17474 | 6,8E+08 | 94376573922 | 8300011085 | Epha4/Hdac4/Psen2/Ascc1/Itga7/Psen1/Spock1/Nefl/Nefm/Syngr1/Tuba1a/Dnaja3/Prkce/Nrxn1/Chrna1/Snta1/Dnajc5/Nlgn1/Sv2a/Epha7/Musk/Pclo/Rph3a/P2rx7/Vamp1/Napa/Kcnc3/Apbb1/Prkaca/Cib2/Syp | 31 |
| CC | GO:0045211 | postsynaptic membrane | 70/1983 | 345/17474 | 8,5E+08 | 1,11965E+12 | 98468367978 | Nrp2/Epha4/Adora1/Rgs7/Kcnh1/Grm1/Grik2/Kcnc2/Rtn4/Canx/Vdac1/Shisa6/Itgb3/Cacng5/Dgkb/Gphn/Chrm3/Actn2/Abhd6/Pcdh8/Cdh9/Cacng2/Cntn1/Faim2/Dnaja3/Fbxo45/Nectin3/Pacsin1/Cdh2/Dcc/Neto1/Cnih2/Dagla/Abhd17b/Htr7/Chrna1/Flrt3/Syndig1/Snta1/Nlgn1/Epha7/Gabbr2/Musk/Ephb2/Fbxo2/Kcnab2/Nsg1/Rph3a/Stx1a/Sema4f/Grm7/Cacna1c/Kcna1/Slc8a2/Grik5/Kcnc3/Kcnc1/Gabra5/Apbb1/Prrt2/Neto2/Nrp1/Nrgn/Hspa8/Grik4/Htr3a/Nptn/Snap91/Clstn2/Atp6ap2 | 70 |
| CC | GO:0048786 | presynaptic active zone | 33/1983 | 122/17474 | 1,3E+09 | 1,58453E+12 | 1,39352E+12 | Rims1/Adora1/Ppfia4/Ppfia2/Canx/Vdac1/Snap47/Flot2/Cntnap1/Erc2/Nrxn1/Cdh2/Stxbp1/Snap25/Syt11/Sv2a/Pclo/Rimbp2/Stx1a/Grm7/Erc1/Napa/Slc17a7/Ppfia3/Ctbp2/Rab3a/Cacna1a/Nectin1/Nptn/Snap91/Bsn/Syp/Syn1 | 33 |

| | | | | | | | | |
|---|---|---|---|---|---|---|---|---|
| CC | GO:0010008 | endosome membrane | 58/1983 | 279/17474 | 3,3E+09 | 3,12544E+11 | 2748688282 | Fzd7/Mreg/Atg9a/Epha4/Bok/Vps4b/Tmem9/Kcnh1/Fig4/Ap3d1/Pip5k1c/Washc4/Rab21/Pmel/Anxa6/Rhob/Rab15/Vps41/Scamp1/Snx18/Abhd6/Chmp7/Slc39a14/Cdip1/Lztr1/Tfrc/Washc1/Ehd3/Rmc1/Myo5b/Ehd1/Dagla/Abhd17b/Pi4k2a/Zfyve27/Lamp5/Syndig1/Atp9a/Rap2b/Tyrp1/Ncdn/Nsg1/Rab35/Snx10/Rab7/Clcn4/Vps35/Vac14/Ldlr/Scamp5/Tmem30a/Snap91/Ephb1/Tmem108/Atp6ap2/Uba1/Rap2c/Slc9a6 | 58 |
| CC | GO:0099568 | cytoplasmic region | 48/1983 | 221/17474 | 6,6E+09 | 5,3549E+11 | 4,7094E+11 | Efhc1/Rims1/Spag16/Kifap3/Uhmk1/Atg5/Grik2/Canx/Drc3/Map2k4/Cfap52/Mapt/Ccdc40/Togaram1/Cltb/Cfap70/Dnah12/Erc2/Dnah5/Rangap1/Spag6l/Mapk1/Tubb4a/Myo5b/Actr1a/Stxbp1/Madd/Abhd12/Cfap206/Dnali1/Hpca/Pclo/Lrrc43/Dnah10/Rimbp2/Dync1i1/Atg7/Erc1/Ppfia3/Dnah3/Ctbp2/Prkaca/Hydin/Map2k1/Mns1/Bsn/Prkar2a/Akap14 | 48 |
| CC | GO:0099569 | presynaptic cytoskeleton | 8/1983 | 12/17474 | 8,8E+09 | 6,8268E+11 | 6,00388E+11 | Rims1/Actg1/Erc2/Nefl/Nefm/Pclo/Erc1/Bsn | 8 |
| CC | GO:0034702 | ion channel complex | 59/1983 | 295/17474 | 9,6E+09 | 7,08143E+11 | 6,22781E+11 | Kcnq5/Kcnh1/Ostm1/Grik2/Mcu/Micu1/Slc17a8/Kcnc2/Kcnmb4/Kcnmb1/Shisa6/Atp5g1/Cacng5/Ttyh2/Hspa2/Nrn1/Abhd6/Cacna2d3/Ptk2b/Kcns2/Kcnv1/Cacng2/Ano6/Cacnb3/Kcng2/Cnih2/Catsperz/Kcnk4/Kcnip2/Olfm1/Ptpa/Scn7a/Chrna1/Lrrc55/Snap25/Abhd12/Cachd1/Kcnab2/Lrrc8d/Cacna1c/Kcna1/Kcna6/Calm3/Grik5/Scn1b/Kcnc3/Cpt1c/Slc17a7/Abcc8/Kcnc1/Gabra5/Micu3/Cacna1a/Kcng4/Kcnk1/Grik4/Scn2b/Htr3a/Porcn | 59 |
| CC | GO:1990351 | transporter complex | 71/1983 | 397/17474 | 6,5E+10 | 0.00305358729369481 | 0.0026854978236568964 | Atp6v1h/Kcnq5/Ndufa10/Atp1b1/Kcnh1/Ostm1/Grik2/Mcu/Micu1/Slc17a8/Kcnc2/Kcnmb4/Pex13/Kcnmb1/Shisa6/Atp5g1/Cacng5/Ttyh2/Hspa2/Nrn1/Abhd6/Cacna2d3/Ptk2b/Kcns2/Atp6v1c1/Kcnv1/Cacng2/Ano6/Cacnb3/Atp6v1a/Atp6v0c/Kcng2/Cnih2/Catsperz/Kcnk4/Kcnip2/Olfm1/Ptpa/Scn7a/Chrna1/Lrrc55/Snap25/Abhd12/Cachd1/Kcnab2/Atp8a1/Lrrc8d/Cacna1c/Kcna1/Kcna6/Calm3/Grik5/Scn1b/Kcnc3/Cpt1c/Slc17a7/Abcc8/Kcnc1/Gabra5/Wdr93/Micu3/Cacna1a/Atp6v0d1/Kcng4/Kcnk1/Grik4/Scn2b/Htr3a/Tmem30a/Porcn/Timm8a1 | 71 |
| CC | GO:0060076 | excitatory synapse | 25/1983 | 98/17474 | 7,1E+07 | 0.003260352378352604 | 0.002867338763984123 | Ppfia4/Cadm3/Kcnh1/Slc17a8/Syt1/Ppp3r1/Shisa6/Pfn1/Synj1/Nrxn1/Neto1/Stxbp1/Rapgef4/Syndig1/Nlgn1/Syt11/Elavl2/Pclo/Ywhah/Chd4/Slc17a7/Cyfip1/Snap91/Bsn/Syp | 25 |
| CC | GO:0016471 | vacuolar proton-transporting V-type ATPase complex | 10/1983 | 23/17474 | 9,8E+10 | 0.004154198831536309 | 0.003653438021622763 | Atp6v1h/Atp6v0a1/Atp6v1c1/Atp6v1a/Atp6v0c/Atp6v1g2/Atp6v1e1/Atp6v1b2/Atp6v0d1/Atp6ap2 | 10 |
| CC | GO:0098878 | neurotransmitter receptor complex | 15/1983 | 46/17474 | 1E+12 | 0.004384075877730976 | 0.0038556049315189227 | Grik2/Shisa6/Cacng5/Nrn1/Abhd6/Ptk2b/Cacng2/Cnih2/Olfm1/Abhd12/Grik5/Cpt1c/Grik4/Htr3a/Porcn | 15 |
| CC | GO:0033176 | proton-transporting V-type ATPase complex | 11/1983 | 28/17474 | 1,3E+12 | 0.005210957607762797 | 0.0045828116142973335 | Atp6v1h/Atp6v0a1/Atp6v1c1/Atp6v1a/Atp6v0c/Atp6v1g2/Atp6v0e2/Atp6v1e1/Atp6v1b2/Atp6v0d1/Atp6ap2 | 11 |
| CC | GO:0000407 | phagophore assembly site | 12/1983 | 33/17474 | 1,6E+12 | 0.00599503951275564 | 0.005272377703917523 | Atg9a/Atg5/Sqstm1/Bcas3/Vmp1/Becn1/Ilrun/Pik3c3/Atg12/Snx7/Rab7/Atg7 | 12 |
| CC | GO:0019897 | extrinsic component of plasma membrane | 37/1983 | 177/17474 | 1,6E+12 | 0.006169374293939608 | 0.0054256975963679404 | Rims1/Zap70/Rgs8/Fyn/Gnaz/Stac2/Jup/Dgkb/Gphn/Snx18/Gng2/Cdh24/Ptk2b/Cdh9/Cdh12/Dnaja3/Ppl/Ap2m1/Cdh2/Gnaq/Anxa1/Olfm1/Stxbp1/Snap25/Src/Gnas/Dnajc6/Fbxo2/Errfi1/Kcnab2/Cabp1/Ap2s1/Gnao1/Cdh8/Snap91/Trf/Cdk16 | 37 |
| CC | GO:0031312 | extrinsic component of organelle membrane | 17/1983 | 59/17474 | 2,1E+12 | 0.007432342820741557 | 0.0065364237370868035 | Atp6v1h/Coq8a/Hid1/Amph/Cadps/Atp6v1c1/Atp6v1a/Atp6v1g2/Bin1/Rph3a/Ubc/Snx10/Syn2/Atp6v1e1/Atp6v1b2/Snap91/Syn1 | 17 |
| CC | GO:0030136 | clathrin-coated vesicle | 27/1983 | 117/17474 | 2,3E+11 | 0.00792344216469577 | 0.006968324348577944 | Clvs2/Slc17a8/Ap1b1/Clint1/Epn2/Vamp2/Vps41/Cltb/Scamp1/Snx18/Cpne6/Ap2m1/Rab14/Dnajc5/Clrn1/Tyrp1/Eps15/Rab35/Tgoln1/Necap1/Ap2s1/Ap2a2/Rab3a/Nrgn/Snap91/Syp/Syn1 | 27 |
| CC | GO:0008328 | ionotropic glutamate receptor complex | 14/1983 | 44/17474 | 2,4E+12 | 0.00805893038592512 | 0.0070874803733602925 | Grik2/Shisa6/Cacng5/Nrn1/Abhd6/Ptk2b/Cacng2/Cnih2/Olfm1/Abhd12/Grik5/Cpt1c/Grik4/Porcn | 14 |
| CC | GO:0098562 | cytoplasmic side of membrane | 40/1983 | 202/17474 | 3,1E+11 | 0.009731223633339909 | 0.008558189884669054 | Ptp4a1/Zap70/Rgs8/Fyn/Gnaz/Map2k2/Rab21/Stac2/Jup/Mapt/Gphn/Traf3/Snx18/Gng2/Ptk2b/Chmp7/Otulinl/Cdip1/Ap2m1/Gnaq/Borcs7/Traf1/Pkp4/Snap25/Src/Gnas/Ptpn3/Errfi1/Kcnab2/Ajap1/Kit/Sppl3/Cabp1/Prmt8/Borcs5/Ap2s1/Ap2a2/Rasa3/Gnao1/Cdk16 | 40 |
| MF | GO:0051082 | unfolded protein binding | 26/1983 | 87/17474 | 2,3E+09 | 25558587528 | 2,24777E+12 | Nudcd3/Erlec1/Canx/Hspa2/Hsp90aa1/Cct5/Dnaja3/Cct8/Tcp1/Hspa1b/Hspa1a/Hsp90ab1/Hspa5/Ndufaf1/Mkks/Dnaja1/Ube4b/Dnajb6/Cct6a/Shq1/Calr/Dnaja2/Hspa8/Dnaja4/Clpx/Vbp1 | 26 |
| MF | GO:0044389 | ubiquitin-like protein ligase binding | 65/1983 | 323/17474 | 2,8E+10 | 28723362184 | 2,5261E+11 | Ube2w/Bag2/Hspd1/Psmd1/Bok/Cxcr4/Grik2/Ube2n/Rtn4/Sqstm1/Ywhae/Becn1/Actg1/Hgs/Trib2/Psma3/Hsp90aa1/Traf3/Nkd2/Ptk2b/Ywhaz/Derl1/Rangap1/Tuba1b/Ube2l3/Dtx3l/Tcp1/Atp6v0c/Hspa1b/Hspa1a/Tubb5/Hsp90ab1/Washc1/Lrpprc/Otub1/Stam/Hspa5/Traf1/Nek6/Ube2l6/Foxo1/Syt11/Ube2j1/Dnaja1/Vcp/Jun/Uchl1/Chek2/Rnf34/Ubc/Flt3/Gpr37/Rad18/Bid/Cdkn1b/Gpi1/Apbb1/Tollip/Mfhas1/Prkaca/Calr/Hspa8/Ubash3b/Prkar2a/Sh3kbp1 | 65 |
| MF | GO:0048156 | tau protein binding | 11/1983 | 22/17474 | 8,4E+09 | 6,61044E+11 | 5813595122 | Bag2/Sgk1/Fyn/Ppp2ca/Hspa2/Hsp90aa1/Ppp2r2a/Hsp90ab1/Bin1/Snca/Apbb1 | 11 |
| MF | GO:0005200 | structural constituent of cytoskeleton | 21/1983 | 68/17474 | 1,2E+11 | 84004102468 | 7,3878E+11 | Tuba4a/Nefh/Camk2b/Actg1/Tubb2a/Nefl/Nefm/Tuba1b/Tuba1a/Tubb5/Tubb4a/Ina/Add3/Vim/Tubb4b/Lmna/Arpc4/Tuba8/Myom2/Tpm1/Tln2 | 21 |
| MF | GO:0042625 | ATPase-coupled ion transmembrane transporter activity | 11/1983 | 23/17474 | 1,4E+11 | 93637834029 | 82350421099 | Atp6v1h/Atp5b/Atp6v0a1/Atp6v1c1/Atp6v1a/Atp6v0c/Atp6v1g2/Atp6v0e2/Atp6v1e1/Atp6v1b2/Atp6v0d1 | 11 |
| MF | GO:0044769 | ATPase activity, coupled to transmembrane movement of ions, rotational mechanism | 11/1983 | 23/17474 | 1,4E+11 | 93637834029 | 82350421099 | Atp6v1h/Atp5b/Atp6v0a1/Atp6v1c1/Atp6v1a/Atp6v0c/Atp6v1g2/Atp6v0e2/Atp6v1e1/Atp6v1b2/Atp6v0d1 | 11 |

| | | | | | | | | | |
|---|---|---|---|---|---|---|---|---|---|
| MF | GO:0046961 | proton-transporting ATPase activity, rotational mechanism | 11/1983 | 23/17474 | 1,4E+11 | 93637834029 | 82350421099 | Atp6v1h/Atp5b/Atp6v0a1/Atp6v1c1/Atp6v1a/Atp6v0c/Atp6v1g2/Atp6v0e2/Atp6v1e1/Atp6v1b2/Atp6v0d1 | 11 |
| MF | GO:0140662 | ATP-dependent protein folding chaperone | 11/1983 | 23/17474 | 1,4E+11 | 93637834029 | 82350421099 | Hspd1/Hspa4/Hspa2/Cct5/Cct8/Tcp1/Hspa5/Cct6a/Hsph1/Hspa8/Clpx | 11 |
| MF | GO:0030674 | protein-macromolecule adaptor activity | 64/1983 | 346/17474 | 5,3E+11 | 0.002638781532677998 | 0.0023206941153263204 | Retreg2/Daw1/Nmnat2/Atp1b1/Serinc1/Map2k2/Socs2/Nefh/Gas2l1/Sqstm1/Skp1/Cdc42se2/Snap47/Vamp2/Sarm1/Poldip2/Jup/Stat3/Grb2/Klhdc2/Ankrd9/Actn2/Rgs14/Nefl/Retreg1/Tonsl/Mapk8ip2/Ap2m1/Dtx3l/Fem1a/Napg/Ddb1/Blnk/Optn/Sh2d3c/Grb14/Rapgef4/Mapk8ip1/Snap25/Trpc4ap/Bank1/Stx18/Pxn/Stx1a/Arpc4/Vhl/Vamp1/Napa/Ap2s1/Akap13/Klhl25/Bet1l/Ap2a2/Irs2/Gatad2a/Rab3a/Amfr/Tradd/Wdr59/Hspa8/Map2k1/Snap91/Ppp2r3a/Ubqln2 | 64 |
| MF | GO:0016247 | channel regulator activity | 33/1983 | 144/17474 | 5,7E+10 | 0.0027800335947714807 | 0.002444919188610572 | Sgk1/Kcnmb4/Kcnmb1/Ywhae/Wnk4/Cacng5/Nrxn3/Rem2/Lynx1/Cacng2/Cacnb3/Nrxn1/Cnih2/Nrxn2/Prkg1/Kcnip2/Lrrc55/Plcb1/Snta1/Ensa/Ptpn3/Kcnab2/Ywhah/Stim2/Cabp1/Stx1a/Grm7/Fxyd7/Scn1b/Abcc8/Scn2b/Stoml1/Fgf13 | 33 |
| MF | GO:0031072 | heat shock protein binding | 33/1983 | 145/17474 | 6,6E+10 | 0.0030971600026885033 | 0.0027238181347922657 | Stau2/Bag2/Adora1/Cd24a/Ogdh/Mapt/Hspa2/Lman2/Sacs/Dnaja3/Tfrc/Iqcg/Fkbp5/Hspa1b/Hspa1a/Hsp90ab1/Eno1b/Stip1/Slc18a2/Hspa5/Dnajc10/Tomm34/Dnaja1/Eno1/Dnajb6/Gpr37/Snca/Cdkn1b/Spn/Dnaja2/Mvd/Hspa8/Dnaja4 | 33 |
| MF | GO:0016208 | AMP binding | 9/1983 | 19/17474 | 9,8E+10 | 0.004154198831536309 | 0.003653438021622763 | Pfkl/Pygl/Pfkp/Cyb5r3/Pfkm/Mpped2/Prkag2/Aprt/Prps1 | 9 |
| MF | GO:0098918 | structural constituent of synapse | 13/1983 | 37/17474 | 1,3E+11 | 0.005149458224226172 | 0.004528725569013848 | Rims1/Ppfia2/Nefh/Camk2b/Actg1/Erc2/Nefl/Ina/Pclo/Rimbp2/Erc1/Ctbp2/Bsn | 13 |
| MF | GO:0009678 | pyrophosphate hydrolysis-driven proton transmembrane transporter activity | 11/1983 | 28/17474 | 1,3E+12 | 0.005210957607762797 | 0.0045828116142973335 | Atp6v1h/Atp5b/Atp6v0a1/Atp6v1c1/Atp6v1a/Atp6v0c/Atp6v1g2/Atp6v0e2/Atp6v1e1/Atp6v1b2/Atp6v0d1 | 11 |
| MF | GO:0044325 | transmembrane transporter binding | 33/1983 | 151/17474 | 1,5E+12 | 0.005826846252557091 | 0.005124459013284505 | Rims1/Bag2/Kcnh1/Fyn/Kcnc2/Ppp2ca/Vdac1/Vamp2/Ywhae/Hsp90aa1/Actn2/Ywhaz/Grina/Ap2m1/Hsp90ab1/Kcnip2/Lrrc55/Snap25/Snta1/Src/Kcnab2/Ywhah/Cabp1/Stx1a/Tcaf1/Cacna1c/Calm3/Abcc8/Kcnc1/Kcng4/Panx1/Nptn/Fgf13 | 33 |
| MF | GO:0005261 | monoatomic cation channel activity | 56/1983 | 305/17474 | 1,8E+12 | 0.006720750923804853 | 0.005910609471190006 | Kcnq5/Kcnh1/Calhm5/Grik2/Mcu/Trpm2/Kcnc2/Kcnmb4/Atp5b/Kcnmb1/Anxa6/Asic2/Cacng5/Psen1/Tmem63c/Cacna2d3/Kcns2/Kcnv1/Cacng2/Cacnb3/Atp6v1a/Atp5a1/Kcng2/Kcnk4/Trpm3/Kcnip2/Scn7a/Chrna1/Lrrc55/Snap25/Slc24a2/Cachd1/Kcnab2/Stim2/P2rx7/Rimbp2/Orai2/Sec61a1/Grm7/Cacna1c/Kcna1/Kcna6/Grik5/Scn1b/Kcnc3/Abcc8/Kcnc1/Rasa3/Tmem38a/Cacna1a/Kcng4/Kcnk1/Panx1/Grik4/Scn2b/Htr3a | 56 |
| MF | GO:0017075 | syntaxin-1 binding | 10/1983 | 25/17474 | 2,3E+12 | 0.007783095264088441 | 0.006844895325632957 | Syt1/Vamp2/Nsf/Cplx2/Sybu/Stxbp1/Snap25/Cplx1/Prrt2/Syp | 10 |
| MF | GO:0035254 | glutamate receptor binding | 19/1983 | 71/17474 | 2,6E+12 | 0.0085696704132066 | 0.007536654115519557 | Il1r1/Fyn/Sqstm1/Canx/Shisa6/Flot2/Nsf/Cacng2/Myo5b/Neto1/Syndig1/Gnas/Sdcbp/Nsg1/Cabp1/Calm3/Homer3/Neto2/Necab2 | 19 |
| MF | GO:0044183 | protein folding chaperone | 16/1983 | 55/17474 | 2,8E+12 | 0.009142561324263915 | 0.008040486869216821 | Hspd1/Hspa4/Hspa2/Hsp90aa1/Cct5/Cct8/Tcp1/Hspa1b/Hspa1a/Hsp90ab1/Hspa5/Dnajb6/Cct6a/Hsph1/Hspa8/Clpx | 16 |

GO_DG_mRNA

| ONTOLOGY | ID | Description | GeneRatio | BgRatio | pvalue | p.adjust | qvalue | geneID | Count |
|---|---|---|---|---|---|---|---|---|---|
| CC | GO:0098984 | neuron to neuron synapse | 113/1933 | 495/17474 | 2,55E+02 | 1,97E+06 | 1,80E+03 | Il1r1/Nck2/Als2/Epha4/C1ql2/Stxbp5/Grm1/Akap7/Grik2/Ank3/Syn3/Camk2b/Adcy1/Cpeb4/Hnrnph1/Rnf112/Abr/Nsf/Cacng5/Dtnb/Dicer1/Actn2/Map1b/Rgs7bp/Synpr/Camk2g/Dlg5/Erc2/Ptk2b/Pdzd2/Cdh9/Rnf19a/Cacng2/Rogdi/Shisa9/Mapk1/Dgcr8/Nectin3/Tiam1/Lrfn2/Sh3gl1/Nrxn1/Mib1/Cdh2/Dcc/Neto1/Dagla/Syt7/Neurl1a/Add3/Ablim1/Itga8/C1ql3/Tanc1/Hnrnpa3/Kcna4/Bdnf/Usp50/Pdyn/Lzts3/Flrt3/Src/Slitrk3/Atp1a1/Kcnd3/Plppr4/Sdcbp/Calb1/Epha7/Mpdz/Lrp8/Grik3/Epb41/Ephb2/Camk2n1/Akap9/Pclo/Slc30a3/Ywhah/Hnrnpd/Ppp1r9a/Lrrc4/Smo/Plxna4/Cald1/Dgki/Lrrtm4/Sema4f/Add2/Grm7/Syn2/Cacna1c/Erc1/Chd4/Ptpro/Slc8a2/Cpeb1/Eef2k/Gsg1l/Sorbs2/Homer3/Disc1/Grik4/Nectin1/Drd2/Map2k1/Snap91/Clstn2/Tmem108/Ctnnb1/Il1rapl1/Dlg3/Ogt | 113 |
| CC | GO:0045211 | postsynaptic membrane | 77/1933 | 345/17474 | 1,08E+05 | 2,78E+10 | 2,54E+10 | Nrp2/Epha4/Cacna1e/Rgs7/Kcnh1/Syne1/Grm1/Grik2/Ank3/Adcy1/Cacng5/Kcnj2/Actn2/Rgs7bp/Pcdh8/Pcdh17/Kctd12/Cdh9/Cacng2/Kcnj4/Cntn1/Shisa9/Nectin3/Tiam1/Cacna1h/Lrfn2/Cdh2/Pcdhb16/Kctd16/Dcc/Neto1/Dagla/Itga8/Chrna1/Kcna4/Lin7c/Flrt3/Slitrk3/Kcnd3/Kcnc4/Plppr4/Epha7/Gabbr2/Grik3/Ephb2/Gabrd/Akap9/Cacna2d1/Gabra4/Gabrb1/Rph3a/Lrrc4/Dgki/Lrrtm4/Sema4f/Grm7/Cacna1c/Kcna1/Ptpro/Slc8a2/Kcnc3/Kcnc1/Gabrb3/Ntrk3/Gsg1l/Prrt2/Glra3/Nrp1/Grik4/Drd2/Snap91/Clstn2/Ctnnb1/Slitrk2/Il1rapl1/Dlg3/Glra2 | 77 |
| CC | GO:0014069 | postsynaptic density | 91/1933 | 437/17474 | 1,50E+07 | 2,88E+10 | 2,63E+10 | Il1r1/Nck2/Als2/Epha4/Grm1/Grik2/Ank3/Syn3/Camk2b/Adcy1/Cpeb4/Hnrnph1/Rnf112/Abr/Nsf/Cacng5/Dtnb/Dicer1/Actn2/Map1b/Rgs7bp/Camk2g/Dlg5/Erc2/Ptk2b/Pdzd2/Cacng2/Shisa9/Mapk1/Dgcr8/Nectin3/Tiam1/Lrfn2/Sh3gl1/Mib1/Cdh2/Dcc/Neto1/Dagla/Neurl1a/Add3/Ablim1/Itga8/Tanc1/Hnrnpa3/Usp50/Lzts3/Flrt3/Src/Slitrk3/Atp1a1/Kcnd3/Plppr4/Sdcbp/Epha7/Mpdz/Lrp8/Grik3/Epb41/Camk2n1/Akap9/Pclo/Hnrnpd/Ppp1r9a/Lrrc4/Smo/Cald1/Dgki/Lrrtm4/Sema4f/Add2/Syn2/Cacna1c/Erc1/Ptpro/Slc8a2/Cpeb1/Eef2k/Gsg1l/Sorbs2/Homer3/Disc1/Grik4/Drd2/Map2k1/Snap91/Clstn2/Tmem108/Ctnnb1/Il1rapl1/Dlg3 | 91 |
| CC | GO:0150034 | distal axon | 81/1933 | 394/17474 | 2,17E+08 | 2,43E+10 | 2,22E+10 | Als2/Epha4/Cxcr4/Adcy10/Grik2/Sirt1/Hap1/Psen1/Dicer1/Amph/Scgn/Cplx2/Map1b/Erc2/Ptk2b/Pcdh9/Slc2a13/Rtn4r/Septin5/Tiam1/Itsn1/Dscam/Dynlt1a/Nrxn1/Dcc/Syt7/Stx3/Slc18a2/Dbh/Olfm1/Tanc1/Slc4a10/Bdnf/Pdyn/Lamp5/Flrt3/Src/Kcnc4/Ptbp2/Unc5c/Calb1/C9orf72/Abitram/Grik3/Hnrnpr/Sri/Pclo/Cad/Rimbp2/Auts2/Orai2/Ppp1r9a/Smo/Dgki/Crhr2/Snca/Grm7/Kcna1/Slc8a2/Kcnc3/Kcnc1/Cyfip1/Gabrb3/Cpeb1/Prrt2/Cngb1/Cdh8/Cdh1/Calb2/Disc1/Pard3/Nrp1/Grik4/Nectin1/Drd2/Snap91/Elk1/Fgf13/Arhgap4/Dlg3/Dcx | 81 |
| CC | GO:0042734 | presynaptic membrane | 55/1933 | 238/17474 | 7,51E+07 | 5,26E+10 | 4,80E+10 | Epha4/Atp2b4/Cacna1e/Kcnh1/Stxbp5/Grm1/Grik2/Ppfia2/Cyth1/Psen1/Rgs7bp/Erc2/Cacna1d/Pcdh17/Kctd12/Cdh9/Cntn1/Itsn1/Kcnj6/Cacna1h/Nrxn1/Cdh2/Kctd16/Syt7/Stx3/Ntng2/Kcna4/Kcnc4/Gabbr2/Grik3/Marcksl1/Ephb2/Cacna2d1/Gabrb1/Stx2/Dgki/Grm7/Cacna1c/Ano2/Kcna1/Napa/Kcnc3/Kcnc1/Prrt2/Glra3/Npy1r/Cntn5/Grik4/Nectin1/Drd2/Snap91/Grm2/Ctnnb1/Gpc4/Atp2b3 | 55 |
| CC | GO:0034702 | ion channel complex | 61/1933 | 295/17474 | 9,55E+08 | 4,09E+11 | 3,73E+11 | Unc80/Cacna1e/Kcnh1/Stxbp5/Grik2/Mcu/Best3/Cacng5/Kcnj2/Hspa2/Ryr2/Nrn1/Cacna1d/Ptk2b/Kcns2/Kcnv1/Cacng2/Ano6/Shisa9/Cacna1h/Kcng3/Kcng2/Kcnip2/Olfm1/Scn3a/Scn9a/Scn7a/Chrna1/Sestd1/Lrrc55/Kcna4/Kcng1/Kcnd3/Kcnc4/Amigo1/Grik3/Gabrd/Akap9/Cacna2d1/Gabra4/Gabrb1/Lrrtm4/Cacna1c/Scnn1a/Ano2/Kcna1/Ryr1/Catsperg1/Kcnc3/Abcc8/Kcnc1/Gabrb3/Glra3/Cngb1/Pkd1l3/Kcnk1/Trpc6/Scn3b/Grik4/Dlg3/Glra2 | 61 |
| CC | GO:0044306 | neuron projection terminus | 48/1933 | 222/17474 | 3,71E+10 | 0.0013001953949089277 | 0.001187754082820252 | Epha4/Grik2/Hap1/Amph/Scgn/Cplx2/Erc2/Ptger4/Septin5/Itsn1/Syt7/Slc18a2/Dbh/Tanc1/Slc4a10/Bdnf/Pdyn/Flrt3/Mme/Kcnc4/Calb1/Grik3/Maco1/Hnrnpr/Sri/Pclo/Cad/Btbd8/Rimbp2/Dgki/Crhr2/Snca/Grm7/Kcna1/Slc8a2/Kcnc3/Kcnc1/Cyfip1/Gabrb3/Prrt2/Cngb1/Cdh8/Cdh1/Calb2/Grik4/Drd2/Snap91/Elk1 | 48 |
| CC | GO:0043679 | axon terminus | 44/1933 | 200/17474 | 5,80E+09 | 0.0017179499241578957 | 0.0015693810672527495 | Epha4/Grik2/Hap1/Amph/Scgn/Cplx2/Erc2/Septin5/Itsn1/Syt7/Slc18a2/Dbh/Tanc1/Slc4a10/Bdnf/Pdyn/Flrt3/Kcnc4/Calb1/Grik3/Hnrnpr/Sri/Pclo/Cad/Rimbp2/Dgki/Crhr2/Snca/Grm7/Kcna1/Slc8a2/Kcnc3/Kcnc1/Cyfip1/Gabrb3/Prrt2/Cngb1/Cdh8/Cdh1/Calb2/Grik4/Drd2/Snap91/Elk1 | 44 |
| CC | GO:0030427 | site of polarized growth | 45/1933 | 220/17474 | 3,22E+10 | 0.006207663828538913 | 0.0056708230824330415 | Als2/Epha4/Cxcr4/Adcy10/Sirt1/Hap1/Psen1/Dicer1/Map1b/Erc2/Ptk2b/Pcdh9/Slc2a13/Rtn4r/Tiam1/Dscam/Dynlt1a/Nrxn1/Dcc/Stx3/Olfm1/Lamp5/Flrt3/Src/Ptbp2/Unc5c/C9orf72/Abitram/Hnrnpr/Pclo/Auts2/Orai2/Ppp1r9a/Smo/Snca/Cyfip1/Cpeb1/Disc1/Pard3/Nrp1/Nectin1/Fgf13/Arhgap4/Dlg3/Dcx | 45 |
| BP | GO:0007416 | synapse assembly | 52/1933 | 213/17474 | 2,51E+07 | 2,43E+10 | 2,22E+10 | C1ql2/Nptx1/Lrrn3/Mef2c/Dlg5/Ptk2b/Pcdh17/Mycbp2/Cdh9/Il1rap/Thbs2/Nrxn1/Cdh2/Nrg2/Pcdhb16/C1ql3/Ntng2/Bdnf/Lzts3/Flrt3/Ghsr/Slitrk3/Amigo1/Negr1/Sdcbp/Epha7/Lingo2/Ephb2/Pclo/Ppp1r9a/Lrrc4/Snca/Lrrtm4/Add2/Lrtm2/Chd4/Gabrb3/Ntrk3/Eef2k/Cdh1/Cntn5/Kirrel3/Nectin1/Clstn2/Ephb1/Ctnnb1/Gpc4/Fgf13/Slitrk2/Il1rapl1/Ogt/Bhlhb9 | 52 |

| | | | | | | | | | |
|---|---|---|---|---|---|---|---|---|---|
| BP | GO:0007409 | axonogenesis | 94/1933 | 482/17474 | 2,52E+07 | 2,43E+10 | 2,22E+10 | Als2/Nrp2/Epha4/Cxcr4/Vangl2/Tgfb2/Stxbp5/Ank3/Adarb1/Arhgef25/Adcy1/B3gnt2/Fstl4/Cntnap1/Nptx1/Crppa/Psen1/Nrn1/Unc5a/Mef2c/Map1b/Mycbp2/Sema5a/Wnt7b/Cntn1/Rtn4r/Sema5b/Epha3/Robo2/Tiam1/Dscam/Tiam2/Cdh2/B4galt6/Wdr36/Dcc/Smad4/Myo5b/Prkg1/Lgi1/Slit1/Ablim1/Olfm1/Ntng2/Tbr1/Chn1/Bdnf/Flrt3/B4galt5/Ythdf1/Bhlhe22/Slitrk3/Amigo1/Plppr4/Unc5c/Usp33/Epha7/C9orf72/Nfib/Ephb2/Draxin/Slit2/Prdm8/Auts2/Smo/Plxna4/Sema4f/Atg7/Cxcl12/Lrtm2/Ntf3/Ptpro/Cyfip1/Ntrk3/Unc5d/Cdh1/Ist1/Disc1/Nrp1/Cntn5/Aplp2/Robo3/Nectin1/Drd2/Islr2/Map2k1/Prtg/Snap91/Ephb1/Fgf13/Slitrk2/Arhgap4/Plxna3/Dcx | 94 |
| BP | GO:0050807 | regulation of synapse organization | 66/1933 | 302/17474 | 3,95E+07 | 3,38E+11 | 3,09E+11 | Nrp2/Epha4/C1ql2/Igsf9/Ppfia2/Camk2b/Neurod2/Cntnap1/Lrrn3/Dact1/Psen1/Rps6ka5/Mef2c/Dlg5/Ptk2b/Pcdh8/Mycbp2/Il1rap/Tiam1/Itsn1/Afdn/Thbs2/Mapk14/Lrfn2/Nrxn1/Cdh2/Nrg2/Myo5b/Neurl1a/C1ql3/Ntng2/Tanc1/Bdnf/Lzts3/Flrt3/Ghsr/Slitrk3/Amigo1/Negr1/Epha7/Lingo2/Lrp8/Ephb2/Ppp1r9a/Snca/Lrrtm4/Lrtm2/Chd4/Ptpro/Cyfip1/Ntrk3/Eef2k/Cdh8/Disc1/Nectin1/Drd2/Nedd4/Snap91/Clstn2/Ephb1/Ctnnb1/Gpc4/Slitrk2/Il1rapl1/Ogt/Bhlhb9 | 66 |
| BP | GO:0050803 | regulation of synapse structure or activity | 67/1933 | 310/17474 | 4,91E+08 | 3,78E+10 | 3,46E+10 | Nrp2/Epha4/C1ql2/Igsf9/Ppfia2/Camk2b/Neurod2/Cntnap1/Lrrn3/Dact1/Psen1/Rps6ka5/Mef2c/Dlg5/Ptk2b/Pcdh8/Mycbp2/Sybu/Il1rap/Tiam1/Itsn1/Afdn/Thbs2/Mapk14/Lrfn2/Nrxn1/Cdh2/Nrg2/Myo5b/Neurl1a/C1ql3/Ntng2/Tanc1/Bdnf/Lzts3/Flrt3/Ghsr/Slitrk3/Amigo1/Negr1/Epha7/Lingo2/Lrp8/Ephb2/Ppp1r9a/Snca/Lrrtm4/Lrtm2/Chd4/Ptpro/Cyfip1/Ntrk3/Eef2k/Cdh8/Disc1/Nectin1/Drd2/Nedd4/Snap91/Clstn2/Ephb1/Ctnnb1/Gpc4/Slitrk2/Il1rapl1/Ogt/Bhlhb9 | 67 |
| BP | GO:0016570 | histone modification | 79/1933 | 409/17474 | 4,90E+08 | 2,90E+12 | 2,65E+12 | Rbbp5/Smyd3/Smyd2/Rcor3/Hdac2/Ddx21/Sirt1/Jmjd1c/Dot1l/Usp15/Usp22/Ncor1/Suz12/Mbtd1/Ezh1/Cbx8/Asxl2/Twist1/Prkd1/Mbip/Mideas/Otub2/Rcor1/Dek/Kat6b/Nipbl/Phf20l1/Hdac7/Naa60/Crebbp/Naa50/Mta3/Epc1/Smad4/Naa40/Rcor2/Mta2/Pcgf6/Taf5/Mllt10/Bmi1/Ehmt1/Tbr1/Mettl8/Gatad2b/Zzz3/Mysm1/Kdm4a/Nfyc/Rlf/Sfpq/Hdac1/Kmt2c/Sart3/Auts2/Baz1b/Trrap/Snca/Smarcad1/Brpf1/Atg7/Kdm5a/Chd4/Prmt8/Pagr1a/Igf2/Sin3b/Smarca5/Sf3b3/Pml/Msl2/Wdr82/Setd2/Eomes/Ctnnb1/Kdm6a/Jade3/Ogt/Usp51 | 79 |
| BP | GO:0003007 | heart morphogenesis | 58/1933 | 273/17474 | 6,97E+07 | 3,54E+11 | 3,23E+11 | Eya1/Nrp2/Vangl2/Tgfb2/Prox1/Cited2/Mdm2/Dvl2/Axin2/Asxl2/Sox11/Twist1/Psen1/Dicer1/Ryr2/Dsp/Mef2c/Pbrm1/Tnnc1/Nipbl/Zfpm2/Ly6e/Arid2/Mrtfb/Robo2/Adamts1/Mib1/Smad4/Rbm20/Pdcd4/Ttn/Jag1/Spry1/Fat4/Npy2r/Notch2/Rbm15/Chd7/Tgfbr1/Jun/Slit2/Pds5a/Tgfbr3/Wnt16/Smo/Lrp6/Ryr1/Parva/Tead1/Npy5r/Npy1r/Nrp1/Epor/Smad3/Tpm1/Nedd4/Ctnnb1/Kdm6a | 58 |
| BP | GO:0010959 | regulation of metal ion transport | 81/1933 | 445/17474 | 4,06E+10 | 0.001358469884536286 | 0.0012409889760141397 | Gpr35/Cxcr4/Atp2b4/Ptgs2/Rgs7/Tgfb2/Akap7/Nkain2/Ank3/Pawr/Gck/Stc2/P2rx5/Inpp5k/Hap1/Wnk4/Kcnj2/Osr1/Prkd1/Hspa2/Ryr2/Actn2/Hecw1/Cacna1d/Ptk2b/Stc1/Fgf14/Kcns2/Cntn1/Ano6/Glp1r/Ehd3/Slc30a6/Prkce/Diaph1/Nedd4l/Neto1/Oga/Kcnip2/Sestd1/Lrrc55/Capn3/Kcng1/Atp1a1/Ptpn22/Gstm7/Amigo1/Chd7/Asph/Nkain3/Tmem38b/Ptpn3/Epb41/Akap9/Sri/Cacna2d1/Ywhah/Stim2/Crhr2/Adcyap1r1/Snca/Cxcl12/Cacna1c/Kcna1/Cracr2a/Abcc8/Kcnc1/Il16/Cemip/Homer3/Tmem38a/Plcg2/Trpc6/Scn3b/Ubash3b/Usp2/Drd2/Pml/Nedd4/Ctnnb1/Fgf13 | 81 |
| BP | GO:2001020 | regulation of response to DNA damage stimulus | 61/1933 | 311/17474 | 5,79E+09 | 0.0017179499241578957 | 0.0015693810672527495 | Eya1/Ackr3/Steap3/Smyd2/Bclaf1/Sirt1/Mdm2/Mrnip/Usp22/Atad5/Mbtd1/Smarcd2/Ddx5/Axin2/Recql5/Cbx8/Twist1/Dpf3/Otub2/Dek/Prkcd/Pbrm1/Shld2/Nkx3-1/Arid2/Ercc4/Arid1b/Smchd1/Epc1/Smarca2/Mms19/Slf2/Taf5/Marchf7/Cd44/Pogz/Trim32/Trp73/Chek2/Bcl7a/Kmt5a/Baz1b/Trrap/Hmgb1/Cxcl12/Rad52/Bid/Trim28/Dpf1/Rnf169/Usp47/Pagr1a/Tex15/Smarca5/Sf3b3/Rfwd3/Atm/Pml/Khdc3/Setd2/Usp51 | 61 |
| BP | GO:0048762 | mesenchymal cell differentiation | 50/1933 | 243/17474 | 1,02E+11 | 0.0028813496391378248 | 0.0026321696041373297 | Rdh10/Nrp2/Epha4/Vangl2/Tgfb2/Cited2/Hdac2/Kitl/Pawr/Anxa6/Rflnb/Ddx5/Axin2/Zfp750/Osr1/Sox11/Twist1/Dicer1/Mef2c/Dlg5/Il17rd/Tasor/Sema5a/Vasn/Mapk1/Sema5b/Epha3/Tiam1/Bambi/Cdh2/Smad4/Pdcd4/Frzb/Jag1/Fam83d/Spry1/Sdcbp/Tgfbr1/Tgfbr3/Wnt16/Smo/Sema4f/Kbtbd8/Lrp6/Trim28/Mapk3/Nrp1/Smad3/Eomes/Ctnnb1 | 50 |
| BP | GO:0051480 | regulation of cytosolic calcium ion concentration | 21/1933 | 69/17474 | 1,05E+11 | 0.0028813496391378248 | 0.0026321696041373297 | Bok/Atp2b4/Kcnh1/Tunar/Ryr2/Scgn/Erc2/Tmem178/Asph/Calb1/Marcksl1/Hcrtr1/Erc1/Slc8a2/Ryr1/Npy1r/Cngb1/Calb2/Trpc6/Cul5/Atp2b3 | 21 |
| BP | GO:0021537 | telencephalon development | 52/1933 | 258/17474 | 1,26E+11 | 0.0030533982477100267 | 0.00278933939421299 | Cxcr4/Btg2/Zbtb18/Prox1/Hdac2/Lamb1/Psen1/Dicer1/Wdr37/Secisbp2/Dnah5/Rtn4r/Robo2/Afdn/Cdh2/Slit1/Emx2/Tbr1/Neurod1/Celf1/Btbd3/Bhlhe22/Fat4/Chd7/Nfib/Lrp8/Hdac1/Ephb2/Draxin/Trp73/Slit2/Prdm8/Smo/Plxna4/Emx1/Atg7/Cxcl12/Kcna1/Lrp6/Disc1/Kirrel3/Cdon/Drd2/Cul5/Dmxl2/Zic1/Tmem108/Eomes/Ctnnb1/Fgf13/Plxna3/Dcx | 52 |
| BP | GO:0055074 | calcium ion homeostasis | 63/1933 | 335/17474 | 1,65E+10 | 0.0037443696789343076 | 0.0034205553958711705 | Trpa1/Bok/Atp2b4/Tgfb2/Kcnh1/Grik2/Mcu/Stc2/Anxa6/Hap1/Wnk4/Ccdc47/Prkd1/Psen1/Slc24a4/Tunar/Ryr2/Scgn/Erc2/Ptk2b/Stc1/Ank/Grina/Gp1bb/Glp1r/Tmem178/Prkce/Diaph1/Capn3/Gstm7/Sypl2/Chd7/Asph/Calb1/Tmem38b/Marcksl1/Hcrtr1/Camk2n1/Sri/Stim2/Gpr12/Snca/Cacna1c/Erc1/Slc8a2/Ryr1/Cemip/Npy1r/Tmem38a/Inpp4b/Cngb1/Calb2/Plcg2/Disc1/Trpc6/Ubash3b/Drd2/Cul5/Pml/Ibtk/Nt5e/Pik3cb/Atp2b3 | 63 |

| | | | | | | | | | |
|---|---|---|---|---|---|---|---|---|---|
| BP | GO:0060560 | developmental growth involved in morphogenesis | 55/1933 | 284/17474 | 2,38E+10 | 0.004891015616327788 | 0.004468039027193951 | Rdh10/Nrp2/Cxcr4/Vangl2/Kif26b/Fstl4/Nrn1/Map1b/Cpne6/Sema5a/Wnt7b/Rtn4r/Sema5b/Tiam1/Dscam/Afdn/Ptk7/Wdr36/Nedd4l/Myo5b/Prkg1/Slit1/Olfm1/Bdnf/Flrt3/Spry1/Csf1/Epha7/C9orf72/Tnc/Draxin/Slit2/Auts2/Smo/Plxna4/Sema4f/Emx1/Cpne9/Cxcl12/Lrp6/Trim28/Cyfip1/Ntrk3/Cdh1/Ist1/Disc1/Nrp1/Islr2/Ppp2r3a/Tmem108/Ctnnb1/Fgf13/Arhgap4/Plxna3/Dcx | 55 |
| BP | GO:0042391 | regulation of membrane potential | 82/1933 | 480/17474 | 3,95E+10 | 0.007416163357680954 | 0.006774811187178909 | Trpa1/Gpr35/Bok/Myoc/Rgs7/Kcnh1/Akap7/Bves/Grik2/Ank3/Pawr/Cntnap1/Kcnj2/Psen1/Kcnk10/Slc24a4/Ryr2/Actn2/Dsp/Mef2c/Tmem161b/Rgs7bp/Cacna1d/Ptk2b/Fgf14/Nup155/Cacng2/Ano6/Kcnj6/Afdn/Cacna1h/Slc29a1/Sh3gl1/Ehd3/Prkce/Nrxn1/Dsg2/Nedd4l/Neto1/Oga/Kcnip2/Scn3a/Scn9a/Scn7a/Chrna1/Kcna4/Bdnf/Src/Npy2r/Atp1a1/Kcnd3/Kcnc4/Ptpn3/Jun/Grik3/Gabrd/Akap9/Pclo/Cacna2d1/Ywhah/Gabra4/Gabrb1/Slc4a4/Rimbp2/Ppp1r9a/Dgki/Snca/Arl6ip5/Cacna1c/Bid/Kcna1/Slc8a2/Gabrb3/Ntrk3/Glra3/Cngb1/Kcnk1/Scn3b/Nedd4/Tmem108/Fgf13/Glra2 | 82 |
| BP | GO:0034329 | cell junction assembly | 78/1933 | 453/17474 | 4,60E+10 | 0.008054820541735536 | 0.007358237094433755 | C1ql2/Myoc/Ptprk/Cntnap1/Nptx1/Lrrn3/Actn1/Actn2/Mef2c/Ocln/Marveld2/Dlg5/Ptk2b/Pcdh17/Mycbp2/Cdh9/Sh3bp1/Hdac7/Il1rap/Epha3/Afdn/Thbs2/Nrxn1/Mpp7/Cdh2/Nrg2/Pcdhb16/Nedd4l/C1ql3/Ntng2/Ptprj/Bdnf/Lzts3/Flrt3/Src/Ghsr/Slitrk3/Pip5k1a/Amigo1/Negr1/Sdcbp/Epha7/Lingo2/Mpdz/Ephb2/Pclo/Limch1/Ppp1r9a/Lrrc4/Snca/Lrrtm4/Add2/Lrtm2/Chd4/Ptpro/Gabrb3/Ntrk3/Eef2k/Cldn22/Cdh8/Cdh1/Cdh15/Pard3/Nrp1/Cntn5/Kirrel3/Nectin1/Smad3/Clstn2/Ephb1/Ctnnb1/Gpc4/Fgf13/Slitrk2/Il1rapl1/Ogt/Bhlhb9/Cldn2 | 78 |
| BP | GO:0021954 | central nervous system neuron development | 25/1933 | 99/17474 | 5,59E+10 | 0.009368433605987578 | 0.008558248482276482 | Nrp2/Epha4/Btg2/Adarb1/Secisbp2/Mycbp2/B4galt6/Dcc/Slc4a10/B4galt5/Bhlhe22/Nhlh2/Nfib/Ephb2/Draxin/Slit2/Prdm8/Plxna4/Atg7/Disc1/Nrp1/Drd2/Ephb1/Plxna3/Dcx | 25 |
| MF | GO:0046873 | metal ion transmembrane transporter activity | 76/1933 | 422/17474 | 1,18E+11 | 0.0030533982477100267 | 0.00278933939421299 | Trpa1/Cnnm4/Atp2b4/Kcnt2/Cacna1e/Kcnh1/Grik2/Mcu/Gm5134/Slc6a15/Anxa6/Cacng5/Kcnj2/Psen1/Kcnk10/Slc24a4/Ryr2/Slc12a7/Slc4a7/Cacna1d/Slc39a2/Slc39a14/Kcns2/Kcnv1/Cacng2/Kcnj4/Kcnj6/Cacna1h/Slc30a6/Kcng3/Slc39a6/Slc23a1/Kcng2/Kcnip2/Cnnm2/Slc18a2/Slc4a10/Scn3a/Scn9a/Scn7a/Lrrc55/Kcna4/Trpm7/Kcng1/Atp1a1/Kcnd3/Kcnc4/Slc30a7/Tmem38b/Grik3/Cacna2d1/Slc30a3/Stim2/Slc4a4/Rimbp2/Orai2/Slc13a4/Grm7/Cacna1c/Scnn1a/Kcna1/Slc8a2/Ryr1/Kcnc3/Abcc8/Kcnc1/Zdhhc13/Tmem38a/Slc9a5/Pkd1l3/Kcnk1/Trpc6/Scn3b/Grik4/Slc38a5/Atp2b3 | 76 |
| MF | GO:0005261 | monoatomic cation channel activity | 59/1933 | 305/17474 | 1,27E+11 | 0.0030533982477100267 | 0.00278933939421299 | Trpa1/Unc80/Kcnt2/Cacna1e/Kcnh1/Grik2/Mcu/Anxa6/P2rx5/Cacng5/Kcnj2/Psen1/Tmem63c/Kcnk10/Slc24a4/Ryr2/Cacna1d/Kcns2/Kcnv1/Cacng2/Kcnj4/Atp6v1a/Kcnj6/Cacna1h/Kcng3/Kcng2/Trpm3/Kcnip2/Scn3a/Scn9a/Scn7a/Chrna1/Lrrc55/Kcna4/Trpm7/Kcng1/Kcnd3/Kcnc4/Tmem38b/Grik3/Cacna2d1/Stim2/Rimbp2/Orai2/Grm7/Cacna1c/Scnn1a/Kcna1/Ryr1/Kcnc3/Abcc8/Kcnc1/Tmem38a/Cngb1/Pkd1l3/Kcnk1/Trpc6/Scn3b/Grik4 | 59 |
| MF | GO:0042393 | histone binding | 46/1933 | 224/17474 | 2,39E+11 | 0.004891015616327788 | 0.004468039027193951 | Rbbp5/Sirt1/Usp15/Pwwp2a/Zzef1/Suz12/Mbtd1/Ezh1/Bptf/Cbx2/Cbx8/Atad2b/Vrk1/Dek/Kat6b/Tdrd3/Atad2/Cbx6/Cbx5/Morc3/Smarca2/Uhrf2/Sfmbt2/Mllt10/Ssrp1/Zzz3/Chd7/Mysm1/Nasp/Kdm4a/Kmt2c/Sart3/Sbno1/Baz1b/Kdm7a/Snca/Kdm5a/Chd4/Spty2d1/Rsf1/Smarca5/Hinfp/Anp32a/Phip/Atrx/Usp51 | 46 |

**GO_CA1_protein**

| ONTOLOGY | ID | Description | GeneRatio | BgRatio | pvalue | p.adjust | qvalue | geneID | Count |
|---|---|---|---|---|---|---|---|---|---|
| BP | GO:0050808 | synapse organization | 55/442 | 459/8933 | 560326 | 250746016,9 | 22678469508 | Adgrl2/Shank3/Cdh6/Igsf9b/Dgkz/Flrt1/Ephb3/Efnb2/Lzts1/Zdhhc2/Sez6/Tpbg/Dab2ip/Adgrl3/Itpka/Igsf21/Slc7a11/Mdga2/Rims4/Ngef/Gabrb2/Pik3r1/Arc/Arhgap33/Cacna2d2/Sdk2/Gria1/L1cam/Camk1/Homer1/Cdkl5/Sema4a/Grm5/Cntnap2/Adgrb3/Shank1/Myh10/Gabrg2/Ntrk2/Magi2/Farp1/Nfasc/Dip2a/Grip2/Adgrl1/Lrrn1/Septin7/Pak1/Taok2/Ppfia3/Rock2/Nrxn1/Cttnbp2/Nf1/Tsc1 | 55 |
| BP | GO:0061640 | cytoskeleton-dependent cytokinesis | 19/442 | 75/8933 | 2,3E+07 | 5226626728 | 4727169602 | Cit/Anln/Septin9/Trim36/Exoc6b/Septin4/Myh10/Septin6/Exoc5/Exoc4/Exoc3/Exoc7/Exoc8/Exoc1/Septin3/Septin7/Exoc2/Rock2/Pdcd6ip | 19 |
| BP | GO:0050804 | modulation of chemical synaptic transmission | 52/442 | 448/8933 | 5326365 | 6438853968 | 5823556249 | Shank3/Chrna4/Dgkz/Sorcs3/Lzts1/Zdhhc2/Asic1/Itpka/Kit/Htr2a/Prkcb/Hcn1/Bcr/Slc7a11/Adcy8/Rims4/Arc/Rgs14/Prkcg/Plcl2/Rin1/Cacna2d2/Pnkd/Gria1/Nr3c1/Syt12/L1cam/Cdh11/Neto2/Gnai1/Homer1/Cdkl5/Grm5/Cntnap2/Shank1/Slc4a8/Gria3/Slc4a10/Ntrk2/Dlgap4/Cyp46a1/Ptpn5/Grip2/Pak1/Braf/Ppfia3/Git1/Kif5b/Nrxn1/Nf1/Ncam1/Iqsec1 | 52 |
| BP | GO:0099177 | regulation of trans-synaptic signaling | 52/442 | 449/8933 | 5,8E+07 | 6438853968 | 5823556249 | Shank3/Chrna4/Dgkz/Sorcs3/Lzts1/Zdhhc2/Asic1/Itpka/Kit/Htr2a/Prkcb/Hcn1/Bcr/Slc7a11/Adcy8/Rims4/Arc/Rgs14/Prkcg/Plcl2/Rin1/Cacna2d2/Pnkd/Gria1/Nr3c1/Syt12/L1cam/Cdh11/Neto2/Gnai1/Homer1/Cdkl5/Grm5/Cntnap2/Shank1/Slc4a8/Gria3/Slc4a10/Ntrk2/Dlgap4/Cyp46a1/Ptpn5/Grip2/Pak1/Braf/Ppfia3/Git1/Kif5b/Nrxn1/Nf1/Ncam1/Iqsec1 | 52 |
| BP | GO:0016358 | dendrite development | 35/442 | 249/8933 | 1733630 | 1,10828E+11 | 1,00238E+11 | Shank3/Flrt1/Ephb3/Cit/Lzts1/Sez6/Tpbg/Dab2ip/Itpka/Alk/Ngef/Arc/Matn2/Arhgap33/Phactr1/Nr3c1/Camk1/Sh3glb1/Trpc5/Hecw2/Cdkl5/Dpysl5/Cntnap2/Adgrb3/Shank1/Ntrk2/Farp1/Dip2a/Slc12a5/Septin7/Pak1/Taok2/Git1/Rock2/Iqsec1 | 35 |
| BP | GO:0006887 | exocytosis | 36/442 | 274/8933 | 6,4E+07 | 23076286025 | 2,08711E+11 | Pex5l/Crhbp/Cadps2/Kit/Syt2/Prkcb/Cdk5r2/Bcr/Rims4/Doc2a/Prkcg/Rab27b/Snx19/Syt12/Exoc6b/Septin4/Vsnl1/Myh10/Slc4a8/Exoc5/Exoc4/Exoc3/Exoc7/Ralb/Exoc8/Exoc1/Rabgef1/Pak1/Exoc2/Rala/Braf/Ppfia3/Git1/Pdcd6ip/Ncam1/Trappc11 | 36 |
| BP | GO:0007611 | learning or memory | 31/442 | 221/8933 | 1,2E+09 | 36633576841 | 3,31329E+11 | Shank3/Sorcs3/Asic1/Tpbg/Kit/Htr2a/Pak6/Nr4a2/Slc7a11/Adcy8/Shc3/Arc/Rgs14/Rin1/Gria1/Kcnab1/Grm5/Cntnap2/Adgrb3/Shank1/Ntrk2/Ndrg4/Slc12a5/Pak1/Adcy3/Braf/Git1/Nrxn1/Nf1/Ncam1/Tsc1 | 31 |
| BP | GO:0050807 | regulation of synapse organization | 32/442 | 245/8933 | 4,1E+08 | 9544231363 | 86321833791 | Adgrl2/Shank3/Flrt1/Ephb3/Lzts1/Tpbg/Dab2ip/Adgrl3/Itpka/Slc7a11/Mdga2/Rims4/Ngef/Pik3r1/Arc/Arhgap33/Camk1/Homer1/Cdkl5/Sema4a/Adgrb3/Myh10/Ntrk2/Magi2/Farp1/Adgrl1/Lrrn1/Taok2/Rock2/Nrxn1/Cttnbp2/Nf1 | 32 |
| BP | GO:0007610 | behavior | 49/442 | 476/8933 | 6,6E+08 | 1,39699E+12 | 1,26349E+12 | Shank3/Chrna4/Crhbp/Sorcs3/Homer2/Sez6/Asic1/Tpbg/Adgrl3/Kit/Htr2a/Alk/Thbs4/Pak6/Hcn1/Nr4a2/Slc7a11/Adcy8/Shc3/Arc/Rgs14/Prkcg/Rin1/Gria1/Nr3c1/Homer1/Oxr1/Kcnab1/Grm5/Cntnap2/Adgrb3/Shank1/Gabrg2/Slc4a10/Ntrk2/Ndrg4/Ptpn5/Mapk10/Slc12a5/Inpp5f/Pak1/Adcy3/Braf/Slc16a1/Git1/Nrxn1/Nf1/Ncam1/Tsc1 | 49 |
| BP | GO:0050803 | regulation of synapse structure or activity | 32/442 | 251/8933 | 7E+08 | 1,42882E+11 | 1,29228E+12 | Adgrl2/Shank3/Flrt1/Ephb3/Lzts1/Tpbg/Dab2ip/Adgrl3/Itpka/Slc7a11/Mdga2/Rims4/Ngef/Pik3r1/Arc/Arhgap33/Camk1/Homer1/Cdkl5/Sema4a/Adgrb3/Myh10/Ntrk2/Magi2/Farp1/Adgrl1/Lrrn1/Taok2/Rock2/Nrxn1/Cttnbp2/Nf1 | 32 |
| BP | GO:1903530 | regulation of secretion by cell | 43/442 | 405/8933 | 1,5E+10 | 2,63771E+11 | 2,38565E+12 | Chrna4/Itpr1/Pex5l/Tbc1d1/Crhbp/Asic1/Cadps2/Htr2a/Rbp4/Syt2/Prkcb/Cdk5r2/Bcr/Adcy8/Rims4/Doc2a/Prkcg/Rab27b/Pnkd/Nr3c1/Syt12/Septin4/Vsnl1/Myh10/Slc4a8/Ntrk2/Bsg/Rap1gds1/Ndufaf2/Rgcc/Exoc1/Rabgef1/Exoc2/Rala/Braf/Slc16a1/Git1/Cd200/Kif5b/Maob/Pdcd6ip/Nf1/Myo18a | 43 |
| BP | GO:0048278 | vesicle docking | 13/442 | 54/8933 | 1,5E+09 | 2,65127E+12 | 2,39791E+12 | Exoc6b/Exoc5/Ndrg4/Exoc4/Exoc3/Exoc7/Ralb/Exoc8/Stx7/Exoc1/Exoc2/Ppfia3/Ncam1 | 13 |
| BP | GO:0007416 | synapse assembly | 24/442 | 176/8933 | 5,5E+09 | 8,2664E+11 | 7,47646E+11 | Adgrl2/Shank3/Flrt1/Ephb3/Efnb2/Lzts1/Zdhhc2/Tpbg/Adgrl3/Mdga2/Gabrb2/Pik3r1/Sdk2/Gria1/Sema4a/Cntnap2/Adgrb3/Gabrg2/Ntrk2/Magi2/Farp1/Adgrl1/Lrrn1/Nrxn1 | 24 |
| BP | GO:0090148 | membrane fission | 9/442 | 29/8933 | 6,8E+09 | 9,45952E+11 | 8,55557E+11 | Sh3glb1/Exoc6b/Exoc5/Exoc4/Exoc3/Exoc7/Exoc8/Exoc1/Exoc2 | 9 |
| BP | GO:0099022 | vesicle tethering | 9/442 | 29/8933 | 6,8E+09 | 9,45952E+11 | 8,55557E+11 | Exoc6b/Exoc5/Exoc4/Exoc3/Exoc7/Exoc8/Exoc1/Exoc2/Trappc11 | 9 |
| BP | GO:0052652 | cyclic purine nucleotide metabolic process | 7/442 | 18/8933 | 1,4E+11 | 0.0015711775550187265 | 0.001421035624592062 | Adcy8/Pde1a/Adcy6/Gucy1b1/Pde2a/Adcy2/Adcy3 | 7 |
| BP | GO:0034329 | cell junction assembly | 35/442 | 330/8933 | 1,5E+11 | 0.001628867343475359 | 0.0014732125694002368 | Adgrl2/Shank3/Cdh6/Flrt1/Ephb3/Efnb2/Lzts1/Zdhhc2/Tpbg/Adgrl3/Bcr/Mdga2/Cdh4/Gabrb2/Pik3r1/Sdk2/Gria1/Cdh11/Epb41l3/Sema4a/Cntnap2/Adgrb3/Gabrg2/Ntrk2/Magi2/Farp1/Vcl/Nfasc/Adgrl1/Lrrn1/Taok2/Rock2/Nrxn1/Pdcd6ip/Tsc1 | 35 |
| BP | GO:0009187 | cyclic nucleotide metabolic process | 7/442 | 19/8933 | 2,1E+11 | 0.002111637681881088 | 0.0019098493118101228 | Adcy8/Pde1a/Adcy6/Gucy1b1/Pde2a/Adcy2/Adcy3 | 7 |
| BP | GO:0031346 | positive regulation of cell projection organization | 34/442 | 322/8933 | 2,2E+11 | 0.002164142364063164 | 0.001957336639770154 | Itpr1/Dab2ip/Itpka/Kit/Alk/Serpini1/Anln/Plxna1/Cdh4/Fnbp1l/Septin9/Pik3r1/Tenm3/L1cam/Camk1/Sh3glb1/Trpc5/Dpysl3/Cdkl5/Trpv2/Ntrk2/Ttbk2/Ndrg4/Ptpn5/Magi2/Septin7/Pak1/Katnb1/Rala/Braf/Nrxn1/Dmd/Nf1/Ppp2r5d | 34 |
| BP | GO:0051301 | cell division | 35/442 | 339/8933 | 2,7E+11 | 0.002586107666792838 | 0.002338978883579762 | Cit/Kit/Thbs4/Map4/Anln/Septin9/Rgs14/Trim36/Sh3glb1/Sirt2/Gnai1/Exoc6b/Septin4/Myh10/Septin6/Exoc5/Exoc4/Exoc3/Exoc7/Ralb/Exoc8/Exoc1/Septin3/Septin7/Katnb1/Exoc2/Rala/Babam2/Git1/Rock2/Ccny/Pdcd6ip/Dync1h1/Kif2a/Pik3r4 | 35 |
| BP | GO:0051932 | synaptic transmission, GABAergic | 11/442 | 51/8933 | 3E+10 | 0.0027536289133683334 | 0.0024904917781596574 | Gabra3/Gabra6/Tpbg/Gabrb2/Plcl2/Cntnap2/Gabrg2/Pak1/Kif5b/Nrxn1/Nf1 | 11 |

| | | | | | | | | | |
|---|---|---|---|---|---|---|---|---|---|
| BP | GO:0035418 | protein localization to synapse | 15/442 | 90/8933 | 3,1E+10 | 0.0027536289133683334 | 0.0024904917781596574 | Zdhhc2/Rab27b/Neto2/Homer1/Shank1/Magi2/Kif5a/Grip2/Mapk10/Git1/Kif5c/Kif5b/Nrxn1/Mapk8ip3/Klc1 | 15 |
| BP | GO:0007420 | brain development | 40/442 | 416/8933 | 3,7E+10 | 0.0031914505884320103 | 0.0028864751573116332 | Adgrl2/Shank3/Ephb3/Arnt2/Tyro3/Sez6/Dab2ip/Adgrl3/Alk/Cdk5r2/Chd5/Synj2/Plxna1/Nr4a2/Bcr/Slc7a11/Phactr1/Ccdc85c/Septin4/Cntnap2/Myh10/Dpcd/Slc4a10/Ntrk2/Rtn4rl1/Ttbk2/Atat1/Ndrg4/Nfasc/Shroom2/Pak1/Kif21b/Git1/Mast1/Nrxn1/Mapk8ip3/Nf1/Ncam1/Tsc1/Samd4b | 40 |
| BP | GO:0022406 | membrane docking | 13/442 | 71/8933 | 3,7E+10 | 0.0031914505884320103 | 0.0028864751573116332 | Exoc6b/Exoc5/Ndrg4/Exoc4/Exoc3/Exoc7/Ralb/Exoc8/Stx7/Exoc1/Exoc2/Ppfia3/Ncam1 | 13 |
| BP | GO:0051965 | positive regulation of synapse assembly | 12/442 | 63/8933 | 4,9E+11 | 0.0038168028884933646 | 0.0034520687106749754 | Adgrl2/Flrt1/Ephb3/Tpbg/Adgrl3/Pik3r1/Sema4a/Adgrb3/Ntrk2/Adgrl1/Lrrn1/Nrxn1 | 12 |
| BP | GO:0019932 | second-messenger-mediated signaling | 20/442 | 151/8933 | 5,1E+10 | 0.0038839677806885705 | 0.003512815317082636 | Itpr1/Pex5l/Homer2/Pde11a/Adcy8/Slc8a1/L1cam/Gnai1/Gucy1a2/Kcnc2/Grm5/Gucy1b1/Pde2a/Ntrk2/Ksr1/Adcy2/Exoc4/Adgrl1/Dmd/Ncam1 | 20 |
| BP | GO:0044089 | positive regulation of cellular component biogenesis | 37/442 | 385/8933 | 7,4E+09 | 0.004880585459054741 | 0.004414196081168006 | Adgrl2/Flrt1/Ephb3/Tpbg/Dab2ip/Adgrl3/Kit/Anln/Fnbp1l/Septin9/Pik3r1/Sh3glb1/Dpysl3/Sema4a/Myd88/Cntnap2/Adgrb3/Ntrk2/Ttbk2/Arhgef10/Atat1/Rgcc/Clu/Ralb/Adgrl1/Lrrn1/Septin7/Pak1/Rala/Braf/Tesk1/Git1/Rock2/Nrxn1/Pdcd6ip/Dync1h1/Tsc1 | 37 |
| BP | GO:0060322 | head development | 41/442 | 446/8933 | 8,3E+10 | 0.005274927905664116 | 0.004770855112561842 | Adgrl2/Shank3/Ephb3/Arnt2/Tyro3/Sez6/Dab2ip/Adgrl3/Alk/Cdk5r2/Chd5/Synj2/Plxna1/Nr4a2/Bcr/Slc7a11/Phactr1/Ccdc85c/Septin4/Cntnap2/Myh10/Dpcd/Slc4a10/Ntrk2/Rtn4rl1/Ttbk2/Atat1/Ndrg4/Nfasc/Shroom2/Pak1/Kif21b/Braf/Git1/Mast1/Nrxn1/Mapk8ip3/Nf1/Ncam1/Tsc1/Samd4b | 41 |
| BP | GO:0051962 | positive regulation of nervous system development | 27/442 | 247/8933 | 8,6E+09 | 0.005404079841574018 | 0.004887665272767328 | Adgrl2/Shank3/Flrt1/Ephb3/Tpbg/Adgrl3/Itpka/Kit/Plxna1/Cdh4/Pik3r1/Rgs14/Mag/L1cam/Sh3glb1/Trpc5/Cdkl5/Sema4a/Grm5/Adgrb3/Trpv2/Ntrk2/Adgrl1/Lrrn1/Pak1/Braf/Nrxn1 | 27 |
| BP | GO:0099640 | axo-dendritic protein transport | 5/442 | 11/8933 | 1E+12 | 0.005920494802148353 | 0.005354731552898658 | Rab27b/Kif5a/Kif5c/Kif5b/Mapk8ip3 | 5 |
| BP | GO:1904321 | response to forskolin | 5/442 | 11/8933 | 1E+12 | 0.005920494802148353 | 0.005354731552898658 | Adcy8/Gnai1/Adcy6/Adcy2/Adcy3 | 5 |
| BP | GO:1904322 | cellular response to forskolin | 5/442 | 11/8933 | 1E+12 | 0.005920494802148353 | 0.005354731552898658 | Adcy8/Gnai1/Adcy6/Adcy2/Adcy3 | 5 |
| BP | GO:0030534 | adult behavior | 17/442 | 126/8933 | 1,5E+12 | 0.007956308141903368 | 0.007196002306525952 | Shank3/Chrna4/Crhbp/Homer2/Sez6/Htr2a/Alk/Nr4a2/Slc7a11/Homer1/Oxr1/Cntnap2/Shank1/Gabrg2/Inpp5f/Nrxn1/Tsc1 | 17 |
| BP | GO:0007157 | heterophilic cell-cell adhesion via plasma membrane cell adhesion molecules | 8/442 | 33/8933 | 1,6E+12 | 0.008047133862164435 | 0.007278148709208411 | Igsf21/Cdh4/Tenm3/L1cam/Lgals1/Alcam/Adgrl1/Nrxn1 | 8 |
| BP | GO:0060074 | synapse maturation | 8/442 | 33/8933 | 1,6E+12 | 0.008047133862164435 | 0.007278148709208411 | Sez6/Dab2ip/Igsf21/Adgrb3/Shank1/Adgrl1/Rock2/Nrxn1 | 8 |
| BP | GO:1903909 | regulation of receptor clustering | 6/442 | 18/8933 | 1,6E+12 | 0.008047133862164435 | 0.007278148709208411 | Shank3/Zdhhc2/Slc7a11/Gsn/Cd81/Grip2 | 6 |
| BP | GO:0048709 | oligodendrocyte differentiation | 12/442 | 72/8933 | 1,9E+12 | 0.009049048680777973 | 0.008184320418133799 | Opalin/Trpc4/Mag/Dusp15/Sirt2/Omg/Cntnap2/Ntrk2/Hdac11/Exoc4/Clu/Nf1 | 12 |
| CC | GO:0097060 | synaptic membrane | 49/442 | 419/8933 | 1,2E+08 | 1,03266E+11 | 9339760230 | Adgrl2/Gabra3/Shank3/Chrna4/Igsf9b/Itpr1/Efnb2/Sorcs3/Lzts1/Gabra6/Kctd8/Adgrl3/Cadps2/Htr2a/Igsf21/Hcn1/Adcy8/Rims4/Gabrb2/Arc/Slc8a1/Prkcg/Gria1/L1cam/Neto2/Ank1/Kcnc2/Fxyd6/Grm5/Cntnap2/Pde2a/Adgrb3/Shank1/Slc4a8/Gria3/Gabrg2/Ntrk2/Ptpn5/Magi2/Farp1/Exoc3/Grip2/Adgrl1/Septin3/Septin7/Rala/Nrxn1/Dmd/Ncam1 | 49 |
| CC | GO:0099572 | postsynaptic specialization | 50/442 | 434/8933 | 1,4E+08 | 1,03266E+11 | 9339760230 | Gabra3/Shank3/Chrna4/Igsf9b/Dgkz/Fam81a/Itpr1/Efnb2/Sorcs3/Homer2/Lzts1/Gabra6/Zdhhc2/Map4/Pak6/Bcr/Sh2d5/Adcy8/Gabrb2/Arc/Rgs14/Slc8a1/Prkcg/Kcnd1/Gria1/Nr3c1/Camk1/Neto2/Epb41l3/Homer1/Cdkl5/Myd88/Grm5/Adgrb3/Shank1/Gria3/Gabrg2/Ntrk2/Dlgap4/Magi2/Exoc4/Grip2/Mapk10/Adgrl1/Pak1/Git1/Dmd/Nf1/Tsc1/Iqsec1 | 50 |
| CC | GO:0150034 | distal axon | 40/442 | 314/8933 | 2,7E+08 | 1,36718E+11 | 1,23653E+11 | Crhbp/Tpbg/Prkcb/Hcn1/Cdk5r2/Synj2/Hcn4/Slc8a1/Prkcg/L1cam/Trpc5/Dpysl3/Septin4/Cdkl5/Kcnc2/Tubb3/Myh10/Slc4a8/Gria3/Septin6/Trpv2/Slc4a10/Ntrk2/Kcna6/Exoc4/Exoc7/Kif5a/Clu/Adgrl1/Septin7/Pak1/Katnb1/Taok2/Git1/Kif5c/Kif5b/Nrxn1/Ncam1/Klc1/Tsc1 | 40 |
| CC | GO:0098984 | neuron to neuron synapse | 50/442 | 453/8933 | 5,6E+07 | 2,28688E+11 | 2,06834E+11 | Shank3/Igsf9b/Dgkz/Fam81a/Itpr1/Efnb2/Sorcs3/Homer2/Lzts1/Gabra6/Zdhhc2/Mal2/Map4/Pak6/Bcr/Sh2d5/Adcy8/Arc/Rgs14/Slc8a1/Prkcg/Kcnd1/Gria1/Nr3c1/Syt12/Camk1/Neto2/Epb41l3/Homer1/Cdkl5/Myd88/Grm5/Cntnap2/Adgrb3/Shank1/Slc4a8/Gria3/Ntrk2/Magi2/Exoc4/Grip2/Mapk10/Adgrl1/Pak1/Git1/Nrxn1/Dmd/Nf1/Tsc1/Iqsec1 | 50 |
| CC | GO:0000145 | exocyst | 8/442 | 15/8933 | 1,6E+09 | 44686765510 | 404164924,2 | Exoc6b/Exoc5/Exoc4/Exoc3/Exoc7/Exoc8/Exoc1/Exoc2 | 8 |
| CC | GO:0032589 | neuron projection membrane | 12/442 | 55/8933 | 1,2E+11 | 0.0013833973150138262 | 0.001251199688615857 | Gabra3/Gabra6/Hcn1/Gria1/Ank1/Epb41l3/Kcnc2/Cntnap2/Gabrg2/Slc12a5/Unc5a/Mapk8ip3 | 12 |
| CC | GO:0030427 | site of polarized growth | 24/442 | 186/8933 | 1,4E+09 | 0.0016153624659925709 | 0.00146099821975688 | Cdk5r2/L1cam/Trpc5/Dpysl3/Cdkl5/Tubb3/Myh10/Trpv2/Ntrk2/Exoc4/Exoc7/Kif5a/Clu/Adgrl1/Pak1/Katnb1/Taok2/Git1/Kif5c/Kif5b/Nrxn1/Ncam1/Klc1/Tsc1 | 24 |
| CC | GO:0005875 | microtubule associated complex | 14/442 | 77/8933 | 2,1E+09 | 0.002111637681881088 | 0.0019098493118101228 | Fnta/Klc2/Kif21a/Kif5a/Katnal1/Katnb1/Kif21b/Dync1i1/Kif5c/Kif5b/Fntb/Dync1h1/Klc1/Kif2a | 14 |
| CC | GO:0043198 | dendritic shaft | 12/442 | 62/8933 | 4,2E+11 | 0.0034082965232865533 | 0.003082599266577313 | Lzts1/Sez6/Asic1/Htr2a/Hcn1/Slc8a1/Gria1/Homer1/Grm5/Gria3/Exoc4/Grip2 | 12 |
| CC | GO:0032809 | neuronal cell body membrane | 8/442 | 30/8933 | 7,5E+10 | 0.004880585459054741 | 0.004414196081168006 | Flrt1/Gabra6/Dab2ip/Adcy8/Gria1/Kcnc2/Slc4a8/Unc5a | 8 |
| CC | GO:0005940 | septin ring | 5/442 | 11/8933 | 1E+12 | 0.005920494802148353 | 0.005354731552898658 | Septin9/Septin4/Septin6/Septin3/Septin7 | 5 |
| CC | GO:0031105 | septin complex | 5/442 | 11/8933 | 1E+12 | 0.005920494802148353 | 0.005354731552898658 | Septin9/Septin4/Septin6/Septin3/Septin7 | 5 |
| CC | GO:0044306 | neuron projection terminus | 21/442 | 174/8933 | 1,3E+12 | 0.007351992868603366 | 0.0066494354789250094 | Flrt1/Crhbp/Tpbg/Prkcb/Hcn1/Synj2/Hcn4/Slc8a1/Prkcg/L1cam/Septin4/Kcnc2/Slc4a8/Gria3/Septin6/Slc4a10/Ntrk2/Kcna6/Septin7/Git1/Dmd | 21 |
| CC | GO:0099023 | vesicle tethering complex | 11/442 | 60/8933 | 1,5E+11 | 0.007956308141903368 | 0.007196002306525952 | Exoc6b/Exoc5/Exoc4/Exoc3/Exoc7/Exoc8/Exoc1/Exoc2/Tgfbrap1/Vps33a/Trappc11 | 11 |

| | | | | | | | | | |
|---|---|---|---|---|---|---|---|---|---|
| CC | GO:0035253 | ciliary rootlet | 5/442 | 12/8933 | 1,7E+11 | 0.008362219900307992 | 0.00756312508478312 | Klc2/Kif5a/Kif5c/Kif5b/Klc1 | 5 |
| CC | GO:0043679 | axon terminus | 19/442 | 154/8933 | 2E+12 | 0.009519350221268595 | 0.00860967988257048 | Crhbp/Tpbg/Prkcb/Hcn1/Synj2/Hcn4/Slc8a1/Prkcg/L1cam/Septin4/Kcnc2/Slc4a8/Gria3/Septin6/Slc4a10/Ntrk2/Kcna6/Septin7/Git1 | 19 |
| MF | GO:0009975 | cyclase activity | 6/442 | 14/8933 | 3E+11 | 0.0027536289133683334 | 0.0024904917781596574 | Adcy8/Gucy1a2/Adcy6/Gucy1b1/Adcy2/Adcy3 | 6 |
| MF | GO:0016849 | phosphorus-oxygen lyase activity | 6/442 | 14/8933 | 3E+11 | 0.0027536289133683334 | 0.0024904917781596574 | Adcy8/Gucy1a2/Adcy6/Gucy1b1/Adcy2/Adcy3 | 6 |
| MF | GO:0030695 | GTPase regulator activity | 34/442 | 342/8933 | 7,5E+09 | 0.004880585459054741 | 0.004414196081168006 | Tbc1d1/Arhgap12/Asap2/Dab2ip/Sh3bp4/Bcr/Rasal1/Ngef/Rgs14/Arhgap33/Garnl3/Rin1/Arhgap26/Dennd5b/Arhgap23/Adap1/Adgrb3/Arhgef10/Vps9d1/Rap1gds1/Arhgdia/Farp1/Rabgef1/Psd3/Git1/Tbc1d17/Ralgapa1/Rap1gap/Rabep1/Ralgapb/Dennd4b/Nf1/Ccz1/Iqsec1 | 34 |
| MF | GO:0060589 | nucleoside-triphosphatase regulator activity | 34/442 | 342/8933 | 7,5E+09 | 0.004880585459054741 | 0.004414196081168006 | Tbc1d1/Arhgap12/Asap2/Dab2ip/Sh3bp4/Bcr/Rasal1/Ngef/Rgs14/Arhgap33/Garnl3/Rin1/Arhgap26/Dennd5b/Arhgap23/Adap1/Adgrb3/Arhgef10/Vps9d1/Rap1gds1/Arhgdia/Farp1/Rabgef1/Psd3/Git1/Tbc1d17/Ralgapa1/Rap1gap/Rabep1/Ralgapb/Dennd4b/Nf1/Ccz1/Iqsec1 | 34 |
| MF | GO:0015276 | ligand-gated monoatomic ion channel activity | 13/442 | 78/8933 | 1E+12 | 0.005920494802148353 | 0.005354731552898658 | Gabra3/Chrna4/Itpr1/Pex5l/Kcnh7/Gabra6/Asic1/Hcn1/Gabrb2/Hcn4/Gria1/Gria3/Gabrg2 | 13 |
| MF | GO:0022834 | ligand-gated channel activity | 13/442 | 78/8933 | 1E+12 | 0.005920494802148353 | 0.005354731552898658 | Gabra3/Chrna4/Itpr1/Pex5l/Kcnh7/Gabra6/Asic1/Hcn1/Gabrb2/Hcn4/Gria1/Gria3/Gabrg2 | 13 |
| MF | GO:0031267 | small GTPase binding | 26/442 | 243/8933 | 1,6E+12 | 0.008171822110945661 | 0.007390921732804721 | Pex5l/Itpka/Sh3bp4/Ndrg1/Rims4/Rin1/Gria1/Dennd5b/Cdkl5/Cyrib/Vps9d1/Exoc5/Arhgdia/Farp1/Vcl/Exoc4/Exoc8/Rabgef1/Pak1/Exoc2/Braf/Git1/Rock2/Rap1gap/Strn3/Wdr44 | 26 |
| MF | GO:0043178 | alcohol binding | 11/442 | 62/8933 | 2E+11 | 0.009321835205085326 | 0.008431039426886935 | Astn2/Itpr1/Trpc4/Rbp4/Syt2/Rlbp1/Plcl2/Trpc5/Osbpl10/Cd81/Osbpl6 | 11 |

**GO_CA3_protein**

| ONTOLOGY | ID | Description | GeneRatio | BgRatio | pvalue | p.adjust | qvalue | geneID | Count |
|---|---|---|---|---|---|---|---|---|---|
| CC | GO:0044304 | main axon | 34/680 | 72/8933 | 0,00106 | 0,543708882 | 0,488956258 | Mbp/Myo1d/Hapln2/Mag/Spock1/Gjc2/Ermn/Sirt2/Nfasc/Dlg2/Mapt/Kcnab2/Cntn2/Cntnap2/Scn8a/Crhbp/Kcnq2/Epb41l3/App/Kcnc1/Kcnc3/Kcnc2/Kcna1/Robo1/Tubb4a/Kcnh1/Scn1b/Scn2a/Cntnap1/Bin1/Kif13b/Ank3/Nav1/Mapk8ip3 | 34 |
| CC | GO:0030673 | axolemma | 10/680 | 17/8933 | 7289741 | 47219772426 | 42464642369 | Myo1d/Mapt/Cntnap2/Epb41l3/Kcnc1/Kcnc3/Kcnc2/Robo1/Kcnh1/Mapk8ip3 | 10 |
| CC | GO:0097060 | synaptic membrane | 63/680 | 419/8933 | 9,3E+07 | 53250598862 | 47888151942 | Grik4/Grin1/Cryab/Kcnq5/Ptprd/Neto1/Shisa6/Chrm2/Nectin3/Dlg2/Actn2/Slc6a9/Flrt3/Nt5e/Syndig1/Kcnab2/Lpar1/Iqsec3/Cntn2/Efnb3/Clstn2/Dgkb/Cntnap2/Gabra5/Chrm3/Scn8a/Cacng2/Grik5/Snph/Syt11/Prkcg/Lrrc4b/Kcnc1/Kcnc3/Kcnc2/Lrrtm1/Grm1/Syt3/Kcna1/Slc6a11/Mtmr2/Cacna1c/Kcnj6/Kcnh1/Abhd6/Srgap2/Rtn4/Scn2a/Epha4/Nrxn3/Cnksr2/Clstn3/Cpe/Cacna1a/Cadm3/Ank3/Lrrtm2/Pi4k2a/Ache/Erbb4/Stxbp5/Grm7/Dcc | 63 |
| CC | GO:0098984 | neuron to neuron synapse | 64/680 | 453/8933 | 7,3E+08 | 2,87631E+12 | 2,58666E+11 | Homer3/Grik4/Nefm/Grin1/Elavl2/Cryab/Ina/Bcas1/Nefh/Ptprd/Neto1/C1ql2/Shisa6/Chrm2/Nectin3/Dlg2/Cabp1/Actn2/Mapt/Slc6a9/Flrt3/Pdyn/Syndig1/Kcnab2/Iqsec3/Efnb3/Clstn2/Dgkb/Cntnap2/Rgs14/Chrm3/Scn8a/Cacng2/Grik5/Syt12/Tanc1/Anks1b/Epb41l3/Frmpd4/Tsc2/Syt11/Prkcg/Lrrc4b/Sdcbp/Grm1/Homer2/Mtmr2/Cacna1c/Srgap2/Rtn4/Dst/Epha4/Cnksr2/Clstn3/Vamp7/Ank3/Plxna4/Lrrtm2/Erbb4/Stxbp5/Grm7/Dcc/Mapk10/Rtn3 | 64 |
| CC | GO:0043209 | myelin sheath | 34/680 | 187/8933 | 1,4E+10 | 4,2247E+12 | 3,79927E+11 | Mobp/Mbp/Plp1/Mog/Tspan2/Cnp/Nefm/Myo1d/Cldn11/Mag/Nefl/Cryab/Ina/Gjc2/Pllp/Ermn/Sirt2/Nefh/Nfasc/Gjc3/Serinc5/Jam3/Ndrg1/Cntn2/Septin4/Dlat/Llgl1/Plec/Igsf8/Ehd1/Tubb4a/Rtn4/Cntnap1/Rdx | 34 |
| CC | GO:0034702 | ion channel complex | 32/680 | 183/8933 | 6,8E+09 | 0.001388724695207015 | 0.0012488772075791882 | Slc17a8/Grik4/Grin1/Kcnq5/Shisa6/Clic4/Ttyh2/Lrrc8d/Ryr3/Scn4b/Kcnab2/Kcnk4/Mcub/Cntnap2/Gabra5/Scn8a/Cacng2/Grik5/Kcnq2/Kcnc1/Kcnc3/Kcnc2/Kcna1/Kcnip2/Cpt1c/Cacna1c/Kcnh1/Abhd6/Scn1b/Scn2a/Cacna1a/Stxbp5 | 32 |
| CC | GO:0043679 | axon terminus | 27/680 | 154/8933 | 3,3E+11 | 0.0035107367515124235 | 0.003157198201995593 | Slc17a8/Grik4/Grin1/Kcnq5/Prkca/Chrm2/Synj2/Flrt3/Pdyn/Kcnab2/Chrm3/Septin4/Crhbp/Grik5/Tanc1/Syt11/Prkcg/Kcnc1/Kcnc3/Kcnc2/P2rx7/Kcna1/Epha4/Bin1/Amph/Gad1/Grm7 | 27 |
| CC | GO:0043204 | perikaryon | 20/680 | 104/8933 | 9,3E+10 | 0.007108023038235778 | 0.006392230219595215 | Slc17a8/Grik4/Nefm/Cryab/Gjc2/Sirt2/Nefh/Hpca/Nell2/Septin4/Crhbp/Grik5/Kcnc2/Kcna1/Epha4/Cpe/Cacna1a/Pi4k2a/Rufy3/Mapk10 | 20 |
| CC | GO:1990351 | transporter complex | 38/680 | 270/8933 | 1,5E+12 | 0.009251135897431704 | 0.008319527121822967 | Slc17a8/Grik4/Grin1/Kcnq5/Shisa6/Clic4/Ttyh2/Lrrc8d/Ryr3/Scn4b/Kcnab2/Kcnk4/Mcub/Cntnap2/Atp8a1/Tmem30a/Gabra5/Scn8a/Cacng2/Grik5/Kcnq2/Kcnc1/Kcnc3/Kcnc2/Atp2a2/Kcna1/Kcnip2/Atp11a/Cpt1c/Cacna1c/Kcnh1/Abhd6/Scn1b/Atp1b3/Scn2a/Cacna1a/Stxbp5/Atp8a2 | 38 |
| BP | GO:0050804 | modulation of chemical synaptic transmission | 70/680 | 448/8933 | 3,3E+07 | 4675042756 | 42042561516 | Ptgs2/Homer3/Grik4/Grin1/Nefl/Ina/Nefh/Ptprd/Neto1/Prkca/Shisa6/Chrm2/Npy2r/Kcnj10/Oxtr/Kit/Nptx1/Mapt/Slc6a9/Slc12a2/Cntn2/Efnb3/Clstn2/Dgkb/Cntnap2/Nptxr/Rgs14/Plcl1/Lnpep/Cacng2/Grik5/Syt12/Mctp1/Camk4/Prkcg/Slc38a2/Sv2c/App/Kcnc3/Atp2a2/Igf1r/Lrrtm1/Grm1/P2rx7/Rapgef2/Wnk1/Mtmr2/Kcnh1/Abhd6/Rtn4/Slc24a2/Cux2/Epha4/Nrxn3/Clstn3/Apba2/Stxbp5l/Dgke/Begain/Cacna1a/Cpeb3/Prkce/Zzef1/Lrrtm2/Erbb4/Stau1/Stxbp5/Grm7/Rab3gap1/Dcc | 70 |
| BP | GO:0099177 | regulation of trans-synaptic signaling | 70/680 | 449/8933 | 3641708 | 4675042756 | 42042561516 | Ptgs2/Homer3/Grik4/Grin1/Nefl/Ina/Nefh/Ptprd/Neto1/Prkca/Shisa6/Chrm2/Npy2r/Kcnj10/Oxtr/Kit/Nptx1/Mapt/Slc6a9/Slc12a2/Cntn2/Efnb3/Clstn2/Dgkb/Cntnap2/Nptxr/Rgs14/Plcl1/Lnpep/Cacng2/Grik5/Syt12/Mctp1/Camk4/Prkcg/Slc38a2/Sv2c/App/Kcnc3/Atp2a2/Igf1r/Lrrtm1/Grm1/P2rx7/Rapgef2/Wnk1/Mtmr2/Kcnh1/Abhd6/Rtn4/Slc24a2/Cux2/Epha4/Nrxn3/Clstn3/Apba2/Stxbp5l/Dgke/Begain/Cacna1a/Cpeb3/Prkce/Zzef1/Lrrtm2/Erbb4/Stau1/Stxbp5/Grm7/Rab3gap1/Dcc | 70 |
| BP | GO:0007272 | ensheathment of neurons | 26/680 | 107/8933 | 7356537 | 47219772426 | 42464642369 | Mbp/Plp1/Tspan2/Cldn11/Mag/Bcas1/Pllp/Sirt2/Nfasc/Gjc3/Serinc5/Jam3/Cd9/Kcnj10/Ndrg1/Arhgef10/Fa2h/Cntn2/Cntnap2/Degs1/Gpc1/Epb41l3/Plec/Abca2/Mtmr2/Cntnap1 | 26 |
| BP | GO:0008366 | axon ensheathment | 26/680 | 107/8933 | 7356537 | 47219772426 | 42464642369 | Mbp/Plp1/Tspan2/Cldn11/Mag/Bcas1/Pllp/Sirt2/Nfasc/Gjc3/Serinc5/Jam3/Cd9/Kcnj10/Ndrg1/Arhgef10/Fa2h/Cntn2/Cntnap2/Degs1/Gpc1/Epb41l3/Plec/Abca2/Mtmr2/Cntnap1 | 26 |
| BP | GO:0061564 | axon development | 58/680 | 382/8933 | 2,2E+09 | 1,03547E+11 | 93119294395 | Mbp/Plp1/Tspan2/Cnp/Nefm/Grin1/Mag/Nefl/Tgfb2/Nefh/Nfasc/Prkca/Sema3g/Ust/Nptx1/Mapt/Flrt3/Slit2/Plxna1/Cntn2/Efnb3/Unc5b/Cntnap2/Ttl/Llgl1/Epb41l3/Tsc2/Slit1/Zdhhc17/App/Spart/Stk25/Igf1r/Slitrk4/Islr2/Robo1/Rtn4rl1/Rtn4/Dst/Scn1b/Ddr1/Epha4/Cntnap1/Arhgef25/Inpp5f/Cacna1a/Camsap2/Kif13b/Ank3/Plxna4/Ache/Unc5c/Stxbp5/Grm7/Rufy3/Dcc/Atp8a2/Mapk8ip3 | 58 |
| BP | GO:0019226 | transmission of nerve impulse | 17/680 | 60/8933 | 1,4E+10 | 4,2247E+12 | 3,79927E+11 | Mag/Nfasc/Jam3/Cntnap2/Scn8a/Cacng2/Kcnq2/Plec/P2rx7/Kcna1/Scn1b/Scn2a/Fgf12/Cntnap1/Cacna1a/Ank3/Grm7 | 17 |
| BP | GO:0045806 | negative regulation of endocytosis | 13/680 | 39/8933 | 3,3E+10 | 8,37881E+11 | 7,53505E+11 | Necab2/Atxn2/Mctp1/Tsc2/Snph/Syt11/Sdcbp/Abca2/Lrrtm1/Mtmr2/Ank3/Lrrtm2/Lrsam1 | 13 |

| | | | | | | | | | |
|---|---|---|---|---|---|---|---|---|---|
| BP | GO:1902074 | response to salt | 36/680 | 217/8933 | 6,3E+09 | 0.0013558119189682608 | 0.001219278816894124 | Grin1/Nefl/Nefh/Cpne4/Chrm2/Cpne7/Clic4/Dlg2/Kcnj10/Hpca/Nptx1/Slc12a2/Nt5e/Rgs8/Chrm3/Slc25a13/Crhbp/Syt12/Plec/Syt11/Cpne6/Prkcg/App/Igf1r/P2rx7/Anxa11/Syt3/Homer2/Wnk1/Kcnh1/Entpd6/Alg2/Prkce/Ahcyl1/Ache/Abat | 36 |
| BP | GO:0007416 | synapse assembly | 31/680 | 176/8933 | 8,1E+09 | 0.0015349465618643711 | 0.0013803742257774809 | Grin1/Elavl2/Ptprd/Prkca/C1ql2/Amigo2/Oxtr/Hapln4/Nptx1/Mapt/Flrt3/Syndig1/Efnb3/Clstn2/Cntnap2/Nptxr/Lrrc4b/Sdcbp/App/Slitrk4/Lrrtm1/Mdga1/Srgap2/Cux2/Nrxn3/Clstn3/Cacna1a/Sdk2/Lrrtm2/Ache/Erbb4 | 31 |
| BP | GO:0034329 | cell junction assembly | 48/680 | 330/8933 | 8,6E+09 | 0.0015691691345187046 | 0.0014111505136336917 | Grin1/Elavl2/Cldn11/Peak1/Ptprd/Nfasc/Prkca/C1ql2/Amigo2/Jam3/Enpp2/Pkp2/Cd9/Oxtr/Hapln4/Actn2/Nptx1/Mapt/Dapk3/Flrt3/Syndig1/Efnb3/Clstn2/Cntnap2/Nptxr/Clasp1/Epb41l3/Plec/Lrrc4b/Sdcbp/App/Slitrk4/Lrrtm1/Mdga1/Rapgef2/Srgap2/Dst/Cux2/Nrxn3/Clstn3/Cntnap1/Cacna1a/Sdk2/Lrrtm2/Myh9/Ache/Erbb4/Myo9a | 48 |
| BP | GO:0043269 | regulation of monoatomic ion transport | 52/680 | 372/8933 | 1,1E+11 | 0.0018959350065003557 | 0.0017050103773929297 | Ptgs2/Homer3/Plp1/Grin1/Tgfb2/Gjc2/Kcnq5/Neto1/Prkca/Ptpn3/Clic4/Kcnj10/Cabp1/Il16/Stac2/Hpca/Scn4b/Actn2/Slc6a9/Kcnab2/Kcnk4/Stim2/Scn8a/Gnas/Cacng2/Kcnq2/App/Kcnc1/Kcnc3/Kcnc2/P2rx7/Kcna1/Homer2/Fhl1/Kcnip2/Wnk1/Saraf/Cacna1c/Kcnj6/Kcnh1/Scn1b/Atp1b3/Scn2a/Fgf12/Slmap/Bin1/Cacna1a/Ank3/Prkce/Ppp3cb/Ahcyl1/Akt3 | 52 |
| BP | GO:0034762 | regulation of transmembrane transport | 54/680 | 393/8933 | 1,3E+10 | 0.002057451101420683 | 0.0018502614630108208 | Slc17a8/Plp1/Grin1/Tgfb2/Gjc2/Kcnq5/Neto1/Prkca/Shisa6/Ptpn3/Clic4/Kcnj10/Cabp1/Stac2/Hpca/Scn4b/Actn2/Slc6a9/Kcnab2/Kcnk4/Stim2/Prkcd/Chrm3/Scn8a/Crhbp/Cacng2/Kcnq2/Slc43a2/Stxbp3/App/Kcnc1/Kcnc3/Kcnc2/P2rx7/Kcna1/Fhl1/Kcnip2/Wnk1/Cacna1c/Kcnj6/Kcnh1/Scn1b/Atp1b3/Scn2a/Fgf12/Slmap/Bin1/Cacna1a/Ank3/Prkce/Ppp3cb/Ahcyl1/Erbb4/Acsl6 | 54 |
| BP | GO:0006935 | chemotaxis | 44/680 | 300/8933 | 1,7E+11 | 0.0022930663015370586 | 0.002062149718616938 | Tgfb2/Nfasc/Rhog/Prkca/Jam3/Enpp2/Nbl1/Sema3g/Il16/Kit/Slc12a2/Smoc2/Flrt3/Camk1d/Lpar1/Slit2/Plxna1/Cntn2/Padi2/Prkcd/Efnb3/Prkcq/Unc5b/Gab1/Dock4/Plec/Tsc2/Slit1/App/Mpp1/Wnk1/Robo1/Rtn4rl1/Scn1b/Fer/Epha4/Arhgef25/Elmo2/Ank3/Plxna4/Wasl/Unc5c/Dcc/Mapk8ip3 | 44 |
| BP | GO:0042330 | taxis | 44/680 | 300/8933 | 1,7E+11 | 0.0022930663015370586 | 0.002062149718616938 | Tgfb2/Nfasc/Rhog/Prkca/Jam3/Enpp2/Nbl1/Sema3g/Il16/Kit/Slc12a2/Smoc2/Flrt3/Camk1d/Lpar1/Slit2/Plxna1/Cntn2/Padi2/Prkcd/Efnb3/Prkcq/Unc5b/Gab1/Dock4/Plec/Tsc2/Slit1/App/Mpp1/Wnk1/Robo1/Rtn4rl1/Scn1b/Fer/Epha4/Arhgef25/Elmo2/Ank3/Plxna4/Wasl/Unc5c/Dcc/Mapk8ip3 | 44 |
| BP | GO:0007422 | peripheral nervous system development | 13/680 | 45/8933 | 1,9E+10 | 0.0025184988113920257 | 0.0022648807022142814 | Hapln2/Sirt2/Nefh/Nfasc/Med12/Ndrg1/Arhgef10/Fa2h/Gpc1/Plec/Cntnap1/Plxna4/Aldh3a2 | 13 |
| BP | GO:0048709 | oligodendrocyte differentiation | 17/680 | 72/8933 | 2E+10 | 0.0025949236432485675 | 0.0023336093933134763 | Plp1/Tspan2/Cnp/Mag/Sirt2/Aspa/Med12/Dusp15/Enpp2/Kcnj10/Il33/Fa2h/Cntn2/Cntnap2/Abca2/Daam2/Cntnap1 | 17 |
| BP | GO:0045161 | neuronal ion channel clustering | 6/680 | 10/8933 | 3,1E+10 | 0.0035107367515124235 | 0.003157198201995593 | Nfasc/Dlg2/Cntn2/Cntnap2/Kcnip2/Ank3 | 6 |
| BP | GO:0007610 | behavior | 61/680 | 476/8933 | 3,1E+10 | 0.0035107367515124235 | 0.003157198201995593 | Ptgs2/Cnp/Grin1/Neto1/Prkca/Npy2r/Kcnj10/Oxtr/Kit/Mapt/Gpr37/Cntn2/Adam11/Efnb3/Ldlr/Cntnap2/Atp8a1/Rgs14/Napepld/Gabra5/Scn8a/Sgsh/Crhbp/Tanc1/Astn1/Camk4/Gls/Tsc2/Syt11/Prkcg/App/Abca2/Ttbk1/Lrrtm1/Grm1/Homer2/Agtpbp1/Cacna1c/Slc24a2/Brinp1/Cux2/Scn2a/Epha4/Nrxn3/Fgf12/Apba2/Inpp5f/Gatm/Cacna1a/Abl2/Specc1/Amph/Cpeb3/Prkce/Ppp3cb/Zzef1/Gad1/Grm7/Abat/Mapk10/Atp8a2 | 61 |
| BP | GO:0042391 | regulation of membrane potential | 46/680 | 327/8933 | 3,1E+11 | 0.0035107367515124235 | 0.003157198201995593 | Atpif1/Bok/Grin1/Neto1/Ptpn3/Npy2r/Pkp2/Kcnj10/Scn4b/Actn2/Mapt/Kcnab2/Kcnk4/Kctd7/Cntnap2/Gabra5/Scn8a/Cacng2/Grik5/Kcnq2/App/Kcnc2/Spart/Atp2a2/P2rx7/Kcna1/Fhl1/Kcnip2/Mtmr2/Cacna1c/Kcnj6/Kcnh1/Cux2/Scn1b/Atp1b3/Scn2a/Fgf12/Cntnap1/Slmap/Bin1/Begain/Cacna1a/Ank3/Prkce/Abat/Rab3gap1 | 46 |
| BP | GO:0032288 | myelin assembly | 8/680 | 20/8933 | 6E+09 | 0.00529149917078613 | 0.004758634113101779 | Mag/Nfasc/Cd9/Gpc1/Epb41l3/Abca2/Mtmr2/Cntnap1 | 8 |
| BP | GO:0045055 | regulated exocytosis | 29/680 | 179/8933 | 7,8E+10 | 0.00655585313754534 | 0.005895664985273696 | Rab15/Prkca/Rasgrp1/Kit/Rab26/Crhbp/Grik5/Syt12/Syt11/Prkcg/Stxbp3/Sv2c/Rab31/Atp2a2/P2rx7/Syt3/Cacna1c/Kcnh1/Fer/Cadps/Stxbp5l/Cacna1a/Vamp7/Ppp3cb/Pi4k2a/Myh9/Vps41/Stxbp5/Rab3gap1 | 29 |
| BP | GO:0090659 | walking behavior | 11/680 | 38/8933 | 8,2E+10 | 0.00659011811443215 | 0.005926479407167292 | Kcnj10/Mapt/Cntn2/Efnb3/Cntnap2/Scn8a/Agtpbp1/Cacna1c/Epha4/Cacna1a/Abl2 | 11 |
| BP | GO:0051960 | regulation of nervous system development | 47/680 | 352/8933 | 9,6E+10 | 0.007286981938082962 | 0.006553167583123036 | Mbp/Mag/Nefl/Gjc2/Sirt2/Ptprd/Aspa/Prkca/Amigo2/Enpp2/Sema3g/Oxtr/Kit/Il33/Mapt/Flrt3/Syndig1/Slit2/Plxna1/Efnb3/Ldlr/Clstn2/Rgs14/Tsc2/Slit1/Lrrc4b/Spart/Stk25/Slitrk4/Lrrtm1/Mdga1/Islr2/Rapgef2/Robo1/Mtmr2/Rtn4/Brinp1/Cux2/Epha4/Daam2/Clstn3/Bin1/Vamp7/Plxna4/Lrrtm2/Ache/Rufy3 | 47 |
| BP | GO:0051051 | negative regulation of transport | 43/680 | 313/8933 | 9,9E+10 | 0.007343084187493564 | 0.006603620218425514 | Ptgs2/Cryab/Tgfb2/Prkca/Npy2r/Oxtr/Cabp1/Actn2/Mapt/Slc6a9/Acsl4/Necab2/Erlec1/Atxn2/Ube2j1/Gnas/Crhbp/Mctp1/Slc43a2/Os9/Tsc2/Snph/Syt11/Stxbp3/Sdcbp/Abca2/Lrrtm1/Wnk1/Mtmr2/Kcnj6/Fgf12/Stxbp5l/Pfkl/Bin1/Ank3/Prkce/Madd/Ppp3cb/Lrrtm2/Lrsam1/Grm7/Abat/Atp9a | 43 |
| BP | GO:0010038 | response to metal ion | 30/680 | 191/8933 | 1,1E+12 | 0.007423906913158772 | 0.006676303926208687 | Cpne4/Cpne7/Clic4/Dlg2/Kcnj10/Hpca/Nptx1/Mapt/Slc12a2/Slc25a13/Crhbp/Syt12/Syt11/Cpne6/Tfrc/App/Smpd3/P2rx7/Anxa11/Syt3/Kcna1/Wnk1/Kcnh1/Khk/Entpd6/Alg2/Gatm/Ank3/Ahcyl1/Abat | 30 |
| BP | GO:0009395 | phospholipid catabolic process | 10/680 | 33/8933 | 1,1E+12 | 0.007635475701723366 | 0.006866567294308493 | Enpp2/Pld1/Prkcd/Ldlr/Napepld/Smpd3/Abhd6/Gpcpd1/Gdpd1/Inpp5f | 10 |

| | | | | | | | | | |
|---|---|---|---|---|---|---|---|---|---|
| BP | GO:0045987 | positive regulation of smooth muscle contraction | 6/680 | 12/8933 | 1,2E+12 | 0.00776117579854578 | 0.006979609127908644 | Ptgs2/Npy2r/Oxtr/Kit/Chrm3/Abat | 6 |
| BP | GO:0022010 | central nervous system myelination | 8/680 | 22/8933 | 1,3E+12 | 0.008386726420996771 | 0.007542165491500605 | Plp1/Mag/Kcnj10/Fa2h/Cntn2/Cntnap2/Abca2/Cntnap1 | 8 |
| BP | GO:0050808 | synapse organization | 57/680 | 459/8933 | 1,4E+11 | 0.008577681866496082 | 0.007713891323388164 | Grin1/Elavl2/Nefl/Hmcn2/Ptprd/Nfasc/Prkca/C1ql2/Shisa6/Amigo2/Oxtr/Cabp1/Hapln4/Nptx1/Mapt/Arhgap22/Flrt3/Syndig1/Iqsec3/Cntn2/Efnb3/Clstn2/Dgkb/Cntnap2/Nptxr/Cacng2/Tanc1/Frmpd4/Tsc2/Cpne6/Lrrc4b/Sdcbp/App/Arhgap39/Igf1r/Slitrk4/Lrrtm1/Mdga1/Mtmr2/Srgap2/Cux2/Epha4/Nrxn3/Cnksr2/Clstn3/Cntnap1/Cacna1a/Abl2/Sdk2/Ank3/Plxna4/Wasl/Lrrtm2/Ache/Erbb4/Stau1/Myo9a | 57 |
| BP | GO:0045109 | intermediate filament organization | 7/680 | 17/8933 | 1,4E+12 | 0.008780976250931834 | 0.007896713537198372 | Nefm/Nefl/Ina/Nefh/Pkp2/Mtm1/Plec | 7 |
| BP | GO:0044283 | small molecule biosynthetic process | 45/680 | 339/8933 | 1,6E+12 | 0.009396340515959108 | 0.008450109330910184 | Ptgs2/Plp1/Fa2h/Gamt/Dhcr24/Qdpr/Nt5e/Degs1/Ephx1/Dhcr7/Msmo1/Slc25a13/Stard3/Bcat1/Gpt2/Aldh1a1/Gls/Slc27a2/Hsd17b7/Coq8a/Ilvbl/Pdk2/Acss2/Abca2/Scd2/Pck2/Psat1/Acaca/Coq5/Prkag2/Coq4/Gls2/Pycr2/Gatm/Mri1/Ip6k1/Myh9/Ppip5k1/Coq6/Gad1/Fasn/Abat/Wdtc1/Nsdhl/Acadvl | 45 |
| MF | GO:0030695 | GTPase regulator activity | 49/680 | 342/8933 | 1E+10 | 0.0017957016936695845 | 0.0016148707692571038 | Arhgef17/Akap13/Kndc1/Rasgrp1/Arhgef10/Dock5/Arhgef4/Arhgap22/Arhgef6/Slit2/Iqsec3/Rgs8/Rgs14/Arap2/Chrm3/Dock4/Psd3/Sgsm2/Dennd11/Llgl1/Agap3/Elmo1/Tsc2/Acap3/Arhgap39/Rasa3/Srgap3/Tbc1d8b/Rapgef2/Arhgap23/Srgap2/Tbc1d9b/Sbf1/Arfgef3/Stxbp5l/Arhgap21/Arhgef25/Dock9/Madd/Chm/Tbck/Dennd5a/Thg1l/Tbc1d5/Arhgef11/Stxbp5/Rab3gap1/Myo9a/Srgap1 | 49 |
| MF | GO:0060589 | nucleoside-triphosphatase regulator activity | 49/680 | 342/8933 | 1E+10 | 0.0017957016936695845 | 0.0016148707692571038 | Arhgef17/Akap13/Kndc1/Rasgrp1/Arhgef10/Dock5/Arhgef4/Arhgap22/Arhgef6/Slit2/Iqsec3/Rgs8/Rgs14/Arap2/Chrm3/Dock4/Psd3/Sgsm2/Dennd11/Llgl1/Agap3/Elmo1/Tsc2/Acap3/Arhgap39/Rasa3/Srgap3/Tbc1d8b/Rapgef2/Arhgap23/Srgap2/Tbc1d9b/Sbf1/Arfgef3/Stxbp5l/Arhgap21/Arhgef25/Dock9/Madd/Chm/Tbck/Dennd5a/Thg1l/Tbc1d5/Arhgef11/Stxbp5/Rab3gap1/Myo9a/Srgap1 | 49 |
| MF | GO:0140326 | ATPase-coupled intramembrane lipid transporter activity | 8/680 | 17/8933 | 1,4E+11 | 0.002205334495385673 | 0.0019832526892342435 | Atp8a1/Tmem30a/Abca2/Atp11a/Atp9b/Abca3/Atp9a/Atp8a2 | 8 |
| MF | GO:0022836 | gated channel activity | 30/680 | 173/8933 | 1,5E+11 | 0.0022710788521702586 | 0.0020423764514879155 | Grik4/Grin1/Kcnq5/Tmem63a/Clic4/Kcnj10/Ryr3/Scn4b/Kcnab2/Kcnk4/Gabra5/Scn8a/Cacng2/Grik5/Kcnq2/Kcnc1/Ano10/Kcnc3/Kcnc2/Rasa3/P2rx7/Kcna1/Kcnip2/Cacna1c/Kcnj6/Kcnh1/Scn1b/Scn2a/Cacna1a/Grm7 | 30 |
| MF | GO:0022839 | monoatomic ion gated channel activity | 30/680 | 173/8933 | 1,5E+11 | 0.0022710788521702586 | 0.0020423764514879155 | Grik4/Grin1/Kcnq5/Tmem63a/Clic4/Kcnj10/Ryr3/Scn4b/Kcnab2/Kcnk4/Gabra5/Scn8a/Cacng2/Grik5/Kcnq2/Kcnc1/Ano10/Kcnc3/Kcnc2/Rasa3/P2rx7/Kcna1/Kcnip2/Cacna1c/Kcnj6/Kcnh1/Scn1b/Scn2a/Cacna1a/Grm7 | 30 |
| MF | GO:0005509 | calcium ion binding | 47/680 | 330/8933 | 1,9E+10 | 0.0025184988113920257 | 0.0022648807022142814 | Grin1/Spock1/Hmcn2/Acan/Pls1/Enpp2/Vcan/Rasgrp1/Sulf2/Cabp7/Cabp1/Ryr3/Hpca/Nell2/Actn2/Dgka/Smoc2/Edil3/Slit2/Necab2/Dgkg/Stim2/Padi2/Ldlr/Clstn2/Dgkb/Slc25a13/Syt12/Mctp1/Syt11/Slit1/Atp2a2/Ehd1/Tbc1d8b/Anxa11/Syt3/Capn1/Kcnip2/Tubb4a/Tbc1d9b/Dst/Clstn3/Cadps/Esyt2/Ppp3cb/Zzef1/Grm7 | 47 |
| MF | GO:0005261 | monoatomic cation channel activity | 29/680 | 170/8933 | 2,9E+10 | 0.0035107367515124235 | 0.003157198201995593 | Grik4/Grin1/Kcnq5/Tmem63a/Kcnj10/Ryr3/Scn4b/Kcnab2/Kcnk4/Stim2/Scn8a/Cacng2/Grik5/Kcnq2/Kcnc1/Kcnc3/Kcnc2/Rasa3/P2rx7/Kcna1/Kcnip2/Cacna1c/Kcnj6/Kcnh1/Slc24a2/Scn1b/Scn2a/Cacna1a/Grm7 | 29 |
| MF | GO:0046873 | metal ion transmembrane transporter activity | 38/680 | 262/8933 | 7,9E+10 | 0.006575137648504199 | 0.00591300750555792 | Slc17a8/Grik4/Grin1/Kcnq5/Kcnj10/Ryr3/Scn4b/Slc6a9/Slc12a2/Kcnab2/Kcnk4/Stim2/Cnnm4/Scn8a/Cacng2/Grik5/Kcnq2/Tfrc/Slc38a2/Zdhhc17/Kcnc1/Kcnc3/Kcnc2/Rasa3/Atp2a2/Kcna1/Slc6a11/Kcnip2/Cacna1c/Kcnj6/Kcnh1/Slc24a2/Scn1b/Atp1b3/Scn2a/Cacna1a/Cnnm2/Grm7 | 38 |
| MF | GO:0016301 | kinase activity | 60/680 | 485/8933 | 1E+12 | 0.007343084187493564 | 0.006603620218425514 | Akap13/Peak1/Tssk4/Prkca/Zap70/Kit/Dapk3/Cerk/Dgka/Camk1d/Ak4/Dgkg/Prkcd/Prkcq/Dgkb/Fam20b/Prag1/Plk3/Map3k5/Camk4/Ak5/Coq8a/Pdk2/Ckm/Prkcg/Hkdc1/Pip4k2c/Ttbk1/Itpk1/Stk25/Igf1r/Dclk2/Map4k3/Acvr1/Wnk1/Nek9/Prkag2/Khk/Fer/Ddr1/Epha4/Pfkl/Dgke/Abl2/Mark4/Mast4/Ksr1/Prkce/Ip6k1/Prkdc/Pi4k2a/Ppip5k1/Cdk14/Erbb4/Gk/Adpgk/Mapk10/Pdk3/Map4k5/Akt3 | 60 |
| MF | GO:0140303 | intramembrane lipid transporter activity | 9/680 | 27/8933 | 1,1E+12 | 0.00748402849273009 | 0.006730371136700026 | Atp8a1/Tmem30a/Abca2/Atg9a/Atp11a/Atp9b/Abca3/Atp9a/Atp8a2 | 9 |

**GO_DG_protein**

| ONTOLOGY | ID | Description | GeneRatio | BgRatio | pvalue | p.adjust | qvalue | geneID | Count |
|---|---|---|---|---|---|---|---|---|---|
| BP | GO:0006397 | mRNA processing | 250/1630 | 377/8933 | 5,6E-84 | 3,29164E-79 | 2,99634E-80 | Khdrbs2/Ptbp3/Rbm4/Rprd1a/Dcps/Prpf38a/Rbmxl1/Prpf38b/Thoc2/Khdrbs1/Sf3a2/Hnrnpa3/Thoc6/Srsf6/Hnrnpu/Wtap/Thoc1/Tardbp/Cirbp/Psip1/Zcchc8/Luc7l2/Hnrnpa0/Hnrnph1/Alkbh5/Thoc5/Rbm4b/U2af2/Celf1/Hnrnpc/Srsf7/Ptbp2/Cpsf4/Snrpe/Khsrp/Srrm1/Plrg1/Rbm39/Srsf3/Sfpq/Dhx9/Snrnp70/Ybx1/Tia1/U2af1/Npm1/Srsf12/Rbm7/Tra2b/Rnps1/Nono/Snu13/Hnrnpa2b1/Dbr1/Srsf2/Tra2a/Rbm42/Sfswap/Snrpd1/Rprd1b/Srsf1/Ddx39b/Prpf40a/Rbmx/Hnrnpl/Ddx5/Srrt/Rbbp6/Rbm3/Hnrnpk/Smu1/Luc7l3/Trmt2a/Snrnp27/Snrnp40/Hnrnpf/Usp39/Isy1/Snrpb/Magoh/Snrpg/Frg1/Xab2/Ncbp1/Fip1l1/Safb/Rbm8a/Hnrnpm/Srsf5/Snrpa/Srsf10/Snrpb2/Rbm22/Sf3b3/Ccar2/Rbm14/Acin1/Sf3b4/Iws1/Cpsf6/Sympk/Wbp11/Prpf19/Cdk13/Yju2/Dhx15/Paf1/Eftud2/Rbm15b/Crnkl1/Sf3a1/Rbm15/Rbm27/Cpsf7/Rbfox3/Snrnp200/Ythdc1/Thoc3/Snrpa1/Srsf4/Eif4a3/Nudt21/Prpf8/Ncl/Sf3a3/Alyref/Sf3b1/Rbm25/Bud31/Cdc5l/Mtrex/Zc3h14/Pnn/Cdc40/Snw1/Sf3b2/Sf1/Rbm10/Ddx41/Ddx17/Rbfox1/Cwc22/Scaf8/Ik/Snrpd3/Tcerg1/Rnf113a2/Pabpn1/Sart1/Cpsf1/Larp7/Virma/Cstf3/Cd2bp2/Son/Sart3/Snrpd2/Rbm6/Ctnnbl1/Rbm17/Sugp1/Dus3l/Rprd2/Cmtr1/Luc7l/Prpf31/Ppp1r8/Thrap3/Srrm2/Ddx23/Cdc73/Gpkow/Smndc1/Scaf1/Tfip11/Ddx39a/Adarb1/Rbm5/Nova2/Hnrnpa1/Cstf1/Xrn2/Rpusd2/Celf2/Cstf2/Aqr/Ncbp2/Prpf4b/Zc3h13/Prpf6/Cpsf2/Bcas2/Sap18/Sltm/Sugp2/Prpf3/Papola/Pqbp1/Cdk9/Dhx8/Lsm2/Pcf11/Prpf4/Cwf19l1/Zmat2/Ddx46/Puf60/Dazap1/Lsm6/Nup98/Cnot6l/Cwc15/Dhx16/Rbm26/Hnrnpll/Rnmt/Rpusd3/Ppie/Prpf40b/Trub1/Phf5a/Ssu72/Lsm4/Ddx1/Paxbp1/Cpsf3/Prpf39/Scaf11/Syncrip/Wdr33/Rbfox2/Lsm8/Celf4/Syf2/Dhx38/Htatsf1/Ppp4r2/Fastkd5/Mettl3/Pus1/Rngtt/Cstf2t/Raly/Rbm28/Akap8l/Adar/Prkaca/Fmr1/Csdc2/Fxr1 | 250 |
| BP | GO:0022613 | ribonucleoprotein complex biogenesis | 161/1630 | 319/8933 | 1,6E-24 | 1,03889E-21 | 9,45689E-23 | Sf3a2/Riox1/Srsf6/Nop58/Fbll1/Psip1/Luc7l2/Rpp30/Nop56/Fbl/Celf1/Bop1/Exosc10/Ptbp2/Mak16/Snrpe/Rrs1/Nol9/Ddx21/Dhx9/Npm1/Srsf12/Nhp2/Pelp1/Snu13/Sfswap/Exosc4/Snrpd1/Exosc2/Srsf1/Ddx39b/Rbmx/Mybbp1a/Exosc1/Luc7l3/Brix1/Usp39/Isy1/Snrpb/Pes1/Snrpg/Rpf2/Frg1/Xab2/Ncbp1/Srsf10/Nifk/Emg1/Krr1/Cpsf6/Wbp11/Prpf19/Wdr18/Crnkl1/Sf3a1/Cpsf7/Nat10/Rps24/Gtpbp4/Snrnp200/Nol11/Ythdc1/Nudt21/Prpf8/Ncl/Sf3a3/Exosc8/Sf3b1/Mtrex/Wdr3/Dkc1/Rbm10/Ddx17/Exosc7/Snrpd3/Pdcd11/Bms1/Rpl7l1/Utp6/Exosc3/Urb1/Heatr1/Sart1/Las1l/Sart3/Snrpd2/Npm3/Ago1/Luc7l/Prpf31/Cdc73/Ddx51/Rbm5/Rpusd2/Celf2/Pak1ip1/Dis3/Dhx29/Prpf6/Esf1/Gar1/Prpf3/Utp18/Lsm2/Rrp12/Tbl3/Nup88/Puf60/Lsm6/Wdr36/Exosc5/Urb2/Nop14/Rrp9/Mettl16/Nvl/Rpl7a/Rps8/Rpl24/Riox2/Wdr43/Lsm4/Noc4l/Gnl3l/Prpf39/Bysl/Scaf11/Mdn1/Ercc2/Pwp1/Utp14a/Celf4/Rpl6/Wdr12/Rps25/Rpl14/Nop2/Rpl10/Ago2/Pa2g4/Rrp7a/Rpl5/Rps9/Eif3a/Rpl27/Rps7/Rps14/Heatr3/Rpl11/Rps16/Rpsa/Adar/Xpo1/Rpl7/Rps19/Rplp0/Rps27/Mrps11/Eif3d/Ruvbl1/Tsr2 | 161 |
| BP | GO:0006403 | RNA localization | 80/1630 | 133/8933 | 3,2E-11 | 8,9191E-12 | 8,11895E-10 | Thoc2/Khdrbs1/Hnrnpa3/Thoc6/Nop58/Hnrnpu/Thoc1/Alkbh5/Thoc5/Fbl/Hnrnpab/Exosc10/Srsf7/Khsrp/Srsf3/Dhx9/Ybx1/Npm1/Hnrnpa2b1/Zc3h11a/Srsf1/Ddx39b/Ssb/Poldip3/Magoh/Ncbp1/Rbm8a/Sarnp/Ahctf1/Nxf1/Iws1/Cpsf6/Yy1/Rbm15b/Nup210/Nup35/Ythdc1/Thoc3/Eif4a3/Fyttd1/Alyref/Dkc1/Chtop/Pabpn1/Phax/Ddx39a/Hnrnpa1/Ncbp2/Prpf6/Fubp3/Nup107/Nup188/Nup43/Nup88/Nup98/Nup160/Nup155/Tpr/Nup133/Ranbp2/Pom121/Nup85/Nup93/Nup37/Xpo5/Nup62/Nup54/Nup153/Seh1l/Nup214/Pcid2/Rae1/Akap8l/Tst/Aaas/Xpo1/Fmr1/G3bp2/Nsun2/Rftn2 | 80 |
| BP | GO:0050657 | nucleic acid transport | 71/1630 | 112/8933 | 2,7E-11 | 6,75169E-10 | 6,14598E-09 | Thoc2/Khdrbs1/Hnrnpa3/Thoc6/Thoc1/Alkbh5/Thoc5/Srsf7/Khsrp/Srsf3/Dhx9/Ybx1/Npm1/Hnrnpa2b1/Zc3h11a/Srsf1/Ddx39b/Ssb/Poldip3/Magoh/Ncbp1/Rbm8a/Sarnp/Ahctf1/Nxf1/Iws1/Cpsf6/Rbm15b/Nup210/Nup35/Ythdc1/Thoc3/Eif4a3/Fyttd1/Alyref/Chtop/Pabpn1/Phax/Ddx39a/Hnrnpa1/Ncbp2/Nup107/Nup188/Nup43/Nup88/Nup98/Nup160/Nup155/Tpr/Nup133/Ranbp2/Pom121/Nup85/Nup93/Nup37/Xpo5/Nup62/Nup54/Nup153/Seh1l/Nup214/Pcid2/Rae1/Akap8l/Tst/Aaas/Xpo1/Fmr1/G3bp2/Nsun2/Rftn2 | 71 |
| BP | GO:0050658 | RNA transport | 71/1630 | 112/8933 | 2,7E-11 | 6,75169E-10 | 6,14598E-09 | Thoc2/Khdrbs1/Hnrnpa3/Thoc6/Thoc1/Alkbh5/Thoc5/Srsf7/Khsrp/Srsf3/Dhx9/Ybx1/Npm1/Hnrnpa2b1/Zc3h11a/Srsf1/Ddx39b/Ssb/Poldip3/Magoh/Ncbp1/Rbm8a/Sarnp/Ahctf1/Nxf1/Iws1/Cpsf6/Rbm15b/Nup210/Nup35/Ythdc1/Thoc3/Eif4a3/Fyttd1/Alyref/Chtop/Pabpn1/Phax/Ddx39a/Hnrnpa1/Ncbp2/Nup107/Nup188/Nup43/Nup88/Nup98/Nup160/Nup155/Tpr/Nup133/Ranbp2/Pom121/Nup85/Nup93/Nup37/Xpo5/Nup62/Nup54/Nup153/Seh1l/Nup214/Pcid2/Rae1/Akap8l/Tst/Aaas/Xpo1/Fmr1/G3bp2/Nsun2/Rftn2 | 71 |

| | | | | | | | | | |
|---|---|---|---|---|---|---|---|---|---|
| BP | GO:0006338 | chromatin remodeling | 100/1630 | 203/8933 | 2,5E-08 | 4,68143E-07 | 4,26145E-07 | Cenpv/Dpf3/Hp1bp3/Hdac1/Baz1b/Macroh2a1/Riox1/Smarca2/Gatad2b/Samd1/Hmgb1/Dek/Smarca5/Top1/Mta2/Brd4/Chd4/Smarcad1/Hdac2/Lmnb2/Mbd3/Pbrm1/Smchd1/Mta3/Ddx21/Kmt2a/Chd3/Rbbp5/Trim28/Npm1/Mta1/Kdm6a/Mecp2/Rbbp7/Cbx8/Rsf1/Kat2a/Mybbp1a/Cbx3/Rbbp4/Atrx/Wdr5/Arid2/Rnf2/Ehmt1/Dmap1/Anp32b/Yy1/Brd7/Hira/Macroh2a2/Bcl7a/Phf2/Smarcd1/Actl6a/Rybp/Kdm5a/Sin3a/Sf3b1/Smarcc2/Smarce1/Ash2l/Suz12/Dpf1/Ssrp1/Kdm1a/Smarcb1/Sart3/Npm3/Tasor/Ddx23/Cfdp1/Lmnb1/Ctr9/Arid1b/Smarcc1/Set/Dnmt1/Nasp/Dnajc9/Dpy30/Kmt2d/Kdm5b/Arid1a/Yeats2/Brd3/Actl6b/Tpr/Morf4l1/Smarca4/Men1/Hcfc1/Setd1a/Bptf/Brd9/Smyd3/Yeats4/Pcid2/Chd6/Ruvbl1 | 100 |
| BP | GO:0002181 | cytoplasmic translation | 67/1630 | 122/8933 | 5,9E-05 | 0,008169773 | 0,000743685 | Rbm4/Hnrnpu/Hnrnpd/Dhx9/Ybx1/Ncbp1/Rps24/Rps26/Cpeb4/Ncbp2/Dhx29/Rpl39/Rpl36/Rpl7a/Rps8/Rpl24/Rpl29/Rpl21/Rpl34/Rps23/Syncrip/Rpl12/Rpl18/Rpl3/Rpl27a/Rpl8/Rpl6/Rpl31/Rpl18a/Rps4x/Rps18/Rps25/Rpl14/Mettl3/Rps13/Rpl22/Rpl15/Eif4a1/Rpl13/Rpl37a/Rpl4/Rps2/Rpl17/Rpl5/Rpl10a/Rps9/Eif3a/Rpl23/Rpl27/Rps15a/Rps20/Rps7/Rps14/Rpl11/Rps16/Rps10/Rpsa/Fmr1/Rpl30/Rpl7/Rps19/Rplp0/Eif3i/Rps3/Eif4a2/Rps11/Eif3d | 67 |
| BP | GO:0006406 | mRNA export from nucleus | 38/1630 | 49/8933 | 0,00209 | 0,274569268 | 0,249937079 | Thoc2/Thoc6/Thoc1/Alkbh5/Thoc5/Srsf3/Hnrnpa2b1/Zc3h11a/Ddx39b/Poldip3/Magoh/Ncbp1/Rbm8a/Sarnp/Nxf1/Iws1/Rbm15b/Ythdc1/Thoc3/Eif4a3/Fyttd1/Alyref/Chtop/Pabpn1/Ddx39a/Nup107/Nup88/Nup160/Nup155/Tpr/Nup133/Nup85/Nup93/Nup214/Pcid2/Rae1/Akap8l/Nsun2 | 38 |
| BP | GO:0022618 | ribonucleoprotein complex assembly | 76/1630 | 153/8933 | 0,00051 | 0,000652382 | 0,059385524 | Sf3a2/Srsf6/Psip1/Luc7l2/Celf1/Bop1/Ptbp2/Snrpe/Rrs1/Dhx9/Srsf12/Snu13/Sfswap/Snrpd1/Srsf1/Ddx39b/Rbmx/Luc7l3/Brix1/Usp39/Isy1/Snrpb/Snrpg/Rpf2/Xab2/Ncbp1/Srsf10/Cpsf6/Prpf19/Crnkl1/Sf3a1/Cpsf7/Snrnp200/Ythdc1/Nudt21/Prpf8/Sf3a3/Sf3b1/Snrpd3/Sart3/Snrpd2/Ago1/Luc7l/Prpf31/Cdc73/Rbm5/Celf2/Dhx29/Prpf6/Prpf3/Lsm2/Puf60/Rpl24/Lsm4/Prpf39/Scaf11/Mdn1/Celf4/Rpl6/Rps25/Nop2/Rpl10/Ago2/Rrp7a/Rpl5/Eif3a/Rps14/Rpl11/Rpsa/Adar/Rps19/Rplp0/Rps27/Mrps11/Eif3d/Ruvbl1 | 76 |
| BP | GO:0071826 | ribonucleoprotein complex subunit organization | 78/1630 | 160/8933 | 0,00725 | 0,089306904 | 0,081294993 | Sf3a2/Srsf6/Psip1/Luc7l2/Celf1/Bop1/Ptbp2/Snrpe/Rrs1/Dhx9/Srsf12/Snu13/Sfswap/Snrpd1/Srsf1/Ddx39b/Rbmx/Luc7l3/Brix1/Usp39/Isy1/Snrpb/Snrpg/Rpf2/Xab2/Ncbp1/Srsf10/Cpsf6/Prpf19/Crnkl1/Sf3a1/Cpsf7/Snrnp200/Ythdc1/Nudt21/Prpf8/Sf3a3/Sf3b1/Snrpd3/Sart3/Snrpd2/Ago1/Luc7l/Prpf31/Cdc73/Tfip11/Rbm5/Celf2/Dhx29/Prpf6/Prpf3/Dhx8/Lsm2/Puf60/Rpl24/Lsm4/Prpf39/Scaf11/Mdn1/Celf4/Rpl6/Rps25/Nop2/Rpl10/Ago2/Rrp7a/Rpl5/Eif3a/Rps14/Rpl11/Rpsa/Adar/Rps19/Rplp0/Rps27/Mrps11/Eif3d/Ruvbl1 | 78 |
| BP | GO:0016570 | histone modification | 102/1630 | 242/8933 | 0,01775 | 2,14244116 | 0,195023824 | Phf20l1/Msl2/Mideas/Hdac1/Baz1b/Riox1/Nfyc/Gatad2b/Fbll1/Dek/Smarca5/Mta2/Fbl/Brd4/Chd4/Smarcad1/Hdac2/Mbd3/Mta3/Ddx21/Pcgf2/Kmt2a/Chd3/Sfpq/Rbbp5/Mta1/Kdm6a/Mecp2/Glyr1/Rbbp7/Cbx8/Kat2a/Mybbp1a/Trrap/Rbbp4/Prkd1/Wdr5/Rnf2/Sf3b3/Ehmt1/Dmap1/Iws1/Sap30bp/Taf4/Paf1/Ep400/Sin3b/Actl6a/Kdm5a/Taf5l/Sin3a/Sf3b1/Snw1/Ash2l/Suz12/Meaf6/Rcor3/Chtop/Ncor2/Kdm1a/Smarcb1/Sart3/Kdm1b/Nfyb/Brms1l/Cdc73/Nelfe/Ctr9/Supt3/Kmt2d/Kdm5b/Cdk9/Msl3/Yeats2/Wdr82/Actl6b/Epc2/Morf4l1/Nelfa/Smarca4/Men1/Hcfc1/Setd1a/Riox2/Smyd3/Rnf40/Yeats4/Rnf20/Paxbp1/Rtf1/Otub2/Atf2/Nsd3/Crebbp/Mrgbp/Prmt1/Prmt8/Skic8/Akap8l/Tbl1xr1/Hdac3/Ruvbl1 | 102 |
| BP | GO:0006974 | cellular response to DNA damage stimulus | 158/1630 | 459/8933 | 0,13792 | 14,83476952 | 13,50391104 | Dpf3/Apex1/H2ax/Cbx5/Vrk1/Baz1b/Hmga1/Smarca2/Pogz/Hmgb1/Thoc1/Cbx1/Tmem109/Hmgn1/Dek/Smarca5/Thoc5/Pds5b/Brd4/Chd4/Smarcad1/Pbrm1/Smchd1/Hdgfl2/Vav3/Sfpq/Dhx9/Fus/Rbbp5/Trim28/Npm1/Mta1/Ints7/Nono/Cbx8/Ddx39b/Kat2a/Mcm6/Ddx5/Rbbp6/Hnrnpk/Cbx3/Smc1a/Trrap/Ppp4r3b/Bclaf1/Xab2/Rad21/Atrx/Arid2/Sf3b3/Ccar2/Dmap1/Ppp1r10/Terf2/Ints3/Prpf19/Taf4/Yy1/Terf2ip/Nsmce3/Brd7/Polr2i/Gtf2h1/Bcl7a/Ep400/Smarcd1/Gtf2h3/Actl6a/Smc3/Taf5l/Smarcc2/Rpa1/Cdc5l/Mtrex/Snw1/Smarce1/Mapk3/Ash2l/Polb/Meaf6/Dpf1/Cdkn2aip/Parp1/Fto/Ssrp1/Kdm1a/Zmynd8/Smarcb1/Pds5a/Ercc4/Foxo3/Nabp2/Xrn2/Pcna/Msh3/Sfr1/Arid1b/Smarcc1/Stn1/Supt3/Tlk1/Nudt16l1/Cdk9/Cdk7/Arid1a/Ercc3/Pwwp3a/Ctc1/Actl6b/Lig3/Mapk14/Rrm2b/Epc2/Morf4l1/Smarca4/Men1/Ccnk/Bax/Wrnip1/Pold1/Lyn/Setd1a/Msh2/Yeats4/Steap3/Ddx1/Casp3/Rfc5/Ercc2/Macrod1/Smarcal1/Exd2/Spred2/Mtch2/Ppp4r3a/Syf2/Otub2/Atf2/Ercc1/Ppp4r2/Eepd1/Nedd4/Mettl3/Mrgbp/Mapk1/Polg2/Upf1/Fmr1/Pnp/Rps3/Htra2/Tnks1bp1/Usp10/Adprs/Usp47/Ruvbl1/Fxr1 | 158 |
| BP | GO:0051168 | nuclear export | 64/1630 | 125/8933 | 0,4651 | 48,27257113 | 4,394193693 | Thoc2/Khdrbs1/Thoc6/Thoc1/Alkbh5/Thoc5/Rrs1/Srsf3/Dhx9/Npm1/Hnrnpa2b1/Zc3h11a/Ddx39b/Ssb/Poldip3/Magoh/Ncbp1/Ctdspl2/Rbm8a/Sarnp/Rbm22/Nxf1/Iws1/Anp32b/Cpsf6/Rbm15b/Ythdc1/Thoc3/Eif4a3/Fyttd1/Alyref/Rbm10/Chtop/Pabpn1/Phax/Ddx39a/Ncbp2/Ube2i/Nup107/Nup188/Nup88/Nup98/Nup160/Nup155/Tpr/Nup133/Ranbp2/Pom121/Nup85/Nup93/Xpo5/Nup62/Mdn1/Nup153/Ranbp3/Nup214/Pcid2/Rae1/Akap8l/Adar/Xpo1/Prkaca/Hdac3/Nsun2 | 64 |

| | | | | | | | | | |
|---|---|---|---|---|---|---|---|---|---|
| BP | GO:0006401 | RNA catabolic process | 78/1630 | 183/8933 | 1,00429 | 91,40553132 | 832,0534825 | Apex1/Dcps/Hnrnpu/Tardbp/Polr2g/Cirbp/Elavl1/Hnrnpa0/Alkbh5/Hnrnpab/Celf1/Hnrnpc/Hnrnpr/Exosc10/Hnrnpd/Khsrp/Dhx9/Fus/Zhx2/Ybx1/Npm1/Rbm7/Zc3h4/Rnps1/Exosc4/Exosc2/Srsf1/Ddx5/Exosc1/Ssb/Magoh/Ncbp1/Rbm8a/Eif4a3/Exosc8/Mtrex/Zc3h14/Dkc1/Rbm10/Dcp1a/Exosc7/Exosc3/Fto/Ago1/Thrap3/Phax/Xrn2/Ncbp2/Dis3/Edc4/Lsm2/Lsm6/Wdr82/Cnot6l/Exosc5/Mettl16/Lsm4/Syncrip/Dcp1b/Pum1/Fastkd5/Mettl3/Pcid2/Ago2/Ythdf1/Angel2/Edc3/Supv3l1/Rc3h2/Cnot1/Upf1/Fmr1/Upf2/Csdc2/Eif3e/Gtpbp1/Nsun2/Fxr1 | 78 |
| BP | GO:0045892 | negative regulation of DNA-templated transcription | 160/1630 | 500/8933 | 156,711 | 13244,3532 | 1205,617433 | Prox1/Rfx3/Nfib/Bhlhe22/Nfix/Prdm8/Maz/Pdcd4/Mideas/Zbtb20/Cbx5/Hdac1/Magee2/Khdrbs1/Hmga1/Macroh2a1/Riox1/Smarca2/Zfp512b/Gatad2b/Samd1/Hnrnpu/Hmgb1/Cbx1/Smad3/Cbx6/Smarca5/Foxk1/Mta2/Hnrnpab/Chd4/Hdac2/Mbd3/Mef2a/Mta3/Pcgf2/Chd3/Sfpq/Zhx2/Trim28/Ybx1/Rps6ka5/Mta1/Mecp2/Zc3h4/Nono/Tle1/Hnrnpa2b1/Rbbp7/Cbx8/Rsf1/Mybbp1a/Hnrnpl/Ddx5/Hnrnpk/Cbx3/Hmg20a/Rbbp4/Hdgf/Bclaf1/Myef2/Ilf3/Nr2f1/Sarnp/Spen/Rnf2/Ccar2/Ehmt1/Pias1/Dmap1/Hexim1/Zbtb18/Yy1/Nsmce3/Brd7/Paf1/Macroh2a2/Rbm15/Sin3b/Rybp/Kdm5a/Sin3a/Smarcc2/Tgfbr1/Snw1/Smarce1/Creb1/Pspc1/Suz12/Phf14/Rbm10/Rcor3/Nr2e1/Scaf8/Parp1/Tcerg1/Cggbp1/Ncor2/Hmgb2/Thap11/Kdm1a/Larp7/Sap130/Zmynd8/Pcbp3/Kdm1b/Tle4/Mageh1/Brms1l/Cdc73/Nelfe/Foxo3/Med25/Ctr9/Pcna/Ube2i/Dnmt1/Ylpm1/Sap18/Mepce/Kdm5b/Arid1a/Yeats2/Wdr82/Zhx3/Nmnat1/Tpr/Nelfa/Smarca4/Men1/Gabpa/Hcfc1/Nrip2/Taf9b/Bptf/Zgpat/Rbfox2/Rtf1/Atf2/Nedd4/Nsd3/Irf2bpl/Rpl10/Pcid2/Crebbp/Pa2g4/Trps1/Nacc1/Dcaf1/Rpl23/Rps14/Tbl1xr1/Cnot1/Xpo1/Prkaca/Hdac3/Pura/Mtdh/Usp47/Prickle1 | 160 |
| BP | GO:0010608 | post-transcriptional regulation of gene expression | 110/1630 | 313/8933 | 334,037 | 240995,4292 | 219375,2208 | Rbm4/Shmt1/Apex1/Dcps/Khdrbs1/Hnrnpu/Tardbp/Polr2g/Cirbp/Elavl1/Hnrnpa0/Alkbh5/Rbm4b/Celf1/Hnrnpc/Hnrnpr/Hnrnpd/Dapk1/Khsrp/Dhx9/Fus/Ybx1/Tia1/Npm1/Mecp2/Pcif1/Exosc2/Srsf1/Ddx39b/Rbm3/Ssb/Poldip3/Magoh/Ilf3/Ncbp1/Nr2f1/Sarnp/Matr3/Nolc1/Nat10/Ythdc1/Eif4a3/Nudt21/Ncl/Exosc8/Zc3h14/Enc1/Mapk3/Dkc1/Rbm10/Dcp1a/Exosc7/Exosc3/Fto/Dus3l/Ago1/Cpeb4/Thrap3/Phax/Lsm14b/Celf2/Ncbp2/Cnot6l/Exosc5/Tpr/Rpusd3/Mettl16/Trub1/Akt2/Xpo5/Tial1/Syncrip/Dcp1b/Pum3/Pum1/Lsm14a/Fastkd5/Mettl3/Rpl10/Pcid2/Ago2/Pa2g4/Ythdf1/Angel2/Mapk1/Rpl5/Ogfod1/Cnot11/Rc3h2/Ddx6/Shmt2/Cnot1/Upf1/Plxnb2/Adar/Fmr1/Csdc2/Eif3b/Rack1/Eif3e/Rps3/Caprin1/Nsun2/Scrib/Eif3d/Epb41l5/Fxr1/Msi1/Mrpl13/Ppp1ca | 110 |
| BP | GO:0033044 | regulation of chromosome organization | 63/1630 | 145/8933 | 1369,02 | 88033,77986 | 80136,08373 | Cenpv/Dpf3/Baz1b/Macroh2a1/Smarca2/Hnrnpu/Smarca5/Numa1/Hnrnpc/Exosc10/Hnrnpd/Pbrm1/Sfpq/Trim28/Hnrnpa2b1/Nek6/Rad21/Atrx/Arid2/Ppp1r10/Terf2/Yy1/Terf2ip/Brd7/Nat10/Bcl7a/Smarcd1/Actl6a/Sin3a/Smarcc2/Smarce1/Mapk3/Dkc1/Ik/Dpf1/Parp1/Smarcb1/Bub3/Tasor/Ercc4/Nabp2/Arid1b/Smarcc1/Stn1/Dnmt1/Ylpm1/Arid1a/Ctc1/Sub1/Actl6b/Tpr/Smarca4/Gnl3/Map3k4/Gnl3l/Ercc1/Pcid2/Mad2l1/Mapk1/Upf1/Psmg2/Cdc23/Ruvbl1 | 63 |
| BP | GO:0019827 | stem cell population maintenance | 47/1630 | 97/8933 | 9171,48 | 548065,1375 | 498897,0576 | Prox1/Nfib/Nfix/Vangl2/Hdac1/Smarca2/Elavl1/Hdac2/Zhx2/Rbbp7/Kat2a/Srrt/Smc1a/Rbbp4/Med24/Paf1/Bcl7a/Smarcd1/Stag2/Actl6a/Smc3/Taf5l/Sin3a/Smarce1/Nr2e1/Sap130/Smarcb1/Brms1l/Cdc73/Foxo3/Ctr9/Zc3h13/Smarcc1/Arid1a/Actl6b/Smarca4/Gnl3/Setd1a/Brd9/Wdr43/Rtf1/Mettl3/Crebbp/Med15/Ddx6/Cnot1/Cdh2 | 47 |
| BP | GO:0044270 | cellular nitrogen compound catabolic process | 94/1630 | 266/8933 | 131290 | 769020,4881 | 700030,0376 | Nt5c1a/Apex1/Dcps/Hnrnpu/Tardbp/Upp1/Polr2g/Cirbp/Elavl1/Hnrnpa0/Alkbh5/Hnrnpab/Celf1/Hnrnpc/Hnrnpr/Exosc10/Hnrnpd/Khsrp/Dhx9/Fus/Zhx2/Ybx1/Npm1/Rbm7/Zc3h4/Rnps1/Exosc4/Exosc2/Srsf1/Ddx5/Exosc1/Ssb/Magoh/Ncbp1/Rbm8a/Cyp3a13/Eif4a3/Exosc8/Mtrex/Zc3h14/Dkc1/Rbm10/Dcp1a/Exosc7/Exosc3/Fto/Ago1/Thrap3/Phax/Xrn2/Ncbp2/Dis3/Edc4/Apaf1/Lsm2/Lsm6/Wdr82/Cnot6l/Nmnat1/Exosc5/Bax/Mettl16/Lsm4/Casp3/Syncrip/Dcp1b/Pum1/Fastkd5/Mettl3/Pcid2/Ago2/Ythdf1/Angel2/Dut/Adal/Gda/Edc3/Supv3l1/Rc3h2/Cnot1/Upf1/Fmr1/Upf2/Pnp/Csdc2/Eif3e/Gtpbp1/Aldh6a1/Nsun2/Acadl/Blvrb/Blvra/Fxr1/Entpd3 | 94 |
| BP | GO:0098727 | maintenance of cell number | 47/1630 | 98/8933 | 145504 | 835731,9409 | 760756,6652 | Prox1/Nfib/Nfix/Vangl2/Hdac1/Smarca2/Elavl1/Hdac2/Zhx2/Rbbp7/Kat2a/Srrt/Smc1a/Rbbp4/Med24/Paf1/Bcl7a/Smarcd1/Stag2/Actl6a/Smc3/Taf5l/Sin3a/Smarce1/Nr2e1/Sap130/Smarcb1/Brms1l/Cdc73/Foxo3/Ctr9/Zc3h13/Smarcc1/Arid1a/Actl6b/Smarca4/Gnl3/Setd1a/Brd9/Wdr43/Rtf1/Mettl3/Crebbp/Med15/Ddx6/Cnot1/Cdh2 | 47 |
| BP | GO:2001020 | regulation of response to DNA damage stimulus | 72/1630 | 184/8933 | 16076,8 | 914522,516 | 832478,7716 | Dpf3/Baz1b/Smarca2/Pogz/Hmgb1/Thoc1/Dek/Smarca5/Thoc5/Brd4/Pbrm1/Smchd1/Hdgfl2/Dhx9/Fus/Trim28/Npm1/Cbx8/Ddx39b/Kat2a/Ddx5/Hnrnpk/Trrap/Ppp4r3b/Bclaf1/Arid2/Sf3b3/Ccar2/Dmap1/Ppp1r10/Taf4/Yy1/Terf2ip/Brd7/Bcl7a/Ep400/Smarcd1/Actl6a/Taf5l/Smarcc2/Smarce1/Meaf6/Dpf1/Parp1/Kdm1a/Smarcb1/Ercc4/Pcna/Arid1b/Smarcc1/Supt3/Nudt16l1/Cdk9/Arid1a/Actl6b/Epc2/Morf4l1/Smarca4/Yeats4/Steap3/Spred2/Mtch2/Otub2/Ercc1/Ppp4r2/Mrgbp/Fmr1/Pnp/Rps3/Usp47/Ruvbl1/Fxr1 | 72 |

| | | | | | | | | | |
|---|---|---|---|---|---|---|---|---|---|
| BP | GO:0043414 | macromolecule methylation | 66/1630 | 165/8933 | 36581,6 | 19674264,37 | 17909244,61 | Trmt10b/Nfyc/Fbll1/Wtap/Smarca5/Mta2/Fbl/Hdac2/Kmt2a/Rbbp5/Trim28/Rab3b/Mecp2/Pcif1/Snrpb/Ilf3/Wdr5/Emg1/Ehmt1/Iws1/Fam98b/Paf1/Rbm15b/Rbm15/Trmt6/Snw1/Mettl1/Ash2l/Suz12/Chtop/Snrpd3/Parp1/Kdm1a/Larp7/Virma/Smarcb1/Trmt61a/Kdm1b/Nfyb/Cmtr1/Nelfe/Ctr9/Zc3h13/Dnmt1/Mepce/Kmt2d/Wdr82/Nelfa/Smarca4/Men1/Hcfc1/Mettl16/Setd1a/Smyd3/Rnf20/Paxbp1/Rtf1/Nop2/Nsd3/Mettl3/Thumpd3/Rab3d/Prmt1/Prmt8/Skic8/Nsun2 | 66 |
| BP | GO:0046700 | heterocycle catabolic process | 92/1630 | 265/8933 | 66397,5 | 34525848,26 | 31428461,57 | Nt5c1a/Apex1/Dcps/Hnrnpu/Tardbp/Upp1/Polr2g/Cirbp/Elavl1/Hnrnpa0/Alkbh5/Hnrnpab/Celf1/Hnrnpc/Hnrnpr/Exosc10/Hnrnpd/Khsrp/Dhx9/Fus/Zhx2/Ybx1/Npm1/Rbm7/Zc3h4/Rnps1/Exosc4/Exosc2/Srsf1/Ddx5/Exosc1/Ssb/Magoh/Ncbp1/Rbm8a/Eif4a3/Exosc8/Mtrex/Zc3h14/Dkc1/Rbm10/Dcp1a/Exosc7/Exosc3/Fto/Ago1/Thrap3/Phax/Xrn2/Ncbp2/Dis3/Edc4/Apaf1/Lsm2/Lsm6/Wdr82/Cnot6l/Nmnat1/Exosc5/Bax/Mettl16/Lsm4/Casp3/Syncrip/Dcp1b/Pum1/Fastkd5/Mettl3/Pcid2/Ago2/Ythdf1/Angel2/Dut/Adal/Gda/Edc3/Supv3l1/Rc3h2/Cnot1/Upf1/Fmr1/Upf2/Pnp/Csdc2/Eif3e/Gtpbp1/Aldh6a1/Nsun2/Blvrb/Blvra/Fxr1/Entpd3 | 92 |
| BP | GO:0019439 | aromatic compound catabolic process | 93/1630 | 274/8933 | 2104461 | 100403131,7 | 913957,5496 | Nt5c1a/Apex1/Dcps/Hnrnpu/Tardbp/Upp1/Polr2g/Cirbp/Elavl1/Hnrnpa0/Alkbh5/Hnrnpab/Celf1/Hnrnpc/Hnrnpr/Exosc10/Hnrnpd/Khsrp/Dhx9/Fus/Zhx2/Ybx1/Npm1/Rbm7/Zc3h4/Rnps1/Exosc4/Exosc2/Srsf1/Ddx5/Exosc1/Ssb/Magoh/Ncbp1/Rbm8a/Eif4a3/Moxd1/Exosc8/Mtrex/Zc3h14/Dkc1/Rbm10/Dcp1a/Exosc7/Exosc3/Fto/Ago1/Thrap3/Phax/Xrn2/Ncbp2/Dis3/Edc4/Apaf1/Lsm2/Lsm6/Wdr82/Cnot6l/Nmnat1/Exosc5/Bax/Mettl16/Lsm4/Casp3/Syncrip/Dcp1b/Pum1/Fastkd5/Mettl3/Pcid2/Ago2/Ythdf1/Angel2/Dut/Adal/Gda/Edc3/Supv3l1/Rc3h2/Cnot1/Upf1/Fmr1/Upf2/Pnp/Csdc2/Eif3e/Gtpbp1/Comt/Nsun2/Blvrb/Blvra/Fxr1/Entpd3 | 93 |
| BP | GO:1901361 | organic cyclic compound catabolic process | 95/1630 | 283/8933 | 264369 | 123149947,3 | 112101905,7 | Nt5c1a/Apex1/Dcps/Hnrnpu/Tardbp/Upp1/Polr2g/Cirbp/Elavl1/Hnrnpa0/Alkbh5/Hnrnpab/Celf1/Hnrnpc/Hnrnpr/Exosc10/Hnrnpd/Khsrp/Dhx9/Fus/Zhx2/Ybx1/Npm1/Rbm7/Zc3h4/Rnps1/Exosc4/Exosc2/Srsf1/Ddx5/Exosc1/Ssb/Magoh/Ncbp1/Rbm8a/Eif4a3/Moxd1/Exosc8/Mtrex/Zc3h14/Dkc1/Rbm10/Dcp1a/Exosc7/Hsd11b1/Exosc3/Fto/Ago1/Thrap3/Phax/Xrn2/Ncbp2/Dis3/Edc4/Apaf1/Lsm2/Lsm6/Wdr82/Cnot6l/Nmnat1/Exosc5/Bax/Mettl16/Lsm4/Casp3/Syncrip/Dcp1b/Pum1/Fastkd5/Mettl3/Pcid2/Ago2/Ythdf1/Angel2/Dut/Adal/Gda/Edc3/Supv3l1/Rc3h2/Cnot1/Upf1/Fmr1/Upf2/Pnp/Csdc2/Eif3e/Gtpbp1/Aldh6a1/Comt/Nsun2/Blvrb/Blvra/Fxr1/Entpd3 | 95 |
| BP | GO:0032259 | methylation | 71/1630 | 191/8933 | 345558 | 15605489,5 | 142054881,1 | Prdm8/Trmt10b/Nfyc/Fbll1/Wtap/Smarca5/Mta2/Fbl/Hdac2/Kmt2a/Rbbp5/Trim28/Rab3b/Mecp2/Pcif1/Trmt2a/Snrpb/Ilf3/Prdm10/Wdr5/Emg1/Ehmt1/Iws1/Fam98b/Paf1/Rbm15b/Rbm15/Trmt6/Snw1/Mettl1/Ash2l/Suz12/Chtop/Snrpd3/Parp1/Kdm1a/Larp7/Virma/Smarcb1/Trmt61a/Kdm1b/Nfyb/Cmtr1/Nelfe/Ctr9/Zc3h13/Dnmt1/Mepce/Kmt2d/Wdr82/Nelfa/Smarca4/Men1/Rnmt/Hcfc1/Mettl16/Setd1a/Smyd3/Rnf20/Paxbp1/Rtf1/Nop2/Nsd3/Mettl3/Thumpd3/Rab3d/Prmt1/Prmt8/Skic8/Comt/Nsun2 | 71 |
| BP | GO:2000036 | regulation of stem cell population maintenance | 24/1630 | 39/8933 | 2,4E+07 | 94361156,82 | 85895818,35 | Hdac1/Smarca2/Elavl1/Hdac2/Rbbp7/Kat2a/Rbbp4/Bcl7a/Smarcd1/Actl6a/Taf5l/Sin3a/Smarce1/Sap130/Smarcb1/Brms1l/Zc3h13/Smarcc1/Arid1a/Actl6b/Smarca4/Brd9/Wdr43/Cnot1 | 24 |
| BP | GO:0000819 | sister chromatid segregation | 54/1630 | 137/8933 | 4167434 | 161140774,9 | 146684496 | Dpf3/Baz1b/Stag1/Macroh2a1/Smarca2/Pogz/Hnrnpu/Smarca5/Numa1/Pds5b/Champ1/Pbrm1/Rrs1/Nek6/Mau2/Smc1a/Rad21/Atrx/Arid2/Brd7/Bcl7a/Smarcd1/Stag2/Actl6a/Smc3/Smarcc2/Smarce1/Ik/Dpf1/Ccsap/Smarcb1/Bub3/Pds5a/Tasor/Top2b/Eml4/Arid1b/Smarcc1/Arid1a/Actl6b/Gtf2b/Tpr/Smarca4/Nup62/Lsm14a/Seh1l/Pcid2/Mad2l1/Map9/Akap8l/Aaas/Psmg2/Cdc23/Prickle1 | 54 |
| BP | GO:0071824 | protein-DNA complex subunit organization | 40/1630 | 89/8933 | 5287547 | 1955070384 | 1779676895 | Cenpv/Hp1bp3/Macroh2a1/Smarca2/Pogz/Hmgb1/Smarca5/Npm1/Rsf1/Gtf2a1/Tbp/Rbbp4/Gtf2f2/Atrx/Arid2/Med24/Pias1/Anp32b/Taf4/Hira/Macroh2a2/Smarcd1/Smarcc2/Smarce1/Creb1/Ssrp1/Smarcb1/Sart3/Med25/Smarcc1/Set/Nasp/Med23/Dnajc9/Arid1a/Gtf2b/Smarca4/Smyd3/Med15/Rpl23 | 40 |
| BP | GO:0018205 | peptidyl-lysine modification | 70/1630 | 198/8933 | 5556163 | 2041631239 | 1858472193 | Phf20l1/Msl2/Baz1b/Nfyc/Dek/Smarca5/Brd4/Hdac2/Mbd3/Senp7/Ddx21/Pcgf2/Kmt2a/Rbbp5/Trim28/Mecp2/Glyr1/Kat2a/Mybbp1a/Trrap/Hmg20a/Senp3/Wdr5/Sf3b3/Ehmt1/Pias1/Dmap1/Iws1/Taf4/Ep400/Actl6a/Taf5l/Sin3a/Sf3b1/Snw1/Ash2l/Suz12/Meaf6/Rnf113a2/Kdm1a/Smarcb1/Uba2/Nfyb/Nelfe/Sae1/Ctr9/Ube2i/Supt3/Kmt2d/Msl3/Yeats2/Wdr82/Actl6b/Epc2/Morf4l1/Nelfa/Smarca4/Men1/Gnl3/Hcfc1/Setd1a/Smyd3/Yeats4/Gnl3l/Rtf1/Atf2/Crebbp/Mrgbp/Skic8/Ruvbl1 | 70 |
| BP | GO:0017148 | negative regulation of translation | 49/1630 | 121/8933 | 7629808 | 271896451,2 | 2475040469 | Rbm4/Shmt1/Dcps/Hnrnpu/Tardbp/Polr2g/Celf1/Hnrnpr/Hnrnpd/Dapk1/Khsrp/Dhx9/Ybx1/Tia1/Pcif1/Exosc2/Ilf3/Eif4a3/Ncl/Exosc8/Enc1/Dcp1a/Exosc7/Exosc3/Fto/Ago1/Cpeb4/Cnot6l/Exosc5/Tpr/Mettl16/Syncrip/Dcp1b/Pum1/Lsm14a/Mettl3/Ago2/Ythdf1/Rc3h2/Ddx6/Cnot1/Upf1/Fmr1/Rack1/Rps3/Caprin1/Scrib/Fxr1/Mrpl13 | 49 |

| | | | | | | | | | |
|---|---|---|---|---|---|---|---|---|---|
| BP | GO:0070316 | regulation of G0 to G1 transition | 18/1630 | 26/8933 | 1,6E+08 | 551566188,9 | 502084022,4 | Dpf3/Smarca2/Pbrm1/Arid2/Brd7/Bcl7a/Smarcd1/Actl6a/Smarcc2/Smarce1/Dpf1/Smarcb1/Arid1b/Smarcc1/Arid1a/Actl6b/Rrm2b/Smarca4 | 18 |
| BP | GO:0030071 | regulation of mitotic metaphase/anaphase transition | 26/1630 | 49/8933 | 3,9E+07 | 12366354659 | 1125694289 | Dpf3/Smarca2/Pbrm1/Nek6/Rad21/Arid2/Brd7/Bcl7a/Smarcd1/Actl6a/Smarcc2/Smarce1/Ik/Dpf1/Smarcb1/Bub3/Arid1b/Smarcc1/Arid1a/Actl6b/Tpr/Smarca4/Pcid2/Mad2l1/Psmg2/Cdc23 | 26 |
| BP | GO:0045023 | G0 to G1 transition | 18/1630 | 27/8933 | 4E+07 | 12769920362 | 11624303863 | Dpf3/Smarca2/Pbrm1/Arid2/Brd7/Bcl7a/Smarcd1/Actl6a/Smarcc2/Smarce1/Dpf1/Smarcb1/Arid1b/Smarcc1/Arid1a/Actl6b/Rrm2b/Smarca4 | 18 |
| BP | GO:0034249 | negative regulation of amide metabolic process | 52/1630 | 139/8933 | 6,3E+07 | 19329391476 | 1759531098 | Rbm4/Shmt1/Hap1/Dcps/Hnrnpu/Tardbp/Polr2g/Celf1/Hnrnpr/Hnrnpd/Dapk1/Khsrp/Dhx9/Ybx1/Tia1/Pcif1/Exosc2/Ilf3/Eif4a3/Ncl/Exosc8/Enc1/Dcp1a/Exosc7/Exosc3/Fto/Ago1/Cpeb4/Cnot6l/Exosc5/Tpr/Mettl16/Syncrip/Dcp1b/Pum1/Lsm14a/Mettl3/Ago2/Ythdf1/Rtn1/Rc3h2/Ddx6/Cnot1/Upf1/Fmr1/Rack1/Rps3/Sorl1/Caprin1/Scrib/Fxr1/Mrpl13 | 52 |
| BP | GO:0031047 | RNA-mediated gene silencing | 31/1630 | 67/8933 | 1,2E+09 | 35673052125 | 32472747352 | Rbm4/Cenpv/Elavl1/Srsf3/Dhx9/Mecp2/Hnrnpa2b1/Ddx5/Srrt/Rbm3/Ncbp1/Nr2f1/Ddx17/Ago1/Ncbp2/Cnot6l/Nup155/Trub1/Xpo5/Tial1/Tsnax/Tsn/Pum1/Mettl3/Ago2/Cnot11/Ddx6/Cnot1/Adar/Fmr1/Fxr1 | 31 |
| BP | GO:2000113 | negative regulation of cellular macromolecule biosynthetic process | 50/1630 | 135/8933 | 1,6E+08 | 4582449174 | 4171347990 | Rbm4/Shmt1/Dcps/Hnrnpu/Tardbp/Polr2g/Celf1/Hnrnpr/Hnrnpd/Dapk1/Khsrp/Dhx9/Ybx1/Tia1/Pcif1/Exosc2/Ilf3/Eif4a3/Ncl/Exosc8/Enc1/Dcp1a/Exosc7/Exosc3/Fto/Ago1/Cpeb4/Cnot6l/Exosc5/Tpr/Mettl16/Syncrip/Dcp1b/Pum1/Lsm14a/Mettl3/Ago2/Ythdf1/Rc3h2/Ddx6/Cnot1/Upf1/Fmr1/Rack1/Rps3/Caprin1/Scrib/Oga/Fxr1/Mrpl13 | 50 |
| BP | GO:0051983 | regulation of chromosome segregation | 30/1630 | 65/8933 | 2,1E+08 | 5809717103 | 5288515124 | Dpf3/Rcc2/Smarca2/Hnrnpu/Numa1/Pbrm1/Nek6/Rad21/Arid2/Brd7/Bcl7a/Smarcd1/Actl6a/Smarcc2/Smarce1/Ik/Dpf1/Smarcb1/Bub3/Arid1b/Smarcc1/Arid1a/Actl6b/Tpr/Smarca4/Pum1/Pcid2/Mad2l1/Psmg2/Cdc23 | 30 |
| BP | GO:0065004 | protein-DNA complex assembly | 32/1630 | 73/8933 | 3,6E+08 | 9384530601 | 8542624561 | Cenpv/Hp1bp3/Macroh2a1/Smarca2/Pogz/Hmgb1/Smarca5/Npm1/Rsf1/Gtf2a1/Tbp/Rbbp4/Gtf2f2/Atrx/Med24/Pias1/Anp32b/Taf4/Hira/Macroh2a2/Creb1/Ssrp1/Sart3/Med25/Set/Nasp/Med23/Dnajc9/Gtf2b/Smarca4/Smyd3/Med15 | 32 |
| BP | GO:0006999 | nuclear pore organization | 11/1630 | 13/8933 | 3,9E+09 | 1,00417E+11 | 914083971,8 | Ahctf1/Nup35/Nup107/Nup98/Nup133/Nup205/Pom121/Nup93/Nup54/Nup153/Seh1l | 11 |
| BP | GO:0098813 | nuclear chromosome segregation | 57/1630 | 167/8933 | 5,6E+08 | 1,37597E+11 | 1,25253E+11 | Dpf3/Baz1b/Rcc2/Stag1/Macroh2a1/Smarca2/Pogz/Hnrnpu/Smarca5/Numa1/Pds5b/Champ1/Pbrm1/Rrs1/Nek6/Mau2/Smc1a/Rad21/Atrx/Arid2/Brd7/Bcl7a/Smarcd1/Stag2/Actl6a/Smc3/Smarcc2/Smarce1/Ik/Dpf1/Ccsap/Smarcb1/Bub3/Pds5a/Tasor/Ercc4/Top2b/Eml4/Arid1b/Smarcc1/Sun1/Arid1a/Actl6b/Gtf2b/Tpr/Smarca4/Nup62/Lsm14a/Seh1l/Pcid2/Mad2l1/Map9/Akap8l/Aaas/Psmg2/Cdc23/Prickle1 | 57 |
| BP | GO:0006260 | DNA replication | 45/1630 | 121/8933 | 5,9E+08 | 1,43276E+11 | 1,30422E+11 | Nfib/Nfix/Nfia/Thoc1/Smarca5/Top1/Ilkap/Dhx9/Npm1/Rbbp7/Mcm6/Rbbp6/Rbbp4/Atrx/Wdr18/Yy1/Gtpbp4/Actl6a/Alyref/Smc3/Sin3a/Rpa1/Polb/Meaf6/Parp1/Ssrp1/Pds5a/Wiz/Pcna/Stn1/Nasp/Cdk9/Ctc1/Lig3/Rrm2b/Wrnip1/Pold1/Rfc5/Smarcal1/Exd2/Polg2/Upf1/Dtd1/Pura/Ruvbl1 | 45 |
| BP | GO:0007059 | chromosome segregation | 69/1630 | 216/8933 | 6,5E+08 | 1,56603E+11 | 1,42554E+11 | Dpf3/Baz1b/Rcc2/Stag1/Macroh2a1/Smarca2/Pogz/Hnrnpu/Smarca5/Top1/Numa1/Pds5b/Brd4/Smarcad1/Champ1/Pbrm1/Rrs1/Nek6/Mau2/Smc1a/Rad21/Atrx/Arid2/Brd7/Bcl7a/Smarcd1/Stag2/Actl6a/Smc3/Smarcc2/Smarce1/Ik/Dpf1/Ccsap/Smarcb1/Bub3/Pds5a/Tasor/Ercc4/Top2b/Eml4/Ube2i/Arid1b/Smarcc1/Sun1/Tlk1/Arid1a/Nup43/Actl6b/Gtf2b/Tpr/Smarca4/Nup37/Nup62/Ercc2/Rcc1/Pum1/Lsm14a/Seh1l/Pcid2/Mad2l1/Map9/Akap8l/Aaas/Hdac3/Rps3/Psmg2/Cdc23/Prickle1 | 69 |
| BP | GO:0044770 | cell cycle phase transition | 86/1630 | 291/8933 | 1,2E+10 | 27894167832 | 2,53917E+11 | Prox1/Nfib/Nfix/Dpf3/Apex1/Nfia/Atp2b4/Rcc2/Macroh2a1/Smarca2/Thoc1/Cirbp/Thoc5/Brd4/Pbrm1/Mta3/Plrg1/Npm1/Mecp2/Ints7/Nek6/Ddx39b/Trrap/Rad21/Arid2/Camk2g/Ccar2/Camk2d/Pias1/Ppp1r10/Anp32b/Ints3/Prpf19/Brd7/Paf1/Bcl7a/Smarcd1/Actl6a/Sin3a/Smarcc2/Cdc5l/Smarce1/Ik/Dpf1/Tpd52l1/Larp7/Smarcb1/Bub3/Cdc73/Hspa2/Lmnb1/Nabp2/Arid1b/Smarcc1/Nasp/Mepce/Cdk7/Arid1a/Ercc3/Ctc1/Actl6b/Mapk14/Rrm2b/Tpr/Smarca4/Men1/Rpl24/Msh2/Cpsf3/Ercc2/Rcc1/Syf2/Atf2/Pcid2/Mad2l1/Crebbp/Rpl17/Cacnb4/Akap8l/Upf1/Camk2b/Appl2/Usp47/Nsun2/Psmg2/Cdc23 | 86 |
| BP | GO:2000736 | regulation of stem cell differentiation | 21/1630 | 41/8933 | 1,7E+10 | 3,79855E+11 | 3457776425 | Hdac1/Gatad2b/Hnrnpu/Mta2/Chd4/Hdac2/Mbd3/Mta3/Chd3/Mta1/Rbbp7/Rbbp4/Cdk13/Nudt21/Men1/Ccnk/Setd1a/Pwp1/Mtch2/Nsun2/Prickle1 | 21 |
| BP | GO:0006334 | nucleosome assembly | 18/1630 | 33/8933 | 2,9E+09 | 6137122112 | 55865479386 | Hp1bp3/Macroh2a1/Smarca2/Smarca5/Npm1/Rsf1/Rbbp4/Atrx/Anp32b/Hira/Macroh2a2/Ssrp1/Sart3/Set/Nasp/Dnajc9/Smarca4/Smyd3 | 18 |
| BP | GO:0140747 | regulation of ncRNA transcription | 12/1630 | 17/8933 | 3,3E+10 | 67638735730 | 61570722038 | Macroh2a1/Zc3h4/Atrx/Macroh2a2/Nol11/Ncl/Larp7/Smarcb1/Mepce/Wdr82/Smarca4/Pwp1 | 12 |
| BP | GO:0098732 | macromolecule deacylation | 33/1630 | 84/8933 | 4,7E+09 | 93772642750 | 85360101119 | Mideas/Hdac1/Gatad2b/Smarca5/Mta2/Chd4/Smarcad1/Hdac2/Mbd3/Mta3/Chd3/Sfpq/Mta1/Rbbp7/Kat2a/Rbbp4/Prkd1/Wdr5/Ccar2/Sap30bp/Sin3b/Sin3a/Rcor3/Ncor2/Brms1l/Msl3/Yeats2/Morf4l1/Spred2/Akap8l/Tbl1xr1/Hdac3/Abhd10 | 33 |
| BP | GO:0010453 | regulation of cell fate commitment | 11/1630 | 15/8933 | 4,8E+09 | 94572154386 | 86087886879 | Hdac1/Gatad2b/Mta2/Chd4/Hdac2/Mbd3/Mta3/Chd3/Mta1/Rbbp7/Rbbp4 | 11 |

| | | | | | | | | | |
|---|---|---|---|---|---|---|---|---|---|
| BP | GO:0042659 | regulation of cell fate specification | 11/1630 | 15/8933 | 4,8E+09 | 94572154386 | 86087886879 | Hdac1/Gatad2b/Mta2/Chd4/Hdac2/Mbd3/Mta3/Chd3/Mta1/Rbbp7/Rbbp4 | 11 |
| BP | GO:0032986 | protein-DNA complex disassembly | 10/1630 | 13/8933 | 6,7E+09 | 1,28446E+12 | 1,16923E+12 | Arid2/Smarcd1/Smarcc2/Smarce1/Ssrp1/Smarcb1/Smarcc1/Arid1a/Smarca4/Rpl23 | 10 |
| BP | GO:0045663 | positive regulation of myoblast differentiation | 16/1630 | 29/8933 | 8,6E+09 | 1,63135E+12 | 1,485E+12 | Dpf3/Smarca2/Pbrm1/Arid2/Brd7/Smarcd1/Actl6a/Smarcc2/Smarce1/Smarcb1/Arid1b/Smarcc1/Arid1a/Actl6b/Mapk14/Smarca4 | 16 |
| BP | GO:0001824 | blastocyst development | 29/1630 | 72/8933 | 9,9E+09 | 1,85728E+12 | 1,69066E+12 | Thoc2/Fbll1/Thoc5/Brd4/Pbrm1/Xab2/Emg1/Matr3/Ints1/Actl6a/Sf3b1/Tgfbr1/Rpl7l1/Plpp4/Smarcb1/Ctr9/Nasp/Necab1/Smarca4/Gabpa/Hcfc1/Bysl/Rtf1/Cdk11b/Nop2/Rpl13/Rrp7a/Tbl1xr1/Cnot1 | 29 |
| BP | GO:0006337 | nucleosome disassembly | 9/1630 | 12/8933 | 2,8E+11 | 4,84046E+11 | 4,40621E+12 | Arid2/Smarcd1/Smarcc2/Smarce1/Ssrp1/Smarcb1/Smarcc1/Arid1a/Smarca4 | 9 |
| BP | GO:0019080 | viral gene expression | 20/1630 | 48/8933 | 1,3E+12 | 0.001979810921055652 | 0.0018021979061458168 | Hdac1/Tardbp/Brd4/Dhx9/Rsf1/Ssb/Hexim1/Snw1/Larp7/Smarcb1/Cdk9/Gtf2b/Smarca4/Pcbp2/Spcs1/Eif3a/Eif3f/Eif3b/Eif3l/Eif3d | 20 |
| BP | GO:0034063 | stress granule assembly | 13/1630 | 25/8933 | 1,4E+12 | 0.002040647004879551 | 0.001857576256532469 | Cirbp/Tia1/Pqbp1/Rps23/Lsm14a/G3bp1/Ythdf1/Ogfod1/Ddx6/Atxn2l/G3bp2/Ubap2l/Prrc2c | 13 |
| BP | GO:0034504 | protein localization to nucleus | 63/1630 | 226/8933 | 2,1E+12 | 0.0030282273273964924 | 0.0027565586646312325 | Hnrnpu/Tardbp/Elavl1/Ints13/Lmnb2/Rrs1/Plrg1/Trim28/Npm1/Srsf1/Ddx5/Hdgf/Rpf2/Hnrnpm/Phip/Prkd1/Rbm22/Nolc1/Dmap1/Paf1/Nup35/Sin3a/Parp1/Larp7/Lmnb1/Banp/Tmem201/Nup107/Mepce/Sun1/Nup188/Nup88/Nup98/Mapk14/Nup155/Tpr/Nup133/Ranbp2/Syne1/Pom121/Nvl/Nup85/Nup93/Nup62/Nup54/Morc3/Dcp1b/Nup153/Atf2/Sun2/Nup214/Tor1a/Nup50/Mapk1/Cacnb4/Heatr3/Rpl11/Adar/Xpo1/Hdac3/Appl2/Hikeshi/Prickle1 | 63 |
| BP | GO:0007623 | circadian rhythm | 37/1630 | 115/8933 | 2,1E+12 | 0.0030745687455254233 | 0.002798742696364554 | Prox1/Rbm4/Hdac1/Cyp7b1/Hnrnpu/Tardbp/Top1/Rbm4b/Hdac2/Hnrnpd/Kmt2a/Sfpq/Dhx9/Mta1/Nono/Mybbp1a/Ddx5/Ccar2/Kdm5a/Sin3a/Creb1/Pspc1/Prkg2/Parp1/Thrap3/Adcy1/Kdm5b/Mycbp2/Btbd9/Mettl3/Nlgn3/Gabrb3/Hdac3/Ass1/Ntrk3/Kcnma1/Ppp1ca | 37 |
| BP | GO:0070988 | demethylation | 14/1630 | 29/8933 | 2,1E+12 | 0.003077850732036386 | 0.00280173024894099 | Apex1/Riox1/Alkbh5/Hnrnpab/Trim28/Kdm6a/Cyp3a13/Phf2/Kdm5a/Fto/Kdm1a/Kdm1b/Riox2/Syncrip | 14 |
| BP | GO:0007346 | regulation of mitotic cell cycle | 73/1630 | 272/8933 | 2,4E+12 | 0.003459174855741394 | 0.0031488449809567226 | Dpf3/Apex1/Rcc2/Smarca2/Smoc2/Hnrnpu/Hmgb1/Ints13/Brd4/Pbrm1/Mta3/Plrg1/Mecp2/Nek6/Trrap/Rad21/Phip/Arid2/Camk2d/Ppp1r10/Anp32b/Ints3/Brd7/Bcl7a/Smarcd1/Actl6a/Sin3a/Smarcc2/Smarce1/Ik/Dpf1/Larp7/Smarcb1/Bub3/Cdc73/Hspa2/Lmnb1/Nabp2/Arid1b/Smarcc1/Mepce/Cdk7/Arid1a/Ercc3/Ctc1/Actl6b/Rrm2b/Tpr/Smarca4/Men1/Rpl24/Msh2/Cpsf3/Nup62/Ercc2/Rcc1/Syf2/Atf2/Cdk11b/Pcid2/Mad2l1/Crebbp/Angel2/Rpl17/Cacnb4/Hdac3/Appl2/Usp47/Insr/Psmg2/Cdc23/Scrib/Nherf1 | 73 |
| BP | GO:0048863 | stem cell differentiation | 38/1630 | 120/8933 | 2,5E+11 | 0.003545873902040312 | 0.003227766072886772 | Sema5a/Hdac1/Gatad2b/Hnrnpu/Mta2/Chd4/Hdac2/Mbd3/Mta3/Chd3/Mta1/Rbbp7/Rbbp4/Nolc1/Cdk13/Nudt21/Aldh1a2/Mapk3/Anxa6/Rdh10/Mapk14/Men1/Ccnk/Setd1a/Phf5a/Ercc2/Pwp1/Mtch2/Sema4f/Pum1/Frzb/Mapk1/Rps7/Cdh2/Nrp2/Nsun2/Prickle1/Sema4b | 38 |
| BP | GO:0045621 | positive regulation of lymphocyte differentiation | 21/1630 | 54/8933 | 2,9E+12 | 0.00405454543007041 | 0.003690803604896662 | Egr3/Smarca2/Pbrm1/Arid2/Brd7/Smarcd1/Atp11c/Actl6a/Smarcc2/Smarce1/Sart1/Smarcb1/Ap3b1/Arid1b/Smarcc1/Arid1a/Actl6b/Smarca4/Pcid2/Itpkb/Pnp | 21 |
| BP | GO:0050927 | positive regulation of positive chemotaxis | 7/1630 | 10/8933 | 4,8E+11 | 0.0062618783217313415 | 0.00570011200561858 | Hmgb1/Smad3/Cdh13/Scg2/Akt2/S1pr1/Ntrk3 | 7 |
| BP | GO:0140694 | non-membrane-bounded organelle assembly | 67/1630 | 251/8933 | 5,1E+11 | 0.006681230502431737 | 0.006081843217401652 | Prox1/Tmod1/Stag1/Pogz/Hnrnpu/Cirbp/Numa1/Bop1/Hdac2/Rrs1/Mef2a/Tia1/Npm1/Kat2a/Smc1a/Brix1/Rpf2/Rbm14/Pgm5/Stag2/Smc3/Tmod4/Ccsap/Tasor/Dhx29/Pqbp1/Cnot6l/Gtf2b/Tpr/Rpl24/Lsm4/Nup62/Rps23/Mdn1/Rcc1/Rpl6/Lsm14a/Rps25/Nop2/Rpl10/G3bp1/Ythdf1/Actn2/Rrp7a/Rpl5/Ogfod1/Map9/Edc3/Rps14/Ddx6/Rpl11/Cnot1/Aaas/Rpsa/Atxn2l/Hdac3/Rps19/Rplp0/Rps3/G3bp2/Rps27/Ubap2l/Dbnl/Mrps11/Prickle1/Prrc2c/Fscn1 | 67 |
| BP | GO:0072331 | signal transduction by p53 class mediator | 28/1630 | 83/8933 | 5,1E+11 | 0.006719053202341685 | 0.0061162727631009686 | Tmem109/Bop1/Rrs1/Npm1/Mybbp1a/Ddx5/Hnrnpk/Rpf2/Atrx/Hexim1/Snw1/Kdm1a/Foxo3/Pak1ip1/Rrm2b/Bax/Msh2/Steap3/Armc10/Spred2/Nop2/Rpl5/Rpl23/Rps20/Rps7/Rpl11/Ppp1r13b/Usp10 | 28 |
| BP | GO:0045445 | myoblast differentiation | 19/1630 | 49/8933 | 5,8E+11 | 0.007547281011728317 | 0.0068701985082385095 | Dpf3/Smarca2/Pbrm1/Ddx5/Arid2/Brd7/Smarcd1/Actl6a/Smarcc2/Smarce1/Ddx17/Smarcb1/Arid1b/Smarcc1/Arid1a/Actl6b/Mapk14/Smarca4/Prickle1 | 19 |
| BP | GO:0046822 | regulation of nucleocytoplasmic transport | 30/1630 | 93/8933 | 7,7E+10 | 0.009826249274064882 | 0.008944715719390045 | Rbm4/Thoc2/Khdrbs1/Tardbp/Thoc5/Inpp4b/Dhx9/Trim28/Ctdspl2/Prkd1/Rbm22/Nolc1/Dmap1/Iws1/Anp32b/Cpsf6/Rbm10/Mapk14/Tpr/Nup62/Nup54/Nup153/Nedd4/Nup214/Mapk1/Akap8l/Xpo1/Prkaca/Hdac3/Nsun2 | 30 |
| CC | GO:0005681 | spliceosomal complex | 124/1630 | 166/8933 | 8,1E-43 | 7,94786E-40 | 7,23484E-41 | Prpf38a/Rbmxl1/Prpf38b/Sf3a2/Hnrnpa3/Hnrnpu/Cirbp/Zcchc8/Luc7l2/Hnrnph1/U2af2/Hnrnpc/Hnrnpr/Ptbp2/Snrpe/Srrm1/Plrg1/Snrnp70/Ybx1/U2af1/Tra2b/Snu13/Hnrnpa2b1/Srsf2/Tra2a/Snrpd1/Srsf1/Ddx39b/Prpf40a/Rbmx/Ddx5/Rbm3/Hnrnpk/Smu1/Luc7l3/Snrnp40/Hnrnpf/Usp39/Isy1/Snrpb/Magoh/Snrpg/Frg1/Xab2/Rbm8a/Hnrnpm/Api5/Snrpa/Cwc27/Snrpb2/Rbm22/Sf3b3/Sf3b4/Prpf19/Yju2/Dhx15/Eftud2/Crnkl1/Sf3a1/Snrnp200/Snrpa1/Eif4a3/Prpf8/Ncl/Sf3a3/Alyref/Sf3b1/Bud31/Cdc5l/Mtrex/Pnn/Cdc40/Snw1/Sf3b2/Sf1/Ddx41/Cwc22/Ik/Snrpd3/Rnf113a2/Sart1/Snrpd2/Ppil2/Ctnnbl1/Rbm17/Sugp1/Luc7l/Prpf31/Ppp1r8/Srrm2/Ddx23/Gpkow/Smndc1/Tfip11/Rbm5/Hnrnpa1/Aqr/Prpf4b/Prpf6/Bcas2/Prpf3/Dhx8/Lsm2/Prpf4/Cwf19l1/Zmat2/Lsm6/Cwc15/Dhx16/Ppie/Prpf40b/Phf5a/Lsm4/Dhx32/Prpf39/Syncrip/Lsm8/Syf2/Dhx38/Htatsf1/Raly/Rbm28/Upf1/Adar | 124 |

| | | | | | | | | | |
|---|---|---|---|---|---|---|---|---|---|
| CC | GO:0016604 | nuclear body | 199/1630 | 443/8933 | 1,9E-25 | 1,11358E-21 | 1,01367E-22 | Rbm4/Hp1bp3/Prdm8/Apex1/Epb41/Zbtb20/H2ax/Cbx5/Sf3a2/Stag1/Thoc6/Srsf6/Nop58/Hnrnpu/Fbll1/Ppp1r16b/Wtap/Thoc1/Tardbp/Zcchc8/Luc7l2/Ewsr1/Alkbh5/Rbm4b/U2af2/Fbl/Ints13/Srsf7/Champ1/Srrm1/Pcgf2/Plrg1/Rbm39/Srsf3/Chd3/Sfpq/Dhx9/Snrnp70/U2af1/Npm1/Ints7/Nono/Hnrnpa2b1/Srsf2/Mau2/Srsf1/Ddx39b/Prpf40a/Ddx5/Srrt/Rbbp6/Poldip3/Luc7l3/Snrnp40/Ppp4r3b/Bclaf1/Pnisr/Rbm8a/Hnrnpm/Srsf5/Api5/Atrx/Sarnp/Ppig/Srsf10/Snrpb2/Rnf2/Brd1/Nolc1/Rbm14/Ehmt1/Acin1/Pias1/Nxf1/Ppp1r10/Terf2/Cpsf6/Sympk/Wbp11/Prpf19/Cdk13/Dhx15/Zbtb18/Terf2ip/Eftud2/Rbm15b/Sf3a1/Rbm15/Srsf11/Ep400/Ddx42/Ythdc1/Snrpa1/Srsf4/Nudt21/Fyttd1/Sf3a3/Taf5l/Pip5k1a/Sf3b1/Rbm25/Rpa1/Cdc5l/Pacsin2/Zc3h14/Pnn/Cdc40/Snw1/Dkc1/Pspc1/Sf3b2/Suz12/Sf1/Rbm10/Ddx17/Cwc22/Tcf20/Ik/Chtop/Snrpd3/Parp1/Ncor2/Fto/Sart1/Virma/Sap130/Cd2bp2/Son/Sart3/Prpf31/Ppp1r8/Thrap3/Srrm2/Nelfe/Tfip11/Ddx39a/Banp/Cstf2/Prpf4b/Zc3h13/Rnf112/Prpf6/Pcna/Fam76a/Ube2i/Bcas2/Ylpm1/Sap18/Sltm/Sugp2/Prpf3/Pqbp1/Cdk9/Dhx8/Prpf4/Ddx46/Nup43/Stk19/Nup98/Cwc15/Mapk14/Nmnat1/Gtf2b/Toe1/Morf4l1/Gtf2e2/Nelfa/Gnl3/Ppie/Setd1a/Phf5a/Ddx1/Scaf11/Morc3/Ppp4r3a/Syf2/Mettl3/Osgep/Crebbp/Dcaf7/Angel2/Ubxn7/Ppih/Actn4/Cacnb4/Gcat/Ncam2/Hace1/Ro60/Akap8l/Adar/Atxn2l/Prkaca/Fmr1/Eif3e/Mtdh/Hikeshi/Adprs/Nbr1 | 199 |
| CC | GO:0000785 | chromatin | 178/1630 | 384/8933 | 2,2E-23 | 1,1973E-19 | 1,08989E-19 | Rfx3/Phf20l1/Bcl11b/H4c1/Msl2/Dpf3/Hp1bp3/H2ax/Cbx5/Hdac1/Baz1b/Hmga1/Stag1/Macroh2a1/Smarca2/Pogz/Gatad2b/Tardbp/Cbx1/Psip1/Hmgn1/Dek/Smad3/Cbx6/Phc2/Smarca5/Mta2/Pds5b/Exosc10/Brd4/Chd4/Smarcad1/Hdac2/Mbd3/Pbrm1/Mef2a/Mta3/Ddx21/Pcgf2/Chd3/Sfpq/Dhx9/Trim28/Mta1/Mecp2/Pelp1/Glyr1/Hnrnpa2b1/Srsf2/Rbbp7/Cbx8/Rsf1/Exosc4/Mau2/Kat2a/Rbmx/Mybbp1a/Hnrnpl/Hnrnpk/Cbx3/Trrap/Tbp/Polr2a/Rbbp4/Ppp4r3b/Prdm10/Rad21/Atrx/Wdr5/Arid2/Rnf2/Brd1/Sf3b3/Ccar2/Ehmt1/Ahctf1/Dmap1/Ppp1r10/Cpsf6/Taf4/Zbtb18/Yy1/Brd7/Hira/Macroh2a2/Bcl7a/Ep400/Smarcd1/Sin3b/Stag2/Actl6a/Smc3/Taf5l/Sin3a/Sf3b1/Smarcc2/Bud31/Enc1/Snw1/Smarce1/Creb1/Ash2l/Suz12/Phf14/Meaf6/Dpf1/Parp1/Exosc3/Ncor2/Hmgb2/Ssrp1/Kdm1a/Sap130/Zmynd8/Smarcb1/Pds5a/Kdm1b/Brms1l/Tasor/Ddx23/Nelfe/Cfdp1/Top2b/Ctr9/Banf1/Pcna/Sfr1/Arid1b/Smarcc1/Supt3/Set/Dnmt1/Nasp/Arid1a/Msl3/Yeats2/Wdr82/Aff4/Brd3/Actl6b/Nmnat1/Epc2/Exosc5/Morf4l1/Smarca4/Men1/Gabpa/Hcfc1/Taf9b/Setd1a/Bptf/Brd9/Wdr43/Rnf40/Yeats4/Rnf20/Rcc1/Hmgn5/Atf2/Tmpo/Ppp4r2/Tox4/Nedd4/Nsd3/Ncoa1/Hlcs/Crebbp/Mrgbp/Chd6/Trps1/Skic8/Ddx6/Akap8l/Upf1/Hdac3/Htra2/Tnks1bp1/Ruvbl1 | 178 |
| CC | GO:0120114 | Sm-like protein family complex | 53/1630 | 79/8933 | 1,2E-06 | 0,001917656 | 1,74562E-05 | Sf3a2/Luc7l2/Snrpe/Snrnp70/Snu13/Rbm42/Snrpd1/Ddx39b/Prpf40a/Luc7l3/Snrnp27/Snrnp40/Usp39/Snrpb/Snrpg/Snrpa/Snrpb2/Sf3b3/Nolc1/Sf3b4/Hexim1/Eftud2/Sf3a1/Snrnp200/Snrpa1/Prpf8/Sf3a3/Sf3b1/Sf3b2/Snrpd3/Sart1/Larp7/Cd2bp2/Sart3/Snrpd2/Luc7l/Prpf31/Ddx23/Prpf6/Mepce/Prpf3/Lsm2/Prpf4/Zmat2/Lsm6/Prpf40b/Phf5a/Lsm4/Prpf39/Lsm8/Htatsf1/Ppih/Fmr1 | 53 |
| CC | GO:0022626 | cytosolic ribosome | 54/1630 | 86/8933 | 5,3E-05 | 0,00750415 | 0,068309373 | Rps24/Rpl7l1/Rps26/Dhx29/Rpl39/Rpl36/Rpl7a/Rps8/Rpl24/Rpl29/Rpl21/Rpl34/Rps23/Rpl12/Rpl18/Rpl3/Rpl27a/Rpl8/Rpl6/Rpl31/Rpl18a/Rps4x/Rps18/Rps25/Rpl14/Rpl10/Rps13/Rpl22/Rpl15/Rpl13/Rpl37a/Rpl4/Rps2/Rpl17/Rpl5/Rpl10a/Rps9/Rpl23/Rpl27/Rps15a/Rps20/Rps7/Rps14/Rpl11/Rps16/Rps10/Rpsa/Rpl30/Rpl7/Rps19/Rplp0/Rps3/Rps27/Rps11 | 54 |
| CC | GO:0070603 | SWI/SNF superfamily-type complex | 45/1630 | 68/8933 | 0,03347 | 3,882947419 | 3,534600014 | Bcl11b/Dpf3/Hdac1/Baz1b/Smarca2/Gatad2b/Dek/Smarca5/Mta2/Chd4/Hdac2/Mbd3/Pbrm1/Mta3/Ddx21/Chd3/Mta1/Rbbp7/Rsf1/Mybbp1a/Trrap/Rbbp4/Arid2/Dmap1/Yy1/Brd7/Bcl7a/Ep400/Smarcd1/Actl6a/Sf3b1/Smarcc2/Smarce1/Suz12/Dpf1/Smarcb1/Cfdp1/Arid1b/Smarcc1/Arid1a/Actl6b/Smarca4/Bptf/Brd9/Ruvbl1 | 45 |
| CC | GO:0032993 | protein-DNA complex | 45/1630 | 78/8933 | 6,69237 | 618,6259501 | 563,1277109 | H4c1/Hp1bp3/H2ax/Macroh2a1/Nfyc/Top1/Chd4/Plrg1/Npm1/Glyr1/Gtf2a1/Mcm6/Hnrnpk/Trrap/Tbp/Gtf2f2/Dmap1/Terf2/Prpf19/Wdr18/Terf2ip/Macroh2a2/Ep400/Gtf2h3/Actl6a/Kdm5a/Rpa1/Cdc5l/Suz12/Meaf6/Parp1/Kdm1b/Nfyb/Bcas2/Stn1/Ercc3/Ctc1/Gtf2b/Epc2/Morf4l1/Pold1/Yeats4/Smarcal1/Mrgbp/Ruvbl1 | 45 |
| CC | GO:0034708 | methyltransferase complex | 43/1630 | 76/8933 | 69,3406 | 5662,451489 | 5154,461019 | Prdm8/Cbx5/Wtap/Tex10/Hdac2/Snrpe/Kmt2a/Rbbp5/Kdm6a/Pelp1/Rbbp7/Snrpd1/Rbbp4/Snrpb/Snrpg/Senp3/Erh/Wdr5/Rnf2/Taf4/Rbm15b/Rbm15/Trmt6/Mettl1/Ash2l/Suz12/Snrpd3/Las1l/Virma/Trmt61a/Snrpd2/Prpf31/Zc3h13/Dpy30/Kmt2d/Wdr82/Men1/Rnmt/Hcfc1/Setd1a/Mettl3/Prmt1/Ruvbl1 | 43 |
| CC | GO:1904949 | ATPase complex | 48/1630 | 95/8933 | 763,697 | 513412,7836 | 46735,34396 | Bcl11b/Dpf3/Hdac1/Baz1b/Smarca2/Gatad2b/Dek/Smarca5/Tmem199/Mta2/Chd4/Hdac2/Mbd3/Pbrm1/Mta3/Ddx21/Chd3/Mta1/Rbbp7/Rsf1/Mybbp1a/Trrap/Ccdc115/Rbbp4/Arid2/Dmap1/Yy1/Brd7/Bcl7a/Ep400/Smarcd1/Actl6a/Sf3b1/Smarcc2/Smarce1/Suz12/Dpf1/Smarcb1/Cfdp1/Arid1b/Smarcc1/Arid1a/Actl6b/Smarca4/Bptf/Brd9/Ruvbl1/Atp6ap2 | 48 |

| | | | | | | | | | |
|---|---|---|---|---|---|---|---|---|---|
| CC | GO:0044815 | DNA packaging complex | 26/1630 | 40/8933 | 84920,9 | 43309634,78 | 3942423,607 | H4c1/Hp1bp3/H2ax/Stag1/Macroh2a1/Glyr1/Smc1a/Trrap/Rad21/Dmap1/Terf2/Terf2ip/Macroh2a2/Ep400/Stag2/Actl6a/Smc3/Meaf6/Kdm1b/Stn1/Ctc1/Epc2/Morf4l1/Yeats4/Mrgbp/Ruvbl1 | 26 |
| CC | GO:0005667 | transcription regulator complex | 83/1630 | 233/8933 | 1350501 | 6713918,015 | 6111598,264 | Rfx3/Apex1/Mideas/Cbx5/Hdac1/Hmga1/Nfyc/Gatad2b/Hnrnpu/Hmgb1/Gtf2f1/Smad3/Mta2/Hnrnpab/Chd4/Tbpl1/Hdac2/Mbd3/Adnp/Mef2a/Mta3/Chd3/Sfpq/Trim28/Mta1/Nono/Tle1/Rbbp7/Gtf2a1/Kat2a/Cbx3/Trrap/Tbp/Rbbp4/Hdgf/Prdm10/Gtf2f2/Arid2/Spen/Med24/Rbm14/Taf4/Yy1/Gtf2h1/Gtf3c2/Gtf2h3/Taf5l/Sin3a/Creb1/Gtf3c4/Dcp1a/Rcor3/Parp1/Ncor2/Thap11/Tle4/Nfyb/Gtf3c3/Foxo3/Tcea1/Med25/Gtf3c5/Supt3/Sap18/Med23/Cdk7/Ercc3/Sub1/Gtf2b/Gtf2e2/Men1/Taf9b/Riox2/Gtf3c1/Ercc2/Atf2/Ncoa1/Hivep2/Pus1/Crebbp/Med15/Tbl1xr1/Hdac3 | 83 |
| CC | GO:0030684 | preribosome | 31/1630 | 58/8933 | 1500577 | 61223538,95 | 55731046,1 | Riox1/Nop58/Fbll1/Nop56/Fbl/Bop1/Mak16/Rrs1/Snu13/Pes1/Emg1/Krr1/Wdr3/Pdcd11/Bms1/Utp6/Rsl1d1/Heatr1/Las1l/Utp18/Tbl3/Wdr36/Nop14/Rrp9/Noc4l/Bysl/Mdn1/Utp14a/Wdr12/Rrp7a/Rps7 | 31 |
| CC | GO:0000428 | DNA-directed RNA polymerase complex | 33/1630 | 69/8933 | 1,7E+08 | 589640313,4 | 536742436,8 | Rprd1a/Gtf2f1/Polr2g/Tbpl1/Gtf2a1/Rprd1b/Kat2a/Trrap/Tbp/Polr2a/Gtf2f2/Polr2b/Taf4/Paf1/Polr2i/Gtf2h1/Gtf2h3/Taf5l/Polr2c/Rprd2/Cdc73/Tcea1/Ctr9/Supt3/Cdk7/Ercc3/Polr2e/Gtf2b/Gtf2e2/Taf9b/Ercc2/Rtf1/Skic8 | 33 |
| CC | GO:0005844 | polysome | 33/1630 | 71/8933 | 4,2E+07 | 1318671000 | 12003702425 | Dhx9/Fus/Larp7/Rps26/Ago1/Fubp3/Rpl39/Rpl36/Rpl7a/Rpl24/Btf3/Rpl29/Rps23/Rpl18/Rpl8/Rpl6/Rpl31/Rpl18a/Rps4x/Rps25/Ago2/Rpl17/Rpl10a/Rpl11/Fmr1/Rpl30/Rpl7/Upf2/Rps3/Eif3h/Fxr1/Msi1/Naa35 | 33 |
| CC | GO:0005643 | nuclear pore | 31/1630 | 65/8933 | 5,1E+08 | 15877759001 | 14453331741 | Ahctf1/Nxf1/Nup210/Nup35/Ube2i/Nup107/Nup188/Nup43/Nup88/Nup98/Nup160/Nup155/Tpr/Nup133/Ranbp2/Nup205/Pom121/Nup85/Nup93/Nup37/Nup62/Nup54/Nup153/Ranbp3/Seh1l/Nup214/Pcid2/Mad2l1/Nup50/Rae1/Aaas | 31 |
| CC | GO:0000974 | Prp19 complex | 13/1630 | 17/8933 | 2,7E+09 | 7326911554 | 6669598860 | U2af2/Plrg1/Polr2a/Isy1/Xab2/Rbm22/Prpf19/Crnkl1/Cdc5l/Ctnnbl1/Bcas2/Cwc15/Syf2 | 13 |
| CC | GO:1905354 | exoribonuclease complex | 14/1630 | 20/8933 | 5,5E+08 | 13656887809 | 1,24317E+11 | Exosc10/Exosc4/Exosc2/Exosc1/Exosc8/Mtrex/Exosc7/Exosc3/Las1l/Dis3/Exosc5/Nvl/Supv3l1/Gtpbp1 | 14 |
| CC | GO:0005732 | sno(s)RNA-containing ribonucleoprotein complex | 12/1630 | 16/8933 | 1,2E+10 | 2,68797E+11 | 24468243053 | Nop58/Fbll1/Rpp30/Nop56/Fbl/Nhp2/Snu13/Nolc1/Dkc1/Gar1/Lsm6/Rrp9 | 12 |
| CC | GO:0000178 | exosome (RNase complex) | 13/1630 | 19/8933 | 2,2E+10 | 46133506184 | 41994772008 | Exosc10/Exosc4/Exosc2/Exosc1/Exosc8/Mtrex/Exosc7/Exosc3/Dis3/Exosc5/Nvl/Supv3l1/Gtpbp1 | 13 |
| CC | GO:0043073 | germ cell nucleus | 21/1630 | 43/8933 | 4,6E+10 | 92666909236 | 84353565291 | H2ax/Hmga1/Hmgn1/Top1/Celf1/Tbpl1/Tbp/Terf2/Taf4/Terf2ip/Rpa1/Smarcb1/Hspa2/Pcna/Smarcc1/Dnmt1/Cdk7/Dazap1/Gtf2b/Smarca4/Tsn | 21 |
| CC | GO:0031248 | protein acetyltransferase complex | 27/1630 | 67/8933 | 2E+10 | 3,47291E+11 | 0.00031613466225740487 | Phf20l1/Msl2/Kat2a/Trrap/Wdr5/Brd1/Sf3b3/Dmap1/Taf4/Ep400/Actl6a/Taf5l/Meaf6/Supt3/Msl3/Yeats2/Actl6b/Epc2/Morf4l1/Hcfc1/Taf9b/Yeats4/Atf2/Crebbp/Mrgbp/Ruvbl1/Naa35 | 27 |
| CC | GO:0005852 | eukaryotic translation initiation factor 3 complex | 9/1630 | 14/8933 | 1,8E+10 | 0.002647958443039223 | 0.0024104045042149156 | Eif3a/Eif3f/Eif3b/Eif3l/Eif3i/Eif3e/Eif3h/Eif3c/Eif3d | 9 |
| CC | GO:0032806 | carboxy-terminal domain protein kinase complex | 11/1630 | 20/8933 | 2,4E+11 | 0.0034050342030285652 | 0.0030995614004295465 | Brd4/Gtf2f2/Cdk13/Gtf2h1/Gtf2h3/Snw1/Cdk9/Cdk7/Ercc3/Ccnk/Ercc2 | 11 |
| MF | GO:0003729 | mRNA binding | 150/1630 | 248/8933 | 6,3E-36 | 5,28646E-33 | 4,8122E-33 | Khdrbs2/Ptbp3/Rbm4/Shmt1/Thoc2/Khdrbs1/Hnrnpa3/Srsf6/Hnrnpu/Tardbp/Cirbp/Luc7l2/Elavl1/Hnrnpa0/Thoc5/Rbm4b/Celf1/Hnrnpc/Hnrnpr/Ptbp2/Hnrnpd/Khsrp/Srsf3/Dhx9/Fus/Snrnp70/Ybx1/Tia1/Mecp2/Tra2b/Rnps1/Nhp2/Hnrnpa2b1/Rbm42/Exosc4/Srsf1/Rbmx/Hnrnpl/Ddx5/Hnrnpk/Ssb/Poldip3/Luc7l3/Fubp1/Myef2/Ilf3/Ncbp1/Rbm8a/Hnrnpm/Srsf5/Spen/Rbm14/Nxf1/Cpsf6/Rbm15b/Rbm15/Cpsf7/Rbfox3/Ythdc1/Srsf4/Eif4a3/Nudt21/Fyttd1/Ncl/Alyref/Exosc8/Sf3b1/Rbm25/Cdc40/Sf1/Dcp1a/Ddx17/Rbfox1/Scaf8/Chtop/Rsl1d1/Cpsf1/Larp7/Pcbp3/Cstf3/Rps26/Luc7l/Cpeb4/Ppp1r8/Srrm3/Srrm2/Lsm14b/Rbm5/Nova2/Hnrnpa1/Celf2/Cstf2/Aqr/Ncbp2/Fubp3/Pcf11/Dazap1/Nup98/Tpr/Hnrnpll/Srbd1/Pcbp2/Gnl3/Ppie/Rpl24/Xpo5/Rnf40/Rnf20/Tial1/Cryz/Syncrip/Dcp1b/Rbfox2/Tsn/Pum3/Celf4/Rpl6/Pum1/Lsm14a/Mettl3/G3bp1/Rps13/Ago2/Ythdf1/Angel2/Cstf2t/Serbp1/Rps2/Rpl5/Elavl3/Slc4a1ap/Eif3a/Edc3/Rc3h2/Rps7/Rps14/Ddx6/Shmt2/Fmr1/Rpl30/Rpl7/Csdc2/Rps3/G3bp2/Copa/Mrps11/Eif3d/Fxr1/Msi1/Mrpl13 | 150 |

| | | | | | | | | | |
|---|---|---|---|---|---|---|---|---|---|
| MF | GO:0003682 | chromatin binding | 143/1630 | 325/8933 | 6,9E-13 | 2,0419E-09 | 1,85872E-09 | H1f10/Bhlhe22/Nfix/Hp1bp3/Prdm8/Apex1/Maz/Nfia/Rbmxl1/Cbx5/Hdac1/Hmga1/Stag1/Macroh2a1/Smarca2/Samd1/Hnrnpu/Gtf2f1/Hdgfl3/Cbx1/Psip1/Hmgn1/Smad3/Cbx6/Smarca5/Top1/Mta2/Brd4/Chd4/Smarcad1/Hdac2/Hnrnpd/Mbd3/Pbrm1/Adnp/Mef2a/Mta3/Pcgf2/Kmt2a/Chd3/Sfpq/Dhx9/Fus/Trim28/Ybx1/Ubtf/Npm1/Mta1/Kdm6a/Mecp2/Zc3h4/Nono/Tle1/Pelp1/Glyr1/Cbx8/Ncoa5/Kat2a/Rbmx/Ddx5/Cbx3/Smc1a/Polr2a/Rad21/Safb/Atrx/Sarnp/Rnf2/Polr2b/Yy1/Paf1/Hira/Macroh2a2/Gtf2h1/Ep400/Smarcd1/Sin3b/Nudt21/Stag2/Actl6a/Kdm5a/Smc3/Sin3a/Sf3b1/Smarcc2/Rpa1/Suz12/Ddx17/Parp1/Ncor2/Ssrp1/Kdm1a/Sbno1/Npm3/Kdm1b/Tle4/Tasor/Ercc4/Nelfe/Top2b/Foxo3/Pcna/Arid1b/Smarcc1/Trim33/Set/Dnmt1/Cdk9/Arid1a/Ercc3/Wdr82/Brd3/Pwwp3a/Nup98/Actl6b/Gtf2b/Tpr/Morf4l1/Nelfa/Smarca4/Men1/Gabpa/Hcfc1/Pold1/Msh2/Rnf20/Ddx1/Nup153/Rcc1/Hmgn5/Atf2/Ercc1/Tox4/Ncoa1/Pus1/Crebbp/Chd6/Actn4/Shmt2/Upf1/Fmr1/Hdac3/Lemd3 | 143 |
| MF | GO:0043565 | sequence-specific DNA binding | 142/1630 | 384/8933 | 0,00054 | 0,067651854 | 0,061582664 | Prox1/Rfx3/Nfib/Egr3/Bhlhe22/Bcl11b/Nfix/Apex1/Maz/Nfia/Zbtb20/Rbmxl1/Hdac1/Mef2d/Usf2/Zfp871/Hmga1/Macroh2a1/Smarca2/Zfp512b/Nfyc/Gatad2b/Hnrnpu/Hmgb1/Tardbp/Smad3/Top1/Foxk1/Mta2/Hnrnpab/Brd4/Tbpl1/Hdac2/Cpsf4/Hnrnpd/Mbd3/Adnp/Mef2a/Mta3/Kmt2a/Chd3/Sfpq/Dhx9/Rbbp5/Ybx1/Ubtf/Npm1/Mta1/Kdm6a/Mecp2/Hnrnpa2b1/Gtf2a1/Rbmx/Mybbp1a/Hnrnpl/Neurod2/Hnrnpk/Cbx3/Tbp/Fubp1/Polr2a/Rbbp4/Hdgf/Myef2/Ccar1/Erh/Safb/Gmeb1/Nr2f1/Emx1/Terf2/Taf4/Zbtb18/Yy1/Terf2ip/Brd7/Macroh2a2/Ncl/Kdm5a/Rpa1/Cdc5l/Creb1/Ash2l/Suz12/Nr2e1/Tcf20/Chtop/Dpf1/Ncor2/Hmgb2/Thap11/Kdm1a/Smarcb1/Nfyb/Ago1/Thrap3/Zscan26/Wiz/Foxo3/Lmnb1/Nabp2/Hnrnpa1/Gtf3c5/Xrn2/Msh3/Stn1/Dnmt1/Sltm/Kmt2d/Kdm5b/Cdk9/Ctc1/Sub1/Zhx3/Gtf2b/Smarca4/Men1/Gabpa/Hcfc1/Zbtb11/Bptf/Msh2/Smyd3/Zkscan16/Zgpat/Tsnax/Tsn/Atf2/Ncoa1/Hivep2/Crebbp/Ago2/Prmt1/Actn4/Trps1/Nacc1/Mrtfa/Tbl1xr1/Upf1/Upf2/Pura/Rps3 | 142 |
| MF | GO:0140640 | catalytic activity, acting on a nucleic acid | 126/1630 | 335/8933 | 0,12267 | 13,43943417 | 1,223375418 | Apex1/Dcps/Hmga1/Trmt10b/Smarca2/Fbll1/Rpp30/Alkbh5/Smarca5/Top1/Fbl/Exosc10/Chd4/Smarcad1/Ddx21/Chd3/Dhx9/Pcif1/Dbr1/Rsf1/Mau2/Ddx39b/Mcm6/Ddx5/Trmt2a/Polr2a/Rbbp4/Isy1/Atrx/Emg1/Terf2/Polr2b/Dhx15/Polr2i/Ep400/Ddx42/Snrnp200/Eif4a3/Smc3/Mtrex/Ints11/Polr2c/Mettl1/Dkc1/Polb/Dcp1a/Ddx41/Ddx17/Fto/Trmt61a/Dus3l/Cmtr1/Ercc4/Ddx23/Top2b/Ddx51/Ddx39a/Ddx50/Xrn2/Aqr/Dis3/Dhx29/Pcna/Msh3/Dnmt1/Mepce/Dhx8/Cdk7/Arid1a/Dhx40/Ddx46/Ercc3/Cnot6l/Dhx16/Lig3/Tatdn1/Polr2e/Toe1/Smarca4/Rnmt/Mettl16/Wrnip1/Trub1/Pold1/Bptf/Msh2/Ddx1/Dhx32/Cpsf3/Rfc5/Tsnax/Dcp1b/Ercc2/Smarcal1/Exd2/Dhx38/Ercc1/Nop2/Rtcb/Mettl3/G3bp1/Pus1/Ago2/Eif4a1/Angel2/Chd6/Thumpd3/Rngtt/Polg2/Elac2/Supv3l1/Ddx6/Cnot1/Upf1/Wars2/Dtd1/Rps3/Ptrh2/Nsun2/Trmu/Eif4a2/Trnt1/Mrpl58/Ruvbl1/Rars2/Rexo2 | 126 |
| MF | GO:0003735 | structural constituent of ribosome | 68/1630 | 142/8933 | 4,25452 | 412,6185516 | 37,56016707 | Rps24/Rpl7l1/Rps26/Rpl39/Mrpl57/Srbd1/Rpl36/Rpl7a/Rps8/Rpl24/Rpl29/Rpl21/Rpl34/Rps23/Rpl12/Rpl18/Rpl3/Rpl27a/Rpl8/Rpl6/Rpl31/Rpl18a/Rps4x/Rps18/Rps25/Rpl14/Rpl10/Rps13/Rpl22/Rpl15/Rpl13/Rpl37a/Rpl4/Rps2/Rpl17/Rpl5/Rpl10a/Rps9/Mrpl37/Rpl23/Rpl27/Rps15a/Rps20/Rps7/Rps14/Mrpl15/Rpl11/Rps16/Rps10/Rpsa/Rpl30/Rpl7/Mrpl9/Rps19/Rplp0/Rps3/Mrps35/Mrpl19/Mrps9/Rps27/Mrpl41/Rps11/Mrpl39/Mrps22/Mrps11/Mrpl3/Mrpl13/Mrps34 | 68 |
| MF | GO:0001098 | basal transcription machinery binding | 32/1630 | 47/8933 | 75,8803 | 5985,434386 | 54484,68439 | Rprd1a/Nop58/Hnrnpu/Gtf2f1/Fbl/Brd4/Dhx9/Pcif1/Gtf2a1/Rprd1b/Tbp/Nolc1/Ccar2/Paf1/Scaf8/Ago1/Rprd2/Ercc4/Cdc73/Scaf1/Ctr9/Pcf11/Gtf2b/Gtf2e2/Smyd3/Wdr43/Elp2/Rtf1/Ercc1/Crebbp/Ago2/Ruvbl1 | 32 |
| MF | GO:0001099 | basal RNA polymerase II transcription machinery binding | 32/1630 | 47/8933 | 75,8803 | 5985,434386 | 54484,68439 | Rprd1a/Nop58/Hnrnpu/Gtf2f1/Fbl/Brd4/Dhx9/Pcif1/Gtf2a1/Rprd1b/Tbp/Nolc1/Ccar2/Paf1/Scaf8/Ago1/Rprd2/Ercc4/Cdc73/Scaf1/Ctr9/Pcf11/Gtf2b/Gtf2e2/Smyd3/Wdr43/Elp2/Rtf1/Ercc1/Crebbp/Ago2/Ruvbl1 | 32 |
| MF | GO:0042393 | histone binding | 66/1630 | 151/8933 | 2823,56 | 206224,6166 | 18772,37711 | Anp32a/H2ax/Cbx5/Vrk1/Baz1b/Ptma/Smarca2/Cbx1/Dek/Cbx6/Smarca5/Brd4/Chd4/Hdgfl2/Kmt2a/Chd3/Rbbp5/Npm1/Ptms/Glyr1/Rbbp7/Cbx8/Rsf1/Cbx3/Rbbp4/Phip/Atrx/Wdr5/Brd1/Anp32b/Brd7/Hira/Zmynd11/Phf2/Ncl/Kdm5a/Smarcc2/Suz12/Phf14/Spin1/Ssrp1/Zmynd8/Sbno1/Sart3/Npm3/Kdm1b/Smarcc1/Set/Nasp/Dnajc9/Kmt2d/Kdm5b/Msl3/Yeats2/Brd3/Smarca4/Bptf/Brd9/Yeats4/Rnf20/Morc3/Pwp1/Rcc1/Chd6/Tbl1xr1/Fmr1 | 66 |
| MF | GO:0003727 | single-stranded RNA binding | 39/1630 | 69/8933 | 10664,2 | 70381,05395 | 64067,02111 | Khdrbs2/Khdrbs1/Hnrnpu/Hmgb1/Polr2g/Cirbp/Cbx6/U2af2/Hnrnpc/Exosc10/Dhx9/Rbm7/Cbx8/Rbmx/Ssb/Hnrnpf/Ilf3/Eif4a3/Zc3h14/Rbm10/Pabpn1/Hnrnpdl/Ago1/Hnrnpa1/Aqr/Zfr/Dazap1/Ppie/Ddx1/Syncrip/Lsm14a/Pus1/Ago2/Rps7/Fmr1/Strbp/Zfr2/Fxr1/Msi1 | 39 |

| | | | | | | | | | |
|---|---|---|---|---|---|---|---|---|---|
| MF | GO:0001067 | transcription regulatory region nucleic acid binding | 106/1630 | 308/8933 | 4124,47 | 254170,7529 | 231368,5584 | Prox1/Rfx3/Nfib/Egr3/Bcl11b/Nfix/Maz/Nfia/Zbtb20/Rbmxl1/Hdac1/Mef2d/Usf2/Zfp871/Hmga1/Macroh2a1/Smarca2/Zfp512b/Nfyc/Hnrnpu/Hmgb1/Tardbp/Smad3/Top1/Foxk1/Brd4/Tbpl1/Hdac2/Adnp/Mef2a/Chd3/Sfpq/Dhx9/Rbbp5/Ubtf/Npm1/Mta1/Kdm6a/Hnrnpa2b1/Gtf2a1/Rbmx/Mybbp1a/Hnrnpl/Neurod2/Hnrnpk/Cbx3/Tbp/Fubp1/Polr2a/Rbbp4/Hdgf/Myef2/Ccar1/Safb/Gmeb1/Nr2f1/Emx1/Taf4/Yy1/Brd7/Macroh2a2/Kdm5a/Cdc5l/Creb1/Ash2l/Suz12/Nr2e1/Tcf20/Hmgb2/Thap11/Kdm1a/Smarcb1/Nfyb/Ago1/Thrap3/Zscan26/Wiz/Foxo3/Gtf3c5/Xrn2/Kmt2d/Cdk9/Sub1/Zhx3/Gtf2b/Smarca4/Men1/Gabpa/Hcfc1/Zbtb11/Bptf/Smyd3/Zkscan16/Zgpat/Atf2/Ncoa1/Hivep2/Crebbp/Ago2/Actn4/Trps1/Nacc1/Mrtfa/Tbl1xr1/Pura/Rps3 | 106 |
| MF | GO:0017069 | snRNA binding | 26/1630 | 38/8933 | 144849 | 835731,9409 | 760756,6652 | Hnrnpu/Ddx21/Snrnp70/Rbm7/Snu13/Ddx39b/Snrpa/Snrpb2/Rbm22/Sf3b3/Hexim1/Eftud2/Snrpa1/Prpf8/Snrpd3/Larp7/Sart3/Prpf31/Ncbp2/Mepce/Cdk9/Prpf4/Toe1/Mettl16/Lsm4/Ro60 | 26 |
| MF | GO:0140098 | catalytic activity, acting on RNA | 86/1630 | 238/8933 | 262685 | 14523760,67 | 13220803,46 | Apex1/Dcps/Trmt10b/Fbll1/Rpp30/Alkbh5/Fbl/Exosc10/Ddx21/Dhx9/Pcif1/Dbr1/Ddx39b/Ddx5/Trmt2a/Polr2a/Isy1/Emg1/Polr2b/Dhx15/Polr2i/Ddx42/Snrnp200/Eif4a3/Mtrex/Ints11/Polr2c/Mettl1/Dcp1a/Ddx41/Ddx17/Fto/Trmt61a/Dus3l/Cmtr1/Ddx23/Ddx51/Ddx39a/Ddx50/Xrn2/Aqr/Dis3/Dhx29/Mepce/Dhx8/Dhx40/Ddx46/Cnot6l/Dhx16/Polr2e/Toe1/Rnmt/Mettl16/Trub1/Ddx1/Dhx32/Cpsf3/Tsnax/Dcp1b/Exd2/Dhx38/Nop2/Rtcb/Mettl3/G3bp1/Pus1/Ago2/Eif4a1/Angel2/Thumpd3/Rngtt/Elac2/Supv3l1/Ddx6/Cnot1/Upf1/Wars2/Dtd1/Ptrh2/Nsun2/Trmu/Eif4a2/Trnt1/Mrpl58/Rars2/Rexo2 | 86 |
| MF | GO:0000993 | RNA polymerase II complex binding | 21/1630 | 28/8933 | 86697,8 | 43837965,04 | 39905168,71 | Rprd1a/Hnrnpu/Brd4/Dhx9/Pcif1/Rprd1b/Ccar2/Paf1/Scaf8/Ago1/Rprd2/Cdc73/Scaf1/Ctr9/Pcf11/Gtf2b/Smyd3/Wdr43/Elp2/Rtf1/Ago2 | 21 |
| MF | GO:0004386 | helicase activity | 45/1630 | 100/8933 | 557387 | 244259152,1 | 22234614,82 | Smarca2/Smarca5/Chd4/Smarcad1/Ddx21/Dhx9/Ddx39b/Mcm6/Ddx5/Atrx/Dhx15/Ep400/Ddx42/Snrnp200/Eif4a3/Mtrex/Ddx41/Ddx17/Ddx23/Ddx51/Ddx39a/Ddx50/Aqr/Dhx29/Dhx8/Dhx40/Ddx46/Ercc3/Dhx16/Smarca4/Wrnip1/Ddx1/Dhx32/Rfc5/Ercc2/Smarcal1/Dhx38/G3bp1/Eif4a1/Chd6/Supv3l1/Ddx6/Upf1/Eif4a2/Ruvbl1 | 45 |
| MF | GO:0140110 | transcription regulator activity | 137/1630 | 462/8933 | 579241 | 251969963,9 | 22936520,68 | Prox1/Rfx3/Nfib/Egr3/Bhlhe22/Bcl11b/Nfix/Prdm8/Apex1/Maz/Nfia/Mideas/Zbtb20/Hdac1/Mef2d/Usf2/Hmga1/Smarca2/Zfp512b/Nfyc/Hnrnpu/Hmgb1/Hdgfl3/Psip1/Smad3/Ewsr1/Gtf2i/Foxk1/Mta2/Brd4/Adnp/Mef2a/Mta3/Hdgfl2/Dhx9/Fus/Zhx2/Trim28/Npm1/Mta1/Mecp2/Tle1/Kat2a/Mybbp1a/Ddx5/Neurod2/Trrap/Ahdc1/Hdgf/Bclaf1/Myef2/Med13l/Ccar1/Gmeb1/Nr2f1/Spen/Emx1/Med24/Rbm14/Pias1/Dmap1/Zbtb18/Yy1/Brd7/Zmynd11/Phf2/Smarcd1/Sin3b/Rybp/Kdm5a/Taf5l/Sin3a/Bud31/Cdc5l/Snw1/Creb1/Ddx17/Rcor3/Nr2e1/Tcf20/Tcerg1/Ncor2/Hmgb2/Thap11/Kdm1a/Zmynd8/Smarcb1/Pcbp3/Tle4/Nfyb/Thrap3/Zscan26/Wiz/Foxo3/Prpf6/Sfr1/Supt3/Ylpm1/Sap18/Kmt2d/Kdm5b/Nup98/Sub1/Zhx3/Mrtfb/Smarca4/Men1/Gabpa/Hcfc1/Riox2/Zbtb11/Wdr43/Rnf20/Ddx1/Zgpat/Rbfox2/Atf2/Ncoa1/Hivep2/Irf2bpl/Pus1/Crebbp/Pa2g4/Actn2/Actn4/Trps1/Raly/Med15/Nacc1/Mrtfa/Actn1/Tbl1xr1/Hdac3/Pura/Dcc/Mtdh/Ruvbl1 | 137 |
| MF | GO:0036002 | pre-mRNA binding | 21/1630 | 30/8933 | 718417 | 307982299,1 | 280352557,2 | Rbm4/Srsf6/Hnrnpu/Tardbp/U2af2/Celf1/U2af1/Rbm7/Tra2b/Hnrnpa2b1/Srsf2/Hnrnpl/Ddx5/Hnrnpk/Rbm22/Prpf8/Sf1/Hnrnpa1/Celf2/Prpf39/Celf4 | 21 |
| MF | GO:0140097 | catalytic activity, acting on DNA | 49/1630 | 116/8933 | 139927 | 57486488,37 | 52329254,21 | Apex1/Hmga1/Smarca2/Smarca5/Top1/Chd4/Smarcad1/Chd3/Dhx9/Rsf1/Mau2/Mcm6/Rbbp4/Atrx/Terf2/Ep400/Smc3/Dkc1/Polb/Fto/Ercc4/Top2b/Pcna/Msh3/Dnmt1/Cdk7/Arid1a/Ercc3/Lig3/Tatdn1/Smarca4/Wrnip1/Pold1/Bptf/Msh2/Ddx1/Rfc5/Ercc2/Smarcal1/Exd2/Ercc1/G3bp1/Chd6/Polg2/Supv3l1/Upf1/Rps3/Ruvbl1/Rexo2 | 49 |
| MF | GO:0003712 | transcription coregulator activity | 88/1630 | 264/8933 | 1826327 | 74003770,45 | 6736473,607 | Prdm8/Apex1/Mideas/Hdac1/Hmga1/Smarca2/Hnrnpu/Hmgb1/Hdgfl3/Psip1/Ewsr1/Mta2/Brd4/Mta3/Hdgfl2/Dhx9/Fus/Trim28/Npm1/Mta1/Mecp2/Tle1/Kat2a/Mybbp1a/Ddx5/Trrap/Hdgf/Bclaf1/Med13l/Ccar1/Spen/Med24/Rbm14/Pias1/Dmap1/Brd7/Zmynd11/Phf2/Smarcd1/Sin3b/Rybp/Kdm5a/Taf5l/Sin3a/Bud31/Snw1/Ddx17/Rcor3/Tcf20/Tcerg1/Ncor2/Hmgb2/Kdm1a/Zmynd8/Smarcb1/Tle4/Thrap3/Prpf6/Sfr1/Supt3/Sap18/Kmt2d/Kdm5b/Nup98/Sub1/Mrtfb/Smarca4/Hcfc1/Riox2/Rnf20/Ddx1/Rbfox2/Ncoa1/Irf2bpl/Pus1/Crebbp/Pa2g4/Actn2/Actn4/Raly/Med15/Mrtfa/Actn1/Tbl1xr1/Hdac3/Dcc/Mtdh/Ruvbl1 | 88 |

| | | | | | | | | | |
|---|---|---|---|---|---|---|---|---|---|
| MF | GO:0008134 | transcription factor binding | 114/1630 | 376/8933 | 4363407 | 1665414076 | 1516006266 | Prox1/Apex1/Nfia/Cbx5/Hdac1/Mef2d/Usf2/Ptma/Hmga1/Nop58/Hnrnpu/Hmgb1/Gtf2f1/Psip1/Smad3/Gtf2i/Mta2/Fbl/Chd4/Hdac2/Mef2a/Dhx9/Fus/Npm1/Mta1/Mecp2/Tle1/Nek6/Gtf2a1/Kat2a/Mybbp1a/Ddx5/Cbx3/Tbp/Hnrnpf/Hdgf/Arid2/Spen/Nolc1/Med24/Ehmt1/Pias1/Dmap1/Taf4/Yy1/Brd7/Hira/Gtf2h1/Ncl/Sin3a/Bud31/Cdc5l/Snw1/Smarce1/Creb1/Mapk3/Suz12/Dcp1a/Pdcd11/Parp1/Tcerg1/Cggbp1/Ncor2/Hmgb2/Kdm1a/Smarcb1/Uba2/Kdm1b/Tle4/Nfyb/Thrap3/Ercc4/Wiz/Foxo3/Med25/Prpf6/Pcna/Ube2i/Dnmt1/Sap18/Cdk9/Arid1a/Yeats2/Grm1/Mapk14/Gtf2b/Gtf2e2/Smarca4/Hcfc1/Setd1a/Bptf/Paxbp1/Ptprn/Rbfox2/Atf2/Ercc1/Ncoa1/Crebbp/Chd6/Ubxn7/Actn4/Sri/Keap1/Dcaf1/Rpl23/Cnot1/Ppp1r13b/Xpo1/Hdac3/Pura/Rps3/Mtdh/Usp11/Ruvbl1 | 114 |
| MF | GO:0008186 | ATP-dependent activity, acting on RNA | 30/1630 | 58/8933 | 7675238 | 271896451,2 | 2475040469 | Ddx21/Dhx9/Ddx39b/Ddx5/Dhx15/Ddx42/Snrnp200/Eif4a3/Mtrex/Ddx41/Ddx17/Ddx23/Ddx51/Ddx39a/Ddx50/Aqr/Dhx29/Dhx8/Dhx40/Ddx46/Dhx16/Ddx1/Dhx32/Dhx38/G3bp1/Eif4a1/Supv3l1/Ddx6/Upf1/Eif4a2 | 30 |
| MF | GO:0031490 | chromatin DNA binding | 26/1630 | 47/8933 | 1,2E+08 | 4237057513 | 385694214,4 | H1f10/Apex1/Macroh2a1/Hnrnpu/Hmgn1/Smad3/Top1/Hdac2/Dhx9/Kdm6a/Mecp2/Atrx/Macroh2a2/Kdm5a/Suz12/Ddx17/Sbno1/Foxo3/Hcfc1/Rcc1/Hmgn5/Tox4/Crebbp/Actn4/Hdac3/Lemd3 | 26 |
| MF | GO:0019843 | rRNA binding | 30/1630 | 59/8933 | 1,3E+08 | 446684100 | 406611126,9 | Hmgb1/Cirbp/Rrs1/Ddx21/Npm1/Brix1/Rpf2/Emg1/Ncl/Rpl12/Rpl3/Rpl8/Rpl6/Rps4x/Rps18/Fastkd5/Rps13/Rpl4/Rpl17/Rpl5/Rps9/Rpl23/Rps14/Rpl11/Tst/Rpl7/Rplp0/Rps3/Rps11/Mrps11 | 30 |
| MF | GO:0070990 | snRNP binding | 11/1630 | 13/8933 | 3,9E+09 | 1,00417E+11 | 914083971,8 | Snrpe/Rbm39/Snrnp70/Snrpd1/Snrpb/Snrpg/Snrpa/Snrpb2/Snrpd3/Snrpd2/Prpf31 | 11 |
| MF | GO:0030515 | snoRNA binding | 16/1630 | 25/8933 | 5,4E+08 | 13565821428 | 1,23488E+11 | Nop58/Nop56/Ddx21/Nhp2/Snu13/Nolc1/Wdr3/Dkc1/Bms1/Utp6/Heatr1/Gar1/Nudt16l1/Tbl3/Rrp9/Bysl | 16 |
| MF | GO:0008094 | ATP-dependent activity, acting on DNA | 31/1630 | 71/8933 | 6,1E+08 | 14866445546 | 1,35327E+11 | Smarca2/Smarca5/Chd4/Smarcad1/Chd3/Dhx9/Rsf1/Mau2/Mcm6/Rbbp4/Atrx/Ep400/Smc3/Top2b/Msh3/Cdk7/Arid1a/Ercc3/Smarca4/Wrnip1/Bptf/Msh2/Ddx1/Rfc5/Ercc2/Smarcal1/G3bp1/Chd6/Supv3l1/Upf1/Ruvbl1 | 31 |
| MF | GO:0031491 | nucleosome binding | 14/1630 | 22/8933 | 3,2E+09 | 65884155332 | 5,99735E+11 | H1f10/Hp1bp3/Macroh2a1/Hmgn1/Smarca5/Glyr1/Hira/Parp1/Ssrp1/Arid1b/Arid1a/Pwwp3a/Rcc1/Hmgn5 | 14 |
| MF | GO:0061980 | regulatory RNA binding | 14/1630 | 25/8933 | 2,5E+11 | 4,43515E+11 | 40372672697 | Rbm4/Elavl1/Ddx21/Dhx9/Ybx1/Mecp2/Hnrnpa2b1/Matr3/Rbm10/Ago1/Hnrnpa1/Pum1/Ago2/Fmr1 | 14 |
| MF | GO:0140223 | general transcription initiation factor activity | 16/1630 | 31/8933 | 2,6E+11 | 4,5812E+11 | 4,17021E+12 | Gtf2f1/Tbpl1/Ubtf/Gtf2a1/Tbp/Gtf2f2/Taf4/Gtf3c2/Gtf2h3/Gtf3c4/Gtf3c3/Gtf3c5/Gtf2b/Gtf2e2/Taf9b/Gtf3c1 | 16 |
| MF | GO:0003725 | double-stranded RNA binding | 23/1630 | 54/8933 | 2,8E+11 | 4,8367E+11 | 4,40279E+11 | Hnrnpu/Hmgb1/Elavl1/Ddx21/Dhx9/Ilf3/Ilf2/Dhx15/Ago1/Adarb1/Zfr/Ddx1/Lsm14a/Ago2/Eif4a1/Sidt1/Supv3l1/Rc3h2/Actn1/Adar/Strbp/Mtdh/Zfr2 | 23 |
| MF | GO:0140034 | methylation-dependent protein binding | 21/1630 | 48/8933 | 3,8E+11 | 6,30512E+11 | 5,73948E+11 | Phf20l1/Cbx5/Cbx1/Cbx6/Hdgfl2/Glyr1/Cbx8/Cbx3/Atrx/Wdr5/Zmynd11/Phf2/Kdm5a/Suz12/Spin1/Zmynd8/Msl3/Bptf/Morc3/Pwp1/Fmr1 | 21 |
| MF | GO:0001091 | RNA polymerase II general transcription initiation factor binding | 12/1630 | 20/8933 | 3,9E+09 | 6,30512E+11 | 5,73948E+11 | Nop58/Hnrnpu/Gtf2f1/Fbl/Gtf2a1/Tbp/Nolc1/Ercc4/Gtf2e2/Ercc1/Crebbp/Ruvbl1 | 12 |
| MF | GO:0032451 | demethylase activity | 11/1630 | 18/8933 | 6,5E+10 | 0.0010261803613931706 | 9,3412E+11 | Riox1/Alkbh5/Rsbn1/Kdm6a/Phf2/Kdm5a/Fto/Kdm1a/Kdm1b/Kdm5b/Riox2 | 11 |
| MF | GO:0032452 | histone demethylase activity | 9/1630 | 14/8933 | 1,8E+10 | 0.002647958443039223 | 0.0024104045042149156 | Riox1/Rsbn1/Kdm6a/Phf2/Kdm5a/Kdm1a/Kdm1b/Kdm5b/Riox2 | 9 |
| MF | GO:0140457 | protein demethylase activity | 9/1630 | 14/8933 | 1,8E+10 | 0.002647958443039223 | 0.0024104045042149156 | Riox1/Rsbn1/Kdm6a/Phf2/Kdm5a/Kdm1a/Kdm1b/Kdm5b/Riox2 | 9 |
| MF | GO:0140658 | ATP-dependent chromatin remodeler activity | 12/1630 | 23/8933 | 2,4E+12 | 0.0034050342030285652 | 0.0030995614004295465 | Smarca2/Smarca5/Chd4/Smarcad1/Chd3/Rsf1/Atrx/Ep400/Arid1a/Smarca4/Smarcal1/Chd6 | 12 |
| MF | GO:0003684 | damaged DNA binding | 16/1630 | 36/8933 | 2,5E+11 | 0.0035737978625148465 | 0.00325318491595067 | Apex1/H2ax/Hmgb1/Rpa1/Polb/Parp1/Hmgb2/Ercc4/Pcna/Msh3/Pold1/Msh2/Ercc2/Ercc1/Crebbp/Rps3 | 16 |
| MF | GO:0106222 | lncRNA binding | 8/1630 | 12/8933 | 3E+12 | 0.004099147160017171 | 0.003731404019545185 | Hnrnpu/Elavl1/Celf1/Suz12/Ddx17/Kdm1a/Celf2/Smarca4 | 8 |
| MF | GO:0008173 | RNA methyltransferase activity | 16/1630 | 37/8933 | 3,7E+12 | 0.005095696906520179 | 0.004638551185679732 | Trmt10b/Fbll1/Fbl/Pcif1/Trmt2a/Emg1/Mettl1/Trmt61a/Cmtr1/Mepce/Rnmt/Mettl16/Nop2/Mettl3/Thumpd3/Nsun2 | 16 |
| MF | GO:0042826 | histone deacetylase binding | 23/1630 | 65/8933 | 7,5E+11 | 0.009526005990873597 | 0.008671407894613951 | Cbx5/Hdac1/Mef2d/Mta2/Chd4/Hdac2/Hnrnpd/Mef2a/Mta3/Sfpq/Mta1/Mecp2/Kat2a/Rbbp4/Nudt21/Nr2e1/Parp1/Ncor2/Brms1l/Top2b/Dnmt1/Akap8l/Hdac3 | 23 |
