## Supplementary Table S2 for "The molecular diversity of hippocampal regions and strata at synaptic resolution revealed by integrated transcriptomic and proteomic profiling"

| subregion | fraction | RTp_Index_ID | Sequence 5'->3' |
|---|---|---|---|
| SO | pure | RT_Index_02 | AGAGGATA |
| SO | unsorted | RT_Index_03 | CTCCTTAC |
| SP | pure | RT_Index_04 | TATGCAGT |
| SP | unsorted | RT_Index_06 | AGGCTTAG |
| SR | pure | RT_Index_07 | ATTAGACG |
| SR | unsorted | RT_Index_08 | CGGAGAGA |
| SLM | pure | RT_Index_09 | CTAGTCGA |
| SLM | unsorted | RT_Index_10 | AGCTAGAA |
| CA1 | pure | RT_Index_12 | TCTTACGC |
| CA1 | unsorted | RT_Index_13 | CTTAATAG |
| CA3 | pure | RT_Index_14 | ATAGCCTT |
| CA3 | unsorted | RT_Index_15 | TAAGGCTC |
| DG | pure | RT_Index_16 | TCGCATAA |
| DG | unsorted | RT_Index_17 | ATGGACTG |

| replicate | Nextera i7 idx | Sequence 5'->3' |
|---|---|---|
| 1 | N711 | AAGAGGCA |
| 2 | N712 | GTAGAGGA |
| 3 | N723 | TAGCGCTC |
| 4 | N714 | GCTCATGA |
| 5 | N715 | ATCTCAGG |
