## Supplementary material for "The molecular diversity of hippocampal regions and strata at synaptic resolution revealed by integrated transcriptomic and proteomic profiling": Table 1

GO_SO_mRNA

| ONTOLOGY | ID | Description | GeneRatio | BgRatio | pvalue | p.adjust | qvalue | geneID | Count |
|---|---|---|---|---|---|---|---|---|---|
| CC | GO:1990531 | phospholipid-translocating ATPase complex | 3/55 | 12/17474 | 6,3624E-06 | 0,007980511 | 0,006778898 | Atp8a2/Atp11b/Atp8a1 | 3 |
| BP | GO:0098935 | dendritic transport | 3/55 | 14/17474 | 1,048E-05 | 0,007980511 | 0,006778898 | Trak2/Kif5a/Kif5b | 3 |

GO_SP_mRNA

| ONTOLOGY | ID | Description | GeneRatio | BgRatio | pvalue | p.adjust | qvalue | geneID | Count |
|---|---|---|---|---|---|---|---|---|---|
| CC | GO:0098984 | neuron to neuron synapse | 128/1581 | 495/17474 | 7,32E-29 | 5,31E-25 | 4,74E-25 | Adgrb3/Inpp4a/Als2/Bmpr2/Map2/Epha4/Tnr/Stxbp5/Grm1/Grik2/Ank3/Slc16a7/Ppp3r1/Actr2/Cpeb4/Hnrnph1/Gria1/Arf1/Rnf112/Nf1/Cdk5r1/Nsf/Tanc2/Baiap2/Dgkb/Nrcam/Lrfn5/Rtn1/Akap5/Dicer1/Cap2/Dapk1/Homer1/Rgs7bp/Erc2/Ptk2b/Slitrk1/Rnf19a/Ywhaz/Mal2/Slc4a8/Scn8a/Rogdi/Grin2a/Mapk1/Dgcr8/Kalrn/Kpna1/Prkn/Pdpk1/Dlgap1/Strn/Slc8a1/Pura/Nr3c1/Dcc/Slc1a1/Sorcs3/Vti1a/Eif3a/Itga8/C1ql3/Tsc1/Tanc1/Lrrc4c/Kcna4/Bdnf/Prnp/Nlgn1/Slitrk3/Gria2/Snx27/Sort1/Plppr4/Lrrc7/Penk/Calb1/Epha7/Map3k7/Unc13b/Sh3gl2/Dnajc6/Dab1/Macf1/Akap9/Adam22/Pclo/Ywhah/Htt/Drd5/Septin11/Mapk10/Nos1/Ppp1r9a/Kcnd2/Plxna4/Dgki/Cntnap2/Add2/Itpr1/Grm7/Syn2/Cacna1c/Grin2b/Plekha5/Chrna7/Cpeb1/Grm5/Dlgap2/Zdhhc2/Sorbs2/Nr3c2/Nptn/Map2k1/Adam10/Clstn2/Gria3/Rbmx/Fmr1/Dmd/Il1rapl1/Arhgef9/Ogt/Drp2/Pak3/Cnksr2/Cdkl5/Frmpd4 | 128 |
| CC | GO:0099572 | postsynaptic specialization | 113/1581 | 480/17474 | 6,04E-22 | 1,46E-18 | 1,3E-18 | Adgrb3/Inpp4a/Als2/Bmpr2/Map2/Epha4/Grm1/Grik2/Ank3/Ppfia2/Slc16a7/Actr2/Cpeb4/Gabrg2/Gabrb2/Hnrnph1/Gria1/Arf1/Rnf112/Nf1/Cdk5r1/Nsf/Tanc2/Baiap2/Dgkb/Nrcam/Lrfn5/Rtn1/Akap5/Dicer1/Cap2/Dapk1/Homer1/Rgs7bp/Erc2/Ptk2b/Slitrk1/Ywhaz/Scn8a/Grin2a/Mapk1/Dgcr8/Kalrn/Kpna1/Prkn/Pdpk1/Dlgap1/Strn/Slc8a1/Pura/Nr3c1/Dcc/Sorcs3/Eif3a/Itga8/Tsc1/Tanc1/Lrrc4c/Prnp/Nlgn1/Slitrk3/Gria2/Snx27/Plppr4/Lrrc7/Epha7/Map3k7/Dnajc6/Dab1/Macf1/Akap9/Adam22/Pclo/Htt/Drd5/Gabra2/Gabrb1/Septin11/Mapk10/Nos1/Ppp1r9a/Kcnd2/Dgki/Add2/Itpr1/Syn2/Cacna1c/Grin2b/Plekha5/Gabra5/Gabrb3/Chrna7/Cpeb1/Grm5/Dlgap2/Zdhhc2/Sorbs2/Nr3c2/Nptn/Map2k1/Adam10/Clstn2/Gria3/Rbmx/Fmr1/Dmd/Il1rapl1/Arhgef9/Drp2/Pak3/Cnksr2/Cdkl5/Frmpd4 | 113 |
| CC | GO:0045211 | postsynaptic membrane | 71/1581 | 345/17474 | 3,01E-11 | 2,19E-08 | 1,95E-08 | Adgrb3/Epha4/Cacna1e/Rgs7/Cnih3/Syne1/Grm1/Grik2/Ank3/Slc16a7/Tenm2/Gabrg2/Gabrb2/Gria1/Dgkb/Nrcam/Lrfn5/Rgs7bp/Kcnma1/Slitrk1/Cntn1/Scn8a/Grin2a/Dlgap1/Strn/Slc8a1/Pcdhb16/Kctd16/Dcc/Sorcs3/Itga8/Lrrc4c/Kcna4/Lin7c/Chrm5/Nlgn1/Nbea/Slitrk3/Gria2/Plppr4/Adgrl2/Lrrc7/Epha7/Usp48/Akap9/Adam22/Cacna2d1/Drd5/Gabra2/Gabrb1/Adgrl3/Kcnd2/Dgki/Grm7/Cacna1c/Grin2b/Gabra5/Gabrb3/Chrna7/Ntrk3/Grm5/Clcn3/Nrp1/Nptn/Adam10/Clstn2/Gria3/Fmr1/Dmd/Il1rapl1/Cnksr2 | 71 |
| CC | GO:0044309 | neuron spine | 50/1581 | 212/17474 | 1,89E-10 | 1,14E-07 | 1,02E-07 | Ncoa2/Als2/Abi2/Epha4/Abl2/Grm1/Atp2b1/Ppfia2/Tenm2/Gria1/Myh10/Cdk5r1/Tanc2/Baiap2/Akap5/Homer1/Rgs7bp/Ptk2b/Ptk2/Grin2a/Strn/Slc8a1/Nr3c1/Myo5b/Slc1a1/Itga8/Kcna4/Zmynd8/Nlgn1/Gria2/Ppp3ca/Lrrc7/Calb1/Drd5/Ppp1r9a/Kcnd2/Dgki/Cntnap2/Grin2b/Cyfip1/Chrna7/Grm5/Dlgap2/Adam10/Nedd4/Gria3/Slc9a6/Fmr1/Ophn1/Frmpd4 | 50 |
| CC | GO:0150034 | distal axon | 75/1581 | 394/17474 | 3,88E-10 | 2,17E-07 | 1,93E-07 | Als2/Map2/Epha4/Arpc5/Grik2/Myh10/Cdk5r1/Nin/Calm1/Dicer1/Amph/Kcnma1/Erc2/Ptk2b/Slc4a8/Grin2a/Synj2/Prkn/Slc8a1/Dcc/Stx3/Slc1a1/Cacna1b/Olfm1/Tsc1/Setx/Tanc1/Itga4/Bdnf/Prnp/Fkbp1a/Gria2/Lrig2/Wdr47/Ptbp2/Penk/Calb1/Unc13b/Ror1/Srsf10/Pclo/Crmp1/Nos1/Ppp1r9a/Dgki/Snca/Aak1/Grm7/Grin2b/Cyfip1/Gabrb3/Chrna7/Cpeb1/Prkcb/Cttn/Gpm6a/Clcn3/Cdh8/Cdh1/Nrp1/Cbl/Myo9a/Tmod2/Rasgrf1/Clasp2/Cck/Usp9x/Elk1/Septin6/Gria3/Slc9a6/Fmr1/Ophn1/Trpc5/Cdkl5 | 75 |
| CC | GO:0043197 | dendritic spine | 48/1581 | 206/17474 | 6,73E-10 | 3,49E-07 | 3,11E-07 | Ncoa2/Als2/Abi2/Epha4/Abl2/Grm1/Atp2b1/Ppfia2/Tenm2/Gria1/Myh10/Cdk5r1/Tanc2/Baiap2/Akap5/Homer1/Rgs7bp/Ptk2b/Ptk2/Grin2a/Strn/Slc8a1/Nr3c1/Myo5b/Slc1a1/Itga8/Kcna4/Zmynd8/Nlgn1/Gria2/Ppp3ca/Lrrc7/Calb1/Drd5/Ppp1r9a/Kcnd2/Dgki/Grin2b/Cyfip1/Chrna7/Grm5/Dlgap2/Adam10/Nedd4/Gria3/Fmr1/Ophn1/Frmpd4 | 48 |
| CC | GO:0044306 | neuron projection terminus | 46/1581 | 222/17474 | 7,27E-08 | 1,82E-05 | 1,62E-05 | Epha4/Grik2/Baiap2/Amph/Kcnma1/Erc2/Slc4a8/Grin2a/Synj2/Prkn/Slc8a1/Slc1a1/Vti1a/Cacna1b/Tanc1/Bdnf/Prnp/Fkbp1a/Gria2/Penk/Calb1/Unc13b/Ror1/Srsf10/Pclo/Btbd8/Nos1/Dgki/Snca/Aak1/Grm7/Grin2b/Cyfip1/Gabrb3/Prkcb/Clcn3/Cdh8/Cdh1/Cck/Elk1/Septin6/Gria3/Slc9a6/Fmr1/Dmd/Ophn1 | 46 |
| CC | GO:0060076 | excitatory synapse | 27/1581 | 98/17474 | 9,67E-08 | 2,22E-05 | 1,98E-05 | Adgrb3/Actr3/Slc16a7/Ppp3r1/Gria1/Baiap2/Akap5/Homer1/Scn8a/Grin2a/Synj1/Kcnj6/Afdn/Kcnj3/Nlgn1/Sort1/Pclo/Ywhah/Plxna4/Dgki/Cntnap2/Grin2b/Cyfip1/Gpm6a/Gria3/Atp2b3/Ogt | 27 |
| CC | GO:0120111 | neuron projection cytoplasm | 18/1581 | 49/17474 | 1,13E-07 | 2,48E-05 | 2,22E-05 | Map2/Uhmk1/Hnrnpu/Grik2/Kif3a/Map2k4/Baiap2/Mapk1/Pura/Madd/Kcnab1/Gria2/Sfpq/Htt/Abhd13/Map2k1/Fmr1/Cdkl5 | 18 |
| CC | GO:0098685 | Schaffer collateral - CA1 synapse | 34/1581 | 143/17474 | 1,21E-07 | 2,57E-05 | 2,3E-05 | Epha4/Tnr/Stxbp5/Grm1/Slc16a7/Ppp3r1/Fbxl20/Baiap2/Rock2/Dgkb/Ptk2b/Synj1/Dcc/Myo5b/Stx3/Slc1a1/Cacna1b/Lrrc4c/Gria2/Snx27/Ppp3ca/Epha7/Sh3gl2/Gabrb1/Adgrl3/Dgki/Itpr1/Syn2/Grm5/Cdh11/Nptn/Myo9a/Slc9a6/Fmr1 | 34 |
| CC | GO:0043679 | axon terminus | 42/1581 | 200/17474 | 1,8E-07 | 3,54E-05 | 3,15E-05 | Epha4/Grik2/Amph/Kcnma1/Erc2/Slc4a8/Grin2a/Synj2/Prkn/Slc8a1/Slc1a1/Cacna1b/Tanc1/Bdnf/Prnp/Fkbp1a/Gria2/Penk/Calb1/Unc13b/Ror1/Srsf10/Pclo/Nos1/Dgki/Snca/Aak1/Grm7/Grin2b/Cyfip1/Gabrb3/Prkcb/Clcn3/Cdh8/Cdh1/Cck/Elk1/Septin6/Gria3/Slc9a6/Fmr1/Ophn1 | 42 |
| CC | GO:0098982 | GABA-ergic synapse | 32/1581 | 134/17474 | 2,53E-07 | 4,36E-05 | 3,89E-05 | Cacna1e/Atp2b1/Gabrg2/Gabrb2/Lrfn5/Htr1a/Erc2/Slitrk1/Epha3/Mdga1/Plcb1/Nlgn1/Nbea/Slitrk3/Calb1/Pclo/Gabra2/Gabrb1/Septin11/Kcnd2/Cntnap2/Itpr1/Grm7/Gabra5/Gabrb3/Clcn3/Nr3c2/Nptn/Cck/Atp2b3/Arhgef9/Ogt | 32 |
| CC | GO:0034702 | ion channel complex | 54/1581 | 295/17474 | 4,13E-07 | 6,25E-05 | 5,58E-05 | Kcnb2/Unc80/Cacna1e/Cnih3/Stxbp5/Ostm1/Grik2/Gabrg2/Gabrb2/Gria1/Akap6/Calm1/Ryr2/Nrn1/Kcnma1/Cacna2d3/Cacna1d/Ptk2b/Kcns2/Kcnv1/Kcnq3/Scn8a/Grin2a/Slc5a3/Cacnb2/Cacna1b/Olfm1/Kcnj3/Scn3a/Scn2a/Sestd1/Kcna4/Ryr3/Trpc4/Kcnab1/Gria2/Akap9/Cacna2d1/Kcnip4/Gabra2/Gabrb1/Kcnd2/Cntnap2/Cacna1c/Grin2b/Gabra5/Gabrb3/Chrna7/Cttn/Micu3/Scn3b/Trpc1/Gria3/Trpc5 | 54 |
| CC | GO:0043198 | dendritic shaft | 21/1581 | 71/17474 | 6,99E-07 | 9,22E-05 | 8,23E-05 | Map2/Epha4/Cnih3/Gria1/Nsf/Baiap2/Akap5/Homer1/Rgs7bp/Slc8a1/Slc1a1/Cacna1b/Kcna4/Zmynd8/Nlgn1/Gria2/Grm7/Cacna1c/Chrna7/Grm5/Gria3 | 21 |
| CC | GO:0008021 | synaptic vesicle | 48/1581 | 256/17474 | 8,78E-07 | 0,00011 | 9,8E-05 | Stxbp5/Atp2b1/Ppfia2/Gria1/Rnf112/Cltc/Rab40b/Calm1/Amph/Tmed9/Rab3c/Mal2/Kcnk9/Slc4a8/Grin2a/Prkn/Rab26/Wdr7/Rab27b/Stx3/Vti1a/Madd/Bdnf/Stx16/Dnajc5/Gria2/Penk/Unc13b/Sh3gl2/Atp8a1/Gabra2/Btbd8/Ica1/Cttnbp2/Dgki/Snca/Syn2/Slc2a3/Grin2b/Clcn4/Sv2b/Clcn3/Npy1r/Dmxl2/Rab8b/Adam10/Septin6/Dmd | 48 |
| CC | GO:0048786 | presynaptic active zone | 29/1581 | 122/17474 | 1,01E-06 | 0,000118 | 0,000105 | Atp2b1/Ppfia2/Gabrb2/Gria1/Zzef1/Kcnma1/Erc2/Cacna1d/Fzd3/Nufip1/Scn8a/Stx3/Cacna1b/Gria2/Gucy1b1/Pnisr/Unc13b/Pclo/Cacna2d1/Gabra2/Gabrb1/Dgki/Cntnap2/Grm7/Grin2b/Gpm6a/Nr3c2/Nptn/Gria3 | 29 |
| CC | GO:0042734 | presynaptic membrane | 45/1581 | 238/17474 | 1,51E-06 | 0,000166 | 0,000148 | Epha4/Cacna1e/Stxbp5/Grm1/Grik2/Atp2b1/Ppfia2/Gabrb2/Gria1/Ap2b1/Cltc/Itsn2/Rgs7bp/Htr1a/Kcnma1/Erc2/Cacna1d/Cntn1/Slc4a8/Scn8a/Grin2a/Kcnj6/Kctd16/Stx3/Kcnj3/Scn2a/Kcna4/Gria2/Unc13b/Dnajc6/Cacna2d1/Gabra2/Gabrb1/Dgki/Cntnap2/Grm7/Cacna1c/Grin2b/Gabra5/Chrna7/Gpm6a/Npy1r/Nptn/Gria3/Atp2b3 | 45 |

| | | | | | | | | | |
|---|---|---|---|---|---|---|---|---|---|
| CC | GO:0030427 | site of polarized growth | 41/1581 | 220/17474 | 6,28E-06 | 0,000543 | 0,000484 | Als2/Map2/Epha4/Arpc5/Myh10/Cdk5r1/Nin/Calm1/Dicer1/Erc2/Ptk2b/Dcc/Stx3/Olfm1/Tsc1/Setx/Itga4/Gria2/Lrig2/Wdr47/Ptbp2/Pclo/Crmp1/Fry/Ppp1r9a/Snca/Cyfip1/Chrna7/Cpeb1/Cttn/Gpm6a/Nrp1/Cbl/Myo9a/Tmod2/Rasgrf1/Clasp2/Usp9x/Fmr1/Trpc5/Cdkl5 | 41 |
| CC | GO:0098791 | Golgi apparatus subcompartment | 49/1581 | 307/17474 | 6,37E-05 | 0,003351 | 0,002991 | Mgat4a/Marchf4/Pam/Relch/Cnst/Tgfb2/Gcc2/Aftph/Nsg2/Arf1/Cpd/5730455P16Rik/Nsf/Atl1/Vps13b/Rbfox1/Smpd4/Birc6/Rab27b/Atp9b/Ccdc186/Arl5b/Tmem87a/Stx16/Nbea/Lrba/Akap9/Pclo/Atp8a1/Caln1/Ica1/Plekha8/Rab30/Syt17/Chid1/Marchf1/Cdh1/Ap1g1/Glg1/Mbtps1/Atp2c2/Sorl1/Adam10/Dop1a/Atp2c1/Clasp2/Slc9a7/Ocrl/Yipf6 | 49 |
| CC | GO:0005635 | nuclear envelope | 62/1581 | 429/17474 | 0,000148 | 0,005986 | 0,005342 | Rb1cc1/Inpp4a/Ptgs2/Tpr/Rgs7/Ahctf1/Syne1/Ranbp2/Ccar1/Mcm3ap/Nav3/Osbpl8/Nup107/Xpot/Nemp1/Fbxw11/Rab40b/Pum2/Akap6/Dync1h1/Ryr2/Nup153/Spin1/Ipo11/Parp8/Xpo7/Lmo7/Eny2/Rogdi/Smpd4/Senp2/Plaat1/Epha3/Tnks2/Mindy3/Tor1b/Scai/Osbpl6/Nup160/Prnp/Tmx4/Phf20/Kpna4/Gabrb1/Ankrd17/Nos1/Clip1/Sun1/Rnf6/Tnpo3/Nup205/Snca/Itpr1/Parp11/Dpy19l3/Nell1/Tnks/Pcm1/Nup93/Ddx19b/Sorl1/Tfdp2 | 62 |
| CC | GO:0031248 | protein acetyltransferase complex | 21/1581 | 98/17474 | 0,000151 | 0,00601 | 0,005363 | Mbtd1/Mbip/Atxn7/Kat6b/Eny2/Taf2/Phf20l1/Yeats2/Naa50/Atf2/Phf20/Naa15/Zzz3/Map3k7/Meaf6/Ep400/Naa25/Trrap/Ing4/Jade3/Ogt | 21 |
| CC | GO:0016442 | RISC complex | 7/1581 | 15/17474 | 0,000164 | 0,006402 | 0,005713 | Dhx9/Dicer1/Ago2/Dcp2/Ago3/Snd1/Ddx6 | 7 |
| CC | GO:0031332 | RNAi effector complex | 7/1581 | 15/17474 | 0,000164 | 0,006402 | 0,005713 | Dhx9/Dicer1/Ago2/Dcp2/Ago3/Snd1/Ddx6 | 7 |
| CC | GO:1990351 | transporter complex | 58/1581 | 397/17474 | 0,000179 | 0,006821 | 0,006087 | Kcnb2/Unc80/Cacna1e/Cnih3/Stxbp5/Ostm1/Grik2/Gabrg2/Gabrb2/Gria1/Akap6/Calm1/Ryr2/Nrn1/Kcnma1/Cacna2d3/Cacna1d/Atp8a2/Ptk2b/Kcns2/Kcnv1/Kcnq3/Scn8a/Grin2a/Slc5a3/Cacnb2/Cacna1b/Olfm1/Kcnj3/Scn3a/Scn2a/Sestd1/Kcna4/Ryr3/Trpc4/Kcnab1/Gria2/Atp6v0b/Akap9/Cacna2d1/Kcnip4/Atp8a1/Gabra2/Gabrb1/Kcnd2/Cntnap2/Cacna1c/Grin2b/Gabra5/Gabrb3/Chrna7/Cttn/Micu3/Scn3b/Tmem30a/Trpc1/Gria3/Trpc5 | 58 |
| CC | GO:0032391 | photoreceptor connecting cilium | 13/1581 | 47/17474 | 0,000194 | 0,007192 | 0,006418 | Kifap3/Ahi1/Cep290/Kif3a/Ttc8/Pcdhb16/Pcdhb22/Topors/Wdr19/D630045J12Rik/Ift122/Rpgrip1l/Lca5 | 13 |
| CC | GO:0000151 | ubiquitin ligase complex | 46/1581 | 302/17474 | 0,000316 | 0,009839 | 0,00878 | Asb1/Tmem183a/Klhl12/Rnf2/Dcaf6/Dyrk2/Cand1/Fbxw11/Rmnd5b/Fbxl20/Med24/Dcaf7/Sel1l/Klhl3/Enc1/Kbtbd7/Rnf19a/Prkn/Fem1c/Bmi1/Ubr1/Anapc1/Phc3/Dcun1d1/Fbxw7/Topors/Zyg11b/Ube4b/Klhl7/Depdc5/Fbxl5/Anapc4/Dcun1d4/Klhl8/Pcgf3/Mkln1/Cul1/Rmnd5a/Usp47/Cul4a/Klhl2/Gan/Nedd4/Armc8/Dcaf1/Fbxl2 | 46 |
| BP | GO:0007409 | axonogenesis | 95/1581 | 482/17474 | 2E-13 | 2,9E-10 | 2,59E-10 | Als2/Bmpr2/Creb1/Map2/Epha4/Actr3/Tnr/Tgfb2/Plxna2/Stxbp5/Kifbp/Ank3/Zdhhc17/Slit3/Tenm2/Myh10/Cdk5r1/Crppa/Ifrd1/Nrcam/Atl1/Nin/Flrt2/Ttc8/D130043K22Rik/Nrn1/Arhgef28/Atp8a2/Fzd3/Mycbp2/Slitrk1/Slitrk5/Matn2/Ext1/Ptk2/Cntn1/Kalrn/Alcam/Epha6/Epha3/Robo1/Robo2/Afg3l2/Dcc/Smad4/Myo5b/Prkg1/Lgi1/Olfm1/Zeb2/Itga4/Lrrc4c/Bdnf/Skil/Slitrk3/Wdr47/Plppr4/Col25a1/Usp33/Epha7/Nr4a3/Megf9/Nfib/Dock7/Dab1/Nrd1/Macf1/Sema3e/Crmp1/Epha5/Rnf6/Plxna4/Braf/Cntnap2/Foxp1/Chl1/Cyfip1/Ntrk3/Cttn/Cdh11/Cdh1/Ist1/Nrp1/Nptn/Map2k1/Prtg/Clasp2/Cck/Xk/Usp9x/Slc9a6/Slitrk4/Pak3/Trpc5/Cdkl5 | 95 |
| BP | GO:0050807 | regulation of synapse organization | 68/1581 | 302/17474 | 1,06E-12 | 1,13E-09 | 1,01E-09 | Adgrb3/Abi2/Epha4/Actr3/Ppfia2/Actr2/Myh10/Nf1/Cdk5r1/Tanc2/Baiap2/Rock2/Dgkb/Lrrn3/Nrcam/Lrfn5/Flrt2/Rps6ka5/Homer1/Ptk2b/Mycbp2/Slitrk1/Slitrk5/Ywhaz/Ptk2/Ube2v2/Opa1/Kalrn/Afdn/Mapk14/Mdga1/Myo5b/C1ql3/Tanc1/Bdnf/Prnp/Zmynd8/Nlgn1/Dhx36/Slitrk3/Adgrl2/Epha7/Adgrl3/Septin11/Ppp1r9a/Cttnbp2/Snca/Grin2b/Cyfip1/Ube3a/Chrna7/Ntrk3/Gpm6a/Cdh8/Adam10/Nedd4/Clstn2/Pgrmc1/Rbmx/Slitrk4/Fmr1/Mecp2/Il1rapl1/Ogt/Bhlhb9/Pak3/Cdkl5/Frmpd4 | 68 |
| BP | GO:0099173 | postsynapse organization | 60/1581 | 249/17474 | 1,09E-12 | 1,13E-09 | 1,01E-09 | Abi2/Epha4/Actr3/Abl2/Ppfia2/Actr2/Arf1/Myh10/Nf1/Cdk5r1/Tanc2/Baiap2/Rock2/Dgkb/Nrcam/Rps6ka5/Homer1/Htr1a/Cpne6/Ptk2b/Grin2a/Opa1/Kalrn/Afdn/Myo5b/C1ql3/Tanc1/Prnp/Zmynd8/Nlgn1/Dhx36/Slitrk3/Snx27/Epha7/Frrs1l/Sh3gl2/Dock7/Adgrl3/Epha5/Ppp1r9a/Cntnap2/Grin2b/Ube3a/Chrna7/Ntrk3/Cttn/Zdhhc2/Sorbs2/Nrp1/Myo9a/Adam10/Rbmx/Il1rapl1/Arhgef9/Ophn1/Bhlhb9/Pak3/Cnksr2/Cdkl5/Frmpd4 | 60 |
| BP | GO:0050803 | regulation of synapse structure or activity | 68/1581 | 310/17474 | 3,83E-12 | 3,47E-09 | 3,1E-09 | Adgrb3/Abi2/Epha4/Actr3/Ppfia2/Actr2/Myh10/Nf1/Cdk5r1/Tanc2/Baiap2/Rock2/Dgkb/Lrrn3/Nrcam/Lrfn5/Flrt2/Rps6ka5/Homer1/Ptk2b/Mycbp2/Slitrk1/Slitrk5/Ywhaz/Ptk2/Ube2v2/Opa1/Kalrn/Afdn/Mapk14/Mdga1/Myo5b/C1ql3/Tanc1/Bdnf/Prnp/Zmynd8/Nlgn1/Dhx36/Slitrk3/Adgrl2/Epha7/Adgrl3/Septin11/Ppp1r9a/Cttnbp2/Snca/Grin2b/Cyfip1/Ube3a/Chrna7/Ntrk3/Gpm6a/Cdh8/Adam10/Nedd4/Clstn2/Pgrmc1/Rbmx/Slitrk4/Fmr1/Mecp2/Il1rapl1/Ogt/Bhlhb9/Pak3/Cdkl5/Frmpd4 | 68 |
| BP | GO:0007612 | learning | 44/1581 | 187/17474 | 2,49E-09 | 1,2E-06 | 1,07E-06 | Adgrb3/Creb1/Ptgs2/Cacna1e/Abl2/Tnr/Zzef1/Nf1/Amph/Atxn1/Gmfb/Grin2a/Kalrn/Synj1/Prkn/Slc1a1/Sorcs3/Tsc1/Tanc1/Bdnf/Meis2/Htt/Drd5/Atp8a1/Kit/Dgki/Braf/Cntnap2/Grm7/Cacna1c/Grin2b/Gabra5/Gabrb3/Ube3a/Chrna7/Grm5/Csmd1/Ddhd2/Kmt2a/Nptn/Pias1/Cck/Pgrmc1/Mecp2 | 44 |
| BP | GO:0106027 | neuron projection organization | 32/1581 | 116/17474 | 6,25E-09 | 2,27E-06 | 2,02E-06 | Abi2/Epha4/Actr3/Abl2/Kifbp/Ppfia2/Actr2/Arf1/Cdk5r1/Tanc2/Baiap2/Homer1/Grin2a/Opa1/Kalrn/Afdn/Myo5b/Tanc1/Prnp/Zmynd8/Nlgn1/Dhx36/Epha5/Ppp1r9a/Cntnap2/Grin2b/Ube3a/Chrna7/Cttn/Adam10/Bhlhb9/Pak3 | 32 |
| BP | GO:0060078 | regulation of postsynaptic membrane potential | 34/1581 | 130/17474 | 9,44E-09 | 3,26E-06 | 2,91E-06 | Rgs7/Grik2/Gabrg2/Gria1/Baiap2/Atxn1/Rgs7bp/Plk2/Ptk2b/Fgf14/Grin2a/Prkn/Afdn/Bdnf/Chrm5/Zmynd8/Nlgn1/Npy2r/Ppp3ca/Unc13b/Pclo/Gabra2/Nos1/Ppp1r9a/Dgki/Cntnap2/Snca/Grin2b/Gabra5/Gabrb3/Chrna7/Nr3c2/Gria3/Mecp2 | 34 |
| BP | GO:0006403 | RNA localization | 38/1581 | 162/17474 | 3,28E-08 | 1,08E-05 | 9,64E-06 | Zc3h11a/Tpr/Dhx9/Hnrnpu/Ahctf1/Ranbp2/Mcm3ap/Cpsf6/Nup107/Xpot/Supt6/Nup153/Eny2/Senp2/Fyttd1/Srsf7/Lrpprc/Thoc1/Dcp2/Tsc1/Nup160/Prpf6/Dhx36/Ncbp1/Htt/G3bp2/Thoc2l/Parp11/Cpeb1/Tnks/Nup93/Ddx19b/Tomm20/Atm/Atr/Setd2/Thoc2/Fmr1 | 38 |
| BP | GO:0031346 | positive regulation of cell projection organization | 76/1581 | 449/17474 | 6,16E-08 | 1,6E-05 | 1,42E-05 | Bmpr2/Abi2/Epha4/Actr3/Abl2/Plxna2/Ahi1/Actr2/Nf1/Baiap2/Kidins220/Nin/Akap5/Ripor2/Enc1/Atp8a2/Ptk2b/Slitrk1/Cntn1/Ube2v2/Opa1/Kalrn/Epha3/Robo1/Robo2/Afdn/Fbxo38/Dcc/Myo5b/Pias2/Vldlr/Camk1d/Setx/Zeb2/Nckap1/Bdnf/Cbfa2t2/Zmynd8/Ss18l1/Nlgn1/Skil/Dhx36/Rapgef2/Fnbp1l/Lrrc7/Ror1/Nrd1/Macf1/Htt/Kit/Cep135/Clip1/Ppp1r9a/Tmem106b/Plxna4/Braf/Itpr1/Lrp6/Cyfip1/Ntrk3/Cpeb1/Gpm6a/Ist1/Nrp1/Nptn/Map2k1/Rab8b/Tmem30a/Tenm1/Fmr1/Dmd/Il1rapl1/Bhlhb9/Pak3/Trpc5/Cdkl5 | 76 |

| | | | | | | | | | |
|---|---|---|---|---|---|---|---|---|---|
| BP | GO:0016570 | histone modification | 70/1581 | 409/17474 | 1,32E-07 | 2,73E-05 | 2,44E-05 | Sf3b1/Trip12/Rbbp5/Kdm5b/Rnf2/Rcor3/Hdac2/Jmjd1c/Usp15/Ncor1/Trim37/Mbtd1/Asxl2/Mbip/Arid4a/Atxn3/Otub2/Rcor1/Arid4b/Atxn7/Kat6b/Ercc6/Nipbl/Ubr5/Eny2/Taf2/Phf20l1/Yeats2/Naa50/Nfkbiz/Paxbp1/Trerf1/Smad4/Setbp1/Kmt5b/Naa40/Lcor/Bmi1/Atf2/Rtf1/Phf20/Tbl1xr1/Ash1l/Gatad2b/Setdb1/Zzz3/Map3k7/Kdm4c/Mysm1/Rlf/Meaf6/Sfpq/Kmt2c/Mtf2/Pcgf3/Ep400/Nos1/Baz1b/Trrap/Snca/Smarcad1/Kdm5a/Ing4/Nsd3/Kmt2a/Setd2/Jade3/Mecp2/Ogt/Huwe1 | 70 |
| BP | GO:0007416 | synapse assembly | 44/1581 | 213/17474 | 1,52E-07 | 3,07E-05 | 2,74E-05 | Adgrb3/Actr3/Gabrg2/Gabrb2/Gria1/Lrrn3/Lrfn5/Akap5/Flrt2/Ptk2b/Mycbp2/Slitrk1/Slitrk5/Ptk2/Ube2v2/Mdga1/Pcdhb16/C1ql3/Bdnf/Nlgn1/Slitrk3/Adgrl2/Epha7/Dock7/Pclo/Gabra2/Adgrl3/Ppp1r9a/Cntnap2/Snca/Add2/Gabrb3/Ntrk3/Zdhhc2/Gpm6a/Cdh1/Nptn/Clstn2/Slitrk4/Mecp2/Il1rapl1/Arhgef9/Ogt/Bhlhb9 | 44 |
| BP | GO:0015931 | nucleobase-containing compound transport | 40/1581 | 187/17474 | 2,1E-07 | 3,91E-05 | 3,49E-05 | Zc3h11a/Tpr/Dhx9/Ahctf1/Ranbp2/Mcm3ap/Cpsf6/Nup107/Xpot/Supt6/Slc35b1/Nup153/Eny2/Abcc5/Senp2/Fyttd1/Srsf7/Lrpprc/Thoc1/Epg5/Tsc1/Nup160/Slc35a3/Slc35a1/Slc25a51/Ncbp1/Htt/G3bp2/Thoc2l/Slc35b4/Parp11/Cpeb1/Tnks/Nup93/Ddx19b/Tomm20/Slc25a36/Setd2/Thoc2/Fmr1 | 40 |
| BP | GO:0048167 | regulation of synaptic plasticity | 47/1581 | 243/17474 | 4,5E-07 | 6,54E-05 | 5,83E-05 | Creb1/Epha4/Ptgs2/Tnr/Grik2/Gria1/Arf1/Zzef1/Nf1/Baiap2/Akap5/Plk2/Erc2/Ptk2b/Fgf14/Grin2a/Mapk1/Lnpep/Nr3c1/Stx3/Slc1a1/Sorcs3/Bdnf/Prnp/Nlgn1/Rapgef2/Penk/Calb1/Unc13b/Htt/Drd5/Kit/Nos1/Ppp1r9a/Dgki/Braf/Snca/Grin2b/Chrna7/Cpeb1/Grm5/Zdhhc2/Kmt2a/Nptn/Rasgrf1/Fmr1/Mecp2 | 47 |
| BP | GO:1990778 | protein localization to cell periphery | 64/1581 | 377/17474 | 6,07E-07 | 8,31E-05 | 7,42E-05 | Kcnb2/Actr3/Cnst/Cnih3/Ank3/Rapgef6/Cltc/Nsf/Rab40b/Rock2/Hectd1/Sec23a/Akap5/Ttc8/Rab3c/Appl1/Exoc5/Grin2a/Pkp2/Kalrn/Zdhhc23/Epha3/Afdn/Rab26/Myo5b/Stx3/Slc1a1/Lgi1/Rab11fip2/C1ql3/Cacnb2/Gpr158/Lin7c/Prnp/Nbea/Rapgef2/Snx27/Tspan5/Lrrc7/Macf1/Adam22/Kcnip4/Wdr19/Mapk10/Tesc/Clip1/Lrp6/C2cd5/Tub/Bag4/Zdhhc2/Cdh1/Atp2c2/Scn3b/Sorl1/Map2k1/Rab8b/Adam10/Atp2c1/Clasp2/Pgrmc1/Ar/Ogt/Wnk3 | 64 |
| BP | GO:0007215 | glutamate receptor signaling pathway | 17/1581 | 49/17474 | 6,55E-07 | 8,81E-05 | 7,86E-05 | Grm1/Cpeb4/Cdk5r1/Homer1/Ptk2b/Grin2a/Kalrn/Gnaq/Slc1a1/Prnp/Plcb1/Gria2/Frrs1l/Grm7/Grin2b/Grm5/Fmr1 | 17 |
| BP | GO:0070936 | protein K48-linked ubiquitination | 20/1581 | 66/17474 | 8,35E-07 | 0,000108 | 9,65E-05 | Tnfaip3/Hace1/Ube2g2/Klhl3/Ube2e2/Kbtbd7/Marchf6/Ubr5/Ttc3/Prkn/Fbxo38/Nedd4l/Topors/Ube3c/Ube2k/Rnf34/Rnf6/Cul1/Ube3a/Birc2 | 20 |
| BP | GO:0022604 | regulation of cell morphogenesis | 50/1581 | 271/17474 | 8,62E-07 | 0,00011 | 9,79E-05 | Epha4/Actr3/Plxna2/Bves/Actr2/Kif3a/Cdc42se2/Gas7/Myh10/Baiap2/Itsn2/Akap5/Cpne6/Ptk2b/Ptk2/Prkdc/Fgd4/Opa1/Kalrn/Ttc3/Brwd1/Prkn/Afdn/Nedd4l/Myo5b/Pias2/Fam171a1/Camsap1/Ss18l1/Dhx36/Macf1/Zmym6/Sema3e/Kit/Mkln1/Plxna4/Cpne9/Cyfip1/Syt17/Zranb1/Palm3/Myo9a/Phip/Mecp2/Il1rapl1/Zmym3/Brwd3/Bhlhb9/Pak3/Cdkl5 | 50 |
| BP | GO:0016579 | protein deubiquitination | 31/1581 | 135/17474 | 9,66E-07 | 0,000117 | 0,000104 | Vcpip1/Usp37/Yod1/Tnfaip3/Usp15/Usp34/Atxn3/Otub2/Atxn7/Eny2/Taf2/Usp25/Usp14/Mindy3/Usp33/Otud6b/Usp45/Mysm1/Usp24/Usp48/Trrap/Usp12/Usp47/Zranb1/Otud4/Usp38/Cyld/Usp28/Usp4/Usp9x/Usp11 | 31 |
| BP | GO:0035249 | synaptic transmission, glutamatergic | 29/1581 | 123/17474 | 1,21E-06 | 0,000137 | 0,000122 | Als2/Ptgs2/Tnr/Cnih3/Grm1/Grik2/Gria1/Zzef1/Nf1/Homer1/Ptk2b/Ext1/Grin2a/Prkn/Nr3c1/Nlgn1/Gria2/Npy2r/Plppr4/Unc13b/Frrs1l/Dgki/Grm7/Grin2b/Grm5/Clcn3/Cdh8/Gria3/Ophn1 | 29 |
| BP | GO:0007626 | locomotory behavior | 47/1581 | 262/17474 | 4,06E-06 | 0,000387 | 0,000346 | Ncoa2/Als2/Epha4/Cacna1e/Abl2/Astn1/Tnr/Grm1/Ncor1/Atxn1/Agtpbp1/Slc4a7/Ppp3cb/Kcnma1/Gmfb/Fgf14/Cntn1/Scn8a/Grin2a/Kalrn/Lsamp/Prkn/Btbd9/Strn/Vps13a/Slc1a1/Cacna1b/Tsc1/Wdr47/Penk/Calb1/Dab1/Adam22/Htt/Adgrl3/Mapk10/Kcnd2/Cntnap2/Snca/Chl1/Cacna1c/Ube3a/Grm5/Ddhd2/Clcn3/Npy1r/Mecp2 | 47 |
| BP | GO:0031503 | protein-containing complex localization | 43/1581 | 233/17474 | 4,89E-06 | 0,000455 | 0,000406 | Hnrnpu/Cnih3/Ppp3r1/Ap2b1/Wdr35/Akap6/Akap5/Kalrn/Synj1/Pura/Myo5b/Stx3/Slc1a1/Lgi1/C1ql3/Ralgapa2/Kif3b/Ralgapb/Nlgn1/Bbs12/Nbea/Ssx2ip/Lrrc7/Mdn1/Sfpq/Akap9/Adam22/Ift172/Wdr19/Mapk10/Ift122/Tub/Caly/Pcm1/Dync2h1/Atm/Nptn/Map2k1/Adam10/Nedd4/Lca5/Ophn1/Ogt | 43 |
| BP | GO:0048588 | developmental cell growth | 49/1581 | 280/17474 | 5,23E-06 | 0,00048 | 0,000428 | Bmpr2/Map2/Actr3/Tnr/Prmt2/Slit3/Map2k4/Itsn2/Ifrd1/Akap6/Nin/D130043K22Rik/Nrn1/Cpne6/Ext1/Tomm70a/Prkn/Afdn/Nr3c1/Nedd4l/Myo5b/Prkg1/Olfm1/Zeb2/Itga4/Bdnf/Epha7/Sh3gl2/Macf1/Sema3e/Rnf6/Plxna4/Foxp1/Cpne9/Cyfip1/Akap13/Ntrk3/Syt17/Cttn/Sorbs2/Cdh1/Ist1/Nrp1/Clasp2/Usp9x/Slc9a6/Mecp2/Trpc5/Cdkl5 | 49 |
| BP | GO:0051962 | positive regulation of nervous system development | 60/1581 | 369/17474 | 5,61E-06 | 0,000502 | 0,000448 | Adgrb3/Bmpr2/Epha4/Actr3/Plxna2/Hdac2/Actr2/Rnf112/Baiap2/Lrrn3/Nin/Akap5/Flrt2/Dicer1/Atxn1/Fzd3/Slitrk1/Slitrk5/Ptk2/Ube2v2/Opa1/Kalrn/Robo1/Robo2/Synj1/Afdn/Myo5b/Pias2/Zeb2/Bdnf/Ss18l1/Nlgn1/Skil/Dhx36/Slitrk3/Adgrl2/Nrd1/Macf1/Kit/Adgrl3/Plxna4/Braf/Cyfip1/Ntrk3/Grm5/Tenm4/Ist1/Il34/Nrp1/Nptn/Map2k1/Clstn2/Slitrk4/Fmr1/Mecp2/Il1rapl1/Bhlhb9/Pak3/Trpc5/Cdkl5 | 60 |
| BP | GO:0050767 | regulation of neurogenesis | 71/1581 | 464/17474 | 7,36E-06 | 0,000614 | 0,000548 | Bmpr2/Map2/Epha4/Actr3/Tnr/Kifap3/Plxna2/Hdac2/Actr2/Rnf112/Nf1/Baiap2/Ifrd1/Nin/Akap5/Dicer1/D130043K22Rik/Atxn1/Fzd3/Slitrk1/Ptk2/Opa1/Kalrn/Robo1/Robo2/Synj1/Afdn/Myo5b/Pias2/Zeb2/Bdnf/Ss18l1/Skil/Dhx36/Rapgef2/Ppp3ca/Epha7/Brinp1/Dock7/Dab1/Nrd1/Macf1/Hes2/Sema3e/Ywhah/Kit/Nos1/Rnf6/Plxna4/Braf/Cyfip1/Ntrk3/Grm5/Tenm4/Hook3/Pcm1/Cdh1/Ist1/Il34/Nrp1/Sorl1/Nptn/Map2k1/Prtg/Fmr1/Mecp2/Il1rapl1/Bhlhb9/Pak3/Trpc5/Cdkl5 | 71 |
| BP | GO:0030705 | cytoskeleton-dependent intracellular transport | 39/1581 | 207/17474 | 8,03E-06 | 0,000654 | 0,000584 | Map2/Hnrnpu/Ank3/Ppfia2/Fbxw11/Kif3a/Tanc2/Wdr35/Dync1h1/Agtpbp1/Pura/Rab27b/Myo5b/Ccdc186/Arhgap21/Dync1i2/Madd/Rasgrp1/Kif3b/Bbs12/Fnbp1l/Ssx2ip/Hook1/Sfpq/Ift172/Htt/Wdr19/Sun1/Ift122/Tub/Caly/Ap3m2/Hook3/Pcm1/Dync2h1/Map2k1/Lca5/Fmr1/Mecp2 | 39 |
| BP | GO:0099504 | synaptic vesicle cycle | 41/1581 | 229/17474 | 1,72E-05 | 0,001196 | 0,001067 | Cacna1e/Stxbp5/Ppfia2/Ppp3r1/Cdk5r1/Ap2b1/Cltc/Fbxl20/Itsn2/Amph/Ppp3cb/Erc2/Fgf14/Slc4a8/Synj1/Synj2/Prkn/Btbd9/Rab27b/Myo5b/Stx3/Cacna1b/Dnajc5/Nlgn1/Unc13b/Sh3gl2/Dnajc6/Pclo/Btbd8/Braf/Snca/Syn2/Cyfip1/Sv2b/Prkcb/Ap3m2/Clcn3/Npy1r/Fmr1/Pls3/Ophn1 | 41 |
| BP | GO:0051650 | establishment of vesicle localization | 36/1581 | 191/17474 | 1,75E-05 | 0,001196 | 0,001067 | Map2/Clasp1/Klhl12/Ahi1/Sar1a/Ppfia2/Fbxw11/Arf1/5730455P16Rik/Tanc2/Trappc12/Dync1h1/Tmed9/Exoc5/Synj1/Prkn/Myo5b/Exoc6/Ccdc186/Itga4/Madd/Rasgrp1/Nlgn1/Fnbp1l/Ap1ar/Pclo/Htt/Exoc1/Snca/Exoc6b/Ap3m2/Trappc11/Map2k1/Clasp2/Mecp2/Shroom2 | 36 |
| BP | GO:0016441 | post-transcriptional gene silencing | 17/1581 | 61/17474 | 1,9E-05 | 0,001262 | 0,001126 | Dhx9/Helz/Pum2/Dicer1/Ago2/Tnrc6b/Trub1/Celf2/Celf1/Ncbp1/Focad/Ago3/Tial1/Elavl1/Ddx6/Fmr1/Mecp2 | 17 |
| BP | GO:0000245 | spliceosomal complex assembly | 14/1581 | 44/17474 | 1,98E-05 | 0,001305 | 0,001165 | Sf3b1/Luc7l3/Prpf39/Luc7l/Celf2/Setx/Celf1/Ptbp2/Ncbp1/Psip1/Srsf10/Srpk2/Rbm5/Rbmx | 14 |

| | | | | | | | | | |
|---|---|---|---|---|---|---|---|---|---|
| BP | GO:0060079 | excitatory postsynaptic potential | 22/1581 | 93/17474 | 2,14E-05 | 0,001387 | 0,001237 | Grik2/Baiap2/Atxn1/Plk2/Ptk2b/Grin2a/Prkn/Afdn/Bdnf/Chrm5/Zmynd8/Nlgn1/Npy2r/Ppp3ca/Pclo/Ppp1r9a/Dgki/Cntnap2/Snca/Grin2b/Chrna7/Mecp2 | 22 |
| BP | GO:1903311 | regulation of mRNA metabolic process | 48/1581 | 287/17474 | 2,24E-05 | 0,001439 | 0,001284 | Khdrbs2/Dhx9/Rc3h1/Hnrnpu/Angel2/Cpsf6/Supt6/Ddx5/Pum2/Rock2/Rbm25/Dicer1/Larp4b/Tut7/Mbnl2/Ago2/Rbfox2/Tnrc6b/Rbfox1/Son/Pnldc1/Srsf7/Dcp2/Celf2/Rc3h2/Celf1/Rbm39/Ythdf3/Dhx36/Rbm15/Virma/Ncbp1/Tut4/Srsf10/Tardbp/Plekhn1/Srpk2/Pan3/Tia1/Cpeb1/Elavl1/Bag4/Cnot7/Sltm/Rbm5/Exosc7/Rbmx/Fmr1 | 48 |
| BP | GO:0034329 | cell junction assembly | 67/1581 | 453/17474 | 3,97E-05 | 0,002382 | 0,002125 | Adgrb3/Abi2/Clasp1/Actr3/Ptprk/Gabrg2/Gabrb2/Gria1/Rock2/Lrrn3/Lrfn5/Akap5/Flrt2/Ptk2b/Mycbp2/Slitrk1/Slitrk5/Cdh12/Ptk2/Ube2v2/Pkp2/Epha3/Afdn/Pdpk1/Mdga1/Pcdhb16/Nedd4l/Slk/C1ql3/Tsc1/Ptprj/Bdnf/Fmn1/Nlgn1/Slitrk3/Rapgef2/Adgrl2/Epha7/Dock7/Macf1/Pclo/Gabra2/Adgrl3/Ppp1r9a/Cntnap2/Snca/Add2/Gabrb3/Ntrk3/Cttn/Zdhhc2/Gpm6a/Cdh8/Cdh11/Cdh1/Nrp1/Nptn/Myo9a/Clstn2/Clasp2/Slitrk4/Mecp2/Il1rapl1/Arhgef9/Ophn1/Ogt/Bhlhb9 | 67 |
| BP | GO:0098742 | cell-cell adhesion via plasma-membrane adhesion molecules | 37/1581 | 206/17474 | 4,03E-05 | 0,002398 | 0,00214 | Kifap3/Tgfb2/Tnfaip3/Tenm2/Nrcam/Lrfn5/Pcdh20/Slitrk1/Cdh12/Alcam/Epha3/Cadm2/Robo1/Robo2/Mapk14/Mdga1/Pcdhb16/Itga4/Lrrc4c/Nlgn1/Fat4/Slitrk3/Dab1/Adgrl3/Tenm4/Cdh8/Cdh11/Cdh1/Fat3/Nptn/Prtg/Clstn2/Pik3cb/Atp2c1/Tenm1/Il1rapl1/Pcdh19 | 37 |
| BP | GO:0034453 | microtubule anchoring | 10/1581 | 26/17474 | 4,9E-05 | 0,002866 | 0,002557 | Clasp1/Cep350/Gcc2/Kif3a/Nin/Ninl/Bccip/Hook3/Pcm1/Clasp2 | 10 |
| BP | GO:0006338 | chromatin remodeling | 53/1581 | 339/17474 | 5,76E-05 | 0,003188 | 0,002845 | Sf3b1/Satb2/Trip12/Rbbp5/Kdm5b/Tpr/Rnf2/Shprh/Tspyl1/Hdac2/Mcm3ap/Hcfc2/Supt6/Dicer1/Jarid2/Zfp369/Brd9/Kat6b/Tasor/Pbrm1/Ercc6/Chd8/Tspyl5/Ubr5/Ubn1/Yeats2/Chd1/Smchd1/Nr3c1/Top1/Gatad2b/Setdb1/Kdm4c/Mysm1/Hp1bp3/Srpk2/Brdt/Kdm2b/Bcl7a/Baz1b/Smarcad1/Rybp/Kdm5a/Chd2/Rsf1/Ctcf/Nfat5/Kmt2a/Setd2/Zfp445/Smarca1/Mecp2/Atrx | 53 |
| BP | GO:0042596 | fear response | 16/1581 | 60/17474 | 5,98E-05 | 0,003257 | 0,002906 | Als2/Cacna1e/Grik2/Fbxl20/Htr1a/Ext1/Slc1a1/Bdnf/Npy2r/Penk/Brinp1/Grm7/Grin2b/Gabra5/Cck/Mecp2 | 16 |
| BP | GO:0001662 | behavioral fear response | 15/1581 | 54/17474 | 6,05E-05 | 0,003257 | 0,002906 | Als2/Cacna1e/Grik2/Fbxl20/Htr1a/Slc1a1/Bdnf/Npy2r/Penk/Brinp1/Grm7/Grin2b/Gabra5/Cck/Mecp2 | 15 |
| BP | GO:0006605 | protein targeting | 46/1581 | 284/17474 | 7,36E-05 | 0,003705 | 0,003306 | Gdap1/Pikfyve/Actr3/Man1a/Gcc2/Ank3/Gnptab/Rab3ip/Dusp18/Cdk5r1/Trim37/Vps41/Sgtb/Ywhaz/Zdhhc23/Tomm70a/Grpel2/Vps13a/Slc1a1/Oga/Sec61a2/Prnp/Ywhab/Tomm34/Nlgn1/Nbea/Fbxw7/Sort1/Zdhhc21/Macf1/Vps13d/Pex1/Pan3/Tcaf1/C2cd5/Zfand6/Bag4/Zdhhc2/Spcs3/Tomm20/Sorl1/Rab8b/Vps13c/Nedd4/Usp9x/Ogt | 46 |
| BP | GO:0060560 | developmental growth involved in morphogenesis | 46/1581 | 284/17474 | 7,36E-05 | 0,003705 | 0,003306 | Bmpr2/Map2/Actr3/Kdm5b/Tnr/Slit3/Itsn2/Ifrd1/Nin/D130043K22Rik/Nrn1/Fgf10/Cpne6/Ext1/Robo1/Prkn/Afdn/Nedd4l/Myo5b/Prkg1/Olfm1/Zeb2/Itga4/Bdnf/Fmn1/Epha7/Sh3gl2/Macf1/Sema3e/Rnf6/Plxna4/Cpne9/Lrp6/Cyfip1/Ntrk3/Syt17/Cttn/Cdh1/Ist1/Nrp1/Clasp2/Usp9x/Slc9a6/Mecp2/Trpc5/Cdkl5 | 46 |
| BP | GO:0006887 | exocytosis | 57/1581 | 376/17474 | 7,4E-05 | 0,003705 | 0,003306 | Clasp1/Cacna1e/Mia3/Stxbp5/Ppfia2/Rab3ip/Arf1/Smcr8/Myh10/Fbxl20/Nsf/Syt16/Vps41/Tmem167/Rab3c/Ppp3cb/Erc2/Exoc5/Ywhaz/Slc4a8/Synj1/Prkn/Pdpk1/Rab26/Fer/Rab27b/Myo5b/Stx3/Exoc6/Rab11fip2/Cacna1b/Lin7c/Rasgrp1/Nlgn1/Unc13b/Nr4a3/Pclo/Kit/Exoc1/Braf/Snca/Exoc6b/Syn2/Cacna1c/Mical3/D6Wsu163e/Sv2b/Syt17/Prkcb/Trappc11/Npy1r/Ap1g1/Snx19/Rab8b/Clasp2/Fmr1/Il1rapl1 | 57 |
| BP | GO:0051648 | vesicle localization | 37/1581 | 212/17474 | 7,57E-05 | 0,003738 | 0,003335 | Map2/Clasp1/Klhl12/Ahi1/Sar1a/Ppfia2/Fbxw11/Arf1/5730455P16Rik/Tanc2/Trappc12/Dync1h1/Tmed9/Exoc5/Synj1/Prkn/Myo5b/Exoc6/Ccdc186/Itga4/Madd/Rasgrp1/Nlgn1/Fnbp1l/Ap1ar/Pclo/Htt/Exoc1/Snca/Exoc6b/Syn2/Ap3m2/Trappc11/Map2k1/Clasp2/Mecp2/Shroom2 | 37 |
| BP | GO:0034249 | negative regulation of amide metabolic process | 36/1581 | 205/17474 | 8,26E-05 | 0,003982 | 0,003553 | Tpr/Dhx9/Rc3h1/Hnrnpu/Cpeb4/Rock2/Rtn1/Tut7/Dapk1/Enc1/Ago2/Tnrc6b/Pnldc1/Dcp2/Tsc1/Rc3h2/Celf1/Prnp/Ythdf3/Dhx36/Trim2/Otud6b/Tut4/Ago3/Tardbp/Plekhn1/Pan3/Tia1/Cpeb1/Spon1/Cnot7/Sorl1/Ddx6/Xrn1/Exosc7/Fmr1 | 36 |
| BP | GO:0043954 | cellular component maintenance | 19/1581 | 82/17474 | 0,000103 | 0,004614 | 0,004117 | Adgrb3/Abl2/Homer1/Appl1/Erc2/Pkp2/Afdn/Tanc1/Prnp/Zmynd8/Nlgn1/Sort1/Adgrl3/Plxna4/Cntnap2/Grin2b/Cttn/Ophn1/Shroom2 | 19 |
| BP | GO:1901888 | regulation of cell junction assembly | 40/1581 | 240/17474 | 0,000113 | 0,004939 | 0,004407 | Adgrb3/Clasp1/Actr3/Rock2/Lrrn3/Lrfn5/Flrt2/Ptk2b/Mycbp2/Slitrk1/Slitrk5/Ptk2/Ube2v2/Pkp2/Epha3/Mdga1/Nedd4l/Slk/Tsc1/Ptprj/Bdnf/Fmn1/Nlgn1/Slitrk3/Rapgef2/Adgrl2/Epha7/Macf1/Adgrl3/Ppp1r9a/Cntnap2/Snca/Ntrk3/Nrp1/Clstn2/Clasp2/Slitrk4/Il1rapl1/Ogt/Bhlhb9 | 40 |
| BP | GO:0032386 | regulation of intracellular transport | 51/1581 | 331/17474 | 0,000115 | 0,004988 | 0,004451 | Map2/Actr3/Yod1/Ptgs2/Tpr/Dhx9/Uhmk1/Bves/Gcc2/Sar1a/Ank3/Ube2g2/Cep290/Ppp1r12a/Cpsf6/Kif3a/Arf1/Supt6/Nf1/Cdk5r1/Akap5/Dync1h1/Nup153/Ice1/Ubr5/Mapk1/Mapk14/Slc1a1/Ctdspl2/Prnp/Fbxw7/Tardbp/Tcaf1/Reep1/C2cd5/Caly/Bag4/Zdhhc2/Pcm1/Cdh1/Arv1/Sorl1/Map2k1/Nedd4/Tmem30a/Dnajc13/Setd2/Thoc2/Mecp2/Snx12/Ogt | 51 |
| BP | GO:0099072 | regulation of postsynaptic membrane neurotransmitter receptor levels | 26/1581 | 132/17474 | 0,000117 | 0,005004 | 0,004465 | Cnih3/Ppp3r1/Ap2b1/Akap5/Kalrn/Synj1/Prkn/Stx3/Lgi1/C1ql3/Nbea/Snx27/Lrrc7/Frrs1l/Adam22/Drd5/Mapk10/Tfr2/Cntnap2/Caly/Map2k1/Adam10/Nedd4/Slc9a6/Ophn1/Ogt | 26 |
| BP | GO:1905475 | regulation of protein localization to membrane | 35/1581 | 201/17474 | 0,000122 | 0,005228 | 0,004665 | Actr3/Cnst/Ank3/Cdk5r1/Ap2b1/Cltc/Hectd1/Akap5/Appl1/Kalrn/Epha3/Slc5a3/Prkn/Myo5b/Stx3/Slc1a1/Rab11fip2/Prnp/Zmynd8/Nbea/Snx27/Frrs1l/Tcaf1/C2cd5/Zdhhc2/Sorbs2/Cdh1/Map2k1/Adam10/Atp2c1/Clasp2/Pgrmc1/Ar/Ogt/Wnk3 | 35 |
| BP | GO:0007623 | circadian rhythm | 37/1581 | 218/17474 | 0,000137 | 0,005691 | 0,005078 | Ncoa2/Creb1/Kdm5b/Dhx9/Hnrnpu/Hdac2/Fbxw11/Ncor1/Ddx5/Rock2/Kcnma1/Mycbp2/Klf10/Prkdc/Nrip1/Btbd9/Setx/Bdnf/Top1/Nlgn1/Npy2r/Fbxw7/Sfpq/Tardbp/Per3/Clock/Mapk10/Kcnd2/Cntnap2/Kdm5a/Gabrb3/Ube3a/Ntrk3/Kmt2a/Usp9x/Ogt/Huwe1 | 37 |
| BP | GO:0035640 | exploration behavior | 12/1581 | 40/17474 | 0,000146 | 0,005986 | 0,005342 | Abl2/Tnr/Atxn3/Htr1a/Slc4a7/Lsamp/Penk/Brinp1/Chl1/Gabrb3/Ube3a/Kmt2a | 12 |
| BP | GO:0010498 | proteasomal protein catabolic process | 68/1581 | 483/17474 | 0,00016 | 0,006343 | 0,00566 | Uggt1/Trip12/Yod1/Edem3/Ascc3/Man1a/Ube2g2/Ascc2/Psme4/Fbxw11/Rmnd5b/Fbxl20/Hectd1/Sel1l/Atxn3/Enc1/Plk2/Kbtbd7/Uggt2/Marchf6/Rnf19a/Phf20l1/Clec16a/Ube2v2/Ubxn7/Usp25/Ltn1/Prkn/Pja2/Dlgap1/Usp14/Fem1c/Csnk1a1/Fbxo38/Nedd4l/Erlin1/Marchf7/Ubr1/Anapc1/Tbl1xr1/Trim2/Fbxw7/Ubxn2b/Topors/Ecpas/Zyg11b/Ube4b/Fbxl5/Anapc4/Ube2k/Clock/Rnf34/Cul1/Rmnd5a/Rybp/Herc2/Cul4a/Sh3rf1/Clgn/Birc2/Pias1/Gclc/Armc8/Fbxl2/Usp9x/Fmr1/Ophn1/Ogt | 68 |
| BP | GO:0035418 | protein localization to synapse | 21/1581 | 99/17474 | 0,000175 | 0,006729 | 0,006005 | Cnih3/Baiap2/Homer1/Grin2a/Kalrn/Dlgap1/Rab27b/Stx3/Lgi1/C1ql3/Nlgn1/Nbea/Slitrk3/Lrrc7/Adam22/Pclo/Mapk10/Zdhhc2/Map2k1/Adam10/Ogt | 21 |

| | | | | | | | | | |
|---|---|---|---|---|---|---|---|---|---|
| BP | GO:0006406 | mRNA export from nucleus | 15/1581 | 59/17474 | 0,000181 | 0,006844 | 0,006107 | Zc3h11a/Tpr/Mcm3ap/Nup107/Supt6/Eny2/Fyttd1/Thoc1/Nup160/Ncbp1/Thoc2l/Nup93/Ddx19b/Setd2/Thoc2 | 15 |
| BP | GO:1990138 | neuron projection extension | 35/1581 | 205/17474 | 0,000182 | 0,00686 | 0,006122 | Bmpr2/Map2/Actr3/Tnr/Slit3/Itsn2/Ifrd1/D130043K22Rik/Nrn1/Cpne6/Prkn/Afdn/Nedd4l/Myo5b/Olfm1/Itga4/Bdnf/Sh3gl2/Macf1/Sema3e/Rnf6/Plxna4/Cpne9/Cyfip1/Ntrk3/Syt17/Cttn/Cdh1/Nrp1/Clasp2/Usp9x/Slc9a6/Mecp2/Trpc5/Cdkl5 | 35 |
| BP | GO:0008380 | RNA splicing | 59/1581 | 407/17474 | 0,000195 | 0,007192 | 0,006418 | Khdrbs2/Sf3b1/Clk1/Dhx9/Hnrnpu/Cdc40/Hnrnph1/Clk4/Supt6/Luc7l3/Ddx5/Prpf39/Rbm25/Ddx46/Srek1/Atxn7/Zc3h13/Mbnl2/Eny2/Rbfox2/Rbfox1/Paxbp1/Son/Luc7l/Srsf7/Thoc1/Celf2/Setx/Celf1/Snrnp200/Rbm39/Prpf6/Ccnl1/Rbm15/Prpf38b/Ptbp2/Zranb2/Virma/Rbm12b2/Ncbp1/Psip1/Sfpq/Srsf10/Tardbp/Ccnl2/Srpk2/Brdt/Zcchc8/Trrap/Rbm28/Zfp638/Tia1/Ppp4r2/Rbm5/Usp4/Thoc2/Htatsf1/Rbmx/Fmr1 | 59 |
| BP | GO:0017148 | negative regulation of translation | 32/1581 | 182/17474 | 0,000195 | 0,007192 | 0,006418 | Tpr/Dhx9/Rc3h1/Hnrnpu/Cpeb4/Rock2/Tut7/Dapk1/Enc1/Ago2/Tnrc6b/Pnldc1/Dcp2/Tsc1/Rc3h2/Celf1/Ythdf3/Dhx36/Trim2/Otud6b/Tut4/Ago3/Tardbp/Plekhn1/Pan3/Tia1/Cpeb1/Cnot7/Ddx6/Xrn1/Exosc7/Fmr1 | 32 |
| BP | GO:0048638 | regulation of developmental growth | 57/1581 | 392/17474 | 0,000229 | 0,008029 | 0,007165 | Bmpr2/Creb1/Map2/Actr3/Tnr/Hdac2/Dusp6/Ncor1/Itsn2/Ifrd1/Akap6/D130043K22Rik/Jarid2/Cpne6/Atp8a2/Mycbp2/Nipbl/Ptk2/Prkdc/Mapk1/Tomm70a/Prkn/Afdn/Mapk14/Nr3c1/Nedd4l/Afg3l2/Myo5b/Tnks2/Olfm1/Mbd5/Celf1/Bdnf/Atrn/Plcb1/Stk4/Epha7/Macf1/Sema3e/Rnf6/Plxna4/Foxp1/Cpne9/Cacna1c/Cyfip1/Ntrk3/Syt17/Cttn/Npy1r/Cdh1/Ist1/Nrp1/Clasp2/Mecp2/Ar/Trpc5/Cdkl5 | 57 |
| BP | GO:0072594 | establishment of protein localization to organelle | 61/1581 | 428/17474 | 0,000245 | 0,008413 | 0,007507 | Gdap1/Ptgs2/Tpr/Tnfaip3/Man1a/Gcc2/Ranbp2/Gnptab/Nup107/Dusp18/Ppp3r1/Nf1/Trim37/Ddx5/Akap5/Ryr2/Vps41/Nup153/Tnpo1/Sgtb/Ipo11/Appl1/Kpna3/Nipbl/Ywhaz/Ubr5/Mapk1/Kpna1/Tomm70a/Mapk14/Grpel2/Vps13a/Sec61a2/Atf2/Tomm34/Stk4/Kpna4/Fbxw7/Sort1/Ppp3ca/Lrrc7/Macf1/Vps13d/Tardbp/Pex1/Tnpo3/Ift122/Zfand6/Sprn/Elavl1/Bag4/Spcs3/Nup93/Cdh1/Tomm20/Sorl1/Rab8b/Vps13c/Nedd4/Phip/Usp9x | 61 |
| BP | GO:0008038 | neuron recognition | 13/1581 | 48/17474 | 0,000245 | 0,008413 | 0,007507 | Epha4/Cdk5r1/Nrcam/Ywhaz/Ext1/Epha3/Robo1/Robo2/Tnfrsf21/Bdnf/Cntnap2/Nrp1/Prtg | 13 |
| BP | GO:0030900 | forebrain development | 57/1581 | 394/17474 | 0,000262 | 0,008874 | 0,007919 | Creb1/Tnr/Zbtb18/Hdac2/Kif3a/Ncor1/Myh10/Nf1/Cdk5r1/Nin/Ttc8/Dicer1/Inhba/Agtpbp1/Fgf10/Slitrk5/Ext1/Mrtfb/Robo1/Robo2/Afdn/Mdga1/Gnaq/Prkg1/Tsc1/Zeb2/Neurod1/Celf1/Plcb1/Fat4/Rapgef2/Wdr47/Nr4a3/Nfib/Dock7/Dab1/Sema3e/Htt/Jakmip1/Epha5/Kdm2b/Sun1/Plxna4/Cntnap2/Neurod6/Lrp6/Hook3/Pcm1/Rpgrip1l/Nrp1/Dync2h1/Dmxl2/Herc1/Setd2/Usp9x/Ophn1/Atrx | 57 |
| BP | GO:0001764 | neuron migration | 34/1581 | 201/17474 | 0,000268 | 0,008986 | 0,008019 | Satb2/Abi2/Nav1/Astn1/Cep85l/Myh10/Cdk5r1/Flrt2/Fzd3/Nipbl/Matn2/Ptk2/Mrtfb/Mdga1/Dcc/Prkg1/Rapgef2/Lrig2/Fktn/Dab1/Sema3e/Adgrl3/Chl1/Ntrk3/Pcm1/Gpm6a/Sh3rf1/Cdh1/Nrp1/Fat3/Cck/Usp9x/Nexmif/Cdkl5 | 34 |
| BP | GO:1902903 | regulation of supramolecular fiber organization | 56/1581 | 386/17474 | 0,000274 | 0,009053 | 0,008078 | Abi2/Map2/Clasp1/Actr3/Camsap2/Arpc5/Hdac2/Washc4/Nav3/Arf1/Ssh2/Taok1/Ppm1e/Baiap2/Cyria/Rock2/Togaram1/Sptb/Ttc8/Htr1a/Atxn7/Gmfb/Ptk2b/Rictor/Prkn/Tnxb/Fer/Eml4/Smad4/Camsap1/Tsc1/Sptan1/Nckap1/Fmn1/Wdr47/Ap1ar/Akap9/Cracd/Clip1/Ppp1r9a/Braf/Snca/Add2/Cyfip1/Chrna7/Akap13/Cttn/Bag4/Colgalt1/Nrp1/Tmod2/Clasp2/Tenm1/Mecp2/Pak3/Shroom2 | 56 |
| BP | GO:0070507 | regulation of microtubule cytoskeleton organization | 29/1581 | 162/17474 | 0,000279 | 0,009069 | 0,008092 | Map2/Clasp1/Camsap2/Tpr/Hnrnpu/Nav3/Taok1/Cdk5r1/Cltc/Ska2/Togaram1/Dync1h1/Ripor2/Htr1a/Atxn7/Epha3/Eml4/Camsap1/Map9/Wdr47/Akap9/Clip1/Hsph1/Snca/Cyld/Ccsap/Cep70/Clasp2/Mecp2 | 29 |
| BP | GO:0032886 | regulation of microtubule-based process | 41/1581 | 259/17474 | 0,000291 | 0,009339 | 0,008334 | Map2/Clasp1/Camsap2/Tpr/Hnrnpu/Nav3/Kif3a/Taok1/Cdk5r1/Cltc/Ska2/Trim37/Rock2/Togaram1/Dync1h1/Ripor2/Htr1a/Plk2/Atxn7/Epha3/Memo1/Eml4/Camsap1/Ccdc39/Ccnl1/Map9/Wdr47/Macf1/Ccnl2/Akap9/Clip1/Hsph1/Snca/Caly/Cyld/Ccsap/Cep295/Chordc1/Cep70/Clasp2/Mecp2 | 41 |
| BP | GO:0010765 | positive regulation of sodium ion transport | 12/1581 | 43/17474 | 0,000311 | 0,009721 | 0,008674 | Akt3/Ank3/Arf1/Fgf14/Cntn1/Nedd4l/Plcb1/Nos1/Tesc/Scn3b/Dmd/Wnk3 | 12 |
| BP | GO:0046834 | lipid phosphorylation | 6/1581 | 12/17474 | 0,000311 | 0,009721 | 0,008674 | Dgkd/Dgke/Dgkb/Dgkh/Dgkg/Dgki | 6 |
| MF | GO:0004843 | cysteine-type deubiquitinase activity | 28/1581 | 92/17474 | 5,05E-09 | 1,93E-06 | 1,72E-06 | Vcpip1/Usp37/Yod1/Tnfaip3/Usp15/Usp34/Usp32/Atxn3/Otub2/Usp25/Usp14/Mindy3/Usp33/Otud6b/Usp45/Mysm1/Usp24/Usp48/Usp12/Usp47/Zranb1/Otud4/Usp38/Cyld/Usp28/Usp4/Usp9x/Usp11 | 28 |
| MF | GO:0042393 | histone binding | 46/1581 | 224/17474 | 9,62E-08 | 2,22E-05 | 1,98E-05 | Rbbp5/Kdm5b/Tspyl1/Mcm3ap/Usp15/Psme4/Zzef1/Supt6/Mbtd1/Zmynd11/Jarid2/Spin1/Brd9/Kat6b/Sfmbt1/Chd8/Tspyl5/Atad2/Cbx5/Yeats2/Chd1/Uhrf2/Trp53bp1/Zmynd8/Tbl1xr1/Zzz3/Mllt3/Mysm1/Kmt2c/Brdt/Mtf2/Sbno1/Baz1b/Kdm7a/Snca/Kdm5a/Ing4/Chd2/Rsf1/Prkcb/Tnks/Kmt2a/Pygo1/Phip/Fmr1/Atrx | 46 |
| MF | GO:0051020 | GTPase binding | 60/1581 | 328/17474 | 9,78E-08 | 2,22E-05 | 1,98E-05 | Als2/Abi2/Rabgap1l/Hace1/Ranbp2/Rab3ip/Xpot/Kif3a/Rapgef6/Cdc42se2/Gria1/Ankfy1/Nsf/Cyria/Rock2/Dock4/Strn3/Nup153/Iqgap2/Tnpo1/Ipo11/Plk2/Exoc5/Xpo7/Mycbp2/Dock9/Acap2/Afdn/Fer/Myo5b/Ric1/Marchf5/Ccdc186/Rab11fip2/Odf2/Gapvd1/Nckap1/Kif3b/Fnbp1l/Usp33/Rragd/Dock7/Ppp1r9a/Tnpo3/Dgki/Braf/Mical3/Dennd5b/Cyfip1/Akap13/Ap1g1/Sorl1/Dmxl2/Map2k1/Rasgrf1/Wdr44/Ocrl/Diaph2/Pak3/Cdkl5 | 60 |
| MF | GO:0019783 | ubiquitin-like protein peptidase activity | 29/1581 | 115/17474 | 2,66E-07 | 4,49E-05 | 4E-05 | Vcpip1/Usp37/Yod1/Tnfaip3/Usp15/Usp34/Usp32/Atxn3/Otub2/Senp2/Usp25/Usp14/Mindy3/Usp33/Otud6b/Usp45/Mysm1/Usp24/Usp48/Usp12/Usp47/Zranb1/Otud4/Usp38/Cyld/Usp28/Usp4/Usp9x/Usp11 | 29 |
| MF | GO:0061659 | ubiquitin-like protein ligase activity | 58/1581 | 327/17474 | 4,66E-07 | 6,63E-05 | 5,92E-05 | Trip12/Asb1/Rnf2/Rc3h1/Shprh/Hace1/Ranbp2/Fbxw11/Rmnd5b/Rnf112/Zzef1/Trim37/Med24/Hectd1/Arel1/Mycbp2/Marchf6/Rnf19a/Ubr5/Dzip3/Ltn1/Prkn/Pja2/Nedd4l/Pias2/Uhrf2/Hectd2/Marchf5/Rc3h2/Ubr1/Trim2/Fbxw7/Trim33/Topors/Nfx1/Ube4b/Ube3c/Ube2k/Rnf34/Rnf6/Mkrn1/Cul1/Herc3/Rmnd5a/Herc2/Ube3a/Trim68/Sh3rf1/Marchf1/Rnf150/Birc2/Cbl/Pias1/Nedd4/Traip/Pja1/Rlim/Huwe1 | 58 |
| MF | GO:0140030 | modification-dependent protein binding | 36/1581 | 169/17474 | 9,3E-07 | 0,000114 | 0,000102 | Tnfaip3/Ank3/Usp15/Psme4/Zzef1/Mbtd1/Zmynd11/Jarid2/Spin1/Brd9/Chd8/Phf20l1/Cbx5/Yeats2/Dzip3/Chd1/Trp53bp1/Zmynd8/Zzz3/Mllt3/Pex1/Tnip2/Brdt/Mtf2/Kdm7a/Kdm5a/Ing4/Zfand6/Rnf169/Zranb1/Kmt2a/Pygo1/Phip/Fmr1/Tab3/Atrx | 36 |
| MF | GO:0140658 | ATP-dependent chromatin remodeler activity | 13/1581 | 33/17474 | 2,64E-06 | 0,000266 | 0,000237 | Smarcal1/Shprh/Ercc6/Chd8/Chd1/Ep400/Smarcad1/Chd2/Rsf1/Chd9/Rad54l2/Smarca1/Atrx | 13 |
| MF | GO:0099507 | ligand-gated monoatomic ion channel activity involved in regulation of presynaptic membrane potential | 11/1581 | 24/17474 | 2,64E-06 | 0,000266 | 0,000237 | Grik2/Gabrb2/Gria1/Kcnma1/Grin2a/Gria2/Gabra2/Gabrb1/Grin2b/Gabra5/Gria3 | 11 |

| | | | | | | | | | |
|---|---|---|---|---|---|---|---|---|---|
| MF | GO:0004386 | helicase activity | 31/1581 | 145/17474 | 4,79E-06 | 0,000451 | 0,000403 | Smarcal1/Dhx9/Shprh/Ascc3/Ddx50/Eif4a1/Dhx33/Ddx5/Helz/Dicer1/Ddx46/Ddx4/Ercc6/Chd8/Chd1/Dhx57/Ythdc2/Setx/Snrnp200/Dhx36/Hfm1/Ep400/Smarcad1/Chd2/Chd9/Ddx19b/Ddx6/Ddx10/Rad54l2/Smarca1/Atrx | 31 |
| MF | GO:0003712 | transcription coregulator activity | 73/1581 | 478/17474 | 5,89E-06 | 0,00052 | 0,000464 | Ncoa2/Nucks1/Kdm5b/Dhx9/Dcaf6/Hnrnpu/Rcor3/Bclaf1/Ncoa7/Ccar1/Jmjd1c/Prmt2/Hcfc2/Ewsr1/Ncor1/Trim37/Med24/Ddx5/Ncoa1/Myt1l/Rcor1/Zmynd11/Nsd1/Kat6b/Sfmbt1/Sub1/Mtdh/Eny2/Rbfox2/Mrtfb/Nfkbiz/Nrip1/Prkn/Trerf1/Lpin2/Nr3c1/Tcf4/Dcc/Pias2/Btaf1/Lcor/Scai/Trp53bp1/Cbfa2t2/Zmynd8/Ss18l1/Prpf6/Tbl1xr1/Med12l/Gon4l/Runx1t1/Phf24/Psip1/Mysm1/Camta1/Kmt2c/Slc30a9/Kdm2b/Trrap/N4bp2l2/Kdm7a/Rybp/Kdm5a/Ing4/Ube3a/Prkcb/Ccnd1/Cnot7/Birc2/Pias1/Rad54l2/Elk1/Mecp2 | 73 |
| MF | GO:0005516 | calmodulin binding | 35/1581 | 189/17474 | 3,35E-05 | 0,002057 | 0,001836 | Map2/Camsap2/Atp2b1/Ewsr1/Myh10/Camkk1/Ddx5/Strn3/Akap5/Slc24a4/Ryr2/Dapk1/Iqgap2/Ppp3cb/Myh7/Kcnq3/Strn/Slc8a1/Myo5b/Camk1d/Camsap1/Sptan1/Scn3a/Pde1a/Ryr3/Plcb1/Ppp3ca/Unc13b/Nos1/Camkk2/Add2/Grm7/Cacna1c/Fbxl2/Phka2 | 35 |
| MF | GO:0030594 | neurotransmitter receptor activity | 21/1581 | 92/17474 | 5,75E-05 | 0,003188 | 0,002845 | Htr5b/Grm1/Grik2/Gabrg2/Gabrb2/Gria1/Htr1a/Ptk2b/Grin2a/Htr4/Chrm5/Gria2/Drd5/Gabra2/Gabrb1/Grin2b/Gabra5/Gabrb3/Chrna7/Grm5/Gria3 | 21 |
| MF | GO:0008066 | glutamate receptor activity | 10/1581 | 27/17474 | 7,15E-05 | 0,003653 | 0,00326 | Grm1/Grik2/Gria1/Ptk2b/Grin2a/Gria2/Grm7/Grin2b/Grm5/Gria3 | 10 |
| MF | GO:0022839 | monoatomic ion gated channel activity | 48/1581 | 302/17474 | 8,38E-05 | 0,003998 | 0,003567 | Kcnb2/Kcnt2/Cacna1e/Grik2/Gabrg2/Gabrb2/Gria1/Tmem63c/Ryr2/Kcnma1/Cacna2d3/Cacna1d/Ptk2b/Nalcn/Kcns2/Kcnv1/Kcnq3/Kcnk9/Scn8a/Grin2a/Kcnj6/Cacnb2/Cacna1b/Kcnj3/Kcnh7/Scn3a/Scn2a/Kcna4/Chrm5/Ryr3/Kcnab1/Gria2/Cacna2d1/Kcnip4/Gabra2/Gabrb1/Kcnd2/Itpr1/Grm7/Cacna1c/Grin2b/Clcn4/Gabra5/Gabrb3/Chrna7/Clcn3/Scn3b/Gria3 | 48 |
| MF | GO:0035198 | miRNA binding | 11/1581 | 33/17474 | 9,52E-05 | 0,004402 | 0,003928 | Rc3h1/Pum2/Dicer1/Tut7/Ago2/Zc3h7a/Matr3/Tut4/Ago3/Elavl1/Fmr1 | 11 |
| MF | GO:0050681 | nuclear androgen receptor binding | 11/1581 | 33/17474 | 9,52E-05 | 0,004402 | 0,003928 | Prmt2/Ddx5/Nsd1/Pias2/Prpf6/Kdm4c/Rnf6/Foxp1/Trim68/Prkcb/Ar | 11 |
| MF | GO:0003729 | mRNA binding | 49/1581 | 314/17474 | 0,000113 | 0,004939 | 0,004407 | Khdrbs2/Sf3b1/Tpr/Dhx9/Rc3h1/Hnrnpu/Angel2/Cdc40/Cpsf6/Cpeb4/Myh10/Dhx33/Luc7l3/Ddx5/Pum2/Rbm25/Larp4b/Ago2/Rbfox2/Rbfox1/Fyttd1/Luc7l/Lrpprc/Cstf2t/Eif3a/Celf2/Rc3h2/Celf1/Myef2/Ythdf3/Dhx36/Rbm15/Ptbp2/Ncbp1/Tardbp/G3bp2/Thoc2l/Zfp638/Tia1/Cpeb1/Tial1/Elavl1/Ddx6/Rbm5/Thoc2/Rbmx/Fmr1/Mecp2/Zc3h12b | 49 |
| MF | GO:0004708 | MAP kinase kinase activity | 7/1581 | 15/17474 | 0,000164 | 0,006402 | 0,005713 | Map2k4/Map2k6/Mapk1/Map3k7/Braf/Map2k1/Pak3 | 7 |
| MF | GO:0004143 | diacylglycerol kinase activity | 6/1581 | 11/17474 | 0,000168 | 0,006496 | 0,005796 | Dgkd/Dgke/Dgkb/Dgkh/Dgkg/Dgki | 6 |
| MF | GO:0004674 | protein serine/threonine kinase activity | 62/1581 | 432/17474 | 0,00018 | 0,006838 | 0,006102 | Clk1/Bmpr2/Pikfyve/Dstyk/Uhmk1/Akt3/Cdk17/Dyrk2/Tbk1/Clk4/Map2k4/Camkk1/Taok1/Map2k6/Rock2/Cdkl1/Map4k5/Rps6ka5/Dapk1/Plk2/Ptk2b/Prkdc/Mapk1/Kalrn/Pdpk1/Mapk14/Map4k3/Map3k2/Csnk1a1/Prkg1/Slk/Camk1d/Acvr1c/Pdk1/Bub1b/Top1/Stk4/Prkab2/Map3k7/Srpk2/Prkag2/Cdkl2/Mapk10/Cdc7/Ksr2/Camkk2/Braf/Aak1/Akap13/Smg1/Prkcb/Nek1/Atm/Map2k1/Csnk1g1/Atr/Pik3cb/Dcaf1/Stk26/Pak3/Wnk3/Cdkl5 | 62 |
| MF | GO:0015085 | calcium ion transmembrane transporter activity | 24/1581 | 124/17474 | 0,000276 | 0,009069 | 0,008092 | Cacna1e/Atp2b1/Slc24a4/Ryr2/Cacna2d3/Cacna1d/Grin2a/Slc8a1/Cacnb2/Cacna1b/Ryr3/Trpc4/Cacna2d1/Stim2/Itpr1/Grm7/Cacna1c/Grin2b/Gpm6a/Atp2c2/Trpc1/Atp2c1/Atp2b3/Trpc5 | 24 |
| MF | GO:0008094 | ATP-dependent activity, acting on DNA | 23/1581 | 117/17474 | 0,00029 | 0,009339 | 0,008334 | Smarcal1/Dhx9/Shprh/Ascc3/Smc6/Rad17/Ercc6/Wapl/Chd8/Nipbl/Chd1/Btaf1/Smc3/Dhx36/Hfm1/Ep400/Smarcad1/Chd2/Rsf1/Chd9/Rad54l2/Smarca1/Atrx | 23 |

GO_SR_mRNA

| ONTOLOGY | ID | Description | GeneRatio | BgRatio | pvalue | p.adjust | qvalue | geneID | Count |
|---|---|---|---|---|---|---|---|---|---|
| CC | GO:0098984 | neuron to neuron synapse | 30/403 | 495/17474 | 1,56E-06 | 0,004009 | 0,003725 | Dst/Inpp4a/Map2/Erbb4/Tnr/Nrcam/Pdpk1/Nrxn1/Dcc/Ptprt/Snx27/Lrrc7/Unc13b/Mpdz/Macf1/Akap9/Grk3/Ppp1r9a/Ptprz1/Itpr1/Grin2b/Eps8/Plekha5/Homer2/Dlg2/Igsf9b/Arhgap32/Fam81a/Dmd/Cdkl5 | 30 |
| CC | GO:0099572 | postsynaptic specialization | 29/403 | 480/17474 | 2,48E-06 | 0,004009 | 0,003725 | Dst/Inpp4a/Map2/Erbb4/Gabrb2/Nrcam/Pdpk1/Dcc/Ptprt/Snx27/Lrrc7/Mpdz/Macf1/Akap9/Gabrb1/Grk3/Ppp1r9a/Ptprz1/Itpr1/Grin2b/Eps8/Plekha5/Homer2/Dlg2/Igsf9b/Arhgap32/Fam81a/Dmd/Cdkl5 | 29 |
| BP | GO:0031503 | protein-containing complex localization | 19/403 | 233/17474 | 2,08E-06 | 0,004009 | 0,003725 | Erbb4/Ipo9/Hnrnpu/Zfp365/Xpo1/Sgcd/Dzip1/Nrxn1/Stx3/Ralgapa2/Ralgapb/Lrrc7/Mdn1/Akap9/Dlg2/Tub/Pcm1/Dync2h1/Ophn1 | 19 |
| BP | GO:0099173 | postsynapse organization | 19/403 | 249/17474 | 5,52E-06 | 0,006674 | 0,006203 | Abl2/Zfp365/Nrcam/Lrrk2/Opa1/Itsn1/Afdn/Stk38/Nrxn1/Ckap5/Snx27/Adgrl3/Epha5/Ppp1r9a/Grin2b/Ube3a/Myo9a/Ophn1/Cdkl5 | 19 |
| BP | GO:0050905 | neuromuscular process | 16/403 | 190/17474 | 8,85E-06 | 0,007516 | 0,006985 | Abl2/Tnr/Agtpbp1/Kcnma1/Atp8a2/Mycbp2/Rbfox2/Nrxn1/Scn1a/Itpr1/Grin2b/Csmd1/Herc1/Myo5a/Mecp2/Dmd | 16 |
| BP | GO:0016441 | post-transcriptional gene silencing | 9/403 | 61/17474 | 1,01E-05 | 0,007516 | 0,006985 | Helz/Pum2/Ago2/Tnrc6b/Focad/Ago3/Ago1/Cnot1/Mecp2 | 9 |

GO_SLM_mRNA

| ONTOLOGY | ID | Description | GeneRatio | BgRatio | pvalue | p.adjust | qvalue | geneID | Count |
|---|---|---|---|---|---|---|---|---|---|
| CC | GO:0062023 | collagen-containing extracellular matrix | 168/3915 | 369/17474 | 2,6356E-23 | 2,1749E-19 | 1,6196E-19 | Col3a1/Fn1/Serpine2/Prelp/Fmod/Prg4/Hmcn1/Lamc2/Angptl1/Tnr/Myoc/Lama2/Lama4/Col13a1/Col18a1/Timp3/Igf1/Ntn4/Lum/Emid1/Efemp1/Col23a1/Sparc/Mfap4/Pmp22/Serpinf1/Serpinf2/Vtn/Lgals9/Col1a1/Clec14a/Smoc1/Fbln5/Nid1/Serpinb9/Serpinb6a/F13a1/Ecm2/Aspn/Omd/Ogn/Tgfbi/Ctsl/Hapln1/Vcan/Thbs4/Ngly1/Nid2/Itih3/Colq/Mmrn2/Lgals3/Ang/Gpc5/Gpc6/Egflam/Angpt1/Col14a1/Lgals1/Fbln1/Plxnb2/Dlg1/Ccdc80/Abi3bp/Adamts1/Runx1/Thbs2/Adamts10/Angptl4/Col11a2/Vegfa/Lama1/Ltbp1/Colec12/Lama3/Lox/Lman1/Efemp2/Acta2/Entpd1/Kazald1/Smc3/Itih2/Itih5/Plxdc2/Entpd2/Egfl7/Lamc3/Angptl2/Itga6/Serping1/F2/Thbs1/Sema6d/Cst3/Tgm2/Matn4/Bmp7/Ctsz/Lama5/Anxa5/Frem2/Bcan/S100a13/S100a3/S100a6/S100a11/F3/Npnt/Ccn1/Col27a1/Tnc/Frem1/Adamtsl1/Col9a2/Col16a1/Tinagl1/Hspg2/Plod1/Vwa1/Sema3c/Fgl2/Reln/Igfbp7/Anxa3/Sparcl1/Eln/Plod3/Gpc2/Col1a2/Ptprz1/Ptn/Rarres2/Loxl3/Anxa4/Vwf/Mgp/Tgfb1/Lgals4/Nphs1/Pcsk6/Mfge8/Ctsc/Serpinh1/Fgfr2/Htra1/Cd151/Ctsd/Igf2/Angpt2/Ncan/Itgb1/Bmper/Vwa5a/Anxa2/Adam10/Col12a1/Plscr1/Plscr2/Col6a6/Lamb2/Rpsa/Ndp/Gpc3/Bgn/Col4a6/Col4a5/Egfl6 | 168 |
| CC | GO:0098857 | membrane microdomain | 152/3915 | 385/17474 | 2,0045E-14 | 7,1919E-12 | 5,3558E-12 | Cavin2/Cflar/Casp8/Erbb4/Irs1/Ptprc/Tnr/Fcgr2b/Fcer1g/Atp1a2/Capn2/Cr1l/Fyn/Smpd2/Gja1/Itgb2/Bsg/Eef2/Stat6/Egfr/Pld2/Myo1c/Cavin1/Pecam1/Adam17/Ripk1/Ptch1/Slc6a3/F2r/Il6st/Itga1/Vcl/Bmpr1a/Tnfrsf10b/Ednrb/Angpt1/Has2/Nfam1/Pacsin2/Mlc1/Tuba1c/Emp2/Dlg1/Cxadr/Ezr/H2-D1/Trem2/Slc5a7/Npc1/Ctnna1/Cd14/Adrb2/Slc22a6/Ahnak/Jak2/Fas/Myof/Sorbs1/Stom/Tfpi/Ctnnd1/Lrp4/Slc1a2/Cd44/Kif18a/Mall/Hck/Sdc4/Ptgis/Pag1/P2ry12/Tlr2/Kirrel/Fcgr1/Gnai3/Bcl10/Lyn/Tgfbr1/Abca1/Tlr4/Tek/Plpp3/Lrp8/Slc2a1/Pdpn/Tnfrsf1b/Prkcz/Nos3/Tlr1/Tlr6/Kdr/Grk3/Selplg/Ptpn11/P2rx7/Stx2/Lat2/Hmgb1/Cav2/Cav1/Smo/Adcyap1r1/Abcg2/Dysf/Ret/Tnfrsf1a/Lrp6/Ldhb/Ehd2/Lipe/Nphs1/Fxyd1/Igf1r/Iqgap1/Serpinh1/Cavin3/Tpp1/Gprc5b/Cln3/Mapk3/Itgam/Ctsd/Ano1/Insr/Erlin2/Dlc1/Casp3/Cpe/Slc27a1/Ednra/Pllp/Cdh1/Itgb1/Birc3/Icam1/Ldlr/Ppp2r1b/Anxa2/Nedd4/Plscr1/Plscr2/Atp1b3/Ephb1/Gnai2/Tgfbr2/Plcd1/Ctnnb1/Lamp2/Dmd/Atp7a/Btk/Gpm6b | 152 |
| CC | GO:0009925 | basal plasma membrane | 115/3915 | 281/17474 | 1,932E-12 | 3,4658E-10 | 2,581E-10 | Slc40a1/Erbb4/Slc41a1/Vangl2/Kcnj10/Slc30a1/Cr1l/Slc26a10/Erbb3/Egfr/Slc22a5/Slc22a21/Slc22a4/Slc47a1/Slc13a5/Myo1c/Slc46a1/Abcc3/Ace/Kcnj16/Numb/Dsp/Marveld2/Erbin/Slc7a7/Abcc4/Slc1a3/Mlc1/Rapgef3/Abcc5/Dlg1/Tfrc/Casr/Cxadr/Ezr/Slc29a1/Epcam/Aqp4/Slc14a1/Slc22a8/Slc22a6/Slc3a2/Slc16a12/Ide/Entpd1/Itga6/Rapgef4/Slc43a3/Cd44/Lin7c/Slc12a6/Slc13a3/Car2/P2ry12/P2ry1/Car14/Slc16a1/Stxbp3/Slc39a8/Mttp/Slc26a7/Tgfbr1/Abca1/Tek/Plpp3/Slc2a1/Cldn19/Epb41/Ldlrap1/Pdpn/Cd38/Slc4a4/Pkd2/Stx2/Cav1/Nod1/Vamp8/Tgfa/Slc6a6/Slc6a13/Slco1c1/Slco1a4/C5ar2/C5ar1/Slc1a5/Ceacam1/Abcc6/Tjp1/Iqgap1/Arrb1/Slco2b1/P2ry6/P2ry2/Stx4a/Cd81/Atp7b/Adam9/Hpgd/Slc27a1/Nod2/Cdh1/Ldlr/Anxa2/Aqp9/Atp1b3/Slco2a1/Slc38a3/Slc26a6/Pth1r/Itga9/Ctnnb1/Shroom4/Msn/Heph/Atp7a | 115 |
| CC | GO:0045178 | basal part of cell | 122/3915 | 304/17474 | 2,0517E-12 | 3,5273E-10 | 2,6268E-10 | Slc40a1/Erbb4/Slc41a1/Vangl2/Kcnj10/Slc30a1/Cr1l/Slc26a10/Erbb3/Egfr/Slc22a5/Slc22a21/Slc22a4/Slc47a1/Slc13a5/Myo1c/Slc46a1/Abcc3/Ace/Kcnj16/Numb/Hfe/Dsp/Edn1/Marveld2/Erbin/Itga1/Slc7a7/Abcc4/Slc1a3/Mlc1/Rapgef3/Abcc5/Dlg1/Tfrc/Casr/Phldb2/Cxadr/Ezr/Slc29a1/Epcam/Aqp4/Slc14a1/Slc22a8/Slc22a6/Slc3a2/Slc16a12/Ide/Entpd1/Itga6/Rapgef4/Slc43a3/Cd44/Lin7c/Slc12a6/Slc13a3/Car2/Clrn1/P2ry12/P2ry1/Car14/Slc16a1/Stxbp3/Slc39a8/Mttp/Slc26a7/Tgfbr1/Abca1/Tek/Plpp3/Slc2a1/Cldn19/Epb41/Ldlrap1/Pdpn/Cd38/Slc4a4/Pkd2/Stx2/Cav1/Nod1/Vamp8/Tgfa/Slc6a6/Slc6a13/Slco1c1/Slco1a4/C5ar2/C5ar1/Slc1a5/Ceacam1/Abcc6/Tjp1/Iqgap1/Prcp/Arrb1/Slco2b1/P2ry6/P2ry2/Stx4a/Cd81/Atp7b/Adam9/Hpgd/Slc27a1/Nod2/Cdh1/Ldlr/Anxa2/Aqp9/Atp1b3/Slco2a1/Slc38a3/Slc26a6/Pth1r/Clasp2/Itga9/Ctnnb1/Shroom4/Msn/Heph/Atp7a | 122 |
| CC | GO:0030055 | cell-substrate junction | 83/3915 | 186/17474 | 1,2253E-11 | 1,7739E-09 | 1,321E-09 | Dst/Map4k4/Tns1/Epb41l5/Ptprc/Hmcn1/Capn2/Lims1/Itgb2/Tns3/Alox8/Lasp1/Grb7/Itga2b/Itgb3/Adam17/Klf11/Syne2/Itgb8/Nedd9/Erbin/Itga1/Fam107a/Vcl/Fermt2/Sorbs3/Lcp1/Parvg/Senp1/Lima1/Tns2/Itga5/Lpp/Itgb5/Phldb2/Ezr/Mdc1/Lama3/Lims2/Fermt3/Lpxn/Pgm5/Jak2/Sorbs1/Nrap/Rsu1/Aif1l/Itga6/Itgav/Hck/Mapre1/Sdc4/Bcar3/Synpo2/Nexn/Cpne3/Tln1/Asap3/Afap1/Arhgap24/Atp6v0a2/Tes/Cav2/Cav1/Vasp/Nphs1/Fes/Iqgap1/Ilk/Parva/Mapk3/Tgfb1i1/Thsd1/Dlc1/Palld/Cyba/Itgb1/Rdx/Clasp2/Limd1/Shroom4/Dmd/Msn | 83 |
| CC | GO:0001726 | ruffle | 68/3915 | 153/17474 | 1,0553E-09 | 8,7084E-08 | 6,4852E-08 | Map2/Vil1/Epb41l5/Arhgap18/S100b/Appl2/Egfr/Plek/Myo1c/Itgb3/Pecam1/Adam17/Fam107a/Spata13/Lcp1/Myo10/Twf1/Lima1/Itga5/Ezr/Fgd2/Aif1/Cd2ap/Jcad/Rock1/Psd2/Pdgfrb/Kank1/Psd4/Aif1l/Itgav/Kif18a/Knstrn/Trpm7/S100a6/S100a11/Tln1/Macf1/Wasf2/Asap3/Pdpn/Cdk6/Hip1r/Ptprz1/Podxl/Frmd4b/Mtmr14/Eps8/Eps8l1/Akt2/Rinl/Iqgap1/Eps8l2/Dlc1/Palld/Plcg2/Itgb1/Dnm2/Tirap/Rdx/Anxa2/Tmod3/Myo6/Rhoa/Clasp2/Mtm1/Amot/Bmx | 68 |
| CC | GO:0005901 | caveola | 49/3915 | 97/17474 | 1,1741E-09 | 9,4065E-08 | 7,005E-08 | Cavin2/Erbb4/Irs1/Atp1a2/Smpd2/Pld2/Cavin1/Ptch1/Slc6a3/F2r/Bmpr1a/Pacsin2/Mlc1/Ctnna1/Adrb2/Slc22a6/Jak2/Myof/Sorbs1/Tfpi/Ctnnd1/Kif18a/Hck/Ptgis/P2ry12/Tgfbr1/Lrp8/Slc2a1/Nos3/Cav2/Cav1/Smo/Adcyap1r1/Lrp6/Ehd2/Lipe/Fxyd1/Igf1r/Cavin3/Cln3/Mapk3/Insr/Dlc1/Slc27a1/Cdh1/Ldlr/Atp1b3/Tgfbr2/Ctnnb1 | 49 |
| CC | GO:0031252 | cell leading edge | 144/3915 | 408/17474 | 1,373E-09 | 1,0895E-07 | 8,1132E-08 | Dst/Map2/Vil1/Epb41l5/Srgap2/Cdc42bpa/Arhgap18/Ctnna3/Adora2a/S100b/Appl2/Kitl/Myo1g/Egfr/Plek/Ccdc88a/Pdlim4/Pld2/Myo1c/Abi3/Itgb3/Pecam1/P4hb/Adam17/Syne2/Carmil1/Dusp22/Fam107a/Spata13/Lcp1/Myo10/Pabpc1/Rac2/Twf1/Rapgef3/Lima1/Itga5/Fgd4/Itgb5/Mylk/Arhgap31/Phldb2/Abi3bp/Itsn1/Ezr/Fgd2/Aif1/Cd2ap/Ptprm/Ston1/Jcad/Rock1/Slc39a6/Ctnna1/Psd2/Pdgfrb/Coro1b/Dock8/Kank1/Plce1/Ablim1/Vim/Psd4/Tsc1/Aif1l/Dpp4/Itgav/Ctnnd1/Slc1a2/Cd44/Kif18a/Knstrn/Trpm7/Lamp5/Pld1/Pik3ca/Clrn1/Rab13/S100a6/S100a11/Unc5c/Pkn2/Wls/Tln1/Macf1/Phactr4/Wasf2/Asap3/Pdpn/Prkcz/Cdk6/Gabrg1/Ptpn13/Pkd2/Hip1r/Stx2/Gper1/Wasf3/Sgce/Ptprz1/Podxl/Dysf/Antxr1/Frmd4b/Mtmr14/Eps8/Eps8l1/Vasp/Akt2/Rinl/Cyfip1/Cib1/Iqgap1/Inppl1/Ilk/Parva/Stx4a/Rnh1/Eps8l2/Insr/Dlc1/Fat1/Palld/Adgre5/Cdh1/Plcg2/Itgb1/Amotl1/Dnm2/Tirap/Rdx/Anxa2/Tmod3/Myo6/Rhoa/Nradd/Clasp2/Ctnnb1/Mtm1/Dmd/Gdpd2/Atp7a/Amot/Bmx | 144 |
| CC | GO:0009897 | external side of plasma membrane | 150/3915 | 430/17474 | 1,58E-09 | 1,1961E-07 | 8,9076E-08 | Il1r1/Serpine2/Tnfrsf11a/Ptprc/Cfh/Fcgr2b/Fcgr3/Fcer1g/Cd84/Capn2/Cr1l/Vsir/Itgb2/Icosl/Stab2/Tcn2/Tnfrsf13b/Atp1b2/Tnfsf13/Asgr1/Clec10a/Gp1ba/Car4/Itga2b/Itgb3/Ace/Pecam1/P4hb/Sdc1/Clec14a/Gm4787/Hfe/Ctsl/Il6st/Itga1/Bmpr1a/Lgals3/Abcc4/Ghr/Osmr/Lifr/Il7r/Ly6a/Ly6c1/Csf2rb2/Csf2rb/Itga5/Tfrc/Heg1/Cd86/Cd80/Cd200r1/H2-K1/H2-Aa/H2-Eb1/H2-D1/H2-Q4/H2-T23/H2-T22/Adgre1/Aqp4/Cd14/Cd74/Cd274/Fas/Ide/Entpd1/Gfra1/Eng/Cd302/Ly75/Itga6/Itgav/F2/Cd44/Cd59a/Thbs1/B2m/Thbd/Cd40/Anxa5/P2ry12/Tlr2/Cd1d1/Il6ra/Rorc/Ctsk/Fcgr1/S1pr1/Vcam1/Enpep/Il11ra1/Abca1/Tlr4/Lepr/Skint6/Csf3r/Pdpn/Tnfrsf14/Abcb1b/Adgra3/Pdgfra/Kdr/Antxr2/Tgfbr3/P2rx7/Abcg2/Antxr1/Cxcl12/Clec4a3/Clec4a2/Cd163/Lag3/Cd9/Vwf/Clec2d/Clec7a/Mill2/Apoe/Ceacam1/Cd33/Siglece/Fcgrt/Flt3l/Mfge8/Anpep/Neu3/Il4ra/Il21r/Itgal/Itgam/Ano1/Insr/Lamp1/Adam9/Clcn3/Adgre5/Cdh5/Itgb1/Tmem123/Icam1/Ldlr/Mcam/Tgfbr2/Itga9/Ccr2/Ccr5/Il13ra1/Il2rg/Amot | 150 |

| | | | | | | | | | |
|---|---|---|---|---|---|---|---|---|---|
| CC | GO:0045177 | apical part of cell | 160/3915 | 484/17474 | 3,2273E-08 | 1,6338E-06 | 1,2167E-06 | Ecrg4/Myo1b/Pard3b/Fn1/Igfbp2/Kcne4/Aspm/Ddr2/Vangl2/Kcnj10/Kcnk2/Hsd11b1/Akap7/Gja1/Nedd1/Erbb3/Egfr/Slc22a5/Slc22a21/Slc22a4/Slc47a1/Atp1b2/Slc46a1/Car4/Itgb3/Adam17/Numb/Lgmn/Hfe/Ptch1/Ctsl/Ocln/Marveld2/Btd/Gjb6/Abcc4/Dab2/Osmr/Pdzd2/Mtdh/Fzd6/Mlc1/Rapgef3/Pfkm/Tmem114/Emp2/Abcc5/Casr/Slc15a2/Upk1b/Cldnd1/Ezr/Itpr3/Cyp4f15/Cyp4f14/Adgrf5/Slc29a1/Slc5a7/Epcam/Pdgfrb/Adrb2/Atp8b1/Pard6g/Tcirg1/Slc22a8/Slc3a2/Trpm6/Fas/Notch1/Dpp4/Cybrd1/Rapgef4/Cd44/Jag1/Car2/Pld1/Slc7a11/P2ry1/Il6ra/Car14/Slc16a1/Dram2/Ahcyl1/Gpsm2/Stxbp3/Vcam1/Enpep/Slc39a8/Tgfbr1/Epb41l4b/Mpdz/Tek/Patj/Dab1/Slc2a1/Svbp/Clic4/Pdpn/Slc2a5/Vamp3/Prkcz/Abcb1a/Abcb1b/Nos3/Prom1/Spp1/P2rx4/Hip1r/Cav1/Podxl/Abcg2/Anxa4/Slc6a6/Cd9/Mgst1/C5ar2/C5ar1/Ceacam1/Fxyd1/Abcc6/Tjp1/Iqgap1/Homer2/Myo7a/Slco2b1/P2ry6/P2ry2/Insc/Stx4a/Ifitm3/Cd81/Ano1/Atp7b/Plat/Nrg1/Pcm1/Fat1/Clcn3/Bst2/Cdh1/Cyba/Kcnk1/Pard3/Amotl1/Ldlr/Il18/Rdx/Myo6/Atp1b3/Slc38a3/Slc26a6/Pth1r/Ctnnb1/Slc6a20a/Shroom4/Clcn5/Msn/Slc16a2/Atp7a/Sytl4 | 160 |
| CC | GO:0030139 | endocytic vesicle | 80/3915 | 205/17474 | 5,3491E-08 | 2,5223E-06 | 1,8784E-06 | Lmbrd1/Rab23/Slc11a1/Srgap2/Rab7b/Appl2/Egfr/Irgm1/Myo1c/Stxbp4/Rab5c/Cdc42ep4/Heatr5a/Ston2/Rin3/Pld4/Syk/Ocln/Scara5/Itsn1/H2-K1/Tap1/H2-D1/H2-T23/Cd2ap/Slc5a7/Rab31/Ston1/Nrxn1/Kif5b/Tcirg1/Unc93b1/Mpeg1/Vim/Gapvd1/Dpp4/Ehd4/B2m/Kif16b/Rin2/Rab22a/Pld1/Rab13/Ctss/Rap1a/Stxbp3/Abca1/Csf3r/Pla2g5/Vamp3/Tlr1/Scarb2/Anxa3/Smo/Nod1/Vamp8/Capg/Dysf/Rab43/Sec61a1/Ehd2/Rinl/Ftl1/Cemip/Stx4a/Lamp1/Clcn3/Nod2/Dnm2/Tirap/Myo1e/Myo6/Slc9a9/Was/Cybb/Lamp2/Amot/Sh3kbp1/Rab9/Tlr7 | 80 |
| CC | GO:0001725 | stress fiber | 42/3915 | 88/17474 | 1,4077E-07 | 5,6943E-06 | 4,2405E-06 | Dst/Mlph/Vangl2/Cnn2/Pdlim4/Daam1/Fam107a/Vcl/Fermt2/Myh6/Lcp1/Lima1/Myh11/Lpp/Mylk/Myl12a/Coro1b/Pgm5/Acta2/Sorbs1/Ablim1/Nebl/Myl9/Synpo2/Pdlim5/Tek/Prkcz/Limch1/Septin11/Ptpn11/Ilk/Dlc1/Pdlim3/Palld/Tpm4/Fhod1/Cyba/Septin7/Pstpip1/Anxa2/Shroom4/Amot | 42 |
| CC | GO:0097517 | contractile actin filament bundle | 42/3915 | 88/17474 | 1,4077E-07 | 5,6943E-06 | 4,2405E-06 | Dst/Mlph/Vangl2/Cnn2/Pdlim4/Daam1/Fam107a/Vcl/Fermt2/Myh6/Lcp1/Lima1/Myh11/Lpp/Mylk/Myl12a/Coro1b/Pgm5/Acta2/Sorbs1/Ablim1/Nebl/Myl9/Synpo2/Pdlim5/Tek/Prkcz/Limch1/Septin11/Ptpn11/Ilk/Dlc1/Pdlim3/Palld/Tpm4/Fhod1/Cyba/Septin7/Pstpip1/Anxa2/Shroom4/Amot | 42 |
| CC | GO:0043235 | receptor complex | 132/3915 | 397/17474 | 3,5445E-07 | 1,2394E-05 | 9,2296E-06 | Ly96/Erbb4/Irs1/Gpr37l1/Cd247/Ddr2/Cr1l/Itgb2/Ptprb/Erbb3/Egfr/Flt4/Pmp22/Chrnb1/Gp1ba/Traf4/Ramp2/Nbr1/Itga2b/Itgb3/Cacng4/Gprc5c/Grin2c/Traf3/Itgb8/Hfe/Ripk1/Syk/Ntrk2/Il6st/Itga1/Bmpr1a/Ghr/Osmr/Lifr/Csf2rb2/Csf2rb/Plxnb2/Acvrl1/Gpr84/Itga5/Tfrc/Itgb5/Gpr156/Cd200r1/Grik1/Itpr3/Cd14/Ticam2/Cd74/Pdgfrb/Csf1r/Adrb2/Lrp5/Jak2/Chuk/Pkd2l1/Gfra1/Apbb1ip/Notch1/Eng/Traf1/Pla2r1/Itga6/Itgav/Calcrl/Nr1h3/Traf6/Cd44/B2m/Mertk/Cd40/Tlr2/Il6ra/Arnt/Notch2/Mttp/Bmpr1b/Lyn/Il11ra1/Tgfbr1/Tlr4/Tek/Lepr/Lrp8/Tie1/Fgfr3/Tlr1/Tlr6/Gabrg1/Gabrb1/Pdgfra/Kdr/Tgfbr3/Diablo/Ephb4/Flt1/Tril/Adcyap1r1/Grid2/Gp9/Ret/Ptpn6/Tnfrsf1a/Eps8/Itpr2/Ceacam1/Tgfb1/Axl/Igf1r/Gprc5b/Il4ra/Itgal/Itgam/Fgfr2/Insr/Ikbkb/Itgb1/Ldlr/Htr3a/Ryk/Plxnb1/Pth1r/Tgfbr2/Itga9/Acvr2b/Il13ra1/Gabrq/Flna/Il2rg/Kctd12b/Tlr7 | 132 |
| CC | GO:0098858 | actin-based cell projection | 82/3915 | 221/17474 | 4,5879E-07 | 1,5328E-05 | 1,1414E-05 | Myo1b/Map2/Vil1/Kcnj10/Utrn/Pcdh15/Kitl/Lyz2/Myo1g/Myo1c/Aoc3/Itgb3/Icam2/Dcxr/Syne2/Ninj1/Ctsl/Spata13/Kif13b/Lcp1/Myo10/Angpt1/Triobp/Twf1/Rapgef3/Fgd4/Dlg1/Cxadr/Ezr/Myo1f/Clic5/Atp8b1/Cd302/Itga6/Itgav/Cd44/Car2/Clrn1/Kcnn3/Rhoc/Vcam1/Cdc14a/Mttp/Unc5c/Tek/Clic4/Pdpn/Prom1/Pdgfra/Ptprz1/Podxl/Antxr1/Slc6a6/Eps8/Ttyh1/Ptprh/Vasp/Tgfb1/Cyfip1/Cib1/Homer2/Myo7a/Fchsd2/Stard10/Eps8l2/Fat1/Palld/Bbs2/Cdh1/Calb2/Itgb1/Rdx/Myo1e/Nedd4/Myo6/Ephb1/Ctnnb1/Fgf13/Mtm1/Dmd/Msn/Atp7a | 82 |
| CC | GO:0032587 | ruffle membrane | 36/3915 | 74/17474 | 6,0112E-07 | 1,9453E-05 | 1,4486E-05 | Map2/Epb41l5/Appl2/Egfr/Plek/Myo1c/Itgb3/Adam17/Fam107a/Spata13/Twf1/Itga5/Aif1/Jcad/Psd2/Kank1/Psd4/Aif1l/Itgav/Macf1/Pdpn/Hip1r/Ptprz1/Eps8/Eps8l1/Akt2/Eps8l2/Dlc1/Plcg2/Itgb1/Dnm2/Tirap/Myo6/Rhoa/Clasp2/Bmx | 36 |
| CC | GO:0016324 | apical plasma membrane | 123/3915 | 378/17474 | 3,013E-06 | 7,6035E-05 | 5,6624E-05 | Ecrg4/Pard3b/Fn1/Igfbp2/Kcne4/Aspm/Ddr2/Vangl2/Kcnj10/Kcnk2/Akap7/Gja1/Erbb3/Egfr/Slc22a5/Slc22a21/Slc22a4/Slc47a1/Atp1b2/Slc46a1/Car4/Itgb3/Adam17/Ocln/Marveld2/Gjb6/Abcc4/Dab2/Osmr/Mtdh/Fzd6/Mlc1/Pfkm/Tmem114/Emp2/Abcc5/Casr/Slc15a2/Upk1b/Cldnd1/Ezr/Cyp4f15/Cyp4f14/Slc29a1/Slc5a7/Epcam/Pdgfrb/Adrb2/Atp8b1/Pard6g/Tcirg1/Slc22a8/Slc3a2/Trpm6/Fas/Notch1/Dpp4/Cybrd1/Rapgef4/Cd44/Jag1/Pld1/P2ry1/Il6ra/Car14/Slc16a1/Dram2/Ahcyl1/Stxbp3/Vcam1/Enpep/Slc39a8/Tgfbr1/Mpdz/Tek/Slc2a1/Pdpn/Slc2a5/Vamp3/Prkcz/Abcb1a/Abcb1b/Prom1/Hip1r/Cav1/Podxl/Abcg2/Anxa4/Slc6a6/Cd9/Ceacam1/Fxyd1/Abcc6/Tjp1/Iqgap1/Myo7a/Slco2b1/P2ry6/P2ry2/Stx4a/Cd81/Ano1/Nrg1/Fat1/Clcn3/Bst2/Cyba/Kcnk1/Pard3/Amotl1/Il18/Rdx/Atp1b3/Slc38a3/Slc26a6/Pth1r/Ctnnb1/Slc6a20a/Shroom4/Msn/Slc16a2/Atp7a/Sytl4 | 123 |
| CC | GO:0034451 | centriolar satellite | 46/3915 | 112/17474 | 7,1096E-06 | 0,00015771 | 0,00011745 | Cspp1/Zbed6/Flii/Pik3r5/Chd3/Spag5/Spag9/Ift43/Pxk/Ccdc66/Dlgap5/Dzip1/Pdzd2/Tbc1d31/Cyth4/Nlrc3/Ccdc14/Cd86/Tap1/Cd2ap/Kif5b/Prkcq/Cntrl/Nek6/Knstrn/Rbm39/Fnip2/Odf2l/Tek/Patj/Tacc3/Ccdc18/Podxl/Dysf/Plk1/Il4ra/Hook3/Pcm1/Bbs9/Tex9/Cep162/Ccdc13/Wdr13/G6pdx/Klhl4/Mid1 | 46 |
| CC | GO:0005777 | peroxisome | 56/3915 | 145/17474 | 7,3029E-06 | 0,00016113 | 0,00011999 | Idh1/Acsl3/Mtarc2/Ddo/Pex13/Slc22a21/Acsl6/Aldh3a2/Nos2/Nbr1/Acox1/Abcd4/Idi1/Eci2/Tkt/Hacl1/Dhrs4/Ephx2/Lacc1/Vwa8/Rida/Serhl/Abcd2/Marf1/Ehhadh/Nudt12/Xdh/Sting1/Hsd17b4/Isoc1/Plaat3/Ide/Phyh/Vim/Acbd5/Cat/Mavs/Pex2/Abcd3/Hsdl2/Scp2/Prdx1/Hmgcl/Crot/Acox3/Pxmp2/Mvk/Cav1/Gstk1/Depp1/Mgst1/Ech1/Cd33/Idh2/Nudt7/Acad11 | 56 |
| CC | GO:0042579 | microbody | 56/3915 | 145/17474 | 7,3029E-06 | 0,00016113 | 0,00011999 | Idh1/Acsl3/Mtarc2/Ddo/Pex13/Slc22a21/Acsl6/Aldh3a2/Nos2/Nbr1/Acox1/Abcd4/Idi1/Eci2/Tkt/Hacl1/Dhrs4/Ephx2/Lacc1/Vwa8/Rida/Serhl/Abcd2/Marf1/Ehhadh/Nudt12/Xdh/Sting1/Hsd17b4/Isoc1/Plaat3/Ide/Phyh/Vim/Acbd5/Cat/Mavs/Pex2/Abcd3/Hsdl2/Scp2/Prdx1/Hmgcl/Crot/Acox3/Pxmp2/Mvk/Cav1/Gstk1/Depp1/Mgst1/Ech1/Cd33/Idh2/Nudt7/Acad11 | 56 |
| CC | GO:0098636 | protein complex involved in cell adhesion | 28/3915 | 57/17474 | 8,3029E-06 | 0,0001775 | 0,00013219 | Tnr/Lama2/Itgb2/Pmp22/Vtn/Itga2b/Itgb3/Itgb8/Nid1/Itga1/Plau/Lgals1/Itga5/Itgb5/Cd80/Jam2/Lama1/Nrxn1/Itga6/Itgav/Lyn/Tnc/Plaur/Itgal/Itgam/Itgb1/Lamb2/Itga9 | 28 |
| CC | GO:0005912 | adherens junction | 57/3915 | 151/17474 | 1,3573E-05 | 0,0002686 | 0,00020002 | Pard3b/Cdh20/Cdh19/Epb41l5/Hmcn1/Ctnna3/Frs2/Pdlim4/Alox8/Cdc42ep4/Dsp/Vcl/Fermt2/Cdh9/Mtss1/Klhl24/Dlg1/Nectin3/Cxadr/Ezr/Vegfa/Ptprm/Jcad/Mpp7/Lama3/Dsc2/Ctnna1/Pgm5/Sorbs1/Frmd4a/Notch1/Itga6/Ctnnd1/Lin7c/Snap23/Jag1/Pdlim5/Lyn/Tln1/Frmd4b/Cdca3/Nectin2/Ceacam1/Tjp1/Pdlim3/Cdh5/Cdh1/Pard3/Itgb1/Esam/Rdx/Myo1e/Ctnnb1/Limd1/Shroom4/Cd99l2/Msn | 57 |
| CC | GO:0097386 | glial cell projection | 22/3915 | 43/17474 | 3,4199E-05 | 0,00060301 | 0,00044906 | Glul/Kcnj10/Kcnk2/Fyn/Atp1b2/Gfap/Lgals3/Gjb2/Lcp1/Mlc1/Ezr/Aif1/Aqp4/Slc1a2/Slc7a11/Grm3/Wasf3/Sirt2/Mt3/Adgrg1/Itgb1/Dmd | 22 |
| CC | GO:0022626 | cytosolic ribosome | 40/3915 | 100/17474 | 5,6237E-05 | 0,00087067 | 0,00064839 | Rpl7/Rpl37a/Rps15/Rpl41/Rpl26/Rpl23a/Rps24/Rpl37/Rpl8/Apod/Rps2/Rps28/Rps18/Rps14/Rpl7a/Rpl35/Rps21/Rps20/Rps8/Rpl22/Ppargc1a/Rplp0/Rpl6/Rpl21/Rpl32/Rps9/Rps5/Rps19/Rpl13a/Rpl18/Rplp2/Rpl18a/Rplp1/Rpl4/Rps27l/Rpl29/Rps27rt/Rpsa/Rpl14/Rpl10 | 40 |
| CC | GO:0030670 | phagocytic vesicle membrane | 14/3915 | 23/17474 | 7,9452E-05 | 0,0011511 | 0,00085722 | Slc11a1/Appl2/H2-K1/Tap1/H2-D1/H2-T23/Rab31/Tcirg1/B2m/Anxa3/Sec61a1/Lamp1/Slc9a9/Lamp2 | 14 |
| CC | GO:0098802 | plasma membrane signaling receptor complex | 63/3915 | 185/17474 | 0,00017976 | 0,00229235 | 0,00170711 | Irs1/Cd247/Itgb2/Erbb3/Pmp22/Chrnb1/Gp1ba/Traf4/Ramp2/Itga2b/Itgb3/Cacng4/Grin2c/Traf3/Itgb8/Hfe/Syk/Il6st/Itga1/Osmr/Csf2rb2/Csf2rb/Acvrl1/Itga5/Tfrc/Itgb5/Gpr156/Grik1/Ticam2/Jak2/Chuk/Apbb1ip/Traf1/Itga6/Itgav/Calcrl/Traf6/B2m/Cd40/Tlr2/Il6ra/Lyn/Tgfbr1/Tlr1/Tlr6/Diablo/Grid2/Gp9/Ptpn6/Eps8/Ceacam1/Tgfb1/Igf1r/Itgal/Itgam/Insr/Ikbkb/Itgb1/Htr3a/Tgfbr2/Itga9/Acvr2b/Flna | 63 |

| | | | | | | | | | |
|---|---|---|---|---|---|---|---|---|---|
| CC | GO:0005769 | early endosome | 102/3915 | 334/17474 | 0,00031324 | 0,0035361 | 0,00263333 | Myo1b/Wdfy1/Bok/Rab29/Rcsd1/Slc30a10/Gja1/Washc4/Rab21/Bloc1s1/Egfr/Havcr2/Pdlim4/Mmgt2/Ankfy1/Aoc3/Rnd2/Tbc1d16/Snx6/Zfyve26/Numb/Rin3/Pld4/Hfe/Ntrk2/Zfyve16/F2r/Pxk/Ap3m1/Wdfy2/Mlc1/Litaf/Atp13a4/Tfrc/Snx4/Fgd2/H2-K1/H2-D1/H2-T22/Slc5a7/Rab31/Ticam2/Cd274/Gpr107/Ehd4/Pla2g4e/Usp8/Lamp5/Kif16b/Snx5/Rab22a/Atp11b/Cd1d1/Snx27/Ctss/Rap1a/Vcam1/Sh3glb1/Wls/Plekhf2/Wasf2/Ldlrap1/Vamp3/Steap2/Pi4k2b/Kdr/Tpcn1/Gper1/Snx8/Hmgb1/Samd9l/Wdr91/Vamp8/Dysf/Marchf8/Ret/Lrp6/Ehd2/Apoe/Ankrd27/Plekhf1/Hps5/St8sia2/Ap3s2/Picalm/Neu3/P2ry2/Cln3/Mapk3/Ifitm2/Ifitm1/Ifitm3/Clcn3/Ldlr/Anxa2/Tbc1d2b/Slc9a9/Ephb1/Cmtm6/Clcn5/Mmgt1/Atp7a | 102 |
| CC | GO:0043296 | apical junction complex | 51/3915 | 146/17474 | 0,0003587 | 0,00394139 | 0,00293515 | Pard3b/F11r/Pmp22/Traf4/Ocln/Marveld2/Cldn10/Mtdh/Cldn5/Dlg1/Nectin3/Cxadr/Jam2/Epcam/Mpp7/Ctnna1/Pard6g/Tjp2/Frmd4a/Ctnnd1/Lin7c/Rab13/Magi3/Usp53/Pkn2/Tgfbr1/Epb41l4b/Mpdz/Patj/Cldn19/Prkcz/Mxra8/Adcyap1r1/Frmd4b/Vasp/Nectin2/Lsr/Tjp1/Arhgap17/Atp7b/Cdh5/Cdh1/Pard3/Amotl1/Esam/Cgnl1/Rhoa/Ctnnb1/Shroom4/Rap2c/Amot | 51 |
| CC | GO:0044291 | cell-cell contact zone | 34/3915 | 89/17474 | 0,000541 | 0,00552513 | 0,00411456 | Dst/Atp1a2/Gja1/Ctnna3/Pecam1/Kcnj2/Dsp/Vcl/Pcdh9/Baiap2l2/Nectin3/Cxadr/Jam2/Dsc2/Ctnna1/Ahnak/Tjp2/Pgm5/Nrap/Pik3ca/Anxa5/Gja5/Camk2d/Slc31a1/Slc2a1/Vamp5/Nectin2/Fxyd1/Tjp1/Fhod1/Itgb1/Ctnnb1/Rap2c/Fgf13 | 34 |
| CC | GO:0031256 | leading edge membrane | 53/3915 | 157/17474 | 0,00070804 | 0,0068577 | 0,00510693 | Map2/Epb41l5/Adora2a/Appl2/Myo1g/Egfr/Plek/Myo1c/Itgb3/Adam17/Syne2/Fam107a/Spata13/Twf1/Itga5/Aif1/Jcad/Slc39a6/Psd2/Dock8/Kank1/Psd4/Aif1l/Itgav/Slc1a2/Cd44/Lamp5/Wls/Macf1/Pdpn/Gabrg1/Hip1r/Gper1/Sgce/Ptprz1/Antxr1/Eps8/Eps8l1/Akt2/Stx4a/Eps8l2/Insr/Dlc1/Adgre5/Plcg2/Itgb1/Dnm2/Tirap/Myo6/Rhoa/Nradd/Clasp2/Bmx | 53 |
| CC | GO:0043230 | extracellular organelle | 38/3915 | 105/17474 | 0,0008993 | 0,00837586 | 0,0062375 | Fn1/Qsox1/Pmel/Cd63/Tmem98/Car4/Itga2b/Ace/Gprc5c/Dicer1/Has2/Vasn/Tfrc/Cd86/Clic1/Iigp1/Ahnak/Cd274/Ide/Arrdc1/Gbp7/Gbp3/Gbp2/Alpl/Sri/Prom1/Gbp9/Gbp6/Podxl/Cd9/Apoe/Anpep/Gprc5b/Cd81/Icam1/Anxa2/Acy1/Lamp2 | 38 |
| CC | GO:0065010 | extracellular membrane-bounded organelle | 38/3915 | 105/17474 | 0,0008993 | 0,00837586 | 0,0062375 | Fn1/Qsox1/Pmel/Cd63/Tmem98/Car4/Itga2b/Ace/Gprc5c/Dicer1/Has2/Vasn/Tfrc/Cd86/Clic1/Iigp1/Ahnak/Cd274/Ide/Arrdc1/Gbp7/Gbp3/Gbp2/Alpl/Sri/Prom1/Gbp9/Gbp6/Podxl/Cd9/Apoe/Anpep/Gprc5b/Cd81/Icam1/Anxa2/Acy1/Lamp2 | 38 |
| CC | GO:0042383 | sarcolemma | 55/3915 | 166/17474 | 0,00094051 | 0,00868127 | 0,00646494 | Dst/Atp1a2/Slc30a1/Utrn/Akap7/Lama2/Bsg/Sgcd/Alox12/Pld2/Car4/Cacng4/Kcnj2/Vcl/Slc38a2/Dlg1/Ezr/Aqp4/Dtna/Adrb2/Plcb3/Ahnak/Pgm5/Fas/Prkcq/Cd59a/Capn3/Anxa5/Vcam1/Camk2d/Slc2a1/Slc2a5/Sri/Nos3/Sgcb/Slc8b1/Cav2/Cav1/Dysf/Alox5/Abcc9/Sspn/Rtn2/Fxyd1/Igf1r/Cib1/Stx4a/Lamp1/Ednra/Itgb1/Rdx/Anxa2/Bgn/Dmd/Msn | 55 |
| BP | GO:0045785 | positive regulation of cell adhesion | 198/3915 | 498/17474 | 8,2829E-19 | 1,7088E-15 | 1,2725E-15 | Map4k4/Il1rl2/Fn1/Igfbp2/Epb41l5/Sox13/Ptprc/Myoc/F11r/Kif26b/Itpkb/Dusp10/Utrn/Lama2/Lims1/Vsir/Itgb2/Icosl/Igf1/Kitl/Smarcc2/Cd63/Havcr2/Irf1/Nlrp3/Pld2/Gp1ba/Vtn/Poldip2/Lgals9/Smarce1/Stat5b/Itgb3/P4hb/Smoc1/Dicer1/Lgals8/Nid1/Gli3/Carmil1/Nedd9/Ninj1/Ecm2/Syk/Edil3/Il6st/Adk/Wnt5a/Fermt2/Pnp/Dock5/Dab2/Egflam/Il7r/Myo10/Angpt1/Has2/Lgals1/Triobp/Fbln1/Arid2/Itga5/Nckap1l/Emp2/Ppm1f/Crkl/Tfrc/Cd86/Cd80/Ccdc80/Abi3bp/Runx1/Rasal3/H2-Ob/H2-Aa/H2-Eb1/Aif1/Vegfa/Vav1/Lama1/Socs5/Rock1/Lims2/Megf10/Cd74/Efemp2/Rela/Gcnt1/Dock8/Smarca2/Jak2/Cd274/Gpam/Prkcq/Rsu1/Apbb1ip/Tsc1/Dpp4/Itga6/Itgav/Mdk/Hsd17b12/Traf6/Cd44/Cd59a/Thbs1/B2m/Rin2/Tgm2/Sdc4/Bmp7/Actl6a/Sox2/P2ry12/Cd1d1/Il6ra/Ptpn22/Csf1/Vav3/Vcam1/Lef1/Npnt/Ccn1/Bcl10/Ripk2/Epb41l4b/Tek/Plpp3/Col16a1/Ptafr/Pdpn/Tnfrsf14/Prkcz/Cdk6/Rhoh/Kdr/Dmp1/Spp1/Ptpn11/Ephb4/Hmgb1/Cav1/Podxl/Ptn/Gimap5/Npy/Ndnf/Dysf/Alox5/Cxcl12/Ret/Ptpn6/Prkd2/Plaur/Ceacam1/Tgfb1/Rras/Tjp1/Klhl25/Cib1/Iqgap1/Ilk/Il4ra/Itgal/Stx4a/Pycard/Dock1/Igf2/Cd81/Adam9/Plekha2/Nrg1/Il15/Adgrg1/Cbfb/Cdh1/Nfat5/Zfhx3/Itgb1/Icam1/Dnm2/Ets1/St3gal4/Il18/Tbx18/Hyal1/Rhoa/Lamb2/Cspg5/Tgfbr2/Ccr2/Ccr5/Sash3/Irak1/Flna/Dmd/Il2rg/Egfl6 | 198 |
| BP | GO:0031589 | cell-substrate adhesion | 154/3915 | 366/17474 | 1,7074E-17 | 2,8179E-14 | 2,0985E-14 | Map4k4/Col3a1/Fn1/Sned1/Bcl2/Epb41l5/Lamc2/Myoc/Utrn/Ccn2/Lims1/Col13a1/Itgb2/Ntn4/Cd63/Myo1g/Atp1b2/Vtn/Poldip2/Col1a1/Itga2b/Itgb3/Pecam1/P4hb/Smoc1/Dicer1/Itgb8/Nid1/Carmil1/Dusp22/Nedd9/Ninj1/Ecm2/Edil3/Pik3r1/Itga1/Fam107a/Nid2/Plau/Vcl/Fermt2/Mmp14/Dock5/Sorbs3/Dab2/Egflam/Adamts12/Angpt1/Has2/Rac2/Lgals1/Triobp/Parvg/Fbln1/Acvrl1/Itga5/Emp2/Ppm1f/Crkl/St6gal1/Apod/Itgb5/Ccdc80/Phldb2/Abi3bp/Vegfa/Ston1/Rock1/Lims2/Efemp2/Fermt3/Lpxn/Kank1/Jak2/Sorbs1/Rsu1/Notch1/Tsc1/Lamc3/Itga6/Itgav/Mdk/Hsd17b12/Thbs1/Trpm7/Bcl2l11/Mertk/Jag1/Rin2/Sdc4/Lama5/Bcan/Efna1/Csf1/Vcam1/Npnt/Ccn1/Frem1/Acer2/Tek/Tesk2/Macf1/Col16a1/Pdpn/Vamp3/Prkcz/Cdk6/Limch1/Kdr/Dmp1/Spp1/Ptpn11/Ptprz1/Ptn/Npy/Ndnf/Antxr1/Adamts9/Vwf/Ttyh1/Bcam/Axl/Rras/Cib1/Iqgap1/Fzd4/Ilk/Parva/Itgal/Itgam/Dock1/Gas6/Angpt2/Thsd1/Adam9/Plekha2/Dlc1/Zfp469/Agt/Itgb1/Dnm2/Ephb1/Rpl29/Rhoa/Lamb2/Cspg5/Clasp2/Itga9/Ctnnb1/Flna/Dmd/Gpm6b/Egfl6/Arhgap6 | 154 |
| BP | GO:0001667 | ameboidal-type cell migration | 178/3915 | 453/17474 | 1,9539E-16 | 2,6873E-13 | 2,0013E-13 | Sox17/Map4k4/Nrp2/Erbb4/Fn1/Tns1/Vil1/Irs1/Sp100/Epb41l5/Adipor1/Glul/Ddr2/Dusp10/Prox1/Akap12/Sash1/Nr2e1/Sgpl1/Col18a1/Itgb2/Syde1/Bsg/Appl2/Igf1/Kitl/Cd63/Tns3/Flt4/Sparc/Adora2b/Pmp22/Alox12/Serpinf1/Rffl/Grn/Itgb3/Pecam1/Smurf2/Gna13/Sox9/Adam17/Pik3cg/Prkd1/Arhgap5/Clec14a/Rhoj/Fut8/Lgmn/Dicer1/Lgals8/Gpld1/Edn1/Mef2c/Isl1/Wnt5a/Sema3g/Vstm4/Mmrn2/Bmp4/Dock5/Rgcc/Ednrb/Sema5a/Angpt1/Has2/Card10/Hdac7/Acvrl1/Sp1/Emp2/Ppm1f/Tbx1/Fstl1/Vegfa/Daam2/Ptprm/Cyp1b1/Jcad/Rock1/Hbegf/Fgf1/Mcc/Sema6a/Smad4/Lrp5/Coro1b/Lpxn/Kank1/Acta2/Notch1/Pkn3/Dpp4/Nfe2l2/Itgav/Pax6/Spred1/Thbs1/Sema6d/Rin2/Sdc4/Cd40/Bmp7/Edn3/Lama5/Fgf2/P2ry12/Efna1/Rab13/Pkn2/Glipr2/Tgfbr1/Klf4/Epb41l4b/Tek/Plpp3/Pik3r3/Svbp/Macf1/Phactr4/Wasf2/Sema3d/Sema3c/Nos3/Paxip1/Fgfbp1/Kdr/Anxa3/Tgfbr3/Ptpn11/P2rx4/Hspb1/Ephb4/Hmgb1/Stard13/Smo/Atoh8/Gata2/Adamts9/Vhl/Cxcl12/Ret/Adipor2/Lrp6/Arhgdib/Prkd2/Apoe/Ceacam1/Tgfb1/Rras/Cib1/Iqgap1/Prcp/Ilk/Pik3c2a/Cln3/Dock1/Igf2/Angpt2/Adam9/Vegfc/Ednra/Cdh5/Plcg2/Agt/Itgb1/Amotl1/Kank2/Bmper/Ets1/Ctsh/Hyal1/Rhoa/Clasp2/Tgfbr2/Arhgap4/Mecp2/Amot | 178 |
| BP | GO:0009611 | response to wounding | 191/3915 | 499/17474 | 3,5786E-16 | 4,2187E-13 | 3,1417E-13 | Dst/Ecrg4/Col3a1/Cflar/Fn1/Slc11a1/Vil1/Serpine2/Bcl2/Cfh/Pla2g4a/Tnr/Ddr2/Fcer1g/F11r/Vangl2/Enpp1/Gja1/Cnn2/Igf1/Elk3/Ddit3/Kremen1/Egfr/Plek/Clec10a/Alox12/Gp1ba/Serpinf2/Vtn/Col1a1/Fkbp10/Grn/Itga2b/Gfap/Itgb3/Pecam1/Gna13/Grin2c/C1qtnf1/Sdc1/Adam17/Zfp36l1/Mtr/F13a1/Dsp/Syk/Dhfr/F2r/Map3k1/Plau/Wnt5a/Prkcd/Mustn1/Fermt2/Lcp1/Slc1a3/Fzd6/Fbln1/Ppara/Ano6/Acvrl1/Abat/Serpind1/Mylk/Phldb2/Abi3bp/Pros1/Cxadr/Cdkn1a/Enpp4/Vegfa/Trem2/Zfp36l2/Nrep/Hbegf/Fgf1/Lox/Adrb2/Smad4/Coro1b/Vegfb/Fermt3/Ahnak/Kank1/Jak2/Papss2/Acta2/Myof/Entpd1/Casp7/Prkcq/Entpd2/Eng/Nfe2l2/Tfpi/Serping1/F2/Mdk/Slc1a2/Cd44/Thbs1/Mertk/Thbd/Ccm2l/Srsf6/Matn4/Sdc4/Anxa5/Fgf2/Slc7a11/P2ry12/P2ry1/Pear1/Il6ra/Cers2/Notch2/Stxbp3/F3/Ccn1/Lyn/Chmp5/Klf4/Epb41l4b/Tnc/Tlr4/Nfib/Nfia/Plpp3/Cldn19/Macf1/Arhgef19/Pdpn/Fgfr3/Tec/Pdgfra/Kdr/Arhgap24/P2rx4/Hmgb1/Cav1/Ptn/Pdia4/Ndnf/Dysf/Gp9/Gata2/Alox5/Adipor2/Ptpn6/Cd9/Vwf/Clec7a/Gpr4/Apoe/Plaur/Ceacam1/Tgfb1/Axl/Zfp36/Tyrobp/Hps5/Igf1r/Abhd2/Prcp/Xylt1/Vkorc1/Cd151/Gas6/Plat/Nrg1/Casp3/Ednra/Mmp2/Plcg2/Wfdc1/Pard3/Itgb1/Yap1/St3gal4/Ubash3b/Neo1/Anxa2/Cd109/Lamb2/Clasp2/Ccr2/Ndp/Flna/Atp7a/Sytl4 | 191 |

| | | | | | | | | | |
|---|---|---|---|---|---|---|---|---|---|
| BP | GO:0043062 | extracellular structure organization | 134/3915 | 315/17474 | 7,5213E-16 | 6,2065E-13 | 4,622E-13 | Col9a1/Col19a1/Col3a1/Cflar/Fn1/Hmcn1/Qsox1/Tnr/Ddr2/Ccn2/Lama2/Nr2e1/P4ha1/Col13a1/Col18a1/Ntn4/Lum/Aebp1/Sh3pxd2b/Col23a1/Mfap4/Pmp22/Serpinf2/Vtn/Col1a1/P3h4/Fkbp10/Ramp2/Gfap/Itgb3/Axin2/Sox9/Smoc1/Fbln5/Nid1/Foxf2/Foxc1/Ecm2/Tgfbi/Adamts16/Adamts6/Colq/Lgals3/Mmp14/Scara3/Rb1/Lcp1/Rgcc/Egflam/Adamts12/Sema5a/Col14a1/Has2/Fbln1/Adamts20/Myh11/Ccdc80/Phldb2/Abi3bp/Adamts1/Adamts10/Col11a2/Lama1/Cyp1b1/Lox/Efemp2/Kazald1/Notch1/Eng/Olfml2a/Dpp4/Hsd17b12/Cst3/Matn4/Slc2a10/Rxfp1/Pbxip1/Ctss/Npnt/Slc39a8/Ccn1/Reck/Tgfbr1/Tmem38b/Col27a1/Adamtsl1/Cyp2j6/Tie1/Col9a2/Col16a1/Hspg2/Pdpn/Tnfrsf1b/Vwa1/Pdgfra/Dmp1/Idua/Eln/Plod3/Col1a2/Cav2/Cav1/Ndnf/Prdm5/Loxl3/Antxr1/Adamts9/Vhl/Tnfrsf1a/Tgfb1/Spint2/Arhgap33os/Adamtsl3/Serpinh1/Csgalnact1/Mmp2/Atxn1l/Crispld2/Zfp469/Agt/Itgb1/Aplp2/Ets1/Anxa2/Myo1e/Nphp3/Lamb2/Clasp2/Crtap/Atp7a/Col4a6/Col4a5/Gpm6b/Egfl6 | 134 |
| BP | GO:0090132 | epithelium migration | 135/3915 | 319/17474 | 9,701E-16 | 7,2775E-13 | 5,4196E-13 | Map4k4/Vil1/Irs1/Sp100/Epb41l5/Adipor1/Glul/Dusp10/Prox1/Sash1/Nr2e1/Col18a1/Itgb2/Bsg/Igf1/Kitl/Cd63/Tns3/Flt4/Sparc/Adora2b/Alox12/Serpinf1/Grn/Itgb3/Pecam1/Sox9/Adam17/Pik3cg/Prkd1/Arhgap5/Clec14a/Rhoj/Lgmn/Dicer1/Lgals8/Gpld1/Mef2c/Wnt5a/Vstm4/Mmrn2/Bmp4/Dock5/Rgcc/Sema5a/Angpt1/Has2/Card10/Hdac7/Acvrl1/Sp1/Emp2/Ppm1f/Fstl1/Vegfa/Daam2/Ptprm/Cyp1b1/Jcad/Rock1/Hbegf/Fgf1/Mcc/Smad4/Coro1b/Lpxn/Kank1/Notch1/Pkn3/Dpp4/Nfe2l2/Itgav/Spred1/Thbs1/Rin2/Cd40/Fgf2/Efna1/Rab13/Pkn2/Glipr2/Tgfbr1/Klf4/Epb41l4b/Tek/Plpp3/Pik3r3/Svbp/Macf1/Nos3/Paxip1/Fgfbp1/Kdr/Anxa3/Tgfbr3/Ptpn11/P2rx4/Hspb1/Ephb4/Hmgb1/Stard13/Atoh8/Gata2/Adamts9/Vhl/Cxcl12/Prkd2/Apoe/Ceacam1/Tgfb1/Rras/Cib1/Prcp/Pik3c2a/Cln3/Dock1/Igf2/Angpt2/Adam9/Vegfc/Cdh5/Plcg2/Agt/Itgb1/Amotl1/Kank2/Bmper/Ets1/Ctsh/Hyal1/Rhoa/Clasp2/Tgfbr2/Mecp2/Amot | 135 |
| BP | GO:0002237 | response to molecule of bacterial origin | 121/3915 | 278/17474 | 2,5051E-15 | 1,3782E-12 | 1,0263E-12 | Ly96/Arid5a/Stat1/Slc11a1/Tnfrsf11a/Cfh/Ncf2/Fcgr2b/Cd84/Dusp10/Tab2/Sash1/Irak3/Havcr2/Irgm1/Igtp/Irgm2/Nlrp3/Cd68/Cxcl16/Nos2/Lgals9/Stat5b/Ace/Adam17/Nfkbia/Ly86/Bmp6/Tifab/Mef2c/F2r/Cd180/Erbin/Wnt5a/Ednrb/Mtdh/Nlrc3/Litaf/Cd86/Cd80/Ifnar1/Akap8/Tap2/Trem2/Ticam1/Eif2ak2/Cd14/Ticam2/Rela/Jak2/Cd274/Chuk/Vim/Mrc1/Card9/Itgav/Spi1/Nr1h3/Traf6/Fbxo3/B2m/Lbp/Tlr2/Ptpn22/Nfkb1/Adh5/Gbp2/Bcl10/Ptgfr/Lyn/Ripk2/Chmp5/Abca1/Tlr4/Ptafr/Tnfrsf1b/Nos3/Hadhb/Tnip2/Tlr1/Tlr6/Gbp6/P2rx7/Hmgb1/Irf5/Nr2c2/Irak2/Mgst1/C5ar1/Tgfb1/Axl/Zfp36/Sirt2/Irf3/Rpl13a/Trim5/Trim12a/Trim12c/Trim30a/Mapk3/Pycard/Adam9/Ednra/Nod2/Nqo1/Plcg2/Irf8/Tirap/Il18/Plscr1/Plscr2/Plscr4/Mapkapk3/Rhoa/Myd88/Cx3cr1/Ldoc1/Irak1/Ogt/Btk/Rps6ka3 | 121 |
| BP | GO:0045446 | endothelial cell differentiation | 64/3915 | 113/17474 | 3,0455E-15 | 1,5707E-12 | 1,1697E-12 | Sox17/F11r/Prox1/Hey2/Col18a1/Btg1/Vezf1/Tmem100/Fzd2/Pecam1/Dicer1/Bmp6/Marveld2/Vcl/Bmp4/Ednrb/Rapgef3/Acvrl1/Cldn5/Heg1/Fstl1/Ezr/Notch4/Vegfa/Xdh/Zeb1/Rock1/Smad4/Tjp2/Notch1/Jag1/Id1/Ppp1r16b/Arhgef26/Rap1a/Tgfbr1/Tie1/Clic4/Pdpn/Abcb1b/Rbpj/Kdr/Plod3/Atoh8/Vhl/Tnfrsf1a/Ceacam1/Tjp1/Fzd4/Ikbkb/Nrg1/Ednra/Cdh5/Agt/S1pr2/Icam1/Rdx/Myd88/Acvr2b/Ctnnb1/Ndp/Rap2c/Dmd/Msn | 64 |
| BP | GO:0003158 | endothelium development | 71/3915 | 132/17474 | 3,6765E-15 | 1,6855E-12 | 1,2552E-12 | Sox17/Slc40a1/F11r/Prox1/Hey2/Col18a1/Bsg/Btg1/Alox12/Vezf1/Tmem100/Fzd2/Pecam1/Dicer1/Bmp6/Marveld2/Vcl/Bmp4/Ednrb/Adamts12/Rapgef3/Acvrl1/Cldn5/Heg1/Fstl1/Ezr/Notch4/Vegfa/Xdh/Zeb1/Rock1/Fgf1/Smad4/Tjp2/Notch1/Jag1/Id1/Ppp1r16b/Arhgef26/Rap1a/Tgfbr1/Tie1/Clic4/Pdpn/Abcb1b/Rbpj/Kdr/Plod3/Stard13/Atoh8/Vhl/Tnfrsf1a/Ceacam1/Tjp1/Fzd4/Ikbkb/Nrg1/Ednra/Cdh5/Agt/S1pr2/Icam1/Rdx/Rhoa/Myd88/Acvr2b/Ctnnb1/Ndp/Rap2c/Dmd/Msn | 71 |
| BP | GO:0071216 | cellular response to biotic stimulus | 100/3915 | 217/17474 | 7,8991E-15 | 3,4307E-12 | 2,5548E-12 | Ly96/Arid5a/Stat1/Cfh/Tmco1/Fcgr2b/Cd84/Sash1/Ddit3/Havcr2/Irgm1/Igtp/Irgm2/Nlrp3/Cd68/Cxcl16/Nos2/Ccdc47/Nfkbia/Ly86/Bmp6/Syk/Tifab/Mef2c/Cd180/Wnt5a/Mtdh/Litaf/Cd86/Cd80/Akap8/Trem2/Ticam1/Cd14/Ticam2/Rela/Jak2/Cd274/Fbh1/Vim/Mrc1/Notch1/Spi1/Nr1h3/Traf6/B2m/Bcl2l11/Lbp/Tlr2/Txnip/Notch2/Ptpn22/Nfkb1/Gbp2/Bcl10/Lyn/Ripk2/Chmp5/Abca1/Tlr4/Zmpste24/Ptafr/Tnfrsf1b/Nos3/Hadhb/Tnip2/Tlr1/Tlr6/Gbp6/Hmgb1/Nr2c2/Irak2/Clec7a/Tgfb1/Axl/Zfp36/Sirt2/Irf3/Trim5/Trim12a/Trim12c/Trim30a/Mapk3/Pycard/Adam9/Nod2/Plcg2/Irf8/Tirap/Il18/Plscr1/Plscr2/Plscr4/Rhoa/Myd88/Cx3cr1/Ldoc1/Irak1/Ogt/Btk | 100 |
| BP | GO:0045765 | regulation of angiogenesis | 124/3915 | 293/17474 | 1,4371E-14 | 5,9296E-12 | 4,4157E-12 | Stat1/Sp100/Chil1/Glul/Sash1/Ccn6/Nr2e1/Itgb2/Btg1/Sparc/Serpinf1/Stat3/Ramp2/Brca1/Grn/Itgb3/Prkd1/Rhoj/Fbln5/Itgb8/Foxc1/Ninj1/Erap1/Isl1/Wnt5a/Stab1/Lgals3/Rgcc/Sema5a/Mtdh/Rapgef3/Acvrl1/Sp1/Itga5/Emp2/Cldn5/Adamts1/Runx1/Thbs2/Notch4/Vegfa/Ptprm/Cyp1b1/Jcad/Rock1/Ecscr/Fgf1/Sema6a/Adrb2/Tcf4/Vegfb/Slc39a12/Eng/Nfe2l2/Mdk/Cd59a/Spred1/Thbs1/Id1/Ppp1r16b/Cd40/Ptgis/Fgf2/Efna1/Shc1/S100a1/Nras/F3/Ddah1/Reck/Klf4/Tek/Tie1/Ago1/Hgf/Nos3/Kdr/Anxa3/Hspb1/Flt1/Hmgb1/Stard13/Tspan12/Ptn/Hipk2/Hk2/Gata2/Adamts9/Alox5/C3ar1/Tnfrsf1a/C5ar1/Prkd2/Gpr4/Ceacam1/Pak4/Rras/Tjp1/Pde3b/Rnh1/Igf2/Angpt2/Tlr3/Vegfc/Klf2/Smad1/Hhip/Gab1/Cdh5/Agt/Itgb1/Bmper/Ets1/Ctsh/Hyal1/Tgfbr2/Cx3cr1/Ctnnb1/Ccr2/Cybb/Mecp2/Foxo4/Cysltr1/Amot | 124 |
| BP | GO:0033002 | muscle cell proliferation | 113/3915 | 260/17474 | 2,3666E-14 | 8,137E-12 | 6,0596E-12 | Cflar/Erbb4/Igfbp5/Rgs5/Ddr2/Kcnk2/Enpp1/Hey2/Gja1/Timp3/Igf1/Frs2/Ddit3/Egfr/Alox12/Serpinf2/Poldip2/Stat5b/Stat3/Itgb3/Gna13/Id2/Sav1/Foxc1/Nqo2/Edn1/Ogn/Mef2c/Pik3r1/Bmpr1a/Bmp4/Ang/Ndrg2/Abcc4/Angpt1/Myc/Irak4/Arid2/Prkdc/Apod/Cxadr/Adamts1/Paxbp1/Cdkn1a/Aif1/Vegfa/Hbegf/Fgf1/Megf10/Pdgfrb/Efemp2/Jak2/Tcf7l2/Notch1/Calcrl/Traf6/Thbs1/Tgm2/Pik3ca/Fgf2/Shc1/Il6ra/Gnai3/S1pr1/Camk2d/Tgfbr1/Klf4/Tlr4/Ptafr/Ldlrap1/Ctnnbip1/Trp73/Hes5/Hgf/Nos3/Ppargc1a/Rbpj/Tgfbr3/Gper1/Flt1/Cav2/Cav1/Vgll4/C3ar1/Ccnd2/Cdkn1b/Apoe/Tgfb1/Igf1r/P2ry6/Ilk/Fgfr2/Nrg1/Hpgd/Ednra/Smad1/Il15/Mmp2/Cyba/Agt/Pdgfd/Yap1/S1pr2/Cnn1/Il18/Ephb1/Gnai2/Rhoa/Tgfbr2/Myd88/Ctnnb1/Apln/Irak1 | 113 |
| BP | GO:0016054 | organic acid catabolic process | 105/3915 | 236/17474 | 3,0515E-14 | 9,685E-12 | 7,2124E-12 | Adhfe1/Hibch/Acadl/Irs1/Dbi/Npl/Echdc1/Ddo/Ilvbl/Aldh1l2/Shmt2/Pex13/Shmt1/Acadvl/Blmh/Nos2/Acox1/Dcxr/Lpin1/Aldh6a1/Abcd4/Gstz1/Aldh5a1/Eci2/Hexb/Mccc2/Hacl1/Glud1/Abcd2/Csad/Abat/Prodh/Ehhadh/Acat3/Eci1/Cbs/Cyp4f15/Cyp4f14/Cyp4f13/Ddah2/Abhd3/Gnpda1/Cdo1/Hsd17b4/Acaa2/Cpt1a/Asrgl1/Gldc/Phyh/Gad2/Sardh/Gad1/Ivd/Lpin3/Pex2/Etfdh/Slc16a1/Dbt/Abcd3/Hadh/Bdh2/Ddah1/Acadm/Decr1/Cpt2/Scp2/Echdc2/Mfsd2a/Hmgcl/Aldh4a1/Pgd/Crot/Nos3/Hadha/Hadhb/Acox3/Idua/Acacb/Acads/Sdsl/Pon3/Aass/Hibadh/Thnsl2/Aldh1l1/Lipe/Bckdha/Akt2/Etfb/Bcat2/Abhd2/Acadsb/Oat/Echs1/Gcdh/Nudt7/Gcsh/Acad8/Etfa/Hexa/Acad11/Amt/Ids/Renbp/Hsd17b10 | 105 |
| BP | GO:0046395 | carboxylic acid catabolic process | 105/3915 | 236/17474 | 3,0515E-14 | 9,685E-12 | 7,2124E-12 | Adhfe1/Hibch/Acadl/Irs1/Dbi/Npl/Echdc1/Ddo/Ilvbl/Aldh1l2/Shmt2/Pex13/Shmt1/Acadvl/Blmh/Nos2/Acox1/Dcxr/Lpin1/Aldh6a1/Abcd4/Gstz1/Aldh5a1/Eci2/Hexb/Mccc2/Hacl1/Glud1/Abcd2/Csad/Abat/Prodh/Ehhadh/Acat3/Eci1/Cbs/Cyp4f15/Cyp4f14/Cyp4f13/Ddah2/Abhd3/Gnpda1/Cdo1/Hsd17b4/Acaa2/Cpt1a/Asrgl1/Gldc/Phyh/Gad2/Sardh/Gad1/Ivd/Lpin3/Pex2/Etfdh/Slc16a1/Dbt/Abcd3/Hadh/Bdh2/Ddah1/Acadm/Decr1/Cpt2/Scp2/Echdc2/Mfsd2a/Hmgcl/Aldh4a1/Pgd/Crot/Nos3/Hadha/Hadhb/Acox3/Idua/Acacb/Acads/Sdsl/Pon3/Aass/Hibadh/Thnsl2/Aldh1l1/Lipe/Bckdha/Akt2/Etfb/Bcat2/Abhd2/Acadsb/Oat/Echs1/Gcdh/Nudt7/Gcsh/Acad8/Etfa/Hexa/Acad11/Amt/Ids/Renbp/Hsd17b10 | 105 |
| BP | GO:0048771 | tissue remodeling | 93/3915 | 200/17474 | 3,4888E-14 | 1,0643E-11 | 7,9258E-12 | Igfbp5/Tmbim1/Inpp5d/Tnfrsf11a/Suco/Ddr2/Enpp1/Gja1/Igf1/Tns3/Egfr/Flt4/Clec10a/Nos2/P3h4/Itgb3/Klf6/Gpr137b/Foxc1/Syk/Mef2c/Thbs4/F2r/Mmp14/Dock5/Cthrc1/Rac2/Acvrl1/Tbx1/Tfrc/Abi3bp/Cbs/Vegfa/Epas1/Rock1/Csf1r/Adrb2/Lrp5/Tcirg1/Itgav/Mdk/Traf6/Lgr4/Rassf2/Jag1/Cst3/Tgm2/Car2/Ctsk/Ctss/Gja5/Notch2/S1pr1/Nfkb1/Chd7/Lepr/Tie1/Abcb1b/Sema3c/Nos3/Fgfr3/Cd38/Rbpj/Spp1/Tgfbr3/Idua/Tmem119/P2rx7/Eln/Pdk4/Cav1/Ptn/Gpnmb/Lrp6/Ceacam1/Axl/Lrrk1/Tpp1/Angpt2/Ednra/Inpp4b/Mmp2/Nfatc3/Agt/Ubash3b/Il18/Pth1r/Acvr2b/Ctnnb1/Ccr2/Ndp/Flna/Atp7a | 93 |

| | | | | | | | | | |
|---|---|---|---|---|---|---|---|---|---|
| BP | GO:0001503 | ossification | 155/3915 | 398/17474 | 4,2975E-14 | 1,2229E-11 | 9,1066E-12 | Nab1/Igfbp5/Tnfrsf11a/Bcl2/Gli2/Suco/Myoc/Pbx1/Ddr2/H3f3a/Ccn2/Enpp1/Asf1a/Gja1/Col13a1/Srgn/Igf1/Kremen1/Fignl1/Sh3pxd2b/Rflnb/Col1a1/Nbr1/Wnt3/Axin2/Sox9/H3f3b/Notum/Asxl2/Id2/Prkd1/Smoc1/Gli3/Gpld1/Prl/Foxc1/Bmp6/Id4/Aspn/Omd/Smad5/Ptch1/Mef2c/Il6st/Wnt5a/Gdf10/Bmpr1a/Fermt2/Bmp4/Mmp14/Nipbl/Ranbp3l/Adamts12/Ank/Cthrc1/Mtss1/Ano6/Hdac7/Sp1/Casr/Cbs/Clic1/Vegfa/Twsg1/Lox/Csf1r/Adrb2/Smad4/Nfatc1/Lrp5/Tcirg1/Rorb/Kazald1/Tcf7l2/Notch1/Lrp4/Mdk/Traf6/Lgr4/Rassf2/Jag1/Id1/Trp53inp2/Zhx3/Bmp7/Sox2/Fgf2/Wwtr1/Igsf10/Csf1/S1pr1/Sgms2/Npnt/Bmpr1b/Ccn1/Map3k7/Tmem38b/Zmpste24/Id3/Hspg2/Alpl/Dhrs3/Ctnnbip1/Isg15/Cdk6/Atraid/Fgfr3/Rbpj/Rest/Bmp2k/Dmp1/Spp1/Tgfbr3/Tmem119/Ptpn11/P2rx7/Sbds/Col1a2/Smo/Ptn/Clec5a/Alox5/Lrp6/Mgp/Tgfb1/Chsy1/P2ry2/Inppl1/Ilk/Xylt1/Mapk3/Fgfr2/Ifitm1/Igf2/Vegfc/Csgalnact1/Smad1/Mmp2/Cbfb/Yap1/Bmp5/Rhoa/Plxnb1/Setd2/Pth1r/Acvr2b/Ctnnb1/Clec3b/Limd1/Ebp/Gpc3/Gdpd2/Hdac8/Chrdl1/Gpm6b | 155 |
| BP | GO:0031349 | positive regulation of defense response | 163/3915 | 425/17474 | 4,4981E-14 | 1,2373E-11 | 9,2141E-12 | Ly96/Wdfy1/Tnfrsf11a/Ikbke/Rab7b/Pla2g4a/Fcgr3/Fcer1g/Aim2/Mndal/Ifi203/Tlr5/Cdk19/Gja1/Appl2/Polr3b/Irak3/Egfr/Havcr2/Irgm1/Irf1/Igtp/Irgm2/Nlrp3/Adora2b/Nlrp1b/Lgals9/Rnf135/Stat5b/Aoc3/Ifi35/Grn/Ace/Cd300a/Pum2/Pik3cg/Nfkbia/Traf3/Ripk1/Ninj1/Syk/Mef2c/Cd180/Erbin/Wnt5a/Il17rb/Mmrn2/Lacc1/Lgals1/Gramd4/Prkdc/Snx4/Parp9/Cd86/Slc15a2/Trem2/Ticam1/Gpr108/Vav1/Colec12/Riok3/Rnf125/Sting1/Cd14/Ticam2/Cd74/Csf1r/Unc93b1/Rela/Slc15a3/Jak2/Pik3ap1/Card9/Nmi/Dpp4/Ifih1/Spi1/Nr1h3/Mdk/Lgr4/Mavs/Tgm2/Lbp/Cd40/Znfx1/Tlr2/Ctss/Fcgr1/Ptpn22/Pde5a/Alpk1/Tifa/Casp6/Gbp5/Bcl10/Lyn/Ripk2/Tlr4/Tlr12/Camk2n1/Pla2g5/Tnip2/Tlr1/Tlr6/Rbm47/Ankrd17/Oasl1/Oas1a/Trim56/Hmgb1/Alox5ap/Cav1/Zc3hav1/Gimap5/Tril/Nod1/Vamp8/Irak2/Il17ra/Lag3/Tnfrsf1a/Clec7a/Gpr4/Nectin2/Rps19/Tyrobp/Lsm14a/Irf3/Ctsc/Trim5/Trim12a/Trim12c/Trim30a/Gprc5b/Nupr1/Mapk3/Pycard/Irf7/Cd81/Lamp1/Tlr3/Ddx60/Ednra/Nod2/Nlrc5/Plcg2/Cyba/Casp4/Ldlr/Ets1/Tirap/Il18/Plscr1/Plscr2/Mapkapk3/Myd88/Ccr2/Ccr5/Irak1/Tlr13/Btk/Rps6ka3/Tlr7 | 163 |
| BP | GO:0007249 | I-kappaB kinase/NF-kappaB signaling | 102/3915 | 231/17474 | 1,2798E-13 | 3,3004E-11 | 2,4578E-11 | Stat1/Casp8/Ikbke/Tab2/Fyn/Ddx21/S100b/Rel/Irf1/Traf4/Trim25/Tmem106a/Prkd1/Nfkbia/Traf3/Ripk1/Edn1/Tifab/Wnt5a/Ripk3/Tnfrsf19/Ednrb/Mtdh/Angpt1/Irak4/Nlrc3/Tfrc/Ube2i/Trem2/Ticam1/Ppm1b/Rock1/Riok3/Ticam2/Cd74/Rela/Chuk/Card9/Traf1/Tank/Traf6/Capn3/Ddrgk1/Cpne1/Tgm2/Cd40/Tnfsf10/Tlr2/S100a13/S100a4/Otud7b/Alpk1/Tifa/Nfkb1/Slc39a8/Gbp7/Gbp3/Bcl10/Cth/Ripk2/Map3k7/Tlr4/Lurap1/Snip1/Hdac1/Tnip2/Tlr6/Rhoh/Ankrd17/Hspb1/Zc3hav1/Nod1/Irak2/Erc1/Ltbr/Irf3/Zfand6/Trim21/Trim68/Trim34a/Trim5/Trim12a/Trim12c/Trim30a/Gprc5b/Pycard/Ikbkb/Tlr3/Ednra/Nod2/Plcg2/Tirap/Hacd3/Rora/Rhoa/Myd88/Cx3cr1/Ctnnb1/Irak1/Btk/Mid2/Tlr7 | 102 |
| BP | GO:0045088 | regulation of innate immune response | 143/3915 | 363/17474 | 1,463E-13 | 3,6203E-11 | 2,696E-11 | Ly96/Casp8/Wdfy1/Ikbke/Rab7b/Cfh/Aim2/Mndal/Ifi203/Tlr5/Dusp10/Appl2/Polr3b/Irak3/Havcr2/Irgm1/Irf1/Igtp/Irgm2/Nlrp3/Nlrp1b/Lgals9/Rnf135/Stat5b/Ifi35/Grn/Cd300a/Pum2/Nfkbia/Traf3/Serpinb9/Ninj1/Syk/Cd180/Erbin/Wnt5a/Lacc1/Gramd4/Nlrc3/Prkdc/Parp14/Parp9/Cd86/Slc15a2/Myo1f/Tap1/Tap2/Trem2/Ticam1/Gpr108/Vav1/Colec12/Riok3/Rnf125/Sting1/Cd14/Ticam2/Atg12/Cd74/Unc93b1/Rela/Slc15a3/Pik3ap1/Card9/Nmi/Dpp4/Ifih1/Nfe2l2/Serping1/Spi1/Nr1h3/Lgr4/Mavs/Samhd1/Lbp/Cd40/Znfx1/Tlr2/Ptpn22/Alpk1/Tifa/Casp6/Gbp5/Bcl10/Lyn/Ripk2/Tlr4/Lrp8/Tlr12/Pla2g5/Isg15/Tnip2/Tlr1/Tlr6/Rbm47/Ankrd17/Oasl1/Oas1a/Ncf1/Trim56/Hmgb1/Cav1/Zc3hav1/Gimap5/Tril/Nod1/Irak2/A2m/Lag3/Clec2d/Clec7a/Apoe/Nectin2/Rps19/Ceacam1/Tgfb1/Tyrobp/Lsm14a/Irf3/Trim21/Trim5/Trim12a/Trim12c/Trim30a/Pycard/Irf7/Igf2/Lamp1/Tlr3/Ddx60/Nod2/Nlrc5/Plcg2/Cyba/Tirap/Plscr1/Plscr2/Mapkapk3/Myd88/Irak1/Tlr13/Rps6ka3/Tlr7 | 143 |
| BP | GO:0042063 | gliogenesis | 143/3915 | 366/17474 | 3,1122E-13 | 7,27E-11 | 5,4139E-11 | Col3a1/Nab1/Fn1/Serpine2/Dbi/Srgap2/Sox13/Gpr37l1/Myoc/Kcnj10/Dusp10/Adgrg6/Ifngr1/Enpp1/Nr2e1/Zfp365/Adora2a/Igf1/Ptprb/Erbb3/Egfr/Efemp1/Pmp22/Atp1b2/Vtn/Tmem98/Stat3/Grn/Gfap/Sox9/Id2/Prkch/Syne2/Dicer1/Gli3/Id4/Ntrk2/Vcan/Hexb/Il6st/Zcchc24/Zfp488/Bmp4/Clu/Rb1/Gpr183/Myc/Sun2/Tspo/Crkl/Nrros/Metrn/Trem2/Daam2/Nr3c1/Csf1r/Apcdd1/Mbd1/Rela/Fas/Tcf7l2/Vim/Notch1/Lamc3/Dlx1/Ckap5/F2/Mdk/Pax6/Fgf2/P2ry12/P2ry1/Tlr2/Cers2/Phgdh/Csf1/Lef1/Lyn/Pou3f2/Tlr4/Nfib/Nfia/Lepr/Dab1/Plpp3/Lrp8/Tal1/Hdac1/C1qa/Tnfrsf1b/Gpr157/Trp73/Hes5/Mxra8/Cdk6/Reln/Fgfr3/Rnf10/Ptpn11/P2rx4/Wasf3/Flt1/Ptprz1/Smo/Ptn/Ezh2/Klf15/Cd9/Lrp6/Sox5/C5ar1/Tgfb1/Akt2/Sirt2/Lgi4/Atf5/Rras/Idh2/Ilk/Sox6/Mapk3/Itgam/Ano1/Arhgef10/Nrg1/Vegfc/Mt3/Adgrg1/Agt/Pard3/Ldlr/Rhoa/Lamb2/Cspg5/Myd88/Cx3cr1/Ctnnb1/Ccr2/Ndp/Mecp2/Dmd/Med12/Gpm6b | 143 |
| BP | GO:0050678 | regulation of epithelial cell proliferation | 156/3915 | 411/17474 | 4,3009E-13 | 9,3398E-11 | 6,9553E-11 | Eya1/Stat1/Cflar/Erbb4/Sgpp2/Gli2/Glul/Dusp10/Prox1/Irf6/Gja1/Lims1/Col18a1/Igf1/Frs2/Egfr/Flt4/Sparc/Adora2b/Serpinf1/Stxbp4/Stat3/Etv4/Grn/Itgb3/Sox9/Adam17/Prkd1/Sav1/Zfp36l1/Dicer1/Prl/Bmp6/Ptch1/Ctsl/Mef2c/Thbs4/Isl1/Plau/Wnt5a/Mmrn2/Bmpr1a/Bmp4/Ang/Rb1/Rgcc/Ednrb/Dab2/Rictor/Sema5a/Rida/Has2/Mtss1/Myc/Acvrl1/Sp1/Nlrc3/Prkdc/Tbx1/Dlg1/Cxadr/Hmgn1/Vegfa/Ptprm/Xdh/Mta3/Jcad/Zeb1/Lims2/Fgf1/Mcc/Nfatc1/Vegfb/Tcf7l2/Nrarp/Notch1/Egfl7/Eng/Mdk/Pax6/Thbs1/B2m/Jag1/Id1/Ppp1r16b/Srsf6/Lama5/Pex2/Cyp7b1/Sox2/Fgf2/Notch2/Nras/F3/Tgfbr1/Nfib/Tek/Tie1/Hspg2/Hes5/Cdk6/Fgfr3/Fgfbp1/Rbpj/Kdr/Tgfbr3/Flt1/Hmgb1/Foxp2/Cav2/Cav1/Smo/Ptn/Fmc1/Atoh8/Dysf/Tgfa/Gkn3/Gata2/Vhl/Alox5/Cxcl12/Ccnd2/Lrp6/Cdkn1b/C5ar2/C5ar1/Prkd2/Apoe/Ceacam1/Tgfb1/Zfp36/Saal1/Nupr1/Fgfr2/Htra1/Igf2/Vegfc/Ednra/Nod2/Cdh1/Wfdc1/Cyba/Yap1/Il18/Aldh1a2/Bmp5/Cd109/Tbx18/Hyal1/Ctnnb1/Wdr13/Apln/Gpc3/Atp7a/Btk | 156 |
| BP | GO:0010632 | regulation of epithelial cell migration | 104/3915 | 242/17474 | 6,0288E-13 | 1,2134E-10 | 9,0363E-11 | Map4k4/Vil1/Irs1/Sp100/Epb41l5/Adipor1/Glul/Dusp10/Prox1/Sash1/Nr2e1/Col18a1/Bsg/Igf1/Cd63/Flt4/Sparc/Adora2b/Alox12/Serpinf1/Grn/Itgb3/Sox9/Adam17/Pik3cg/Prkd1/Arhgap5/Rhoj/Lgmn/Dicer1/Gpld1/Mef2c/Wnt5a/Mmrn2/Bmp4/Dock5/Rgcc/Sema5a/Angpt1/Has2/Card10/Hdac7/Acvrl1/Sp1/Emp2/Ppm1f/Vegfa/Ptprm/Jcad/Hbegf/Fgf1/Mcc/Notch1/Nfe2l2/Spred1/Thbs1/Rin2/Cd40/Fgf2/Efna1/Glipr2/Klf4/Epb41l4b/Tek/Plpp3/Svbp/Macf1/Nos3/Fgfbp1/Kdr/Anxa3/Tgfbr3/P2rx4/Hspb1/Hmgb1/Stard13/Atoh8/Gata2/Adamts9/Prkd2/Apoe/Ceacam1/Tgfb1/Rras/Cib1/Prcp/Pik3c2a/Dock1/Igf2/Angpt2/Adam9/Vegfc/Plcg2/Agt/Amotl1/Bmper/Ets1/Ctsh/Hyal1/Rhoa/Clasp2/Tgfbr2/Mecp2/Amot | 104 |
| BP | GO:0034440 | lipid oxidation | 63/3915 | 121/17474 | 9,1177E-13 | 1,7914E-10 | 1,3341E-10 | Acadl/Irs1/Dbi/Adipor1/Fmo1/Fmo2/Echdc1/Ilvbl/Aldh1l2/Appl2/Pex13/Alox8/Acadvl/Alox12/Sox9/Acox1/Abcd4/Eci2/Hacl1/Ppara/Abcd2/Ehhadh/Apod/Acat3/Eci1/Cyp4f13/Pla2g7/Hsd17b4/Acaa2/Cpt1a/Phyh/Ivd/Pex2/Etfdh/Abcd3/Hadh/Bdh2/Adh5/Acadm/Decr1/Cpt2/Scp2/Echdc2/Mfsd2a/Crot/Hadha/Hadhb/Acox3/Ppargc1a/Acacb/Acads/Pdk4/Alox5/Adipor2/Akt2/Etfb/Klhl25/Echs1/Cyp4v3/Gcdh/Etfa/Acad11/Hsd17b10 | 63 |
| BP | GO:0002833 | positive regulation of response to biotic stimulus | 126/3915 | 316/17474 | 1,5266E-12 | 2,7995E-10 | 2,0848E-10 | Ly96/Wdfy1/Ikbke/Rab7b/Aim2/Mndal/Ifi203/Tlr5/Sash1/Appl2/Polr3b/Irak3/Havcr2/Irgm1/Irf1/Igtp/Irgm2/Nlrp3/Nlrp1b/Lgals9/Rnf135/Stat5b/Ifi35/Grn/Cd300a/Pum2/Nfkbia/Traf3/Ly86/Bmp6/Ninj1/Syk/Cd180/Erbin/Wnt5a/Mmrn2/Lacc1/Gramd4/Prkdc/Parp9/Cd86/Slc15a2/Trem2/Ticam1/Gpr108/Vav1/Colec12/Riok3/Rnf125/Sting1/Cd14/Ticam2/Cd74/Unc93b1/Rela/Slc15a3/Cd274/Pik3ap1/Card9/Nmi/Dpp4/Ifih1/Spi1/Nr1h3/Traf6/Lgr4/Mavs/Lbp/Cd40/Znfx1/Tlr2/Ptpn22/Alpk1/Tifa/Casp6/Gbp5/Bcl10/Lyn/Ripk2/Tlr4/Tlr12/Pla2g5/Tnip2/Tlr1/Tlr6/Rbm47/Ankrd17/Oasl1/Oas1a/Trim56/Hmgb1/Cav1/Zc3hav1/Gimap5/Tril/Nod1/Irak2/Lag3/Clec7a/Nectin2/Rps19/Tyrobp/Lsm14a/Irf3/Trim5/Trim12a/Trim12c/Trim30a/Pycard/Irf7/Lamp1/Tlr3/Ddx60/Nod2/Nlrc5/Plcg2/Cyba/Tirap/Plscr1/Plscr2/Mapkapk3/Myd88/Irak1/Tlr13/Rps6ka3/Tlr7 | 126 |
| BP | GO:0032675 | regulation of interleukin-6 production | 76/3915 | 160/17474 | 1,9777E-12 | 3,4723E-10 | 2,5858E-10 | Arid5a/Il1rl2/Inpp5d/Rab7b/Prg4/Fcer1g/Cd84/Capn2/Bsg/Irak3/Havcr2/Adora2b/Nos2/Lgals9/Stat3/Tmem106a/Syk/F2r/Isl1/Wnt5a/Nckap1l/Nlrc3/Il1rap/Zbtb20/Cd200r1/Aif1/Trem2/Ticam1/Socs5/Aqp4/Ticam2/Cd74/Unc93b1/Afap1l2/Card9/Ifih1/Traf6/Mavs/Lbp/Tlr2/Il6ra/Ptpn22/Bank1/Ccn1/Ripk2/Tlr4/Laptm5/Ptafr/Hgf/Tlr1/Tlr6/Ptpn11/P2rx7/Hmgb1/Nod1/Il17rc/Il17ra/Ptpn6/Tnfrsf1a/C5ar2/Tyrobp/Arrb1/Trim30a/Pycard/Gas6/Tlr3/Klf2/Nod2/Plcg2/Cyba/Tirap/Myd88/Ccr5/Elf4/Btk/Tlr7 | 76 |

| | | | | | | | | | |
|---|---|---|---|---|---|---|---|---|---|
| BP | GO:0007162 | negative regulation of cell adhesion | 123/3915 | 308/17474 | 2,4835E-12 | 4,1824E-10 | 3,1146E-10 | Serpine2/Epb41l5/Ptprc/Tnr/Rc3h1/Myoc/Vsir/Specc1l/Adora2a/Erbb3/Havcr2/Irf1/Alox12/Lgals9/Col1a1/Cd300a/C1qtnf1/Gli3/Hfe/Dusp22/Bmp6/Pik3r1/Fam107a/Prkcd/Bmp4/Lgals3/Mmp14/Rgcc/Sema5a/Angpt1/Lgals1/Fbln1/Ppara/Plxnb2/Acvrl1/Nckap1l/Abat/Ppm1f/Apod/Dlg1/Cd86/Cd80/Phldb2/Jam2/Runx1/Myo1f/H2-Aa/Notch4/Vegfa/Twsg1/Cyp1b1/Socs5/Epcam/Sema6a/Cd74/Adrb2/Lpxn/Kank1/Jak2/Cd274/Nrarp/Notch1/Ass1/Spi1/Mdk/Cd44/Thbs1/Jag1/Ccm2l/Sdc4/Pag1/Tnfaip8l2/Ptpn22/Pde5a/Klf4/Akna/Tnc/Acer2/Dab1/Laptm5/Hspg2/Pla2g5/Tnfrsf14/Fgl2/Ptpn11/Hspb1/Hmgb1/Ptprz1/Podxl/Gimap5/Gpnmb/Loxl3/Nat8f5/Nat8/Nat8f2/Nat8f1/Cxcl12/Wnk1/Ptpn6/Lag3/Cd9/Ceacam1/Tgfb1/Spint2/Cd37/Fzd4/Lrrc32/Swap70/Pde3b/Il4ra/Angpt2/Dlc1/Casp3/Mmp2/Cbfb/Cdh1/Ubash3b/Rdx/Adam10/Rhoa/Plxnb1/Clasp2/Arhgap6 | 123 |
| BP | GO:0050900 | leukocyte migration | 140/3915 | 367/17474 | 4,486E-12 | 6,9846E-10 | 5,2014E-10 | Il1r1/Fcgr3/Fcer1g/F11r/Enpp1/Itgb2/Bsg/Cnn2/Kitl/Myo1g/Stk10/Cxcl16/Gp1ba/Lgals9/Ccl9/Ccl6/Stat5b/Aoc3/Itga2b/Itgb3/Pecam1/Cd300a/Adam17/Rin3/Lgmn/Nedd9/Edn1/Ninj1/Syk/Cxcl14/Thbs4/Pik3r1/Itga1/Wnt5a/Lgals3/Mmp14/Ripk3/Lrch1/Ednrb/Gpr183/Rac2/Irak4/Ano6/Nckap1l/Emp2/Crkl/Apod/Cd200r1/Cxadr/Jam2/Myo1f/Aif1/Pla2g7/Vegfa/Trem2/Vav1/Rock1/Cd74/Csf1r/Vegfb/Gcnt1/Dock8/Dpp4/Itga6/Spi1/Mdk/Thbs1/Lbp/Edn3/Cyp7b1/P2ry12/Tlr2/Ptpn22/Csf1/Vav3/S1pr1/Vcam1/Lyn/Fut9/Csf3r/Ptafr/Padi2/Tnfrsf14/Spp1/Cmklr1/Selplg/Slc8b1/P2rx4/Sbds/Flt1/Hmgb1/Podxl/Ptn/Arhgef5/Rarres2/Dysf/Il17rc/Hrh1/Alox5/Cxcl12/Wnk1/Il17ra/C3ar1/Cd9/Eps8/C5ar2/C5ar1/Rps19/Tgfb1/Rpl13a/Swap70/Lyve1/Mapk3/Itgal/Hsd3b7/Pycard/Itgam/Cd81/Gas6/Mtus1/Vegfc/Ednra/Nod2/Mmp2/Cklf/Itgb1/Pdgfd/Icam1/St3gal4/Tirap/Adam10/Rhoa/Itga9/Oxsr1/Myd88/Cx3cr1/Ccr2/Cd99l2/Msn/Mospd2 | 140 |
| BP | GO:0050673 | epithelial cell proliferation | 174/3915 | 487/17474 | 8,2989E-12 | 1,2451E-09 | 9,2725E-10 | Eya1/Stat1/Cflar/Erbb4/Igfbp5/Sgpp2/Gli2/Glul/Dusp10/Prox1/Irf6/Gja1/Fabp7/Lims1/Col18a1/Igf1/Frs2/Stat6/Egfr/Flt4/Sparc/Adora2b/Serpinf1/Stxbp4/Stat3/Etv4/Grn/Itgb3/Sox9/Adam17/Id2/Prkd1/Sav1/Zfp36l1/Dicer1/Prl/Bmp6/Ptch1/Ctsl/Mef2c/Thbs4/Fst/Isl1/Plau/Adk/Wnt5a/Vstm4/Mmrn2/Bmpr1a/Bmp4/Ang/Bcl2l2/Rb1/Rgcc/Ednrb/Dab2/Rictor/Sema5a/Rida/Has2/Mtss1/Myc/Acvrl1/Sp1/Nlrc3/Prkdc/Tbx1/Dlg1/Cxadr/Hmgn1/Vegfa/Ptprm/Xdh/Mta3/Jcad/Zeb1/Lims2/Fgf1/Mcc/Nfatc1/Vegfb/Chuk/Tcf7l2/Nrarp/Notch1/Egfl7/Eng/Mdk/Pax6/Lgr4/Thbs1/B2m/Jag1/Id1/Ppp1r16b/Srsf6/Lama5/Pex2/Cyp7b1/Sox2/Fgf2/Notch2/Nras/F3/Enpep/Tgfbr1/Nfib/Tek/Tie1/Rps6ka1/Hspg2/Hes5/Cdk6/Hgf/Fgfr3/Fgfbp1/Rbpj/Kdr/Tgfbr3/Ncf1/Flt1/Hmgb1/Foxp2/Cav2/Cav1/Smo/Ptn/Fmc1/Atoh8/Dysf/Alms1/Tgfa/Gkn3/Gata2/Vhl/Alox5/Cxcl12/Ccnd2/Lrp6/Cdkn1b/C5ar2/C5ar1/Prkd2/Apoe/Ceacam1/Tgfb1/Zfp36/Saal1/Nupr1/Fgfr2/Htra1/Mki67/Igf2/Vegfc/Ednra/Nod2/Cdh1/Wfdc1/Cyba/Yap1/Bmper/Il18/Aldh1a2/Bmp5/Cd109/Tbx18/Hyal1/Ctnnb1/Wdr13/Apln/Gpc3/Prkx/Atp7a/Btk | 174 |
| BP | GO:0032760 | positive regulation of tumor necrosis factor production | 56/3915 | 107/17474 | 1,2643E-11 | 1,7988E-09 | 1,3396E-09 | Ly96/Arid5a/Ptprc/Fcgr3/Fcer1g/Cd84/Akap12/Ifngr1/Havcr2/Lgals9/Stat3/Tmem106a/Ripk1/Syk/Pik3r1/Isl1/Wnt5a/Clu/Zbtb20/Ticam1/Cd14/Csf1r/Jak2/Card9/Ifih1/Thbs1/Mavs/Lbp/Tlr2/Ccn1/Ripk2/Tlr4/Cyp2j6/Ptafr/Tlr1/Oas1a/Ptpn11/Hspb1/Hmgb1/Nod1/Tnfrsf1a/Clec7a/Tyrobp/Pycard/Tlr3/Nod2/Plcg2/Cyba/Tirap/Il18/Myd88/Ccr2/Ccr5/Cybb/Sash3/Btk | 56 |
| BP | GO:0032635 | interleukin-6 production | 77/3915 | 168/17474 | 1,3346E-11 | 1,8667E-09 | 1,3901E-09 | Arid5a/Il1rl2/Inpp5d/Rab7b/Prg4/Fcer1g/Cd84/Capn2/Bsg/Irak3/Havcr2/Adora2b/Nos2/Lgals9/Stat3/Tmem106a/Syk/F2r/Isl1/Wnt5a/Nckap1l/Nlrc3/Il1rap/Zbtb20/Cd200r1/Aif1/Trem2/Ticam1/Socs5/Aqp4/Ticam2/Cd74/Unc93b1/Afap1l2/Card9/Ifih1/Traf6/Mavs/Lbp/Tlr2/Il6ra/Ptpn22/Bank1/Ccn1/Ripk2/Tlr4/Laptm5/Ptafr/Hgf/Tlr1/Tlr6/Ptpn11/P2rx7/Hmgb1/Nod1/Il17rc/Il17ra/Ptpn6/Tnfrsf1a/C5ar2/Tyrobp/Arrb1/Trim30a/Pycard/Gas6/Tlr3/Klf2/Nod2/Plcg2/Cyba/Tirap/Il18/Myd88/Ccr5/Elf4/Btk/Tlr7 | 77 |
| BP | GO:0006909 | phagocytosis | 95/3915 | 224/17474 | 1,4899E-11 | 2,0491E-09 | 1,526E-09 | Gulp1/Slc11a1/Rab7b/Ptprc/Ncf2/Fcgr2b/Fcgr3/Fcer1g/Itgb2/Cnn2/Appl2/Myo1g/Dock2/Rack1/Itgb3/Pecam1/Cd300a/Pld4/Elmo1/Syk/Lman2/Xkr6/Ncf4/Ano6/Bin2/Nckap1l/Pros1/C4b/C2/Aif1/Trem2/Pot1b/Vav1/Rab31/Colec12/Ticam2/Megf10/Il15ra/Cd302/Itgav/Nr1h3/Thbs1/Spg11/Mertk/Hck/Tgm2/Lbp/Pik3ca/Tlr2/Pear1/Fcgr1/Rap1a/Abca1/Tlr4/Lepr/Xkr8/Pla2g5/Anxa3/P2rx7/Hmgb1/Dysf/Arhgap25/Gata2/Atg7/Clec7a/Tgfb1/Axl/Tyrobp/Siglece/Mfge8/Myo7a/P2ry6/Cln3/Itgal/Pycard/Itgam/Dock1/Gas6/Clcn3/Il15/Nod2/Plcg2/Irf8/Cyba/Itgb1/Dnm2/Ldlr/Plscr1/Plscr2/Myd88/Ccr2/Srpx/Cfp/Il2rg/Btk | 95 |
| BP | GO:0048660 | regulation of smooth muscle cell proliferation | 82/3915 | 186/17474 | 3,4729E-11 | 4,2773E-09 | 3,1853E-09 | Igfbp5/Rgs5/Ddr2/Timp3/Igf1/Frs2/Egfr/Alox12/Serpinf2/Stat5b/Itgb3/Gna13/Id2/Nqo2/Edn1/Ogn/Mef2c/Pik3r1/Bmpr1a/Bmp4/Ang/Ndrg2/Abcc4/Myc/Irak4/Prkdc/Apod/Adamts1/Cdkn1a/Aif1/Vegfa/Hbegf/Pdgfrb/Efemp2/Jak2/Tcf7l2/Calcrl/Traf6/Thbs1/Tgm2/Pik3ca/Fgf2/Shc1/Il6ra/Gnai3/S1pr1/Camk2d/Klf4/Tlr4/Ptafr/Ldlrap1/Ctnnbip1/Hes5/Nos3/Ppargc1a/Gper1/Flt1/Cav1/C3ar1/Cdkn1b/Apoe/Tgfb1/Igf1r/P2ry6/Ilk/Fgfr2/Hpgd/Il15/Mmp2/Cyba/Agt/Pdgfd/S1pr2/Cnn1/Il18/Gnai2/Rhoa/Tgfbr2/Myd88/Ctnnb1/Apln/Irak1 | 82 |
| BP | GO:0044242 | cellular lipid catabolic process | 93/3915 | 222/17474 | 5,4993E-11 | 6,4828E-09 | 4,8278E-09 | Idh1/Acadl/Irs1/Neu4/Dbi/Pla2g4a/Prdx6/Echdc1/Smpd2/Smpdl3a/Psap/Sgpl1/Ilvbl/Aldh1l2/Pex13/Gm2a/Acadvl/Pld2/Acox1/Lpin1/Pik3cg/Abcd4/Galc/Gpld1/Eci2/Hexb/Prkcd/Hacl1/Smpd5/Naga/Abcd2/Ehhadh/Acat3/Eci1/Cyp4f15/Cyp4f14/Cyp4f13/Pla2g7/Cyp1b1/Abhd3/Hsd17b4/Acaa2/Cpt1a/Plcb3/Phyh/Pnpla7/Ivd/Pla2g4e/Gpcpd1/Lpin3/Pex2/Pld1/Etfdh/Abcd3/Hadh/Bdh2/Acadm/Decr1/Acer2/Cpt2/Scp2/Echdc2/Mfsd2a/Pla2g5/Crot/Hadha/Hadhb/Plb1/Acox3/Acacb/Acads/Pla2g1b/Mgll/Apoc1/Lipe/Akt2/Sirt2/Etfb/Abhd2/Neu3/Echs1/Pnpla2/Asah1/Gcdh/Mt3/Nudt7/Plcg2/Ldlr/Bco2/Etfa/Hexa/Acad11/Hsd17b10 | 93 |
| BP | GO:0060326 | cell chemotaxis | 111/3915 | 282/17474 | 8,228E-11 | 9,4302E-09 | 7,0226E-09 | Fcgr3/Fcer1g/Itgb2/Bsg/Cxcl16/Ccl9/Ccl6/Adam17/Prkd1/Rin3/Lgmn/Nedd9/Edn1/Ninj1/Syk/Cxcl14/Thbs4/Itga1/Wnt5a/Prkcd/Lgals3/Ednrb/Gpr183/Sema5a/Rac2/Ano6/Bin2/Nckap1l/Crkl/Cxadr/Aif1/Pla2g7/Vegfa/Vav1/Hbegf/Fgf1/Lox/Cd74/Pdgfrb/Csf1r/Coro1b/Vegfb/Prkcq/Notch1/Eng/Dpp4/Spi1/Mdk/Thbs1/Lbp/Edn3/Cyp7b1/Fgf2/Rab13/Csf1/Vav3/S1pr1/Vcam1/Lef1/Lyn/Csf3r/Padi2/Hgf/Pdgfra/Kdr/Spp1/Cmklr1/Slc8b1/P2rx4/Sbds/Hspb1/Flt1/Hmgb1/Ptn/Arhgef5/Rarres2/Dysf/Il17rc/Hrh1/Alox5/Cxcl12/Wnk1/Il17ra/C3ar1/C5ar2/C5ar1/Prkd2/Rps19/Rpl13a/Swap70/Parva/Mapk3/Hsd3b7/Itgam/Gas6/Mtus1/Vegfc/Ednra/Gab1/Nod2/Mmp2/Cklf/Pdgfd/Tirap/Adam10/Ephb1/Itga9/Oxsr1/Cx3cr1/Ccr2/Mospd2 | 111 |
| BP | GO:0072001 | renal system development | 127/3915 | 336/17474 | 8,9917E-11 | 9,8933E-09 | 7,3676E-09 | Sox17/Eya1/Tfap2b/Stat1/Cflar/Erbb4/Bcl2/Gli2/Cfh/Pbx1/Vangl2/Kif26b/Cenpf/Enpp1/Psap/Sgpl1/Bicc1/Timeless/Gdf11/Hs3st3b1/Hs3st3a1/Lhx1/Itgb3/Ace/Sox9/Id2/Ttc8/Nid1/Gli3/Foxc1/Bmp6/Smad5/Ptch1/Adamts16/Mef2c/Vcan/Gcnt4/Adamts6/Wnt5a/Bmp4/Ednrb/Angpt1/Mtss1/Myc/Ift27/Mpst/Tns2/Dlg1/Adamts1/Vegfa/Epcam/Fgf1/Pdgfrb/Smad4/Smad2/Gcnt1/Plce1/Emx2/Notch1/Tsc1/Cntrl/Ctnnd1/Nup160/Lrp4/Cd44/Cat/Lgr4/Bcl2l11/Jag1/Bmp7/Lama5/Mecom/Fgf2/Frem2/Wwtr1/Tiparp/Il6ra/Notch2/Enpep/Npnt/Tet2/Tgfbr1/Invs/Tek/Nfia/Zmpste24/Heyl/Id3/Ctnnbip1/Hes5/Cc2d2a/Prom1/Pdgfra/Spp1/Pkd2/Cux1/C1galt1/Smo/Anxa4/Sec61a1/Klf15/Ret/C3ar1/Gpr4/Tgfb1/Tshz3/Ilk/Fgfr2/Angpt2/Ednra/Smad1/Sall1/Agt/Pdgfd/Yap1/Kank2/Bmper/Myo1e/Aldh1a2/Tbx18/Nphp3/Rhoa/Lamb2/Acvr2b/Ctnnb1/Gpc3/Amer1 | 127 |
| BP | GO:0002685 | regulation of leukocyte migration | 94/3915 | 228/17474 | 1,2375E-10 | 1,3437E-08 | 1,0006E-08 | Il1r1/Cnn2/Kitl/Stk10/Gp1ba/Lgals9/Aoc3/Itga2b/Itgb3/Pecam1/Cd300a/Adam17/Rin3/Lgmn/Nedd9/Edn1/Ninj1/Cxcl14/Thbs4/Pik3r1/Wnt5a/Lgals3/Mmp14/Ripk3/Lrch1/Rac2/Ano6/Nckap1l/Crkl/Apod/Cd200r1/Jam2/Myo1f/Aif1/Pla2g7/Vegfa/Trem2/Cd74/Csf1r/Vegfb/Gcnt1/Dock8/Dpp4/Spi1/Mdk/Thbs1/Lbp/Edn3/P2ry12/Tlr2/Ptpn22/Csf1/Lyn/Fut9/Ptafr/Padi2/Tnfrsf14/Cmklr1/Slc8b1/P2rx4/Hmgb1/Ptn/Rarres2/Dysf/Cxcl12/Wnk1/C3ar1/Cd9/C5ar2/C5ar1/Tgfb1/Swap70/Lyve1/Mapk3/Pycard/Cd81/Gas6/Mtus1/Vegfc/Ednra/Nod2/Pdgfd/Icam1/St3gal4/Tirap/Adam10/Rhoa/Oxsr1/Myd88/Cx3cr1/Ccr2/Cd99l2/Msn/Mospd2 | 94 |

| | | | | | | | | |
|---|---|---|---|---|---|---|---|---|
| BP | GO:0050867 | positive regulation of cell activation | 145/3915 | 400/17474 | 1,4362E-10 | 1,5077E-08 | 1,1228E-08 | Il1rl2/Igfbp2/Inpp5d/Bcl2/Sox13/Ptprc/Pla2g4a/Ddr2/Fcer1g/Itpkb/Dusp10/Ccn2/Vsir/Itgb2/Icosl/Igf1/Kitl/Stat6/Smarcc2/Plek/Havcr2/Irf1/Nlrp3/Adora2b/Tnfsf13/Pld2/Gp1ba/Lgals9/Smarce1/Stat5b/Lgals8/Gli3/Syk/Mef2c/Il6st/Adk/Wnt5a/Shld2/Pnp/Mmp14/Gpr183/Il7r/Lgals1/Arid2/Nckap1l/Prkdc/Tfrc/Snx4/Cd86/Cd80/Runx1/Cdkn1a/Rasal3/H2-Ob/H2-Aa/H2-Eb1/Aif1/Trem2/Ticam1/Socs5/Ticam2/Cd74/Pdgfrb/Csf1r/Kmt5b/Dock8/Smarca2/Jak2/Cd274/Acta2/Gpam/Prkcq/Il15ra/Rif1/Dpp4/Spi1/Mdk/Traf6/Cd59a/Thbs1/Capn3/B2m/Shld1/Lbp/Cd40/Actl6a/Cd1d1/Ptpn22/Vav3/Vcam1/Lef1/Bcl10/Ripk2/Exosc3/Tlr4/Ptafr/Rps6ka1/Pdpn/Tnfrsf14/Prkcz/Paxip1/Tnip2/Cd38/Tlr6/Rhoh/Ephb4/Hmgb1/Cav1/Smo/Gimap5/Vamp8/Gp9/Gata2/Clec7a/Nectin2/Tgfb1/Axl/Tyrobp/Flt3l/Klhl25/Ctsc/Il4ra/Itgal/Stx4a/Pycard/Itgam/Igf2/Cd81/Lamp1/Gas6/Il15/Nod2/Cbfb/Tirap/Il18/Rhoa/Tgfbr2/Myd88/Ccr2/Sash3/Atp11c/Flna/Il2rg/Btk/Sh3kbp1 | 145 |
| BP | GO:0033674 | positive regulation of kinase activity | 150/3915 | 423/17474 | 4,2655E-10 | 4,0458E-08 | 3,0129E-08 | Erbb4/Slc11a1/Irs1/Tnfrsf11a/Ralb/Chil1/Ptprc/Ddr2/Vangl2/Prox1/Tab2/Sash1/Igf1/Kitl/Erbb3/Egfr/Ccdc88a/Irgm1/Flt4/Igtp/Irgm2/Pik3r5/Traf4/Tom1l1/Abi3/Itgb3/Ace/Axin2/Cd300a/Adam17/Pik3cg/Nedd9/Edn1/Syk/Ntrk2/Mef2c/Map3k1/Nek10/Wnt5a/Prkcd/Fermt2/Ang/Ripk3/Clu/Rgcc/Bora/Ghr/Angpt1/Card10/Dazap2/Emp2/Fgd4/Dlg1/Cd86/Snx9/Cdkn1a/Fgd2/Vegfa/Trem2/Spdya/Nrxn1/Hbegf/Fgf1/Cd74/Pdgfrb/Csf1r/Adrb2/Zfp91/Jak2/F2/Traf6/Thbs1/Mertk/Rassf2/Tpx2/Sdc4/Cd40/Edn3/Pik3ca/Fgf2/Efna1/Il6ra/Jtb/Magi3/Rap1a/Csf1/Vav3/Pde5a/Cenpe/Sh3glb1/Ccn1/Lyn/Map3k7/Tlr4/Tek/Dab1/Lrp8/Tie1/Clspn/Prkcz/Reln/Fgfr3/Tlr1/Tlr6/Pdgfra/Kdr/Pkd2/P2rx7/Ephb4/Mmd2/Flt1/Arhgef5/Ezh2/Tgfa/Camk1/Ret/Ccnd2/Cdkn1b/Ceacam1/Tgfb1/Axl/Igf1r/Cib1/Iqgap1/Cemip/Fzd4/Ilk/Tead1/Gprc5b/Fgfr2/Cd81/Insr/Gas6/Tcim/Adam9/Nrg1/Vegfc/Slc27a1/Gab1/Nod2/Mt3/Agt/Pdgfd/Tirap/Il18/Ryk/Rhoa/Tgfbr2/Acvr2b/Irak1 | 150 |
| BP | GO:0019221 | cytokine-mediated signaling pathway | 136/3915 | 376/17474 | 6,322E-10 | 5,6096E-08 | 4,1774E-08 | Il1r1/Stat1/Tnfrsf11a/Ikbke/Adipor1/Ptprc/Fcer1g/Aim2/Ifngr1/Lims1/Appl2/Irak3/Stat6/Irgm1/Irf1/Igtp/Irgm2/Traf4/Rffl/Ccl9/Ccl6/Srsf1/Stat5b/Stat3/Adam17/Nfkbia/Traf3/Klf6/Foxc1/Ripk1/Edn1/Syk/Naip2/Naip5/Naip6/Il6st/Wnt5a/Ghr/Osmr/Lifr/Il7r/Angpt1/Csf2rb2/Csf2rb/Irak4/Il1rap/Parp14/Parp9/Ifnar2/Il10rb/Ifnar1/Crebrf/Trem2/Socs5/Sting1/Ticam2/Iigp1/Cd74/Csf1r/Smad4/Rela/Tjp2/Jak2/Chuk/Wbp1l/Il15ra/Notch1/Traf1/Nmi/Spi1/Traf6/Mavs/Cpne1/Samhd1/Il6ra/H2bc21/Hipk1/Rbm15/Csf1/Lyn/Ripk2/Il11ra1/Lepr/Csf3r/Laptm5/Padi2/Isg15/Tnip2/Rbm47/Oasl2/Oasl1/Oas1a/Trim56/Cav1/Irf5/Irak2/Cxcl12/Adipor2/Wnk1/Il17ra/Ptpn6/Tnfrsf1a/Ceacam1/Axl/Lsm14a/Irf3/Cib1/Ilk/Il4ra/Il21r/Mapk3/Pycard/Ifitm2/Ifitm1/Ifitm3/Sigirr/Irf7/Gas6/Ikbkb/Slc27a1/Gab1/Il15/Bbs2/Mt3/Nlrc5/Casp4/Yap1/Il18/Ccrl2/Oxsr1/Myd88/Cx3cr1/Ccr2/Il13ra1/Irak1/Il2rg | 136 |
| BP | GO:0070374 | positive regulation of ERK1 and ERK2 cascade | 89/3915 | 218/17474 | 6,9479E-10 | 6,0351E-08 | 4,4944E-08 | Cflar/Erbb4/Tnfrsf11a/Chil1/Ptprc/Slc30a10/Akap12/Ccn2/Igf1/Egfr/Havcr2/Flt4/Serpinf2/Ccl9/Ccl6/Itgb3/Dnajc27/Ntsr2/Adam17/Nqo2/Prxl2c/F2r/Fermt2/Bmp4/Gpr183/Angpt1/Crkl/Casr/Vegfa/Trem2/Nrxn1/Fgf1/Cd74/Pdgfrb/Csf1r/Vegfb/Acta2/Fgfbp3/Card9/Notch1/Cd44/Fgf2/P2ry1/Tlr2/Shc1/Notch2/Ptpn22/Rap1a/Camk2d/Npnt/Ripk2/Glipr2/Tlr4/Tek/Pla2g5/Prkcz/Fgfr3/Pdgfra/Kdr/Ptpn11/Gper1/Hmgb1/Gpnmb/Npy/Mturn/Nod1/Cxcl12/C5ar2/C5ar1/Prkd2/Apoe/Tgfb1/Cib1/Arrb1/P2ry6/Cavin3/Mapk3/Pycard/Fgfr2/Gas6/Nrg1/Ednra/Nod2/Mt3/Pdgfd/Icam1/Bmper/Tirap/Gnai2 | 89 |
| BP | GO:0032970 | regulation of actin filament-based process | 143/3915 | 401/17474 | 7,0432E-10 | 6,0542E-08 | 4,5086E-08 | Vil1/Lmod1/Myoc/F11r/Vangl2/Atp1a2/Prox1/Lats1/Ccn2/Arhgap18/Pln/Gja1/Ctnna3/Specc1l/Cnn2/Washc4/Plek/Ccdc88a/Sh3pxd2b/Pdlim4/Flii/Arhgef15/Serpinf2/Myo1c/Itgb3/Pecam1/Kcnj2/Cfl2/Ttc8/Carmil1/Dsp/Nedd9/Edn1/Mef2c/Pik3r1/Map3k1/Fam107a/Prkcd/Fermt2/Gmfb/Sorbs3/Rgcc/Rictor/Sema5a/Mtss1/Baiap2l2/Twf1/Rapgef3/Lima1/Nckap1l/Ppm1f/Dlg1/Pak2/Hcls1/Phldb2/Abi3bp/Snx9/Myo1f/Kank3/Cd2ap/Daam2/Svil/Rock1/Dsc2/Pdgfrb/Csf1r/Smad4/Coro1b/Kank1/Acta2/Add3/Prkcq/Phpt1/Tsc1/Mdk/Id1/Hck/Sdc4/Pik3ca/Tlr2/Arfip1/Kirrel/Gja5/Notch2/Casq2/Rhoc/Capza1/S1pr1/Synpo2/Tgfbr1/Tek/Wasf2/Asap3/Arhgef19/Pdpn/Akap9/Sri/Limch1/Hip1r/Eln/Arpc1b/Wasf3/Cav1/Arhgef5/Capg/Alms1/Arhgdib/Eps8/Vasp/Tgfb1/Plekhg2/Nphs1/Fxyd1/Rhpn2/Cyfip1/Tjp1/Fes/Iqgap1/Fchsd2/Arap1/Ilk/Swap70/Arhgap17/Pycard/Rnh1/Arhgef10/Dlc1/Bst2/Fhod1/Pard3/Icam1/Kank2/Esam/Rdx/Cgnl1/Tmod3/Rhoa/Clasp2/Was/Fgf13/Flna/Gpm6b/Arhgap6 | 143 |
| BP | GO:0045601 | regulation of endothelial cell differentiation | 27/3915 | 40/17474 | 1,4006E-09 | 1,1008E-07 | 8,1973E-08 | F11r/Btg1/Vezf1/Tmem100/Dicer1/Bmp6/Vcl/Bmp4/Acvrl1/Cldn5/Notch4/Vegfa/Xdh/Zeb1/Rock1/Notch1/Jag1/Id1/Tgfbr1/Atoh8/Vhl/Tnfrsf1a/Ceacam1/Ikbkb/Cdh5/S1pr2/Ctnnb1 | 27 |
| BP | GO:0030099 | myeloid cell differentiation | 154/3915 | 444/17474 | 1,5143E-09 | 1,1679E-07 | 8,6972E-08 | Stat1/Cflar/Casp8/Inpp5d/Tnfrsf11a/Rab7b/Fcer1g/Mndal/Ifi203/Itpkb/Lbr/Enpp1/Kitl/Ikzf1/Meis1/Hba-a1/Hba-a2/Ncor1/Gp1ba/Ccl9/Cdk5rap3/Stat5b/Stat3/Itgb3/Asxl2/Id2/Nfkbia/Zfp36l1/Ncapg2/Itgb8/Gpr137b/Ripk1/Nedd9/Ninj1/Sfxn1/Tifab/Smad5/Mef2c/Pik3r1/Prxl2a/Bmp4/Rb1/Gpr183/Dab2/Myc/Senp1/Rarg/Sp1/Nckap1l/Prkdc/Nrros/Tfrc/Hcls1/Gabpa/Runx1/Glo1/Pknox1/Adgrf5/Vegfa/Trem2/Epas1/Nr3c1/Lox/Rps14/Cd74/Csf1r/Nfatc1/Tcirg1/Tjp2/Jak2/Fas/Chuk/Pip4k2a/Spi1/Traf6/Lmo2/B2m/Rassf2/Jag1/Car2/Tlr2/Notch2/Cd101/Rbm15/Csf1/Lef1/Tet2/Lyn/Ptbp3/Tlr4/Tal1/Pabpc4/Csf3r/Wasf2/C1qc/Ctnnbip1/Prdm16/Isg15/Cdk6/Fgfr3/Rbpj/Tgfbr3/Ptpn11/Hmgb1/Bpgm/Clec5a/Gimap5/Mturn/Gp9/Gata2/Ptpn6/Ltbr/Clec2d/Rps19/Ceacam1/Tgfb1/Zfp36/Tyrobp/Cebpg/Lrrk1/Cib1/Fes/Ucp2/Sox6/Itgam/Cd81/Kat6a/Casp3/Klf2/Inpp4b/Il15/Dnase2a/Cbfb/Maf/Irf8/Abcb10/Fli1/Ets1/Tirap/Ubash3b/Tmod3/Cd109/Plscr1/Rbp1/Tgfbr2/Myd88/Ctnnb1/Ndp/Gpc3/Hmgb3/Flna/G6pdx/Heph/Alas2 | 154 |
| BP | GO:0051090 | regulation of DNA-binding transcription factor activity | 152/3915 | 437/17474 | 1,5483E-09 | 1,183E-07 | 8,8101E-08 | Cflar/Sp100/Tnfrsf11a/Ikbke/Rab7b/Pbx1/Ddr2/Fcgr2b/Cd84/Aim2/Prox1/Enpp1/Itgb2/Irak3/Ddit3/Adcy1/Havcr2/Nlrp3/Trim25/Spop/Cdk5rap3/Stat3/Fzd2/Trib2/Id2/Prkd1/Nfkbia/Sav1/Prkch/Traf3/Siva1/Gli3/Sfrp4/Ripk1/Edn1/Syk/Ptch1/Erbin/Wnt5a/Ripk3/Clu/Rb1/Rgcc/Zic2/Dap/Mtdh/Fzd6/Nfam1/Nlrc3/Il1rap/Tfrc/Hcls1/Trim26/Vegfa/Ticam1/Eif2ak2/Cyp1b1/Sting1/Ticam2/Camk2a/Lrp5/Rela/Jak2/Chuk/Tcf7l2/Prkcq/Card9/Traf1/Spi1/Traf6/Cat/Capn3/Ddrgk1/Mavs/Id1/Hck/Cd40/Ptgis/Tlr2/Bcl10/Cth/Ripk2/Klf4/Tlr4/Plpp3/Lrp8/Heyl/Id3/Ctnnbip1/Prkcz/Sri/Reln/Paxip1/Tacc3/Cytl1/Ppargc1a/Tlr6/Anxa3/Pkd2/Tgfbr3/Cmklr1/Cav1/Smo/Hipk2/Arhgef5/Ezh2/Mturn/Nod1/Anxa4/Irak2/Erc1/Kdm5a/Lrp6/Prkd2/Tgfb1/Cebpg/Igf1r/Cib1/Fzd4/Arrb1/Trim21/Trim68/Trim34a/Trim5/Trim12a/Trim12c/Trim30a/Sox6/Nupr1/Mapk3/Pycard/Sigirr/Gas6/Ikbkb/Kat6a/Tlr3/Nwd1/Nod2/Nlrc5/Plcg2/Icam1/Tirap/Il18/Hmgn3/Myd88/Cx3cr1/Ctnnb1/Ndp/Zic3/Irak1/Flna/Mid2 | 152 |
| BP | GO:0007159 | leukocyte cell-cell adhesion | 140/3915 | 395/17474 | 1,6913E-09 | 1,2574E-07 | 9,3635E-08 | Il1rl2/Igfbp2/Sox13/Ptprc/Cfh/Rc3h1/F11r/Itpkb/Dusp10/Vsir/Adora2a/Itgb2/Icosl/Igf1/Kitl/Smarcc2/Stk10/Havcr2/Irf1/Nlrp3/Gp1ba/Lgals9/Smarce1/Stat5b/Pecam1/Cd300a/Lgals8/Gli3/Hfe/Dusp22/Syk/Il6st/Adk/Bmp4/Lgals3/Pnp/Il7r/Has2/Rac2/Lgals1/Ppara/Arid2/Itga5/Nckap1l/Dlg1/Tfrc/Cd86/Cd80/Jam2/Runx1/Rasal3/H2-Ob/H2-Aa/H2-Eb1/Aif1/Twsg1/Socs5/Rock1/Cd74/Rela/Fermt3/Gcnt1/Dock8/Smarca2/Jak2/Cd274/Gpam/Prkcq/Nrarp/Ass1/Dpp4/Mdk/Traf6/Cd44/Cd59a/Thbs1/B2m/Sdc4/Bmp7/Pag1/Actl6a/Cd1d1/Tnfaip8l2/Ptpn22/Vcam1/Pde5a/Lef1/Slc39a8/Bcl10/Ripk2/Fut9/Klf4/Laptm5/Ptafr/Pla2g5/Tnfrsf14/Prkcz/Fgl2/Rhoh/Selplg/Hspb1/Ephb4/Hmgb1/Cav1/Gimap5/Gpnmb/Loxl3/Alox5/Cxcl12/Wnk1/Ptpn6/Lag3/Ceacam1/Tgfb1/Cd37/Klhl25/Lrrc32/Il4ra/Itgal/Pycard/Itgam/Igf2/Cd81/Casp3/Il15/Cbfb/Nfat5/Itgb1/Icam1/Ets1/St3gal4/Il18/Rhoa/Tgfbr2/Cx3cr1/Ccr2/Sash3/Irak1/Msn/Il2rg | 140 |

| | | | | | | | | | |
|---|---|---|---|---|---|---|---|---|---|
| BP | GO:0009615 | response to virus | 123/3915 | 336/17474 | 1,7935E-09 | 1,3214E-07 | 9,8407E-08 | Stat1/Bcl2/Gli2/Ikbke/Rab7b/Ptprc/Ivns1abp/Xpr1/Aim2/Mndal/Ifi203/Traf3ip3/Ifngr1/Ddx21/Polr3b/Irak3/Irgm1/Irf1/Igtp/Irgm2/Nlrp3/Nlrp1b/Lgals9/Rnf135/Slfn8/Trim25/Ifi27l2a/Dicer1/Traf3/Itgb8/Lgals8/Serinc5/Erbin/Wdfy4/Ripk3/Bnip3l/Ddx17/Apobec3/Polr3h/Gramd4/Rtp4/Parp9/Cd86/Cxadr/Ifnar2/Il10rb/Ifnar1/Rrp1b/Marchf2/Trim26/Ticam1/Eif2ak2/Ppm1b/Riok3/Rnf125/Sting1/Ticam2/Atg12/Unc93b1/Ifit3/Ifit1/Chuk/Gpam/Card9/Nmi/Ifih1/Mavs/Samhd1/Serinc3/Cd40/Znfx1/Vapb/Ptpn22/Nfkb1/Gbp7/Gbp3/Zmpste24/Isg15/Fgl2/Gbp4/Oasl2/Oasl1/Oas1a/Trim56/Irf5/Zc3hav1/Vamp8/Atg7/Il17ra/Apobec1/Abcc9/Exosc5/Tgfb1/Hnrnpul1/Lsm14a/Irf3/Cd37/Trim21/Trim34a/Trim5/Trim12a/Trim12c/Trim30a/Pycard/Htra1/Ifitm2/Ifitm1/Ifitm3/Irf7/Tlr3/Ddx60/Bst2/Il15/Elmod2/Mlkl/Plscr1/Plscr2/Hyal1/Setd2/Myd88/Tlr13/Mid2/Tlr7 | 123 |
| BP | GO:0050731 | positive regulation of peptidyl-tyrosine phosphorylation | 77/3915 | 184/17474 | 2,5638E-09 | 1,8397E-07 | 1,37E-07 | Erbb4/Ptprc/Fyn/Igf1/Kitl/Erbb3/Vtn/Abi3/Itgb3/Ace/Pecam1/Adam17/Bmp6/Nedd9/Syk/Thbs4/Il6st/Isl1/Ghr/Rictor/Angpt1/Itga5/Pak2/Parp14/Parp9/Hcls1/Cd80/Vegfa/Trem2/Hbegf/Cd74/Csf1r/Vegfb/Jak2/Gfra1/Ctnnd1/Lrp4/Cd44/Ehd4/Cd40/Efna1/Il6ra/Bank1/Lyn/Ripk2/Tlr4/Plpp3/Lrp8/Tnfrsf14/Hes5/Hgf/Reln/Yes1/Fgfr3/Pdgfra/Ptpn11/Cav2/Ptprz1/Tgfa/Tnfrsf1a/Tgfb1/Lrrk1/Iqgap1/Gprc5b/Ctf1/Igf2/Cd81/Gas6/Nrg1/Il15/Nod2/Arl2bp/Plcg2/Agt/Itgb1/Icam1/Il18 | 77 |
| BP | GO:0002460 | adaptive immune response based on somatic recombination of immune receptors built from immunoglobulin superfamily domains | 119/3915 | 324/17474 | 2,6193E-09 | 1,8633E-07 | 1,3876E-07 | Arid5a/Il1r1/Slc11a1/Inpp5d/Ptprc/Dennd1b/Cfh/Pla2g4a/Rc3h1/Fcgr2b/Fcgr3/Fcer1g/Cr1l/Enpp1/Vsir/Icosl/Stat6/Myo1g/Havcr2/Irf1/Nlrp3/Tnfsf13/Tmem98/Stat3/Adam17/Hfe/Dusp22/Serpinb9/Mef2c/Prkcd/Shld2/Pnp/Ripk3/Il7r/Csf2rb2/Csf2rb/Nckap1l/Emp2/Klhl6/Dlg1/Tfrc/H2-K1/Tap2/H2-Ob/H2-Aa/H2-Eb1/C4b/C2/H2-D1/H2-Q4/H2-T23/H2-T22/Vegfa/Trem2/Socs5/Cd74/Kmt5b/Tcirg1/Unc93b1/Jak2/Cd274/Fas/Prkcq/Card9/Notch1/Rif1/Dpp4/Serping1/Traf6/B2m/Shld1/Jag1/Cd40/Cd1d1/Il6ra/Rorc/Fcgr1/Lef1/Bcl10/Ripk2/Map3k7/Bach2/Exosc3/Csf3r/C1qb/C1qc/C1qa/Tnfrsf1b/Prkcz/Fgl2/Paxip1/P2rx7/Hmgb1/Gimap5/Loxl3/Il17ra/C3ar1/Ptpn6/Cracr2a/Mill2/Nectin2/Tgfb1/Ctsc/Swap70/Il4ra/Il21r/Irf7/Cd81/Nod2/Icam1/Il18/Rora/Ctsh/Parp3/Myd88/Ccr2/Was/Sash3/Btk | 119 |
| BP | GO:0051249 | regulation of lymphocyte activation | 167/3915 | 494/17474 | 2,684E-09 | 1,893E-07 | 1,4097E-07 | Il1rl2/Igfbp2/Inpp5d/Bcl2/Sox13/Ptprc/Rc3h1/Fcgr2b/Itpkb/Dusp10/Vsir/Adora2a/Icosl/Fbxo7/Igf1/Kitl/Stat6/Smarcc2/Havcr2/Irf1/Nlrp3/Tnfrsf13b/Tnfsf13/Lgals9/Smarce1/Stat5b/Cd300a/Id2/Zfp36l1/Dicer1/Lgals8/Gli3/Hfe/Dusp22/Nedd9/Syk/Ctla2a/Mef2c/Il6st/Adk/Duxbl1/Shld2/Bmp4/Lgals3/Pnp/Mmp14/Ripk3/Gpr183/Il7r/Rac2/Lgals1/Nfam1/Arid2/Nckap1l/Prkdc/Dlg1/Tfrc/Cd86/Cd80/Runx1/Cdkn1a/Rasal3/H2-Ob/H2-Aa/H2-Eb1/Aif1/Ticam1/Twsg1/Zfp36l2/Socs5/Zeb1/Ticam2/Cd74/Csf1r/Kmt5b/Dock8/Smarca2/Jak2/Cd274/Fas/Gpam/Prkcq/Il15ra/Nrarp/Rif1/Dpp4/Spi1/Mdk/Traf6/Cd44/Cd59a/B2m/Mertk/Shld1/Sdc4/Cd40/Pag1/Actl6a/Cd1d1/Rorc/Tnfaip8l2/Ptpn22/Vav3/Vcam1/Pde5a/Lef1/Bank1/Bcl10/Lyn/Ripk2/Exosc3/Tlr4/Laptm5/Themis2/Pla2g5/Tnfrsf1b/Tnfrsf14/Prkcz/Fgl2/Paxip1/Tnip2/Cd38/Rhoh/Hspb1/Ephb4/Hmgb1/Cav1/Gimap5/Gpnmb/Loxl3/Ptpn6/Lag3/Clec7a/Pglyrp1/Ceacam1/Tgfb1/Axl/Tyrobp/Flt3l/Cd37/Klhl25/Lrrc32/Il4ra/Itgal/Pycard/Igf2/Cd81/Lamp1/Gas6/Casp3/Il15/Nod2/Cbfb/Tirap/Il18/Parp3/Rhoa/Tgfbr2/Myd88/Ctnnb1/Ccr2/Sash3/Atp11c/Hmgb3/Il2rg/Btk/Sh3kbp1 | 167 |
| BP | GO:0072593 | reactive oxygen species metabolic process | 91/3915 | 230/17474 | 2,9884E-09 | 2,038E-07 | 1,5177E-07 | Cflar/Bcl2/Ncf2/Prdx6/Slc30a10/Ccn2/Ccn6/Fyn/Sesn1/Itgb2/Egfr/Hba-a1/Sh3pxd2b/Nos2/Stat3/Brca1/Acox1/Dcxr/Hbp1/Fbln5/Nqo2/Ripk1/Edn1/Syk/Dhfr/Nnt/Plau/Prkcd/Ripk3/Ncf4/Tspo/Ppara/Abcd2/Nrros/Cdkn1a/Cbs/Ier3/Vav1/Xdh/Cyp1b1/Pdgfrb/Rfk/Pla2r1/Nfe2l2/F2/Cat/Thbs1/Foxo1/Shc1/Gnai3/Ccn1/Map3k7/Tlr4/Prdx1/Slc25a33/Nos3/Tlr6/Aldh2/P2rx7/Ncf1/Pon3/Pdk4/Hk2/Alox5/Tgfb1/Sirt2/Tyrobp/Prcp/Ndufc2/Ucp2/Hbb-bt/Trim30a/Itgam/Sirt3/Insr/Clcn3/Mt3/Nqo1/Plcg2/Cyba/Agt/Birc3/Bco2/Il18/Gnai2/Rhoa/Tgfbr2/Cybb/G6pdx/Ogt/Atp7a | 91 |
| BP | GO:0007015 | actin filament organization | 152/3915 | 443/17474 | 4,4913E-09 | 2,9415E-07 | 2,1905E-07 | Myo1b/Vil1/Bcl2/Lmod1/Hmcn1/Myoc/F11r/Prox1/Lats1/Ccn2/Arhgap18/Mical1/Gja1/Ctnna3/Pcdh15/Specc1l/Washc4/Hsp90b1/Myo1g/Plek/Ccdc88a/Sh3pxd2b/Pdlim4/Flii/Arhgef15/Serpinf2/Myo1c/Rflnb/Tnfaip1/Myo19/Pecam1/Cfl2/Ttc8/Elmo1/Carmil1/Nedd9/Phactr1/Zbed3/Pik3r1/Map3k1/Fam107a/Prkcd/Fermt2/Gmfb/Sorbs3/Lcp1/Rgcc/Rictor/Sema5a/Mtss1/Rac2/Baiap2l2/Twf1/Rapgef3/Lima1/Nckap1l/Emp2/Ppm1f/Dlg1/Pak2/Itgb5/Hcls1/Phldb2/Snx9/Ezr/Myo1f/Kank3/Aif1/Cd2ap/Daam2/Svil/Ctnna1/Spire1/Coro1b/Kank1/Sorbs1/Add3/Nrap/Prkcq/Nebl/Tsc1/Aif1l/Nostrin/Wipf1/Ttn/Id1/Sdc4/Pik3ca/Clrn1/Tlr2/Arfip1/Kirrel/Rhoc/Capza1/S1pr1/Synpo2/Tpm2/Tgfbr1/Sh3d21/Wasf2/Asap3/Limch1/Hip1r/Eln/Hip1/Arpc1b/Wasf3/Cald1/Arhgef5/Capg/Alms1/Arhgap25/Eps8/Vasp/Plekhg2/Nphs1/Rhpn2/Cyfip1/Tjp1/Myo7a/Arrb1/Fchsd2/Arap1/Inppl1/Swap70/Arhgap17/Pycard/Rnh1/Arhgef10/Dlc1/Fat1/Pdlim3/Tpm4/Fhod1/Casp4/Icam1/Kank2/Esam/Rdx/Pstpip1/Myo1e/Cgnl1/Tmod3/Myo6/Rhoa/Clasp2/Shroom4/Was/Flna/Gdpd2/Sh3kbp1/Arhgap6 | 152 |
| BP | GO:0001649 | osteoblast differentiation | 82/3915 | 203/17474 | 5,7837E-09 | 3,7179E-07 | 2,7687E-07 | Igfbp5/Gli2/Suco/Myoc/Ddr2/H3f3a/Enpp1/Asf1a/Gja1/Igf1/Fignl1/Sh3pxd2b/Col1a1/Nbr1/Wnt3/Axin2/Sox9/H3f3b/Id2/Prkd1/Smoc1/Gli3/Bmp6/Id4/Smad5/Ptch1/Mef2c/Il6st/Gdf10/Bmpr1a/Fermt2/Bmp4/Ranbp3l/Cthrc1/Hdac7/Clic1/Vegfa/Twsg1/Lox/Smad4/Nfatc1/Lrp5/Tcirg1/Rorb/Notch1/Lgr4/Rassf2/Jag1/Id1/Trp53inp2/Zhx3/Bmp7/Sox2/Fgf2/Wwtr1/Npnt/Bmpr1b/Ccn1/Map3k7/Id3/Ctnnbip1/Cdk6/Atraid/Rest/Spp1/Tgfbr3/Tmem119/Smo/Clec5a/Ilk/Ifitm1/Igf2/Vegfc/Smad1/Cbfb/Yap1/Pth1r/Acvr2b/Ctnnb1/Limd1/Gdpd2/Hdac8 | 82 |
| BP | GO:0032956 | regulation of actin cytoskeleton organization | 126/3915 | 354/17474 | 8,2001E-09 | 4,9392E-07 | 3,6782E-07 | Vil1/Lmod1/Myoc/F11r/Vangl2/Prox1/Lats1/Ccn2/Arhgap18/Gja1/Specc1l/Washc4/Plek/Ccdc88a/Sh3pxd2b/Pdlim4/Flii/Arhgef15/Serpinf2/Myo1c/Itgb3/Pecam1/Cfl2/Ttc8/Carmil1/Nedd9/Edn1/Mef2c/Pik3r1/Map3k1/Fam107a/Prkcd/Fermt2/Gmfb/Sorbs3/Rgcc/Rictor/Sema5a/Mtss1/Baiap2l2/Twf1/Rapgef3/Lima1/Nckap1l/Ppm1f/Dlg1/Pak2/Hcls1/Phldb2/Abi3bp/Snx9/Myo1f/Kank3/Cd2ap/Daam2/Svil/Rock1/Pdgfrb/Csf1r/Smad4/Coro1b/Kank1/Add3/Prkcq/Phpt1/Tsc1/Mdk/Id1/Hck/Sdc4/Pik3ca/Tlr2/Arfip1/Kirrel/Notch2/Rhoc/Capza1/S1pr1/Synpo2/Tgfbr1/Tek/Wasf2/Asap3/Arhgef19/Limch1/Hip1r/Eln/Arpc1b/Wasf3/Arhgef5/Capg/Alms1/Arhgdib/Eps8/Vasp/Tgfb1/Plekhg2/Nphs1/Rhpn2/Cyfip1/Tjp1/Fes/Iqgap1/Fchsd2/Arap1/Ilk/Swap70/Arhgap17/Pycard/Rnh1/Arhgef10/Dlc1/Bst2/Fhod1/Icam1/Kank2/Esam/Rdx/Cgnl1/Tmod3/Rhoa/Clasp2/Was/Flna/Gpm6b/Arhgap6 | 126 |
| BP | GO:0044282 | small molecule catabolic process | 121/3915 | 337/17474 | 9,0535E-09 | 5,3748E-07 | 4,0026E-07 | Adhfe1/Hibch/Acadl/Cyp27a1/Irs1/Dbi/Npl/Echdc1/Ddo/Ilvbl/Aldh1l2/Shmt2/Pex13/Shmt1/Acadvl/Blmh/Nos2/Acox1/Dcxr/Lpin1/Aldh6a1/Abcd4/Gstz1/Aldh5a1/Nqo2/Eci2/Hexb/Mccc2/Hacl1/Glud1/Pnp/Oxct1/Abcd2/Pfkm/Csad/Abat/Prodh/Ehhadh/Acat3/Eci1/Glo1/Cbs/Cyp4f15/Cyp4f14/Cyp4f13/Ddah2/Enpp4/Xdh/Abhd3/Gnpda1/Cdo1/Hsd17b4/Acaa2/Cpt1a/Asrgl1/Aldh1a1/Gldc/Phyh/Gad2/Sardh/Gad1/Ivd/Lpin3/Pex2/Etfdh/Slc16a1/Dbt/Abcd3/Hadh/Bdh2/Adh5/Ddah1/Acadm/Decr1/Cpt2/Scp2/Echdc2/Mfsd2a/Hmgcl/Aldh4a1/Pgd/Crot/Nos3/Hadha/Hadhb/Khk/Acox3/Idua/Acacb/Acads/Sdsl/Pon3/Aass/Hibadh/Thnsl2/Aldh1l1/Dera/Apoe/Lipe/Bckdha/Akt2/Etfb/Dhdh/Bcat2/Abhd2/Hsd3b7/Acadsb/Oat/Echs1/Gcdh/Mt3/Nudt7/Gcsh/Acad8/Etfa/Hexa/Acad11/Amt/Ids/Renbp/Hsd17b10 | 121 |

| | | | | | | | | | |
|---|---|---|---|---|---|---|---|---|---|
| BP | GO:0002697 | regulation of immune effector process | 144/3915 | 419/17474 | 1,0138E-08 | 5,9753E-07 | 4,4498E-07 | Arid5a/Il1r1/Ptprc/Dennd1b/Cfh/Rc3h1/Fcgr2b/Fcgr3/Fcer1g/Cd84/Dusp10/Cr1l/Vsir/Ddx21/Itgb2/Appl2/Irak3/Stat6/Havcr2/Irf1/Nlrp3/Adora2b/Tnfsf13/Pld2/Lgals9/Stat5b/Grn/Cd300a/Hfe/Dusp22/Serpinb9/Syk/Wnt5a/Shld2/Lgals3/Pnp/Ripk3/Lacc1/Il7r/Angpt1/Rac2/Tmbim6/Nckap1l/Litaf/Prkdc/Tfrc/Snx4/Cd86/H2-K1/Tap1/Tap2/C4b/C2/H2-D1/H2-Q4/H2-T23/H2-T22/Trem2/Ticam1/Vav1/Socs5/Ticam2/Cd74/Kmt5b/Card9/Rif1/Dpp4/Serping1/Spi1/Traf6/Cd59a/B2m/Mavs/Shld1/Lbp/Cd40/Tlr2/Cd1d1/Fcgr1/Ptpn22/Rap1a/Lyn/Ripk2/Map3k7/Exosc3/Tlr4/Tek/Laptm5/Ptafr/Pla2g5/Tnfrsf1b/Tnfrsf14/Prkcz/Fgl2/Paxip1/Ankrd17/P2rx7/Ncf1/Hmgb1/Gimap5/Tril/Nod1/Vamp8/Loxl3/Gata2/A2m/Ptpn6/Lag3/Clec2d/Mill2/Pglyrp1/Nectin2/Rps19/Ceacam1/Tgfb1/Axl/Tyrobp/Cd37/Fes/Gprc5b/Il4ra/Stx4a/Pycard/Itgam/Igf2/Cd81/Lamp1/Tlr3/Ddx60/Bst2/Nod2/Plcg2/Casp4/Tirap/Il18/Parp3/Myd88/Cx3cr1/Ccr2/Was/Cfp/Sash3/Btk/Tlr7 | 144 |
| BP | GO:0048754 | branching morphogenesis of an epithelial tube | 77/3915 | 190/17474 | 1,4067E-08 | 8,175E-07 | 6,0879E-07 | Eya1/Bcl2/Gli2/Pbx1/Vangl2/Igf1/Tmtc3/Timeless/Hs3st3b1/Hs3st3a1/Lhx1/Mks1/Etv4/Gna13/Sox9/Dicer1/Gli3/Edn1/Ptch1/Adamts16/Wnt5a/Bmp4/Mmp14/Sema5a/Angpt1/Myc/Tbx1/Dlg1/Casr/Notch4/Vegfa/Lama1/Fgf1/Smad4/Nfatc1/Lrp5/Nrarp/Notch1/Eng/Mdk/Cd44/Lgr4/Bmp7/Ctsz/Lama5/Sox2/Fgf2/Rbm15/Csf1/Lef1/Npnt/Tnc/Tek/Tie1/Clic4/Ctnnbip1/Pdgfra/Kdr/Tbx3/Flt1/Smo/Cxcl12/Lrp6/Mgp/Tgfb1/Ilk/Fgfr2/Ednra/Hhip/Sall1/Nfatc3/Agt/Yap1/Ctsh/Tgfbr2/Ctnnb1/Gpc3 | 77 |
| BP | GO:0048762 | mesenchymal cell differentiation | 93/3915 | 243/17474 | 1,4211E-08 | 8,1919E-07 | 6,1005E-07 | Nrp2/Erbb4/Fn1/Bcl2/Epb41l5/Adipor1/Vangl2/Hey2/Gja1/Timp3/Kitl/Rflnb/Tmem100/Col1a1/Axin2/Sox9/Dicer1/Foxc1/Edn1/Fam172a/Mef2c/Isl1/Il17rd/Tasor/Wnt5a/Sema3g/Bmpr1a/Fermt2/Bmp4/Rgcc/Ednrb/Zic5/Zic2/Dab2/Ranbp3l/Sema5a/Has2/Ddx17/Vasn/Emp2/Tbx1/Phldb2/Notch4/Vegfa/Trip10/Rock1/Sema6a/Smad4/Smad2/Nfatc1/Tcf7l2/Notch1/Eng/Mdk/Pax6/Spred1/Sema6d/Jag1/Bmp7/Edn3/Lama5/Wwtr1/Efna1/Lef1/Glipr2/Tgfbr1/Akna/Heyl/Phactr4/Pdpn/Sema3d/Sema3c/Rbpj/Tgfbr3/Tbx3/Smo/Ezh2/Loxl3/Ret/Lrp6/Plaur/Tgfb1/Fuz/Mapk3/Tgfb1i1/Fgfr2/Ednra/Aldh1a2/Clasp2/Tgfbr2/Ctnnb1/Flna/Amer1 | 93 |
| BP | GO:0002443 | leukocyte mediated immunity | 143/3915 | 419/17474 | 1,8855E-08 | 1,0373E-06 | 7,7247E-07 | Arid5a/Il1r1/Slc11a1/Inpp5d/Ptprc/Dennd1b/Cfh/Fcgr2b/Fcgr3/Fcer1g/Cd84/Cr1l/Enpp1/Vsir/Ddx21/Itgb2/Icosl/Stat6/Myo1g/Havcr2/Nlrp3/Adora2b/Tnfsf13/Pld2/Lgals9/Stat5b/Ace/Cd300a/Hfe/Dusp22/Serpinb9/Syk/Prkcd/Shld2/Pnp/Ripk3/Il7r/Csf2rb2/Csf2rb/Rac2/Irak4/Nckap1l/Emp2/Dlg1/Tfrc/Snx4/Myo1f/H2-K1/Tap1/Tap2/H2-Ob/H2-Aa/H2-Eb1/C4b/C2/H2-D1/H2-Q4/H2-T23/H2-T22/Trem2/Ticam1/Vav1/Kif5b/Ticam2/Cd74/Kmt5b/Tcirg1/Unc93b1/Fas/Ptgds/Card9/Rif1/Dpp4/Serping1/Spi1/F2/Traf6/Snap23/B2m/Mavs/Shld1/Jag1/Cd40/Tlr2/Cd1d1/Fcgr1/Bcl10/Lyn/Map3k7/Exosc3/Tlr4/Prdx1/Csf3r/Ptafr/C1qb/C1qc/C1qa/Tnfrsf1b/Prkcz/Fgl2/Paxip1/Anxa3/P2rx7/Ncf1/Lat2/Hmgb1/Gimap5/Vamp8/Gata2/Ptpn6/Lag3/Clec2d/Mill2/Nectin2/Ceacam1/Tgfb1/Tyrobp/Cebpg/Fes/Ctsc/Swap70/Il4ra/Il21r/Stx4a/Itgam/Irf7/Igf2/Cd81/Lamp1/Tlr3/Bst2/Nod2/Plcg2/Icam1/Il18/Ctsh/Parp3/Myd88/Cx3cr1/Ccr2/Was/Sash3/Btk | 143 |
| BP | GO:0060348 | bone development | 94/3915 | 249/17474 | 2,6085E-08 | 1,3978E-06 | 1,0409E-06 | Col9a1/Rab23/Col3a1/Nab1/Ptprc/Cfh/Myoc/Ddr2/Enpp1/Gja1/Col13a1/Timp3/Igf1/Meis1/Sh3pxd2b/Sparc/Gp1ba/Rflnb/Col1a1/Axin2/Sox9/Notum/Asxl2/Foxn3/Fbln5/Trip11/Gli3/Sfrp4/Foxc1/Bmp6/Ninj1/Ogn/Tifab/Smad5/Mef2c/Bmp4/Mmp14/Ranbp3l/Adamts12/Has2/Ano6/Rarg/Bbx/Cbs/Lox/Lrp5/Tjp2/Papss2/Pip4k2a/Thbs1/Notch2/Bmpr1b/Map3k7/Tmem38b/Col27a1/Frem1/Bnc2/Lepr/Tal1/Zmpste24/Wasf2/Hspg2/Alpl/Dhrs3/Insig1/Fgfr3/Kdr/Idua/Tmem119/Ptpn11/Sbds/Gp9/Ptpn6/Tulp3/Lrp6/Tyrobp/Chsy1/Lrrk1/Serpinh1/Inppl1/Xylt1/Vkorc1/Fgfr2/Csgalnact1/Slc10a7/Smad1/Fli1/Rhoa/Plxnb1/Tgfbr2/Ebp/Bgn/Flna/Amer1 | 94 |
| BP | GO:0060541 | respiratory system development | 94/3915 | 249/17474 | 2,6085E-08 | 1,3978E-06 | 1,0409E-06 | Eya1/Col3a1/Igfbp5/Gli2/Chil1/Pou2f1/Vangl2/Hsd11b1/Ccn2/Igf1/Tmtc3/Timeless/Tns3/Flt4/Sparc/Ace/Sox9/Sav1/Numb/Dicer1/Gli3/Wnt5a/Bmpr1a/Bmp4/Mmp14/Rarg/Sp1/Heg1/Vegfa/Lama1/Smchd1/Epas1/Fgf1/Lox/Pdgfrb/Smad2/Notch1/Rbbp9/Id1/Srsf6/Ctsz/Lama5/Myt1/Sox2/Fgf2/Slc7a11/Rxfp1/Gpsm2/Lef1/Chd7/Wwp1/Tmem38b/Tnc/Nfib/Pdpn/Trp73/Nos3/Fgfr3/Rbpj/Pdgfra/Kdr/Hopx/Cux1/Plod3/Hmgb1/Foxp2/Loxl3/Tulp3/Lrp6/Mgp/Tgfb1/Tshz3/Fuz/Rcn3/Rpl13a/Insc/Mapk3/Fgfr2/Dhcr7/Ano1/Asah1/Klf2/Hhip/Atxn1l/Yap1/Aldh1a2/Ctsh/Nphp3/Tgfbr2/Acvr2b/Ctnnb1/Gpc3/Zic3/Atp7a | 94 |
| BP | GO:0032102 | negative regulation of response to external stimulus | 137/3915 | 400/17474 | 2,9237E-08 | 1,5294E-06 | 1,1389E-06 | Serpine2/Rab7b/Tnr/Fcgr2b/Mndal/Ifi203/Dusp10/Gja1/Fabp7/Adora2a/Igf1/Irak3/Kremen1/Havcr2/Nlrp3/Alox12/Gp1ba/Serpinf1/Serpinf2/Vtn/Lgals9/Grn/Wnt3/C1qtnf1/Rin3/Serpinb9/Ctla2a/Slc6a3/Erbin/Isl1/Nr1d2/Plau/Wnt5a/Prkcd/Sema3g/Rb1/Sema5a/Ppara/Gramd4/Nlrc3/Abat/St6gal1/Parp14/Cd200r1/Phldb2/Pros1/Tap1/Tap2/Aif1/Ier3/Trem2/Ppm1b/Socs5/Nrxn1/Riok3/Rnf125/Gpr17/Atg12/Sema6a/Adrb2/Coro1b/Notch1/Nmi/Dpp4/Calcrl/Tfpi/Serping1/Nr1h3/F2/Mdk/Cd44/Thbs1/Sema6d/Thbd/Samhd1/Ptgis/Anxa5/Fgf2/Tnfaip8l2/Cers2/Ptpn22/Nfkb1/Slc39a8/Cldn19/Pla2g5/Padi2/Tnfrsf1b/Isg15/Sema3d/Hgf/Sema3c/Fgl2/Pdgfra/Mvk/Oas1a/Gper1/Npy/Grid2/Alox5/A2m/Lpcat3/Tnfrsf1a/Cd9/Clec2d/C5ar2/Pglyrp1/Apoe/Plaur/Rps19/Ceacam1/Tgfb1/Zfp36/Sirt2/Siglece/Trim21/Xylt1/Htra1/Igf2/Angpt2/Plat/Nod2/Bbs2/Nlrc5/Cdh5/Wfdc1/Ldlr/Ets1/Ubash3b/Neo1/Uaca/Rora/Anxa2/Cd109/Ryk/Clasp2/Cx3cr1/Elf4 | 137 |
| BP | GO:0051607 | defense response to virus | 102/3915 | 277/17474 | 3,0233E-08 | 1,5496E-06 | 1,154E-06 | Stat1/Bcl2/Rab7b/Ptprc/Aim2/Mndal/Ifi203/Traf3ip3/Ifngr1/Ddx21/Polr3b/Irf1/Nlrp3/Nlrp1b/Rnf135/Slfn8/Trim25/Dicer1/Traf3/Serinc5/Erbin/Ripk3/Bnip3l/Ddx17/Apobec3/Polr3h/Gramd4/Rtp4/Parp9/Cd86/Cxadr/Ifnar2/Il10rb/Ifnar1/Marchf2/Trim26/Ticam1/Eif2ak2/Ppm1b/Riok3/Rnf125/Sting1/Ticam2/Atg12/Unc93b1/Ifit3/Ifit1/Chuk/Gpam/Card9/Ifih1/Mavs/Samhd1/Serinc3/Cd40/Znfx1/Vapb/Ptpn22/Gbp7/Gbp3/Zmpste24/Isg15/Fgl2/Gbp4/Oasl2/Oasl1/Oas1a/Trim56/Irf5/Zc3hav1/Vamp8/Atg7/Apobec1/Abcc9/Exosc5/Lsm14a/Irf3/Cd37/Trim21/Trim34a/Trim5/Trim12a/Trim12c/Trim30a/Pycard/Htra1/Ifitm2/Ifitm1/Ifitm3/Irf7/Tlr3/Ddx60/Bst2/Il15/Elmod2/Mlkl/Plscr1/Plscr2/Setd2/Myd88/Mid2/Tlr7 | 102 |
| BP | GO:0140546 | defense response to symbiont | 102/3915 | 277/17474 | 3,0233E-08 | 1,5496E-06 | 1,154E-06 | Stat1/Bcl2/Rab7b/Ptprc/Aim2/Mndal/Ifi203/Traf3ip3/Ifngr1/Ddx21/Polr3b/Irf1/Nlrp3/Nlrp1b/Rnf135/Slfn8/Trim25/Dicer1/Traf3/Serinc5/Erbin/Ripk3/Bnip3l/Ddx17/Apobec3/Polr3h/Gramd4/Rtp4/Parp9/Cd86/Cxadr/Ifnar2/Il10rb/Ifnar1/Marchf2/Trim26/Ticam1/Eif2ak2/Ppm1b/Riok3/Rnf125/Sting1/Ticam2/Atg12/Unc93b1/Ifit3/Ifit1/Chuk/Gpam/Card9/Ifih1/Mavs/Samhd1/Serinc3/Cd40/Znfx1/Vapb/Ptpn22/Gbp7/Gbp3/Zmpste24/Isg15/Fgl2/Gbp4/Oasl2/Oasl1/Oas1a/Trim56/Irf5/Zc3hav1/Vamp8/Atg7/Apobec1/Abcc9/Exosc5/Lsm14a/Irf3/Cd37/Trim21/Trim34a/Trim5/Trim12a/Trim12c/Trim30a/Pycard/Htra1/Ifitm2/Ifitm1/Ifitm3/Irf7/Tlr3/Ddx60/Bst2/Il15/Elmod2/Mlkl/Plscr1/Plscr2/Setd2/Myd88/Mid2/Tlr7 | 102 |
| BP | GO:0002274 | myeloid leukocyte activation | 92/3915 | 243/17474 | 3,114E-08 | 1,5862E-06 | 1,1812E-06 | Fn1/Slc11a1/Pla2g4a/Fcgr3/Fcer1g/Cd84/Ifngr1/Itgb2/Dock2/Havcr2/Adora2b/Pld2/Lgals9/Ifi35/Tmem106a/Grn/Cd300a/Itgb8/Gpr137b/Syk/Wnt5a/Prkcd/Clu/Rac2/Snx4/Myo1f/Aif1/Adgrf5/Trem2/Ticam1/Csf1r/Jak2/Ptgds/Nmi/Spi1/Nr1h3/Traf6/Thbs1/Snap23/Lbp/Tlr2/Cd1d1/Notch2/Bcl10/Lyn/Tlr4/Ptafr/C1qa/Pla2g5/Tnip2/Rbpj/Tlr1/Tlr6/Rhoh/Anxa3/Lat2/Hmgb1/Gimap5/Npy/Vamp8/Dysf/Gata2/Ptpn6/Ltbr/C5ar1/Nectin2/Tgfb1/Tyrobp/Flt3l/Cd37/Fes/Ctsc/Il4ra/Stx4a/Pycard/Itgam/Adam9/Slc7a2/Tlr3/Il15/Plcg2/Ldlr/Il18/Rora/Adam10/Plscr1/Plscr2/Tgfbr2/Myd88/Cx3cr1/Ccr2/Btk | 92 |

| | | | | | | | | |
|---|---|---|---|---|---|---|---|---|
| BP | GO:1903706 | regulation of hemopoiesis | 142/3915 | 419/17474 | 3,4699E-08 | 1,7354E-06 | 1,2923E-06 | Il1rl2/Stat1/Casp8/Inpp5d/Rab7b/Sox13/Ptprc/Rc3h1/Itpkb/Dusp10/Vsir/Fbxo7/Kitl/Smarcc2/Ikzf1/Meis1/Irf1/Nlrp3/Lgals9/Ccl9/Smarce1/Stat5b/Stat3/Itgb3/Asxl2/Id2/Nfkbia/Zfp36l1/Dicer1/Gpr137b/Gli3/Ripk1/Nedd9/Ninj1/Syk/Ctla2a/Mef2c/Pik3r1/Duxbl1/Prxl2a/Bmp4/Pnp/Mmp14/Rb1/Il7r/Myc/Nfam1/Arid2/Senp1/Rarg/Nckap1l/Prkdc/Hcls1/Gabpa/Runx1/H2-Aa/Trem2/Zfp36l2/Socs5/Zeb1/Lox/Cd74/Csf1r/Tjp2/Smarca2/Fas/Il15ra/Nrarp/Spi1/Mdk/Traf6/Cd44/Lmo2/B2m/Rassf2/Jag1/Car2/Actl6a/Cd1d1/Rorc/Notch2/Cd101/Rbm15/Csf1/Lef1/Lyn/Ripk2/Tal1/Csf3r/C1qc/Ctnnbip1/Prdm16/Prkcz/Isg15/Cdk6/Fgl2/Fgfr3/Rhoh/Hspb1/Hmgb1/Ptn/Gimap5/Tmem176b/Tmem176a/Mturn/Loxl3/Gata2/Atg7/Ptpn6/Lag3/Clec2d/Pglyrp1/Ceacam1/Tgfb1/Axl/Zfp36/Tyrobp/Flt3l/Klhl25/Cib1/Fes/Il4ra/Itgam/Gas6/Kat6a/Tcim/Inpp4b/Il15/Cbfb/Abcb10/Ets1/Ubash3b/Il18/Rbp1/Rhoa/Tgfbr2/Ctnnb1/Ccr2/Sash3/Atp11c/Hmgb3/Il2rg | 142 |
| BP | GO:0001570 | vasculogenesis | 46/3915 | 97/17474 | 4,6477E-08 | 2,2429E-06 | 1,6703E-06 | Sox17/Enpp1/Hey2/Sgpl1/Tmem100/Ramp2/Gjc1/Zfp36l1/Itgb8/Ntrk2/Angpt1/Has2/Hdac7/Emp2/Heg1/Xdh/Fgf1/Pdgfrb/Notch1/Egfl7/Eng/Spred1/Rin2/Tiparp/Rap1a/Tek/Tie1/Paxip1/Kdr/Tgfbr3/Myo18b/Asb4/Cav1/Smo/Ceacam1/Tgfb1/Rras/Tead2/Fzd4/Fgfr2/Yap1/Myo1e/Setd2/Tgfbr2/Ctnnb1/Amot | 46 |
| BP | GO:0001894 | tissue homeostasis | 97/3915 | 262/17474 | 4,818E-08 | 2,285E-06 | 1,7016E-06 | Col9a1/Col3a1/Inpp5d/Tnfrsf11a/Bcl2/Ccn2/Enpp1/Pbld2/Pcdh15/Adora2a/Bsg/Tns3/Egfr/Slc22a5/Slc22a21/Atp1b2/Acaca/Mks1/Itgb3/Sox9/Gpr137b/Foxc1/Syk/F2r/Ocln/Vstm4/Rb1/Ank/Angpt1/Col14a1/Rac2/Atf1/Cldn5/Tfrc/Cxadr/Col11a2/Vegfa/Pdgfrb/Csf1r/Adrb2/Tcirg1/Aldh1a1/Tjp2/Notch1/Ptgs1/Itgav/Traf6/Trp53inp2/Car2/Wwtr1/Clrn1/Ctsk/Ctss/Dram2/Csf1/S1pr1/Abca4/Slc39a8/Tlr4/Nfib/Mfsd2a/Alpl/Nos3/Cytl1/Cd38/Prom1/Spp1/Idua/Tmem119/Ptpn11/P2rx7/Pdk4/Tspan12/Smo/Gata2/Vhl/Lrp6/Ceacam1/Lsr/Flt3l/Abcc6/Tjp1/Lrrk1/Tpp1/Clcn3/Nod2/Bbs2/Itgb1/Yap1/Ubash3b/Nphp3/Pth1r/Ctnnb1/Ccr2/Ndp/Xiap/Sash3 | 97 |
| BP | GO:0060249 | anatomical structure homeostasis | 97/3915 | 262/17474 | 4,818E-08 | 2,285E-06 | 1,7016E-06 | Col9a1/Col3a1/Inpp5d/Tnfrsf11a/Bcl2/Ccn2/Enpp1/Pbld2/Pcdh15/Adora2a/Bsg/Tns3/Egfr/Slc22a5/Slc22a21/Atp1b2/Acaca/Mks1/Itgb3/Sox9/Gpr137b/Foxc1/Syk/F2r/Ocln/Vstm4/Rb1/Ank/Angpt1/Col14a1/Rac2/Atf1/Cldn5/Tfrc/Cxadr/Col11a2/Vegfa/Pdgfrb/Csf1r/Adrb2/Tcirg1/Aldh1a1/Tjp2/Notch1/Ptgs1/Itgav/Traf6/Trp53inp2/Car2/Wwtr1/Clrn1/Ctsk/Ctss/Dram2/Csf1/S1pr1/Abca4/Slc39a8/Tlr4/Nfib/Mfsd2a/Alpl/Nos3/Cytl1/Cd38/Prom1/Spp1/Idua/Tmem119/Ptpn11/P2rx7/Pdk4/Tspan12/Smo/Gata2/Vhl/Lrp6/Ceacam1/Lsr/Flt3l/Abcc6/Tjp1/Lrrk1/Tpp1/Clcn3/Nod2/Bbs2/Itgb1/Yap1/Ubash3b/Nphp3/Pth1r/Ctnnb1/Ccr2/Ndp/Xiap/Sash3 | 97 |
| BP | GO:0061351 | neural precursor cell proliferation | 80/3915 | 207/17474 | 8,7321E-08 | 3,895E-06 | 2,9006E-06 | Gli2/Btg2/Gpr37l1/Aspm/Prox1/Nr2e1/Fabp7/Lims1/Appl2/Igf1/Frs2/Acsl6/Kctd11/Lhx1/Grn/Id2/Numb/Gli3/Id4/Wnt5a/Fzd6/Plxnb2/Nde1/Tead3/Vegfa/Lims2/Ctnna1/Emx2/Rsu1/Notch1/Lhx2/Mdk/Pax6/Ctsz/Sox2/Fgf2/Foxo1/Nes/Lef1/Gng5/Lyn/Pou3f2/Akna/Nfib/Nfia/Dock7/Tacc3/Fgfr3/Ptprz1/Smo/Ptn/Gata2/Lrp6/Sox5/C5ar1/Tgfb1/Sirt2/Spint2/Atf5/Ilk/Fgfr2/Tacc1/Hook3/Pcm1/Vegfc/Hhip/Sall1/Adgrg1/Itgb1/Rora/Ephb1/Ryk/Gnai2/Rhoa/Cx3cr1/Ctnnb1/Ccr5/Fgf13/Flna/Slc16a2 | 80 |
| BP | GO:1904019 | epithelial cell apoptotic process | 60/3915 | 142/17474 | 9,3782E-08 | 4,1355E-06 | 3,0797E-06 | Cflar/Casp8/Bok/Bcl2/Lims1/Col18a1/Igf1/Spop/Ramp2/Pik3cg/Sav1/Zfp36l1/Sfrp4/Cast/Bmp4/Rb1/Rgcc/Sema5a/Angpt1/Ppara/Ano6/Atf7/Angptl4/Ecscr/Jak2/Fas/Tcf7l2/Pla2r1/Nfe2l2/Mdk/Thbs1/Id1/Srsf6/Cd40/Hipk1/Usp53/Casp6/Lef1/Tek/Tnip2/Ppargc1a/Kdr/Pla2g1b/Gper1/Ndnf/Alms1/Gata2/Atg7/Cdkn1b/Zfp36/Pak4/Igf1r/Nupr1/Gas6/Casp3/Ednra/Cdh5/Yap1/Icam1/Tgfbr2 | 60 |
| BP | GO:2000377 | regulation of reactive oxygen species metabolic process | 65/3915 | 158/17474 | 9,4216E-08 | 4,1355E-06 | 3,0797E-06 | Cflar/Bcl2/Ncf2/Slc30a10/Ccn6/Fyn/Itgb2/Egfr/Stat3/Brca1/Dcxr/Hbp1/Fbln5/Nqo2/Ripk1/Syk/Dhfr/Nnt/Plau/Prkcd/Ripk3/Tspo/Ppara/Abcd2/Cdkn1a/Ier3/Xdh/Cyp1b1/Pdgfrb/Nfe2l2/F2/Thbs1/Foxo1/Shc1/Gnai3/Map3k7/Tlr4/Slc25a33/Tlr6/Aldh2/Pon3/Hk2/Alox5/Tgfb1/Sirt2/Tyrobp/Prcp/Ndufc2/Trim30a/Itgam/Sirt3/Insr/Clcn3/Mt3/Plcg2/Cyba/Agt/Birc3/Bco2/Il18/Gnai2/Rhoa/Tgfbr2/G6pdx/Ogt | 65 |
| BP | GO:1904035 | regulation of epithelial cell apoptotic process | 49/3915 | 108/17474 | 9,9114E-08 | 4,2822E-06 | 3,1889E-06 | Cflar/Bok/Bcl2/Lims1/Col18a1/Igf1/Spop/Ramp2/Sav1/Zfp36l1/Sfrp4/Cast/Bmp4/Rb1/Rgcc/Sema5a/Angpt1/Ppara/Ano6/Angptl4/Ecscr/Jak2/Tcf7l2/Pla2r1/Nfe2l2/Mdk/Thbs1/Id1/Srsf6/Cd40/Tek/Tnip2/Ppargc1a/Kdr/Pla2g1b/Gper1/Ndnf/Alms1/Gata2/Atg7/Cdkn1b/Zfp36/Pak4/Igf1r/Nupr1/Gas6/Cdh5/Yap1/Icam1 | 49 |
| BP | GO:0014910 | regulation of smooth muscle cell migration | 47/3915 | 102/17474 | 1,0064E-07 | 4,3029E-06 | 3,2043E-06 | Igfbp5/Bcl2/Ddr2/Igf1/Vtn/Stat5b/Itgb3/Ace/Gna13/Mef2c/Il6st/Plau/Bmpr1a/Dock5/Has2/Myc/Rapgef3/Adamts1/Aif1/Cyp1b1/Pdgfrb/Coro1b/Egfl7/Rapgef4/Nfe2l2/Mdk/Sema6d/S100a11/F3/Camk2d/Tlr4/Dock7/Ppargc1a/Cav1/Smo/Abhd2/Iqgap1/P2ry6/P2ry2/Ilk/Agt/Pdgfd/S1pr2/Il18/Rhoa/Foxo4/Atp7a | 47 |
| BP | GO:1902903 | regulation of supramolecular fiber organization | 131/3915 | 386/17474 | 1,0654E-07 | 4,5316E-06 | 3,3747E-06 | Map2/Vil1/Lmod1/Myoc/Pfdn2/F11r/Prox1/Lats1/Ccn2/Arhgap18/Gja1/Specc1l/Washc4/Aebp1/Plek/Ccdc88a/Sh3pxd2b/Pdlim4/Flii/Arhgef15/Serpinf2/Myo1c/Pecam1/Cfl2/Ttc8/Carmil1/Edn1/Mef2c/Pik3r1/Map3k1/Atxn7/Prkcd/Fermt2/Gmfb/Clu/Sorbs3/Rb1/Rgcc/Rictor/Sema5a/Mtss1/Baiap2l2/Twf1/Rapgef3/Lima1/Nckap1l/Ppm1f/Dlg1/Pak2/Hcls1/Phldb2/Snx9/Kank3/Trem2/Daam2/Eml4/Svil/Smad4/Coro1b/Efemp2/Kank1/Add3/Prkcq/Slc39a12/Tsc1/Ckap5/Usp8/Id1/Tpx2/Mapre1/Sdc4/Pik3ca/Tlr2/Arfip1/Kirrel/Rhoc/Capza1/S1pr1/Synpo2/Tgfbr1/Cdk5rap2/Wasf2/Asap3/Hspg2/Akap9/Limch1/Hip1r/Eln/Arpc1b/Wasf3/Cav1/Arhgef5/Capg/Dysf/Alms1/Cdkn1b/Eps8/Vasp/Apoe/Plekhg2/Nphs1/Rhpn2/Cyfip1/Tjp1/Cib1/Fes/Fchsd2/Arap1/Swap70/Pycard/Rnh1/Arhgef10/Dlc1/Cdh5/Fhod1/Icam1/Ldlr/Kank2/Esam/Rdx/Cgnl1/Tmod3/Rhoa/Clasp2/Was/Mid1ip1/Fgf13/Mecp2/Flna/Arhgap6/Mid1 | 131 |
| BP | GO:0007596 | blood coagulation | 71/3915 | 178/17474 | 1,0749E-07 | 4,5486E-06 | 3,3874E-06 | Serpine2/Cfh/Pla2g4a/Fcer1g/F11r/Plek/Alox12/Gp1ba/Serpinf2/Vtn/Itga2b/Itgb3/Gna13/C1qtnf1/F13a1/Syk/F2r/Plau/Prkcd/Fzd6/Fbln1/Ano6/Abat/Serpind1/Pros1/Enpp4/Adrb2/Fermt3/Jak2/Papss2/Entpd1/Prkcq/Entpd2/Nfe2l2/Tfpi/Serping1/F2/Thbs1/Mertk/Thbd/Anxa5/Slc7a11/P2ry12/P2ry1/Pear1/Il6ra/Stxbp3/F3/Lyn/Tlr4/Pdpn/Tec/Pdgfra/Cav1/Pdia4/Gp9/Ptpn6/Cd9/Vwf/Apoe/Plaur/Ceacam1/Axl/Hps5/Vkorc1/Gas6/Plat/St3gal4/Ubash3b/Anxa2/Flna | 71 |
| BP | GO:0003018 | vascular process in circulatory system | 87/3915 | 232/17474 | 1,2033E-07 | 5,0404E-06 | 3,7536E-06 | Atp1a2/Akap12/Gja1/Adora2a/Egfr/Slc22a5/Adora2b/Serpinf2/Scpep1/Ramp2/Ace/Foxc1/Bmp6/Edn1/F2r/Ocln/Itga1/Vstm4/Fermt2/Dock5/Ednrb/Angpt1/Cldn5/Casr/Cbs/Vegfa/Kat2b/Ptprm/Rock1/Nr3c1/Adrb2/Tjp2/Acta2/Fgfbp3/Add3/Ptgs1/Edn3/P2ry1/Gucy1a1/Shc1/Gja5/Gclm/Pde5a/Ddah1/Slc2a1/Mfsd2a/Ctnnbip1/Abcb1a/Abcb1b/Nos3/Slc5a6/Cd38/Kdr/Tgfbr3/P2rx4/Plod3/Gper1/Cav1/Tbxas1/Abcg2/Mgll/Hrh1/Alox5/Pde3a/Slc1a5/Gpr4/Apoe/Ceacam1/Tgfb1/Tjp1/P2ry2/Slc27a1/Klf2/Ednra/Mmp2/Bbs2/Cdh5/Agt/Arhgap42/Icam1/Il18/Rhoa/Apln/G6pdx/Slc16a2/Cysltr1/Amot | 87 |
| BP | GO:0140895 | cell surface toll-like receptor signaling pathway | 30/3915 | 54/17474 | 1,254E-07 | 5,2264E-06 | 3,8921E-06 | Ly96/Wdfy1/Rab7b/Tlr5/Appl2/Ifi35/Nfkbia/Ninj1/Trem2/Cd14/Ticam2/Pik3ap1/Nmi/Nr1h3/Lbp/Tlr2/Ptpn22/Lyn/Ripk2/Tlr4/Tnip2/Tlr1/Tlr6/Oas1a/Hmgb1/Tril/Nod2/Cyba/Tirap/Irak1 | 30 |
| BP | GO:0019216 | regulation of lipid metabolic process | 126/3915 | 369/17474 | 1,2617E-07 | 5,2318E-06 | 3,8962E-06 | Ormdl1/Idh1/Acadl/Erbb4/Acsl3/Irs1/Sctr/Dbi/Adipor1/Pla2g4a/Fmo1/Fmo2/Capn2/Prox1/Ppp2r5a/Psap/Appl2/Igf1/Ormdl2/Pik3ip1/Adora2b/Ncor1/Acadvl/Stat5b/Brca1/Sox9/Rnf213/Lpin1/Id2/Pik3cg/Gpld1/Bmp6/Golm1/Lpcat1/Pde8b/Nr1d2/Prkcd/Ephx2/Dab2/Tspo/Ppara/Abcd2/Apod/Zbtb20/Lmf1/Angptl4/Adgrf5/Trem2/Stard4/Fgf1/Nr3c1/Cd74/Pdgfrb/Cpt1a/Sorbs1/Gpam/Tcf7l2/Pip4k2a/Nr1h3/F2/Pex2/Fabp5/Dnajc19/Fgf2/Rorc/Cers2/Fmo5/Vav3/Nfkb1/Sh3glb1/Ccn1/Lyn/Tek/Scp2/Zfp69/Zmpste24/Mfsd2a/Insig1/Fgfr3/Ppargc1a/Pdgfra/Rest/Igfbp7/Acacb/Hcar1/Mlxipl/Gper1/Flt1/Pdk4/Cav1/Fmc1/Rarres2/Gimap5/Hrh1/Atg7/Ankrd26/Apobec1/Lpcat3/Crebl2/Apoc1/Apoe/Ceacam1/Tgfb1/Akt2/Lsr/Igf1r/Klhl25/Thrsp/Pde3b/Sirt3/Pnpla2/Igf2/Cd81/Dhcr7/Erlin2/Asah1/Hpgd/Slc27a1/Nod2/Plcg2/Agt/Ldlr/Rora/Bmp5/Elovl5/Mid1ip1 | 126 |

| | | | | | | | | | |
|---|---|---|---|---|---|---|---|---|---|
| BP | GO:0060537 | muscle tissue development | 151/3915 | 461/17474 | 1,4948E-07 | 5,9018E-06 | 4,3951E-06 | Eya1/Tfap2b/Col19a1/Col3a1/Cflar/Casp8/Erbb4/Igfbp5/Bcl2/Btg2/Csrp1/Tnni1/Kcnk2/Cenpf/Prox1/Atf3/Hey2/Pln/Gja1/S100b/Igf1/Csrp2/Frs2/Erbb3/Meis1/Sgcd/Pmp22/Hlf/Gjc1/Sox9/Id2/Cfl2/Sav1/Dicer1/Foxc1/Dsp/Edn1/Mef2c/Nln/Isl1/Nr1d2/Wnt5a/Bmpr1a/Bmp4/Myh6/Rb1/Angpt1/Col14a1/Myc/Ddx17/Ppara/Arid2/Hdac7/Myh11/Tbx1/Dlg1/Heg1/Mylk/Boc/Cxadr/Vegfa/Med20/Trip10/Foxn2/Svil/Fgf1/Nr3c1/Lox/Megf10/Pdgfrb/Adrb2/Smad4/Efemp2/Pgm5/Tll2/Tcf7l2/Nrap/Nebl/Notch1/Tsc1/Eng/Ttn/Spg11/Gpcpd1/Ccm2l/Bmp7/Fgf2/Mbnl1/Tiparp/Gja5/Nras/S1pr1/Camk2d/Lef1/Pdlim5/Acadm/Chd7/Myorg/Tgfbr1/Zmpste24/Heyl/Hspg2/Trp73/Sema3c/Ppargc1a/Rbpj/Sgcb/Pdgfra/Tgfbr3/Myo18b/Tbx3/Eln/Foxp2/Cav2/Cav1/Smo/Vamp5/Dysf/Adamts9/Atg7/Vgll4/Ccnd2/Lrp6/Tgfb1/Sirt2/Nphs1/Tshz3/Vrk3/Sox6/Nupr1/Fgfr2/Igf2/Nrg1/Ednra/Smad1/Nfatc3/Agt/Itgb1/Yap1/Aldh1a2/Bmp5/Tbx18/Ephb1/Rhoa/Tgfbr2/Ctnnb1/Zic3/Mtm1/Srpk3/G6pdx/Dmd | 151 |
| BP | GO:0001763 | morphogenesis of a branching structure | 91/3915 | 247/17474 | 1,6032E-07 | 6,2996E-06 | 4,6913E-06 | Eya1/Bcl2/Gli2/Pbx1/Vangl2/Prox1/Col13a1/Igf1/Ntn4/Tmtc3/Frs2/Timeless/Hs3st3b1/Hs3st3a1/Lhx1/Mks1/Etv4/Gna13/Sox9/Btbd7/Dicer1/Gli3/Edn1/Ptch1/Adamts16/Wnt5a/Bmp4/Mmp14/Sema5a/Angpt1/Myc/Tbx1/Dlg1/Casr/Notch4/Vegfa/Lama1/Fgf1/Smad4/Nfatc1/Lrp5/Nrarp/Notch1/Eng/Mdk/Cd44/Lgr4/Tgm2/Bmp7/Ctsz/Lama5/Sox2/Fgf2/Rbm15/Csf1/Lef1/Npnt/Tnc/Tek/Tie1/Clic4/Ctnnbip1/Hgf/Sema3c/Pdgfra/Kdr/Tbx3/Flt1/Smo/Cxcl12/Lrp6/Mgp/Tgfb1/Spint2/Ilk/Fgfr2/Cpe/Ednra/Hhip/Sall1/Nfatc3/Cdh1/Agt/Yap1/Ctsh/Setd2/Tgfbr2/Ctnnb1/Gpc3/Fgf13/Mecp2 | 91 |
| BP | GO:0061081 | positive regulation of myeloid leukocyte cytokine production involved in immune response | 24/3915 | 39/17474 | 1,6626E-07 | 6,4411E-06 | 4,7967E-06 | Fcer1g/Ddx21/Syk/Wnt5a/Ticam1/Ticam2/Cd74/Card9/Mavs/Tlr2/Ripk2/Tlr4/Laptm5/P2rx7/Nod1/Gprc5b/Pycard/Tlr3/Nod2/Plcg2/Casp4/Tirap/Myd88/Tlr7 | 24 |
| BP | GO:0048872 | homeostasis of number of cells | 124/3915 | 364/17474 | 1,834E-07 | 7,039E-06 | 5,242E-06 | Slc40a1/Stat1/Cflar/Dock10/Inpp5d/Bcl2/Cfh/Rc3h1/Fcgr2b/Fcer1g/Itpkb/Myct1/Enpp1/Kitl/Ikzf1/Hba-a1/Hba-a2/Tnfrsf13b/Ncor1/Cdk5rap3/Stat5b/Stat3/Sox9/Adam17/Id2/Zfp36l1/Siva1/Ncapg2/Sfxn1/Smad5/Mef2c/F2r/Gcnt4/Rps24/Bmp4/Ripk3/Rb1/Gpr183/Il7r/Col14a1/Senp1/Sp1/Nckap1l/Prkdc/Hcls1/Fstl1/Pknox1/Adgrf5/Vegfa/Epas1/Rps14/Cd74/Csf1r/Tcirg1/Smarca2/Jak2/Fas/Gpam/Card9/Notch1/Spi1/Cd44/Lmo2/Lgr4/B2m/Bcl2l11/Mertk/Rassf2/Mecom/Slc7a11/P2ry14/Csf1/Tet2/Emcn/Bcl10/Lyn/Ptbp3/Tal1/Prdx1/Xkr8/Isg15/Cdk6/Nos3/Tgfbr3/Ptpn11/P2rx7/Hmgb1/Smo/Bpgm/Arhgef5/Ezh2/Gimap5/Gata2/Vhl/Lpcat3/Rps19/Tgfb1/Axl/Zfp36/Cebpg/Flt3l/Ampd3/Sox6/Itgam/Ikbkb/Casp3/Klf2/Dnase2a/Adgrg1/Abcb10/Ets1/Il18/Tmod3/Cx3cr1/Ccr2/Dock11/Xiap/Sash3/G6pdx/Zfx/Heph/Btk/Tsc22d3/Alas2 | 124 |
| BP | GO:0050866 | negative regulation of cell activation | 83/3915 | 221/17474 | 2,1584E-07 | 8,2079E-06 | 6,1124E-06 | Fn1/Serpine2/Inpp5d/Rc3h1/Fcgr2b/Cd84/Vsir/Adora2a/Fbxo7/Havcr2/Irf1/Tnfrsf13b/Alox12/Lgals9/Grn/Cd300a/C1qtnf1/Id2/Gli3/Hfe/Dusp22/Prkcd/Bmp4/Lgals3/Nckap1l/Abat/Dlg1/Cd86/Cd80/Runx1/H2-Aa/Adgrf5/Trem2/Twsg1/Socs5/Cd74/Adrb2/Cd274/Fas/Nrarp/Spi1/Nr1h3/Mdk/Cd44/Mertk/Sdc4/Pag1/Tnfaip8l2/Ptpn22/Pde5a/Bank1/Lyn/Laptm5/Pla2g5/Tnfrsf14/Fgl2/Pdgfra/Hspb1/Gper1/Hmgb1/Gimap5/Gpnmb/Loxl3/Ptpn6/Lag3/Cd9/Pglyrp1/Apoe/Ceacam1/Tgfb1/Axl/Tyrobp/Cd37/Lrrc32/Il4ra/Casp3/Cbfb/Ldlr/Ubash3b/Parp3/Ccr2/Hmgb3/Btk | 83 |
| BP | GO:0061448 | connective tissue development | 102/3915 | 287/17474 | 2,2925E-07 | 8,6442E-06 | 6,4373E-06 | Col9a1/Arid5a/Col3a1/Cflar/Klf7/Gli2/Cfh/Prrx1/Ccn2/Enpp1/Igf1/Efemp1/Sh3pxd2b/Rflnb/Col1a1/Itgb3/Axin2/Sox9/Pum2/Mboat2/Id2/Sptlc2/Trip11/Dicer1/Itgb8/Gli3/Gpld1/Bmp6/Edn1/Id4/Ogn/Smad5/Mef2c/Wnt5a/Mustn1/Bmpr1a/Bmp4/Rb1/Adamts12/Trps1/Rarg/Cbs/Col11a2/Twsg1/Zeb1/Lox/Pdgfrb/Lrp5/Rela/Plaat3/Notch1/Spi1/Mdk/Thbs1/Ddrgk1/Bmp7/Pik3ca/Fgf2/Rxfp1/Pbxip1/Il6ra/Rorc/Ctsk/Csf1/Bmpr1b/Ccn1/Tgfbr1/Col27a1/Nfib/Nfia/Zmpste24/Hspg2/Hes5/Paxip1/Fgfr3/Cytl1/Ppargc1a/Idua/Ptpn11/Creb3l2/C3ar1/Lrp6/Mgp/Sox5/Gpr4/Tgfb1/Chsy1/Serpinh1/Sox6/Mapk3/Csgalnact1/Smad1/Bbs2/Maf/Pdgfd/Bmp5/Pth1r/Tgfbr2/Ctnnb1/Bgn/Amer1/Atp7a | 102 |
| BP | GO:0007599 | hemostasis | 71/3915 | 181/17474 | 2,3235E-07 | 8,6757E-06 | 6,4608E-06 | Serpine2/Cfh/Pla2g4a/Fcer1g/F11r/Plek/Alox12/Gp1ba/Serpinf2/Vtn/Itga2b/Itgb3/Gna13/C1qtnf1/F13a1/Syk/F2r/Plau/Prkcd/Fzd6/Fbln1/Ano6/Abat/Serpind1/Pros1/Enpp4/Adrb2/Fermt3/Jak2/Papss2/Entpd1/Prkcq/Entpd2/Nfe2l2/Tfpi/Serping1/F2/Thbs1/Mertk/Thbd/Anxa5/Slc7a11/P2ry12/P2ry1/Pear1/Il6ra/Stxbp3/F3/Lyn/Tlr4/Pdpn/Tec/Pdgfra/Cav1/Pdia4/Gp9/Ptpn6/Cd9/Vwf/Apoe/Plaur/Ceacam1/Axl/Hps5/Vkorc1/Gas6/Plat/St3gal4/Ubash3b/Anxa2/Flna | 71 |
| BP | GO:0050817 | coagulation | 71/3915 | 181/17474 | 2,3235E-07 | 8,6757E-06 | 6,4608E-06 | Serpine2/Cfh/Pla2g4a/Fcer1g/F11r/Plek/Alox12/Gp1ba/Serpinf2/Vtn/Itga2b/Itgb3/Gna13/C1qtnf1/F13a1/Syk/F2r/Plau/Prkcd/Fzd6/Fbln1/Ano6/Abat/Serpind1/Pros1/Enpp4/Adrb2/Fermt3/Jak2/Papss2/Entpd1/Prkcq/Entpd2/Nfe2l2/Tfpi/Serping1/F2/Thbs1/Mertk/Thbd/Anxa5/Slc7a11/P2ry12/P2ry1/Pear1/Il6ra/Stxbp3/F3/Lyn/Tlr4/Pdpn/Tec/Pdgfra/Cav1/Pdia4/Gp9/Ptpn6/Cd9/Vwf/Apoe/Plaur/Ceacam1/Axl/Hps5/Vkorc1/Gas6/Plat/St3gal4/Ubash3b/Anxa2/Flna | 71 |
| BP | GO:0050920 | regulation of chemotaxis | 84/3915 | 225/17474 | 2,4636E-07 | 9,0755E-06 | 6,7586E-06 | Fn1/Wnt3/Adam17/Prkd1/Rin3/Lgmn/Tubb2b/Nedd9/Edn1/Ninj1/Cxcl14/Thbs4/Wnt5a/Sema3g/Gpr183/Sema5a/Rac2/Ano6/Nckap1l/Ppm1f/St6gal1/Casr/Aif1/Pla2g7/Vegfa/Trem2/Fgf1/Sema6a/Cd74/Pdgfrb/Csf1r/Coro1b/Vegfb/Notch1/Dpp4/Spi1/Mdk/Thbs1/Sema6d/Lbp/Edn3/Fgf2/P2ry12/Csf1/S1pr1/Lyn/Padi2/Sema3d/Sema3c/Pdgfra/Kdr/Cmklr1/Slc8b1/P2rx4/Hspb1/Hmgb1/Ptn/Rarres2/Dysf/Cxcl12/Wnk1/C3ar1/C5ar2/C5ar1/Prkd2/Tgfb1/Akt2/Swap70/Mapk3/Stx4a/Gas6/Angpt2/Mtus1/Vegfc/Ednra/Nod2/Pdgfd/Tirap/Adam10/Ryk/Oxsr1/Cx3cr1/Ccr2/Mospd2 | 84 |
| BP | GO:0060411 | cardiac septum morphogenesis | 38/3915 | 78/17474 | 2,8148E-07 | 1,0233E-05 | 7,6202E-06 | Nrp2/Vangl2/Prox1/Hey2/Mks1/Fzd2/Id2/Sav1/Isl1/Wnt5a/Bmpr1a/Bmp4/Tbx1/Smad4/Nfatc1/Notch1/Eng/Jag1/Bmp7/Gja5/Notch2/Rbm15/Ccn1/Chd7/Tgfbr1/Heyl/Dhrs3/Sema3c/Nos3/Rbpj/Tgfbr3/Tbx3/Smo/Lrp6/Parva/Fgfr2/Bmp5/Tgfbr2 | 38 |
| BP | GO:0002532 | production of molecular mediator involved in inflammatory response | 44/3915 | 96/17474 | 3,0748E-07 | 1,0984E-05 | 8,18E-06 | Fcer1g/Dusp10/Appl2/Nos2/Stat3/Grn/Cd300a/Adam17/Pld4/Syk/Ephx2/Ppara/Abcd2/Nlrc3/Apod/Snx4/Trem2/Ticam1/Ticam2/Card9/F2/Snap23/Lbp/Pbxip1/Gbp5/Lyn/Tlr4/Tlr6/Ncf1/Alox5ap/Ezh2/Vamp8/Il17rc/Alox5/Il17ra/Clec7a/Rps19/Il4ra/Pycard/Slc7a2/Adcy7/Nod2/Myd88/Btk | 44 |
| BP | GO:0060760 | positive regulation of response to cytokine stimulus | 33/3915 | 64/17474 | 3,1696E-07 | 1,1225E-05 | 8,3596E-06 | Il1r1/Ikbke/Irgm1/Igtp/Irgm2/Adam17/Ripk1/Edn1/Wnt5a/Parp14/Parp9/Crebrf/Trem2/Sting1/Ticam2/Cd74/Ifih1/Mavs/Cpne1/Tlr2/Csf1/Ripk2/Tlr4/Laptm5/Rbm47/Trim56/Axl/Lsm14a/Irf3/Irf7/Gas6/Nlrc5/Casp4 | 33 |
| BP | GO:1901652 | response to peptide | 140/3915 | 427/17474 | 3,9539E-07 | 1,3595E-05 | 1,0124E-05 | Col3a1/Stat1/Irs1/Adipor1/Fcgr2b/Eprs/Slc30a10/Enpp1/Fyn/Gja1/Appl2/Igf1/Stat6/Timeless/Cdk2/Grb10/Irf1/Adora2b/Pld2/Serpinf1/Myo1c/Stxbp4/Stat5b/Stat3/Lpin1/Snx6/Nfkbia/Zfp36l1/Lgmn/Serpina1c/Gpld1/Edn1/Gkap1/Zbed3/Pik3r1/Erbin/Prkcd/Rb1/Tbc1d4/Ednrb/Ghr/Ppara/Tns2/Sp1/Prkdc/Srsf3/Trem2/Kat2b/Rab31/Rock1/Cdo1/Camk2a/Adrb2/Rela/Kank1/Jak2/Ide/Sorbs1/Chuk/Gpam/Prkcq/Vim/Pip4k2a/Card9/Notch1/Tsc1/Nfe2l2/Zfp106/Gatm/Jag1/Snx5/Lpin3/Car2/Pld1/Pik3ca/Anxa5/Foxo1/Shc1/Rab13/Ptpn22/Ahcyl1/Stxbp3/Vcam1/Bcar3/Nfkb1/Ripk2/Ggh/Chmp5/Tlr4/Dennd4c/Leprot/Pik3r3/Slc2a1/Agtrap/Slc25a33/Prkcz/Insig1/Hadha/Khk/Tlr6/Tgfbr3/Ptpn11/Gper1/Pdk4/Cav2/Cav1/Irf5/Rarres2/Nod1/Klf15/Ankrd26/Ceacam1/Akt2/Cyfip1/Igf1r/Inppl1/Pde3b/Mapk3/Ctsd/Igf2/Ano1/Insr/Ikbkb/Eif4ebp1/Slc27a1/Ednra/Nod2/Mmp2/Agt/Sesn3/Icam1/Il18/Tcf12/Gnai2/Slc26a6/Plcd1/Ctnnb1/Ldoc1/Foxo4/Ogt | 140 |

| | | | | | | | | | |
|---|---|---|---|---|---|---|---|---|---|
| BP | GO:0001818 | negative regulation of cytokine production | 100/3915 | 283/17474 | 4,0946E-07 | 1,3848E-05 | 1,0312E-05 | Fn1/Slc11a1/Inpp5d/Ptprc/Prg4/Fcgr2b/Cd84/Vsir/Srgn/Appl2/Igf1/Irak3/Ddit3/Rel/Havcr2/Nlrp3/Lgals9/Trib2/Dicer1/Hfe/Erbin/Ndrg2/Rgcc/Ubr5/Angpt1/Tspo/Fbln1/Ppara/Abcd2/Hdac7/Nckap1l/Nlrc3/Apod/Cd200r1/Ezr/Akap8/Cd2ap/Trem2/Twsg1/Ppm1b/Socs5/Aqp4/Rnf125/Ticam2/Atg12/Cd274/Nmi/F2/Cd59a/Lgr4/Thbs1/Mertk/Lbp/Slc2a10/Tlr2/Otud7b/Ptpn22/Lef1/Nfkb1/Bank1/Gbp7/Gbp3/Klf4/Tlr4/Laptm5/Hgf/Tlr6/Gbp4/Cmklr1/Oas1a/Flt1/Hmgb1/Ezh2/Gimap5/Gpnmb/Anxa4/Ptpn6/Lag3/Tnfrsf1a/C5ar2/Pglyrp1/Ceacam1/Tgfb1/Axl/Zfp36/Tyrobp/Homer2/Lrrc32/Arrb1/Trim30a/Pycard/Sigirr/Gas6/Bst2/Klf2/Adcy7/Nod2/Cx3cr1/Elf4/Btk | 100 |
| BP | GO:0071396 | cellular response to lipid | 155/3915 | 484/17474 | 4,9065E-07 | 1,626E-05 | 1,2109E-05 | Ly96/Arid5a/Stat1/Irs1/Cfh/Brinp3/Cd84/Atp1a2/Smyd3/Lats1/Sash1/Igf1/Egfr/Havcr2/Irgm1/Igtp/Irgm2/Nlrp3/Ncor1/Cd68/Cxcl16/Nos2/Acaca/Brca1/Sox9/Nfkbia/Zfp36l1/Gpld1/Ly86/Bmp6/Syk/Tifab/Ntrk2/Ptch1/Mef2c/Cd180/Isl1/Fam107a/Bmp4/Dab2/Mtdh/Ubr5/Fbxo32/Ddx17/Mlc1/Tfap4/Litaf/Ube2l3/Tbx1/Cd86/Cd80/Crebrf/Akap8/Trem2/Ticam1/Lbh/Zfp36l2/Npc1/Cd14/Nr3c1/Ticam2/Rela/Rorb/Jak2/Cd274/Lcor/Vim/Mrc1/Spi1/Nr1h3/Traf6/B2m/Ddrgk1/Lbp/Bmp7/Cyp7b1/Rorc/Ptpn22/Nfkb1/Gbp2/Bcl10/Ptgfr/Lyn/Ripk2/Chmp5/Abca1/Tlr4/Heyl/Hdac1/Ptafr/Id3/Padi2/Srarp/Tnfrsf1b/Nos3/Insig1/Hadhb/Yes1/Tnip2/Rest/Spp1/Gbp6/Gper1/Hmgb1/Pdk4/Smo/Kdm3a/Nr2c2/Irak2/Ret/Phc1/Lrp6/Mgst1/Tgfb1/Axl/Zfp36/Irf3/Tead2/Abhd2/Fes/P2ry6/Trim68/Trim5/Trim12a/Trim12c/Trim30a/Tead1/Mapk3/Pycard/Adam9/Eif4ebp1/Nod2/Mmp2/Plcg2/Irf8/Yap1/Ldlr/Kank2/Il18/Trip4/Rora/Aldh1a2/Nedd4/Plscr1/Plscr2/Plscr4/Rhoa/Myd88/Cx3cr1/Kdm6a/Ldoc1/Irak1/Msn/Ogt/Hdac8 | 155 |
| BP | GO:1903034 | regulation of response to wounding | 67/3915 | 171/17474 | 5,2765E-07 | 1,7417E-05 | 1,297E-05 | Vil1/Serpine2/Tnr/Ddr2/Gja1/Kremen1/Alox12/Gp1ba/Serpinf2/Vtn/Grn/C1qtnf1/Adam17/F2r/Plau/Prkcd/Fermt2/Ano6/Abat/Mylk/Phldb2/Pros1/Enpp4/Hbegf/Adrb2/Vegfb/Kank1/Acta2/Nfe2l2/Tfpi/Serping1/F2/Mdk/Thbs1/Thbd/Srsf6/Anxa5/Fgf2/Cers2/F3/Klf4/Cldn19/Pdgfra/Hmgb1/Cav1/Ptn/Alox5/Cd9/Clec7a/Apoe/Plaur/Ceacam1/Igf1r/Xylt1/Vkorc1/Plat/Nrg1/Wfdc1/Itgb1/St3gal4/Ubash3b/Neo1/Anxa2/Cd109/Clasp2/Flna/Atp7a | 67 |
| BP | GO:0002366 | leukocyte activation involved in immune response | 99/3915 | 281/17474 | 5,4617E-07 | 1,7956E-05 | 1,3372E-05 | Slc11a1/Dock10/Ptprc/Rc3h1/Fcer1g/Cd84/Enpp1/Itgb2/Icosl/Stat6/Dock2/Havcr2/Irf1/Nlrp3/Adora2b/Tnfsf13/Pld2/Lgals9/Tmem98/Stat3/Ifi35/Grn/Cd300a/Lgals8/Syk/Cd180/Shld2/Lgals3/Lcp1/Gpr183/Rac2/Mfng/Lgals1/Nckap1l/Tfrc/Snx4/Myo1f/Trem2/Ticam1/Socs5/Cd74/Kmt5b/Apbb1ip/Ptgds/Tsc1/Nmi/Rif1/Spi1/Mdk/Snap23/Shld1/Lbp/Cd40/Il6ra/Rorc/Notch2/Lef1/Lyn/Ripk2/Exosc3/Tlr4/Ptafr/Prkcz/Fgl2/Paxip1/Anxa3/Lat2/Lfng/Hmgb1/Vamp8/Loxl3/Dysf/Gata2/Ptpn6/Cracr2a/Pglyrp1/Tgfb1/Tyrobp/Fes/Swap70/Il4ra/Itgal/Stx4a/Pycard/Itgam/Cd81/Lamp1/Plcg2/Irf8/Icam1/Il18/Rora/Parp3/Myd88/Ccr2/Dock11/Atp7a/Itm2a/Btk | 99 |
| BP | GO:0022612 | gland morphogenesis | 62/3915 | 155/17474 | 5,9816E-07 | 1,9433E-05 | 1,4472E-05 | Cflar/Igfbp5/Bcl2/Gli2/Prox1/Lims1/Igf1/Ntn4/Frs2/Stat6/Egfr/Etv4/Sox9/Btbd7/Gli3/Id4/Ptch1/Plau/Wnt5a/Bmp4/Rarg/Twsg1/Lama1/Lims2/Fgf1/Nr3c1/Lrp5/Notch1/Ctnnd1/Mdk/Cd44/Pax6/Tgm2/Bmp7/Lama5/Cyp7b1/Rxfp1/Notch2/Csf1/Tet2/Nfkb1/Tnc/Nfib/Rps6ka1/Hgf/Sema3c/Tbx3/Cav1/Smo/Ptn/Tgfa/Lrp6/Ceacam1/Tgfb1/Igf1r/Fgfr2/Mki67/Mmp2/Cdh1/Il18/Tgfbr2/Msn | 62 |
| BP | GO:0034341 | response to type II interferon | 57/3915 | 140/17474 | 8,5037E-07 | 2,6127E-05 | 1,9457E-05 | Stat1/Slc11a1/Sp100/Rab7b/Eprs/Ifngr1/Irgm1/Irf1/Slc22a5/Slc22a21/Igtp/Irgm2/Cxcl16/Myo1c/Nos2/Ccl9/Ccl6/Stxbp4/Cdc42ep4/Parp14/Parp9/H2-Aa/H2-Eb1/Kif5b/Aqp4/Cd74/Jak2/Vim/Mrc1/Nmi/Kif16b/Cd40/Tlr2/Stxbp3/Gbp5/Gbp7/Gbp3/Gbp2/Tlr4/Vamp3/Gbp9/Gbp4/Gbp6/Vamp8/Capg/Rab43/Rpl13a/Trim21/Stx4a/Ifitm2/Ifitm1/Ifitm3/Bst2/Nlrc5/Irf8/Slc26a6/Was | 57 |
| BP | GO:0002683 | negative regulation of immune system process | 155/3915 | 488/17474 | 8,5169E-07 | 2,6127E-05 | 1,9457E-05 | Col3a1/Fn1/Inpp5d/Rab7b/Ptprc/Rc3h1/Fcgr2b/Fcer1g/Cd84/Mndal/Ifi203/Dusp10/Cr1l/Vsir/Adora2a/Cnn2/Fbxo7/Igf1/Kitl/Irak3/Stat6/Havcr2/Irf1/Tnfrsf13b/Cd68/Lgals9/Grn/Sox9/Cd300a/Id2/Rin3/Gpr137b/Gli3/Hfe/Dusp22/Serpinb9/Pik3r1/Erbin/Bmp4/Lgals3/Lrch1/Elf1/Il7r/Angpt1/Myc/Gramd4/Tmbim6/Nckap1l/Nlrc3/Prkdc/Apod/Dlg1/Parp14/Cd86/Cd80/Cd200r1/Runx1/Ezr/Tap1/Tap2/H2-Ob/H2-Aa/Adgrf5/Trem2/Gpr108/Twsg1/Socs5/Riok3/Rnf125/Gpr17/Ticam2/Atg12/Cd74/Lpxn/Tjp2/Cd274/Fas/Pik3ap1/Gpam/Nrarp/Phpt1/Nmi/Dpp4/Serping1/Spi1/Nr1h3/Mdk/Cd44/Cd59a/Lgr4/Thbs1/Mertk/Samhd1/Sdc4/Pag1/Tnfaip8l2/Ptpn22/Pde5a/Bank1/Lyn/Laptm5/C1qc/Pla2g5/Padi2/Prdm16/Tnfrsf14/Isg15/Cdk6/Fgl2/Tlr6/Oas1a/Hspb1/Gper1/Hmgb1/Gimap5/Tmem176b/Tmem176a/Gpnmb/Npy/Loxl3/Gata2/Cxcl12/A2m/Ptpn6/Lag3/Tapbpl/Clec2d/C5ar2/Pglyrp1/Rps19/Ceacam1/Tgfb1/Axl/Tyrobp/Cd37/Lrrc32/Trim21/Trim30a/Il4ra/Igf2/Casp3/Bst2/Inpp4b/Nod2/Nlrc5/Cbfb/Ldlr/Ubash3b/Parp3/Cx3cr1/Ctnnb1/Ccr2/Hmgb3/Btk/Tsc22d3 | 155 |
| BP | GO:0035456 | response to interferon-beta | 31/3915 | 61/17474 | 1,084E-06 | 3,247E-05 | 2,418E-05 | Stat1/Ikbke/Aim2/Mndal/Ifi203/Traf3ip3/Irgm1/9930111J21Rik1/Ifi47/Irf1/Igtp/Irgm2/Xaf1/Ifnar2/Sting1/Iigp1/Camk2a/Ifit3/Ifit1/Mavs/Gbp7/Gbp3/Gbp2/Gbp6/Oas1a/Ifitm2/Ifitm1/Ifitm3/Bst2/Plscr1/Plscr2 | 31 |
| BP | GO:0035909 | aorta morphogenesis | 22/3915 | 37/17474 | 1,2643E-06 | 3,7262E-05 | 2,7749E-05 | Eya1/Tfap2b/Col3a1/Prox1/Hey2/Fkbp10/Bmpr1a/Acvrl1/Tbx1/Mylk/Notch4/Pdgfrb/Efemp2/Notch1/Eng/Jag1/Chd7/Rbpj/Ephb4/Adamts9/Kat6a/Tgfbr2 | 22 |
| BP | GO:0014013 | regulation of gliogenesis | 54/3915 | 132/17474 | 1,3705E-06 | 3,9497E-05 | 2,9413E-05 | Serpine2/Gpr37l1/Dusp10/Nr2e1/Zfp365/Igf1/Egfr/Tmem98/Gfap/Id2/Prkch/Dicer1/Id4/Il6st/Zcchc24/Zfp488/Bmp4/Rb1/Myc/Tspo/Trem2/Daam2/Csf1r/Mbd1/Rela/Fas/Vim/Notch1/Dlx1/F2/Mdk/Tlr2/Cers2/Lyn/Dab1/Hdac1/Tnfrsf1b/Trp73/Hes5/Fgfr3/Rnf10/Flt1/Ptprz1/Ptn/Ezh2/Tgfb1/Sirt2/Atf5/Idh2/Nrg1/Vegfc/Ldlr/Ctnnb1/Mecp2 | 54 |
| BP | GO:0034103 | regulation of tissue remodeling | 38/3915 | 82/17474 | 1,3776E-06 | 3,9497E-05 | 2,9413E-05 | Tmbim1/Inpp5d/Tnfrsf11a/Suco/Ddr2/Gja1/Egfr/Flt4/Itgb3/Klf6/Gpr137b/Syk/Thbs4/Tfrc/Abi3bp/Vegfa/Rock1/Csf1r/Itgav/Mdk/Cst3/Car2/S1pr1/Lepr/Fgfr3/Cd38/Spp1/Idua/Tmem119/P2rx7/Pdk4/Gpnmb/Lrp6/Ceacam1/Inpp4b/Agt/Ubash3b/Il18 | 38 |
| BP | GO:0050764 | regulation of phagocytosis | 49/3915 | 116/17474 | 1,3785E-06 | 3,9497E-05 | 2,9413E-05 | Slc11a1/Ptprc/Fcgr2b/Fcgr3/Fcer1g/Cnn2/Appl2/Dock2/Rack1/Cd300a/Syk/Lman2/Ano6/Nckap1l/Pros1/C4b/C2/Trem2/Pot1b/Rab31/Il15ra/Itgav/Mertk/Hck/Tgm2/Lbp/Tlr2/Fcgr1/Rap1a/Pla2g5/Hmgb1/Dysf/Gata2/Atg7/Clec7a/Tgfb1/Siglece/Pycard/Gas6/Il15/Nod2/Plcg2/Cyba/Dnm2/Plscr1/Plscr2/Cfp/Il2rg/Btk | 49 |
| BP | GO:0010721 | negative regulation of cell development | 103/3915 | 301/17474 | 1,545E-06 | 4,3812E-05 | 3,2627E-05 | Map2/Ctdsp1/Inpp5d/Btg2/Gpr37l1/Tnr/Rc3h1/Dusp10/Prox1/Nr2e1/Appl2/Fbxo7/Igf1/Irf1/Kctd11/Rflnb/Tmem98/Wnt3/Id2/Dicer1/Gpr137b/Gli3/Id4/Pik3r1/Wnt5a/Sema3g/Bmpr1a/Bmp4/Rb1/Ednrb/Sema5a/Myc/Tspo/Dip2b/Runx1/Vegfa/Trem2/Daam2/Socs5/Ctnna1/Sema6a/Cd74/Mbd1/Tjp2/Fas/Nrarp/Notch1/Lhx2/Dlx1/Lrp4/F2/Mdk/Cd44/Pax6/B2m/Sema6d/Id1/Bmp7/Cers2/Lyn/Ttpa/Dab1/C1qc/Prdm16/Hes5/Cdk6/Sema3d/Sema3c/Fgl2/Fgfr3/Rest/Rnf10/Hspb1/Hmgb1/Ptn/Tmem176b/Tmem176a/Loxl3/Gata2/Lag3/Clec2d/Pglyrp1/Ceacam1/Tgfb1/Sirt2/Atf5/Idh2/Il4ra/Hook3/Pcm1/Inpp4b/Mt3/Cbfb/Cdh1/Ldlr/Ubash3b/Ryk/Ctnnb1/Ccr5/Fgf13/Hmgb3/Arhgap4/Mecp2 | 103 |
| BP | GO:0016053 | organic acid biosynthetic process | 100/3915 | 291/17474 | 1,7592E-06 | 4,9211E-05 | 3,6648E-05 | Acadl/Cyp27a1/Acsl3/Pla2g4a/Glul/Fmo1/Prox1/Dse/Ggt1/Ggt5/Ilvbl/Lta4h/Shmt2/Ltc4s/Shmt1/Alox8/Acadvl/Alox12/Acaca/Brca1/Adi1/Mtr/Elovl2/Edn1/Syk/Dhfr/Elovl7/Slc1a3/Abcd2/Csad/Abat/Hacd2/Cbs/Abhd3/Stard4/Cd74/Fads2/Fads1/Psat1/Aldh1a1/Scd2/Scd1/Sephs1/Gad2/Ptgds/Ptges/Ass1/Ptgs1/Gad1/Nr1h3/Gatm/Acss1/Acss2/Ptgis/Pex2/Fabp5/Cyp7b1/Phgdh/Gstm7/Gstm1/Abcd3/Cth/Hacd4/Elovl1/Pla2g5/Prxl2b/Insig1/Ugdh/Acacb/Pla2g1b/Prkab1/Sdsl/Psph/Mlxipl/Plod3/Alox5ap/Pdk4/Aass/Tbxas1/Hpgds/Thnsl2/Mgll/Alox5/Ldhb/Apoc1/Ceacam1/Bcat2/Klhl25/Abhd2/Cln3/Erlin2/Agt/Acsbg1/Hacd3/Aldh1a2/Elovl5/Plod2/Rbp1/Mid1ip1/Hsd17b10 | 100 |

| | | | | | | | | | |
|---|---|---|---|---|---|---|---|---|---|
| BP | GO:0042692 | muscle cell differentiation | 139/3915 | 434/17474 | 1,8738E-06 | 5,1692E-05 | 3,8495E-05 | Cflar/Igfbp5/Bcl2/Zbed6/Lmod1/Csrp1/Cfh/Vangl2/Smyd3/H3f3a/Capn2/Prox1/Lama2/Hey2/Asf1a/Ptbp1/Igf1/Tmtc3/Csrp2/Frs2/Meis1/Dock2/Sgcd/Flii/Pmp22/Chrnb1/Spag9/Ramp2/Sox9/H3f3b/Id2/Cfl2/Dpf3/Dicer1/Nid1/Edn1/Ninj1/Mef2c/Nln/Isl1/Bmpr1a/Bmp4/Mmp14/Myh6/Dock5/Rb1/Ednrb/Col14a1/Myc/Ppara/Myh11/Tbx1/Cxadr/Tmem204/Vegfa/Trip10/Lama1/Zeb1/Nr3c1/Lox/Megf10/Pdgfrb/Csf1r/Smad4/Nfatc1/Efemp2/Pgm5/Smarca2/Myof/Chuk/Nrap/Nebl/Notch1/Tsc1/Eng/Ttn/Capn3/Spg11/Myl9/Ptgfrn/Casq2/Cd53/Sypl2/Camk2d/Npnt/Pdlim5/Acadm/Cth/Myorg/Zmpste24/Tcf23/Rbpj/Sgcb/Pdgfra/Hopx/Ankrd17/Myo18b/Tmem119/Tbx3/Gper1/Cav2/Smo/Ezh2/Dysf/Camk1/Atg7/Cxcl12/Cd9/Ccnd2/Ehd2/Tgfb1/Nphs1/Tshz3/Flt3l/Sox6/Il4ra/Fgfr2/Dock1/Igf2/Cd81/Atp11a/Nrg1/Ednra/Smad1/Nfatc3/Agt/Itgb1/Neo1/Rbpms2/Rora/Bnip2/Tmod3/Tbx18/Lamb2/Ctnnb1/Mecp2/G6pdx/Dmd/Foxo4 | 139 |
| BP | GO:0030193 | regulation of blood coagulation | 33/3915 | 68/17474 | 1,8792E-06 | 5,1692E-05 | 3,8495E-05 | Serpine2/Alox12/Gp1ba/Serpinf2/Vtn/C1qtnf1/F2r/Plau/Prkcd/Ano6/Abat/Pros1/Enpp4/Adrb2/Nfe2l2/Tfpi/Serping1/F2/Thbs1/Thbd/Anxa5/F3/Pdgfra/Cav1/Cd9/Apoe/Plaur/Ceacam1/Vkorc1/Plat/St3gal4/Ubash3b/Anxa2 | 33 |
| BP | GO:0098754 | detoxification | 33/3915 | 68/17474 | 1,8792E-06 | 5,1692E-05 | 3,8495E-05 | Prdx6/Mtarc2/Slc30a10/Slc30a1/Sesn1/Slc47a1/Nxn/Rdh11/Fbln5/Dhfr/Nnt/Ralbp1/Aldh1a1/Nfe2l2/Cat/Selenot/Gstm5/Gstm7/Gstm1/Slc39a8/Adh5/Prdx1/Abcb1a/Abcb1b/Nos3/Aldh2/Abcg2/Txnrd3/Mt3/Mt2/Mt1/Nqo1/Atp7a | 33 |
| BP | GO:1901653 | cellular response to peptide | 109/3915 | 325/17474 | 2,2922E-06 | 6,1816E-05 | 4,6034E-05 | Irs1/Adipor1/Fcgr2b/Eprs/Slc30a10/Enpp1/Fyn/Gja1/Appl2/Igf1/Stat6/Timeless/Cdk2/Grb10/Irf1/Myo1c/Stxbp4/Stat5b/Stat3/Lpin1/Snx6/Zfp36l1/Lgmn/Gpld1/Edn1/Gkap1/Pik3r1/Prkcd/Rb1/Tbc1d4/Ghr/Tns2/Sp1/Prkdc/Srsf3/Trem2/Kat2b/Rab31/Rock1/Camk2a/Adrb2/Rela/Kank1/Jak2/Ide/Sorbs1/Gpam/Prkcq/Vim/Pip4k2a/Nfe2l2/Zfp106/Snx5/Lpin3/Car2/Pik3ca/Anxa5/Foxo1/Shc1/Rab13/Ptpn22/Ahcyl1/Vcam1/Bcar3/Nfkb1/Ripk2/Chmp5/Tlr4/Dennd4c/Leprot/Pik3r3/Agtrap/Slc25a33/Prkcz/Insig1/Tlr6/Ptpn11/Gper1/Pdk4/Cav2/Cav1/Rarres2/Nod1/Ankrd26/Ceacam1/Akt2/Cyfip1/Igf1r/Inppl1/Pde3b/Mapk3/Ctsd/Igf2/Ano1/Insr/Eif4ebp1/Slc27a1/Ednra/Nod2/Agt/Sesn3/Icam1/Il18/Gnai2/Slc26a6/Ctnnb1/Ldoc1/Foxo4/Ogt | 109 |
| BP | GO:0061298 | retina vasculature development in camera-type eye | 15/3915 | 21/17474 | 2,3442E-06 | 6,3011E-05 | 4,6924E-05 | Arhgef15/Rhoj/Vstm4/Acvrl1/Lama1/Cyp1b1/Pdgfrb/Lrp5/Rom1/Clic4/Pdgfra/Tgfb1/Fzd4/Acvr2b/Ndp | 15 |
| BP | GO:0010952 | positive regulation of peptidase activity | 71/3915 | 191/17474 | 2,4327E-06 | 6,5178E-05 | 4,8538E-05 | Stat1/Casp8/Bok/Aim2/Mndal/Ifi203/Perp/Ccn2/Fyn/Vsir/Rack1/Nlrp3/Alox12/Nlrp1b/Atp2a3/Stat3/Grn/Lgmn/Syk/Ctsl/F2r/Sox7/Dap/Myc/Gramd4/Senp1/Tfap4/Ppm1f/Xdh/Jak2/Fas/Card9/Tank/Bcl2l11/Ddrgk1/Tnfsf10/Efna1/Ctsk/Ctss/F3/Casp6/Ccn1/Bcl10/Lyn/Asph/Ripk2/Casp8ap2/Anp32b/Acer2/Laptm5/Rest/Diablo/Hip1r/Hip1/Gper1/Hmgb1/Casp2/Nod1/Antxr1/Ret/Rcn3/Picalm/Arrb1/Pycard/Ctsd/Dlc1/Casp12/Uaca/Rps27l/Ctsh/Rhoa | 71 |
| BP | GO:0034109 | homotypic cell-cell adhesion | 40/3915 | 90/17474 | 2,7422E-06 | 7,112E-05 | 5,2963E-05 | Serpine2/Cfh/F11r/Ctnna3/Plek/Alox12/Gp1ba/Itga2b/Itgb3/C1qtnf1/Dsp/Syk/Prkcd/Lgals1/Abat/Cxadr/Dsc2/Megf10/Adrb2/Fermt3/Tjp2/Jak2/Prkcq/Ccm2l/Slc7a11/P2ry12/Pear1/Il6ra/Stxbp3/Lyn/Plpp3/Pdpn/Pdgfra/Pdia4/Ptpn6/Cd9/Plaur/Ceacam1/Ubash3b/Rdx | 40 |
| BP | GO:0099024 | plasma membrane invagination | 32/3915 | 66/17474 | 2,7493E-06 | 7,112E-05 | 5,2963E-05 | Gulp1/Fcgr2b/Fcgr3/Fcer1g/Itgb2/Appl2/Cd300a/Elmo1/Xkr6/Ano6/Bin2/Nckap1l/Snx9/Aif1/Trem2/Rab31/Megf10/Spire1/Thbs1/Lbp/Fcgr1/Abca1/Xkr8/Arhgap25/Gata2/Clec7a/Siglece/Itgam/Clcn3/Plcg2/Snx33/Rhoa | 32 |
| BP | GO:0006575 | cellular modified amino acid metabolic process | 47/3915 | 112/17474 | 2,7994E-06 | 7,1964E-05 | 5,3592E-05 | Acadl/Aldh9a1/Iyd/Aldh1l2/Shmt2/Slc22a5/Slc22a4/Shmt1/Mboat2/Dio2/Ckb/Mtr/Mboat1/Ctsl/Ptdss1/Dhfr/Serinc5/Gcnt4/Chdh/Ggact/Cpq/Csad/Cbs/Cdo1/Cpt1a/Sardh/Ass1/Bbox1/Gatm/Serinc3/Slco4a1/Ctsk/Acadm/Ggh/Cpt2/Mfsd2a/Crot/Reln/Plod3/Aldh1l1/Lpcat3/Slco1c1/Folh1/Slc27a1/Plscr1/Plod2/Slc16a2 | 47 |
| BP | GO:0051216 | cartilage development | 76/3915 | 209/17474 | 2,8746E-06 | 7,3667E-05 | 5,486E-05 | Col9a1/Arid5a/Col3a1/Gli2/Cfh/Prrx1/Ccn2/Enpp1/Efemp1/Rflnb/Col1a1/Axin2/Sox9/Mboat2/Trip11/Dicer1/Itgb8/Gli3/Gpld1/Bmp6/Edn1/Ogn/Smad5/Mef2c/Wnt5a/Mustn1/Bmpr1a/Bmp4/Rb1/Adamts12/Trps1/Rarg/Cbs/Col11a2/Twsg1/Zeb1/Rela/Mdk/Thbs1/Ddrgk1/Bmp7/Fgf2/Pbxip1/Ctsk/Bmpr1b/Ccn1/Tgfbr1/Col27a1/Nfib/Nfia/Zmpste24/Hspg2/Hes5/Fgfr3/Cytl1/Idua/Ptpn11/Creb3l2/Lrp6/Mgp/Sox5/Tgfb1/Chsy1/Serpinh1/Sox6/Mapk3/Csgalnact1/Smad1/Bbs2/Maf/Bmp5/Pth1r/Tgfbr2/Ctnnb1/Bgn/Atp7a | 76 |
| BP | GO:0043331 | response to dsRNA | 23/3915 | 41/17474 | 2,954E-06 | 7,4774E-05 | 5,5685E-05 | Stat1/Rftn2/Ralb/Ddx21/Irak3/Nfkbia/Ticam1/Riok3/Sting1/Ticam2/Slc3a2/Card9/Ifih1/Mavs/Nfkb1/Ripk2/P2rx7/Cav1/Zc3hav1/Irf3/Mapk3/Tlr3/Nod2 | 23 |
| BP | GO:2000177 | regulation of neural precursor cell proliferation | 50/3915 | 122/17474 | 3,1195E-06 | 7,8243E-05 | 5,8268E-05 | Gli2/Btg2/Gpr37l1/Aspm/Prox1/Nr2e1/Lims1/Appl2/Igf1/Kctd11/Lhx1/Id2/Gli3/Id4/Wnt5a/Vegfa/Lims2/Ctnna1/Rsu1/Notch1/Lhx2/Mdk/Pax6/Ctsz/Sox2/Fgf2/Foxo1/Nes/Gng5/Lyn/Fgfr3/Ptprz1/Smo/Ptn/Gata2/Tgfb1/Sirt2/Spint2/Ilk/Vegfc/Sall1/Adgrg1/Itgb1/Gnai2/Rhoa/Cx3cr1/Ctnnb1/Ccr5/Flna/Slc16a2 | 50 |
| BP | GO:0010035 | response to inorganic substance | 150/3915 | 480/17474 | 3,5474E-06 | 8,7382E-05 | 6,5073E-05 | Slc40a1/Cflar/Fn1/D2hgdh/Bcl2/Slc41a1/Gpr37l1/Nek7/Glrx2/Pla2g4a/Ddr2/Kcnj10/Itpkb/Slc30a10/Slc30a1/Enpp1/Fyn/Pln/Psap/Pcdh15/Adora2a/Fbxo7/Stat6/Timeless/Egfr/Rack1/Pld2/Slc13a5/Id2/Ngb/Fbln5/Lgmn/Serpina1c/Crip1/Net1/Mtr/Hfe/Gpld1/Ripk1/Bmp6/Edn1/Hk3/Ptch1/Mef2c/Dhfr/Prkcd/Glud1/Ripk3/Ednrb/Slc1a3/Ank/Slc38a2/Atf1/Abat/Spidr/Tfrc/Mylk/Casr/Itpr3/Slc25a23/Xdh/Cyp1b1/Nrxn1/Npc1/Pdgfrb/Tcirg1/Rela/Sipa1/Rasgrp2/Fas/Chuk/Pkd2l1/Ass1/Cybrd1/Nfe2l2/Ttn/Itgav/Cat/Thbs1/Capn3/B2m/Gatm/Kcnip3/Cpne1/Cp/Kcnmb2/Pik3ca/Sox2/Anxa5/Foxo1/S100a16/Syt6/Ahcyl1/Slc25a24/Casp6/Slc39a8/Asph/Cpne3/Ggh/Aco1/Prdx1/Clic4/Alpl/Hgf/Nos3/Khk/Fgfr3/Kdr/Pkd2/P2rx7/P2rx4/Rasa4/Alox5ap/Cav1/Ezh2/Pde1c/Slc6a1/Wnk1/Apobec1/Cdkn1b/Lig1/Axl/Abcc6/Iqgap1/Dlg2/Ucp3/Ucp2/Chp2/Mapk3/Atp7b/Adam9/Fnta/Casp3/Ednra/Adcy7/Mmp2/Mt3/Mt2/Mt1/Lcat/Nfatc3/Cdh1/Nfat5/Nqo1/Plcg2/Plscr1/Plscr2/Plcd1/Mecp2/Atp7a | 150 |
| BP | GO:0033627 | cell adhesion mediated by integrin | 39/3915 | 88/17474 | 3,9606E-06 | 9,6126E-05 | 7,1585E-05 | Itgb2/Vtn/Itga2b/Itgb3/Adam17/Itgb8/Syk/Itga1/Plau/Itga5/Nckap1l/Crkl/Itgb5/Cyp1b1/Fermt3/Lpxn/Dpp4/Itga6/Itgav/P2ry12/Efna1/Npnt/Lyn/Acer2/Plpp3/Col16a1/Ptpn11/Podxl/Ret/Wnk1/Ptpn6/Cib1/Swap70/Itgal/Itgam/Adam9/Itgb1/Icam1/Itga9 | 39 |
| BP | GO:0052547 | regulation of peptidase activity | 131/3915 | 410/17474 | 4,2055E-06 | 0,00010137 | 7,5492E-05 | Stat1/Cflar/Casp8/Serpine2/Bok/Aim2/Mndal/Ifi203/Perp/Ccn2/Fyn/Mical1/Vsir/Adora2a/Timp3/Igf1/Rack1/Nlrp3/Alox12/Nlrp1b/Atp2a3/Serpinf1/Serpinf2/Vtn/Rffl/Stat3/Grn/Tmed10/Lgmn/Serpina1c/Serpinb6b/Serpinb9/Serpinb6a/Syk/Ctsl/Cast/F2r/Naip2/Naip5/Naip6/Itih3/Sox7/Dap/Myc/Gramd4/Senp1/Tfap4/Ppm1f/Serpind1/Pak2/Psmb9/Psmb8/Vegfa/Xdh/Birc6/Dele1/Ubxn1/Jak2/Fas/Itih2/Itih5/Card9/Tank/Tfpi/Serping1/Cd44/Thbs1/Bcl2l11/Ddrgk1/Cst3/Tnfsf10/Sox2/Lxn/Efna1/Ctsk/Ctss/F3/Casp6/Ccn1/Bcl10/Lyn/Asph/Ripk2/Casp8ap2/Reck/Anp32b/Klf4/Acer2/Svbp/Hdac1/Laptm5/Rps6ka1/Hgf/Rest/Diablo/Hip1r/Hip1/Gper1/Hmgb1/Cav1/Casp2/Nod1/Antxr1/Timp4/Ret/A2m/Plaur/Akt2/Spint2/Rcn3/Picalm/Serpinh1/Arrb1/Pycard/Ctsd/Gas6/Dlc1/Spock3/Mt3/Cdh1/Wfdc1/Casp12/Uaca/Rps27l/Cd109/Ctsh/Rhoa/Xiap/Gpc3/Renbp/Rps6ka3 | 131 |
| BP | GO:0007044 | cell-substrate junction assembly | 42/3915 | 99/17474 | 6,6837E-06 | 0,00015096 | 0,00011242 | Dst/Map4k4/Fn1/Tns1/Bcl2/Epb41l5/Myoc/Lims1/Poldip2/Itgb3/Dusp22/Fam107a/Vcl/Fermt2/Mmp14/Acvrl1/Itga5/Ppm1f/Apod/Phldb2/Vegfa/Ston1/Rock1/Lama3/Sorbs1/Tsc1/Itgav/Sdc4/Tln1/Tek/Tesk2/Macf1/Col16a1/Limch1/Kdr/Ptpn11/Iqgap1/Thsd1/Dlc1/Clasp2/Gpm6b/Arhgap6 | 42 |
| BP | GO:0030278 | regulation of ossification | 54/3915 | 138/17474 | 6,7271E-06 | 0,00015126 | 0,00011264 | Bcl2/Pbx1/Ddr2/Enpp1/Gja1/Srgn/Kremen1/Rflnb/Nbr1/Sox9/Notum/Asxl2/Prl/Bmp6/Omd/Mef2c/Wnt5a/Bmpr1a/Bmp4/Nipbl/Ank/Ano6/Csf1r/Adrb2/Notch1/Lrp4/Mdk/Bmp7/Csf1/S1pr1/Sgms2/Bmpr1b/Ccn1/Zmpste24/Dhrs3/Isg15/Atraid/Fgfr3/Rbpj/Bmp2k/Tmem119/Ptpn11/P2rx7/Ptn/Alox5/Lrp6/Mgp/Tgfb1/Chsy1/P2ry2/Mapk3/Setd2/Acvr2b/Gpm6b | 54 |

| | | | | | | | | | |
|---|---|---|---|---|---|---|---|---|---|
| BP | GO:0071559 | response to transforming growth factor beta | 83/3915 | 238/17474 | 6,8217E-06 | 0,00015297 | 0,00011392 | Col3a1/Cflar/Epb41l5/Lats1/Fyn/Tet1/Pbld2/Appl2/Sap30l/Stat3/Smurf2/Sox9/Adam17/Snx6/Fut8/Zfp36l1/Itgb8/Dusp22/Aspn/Smad5/Mef2c/Map3k1/Il17rd/Fermt2/Dab2/Ppara/Acvrl1/Vasn/Crkl/Cldn5/Nrros/Itgb5/Runx1/Twsg1/Ltbp1/Zfp36l2/Zeb1/Rock1/Nrep/Nr3c1/Lox/Smad4/Smad2/Lpxn/Eng/Dlx1/Spi1/Spred1/Thbs1/Slc2a10/Npnt/Bmpr1b/Map3k7/Tgfbr1/Hdac1/Prdm16/Nos3/Yes1/Htra3/Ppargc1a/Tgfbr3/Col1a2/Cav2/Cav1/Hipk2/Sox5/Tgfb1/Igf1r/Lrrc32/Sox6/Tgfb1i1/Htra1/Adam9/Hpgd/Smad1/Cdh5/Zfhx3/Pdgfd/Dnm2/Cd109/Tgfbr2/Ndp/Ogt | 83 |
| BP | GO:0002577 | regulation of antigen processing and presentation | 14/3915 | 20/17474 | 7,5493E-06 | 0,00016481 | 0,00012273 | Slc11a1/Cd68/Hfe/Tap2/H2-Ob/Trem2/Cd74/Thbs1/Fgl2/Nod1/Tapbpl/Pycard/Nod2/Was | 14 |
| BP | GO:0032943 | mononuclear cell proliferation | 106/3915 | 322/17474 | 8,0274E-06 | 0,00017312 | 0,00012892 | Igfbp2/Slc11a1/Inpp5d/Bcl2/Ptprc/Rc3h1/Fcgr2b/Fyn/Gja1/Vsir/Itgb2/Icosl/Igf1/Kitl/Dock2/Havcr2/Irf1/Tnfrsf13b/Lgals9/Stat5b/Cd300a/Syk/Mef2c/Cd180/Il6st/Adk/Prkcd/Bmp4/Lgals3/Pnp/Ripk3/Gpr183/Il7r/Myc/Rac2/Lmbr1l/Nckap1l/Emp2/St6gal1/Dlg1/Tfrc/Cd86/Cd80/Ifnar2/Cdkn1a/Rasal3/H2-Aa/Aif1/Ticam1/Twsg1/Cd74/Csf1r/Dock8/Jak2/Cd274/Gpam/Prkcq/Traf6/Cd44/Cd59a/Sdc4/Cd40/Cd1d1/Ptpn22/Csf1/Vav3/Vcam1/Pde5a/Lef1/Lyn/Ripk2/Tlr4/Laptm5/Pla2g5/Tnfrsf1b/Tnfrsf14/Cd38/P2rx7/Hmgb1/Gpnmb/Cxcl12/Ptpn6/Ceacam1/Tgfb1/Tyrobp/Flt3l/Cd37/Lrrc32/Itgal/Pycard/Itgam/Cd151/Igf2/Cd81/Casp3/Il15/Tirap/Il18/Tgfbr2/Myd88/Ctnnb1/Ccr2/Sash3/Elf4/Msn/Btk | 106 |
| BP | GO:0006911 | phagocytosis, engulfment | 28/3915 | 57/17474 | 8,3029E-06 | 0,0001775 | 0,00013219 | Gulp1/Fcgr2b/Fcgr3/Fcer1g/Itgb2/Appl2/Cd300a/Elmo1/Xkr6/Ano6/Bin2/Nckap1l/Aif1/Trem2/Rab31/Megf10/Thbs1/Lbp/Fcgr1/Abca1/Xkr8/Arhgap25/Gata2/Clec7a/Siglece/Itgam/Clcn3/Plcg2 | 28 |
| BP | GO:0034446 | substrate adhesion-dependent cell spreading | 43/3915 | 103/17474 | 8,5475E-06 | 0,00018179 | 0,00013538 | Fn1/Lamc2/Myoc/Lims1/Ntn4/Itgb3/P4hb/Carmil1/Nedd9/Fermt2/Dock5/Dab2/Has2/Triobp/Parvg/Fbln1/Crkl/St6gal1/Lims2/Fermt3/Lpxn/Kank1/Lamc3/Itgav/Mdk/Mertk/Lama5/Efna1/Tek/Pdpn/Vamp3/Antxr1/Axl/Cib1/Fzd4/Ilk/Parva/Dock1/Dnm2/Rhoa/Lamb2/Cspg5/Flna | 43 |
| BP | GO:0032963 | collagen metabolic process | 47/3915 | 116/17474 | 8,7843E-06 | 0,00018539 | 0,00013806 | Tram2/Fn1/Suco/Ddr2/Ccn2/Vsir/Mfap4/Adora2b/Serpinf2/Col1a1/P3h4/Retreg3/Mrc2/Dicer1/Ctsl/F2r/Prkcd/Bmp4/Mmp14/Rgcc/Rapgef3/Tns2/Ltbp1/Pdgfrb/Vim/Notch1/Eng/F2/Cst3/Id1/Il6ra/Ctsk/Ctss/Rap1a/Cyp2j6/Fgfr3/Tgfbr3/Idua/P2rx7/Plod3/Col1a2/Tgfb1/Rcn3/Serpinh1/Mmp2/Itgb1/Il18 | 47 |
| BP | GO:0048002 | antigen processing and presentation of peptide antigen | 30/3915 | 63/17474 | 8,846E-06 | 0,00018622 | 0,00013868 | Slc11a1/Fcgr2b/Fcgr3/Fcer1g/Hfe/Ctsl/Erap1/H2-K1/Tap1/Tap2/H2-Ob/H2-Aa/H2-Eb1/H2-D1/H2-Q4/H2-T23/H2-T22/Trem2/Cd74/Unc93b1/Ide/Traf6/B2m/Ctss/Fcgr1/Marchf8/Tapbpl/Mill2/Pycard/Ifi30 | 30 |
| BP | GO:0050818 | regulation of coagulation | 33/3915 | 72/17474 | 8,9745E-06 | 0,00018749 | 0,00013962 | Serpine2/Alox12/Gp1ba/Serpinf2/Vtn/C1qtnf1/F2r/Plau/Prkcd/Ano6/Abat/Pros1/Enpp4/Adrb2/Nfe2l2/Tfpi/Serping1/F2/Thbs1/Thbd/Anxa5/F3/Pdgfra/Cav1/Cd9/Apoe/Plaur/Ceacam1/Vkorc1/Plat/St3gal4/Ubash3b/Anxa2 | 33 |
| BP | GO:0060856 | establishment of blood-brain barrier | 13/3915 | 18/17474 | 9,4855E-06 | 0,00019667 | 0,00014646 | Lrp5/Ttpa/Reck/Mfsd2a/Mxra8/Abcb1b/Bpgm/Lrp6/Lsr/Fzd4/Ctnnb1/Ndp/Dmd | 13 |
| BP | GO:0043114 | regulation of vascular permeability | 24/3915 | 46/17474 | 9,8214E-06 | 0,00020211 | 0,00015051 | Akap12/Adora2a/Ramp2/Bmp6/Ocln/Fermt2/Angpt1/Cldn5/Vegfa/Nr3c1/Tjp2/Fgfbp3/Ddah1/Ctnnbip1/Hrh1/Pde3a/Gpr4/Apoe/Ceacam1/Tgfb1/Tjp1/Cdh5/Il18/Amot | 24 |
| BP | GO:0048863 | stem cell differentiation | 88/3915 | 258/17474 | 9,9794E-06 | 0,00020485 | 0,00015255 | Sox17/Nrp2/Erbb4/Fn1/Bcl2/Ptprc/Vsir/Mbd3/Kitl/Chd3/Msi2/Stat3/Etv4/Wnt3/Sox9/Dicer1/Foxc1/Edn1/Fam172a/Mef2c/Isl1/Sema3g/Bmpr1a/Bmp4/Ednrb/Zic5/Zic2/Sema5a/Lmbr1l/Prkdc/Tbx1/Lbh/Eif2ak2/Mta3/Zfp36l2/Epcam/Sema6a/Smad4/Notch1/Phf19/Nfe2l2/Pax6/Sema6d/Shc4/Jag1/Bmp7/Edn3/Lama5/Sox2/Fgf2/Tal1/Hdac1/Phactr4/Hes5/Cdk6/Sema3d/Sema3c/Pus7/Rbpj/Pdgfra/Rest/Tbx3/Smo/Ptn/Ezh2/Kdm3a/Tcf7l1/Ret/A2m/Lrp6/Sox5/Zfp36/Tead2/Chd2/Sox6/Mapk3/Fgfr2/Nrg1/Gatad2a/Ednra/Itgb1/Yap1/Aldh1a2/Tbx18/Setd2/Ctnnb1/Zic3/Foxo4 | 88 |
| BP | GO:0046651 | lymphocyte proliferation | 104/3915 | 317/17474 | 1,1465E-05 | 0,00023245 | 0,00017311 | Igfbp2/Slc11a1/Inpp5d/Bcl2/Ptprc/Rc3h1/Fcgr2b/Fyn/Gja1/Vsir/Itgb2/Icosl/Igf1/Kitl/Dock2/Havcr2/Irf1/Tnfrsf13b/Lgals9/Stat5b/Cd300a/Syk/Mef2c/Cd180/Il6st/Adk/Prkcd/Bmp4/Lgals3/Pnp/Ripk3/Gpr183/Il7r/Myc/Rac2/Lmbr1l/Nckap1l/Emp2/Dlg1/Tfrc/Cd86/Cd80/Ifnar2/Cdkn1a/Rasal3/H2-Aa/Aif1/Ticam1/Twsg1/Cd74/Csf1r/Dock8/Jak2/Cd274/Gpam/Prkcq/Traf6/Cd44/Cd59a/Sdc4/Cd40/Cd1d1/Ptpn22/Vav3/Vcam1/Pde5a/Lef1/Lyn/Ripk2/Tlr4/Laptm5/Pla2g5/Tnfrsf1b/Tnfrsf14/Cd38/P2rx7/Hmgb1/Gpnmb/Cxcl12/Ptpn6/Ceacam1/Tgfb1/Tyrobp/Flt3l/Cd37/Lrrc32/Itgal/Pycard/Itgam/Cd151/Igf2/Cd81/Casp3/Il15/Tirap/Il18/Tgfbr2/Myd88/Ctnnb1/Ccr2/Sash3/Elf4/Msn/Btk | 104 |
| BP | GO:0120161 | regulation of cold-induced thermogenesis | 56/3915 | 147/17474 | 1,1764E-05 | 0,00023794 | 0,00017719 | Plcl1/Acadl/Adipor1/G0s2/Lama4/Gja1/Appl2/Ddit3/Stat6/Grb10/Pctp/Lpin1/Adam17/Nova1/Dio2/Syk/Rb1/Lnpep/Vegfa/Epas1/Adrb2/Aldh1a1/Jak2/Scd1/Notch1/Nr1h3/Lgr4/Gatm/Id1/Fabp5/Hadh/Decr1/Tlr4/Dock7/Lepr/Oma1/Acot11/Cpt2/Alpl/Prdm16/Ppargc1a/Rbpj/Cmklr1/Prkab1/Cav1/Kdm3a/Alms1/Adipor2/Igf1r/Ucp2/Il4ra/Il15/Il18/Acvr2b/Ccr2/Ogt | 56 |
| BP | GO:0071560 | cellular response to transforming growth factor beta stimulus | 81/3915 | 234/17474 | 1,1985E-05 | 0,0002418 | 0,00018007 | Col3a1/Cflar/Epb41l5/Lats1/Fyn/Tet1/Pbld2/Appl2/Sap30l/Stat3/Smurf2/Sox9/Adam17/Snx6/Fut8/Zfp36l1/Itgb8/Dusp22/Aspn/Smad5/Mef2c/Map3k1/Il17rd/Fermt2/Dab2/Ppara/Acvrl1/Vasn/Crkl/Cldn5/Nrros/Itgb5/Runx1/Twsg1/Ltbp1/Zfp36l2/Zeb1/Nrep/Nr3c1/Lox/Smad4/Smad2/Lpxn/Eng/Dlx1/Spi1/Spred1/Thbs1/Slc2a10/Npnt/Bmpr1b/Map3k7/Tgfbr1/Hdac1/Prdm16/Nos3/Yes1/Htra3/Ppargc1a/Tgfbr3/Col1a2/Cav2/Cav1/Hipk2/Sox5/Tgfb1/Igf1r/Lrrc32/Sox6/Tgfb1i1/Htra1/Adam9/Hpgd/Smad1/Cdh5/Pdgfd/Dnm2/Cd109/Tgfbr2/Ndp/Ogt | 81 |
| BP | GO:0032944 | regulation of mononuclear cell proliferation | 86/3915 | 252/17474 | 1,2227E-05 | 0,00024549 | 0,00018282 | Igfbp2/Inpp5d/Bcl2/Ptprc/Rc3h1/Fcgr2b/Vsir/Icosl/Igf1/Kitl/Havcr2/Irf1/Tnfrsf13b/Lgals9/Stat5b/Cd300a/Syk/Mef2c/Il6st/Adk/Bmp4/Lgals3/Pnp/Ripk3/Gpr183/Rac2/Nckap1l/St6gal1/Dlg1/Tfrc/Cd86/Cd80/Cdkn1a/Rasal3/H2-Aa/Aif1/Ticam1/Twsg1/Cd74/Csf1r/Jak2/Cd274/Gpam/Prkcq/Traf6/Cd44/Cd59a/Sdc4/Cd40/Cd1d1/Ptpn22/Csf1/Vav3/Vcam1/Pde5a/Lyn/Ripk2/Tlr4/Laptm5/Pla2g5/Tnfrsf1b/Tnfrsf14/Cd38/Hmgb1/Gpnmb/Ptpn6/Ceacam1/Tgfb1/Tyrobp/Flt3l/Cd37/Lrrc32/Itgal/Pycard/Igf2/Cd81/Casp3/Il15/Tirap/Il18/Tgfbr2/Myd88/Ctnnb1/Ccr2/Sash3/Btk | 86 |
| BP | GO:0010464 | regulation of mesenchymal cell proliferation | 22/3915 | 41/17474 | 1,2775E-05 | 0,00025525 | 0,00019008 | Irs1/Prrx1/Sox9/Arhgap5/Wnt5a/Bmpr1a/Bmp4/Myc/Tbx1/Zeb1/Nfib/Ctnnbip1/Kdr/Hmgb1/Foxp2/Smo/Ptn/Lrp6/Fgfr2/Tbx18/Tgfbr2/Ctnnb1 | 22 |
| BP | GO:0061082 | myeloid leukocyte cytokine production | 30/3915 | 64/17474 | 1,3101E-05 | 0,00025988 | 0,00019354 | Fcer1g/Ddx21/Irak3/Nlrp3/Syk/Wnt5a/Litaf/Ticam1/Ticam2/Cd74/Card9/Mavs/Tlr2/Ripk2/Tlr4/Laptm5/P2rx7/Nod1/Tgfb1/Axl/Gprc5b/Pycard/Gas6/Tlr3/Nod2/Plcg2/Casp4/Tirap/Myd88/Tlr7 | 30 |
| BP | GO:0040013 | negative regulation of locomotion | 112/3915 | 348/17474 | 1,4223E-05 | 0,00028012 | 0,0002086 | Col3a1/Igfbp5/Sp100/Bcl2/Srgap2/Adipor1/Dusp10/Tet1/Adora2a/Cnn2/Cd63/Sap30l/Atp1b2/Serpinf1/Stat3/Wnt3/Gna13/Cd300a/Rin3/Gm266/Dusp22/Nedd9/Mef2c/Wnt5a/Sema3g/Mmrn2/Bmpr1a/Lrch1/Rgcc/Sema5a/Card10/Fbln1/Arid2/Acvrl1/Cldn5/St6gal1/Apod/Cd200r1/Phldb2/Aif1/Ptprm/Cyp1b1/Ctnna1/Mcc/Sema6a/Cd74/Coro1b/Kank1/Fas/Notch1/Egfl7/Pip5kl1/Eng/Dpp4/Nfe2l2/Spred1/Thbs1/Sema6d/Jag1/Ptprt/Fgf2/Cers2/Trp53inp1/Reck/Rnf20/Klf4/Tie1/Svbp/Cldn19/Hdac1/Clic4/Padi2/Sema3d/Sema3c/Ppargc1a/Limch1/Tgfbr3/Hmgb1/Stard13/Cav1/Ptn/Adamts9/Cxcl12/Adipor2/Arhgdib/C5ar2/Apoe/Tgfb1/Spint2/Fuz/Rras/Abhd2/Idh2/Ilk/Ifitm1/Angpt2/Nrg1/Dlc1/Bst2/Adgrg1/Cdh1/S1pr2/Ryk/Rhoa/Clasp2/Cx3cr1/Ccr5/Was/Rap2c/Arhgap4/Mecp2/Ogt | 112 |

| | | | | | | | | | |
|---|---|---|---|---|---|---|---|---|---|
| BP | GO:1990845 | adaptive thermogenesis | 61/3915 | 165/17474 | 1,468E-05 | 0,00028843 | 0,00021479 | Plcl1/Acadl/Sctr/Pm20d1/Adipor1/G0s2/Lama4/Gja1/Appl2/Ddit3/Stat6/Grb10/Pctp/Lpin1/Adam17/Nova1/Dio2/Ckb/Syk/Rb1/Lnpep/Clic5/Vegfa/Epas1/Adrb2/Aldh1a1/Jak2/Scd1/Notch1/Nr1h3/Lgr4/Gatm/Id1/Fabp5/Hadh/Decr1/Tlr4/Dock7/Lepr/Oma1/Acot11/Cpt2/Alpl/Prdm16/Ppargc1a/Rbpj/Cmklr1/Prkab1/Cav1/Kdm3a/Alms1/Adipor2/Igf1r/Ucp3/Ucp2/Il4ra/Il15/Il18/Acvr2b/Ccr2/Ogt | 61 |
| BP | GO:0051495 | positive regulation of cytoskeleton organization | 71/3915 | 200/17474 | 1,5495E-05 | 0,00030372 | 0,00022618 | Vil1/Lmod1/Myoc/Pfdn2/Prox1/Ccn2/Plek/Ccdc88a/Sh3pxd2b/Pdlim4/Arhgef15/Serpinf2/Myo1c/Spag5/Cfl2/Carmil1/Edn1/Map3k1/Fermt2/Cenpj/Sorbs3/Rgcc/Rictor/Sema5a/Mtss1/Baiap2l2/Rapgef3/Nckap1l/Ppm1f/Dlg1/Snx9/Daam2/Add3/Prkcq/Tsc1/Ckap5/Id1/Hck/Mapre1/Sdc4/Kirrel/Nes/Rhoc/Gpsm2/Sass6/Synpo2/Tgfbr1/Cdk5rap2/Tek/Wasf2/Akap9/Limch1/P2rx7/Wasf3/Cav1/Arhgef5/Cdkn1b/Vasp/Nphs1/Cyfip1/Fes/Fchsd2/Swap70/Pycard/Arhgef10/Fhod1/Icam1/Rhoa/Was/Mecp2/Flna | 71 |
| BP | GO:0009595 | detection of biotic stimulus | 20/3915 | 36/17474 | 1,5951E-05 | 0,00031191 | 0,00023228 | Ly96/Nlrp3/Naip2/Naip5/Naip6/Tspo/Trem2/Itgav/Lbp/Tlr2/Tlr4/Tlr1/Tlr6/Smo/Nod1/Clec7a/Pglyrp1/Nod2/Yap1/Srpx | 20 |
| BP | GO:2000146 | negative regulation of cell motility | 102/3915 | 313/17474 | 1,8979E-05 | 0,00036254 | 0,00026999 | Col3a1/Igfbp5/Sp100/Bcl2/Srgap2/Adipor1/Dusp10/Tet1/Cnn2/Cd63/Sap30l/Atp1b2/Serpinf1/Stat3/Gna13/Cd300a/Rin3/Gm266/Dusp22/Nedd9/Mef2c/Mmrn2/Bmpr1a/Lrch1/Rgcc/Card10/Fbln1/Arid2/Acvrl1/Cldn5/Apod/Cd200r1/Phldb2/Aif1/Ptprm/Cyp1b1/Ctnna1/Mcc/Cd74/Coro1b/Kank1/Fas/Notch1/Egfl7/Pip5kl1/Eng/Dpp4/Nfe2l2/Spred1/Thbs1/Sema6d/Jag1/Ptprt/Fgf2/Cers2/Trp53inp1/Reck/Rnf20/Klf4/Tie1/Svbp/Cldn19/Hdac1/Clic4/Padi2/Ppargc1a/Limch1/Tgfbr3/Hmgb1/Stard13/Cav1/Ptn/Adamts9/Cxcl12/Adipor2/Arhgdib/C5ar2/Apoe/Tgfb1/Spint2/Fuz/Rras/Abhd2/Idh2/Ilk/Ifitm1/Angpt2/Nrg1/Dlc1/Bst2/Adgrg1/Cdh1/S1pr2/Rhoa/Clasp2/Cx3cr1/Ccr5/Was/Rap2c/Arhgap4/Mecp2/Ogt | 102 |
| BP | GO:0042552 | myelination | 58/3915 | 156/17474 | 1,9402E-05 | 0,00036891 | 0,00027473 | Nab1/Ormdl1/Myoc/Kcnj10/Adgrg6/Enpp1/Fyn/Psap/S100b/Igf1/Pmp22/Tmem98/Lpin1/Galc/Dicer1/Id4/Ntrk2/Serinc5/Hexb/Zfp488/Clu/Abcd2/Rarg/Cldn5/Dlg1/Jam2/Zfp24/Tcf7l2/Tsc1/Ckap5/Mall/Tlr2/Pou3f2/Mpdz/Tnfrsf1b/Hes5/Hgf/Fgfr3/Rnf10/Gjc3/Wasf3/Ptprz1/Ptn/Cd9/Tgfb1/Akt2/Sirt2/Lgi4/Cyfip1/Ctsc/Ilk/Arhgef10/Nrg1/Pllp/Pard3/Hexa/Ctnnb1/Gpm6b | 58 |
| BP | GO:0042113 | B cell activation | 91/3915 | 273/17474 | 1,9843E-05 | 0,0003747 | 0,00027904 | Dock10/Inpp5d/Bcl2/Ptprc/Rc3h1/Fcgr2b/Enpp1/Icosl/Stat6/Ikzf1/Tnfrsf13b/Tnfsf13/Stat5b/Cd300a/Adam17/Id2/Zfp36l1/Lgals8/Syk/Mef2c/Pik3r1/Cd180/Prkcd/Shld2/Pnp/Mmp14/Gpr183/Il7r/Mfng/Lgals1/Nfam1/Hdac7/Nckap1l/Prkdc/Tfrc/Cd86/Cdkn1a/Ticam1/Zfp36l2/Cd74/Nfatc1/Kmt5b/Tcirg1/Fas/Blnk/Rif1/Dpp4/Spi1/Shld1/Cd40/Fcrl1/Notch2/Vav3/Lef1/Bank1/Lyn/Exosc3/Tlr4/Txlna/Laptm5/Themis2/Paxip1/Tnip2/Cd38/Rbpj/Lat2/Lfng/Ezh2/Skap2/Ptpn6/Tgfb1/Tyrobp/Cebpg/Swap70/Cd81/Casp3/Lyl1/Nod2/Plcg2/Irf8/Tirap/Parp3/Cmtm7/Dock11/Sash3/Atp11c/Hmgb3/Il2rg/Itm2a/Btk/Sh3kbp1 | 91 |
| BP | GO:0150115 | cell-substrate junction organization | 43/3915 | 106/17474 | 2,0093E-05 | 0,00037684 | 0,00028064 | Dst/Map4k4/Fn1/Tns1/Bcl2/Epb41l5/Myoc/Lims1/Poldip2/Itgb3/Dusp22/Pik3r1/Fam107a/Vcl/Fermt2/Mmp14/Acvrl1/Itga5/Ppm1f/Apod/Phldb2/Vegfa/Ston1/Rock1/Lama3/Sorbs1/Tsc1/Itgav/Sdc4/Tln1/Tek/Tesk2/Macf1/Col16a1/Limch1/Kdr/Ptpn11/Iqgap1/Thsd1/Dlc1/Clasp2/Gpm6b/Arhgap6 | 43 |
| BP | GO:0007405 | neuroblast proliferation | 37/3915 | 87/17474 | 2,1655E-05 | 0,00040337 | 0,00030039 | Btg2/Aspm/Prox1/Nr2e1/Frs2/Acsl6/Kctd11/Numb/Gli3/Id4/Plxnb2/Nde1/Tead3/Vegfa/Ctnna1/Notch1/Pax6/Sox2/Fgf2/Lef1/Akna/Dock7/Fgfr3/Smo/Ptn/Gata2/Lrp6/Sox5/Tgfb1/Fgfr2/Vegfc/Hhip/Sall1/Itgb1/Cx3cr1/Ctnnb1/Fgf13 | 37 |
| BP | GO:0071887 | leukocyte apoptotic process | 51/3915 | 133/17474 | 2,2998E-05 | 0,00042744 | 0,00031831 | Casp8/Bcl2/Fcgr2b/Fcer1g/Itpkb/Kitl/Adam17/Siva1/Gli3/Serpinb9/Ripk1/Mef2c/Wnt5a/Bmp4/Lgals3/Pnp/Ripk3/Il7r/Myc/Lmbr1l/Crkl/St6gal1/Hcls1/Cd74/Dock8/Cd274/Fas/Gpam/Casp7/Prkcq/Cd44/Bcl2l11/Mertk/Slc7a11/Efna1/Rorc/Bcl10/Lyn/P2rx7/Gimap8/Vhl/Cxcl12/Prkd2/Axl/Itgam/Gas6/Nod2/Il18/Plekho2/Ccr5/Tsc22d3 | 51 |
| BP | GO:0060070 | canonical Wnt signaling pathway | 97/3915 | 296/17474 | 2,3403E-05 | 0,00043204 | 0,00032174 | Sox17/Ccnyl1/Sox13/Aspm/Lats1/Bicc1/Ddit3/Kremen1/Znrf3/Egfr/Tmem88/Mks1/Col1a1/Fzd2/Wnt3/Axin2/Sox9/Notum/Sdc1/Nid1/Gli3/Sfrp4/Edn1/Zbed3/Map3k1/Isl1/Wnt5a/Amer2/Sox7/Scel/Ednrb/Gpc5/Dab2/Sema5a/Ubr5/Fzd6/Cthrc1/Lmbr1l/Daam2/Ppm1b/Mcc/Lrp5/Tle4/Kank1/Tcf7l2/Nrarp/Notch1/Lypd6/Ctnnd1/Lrp4/Mdk/Lgr4/Usp8/Sox2/Fgf2/Jade1/Foxo1/Wwtr1/Lef1/Dkk2/Nfkb1/Wls/Reck/Invs/Klf4/Plpp3/Rspo1/Hdac1/Ctnnbip1/Fgfr3/Rbpj/Cav1/Tcf7l1/Wnk1/Lrp6/Apoe/Tgfb1/Fuz/Lrrk1/Fzd4/Ilk/Gprc5b/Fgfr2/Ednra/Nkd1/Cdh1/Yap1/Tbx18/Ppp2r3a/Nphp3/Rbms3/Ctnnb1/Limd1/Ndp/Xiap/Gpc3/Amer1 | 97 |
| BP | GO:0001666 | response to hypoxia | 74/3915 | 213/17474 | 2,4273E-05 | 0,00044412 | 0,00033074 | Bcl2/Kcnk2/Cr1l/Pld2/Nos2/P4hb/Adam17/Egln3/Zfp36l1/Ngb/Edn1/Camk2g/Plau/Ang/Bnip3l/Rgcc/Myc/Ppara/Tmbim6/Vasn/Abat/Cbs/Vegfa/Trem2/Epas1/Adrb2/Smad4/Acaa2/Pygm/Vegfb/Notch1/Eng/Dpp4/Nfe2l2/Cat/Ptgis/Arnt/Camk2d/Ddah1/Tek/Plekhn1/Tacc3/Cd38/Ppargc1a/Rbpj/Kdr/Rest/Flt1/Cav1/Ndnf/Vhl/Itpr2/Hif3a/Sirt2/Fzd4/Ucp3/Ucp2/Plat/Eif4ebp1/Vegfc/Ednra/Mmp2/Mt3/Nfatc3/Il18/Rora/Limd1/Cybb/Ndp/Fundc1/Kdm6a/Mecp2/Ogt/Alas2 | 74 |
| BP | GO:0072089 | stem cell proliferation | 50/3915 | 131/17474 | 3,1799E-05 | 0,00056676 | 0,00042206 | Sox17/Gli2/Ptprc/Prg4/Prrx1/Pbx1/Irf6/Nr2e1/Kitl/Znrf3/Rnf43/Wnt3/Ace/Axin2/Sox9/Zfp36l1/Gli3/Ptch1/Wnt5a/Fermt2/Bmp4/Dab2/Fbln1/Rarg/Eif2ak2/Epcam/Nfatc1/Notch1/Mecom/Fgf2/Nes/Nfib/Cdkn2c/Ago3/Abcb1a/Abcb1b/Rbpj/Kdr/Tbx3/Lrp6/Tgfb1/Fgfr2/Mki67/Vegfc/Atxn1l/Yap1/Cd109/Tbx18/Tgfbr2/Ctnnb1 | 50 |
| BP | GO:0044546 | NLRP3 inflammasome complex assembly | 18/3915 | 32/17474 | 3,3864E-05 | 0,00060081 | 0,00044743 | Nek7/Aim2/Nlrp3/Prkd1/Nlrc3/Trem2/Eif2ak2/Mavs/Ptpn22/Gbp5/Tlr4/Tlr6/Sirt2/Trim30a/Pycard/Plcg2/Myd88/Btk | 18 |
| BP | GO:0019724 | B cell mediated immunity | 54/3915 | 145/17474 | 3,5009E-05 | 0,00061599 | 0,00045872 | Inpp5d/Ptprc/Cfh/Fcgr2b/Fcgr3/Fcer1g/Cr1l/Enpp1/Icosl/Stat6/Tnfsf13/Prkcd/Shld2/Csf2rb2/Csf2rb/Tfrc/H2-Ob/H2-Aa/H2-Eb1/C4b/C2/Trem2/Cd74/Kmt5b/Tcirg1/Fas/Card9/Rif1/Serping1/B2m/Shld1/Cd40/Fcgr1/Bcl10/Exosc3/Csf3r/C1qb/C1qc/C1qa/Fgl2/Paxip1/Gimap5/Ptpn6/Nectin2/Tgfb1/Swap70/Il4ra/Il21r/Irf7/Cd81/Nod2/Parp3/Myd88/Btk | 54 |
| BP | GO:0006739 | NADP metabolic process | 24/3915 | 49/17474 | 3,8551E-05 | 0,00066414 | 0,00049459 | Idh1/Ncf2/Fmo1/Fmo2/Hsd11b1/Aldh1l2/Fdxr/Dcxr/Nnt/Nudt13/Tkt/Nadk2/Nudt12/Me2/Fmo5/Pgd/Nadk/Ncf1/Aldh1l1/Idh2/Nqo1/Me1/G6pdx/Prps2 | 24 |
| BP | GO:0002440 | production of molecular mediator of immune response | 89/3915 | 270/17474 | 3,9002E-05 | 0,00066522 | 0,00049539 | Arid5a/Il1r1/Slc11a1/Ptprc/Dennd1b/Fcgr2b/Fcer1g/Enpp1/Vsir/Ddx21/Icosl/Irak3/Stat6/Nlrp3/Tnfsf13/Hfe/Syk/Wnt5a/Shld2/Lacc1/Il7r/Angpt1/Tmbim6/Litaf/Prkdc/Dlg1/Tfrc/Cd86/H2-Ob/H2-Aa/H2-Eb1/Ticam1/Ticam2/Cd74/Kmt5b/Tcirg1/Fas/Card9/Rif1/Traf6/B2m/Mavs/Shld1/Samhd1/Cd40/Tlr2/Ptpn22/Ripk2/Map3k7/Exosc3/Tlr4/Tek/Laptm5/Tnfrsf1b/Tnfrsf14/Prkcz/Fgl2/Paxip1/P2rx7/Cyren/Ezh2/Gimap5/Tril/Nod1/Tgfb1/Axl/Lgals4/Cd37/Swap70/Gprc5b/Il4ra/Stx4a/Pycard/Cd81/Gas6/Tlr3/Bst2/Nod2/Plcg2/Casp4/Tirap/Il18/Parp3/Myd88/Ccr2/Sash3/Itm2a/Btk/Tlr7 | 89 |
| BP | GO:0006907 | pinocytosis | 14/3915 | 22/17474 | 3,9178E-05 | 0,00066522 | 0,00049539 | Appl2/Dock2/Ankfy1/Carmil1/Nr1h3/Ehd4/Snx5/Cav1/Axl/Cln3/Pycard/Dnm2/Snx33/Mapkapk3 | 14 |
| BP | GO:0140632 | inflammasome complex assembly | 19/3915 | 35/17474 | 4,0188E-05 | 0,00067541 | 0,00050298 | Nek7/Aim2/Nlrp3/Nlrp1b/Prkd1/Nlrc3/Trem2/Eif2ak2/Mavs/Ptpn22/Gbp5/Tlr4/Tlr6/Sirt2/Trim30a/Pycard/Plcg2/Myd88/Btk | 19 |
| BP | GO:0010712 | regulation of collagen metabolic process | 28/3915 | 61/17474 | 4,0694E-05 | 0,00068115 | 0,00050725 | Fn1/Suco/Ddr2/Ccn2/Vsir/Mfap4/Adora2b/Serpinf2/Dicer1/F2r/Bmp4/Rgcc/Rapgef3/Ltbp1/Pdgfrb/Vim/Notch1/Eng/F2/Cst3/Il6ra/Rap1a/Cyp2j6/Fgfr3/Idua/Tgfb1/Itgb1/Il18 | 28 |
| BP | GO:0072091 | regulation of stem cell proliferation | 40/3915 | 99/17474 | 4,2944E-05 | 0,00071304 | 0,000531 | Sox17/Gli2/Ptprc/Prrx1/Pbx1/Irf6/Nr2e1/Kitl/Ace/Sox9/Zfp36l1/Gli3/Ptch1/Wnt5a/Fermt2/Bmp4/Fbln1/Rarg/Eif2ak2/Epcam/Nfatc1/Notch1/Mecom/Fgf2/Nfib/Cdkn2c/Ago3/Rbpj/Kdr/Tbx3/Lrp6/Tgfb1/Fgfr2/Vegfc/Atxn1l/Yap1/Cd109/Tbx18/Tgfbr2/Ctnnb1 | 40 |

| | | | | | | | | | |
|---|---|---|---|---|---|---|---|---|---|
| BP | GO:0006809 | nitric oxide biosynthetic process | 35/3915 | 83/17474 | 4,4343E-05 | 0,0007333 | 0,00054609 | Tlr5/Mtarc2/Itgb2/Igf1/Nos2/Clu/Tspo/Ddah2/Aif1/Ticam1/Cyp1b1/Jak2/Ass1/Ptgis/Tlr2/Ddah1/Klf4/Tlr4/Nos3/Tlr6/Pkd2/P2rx4/Cav1/Gimap5/Spr/Hrh1/Akt2/Insr/Slc7a2/Klf2/Agt/Icam1/Dnm2/Rora/Cx3cr1 | 35 |
| BP | GO:1903510 | mucopolysaccharide metabolic process | 35/3915 | 83/17474 | 4,4343E-05 | 0,0007333 | 0,00054609 | Dse/Stab2/Gns/Hexb/Itih3/Ednrb/Egflam/Angpt1/Has2/Abcc5/Itih2/Itih5/Cd44/Fgf2/Nfkb1/Slc35d1/Ugdh/Idua/Gusb/Ndnf/Tgfb1/Chsy1/Cemip/Lyve1/Xylt1/Spock3/Csgalnact1/Ednra/Slc10a7/Il15/Hexa/Hyal1/Chst7/Ids/Bgn | 35 |
| BP | GO:2000106 | regulation of leukocyte apoptotic process | 42/3915 | 106/17474 | 4,8276E-05 | 0,0007842 | 0,00058399 | Casp8/Bcl2/Fcgr2b/Fcer1g/Itpkb/Kitl/Adam17/Serpinb9/Mef2c/Wnt5a/Bmp4/Lgals3/Pnp/Ripk3/Il7r/Myc/St6gal1/Hcls1/Cd74/Dock8/Cd274/Gpam/Prkcq/Cd44/Bcl2l11/Mertk/Slc7a11/Efna1/Rorc/Bcl10/Lyn/P2rx7/Gimap8/Vhl/Cxcl12/Prkd2/Axl/Gas6/Nod2/Il18/Ccr5/Tsc22d3 | 42 |
| BP | GO:0055094 | response to lipoprotein particle | 17/3915 | 30/17474 | 4,9308E-05 | 0,00079766 | 0,00059402 | Fcer1g/Itgb2/Cd68/Syk/Hmgcs1/Trem2/Ticam1/Socs5/Npc1/Tlr4/Tlr6/Cd9/Apoe/Cd81/Itgb1/Ldlr/Myd88 | 17 |
| BP | GO:0150076 | neuroinflammatory response | 32/3915 | 74/17474 | 5,1398E-05 | 0,00082197 | 0,00061212 | Ifngr1/Adora2a/Igf1/Adcy1/Egfr/Grn/Clu/Cd200r1/Aif1/Trem2/Nr3c1/Csf1r/Jak2/Tlr2/Tlr4/C1qa/Tnfrsf1b/Tlr1/Tlr6/Smo/Bpgm/C5ar1/Tyrobp/Ctsc/Nupr1/Itgam/Tlr3/Plcg2/Agt/Itgb1/Ldlr/Cx3cr1 | 32 |
| BP | GO:0031099 | regeneration | 56/3915 | 154/17474 | 5,5066E-05 | 0,00086635 | 0,00064517 | Map4k4/Cflar/Erbb4/Bcl2/Tnr/Prrx1/Dusp10/Enpp1/Gja1/Igf1/Kremen1/Egfr/Vtn/Grn/Gfap/Ace/Adam17/Dicer1/Mtr/Ninj1/Dhfr/Plau/Mustn1/Lifr/Cdkn1a/Nrep/Jak2/Fas/Mdk/Capn3/Igsf10/Cers2/Ptgfrn/Spaar/Klf4/Tnc/Nfib/Hgf/Tec/Hopx/Ptn/Ezh2/Dysf/Cxcl12/Cd9/Cdkn1b/Tgfb1/Igf1r/Xylt1/Cd81/Nrg1/Mmp2/Yap1/Neo1/Lamb2/Flna | 56 |
| BP | GO:0035265 | organ growth | 71/3915 | 207/17474 | 5,6146E-05 | 0,00087067 | 0,00064839 | Col9a1/Erbb4/Bcl2/Ddr2/Kcnk2/Prox1/Lats1/Enpp1/Hey2/Gja1/Psap/Igf1/Meis1/Sox9/Sav1/Fbln5/Foxc1/Edn1/Mef2c/Bmpr1a/Myh6/Col14a1/Ppara/Arid2/Rarg/Heg1/Cxadr/Trip10/Fgf1/Nr3c1/Pdgfrb/Smad2/Tcf7l2/Notch1/Ttn/Thbs1/Bcl2l11/Ccm2l/Fgf2/Serp1/S1pr1/Camk2d/Pdlim5/Tgfbr1/Tmem38b/Col27a1/Bnc2/Lepr/Zmpste24/Trp73/Rbpj/Pdgfra/Kdr/Tgfbr3/Acacb/Ptpn11/Smo/Vgll4/Ankrd26/Ccnd2/Fgfr2/Igf2/Nrg1/Smad1/Agt/Yap1/Tgfbr2/Acvr2b/Ctnnb1/Sash3/G6pdx | 71 |
| BP | GO:0009791 | post-embryonic development | 46/3915 | 120/17474 | 5,7106E-05 | 0,00087789 | 0,00065376 | Bcl2/Enpp1/Sgpl1/Tmtc3/Lhx1/Ccdc47/Dicer1/Aldh5a1/Bmp4/Selenop/Heg1/Morc3/Hmgn1/Mmut/Vegfa/Smad2/Jak2/Tcf7l2/Alx4/Bcl2l11/Myt1/Mecom/Serp1/Tiparp/Tet2/Acadm/Aco1/Tgfbr1/Invs/Klf4/Trp73/Sema3c/Kdr/Flt1/Foxp2/Atg7/Atf5/Myo7a/Inppl1/Sox6/Fgfr2/Dhcr7/Myo1e/Acvr2b/Mecp2/Zfx | 46 |
| BP | GO:0051893 | regulation of focal adhesion assembly | 29/3915 | 65/17474 | 5,7129E-05 | 0,00087789 | 0,00065376 | Map4k4/Epb41l5/Myoc/Lims1/Poldip2/Dusp22/Fam107a/Vcl/Fermt2/Mmp14/Acvrl1/Ppm1f/Apod/Phldb2/Vegfa/Rock1/Tsc1/Sdc4/Tek/Macf1/Col16a1/Limch1/Kdr/Ptpn11/Iqgap1/Dlc1/Clasp2/Gpm6b/Arhgap6 | 29 |
| BP | GO:0090109 | regulation of cell-substrate junction assembly | 29/3915 | 65/17474 | 5,7129E-05 | 0,00087789 | 0,00065376 | Map4k4/Epb41l5/Myoc/Lims1/Poldip2/Dusp22/Fam107a/Vcl/Fermt2/Mmp14/Acvrl1/Ppm1f/Apod/Phldb2/Vegfa/Rock1/Tsc1/Sdc4/Tek/Macf1/Col16a1/Limch1/Kdr/Ptpn11/Iqgap1/Dlc1/Clasp2/Gpm6b/Arhgap6 | 29 |
| BP | GO:1990748 | cellular detoxification | 24/3915 | 50/17474 | 5,8469E-05 | 0,00089515 | 0,00066662 | Prdx6/Mtarc2/Sesn1/Nxn/Rdh11/Fbln5/Dhfr/Nnt/Aldh1a1/Nfe2l2/Cat/Selenot/Gstm5/Gstm7/Gstm1/Adh5/Prdx1/Nos3/Aldh2/Abcg2/Txnrd3/Mt3/Nqo1/Atp7a | 24 |
| BP | GO:0048738 | cardiac muscle tissue development | 85/3915 | 258/17474 | 5,8699E-05 | 0,000897 | 0,000668 | Casp8/Erbb4/Tnni1/Kcnk2/Prox1/Hey2/Pln/Gja1/Igf1/Frs2/Erbb3/Meis1/Sgcd/Gjc1/Id2/Sav1/Dicer1/Foxc1/Dsp/Edn1/Mef2c/Isl1/Wnt5a/Bmpr1a/Bmp4/Myh6/Angpt1/Col14a1/Ppara/Arid2/Myh11/Heg1/Cxadr/Vegfa/Trip10/Fgf1/Nr3c1/Pdgfrb/Smad4/Nrap/Nebl/Notch1/Tsc1/Eng/Ttn/Ccm2l/Bmp7/Fgf2/Gja5/S1pr1/Camk2d/Pdlim5/Acadm/Chd7/Tgfbr1/Zmpste24/Hspg2/Trp73/Rbpj/Sgcb/Pdgfra/Tgfbr3/Myo18b/Tbx3/Adamts9/Atg7/Vgll4/Ccnd2/Lrp6/Tgfb1/Sox6/Fgfr2/Nrg1/Ednra/Smad1/Agt/Itgb1/Yap1/Aldh1a2/Bmp5/Tbx18/Tgfbr2/Ctnnb1/Zic3/G6pdx | 85 |
| BP | GO:0071402 | cellular response to lipoprotein particle stimulus | 18/3915 | 33/17474 | 5,8926E-05 | 0,00089715 | 0,00066811 | Fcer1g/Itgb2/Cd68/Adam17/Syk/Hmgcs1/Trem2/Ticam1/Socs5/Npc1/Tlr4/Tlr6/Cd9/Apoe/Cd81/Itgb1/Ldlr/Myd88 | 18 |
| BP | GO:0061383 | trabecula morphogenesis | 26/3915 | 56/17474 | 5,9702E-05 | 0,00090397 | 0,00067319 | Slc40a1/Adgrg6/Enpp1/Hey2/Col1a1/Bmpr1a/Heg1/Adamts1/Vegfa/Nfatc1/Notch1/Eng/Ccm2l/Bmp7/S1pr1/Chd7/Tgfbr1/Tek/Fgfr3/Rbpj/Tgfbr3/Nrg1/Mmp2/Bmp5/Rhoa/Plxnb1 | 26 |
| BP | GO:0015849 | organic acid transport | 109/3915 | 348/17474 | 6,5572E-05 | 0,00098382 | 0,00073265 | Slc11a1/Acsl3/Septin2/Tnfrsf11a/Pla2g4a/Kcnj10/Eprs/Gja1/Fabp7/Psap/Slc16a9/Adora2a/Slc1a4/Slc22a4/Acsl6/Slc16a13/Slc13a5/Slc46a1/Nos2/Abcc3/Gfap/Ace/Slc38a6/Abcd4/Edn1/Syk/Sfxn1/Ntrk2/Slc38a9/Prkcd/Slc7a7/Abcc4/Slc1a3/Myc/Abcd2/Slc38a2/Abat/Abcc5/Casr/Atp5j/Grik1/Nr3c1/Atp8b1/Lrp5/Slc22a8/Slc22a6/Slc3a2/Slc16a12/Lrrc8a/Ptges/Pla2r1/Slc38a11/Slc1a2/Thbs1/Slc13a3/Fabp5/Slc7a11/Crabp2/Slc16a1/Slc16a4/Abcd3/Slc26a7/Plin2/Slc2a1/Mfsd2a/Pla2g5/Abcb1a/Abcb1b/Crot/Slc5a6/Lrrc8c/Pla2g1b/P2rx7/Abcg2/Sfxn5/Slc6a6/Slc6a11/Slc6a1/Slc25a18/Slc6a13/Slc6a12/Slco1c1/Slco1a4/Slc1a5/Apoe/Ceacam1/Akt2/Fxyd1/Slc7a10/Abcc6/Slco2b1/Ucp2/P2ry2/Stard10/Cln3/Slc25a15/Slc7a2/Slc27a1/Agt/Itgb1/Aqp9/Myo6/Rbp1/Slco2a1/Slc38a3/Slc26a6/Slc6a20a/Slc38a5/Slc16a2 | 109 |
| BP | GO:0097191 | extrinsic apoptotic signaling pathway | 79/3915 | 237/17474 | 6,7668E-05 | 0,00100612 | 0,00074926 | Sgk3/Eya1/Cflar/Casp8/Tmbim1/Sp100/Bok/Bcl2/Ptprc/Pea15a/Atf3/G0s2/Eya4/Fyn/Timp3/Igf1/Kitl/Erbb3/Tnfsf12/Gabarap/Rffl/Brca1/Siva1/Ripk1/Pik3r1/Bmp4/Lgals3/Bcl2l2/Tnfrsf10b/Dab2/Trps1/Deptor/Pak2/Vegfa/Birc6/Ctnna1/Dele1/Rela/Jak2/Fas/Tcf7l2/Casp7/Traf1/Itga6/Itgav/Spi1/Thbs1/Bcl2l11/Tnfsf10/Hipk1/Gclm/Bmpr1b/Bcl10/Casp8ap2/Tgfbr1/Mknk1/Tnfrsf1b/Hgf/Fgfr3/P2rx7/Gper1/Cav1/Casp2/Ret/Ltbr/Zfp110/Tgfb1/Stx4a/Pycard/Nrg1/Tlr3/Casp3/Agt/Yap1/Icam1/Il18/Ppp2r1b/Bcl2a1b/Srpx | 79 |
| BP | GO:0006066 | alcohol metabolic process | 102/3915 | 322/17474 | 6,8073E-05 | 0,00101032 | 0,00075239 | Idh1/Acadl/Cyp27a1/Sgpp2/Pla2g4a/Soat1/Itpkb/Lbr/Enpp1/Lss/Plek/Aldh3a2/Pmp22/Acadvl/Pctp/Fdxr/Sptssa/Dhrs7/Rdh11/Npc2/Sptlc2/Degs2/Bmp6/Pcbd2/Dhfr/Hmgcs1/Abhd4/Ephx2/Ednrb/Dab2/Aco2/Lima1/Lmf1/Cyp1b1/Npc1/Stard4/Fgf1/Lrp5/Plcb3/Aldh1a1/Myof/Cat/Pex2/Cyp7b1/Fgf2/P2ry1/Dpm3/Pmvk/Fmo5/Hmgcs2/Nfkb1/Mttp/Adh5/Abca1/Acer2/Lepr/Plpp3/Scp2/Ptafr/Ldlrap1/Dhrs3/Cyp51/Insig1/Plb1/Rest/Naaa/Grk3/Mvk/Gper1/Tpk1/Adcyap1r1/Retsat/Dysf/Spr/Lpcat3/Apoc1/Apoe/Lipe/Idh2/P2ry6/Inppl1/Hsd3b7/Dhcr7/Erlin2/Asah1/Msmo1/Mt3/Lcat/Plcg2/Ldlr/Aplp2/Slc37a2/Sc5d/Pts/Bmp5/Rbp1/Pth1r/Plcd1/Ebp/Nsdhl/Mecp2/G6pdx | 102 |
| BP | GO:2001057 | reactive nitrogen species metabolic process | 37/3915 | 91/17474 | 6,9859E-05 | 0,00103496 | 0,00077074 | Tlr5/Mtarc2/Itgb2/Igf1/Nos2/Tmem106a/Clu/Tspo/Ddah2/Aif1/Ticam1/Cyp1b1/Jak2/Ass1/Ptgis/Tlr2/Ddah1/Klf4/Tlr4/Nos3/Tlr6/Pkd2/P2rx4/Cav1/Gimap5/Npy/Spr/Hrh1/Akt2/Insr/Slc7a2/Klf2/Agt/Icam1/Dnm2/Rora/Cx3cr1 | 37 |
| BP | GO:0035455 | response to interferon-alpha | 16/3915 | 28/17474 | 7,1744E-05 | 0,00105719 | 0,00078729 | Ro60/Traf3ip3/Myc/Ifnar2/Ifnar1/Eif2ak2/Ifit3/Ifit1/Oas1a/Axl/Ifitm2/Ifitm1/Ifitm3/Gas6/Bst2/Plscr1 | 16 |
| BP | GO:0045216 | cell-cell junction organization | 71/3915 | 209/17474 | 7,929E-05 | 0,0011511 | 0,00085722 | Cdh20/Cdh19/F11r/Perp/Gja1/Lims1/Specc1l/Pmp22/Myo1c/Ramp2/Gjc1/Ace/Pecam1/Prkch/Numb/Dsp/Bmp6/F2r/Ocln/Marveld2/Vcl/Fermt2/Gjb2/Gjb6/Cldn10/Cdh9/Mtss1/Hdac7/Cldn5/Dlg1/Pak2/Heg1/Cxadr/Vegfa/Mpp7/Rock1/Lims2/Ctnna1/Csf1r/Tjp2/Ctnnd1/Kirrel/Rab13/Gja5/Hipk1/Rhoc/Pkn2/Tgfbr1/Mpdz/Patj/Cldn19/Hopx/Cav1/Cd9/Ceacam1/Tgfb1/Nphs1/Lsr/Tjp1/Ikbkb/Cdh5/Cdh1/Agt/Pard3/Itgb1/Esam/Rdx/Adam10/Rhoa/Ctnnb1/Flna | 71 |

| | | | | | | | | | |
|---|---|---|---|---|---|---|---|---|---|
| BP | GO:0046942 | carboxylic acid transport | 108/3915 | 346/17474 | 8,2441E-05 | 0,00118726 | 0,00088415 | Slc11a1/Acsl3/Septin2/Tnfrsf11a/Pla2g4a/Kcnj10/Eprs/Gja1/Fabp7/Psap/Slc16a9/Adora2a/Slc1a4/Slc22a4/Acsl6/Slc16a13/Slc13a5/Slc46a1/Nos2/Abcc3/Gfap/Ace/Slc38a6/Abcd4/Edn1/Syk/Sfxn1/Ntrk2/Slc38a9/Prkcd/Slc7a7/Abcc4/Slc1a3/Myc/Abcd2/Slc38a2/Abat/Abcc5/Casr/Atp5j/Grik1/Nr3c1/Atp8b1/Lrp5/Slc22a8/Slc22a6/Slc3a2/Slc16a12/Lrrc8a/Ptges/Pla2r1/Slc38a11/Slc1a2/Thbs1/Slc13a3/Fabp5/Slc7a11/Crabp2/Slc16a1/Slc16a4/Abcd3/Slc26a7/Plin2/Slc2a1/Mfsd2a/Pla2g5/Abcb1a/Abcb1b/Crot/Slc5a6/Lrrc8c/Pla2g1b/P2rx7/Abcg2/Sfxn5/Slc6a6/Slc6a11/Slc6a1/Slc25a18/Slc6a13/Slc6a12/Slco1c1/Slco1a4/Slc1a5/Apoe/Ceacam1/Akt2/Slc7a10/Abcc6/Slco2b1/Ucp2/P2ry2/Stard10/Cln3/Slc25a15/Slc7a2/Slc27a1/Agt/Itgb1/Aqp9/Myo6/Rbp1/Slco2a1/Slc38a3/Slc26a6/Slc6a20a/Slc38a5/Slc16a2 | 108 |
| BP | GO:0043583 | ear development | 80/3915 | 242/17474 | 8,3826E-05 | 0,0012051 | 0,00089744 | Eya1/Bcl2/Gli2/Prrx1/Vangl2/Prox1/Eya4/Enpp1/Hey2/Psap/Pcdh15/Igf1/Lrig3/Otx1/Mks1/Fzd2/Sox9/Ttc8/Trip11/Dicer1/Gli3/Bloc1s5/Edn1/Tifab/Cxcl14/Wnt5a/Bmp4/Gjb6/Nipbl/Fzd6/Cthrc1/Myc/Ift27/Triobp/Tbx1/H2-K1/H2-T23/Clic5/Zeb1/Atp8b1/Tshz1/Notch1/Bcl2l11/Jag1/Sdc4/Sox2/Fgf2/Frem2/Clrn1/Chd7/Anp32b/C1qb/Hes5/Insig1/Fgfr3/Cytl1/Rbpj/Rest/Igfbp7/Tbx3/Ptpn11/Cux1/Alms1/Gata2/Cecr2/Cdkn1b/Myo7a/Mapk3/Fgfr2/Ednra/Cdh1/Maf/Bmper/Adam10/Bmp5/Myo6/Tbx18/Tmie/Zic3/Pou3f4 | 80 |
| BP | GO:0060977 | coronary vasculature morphogenesis | 17/3915 | 31/17474 | 8,643E-05 | 0,00123823 | 0,00092211 | Hey2/Sgcd/Ace/Angpt1/Arid2/Tbx1/Vegfa/Fgf1/Pdgfrb/Notch1/Spred1/Fgf2/Tgfbr1/Tgfbr3/Fgfr2/Setd2/Ctnnb1 | 17 |
| BP | GO:0050870 | positive regulation of T cell activation | 78/3915 | 235/17474 | 8,7912E-05 | 0,00125078 | 0,00093146 | Il1rl2/Igfbp2/Sox13/Ptprc/Itpkb/Dusp10/Vsir/Icosl/Igf1/Kitl/Smarcc2/Havcr2/Irf1/Nlrp3/Lgals9/Smarce1/Stat5b/Lgals8/Gli3/Syk/Il6st/Adk/Pnp/Il7r/Lgals1/Arid2/Nckap1l/Tfrc/Cd86/Cd80/Runx1/Rasal3/H2-Ob/H2-Aa/H2-Eb1/Aif1/Socs5/Cd74/Dock8/Smarca2/Jak2/Cd274/Gpam/Prkcq/Dpp4/Mdk/Traf6/Cd59a/B2m/Actl6a/Cd1d1/Ptpn22/Vcam1/Lef1/Bcl10/Ripk2/Tnfrsf14/Prkcz/Rhoh/Ephb4/Hmgb1/Cav1/Gimap5/Tgfb1/Klhl25/Il4ra/Itgal/Pycard/Igf2/Cd81/Il15/Cbfb/Il18/Rhoa/Tgfbr2/Ccr2/Sash3/Il2rg | 78 |
| BP | GO:0140888 | interferon-mediated signaling pathway | 30/3915 | 70/17474 | 0,00010646 | 0,00148149 | 0,00110327 | Stat1/Ikbke/Ifngr1/Irgm1/Irf1/Igtp/Irgm2/Wnt5a/Parp14/Parp9/Ifnar2/Ifnar1/Sting1/Jak2/Nmi/Mavs/Samhd1/Isg15/Rbm47/Oas1a/Trim56/Lsm14a/Irf3/Ifitm2/Ifitm1/Ifitm3/Irf7/Nlrc5/Myd88/Irak1 | 30 |
| BP | GO:0150116 | regulation of cell-substrate junction organization | 30/3915 | 70/17474 | 0,00010646 | 0,00148149 | 0,00110327 | Map4k4/Epb41l5/Myoc/Lims1/Poldip2/Dusp22/Pik3r1/Fam107a/Vcl/Fermt2/Mmp14/Acvrl1/Ppm1f/Apod/Phldb2/Vegfa/Rock1/Tsc1/Sdc4/Tek/Macf1/Col16a1/Limch1/Kdr/Ptpn11/Iqgap1/Dlc1/Clasp2/Gpm6b/Arhgap6 | 30 |
| BP | GO:1905039 | carboxylic acid transmembrane transport | 51/3915 | 140/17474 | 0,00010983 | 0,00152584 | 0,00113629 | Slc11a1/Septin2/Kcnj10/Slc16a9/Slc1a4/Slc22a4/Acsl6/Slc46a1/Gfap/Slc38a6/Abcd4/Sfxn1/Slc38a9/Prkcd/Slc7a7/Slc1a3/Myc/Abcd2/Slc38a2/Abcc5/Slc3a2/Slc16a12/Lrrc8a/Slc38a11/Slc1a2/Thbs1/Slc13a3/Slc7a11/Slc16a1/Abcd3/Slc2a1/Abcb1a/Abcb1b/Slc5a6/Lrrc8c/Slc6a6/Slc25a18/Slc6a13/Slc1a5/Akt2/Slc7a10/Cln3/Slc25a15/Slc7a2/Slc27a1/Agt/Itgb1/Slc38a3/Slc6a20a/Slc38a5/Slc16a2 | 51 |
| BP | GO:0072337 | modified amino acid transport | 22/3915 | 46/17474 | 0,00012619 | 0,0017183 | 0,00127961 | Slc11a1/Slc16a9/Slc1a4/Slc22a5/Slc22a21/Slc22a4/Slc46a1/Slc38a9/Slc7a7/Slc38a2/Abcc5/Slc16a12/Slc7a11/Slc5a6/Slc6a13/Slc6a12/Cln3/Slc25a15/Slc7a2/Agt/Slc25a20/Slc6a20a | 22 |
| BP | GO:0046631 | alpha-beta T cell activation | 63/3915 | 183/17474 | 0,00012675 | 0,00171834 | 0,00127965 | Bcl2/Ptprc/Rc3h1/Itpkb/Psap/Vsir/Adora2a/Stat6/Dock2/Irf1/Nlrp3/Ncor1/Lgals9/Tmem98/Stat3/Cd300a/Gli3/Hfe/Syk/Ctsl/Wdfy4/Pnp/Gpr183/Myc/Nckap1l/Cd80/Runx1/Rasal3/Twsg1/Socs5/Tcirg1/Jak2/Cd274/Prkcq/Cd44/Cd1d1/Il6ra/Rorc/Ptpn22/Lef1/Ripk2/Rpl22/Tnfrsf14/Prkcz/Hmgb1/Gimap5/Loxl3/Cracr2a/Tgfb1/Klhl25/Il4ra/Cd81/Il15/Cbfb/Il18/Rora/Rhoa/Tgfbr2/Ccr2/Sash3/Elf4/Il2rg/Atp7a | 63 |
| BP | GO:0050868 | negative regulation of T cell activation | 48/3915 | 131/17474 | 0,00014659 | 0,00193855 | 0,00144364 | Rc3h1/Vsir/Adora2a/Havcr2/Irf1/Lgals9/Cd300a/Gli3/Hfe/Dusp22/Bmp4/Lgals3/Nckap1l/Dlg1/Cd86/Cd80/Runx1/H2-Aa/Twsg1/Socs5/Cd74/Cd274/Nrarp/Mdk/Cd44/Sdc4/Pag1/Tnfaip8l2/Ptpn22/Pde5a/Laptm5/Pla2g5/Tnfrsf14/Fgl2/Hspb1/Hmgb1/Gimap5/Gpnmb/Loxl3/Ptpn6/Lag3/Ceacam1/Tgfb1/Cd37/Lrrc32/Il4ra/Casp3/Cbfb | 48 |
| BP | GO:0009081 | branched-chain amino acid metabolic process | 14/3915 | 24/17474 | 0,00015138 | 0,00198909 | 0,00148128 | Hibch/Ilvbl/Aldh6a1/Mccc2/Ivd/Dbt/Hmgcl/Sdsl/Hibadh/Bckdha/Bcat2/Acadsb/Acad8/Hsd17b10 | 14 |
| BP | GO:0198738 | cell-cell signaling by wnt | 128/3915 | 428/17474 | 0,00016041 | 0,0020813 | 0,00154995 | Sox17/Ccnyl1/Sox13/Aspm/Myoc/Vangl2/Lats1/Enpp1/Bicc1/Wif1/Ddit3/Kremen1/Znrf3/Grb10/Egfr/Tmem88/Nxn/Rnf43/Mks1/Col1a1/Fzd2/Wnt3/Axin2/Sox9/Rnf213/Notum/Sdc1/Hbp1/Daam1/Nid1/Gli3/Sfrp4/Edn1/Zbed3/Map3k1/Isl1/Wnt5a/Fermt2/Ndrg2/Amer2/Sox7/Scel/Ednrb/Gpc5/Dab2/Sema5a/Ubr5/Fzd6/Cthrc1/Myc/Lmbr1l/Daam2/Ppm1b/Mcc/Apcdd1/Mbd2/Lrp5/Tle4/Kank1/Tcf7l2/Grk5/Nrarp/Notch1/Lypd6/Ctnnd1/Lrp4/Mdk/Cd44/Lgr4/Usp8/Csnk2a1/Sox2/Fgf2/Jade1/Foxo1/Wwtr1/Tlr2/Pbxip1/Lef1/Dkk2/Nfkb1/Wls/Reck/Invs/Klf4/Plpp3/Macf1/Rspo1/Hdac1/Ctnnbip1/Fgfr3/Rbpj/Pkd2/Mdfic/Cav1/Tcf7l1/Klf15/Vgll4/Wnk1/Lrp6/Apoe/Tgfb1/Fuz/Lrrk1/Fzd4/Ilk/Gprc5b/Tgfb1i1/Fgfr2/Cpe/Ednra/Nkd1/Sall1/Cdh1/Yap1/Amotl1/Tbx18/Ppp2r3a/Ryk/Nphp3/Rbms3/Ctnnb1/Limd1/Ndp/Xiap/Gpc3/Amer1/Med12 | 128 |
| BP | GO:2000736 | regulation of stem cell differentiation | 33/3915 | 81/17474 | 0,00016103 | 0,00208278 | 0,00155104 | Sox17/Vsir/Mbd3/Chd3/Stat3/Sox9/Dicer1/Foxc1/Prkdc/Lbh/Eif2ak2/Mta3/Zfp36l2/Notch1/Nfe2l2/Jag1/Fgf2/Hdac1/Hes5/Cdk6/Pus7/Pdgfra/Rest/Tbx3/Ptn/Ezh2/Kdm3a/Sox5/Zfp36/Tead2/Sox6/Gatad2a/Yap1 | 33 |
| BP | GO:0032964 | collagen biosynthetic process | 27/3915 | 62/17474 | 0,00016785 | 0,00215081 | 0,00160171 | Tram2/Fn1/Suco/Ddr2/Ccn2/Adora2b/Serpinf2/Col1a1/P3h4/Dicer1/F2r/Bmp4/Rgcc/Rapgef3/Ltbp1/Pdgfrb/Vim/Notch1/Eng/F2/Il6ra/Rap1a/Cyp2j6/Tgfb1/Rcn3/Serpinh1/Il18 | 27 |
| BP | GO:0008203 | cholesterol metabolic process | 48/3915 | 132/17474 | 0,00018089 | 0,00229235 | 0,00170711 | Acadl/Cyp27a1/Soat1/Lbr/Lss/Pmp22/Acadvl/Pctp/Fdxr/Npc2/Hmgcs1/Ephx2/Lima1/Lmf1/Npc1/Stard4/Fgf1/Lrp5/Cat/Pex2/Cyp7b1/Pmvk/Fmo5/Hmgcs2/Mttp/Abca1/Lepr/Scp2/Ldlrap1/Cyp51/Insig1/Mvk/Lpcat3/Apoc1/Apoe/Lipe/Hsd3b7/Dhcr7/Erlin2/Msmo1/Mt3/Lcat/Ldlr/Aplp2/Sc5d/Ebp/Nsdhl/G6pdx | 48 |
| BP | GO:0038084 | vascular endothelial growth factor signaling pathway | 20/3915 | 41/17474 | 0,00018091 | 0,00229235 | 0,00170711 | Nrp2/Cd63/Flt4/Myo1c/Prkd1/Foxc1/Vegfa/Xdh/Jcad/Sema6a/Pdgfrb/Tcf4/Vegfb/Pik3ca/Pdgfra/Kdr/Hspb1/Prkd2/Vegfc/Gab1 | 20 |
| BP | GO:0090287 | regulation of cellular response to growth factor stimulus | 97/3915 | 311/17474 | 0,00019042 | 0,00237361 | 0,00176763 | Tfap2b/Cflar/Lats1/Vsir/Tet1/Pbld2/Cd63/Grb10/Sap30l/Myo1c/Smurf2/Adam17/Snx6/Sfrp4/Aspn/Fst/Il17rd/Wnt5a/Mmrn2/Bmp4/Dab2/Adamts12/Angpt1/Ppara/Acvrl1/Vasn/Nrros/Fstl1/Tmem204/Vegfa/Twsg1/Xdh/Ltbp1/Nrxn1/Jcad/Zeb1/Nrep/Fgf1/Sema6a/Lox/Tcf4/Smad4/Smad2/Vegfb/Fgfbp3/Myof/Tcf7l2/Notch1/Eng/Dlx1/Cd59a/Spred1/Thbs1/Slc2a10/Dok5/Fgf2/Notch2/Npnt/Sh3glb1/Ccn1/Tek/Hdac1/Prdm16/Hes5/Elapor2/Htra3/Fgfbp1/Rbpj/Kdr/Tgfbr3/Cav2/Cav1/Hipk2/Prkd2/Tgfb1/Fuz/Cyfip1/Fzd4/Ilk/Tgfb1i1/Htra1/Vegfc/Hhip/Mt3/Cdh5/Agt/Dnm2/Bmper/Neo1/Nedd4/Cd109/Ctnnb1/Apln/Gpc3/Dmd/Ogt/Chrdl1 | 97 |
| BP | GO:0060415 | muscle tissue morphogenesis | 33/3915 | 82/17474 | 0,00021211 | 0,00262817 | 0,0019572 | Col3a1/Tnni1/Vangl2/Prox1/Hey2/Fzd2/Foxc1/Dsp/Isl1/Wnt5a/Bmpr1a/Myh6/Angpt1/Tbx1/Heg1/Mylk/Smad4/Efemp2/Notch1/Eng/Ttn/Ccm2l/S1pr1/Chd7/Tgfbr1/Rbpj/Tgfbr3/Lrp6/Tgfb1/Fgfr2/Nrg1/Ednra/Ctnnb1 | 33 |

| | | | | | | | | | |
|---|---|---|---|---|---|---|---|---|---|
| BP | GO:0006644 | phospholipid metabolic process | 107/3915 | 350/17474 | 0,00021679 | 0,00266213 | 0,00198249 | Ormdl1/Pgap1/Idh1/Erbb4/Plcd4/Acsl3/Inpp5d/Dbi/Pla2g4a/Prdx6/Pigc/Pigm/Itpkb/Capn2/Smpd2/Smpdl3a/Chpt1/Dgka/Plek/Acsl6/Alox8/Pld2/Pitpnc1/Abca8a/Lpin1/Mboat2/Pik3cg/Pigh/Sptlc2/Aspg/Idi1/Gpld1/Mboat1/Ptdss1/Lpcat1/Serinc5/Hexb/Pik3r1/Hmgcs1/Prkcd/Socs4/Abhd4/Smpd5/Pcyt1a/Adgrf5/Pla2g7/Socs5/Abhd3/Pdgfrb/Csf1r/Plcb3/Plaat3/Sgms1/Gpam/Pip4k2a/Pnpla7/Agpat2/Pip5kl1/Ptpmt1/Nr1h3/Plcb2/Pla2g4e/Gpcpd1/Serinc3/Fabp5/Pld1/Pik3ca/Dnajc19/Fgf2/Dpm3/Pmvk/Hmgcs2/Sgms2/Sh3glb1/Tmem38b/Plpp3/Scp2/Pik3r3/Mfsd2a/Pla2g5/Hadha/Plb1/Pi4k2b/Naaa/Mvk/Pla2g1b/Pyurf/Hdhd5/Lpcat3/Apoc1/Inppl1/Pgap2/Pik3c2a/Cln3/Plpp4/Agpat5/Slc27a1/Inpp4b/Lpcat2/Lcat/Plcg2/Ldlr/Plscr1/Plcd1/Mtm1/Mecp2/Pcyt1b | 107 |
| BP | GO:1900015 | regulation of cytokine production involved in inflammatory response | 25/3915 | 57/17474 | 0,00025272 | 0,00301802 | 0,00224752 | Appl2/Nos2/Stat3/Pld4/Ppara/Abcd2/Nlrc3/Apod/Trem2/Ticam1/Ticam2/Card9/F2/Gbp5/Tlr4/Tlr6/Ezh2/Il17rc/Alox5/Il17ra/Clec7a/Pycard/Adcy7/Nod2/Myd88 | 25 |
| BP | GO:0010463 | mesenchymal cell proliferation | 24/3915 | 54/17474 | 0,00026152 | 0,00309178 | 0,00230245 | Irs1/Prrx1/Sox9/Arhgap5/Wnt5a/Bmpr1a/Bmp4/Myc/Tbx1/Zeb1/Bmp7/Nfib/Ctnnbip1/Kdr/Hmgb1/Foxp2/Smo/Ptn/Lrp6/Fgfr2/Tbx18/Tgfbr2/Ctnnb1/Gpc3 | 24 |
| BP | GO:0030900 | forebrain development | 118/3915 | 394/17474 | 0,00026622 | 0,00313235 | 0,00233266 | Efhc1/Col3a1/Pgap1/Nrp2/Erbb4/Gli2/Dbi/Srgap2/Btg2/Aspm/Tnr/Atp1a2/Prox1/Fyn/Nr2e1/Fabp7/Rfx4/Frs2/Ikzf1/Egfr/Otx1/Pex13/Ncor1/Atp1b2/Lhx1/Id2/Syne2/Numb/Ttc8/Dicer1/Gli3/Tubb2b/Phactr1/Id4/Secisbp2/Ntrk2/Slc6a3/Isl1/Wnt5a/Bmpr1a/Bmp4/Zic5/Sema5a/Sun2/Nde1/Crkl/D16Ertd472e/Twsg1/Csf1r/Sall3/Tcf7l2/Emx2/Notch1/Tsc1/Lhx2/Dlx1/Ccdc141/Mdk/Slc1a2/Pax6/B2m/Btbd3/Mecom/Sox2/Fgf2/Slc7a11/P2ry12/Bcan/Nhlh2/Lef1/Chd7/Pou3f2/Akna/Nfib/Dock7/Dab1/Lrp8/Mfsd2a/Hdac1/Trp73/Hes5/Reln/Tacc3/Fgfr3/Rbpj/Tbx3/Foxp2/Smo/Ndnf/Gata2/Atg7/Cxcl12/Lrp6/Axl/Tyrobp/Atf5/Itgam/Fgfr2/Tacc1/Hook3/Fut10/Nrg1/Dlc1/Pcm1/Sall1/Bbs2/Adgrg1/Aplp2/Aldh1a2/Zic1/Ryk/Rhoa/Setd2/Ctnnb1/Zic3/Fgf13/Atp7a/Pou3f4 | 118 |
| BP | GO:0010939 | regulation of necrotic cell death | 19/3915 | 39/17474 | 0,00026685 | 0,00313235 | 0,00233266 | Cflar/Casp8/Bok/Ddr2/Ripk1/Ripk3/Tspo/Casp6/Map3k7/Mutyh/Cav1/Casp2/Slc6a6/Slc6a13/Ndufc2/Nupr1/Asah1/Mt3/Birc3 | 19 |
| BP | GO:0006691 | leukotriene metabolic process | 14/3915 | 25/17474 | 0,00027322 | 0,00314024 | 0,00233854 | Pla2g4a/Ggt5/Lta4h/Ltc4s/Syk/Cyp4f15/Cyp4f14/Cyp4f13/Tlr2/Tlr4/Pla2g5/Ncf1/Alox5ap/Alox5 | 14 |
| BP | GO:0048010 | vascular endothelial growth factor receptor signaling pathway | 22/3915 | 48/17474 | 0,00027323 | 0,00314024 | 0,00233854 | Grb10/Flt4/Prkd1/Clec14a/Foxc1/Mmrn2/Bmp4/Angpt1/Tmem204/Vegfa/Vegfb/Myof/Cd59a/Tek/Kdr/Hspb1/Flt1/Prkd2/Fzd4/Vegfc/Mt3/Nedd4 | 22 |
| BP | GO:0140241 | translation at synapse | 22/3915 | 48/17474 | 0,00027323 | 0,00314024 | 0,00233854 | Rpl7/Rpl37a/Eef2/Rpl26/Rpl23a/Rps24/Rpl37/Rpl8/Rps28/Rps14/Rpl7a/Rpl35/Rpl22/Rpl6/Rpl32/Rps5/Rpl13a/Rplp2/Rpl4/Rpl29/Rpl14/Rpl10 | 22 |
| BP | GO:0140242 | translation at postsynapse | 22/3915 | 48/17474 | 0,00027323 | 0,00314024 | 0,00233854 | Rpl7/Rpl37a/Eef2/Rpl26/Rpl23a/Rps24/Rpl37/Rpl8/Rps28/Rps14/Rpl7a/Rpl35/Rpl22/Rpl6/Rpl32/Rps5/Rpl13a/Rplp2/Rpl4/Rpl29/Rpl14/Rpl10 | 22 |
| BP | GO:0051098 | regulation of binding | 120/3915 | 402/17474 | 0,00027442 | 0,00314958 | 0,00234549 | Plcl1/Map2/Dnajb2/Sp100/Hjurp/Ralb/Epb41l5/Mndal/Ifi203/Traf3ip3/Hey2/Pln/Igf1/Ddit3/Tns3/Rack1/Atp2a3/Vtn/Poldip2/Lgals9/Ramp2/Ace/Id2/Hfe/Foxc1/Isl1/Wnt5a/Bmp4/Hmbox1/Rb1/Cln5/Dzip1/Dab2/Cthrc1/Angpt1/Myc/Mfng/H1f0/Ppara/Rapgef3/Tmbim6/Dazap2/Tfap4/Cldn5/Parp9/Cdkn1a/Kat2b/Ticam1/Rock1/Sting1/Lox/Adrb2/Mbd2/Smad4/Smad2/Jak2/Ide/Tcf7l2/Fbh1/Lhx2/Traf6/Pax6/B2m/Ddrgk1/Mavs/Id1/Ccm2l/Mapre1/Cpne1/Ctsz/P2ry1/Nes/Snapin/Hipk1/1810037I17Rik/Lef1/Ripk2/Tgfbr1/Klf4/Tlr4/Nfib/Zmpste24/Epb41/Ldlrap1/Ctnnbip1/Sri/Hopx/Rest/Tgfbr3/Hip1r/Gper1/Lfng/Hmgb1/Cav1/Smo/Cald1/Hipk2/Dysf/Camk1/Apoe/Plaur/Tgfb1/Sirt2/Rsf1/Arrb1/Trim21/Plk1/Mapk3/Ctbp2/Aktip/Map1lc3b/Aplp2/Ubash3b/Anxa2/Plscr1/Atr/Nphp3/Ctnnb1/Ldoc1/Mecp2 | 120 |
| BP | GO:0062012 | regulation of small molecule metabolic process | 108/3915 | 356/17474 | 0,00028019 | 0,00320243 | 0,00238485 | Acadl/Irs1/Dbi/Adipor1/Pla2g4a/Fmo1/Fmo2/Prox1/Hsd11b1/Appl2/Igf1/Plek/Ppp4r3b/Adora2b/Ncor1/Acadvl/Nos2/Stat3/Brca1/Sox9/C1qtnf1/Gpld1/Ppp1r3g/Bmp6/Prxl2c/Nln/Slc7a7/Ephx2/Lacc1/Clybl/Dab2/Atpsckmt/Myc/Gpt/Tspo/Ppara/Abcd2/Zbtb20/Lmf1/Ier3/Guca1b/Trem2/Kat2b/Stard4/Fgf1/Nr3c1/Cd74/Me2/Cpt1a/Sorbs1/Entpd1/Tcf7l2/Nr1h3/Pex2/Fabp5/Slc7a11/Foxo1/P2ry1/Rorc/Fmo5/Nfkb1/Acadm/Cyp2j6/Lepr/Scp2/Zmpste24/Mfsd2a/Ptafr/Nos3/Insig1/Ppargc1a/Rest/Slc4a4/Acacb/P2rx7/Mlxipl/Gper1/Hmgb1/Pdk4/Cav1/Fam3c/Adcyap1r1/Ankrd26/Lpcat3/Apoc1/Apoe/Ceacam1/Akt2/Klhl25/Ndufc2/P2ry6/Cln3/Nupr1/Igf2/Dhcr7/Insr/Erlin2/Gpt2/Agt/Ldlr/Rora/Bmp5/Elovl5/Me1/Pth1r/Plcd1/Mid1ip1/Ogt | 108 |
| BP | GO:0070265 | necrotic cell death | 27/3915 | 64/17474 | 0,00031587 | 0,00354416 | 0,00263933 | Cflar/Casp8/Bok/Ddr2/Pygl/Ripk1/Ninj1/Ripk3/Tspo/Fas/Trpm7/Casp6/Map3k7/Mutyh/Cav1/Casp2/Slc6a6/Slc6a13/Irf3/Ndufc2/Nupr1/Asah1/Tlr3/Mt3/Mlkl/Tmem123/Birc3 | 27 |
| BP | GO:0006790 | sulfur compound metabolic process | 88/3915 | 281/17474 | 0,00031696 | 0,00354416 | 0,00263933 | Acsl3/Fmo1/Vangl2/Enpp1/Dse/Amd1/Mical1/Ggt1/Ggt5/Gns/Cs/Suox/Acsl6/Hs3st3b1/Hs3st3a1/Mettl16/Blmh/Acaca/Stat5b/Arsg/Sgsh/Adi1/Mtr/Hexb/Hmgcs1/Btd/Ednrb/Ghr/Egflam/Angpt1/Tst/Mpst/Csad/Sp1/Ehhadh/Nubp2/Cbs/Mmut/Xdh/Cdo1/Hsd17b4/Acaa2/Papss2/Mms19/Gpam/Tcf7l2/Sqor/Acss1/Acss2/Slc7a11/Pmvk/Hmgcs2/Phgdh/Ahcyl1/Gstm1/Gclm/Papss1/Cth/Slc35d1/Hmgcl/Ugdh/Idua/Acacb/Mvk/Gusb/Pdk4/Ahcyl2/Tpk1/Ndnf/Suclg2/Mcee/Chsy1/Xylt1/Sult1a1/Acadsb/Spock3/Csgalnact1/Ednra/Slc10a7/Gcdh/Nudt7/Hexa/Ciao2a/Hyal1/Ctnnb1/Chst7/Ids/Bgn | 88 |
| BP | GO:0048638 | regulation of developmental growth | 117/3915 | 392/17474 | 0,00032662 | 0,00363239 | 0,00270504 | Map2/Erbb4/Fn1/Vil1/Bcl2/Tnr/Dusp10/Kcnk2/Prox1/Lats1/Hey2/Gja1/Sgpl1/Igf1/Rab21/Ikzf1/Sh3pxd2b/Ncor1/Spag9/Stat5b/Stat3/Rnd2/Grn/Wnt3/Sav1/Lgmn/Foxc1/Edn1/Ptch1/Slc6a3/Mef2c/Wnt5a/Sema3g/Colq/Bmpr1a/Bmp4/Myh6/Ghr/Nipbl/Sema5a/Col14a1/Ppara/Dip2b/Prkdc/Dlg1/Cxadr/Ezr/Cdkn1a/Vegfa/Trip10/Nr3c1/Sema6a/Adrb2/Tll2/Gpam/Tcf7l2/Notch1/Capn3/Sema6d/Bcl2l11/Pik3ca/Fgf2/Serp1/Crabp2/Csf1/Sh3glb1/Chd7/Pou3f2/Spaar/Tgfbr1/Zmpste24/Mfsd2a/Macf1/Trp73/Sema3d/Sema3c/Fgfr3/Rbpj/Hopx/Tgfbr3/Acacb/Smo/Spr/Alms1/Mgll/Vgll4/Cxcl12/Ankrd26/Ccnd2/Cdkn1b/Apoe/Cyfip1/Igf1r/Picalm/Ilk/Fgfr2/Igf2/Insr/Nrg1/Nkd1/Bbs2/Mt3/Cdh1/Agt/Yap1/Dnm2/Rpl4/Ryk/Clasp2/Tgfbr2/Ccr5/Sash3/Fgf13/Mtm1/Arhgap4/Mecp2/G6pdx | 117 |
| BP | GO:0043254 | regulation of protein-containing complex assembly | 125/3915 | 424/17474 | 0,00036222 | 0,00397483 | 0,00296005 | Map2/Vil1/Hjurp/Ralb/Ikbke/Lmod1/Nek7/Aida/Lats1/Arhgap18/Sar1a/Ikzf1/Plek/Rack1/Irgm1/Igtp/Irgm2/Flii/Xaf1/Myo1c/Pecam1/Sox9/H3f3b/Stxbp6/Prkd1/Carmil1/Syk/Map3k1/Isl1/Prkcd/Fermt2/Lgals3/Clu/Lcp1/Rictor/Baiap2l2/Twf1/Nckap1l/Nlrc3/Spidr/Dlg1/Tfrc/Hcls1/Snx9/Kank3/Vegfa/Trem2/Daam2/Ticam1/Eif2ak2/Svil/Mpp7/Riok3/Kank1/Fas/Add3/Slc39a12/Ckap5/Bmf/Bcl2l11/Mavs/Hck/Mapre1/P2ry12/Fnip2/Tlr2/Kirrel/Ptpn22/Rhoc/Capza1/Gbp5/Sh3glb1/Map3k7/Abca1/Tlr4/Cdk5rap2/Tal1/Ctnnbip1/Prkcz/Isg15/Akap9/Tlr6/Hrk/Ptpn11/Hip1r/Eln/Arpc1b/Hmgb1/Cav1/Arhgef5/Capg/Cdkn1b/Eps8/Vasp/Apoe/Tgfb1/Plekhg2/Sirt2/Nphs1/Ankrd27/Rpl13a/Svip/Cyfip1/Fes/Fchsd2/Trim30a/Pycard/Eif4ebp1/Nrg1/Cdh5/Lcat/Plcg2/Icam1/Tirap/Esam/Rdx/Tmod3/Atr/Rhoa/Clasp2/Myd88/Lamp2/Mecp2/Msn/Btk | 125 |
| BP | GO:0035458 | cellular response to interferon-beta | 22/3915 | 49/17474 | 0,00039179 | 0,00425961 | 0,00317213 | Stat1/Aim2/Mndal/Ifi203/Traf3ip3/Irgm1/9930111J21Rik1/Ifi47/Irf1/Igtp/Irgm2/Sting1/Iigp1/Camk2a/Ifit3/Ifit1/Mavs/Gbp7/Gbp3/Gbp2/Gbp6/Oas1a | 22 |
| BP | GO:1900026 | positive regulation of substrate adhesion-dependent cell spreading | 20/3915 | 43/17474 | 0,00040381 | 0,00433209 | 0,00322611 | Myoc/Lims1/Itgb3/P4hb/Carmil1/Nedd9/Fermt2/Dock5/Dab2/Has2/Triobp/Crkl/Lims2/Mdk/Cib1/Ilk/Dock1/Dnm2/Cspg5/Flna | 20 |

| | | | | | | | | | |
|---|---|---|---|---|---|---|---|---|---|
| BP | GO:0015748 | organophosphate ester transport | 44/3915 | 122/17474 | 0,00040423 | 0,00433209 | 0,00322611 | Dbi/Gja1/Pctp/Pitpnc1/Prkcd/Xkr6/Abcc4/Ano6/Abcc5/Slc35b2/Slc25a23/Atp8b1/Prelid3a/Slc66a2/Lrrc8a/Pltp/Atp11b/Slc25a24/Abca4/Mttp/Abca1/Scp2/Mfsd2a/Xkr8/Slc25a33/Abcb1a/Abcb1b/Atp8a1/Atp10d/Scarb2/Lrrc8c/P2rx7/Apoc1/Apoe/Abcc6/Atp10a/Atp11a/Ldlr/Slc37a2/Plscr1/Plscr2/Plscr4/Slc25a36/Atp11c | 44 |
| BP | GO:0010977 | negative regulation of neuron projection development | 55/3915 | 161/17474 | 0,00040871 | 0,00436877 | 0,00325342 | Map4k4/Map2/Neu4/Rab29/Tnr/Katna1/Zfp365/Kremen1/Efemp1/Pmp22/Rap1gap2/Gfap/Wnt3/Adam17/Wnt5a/Sema3g/Dab2/Sema5a/Lgals1/Dip2b/H2-K1/H2-D1/Sema6a/Kank1/Il15ra/Vim/Tsc1/Lrp4/B2m/Sema6d/Ctsz/Efna1/Snapin/Cers2/Dab1/Sema3d/Sema3c/Cd38/Ptprz1/Apoe/Cib1/Inppl1/Dennd5a/Xylt1/Mt3/Cdh1/Ntm/Neo1/Ryk/Rhoa/Ccr5/Thoc2/Fgf13/Arhgap4/Flna | 55 |
| BP | GO:0060021 | roof of mouth development | 37/3915 | 98/17474 | 0,00041239 | 0,0043874 | 0,0032673 | Prrx1/Irf6/Sgpl1/Gdf11/Fzd2/Itgb8/Gli3/Foxf2/Mef2c/Wnt5a/Bmpr1a/Tbx1/Dlg1/Col11a2/Smad4/Smad2/Tshz1/Alx4/Tiparp/Lef1/Chd7/Asph/Anp32b/Tgfbr1/Bnc2/Dhrs3/Prdm16/Insig1/Pdgfra/Tgfbr3/Tbx3/Loxl3/Lrp6/Fuz/Lrrc32/Tgfbr2/Acvr2b | 37 |
| BP | GO:0032060 | bleb assembly | 9/3915 | 13/17474 | 0,00041258 | 0,0043874 | 0,0032673 | Prdx6/Pmp22/Ano6/Emp2/Mylk/Rock1/P2rx7/Emp1/Emp3 | 9 |
| BP | GO:0030048 | actin filament-based movement | 45/3915 | 126/17474 | 0,00044762 | 0,0047478 | 0,00353569 | Myo1b/Vil1/Kcne4/Epb41l5/Atp1a2/Pln/Ctnna3/Myo1g/Sgcd/Myo1c/Myo19/Kcnj2/Syne2/Dsp/Myh6/Sun2/Emp2/Dlg1/Myo1f/Rock1/Dsc2/Acta2/Wipf1/Pik3ca/Gja5/Casq2/Fnbp1l/Camk2d/Wasf2/Pdpn/Akap9/Sri/Limch1/Cav1/Abcc9/Fxyd1/Myo7a/Parva/Pard3/Myo1e/Myo6/Was/Fgf13/Flna/Kcne1l | 45 |
| BP | GO:0050792 | regulation of viral process | 59/3915 | 176/17474 | 0,00044948 | 0,00476139 | 0,00354581 | Stat1/Sp100/Bcl2/Mndal/Ifi203/Prox1/Bsg/Trim25/P4hb/Dicer1/Map3k1/Tasor/Lgals1/Apobec3/Fbln1/Ppara/Tmem39a/Trim26/Eif2ak2/Atg12/Cd74/Fam111a/Notch1/Stom/Ifih1/Mavs/Znfx1/Vapb/Gbp7/Gbp3/Pkn2/Isg15/Oasl2/Oasl1/Oas1a/Trim56/Hmgb1/Mdfic/Zc3hav1/Resf1/Nectin2/Axl/Zfp36/Rsf1/Trim21/Trim68/Trim34a/Trim5/Trim12a/Trim12c/Trim30a/Ifitm2/Ifitm1/Ifitm3/Bst2/Hacd3/Plscr1/Plscr2/Mid2 | 59 |
| BP | GO:0009100 | glycoprotein metabolic process | 106/3915 | 353/17474 | 0,00047812 | 0,00503251 | 0,00374771 | B3gat2/Uggt1/Chst10/Poglut2/Neu4/St8sia4/Bcl2/Mgat5/Soat1/Vangl2/Dse/Tet1/Igf1/Galnt4/Tmtc3/Tmtc2/Rxylt1/Gfpt2/Galnt10/Hs3st3b1/Hs3st3a1/Tmem106a/St6galnac2/Sgsh/Fut8/Entpd5/Pomt2/Trip11/Ctsl/Hexb/Ngly1/Mustn1/Galnt15/Egflam/Adamts12/Pmm1/A4galt/Gxylt1/Pmm2/B3gnt5/St6gal1/St3gal6/Lmf1/Galnt14/Npc1/Galnt1/Hbegf/Vegfb/Gcnt1/Tcf7l2/St6galnac6/Ganc/Cst3/Pofut1/Edem2/Rpn2/Slc2a10/Serp1/Dpm3/Tet2/Manba/Slc39a8/St6galnac3/Dpy19l4/Fut9/Erp44/Acer2/Slc35d1/Fbxo2/Ost4/Cytl1/Ugdh/Idua/Gusb/Plod3/Gal3st4/C1galt1/Ndnf/Gfpt1/Eogt/Edem1/Pcsk6/Chsy1/St8sia2/Xylt1/Insr/Fut10/Aga/Spock3/Csgalnact1/Il15/Mt3/St3gal4/Stt3a/Poglut3/Hexa/Hyal1/Ctnnb1/Chst7/Ids/Bgn/Ogt/Magt1/Atp7a/Itm2a/Btk | 106 |
| BP | GO:0035924 | cellular response to vascular endothelial growth factor stimulus | 25/3915 | 59/17474 | 0,00048005 | 0,00503991 | 0,00375322 | Nrp2/Cd63/Flt4/Myo1c/Ramp2/Prkd1/Foxc1/Adamts12/Vegfa/Xdh/Jcad/Sema6a/Pdgfrb/Tcf4/Rela/Vegfb/Notch1/Pik3ca/Pdgfra/Kdr/Hspb1/Flt1/Prkd2/Vegfc/Gab1 | 25 |
| BP | GO:0006027 | glycosaminoglycan catabolic process | 15/3915 | 29/17474 | 0,00052303 | 0,00540184 | 0,00402275 | Stab2/Gns/Sgsh/Hexb/Cd44/Fgf2/Idua/Gusb/Pglyrp1/Tgfb1/Cemip/Lyve1/Hexa/Hyal1/Ids | 15 |
| BP | GO:0043550 | regulation of lipid kinase activity | 23/3915 | 53/17474 | 0,0005307 | 0,00547421 | 0,00407665 | Irs1/Ppp2r5a/Pik3ip1/Pik3r5/Pik3r1/Rb1/Pdgfrb/Pip4k2a/F2/Fgf2/Vav3/Sh3glb1/Lyn/Tek/Pik3r3/Fgfr3/Pdgfra/Flt1/Wdr91/Tgfb1/Cd81/Nod2/Rbl2 | 23 |
| BP | GO:0005977 | glycogen metabolic process | 31/3915 | 79/17474 | 0,00054742 | 0,00557688 | 0,00415311 | Irs1/Enpp1/Igf1/Grb10/Ugp2/Pygl/Ppp1r3g/Il6st/Pfkm/Gbe1/Pygm/Ppp1r3c/Sorbs1/Tcf7l2/Pygb/Ppp1r3d/Agl/Acadm/Lepr/Khk/Pgm2/Phkg1/Hmgb1/Gfpt1/Akt2/Gys1/Igf2/Insr/Ppp1r3b/Phkb/Phka1 | 31 |
| BP | GO:0006073 | cellular glucan metabolic process | 31/3915 | 79/17474 | 0,00054742 | 0,00557688 | 0,00415311 | Irs1/Enpp1/Igf1/Grb10/Ugp2/Pygl/Ppp1r3g/Il6st/Pfkm/Gbe1/Pygm/Ppp1r3c/Sorbs1/Tcf7l2/Pygb/Ppp1r3d/Agl/Acadm/Lepr/Khk/Pgm2/Phkg1/Hmgb1/Gfpt1/Akt2/Gys1/Igf2/Insr/Ppp1r3b/Phkb/Phka1 | 31 |
| BP | GO:0060740 | prostate gland epithelium morphogenesis | 16/3915 | 32/17474 | 0,0005599 | 0,00569001 | 0,00423735 | Gli2/Igf1/Frs2/Sox9/Id4/Wnt5a/Bmp4/Rarg/Notch1/Cd44/Bmp7/Cyp7b1/Tnc/Igf1r/Fgfr2/Mmp2 | 16 |
| BP | GO:0030513 | positive regulation of BMP signaling pathway | 19/3915 | 41/17474 | 0,00059337 | 0,00594239 | 0,0044253 | Vsir/Bmp4/Acvrl1/Twsg1/Smad4/Smad2/Notch1/Eng/Notch2/Ccn1/Hes5/Elapor2/Rbpj/Kdr/Tgfbr3/Ilk/Cdh5/Neo1/Gpc3 | 19 |
| BP | GO:0031102 | neuron projection regeneration | 26/3915 | 63/17474 | 0,00060892 | 0,0060613 | 0,00451385 | Map4k4/Bcl2/Tnr/Prrx1/Enpp1/Kremen1/Grn/Gfap/Adam17/Mtr/Dhfr/Nrep/Jak2/Fas/Cers2/Klf4/Tnc/Hgf/Ptn/Igf1r/Xylt1/Nrg1/Mmp2/Neo1/Lamb2/Flna | 26 |
| BP | GO:0010563 | negative regulation of phosphorus metabolic process | 130/3915 | 449/17474 | 0,00062491 | 0,00620544 | 0,00462119 | Ctdsp1/Mgat5/Fam72a/Ptprc/Ddr2/Smyd3/Aida/Dusp10/Ppp2r5a/Cnksr3/Lats1/Enpp1/Mical1/Adora2a/Fbxo7/Igf1/Ptprb/Irak3/Ctdsp2/Pik3ip1/Grb10/Plek/Rack1/Irf1/Ncor1/Spag9/Cdk5rap3/Ppp1r1b/Stat3/Pecam1/Cd300a/Lpin1/Prkch/Dusp22/Lpcat1/Cmya5/Zbed3/Ocln/Nnt/Atxn7/Prkcd/Bmp4/Socs4/Rb1/Dnajc3/Angpt1/Deptor/Tspo/Fbln1/Ppara/Nckap1l/Tfap4/Prkdc/Ppm1f/Crkl/Heg1/Parp14/Pcp4/Stk38/Cdkn1a/Ier3/Kat2b/Xdh/Socs5/Rock1/Entpd1/Pip4k2a/Pip5kl1/Eng/Pax6/Spred1/Rassf2/Ppp1r16b/Ptprt/Pkig/Bmp7/Wwtr1/Kirrel/Ptpn22/Lyn/Cdk5rap2/Plpp3/Cdkn2c/Camk2n1/Prkcz/Hgf/Ppargc1a/Ppef2/Ptpn13/Gbp4/Mlxipl/Hspb1/Cav1/Wnk1/Ptpn6/Lrp6/Dusp16/Cdkn1b/Ptprh/Apoc1/Apoe/Ceacam1/Tgfb1/Sirt2/Kirrel2/Lrrk1/Cib1/Iqgap1/Arrb1/Inppl1/Ilk/Swap70/Plk1/Nupr1/Mvp/Pycard/Insr/Ikbkb/Casp3/Slc27a1/Agt/Pard3/Birc3/Ubash3b/Il18/Cd109/Ibtk/Gnai2/Dmd/Ogt | 130 |
| BP | GO:0045936 | negative regulation of phosphate metabolic process | 130/3915 | 449/17474 | 0,00062491 | 0,00620544 | 0,00462119 | Ctdsp1/Mgat5/Fam72a/Ptprc/Ddr2/Smyd3/Aida/Dusp10/Ppp2r5a/Cnksr3/Lats1/Enpp1/Mical1/Adora2a/Fbxo7/Igf1/Ptprb/Irak3/Ctdsp2/Pik3ip1/Grb10/Plek/Rack1/Irf1/Ncor1/Spag9/Cdk5rap3/Ppp1r1b/Stat3/Pecam1/Cd300a/Lpin1/Prkch/Dusp22/Lpcat1/Cmya5/Zbed3/Ocln/Nnt/Atxn7/Prkcd/Bmp4/Socs4/Rb1/Dnajc3/Angpt1/Deptor/Tspo/Fbln1/Ppara/Nckap1l/Tfap4/Prkdc/Ppm1f/Crkl/Heg1/Parp14/Pcp4/Stk38/Cdkn1a/Ier3/Kat2b/Xdh/Socs5/Rock1/Entpd1/Pip4k2a/Pip5kl1/Eng/Pax6/Spred1/Rassf2/Ppp1r16b/Ptprt/Pkig/Bmp7/Wwtr1/Kirrel/Ptpn22/Lyn/Cdk5rap2/Plpp3/Cdkn2c/Camk2n1/Prkcz/Hgf/Ppargc1a/Ppef2/Ptpn13/Gbp4/Mlxipl/Hspb1/Cav1/Wnk1/Ptpn6/Lrp6/Dusp16/Cdkn1b/Ptprh/Apoc1/Apoe/Ceacam1/Tgfb1/Sirt2/Kirrel2/Lrrk1/Cib1/Iqgap1/Arrb1/Inppl1/Ilk/Swap70/Plk1/Nupr1/Mvp/Pycard/Insr/Ikbkb/Casp3/Slc27a1/Agt/Pard3/Birc3/Ubash3b/Il18/Cd109/Ibtk/Gnai2/Dmd/Ogt | 130 |
| BP | GO:0035296 | regulation of tube diameter | 57/3915 | 171/17474 | 0,00065085 | 0,0064089 | 0,00477271 | Atp1a2/Gja1/Adora2a/Egfr/Adora2b/Serpinf2/Scpep1/Ace/Foxc1/Edn1/F2r/Itga1/Vstm4/Dock5/Ednrb/Casr/Cbs/Vegfa/Kat2b/Ptprm/Rock1/Adrb2/Acta2/Add3/Ptgs1/Edn3/P2ry1/Gucy1a1/Shc1/Gja5/Gclm/Pde5a/Nos3/Cd38/Kdr/Tgfbr3/P2rx4/Plod3/Gper1/Cav1/Tbxas1/Mgll/Hrh1/Alox5/Apoe/P2ry2/Klf2/Ednra/Mmp2/Bbs2/Agt/Arhgap42/Icam1/Rhoa/Apln/G6pdx/Cysltr1 | 57 |
| BP | GO:0097746 | blood vessel diameter maintenance | 57/3915 | 171/17474 | 0,00065085 | 0,0064089 | 0,00477271 | Atp1a2/Gja1/Adora2a/Egfr/Adora2b/Serpinf2/Scpep1/Ace/Foxc1/Edn1/F2r/Itga1/Vstm4/Dock5/Ednrb/Casr/Cbs/Vegfa/Kat2b/Ptprm/Rock1/Adrb2/Acta2/Add3/Ptgs1/Edn3/P2ry1/Gucy1a1/Shc1/Gja5/Gclm/Pde5a/Nos3/Cd38/Kdr/Tgfbr3/P2rx4/Plod3/Gper1/Cav1/Tbxas1/Mgll/Hrh1/Alox5/Apoe/P2ry2/Klf2/Ednra/Mmp2/Bbs2/Agt/Arhgap42/Icam1/Rhoa/Apln/G6pdx/Cysltr1 | 57 |

| | | | | | | | | | |
|---|---|---|---|---|---|---|---|---|---|
| BP | GO:0034612 | response to tumor necrosis factor | 60/3915 | 182/17474 | 0,00065576 | 0,0064344 | 0,0047917 | Stat1/Casp8/Tnfrsf11a/Aim2/Lims1/Irf1/Slc22a5/Slc22a21/Cxcl16/Traf4/Nos2/Rffl/Ccl9/Ccl6/Brca1/Adam17/Nfkbia/Zfp36l1/Traf3/Ripk1/Syk/Naip2/Naip5/Naip6/Erbin/Adamts12/Zfp36l2/Cd14/Rela/Tjp2/Jak2/Chuk/Traf1/Tank/Nfe2l2/Traf6/Cpne1/Cd40/H2bc21/Hipk1/Npnt/Nfkb1/Gbp3/Laptm5/Gper1/Tnfrsf1a/Zfp36/Cib1/Zfand6/Ilk/Mapk3/Pycard/Insr/Gas6/Ikbkb/Adam9/Asah1/Birc3/Rora/Adam10 | 60 |
| BP | GO:0006112 | energy reserve metabolic process | 34/3915 | 90/17474 | 0,00068467 | 0,00671005 | 0,00499697 | Irs1/Enpp1/S100b/Igf1/Grb10/Ugp2/Pygl/Ppp1r3g/Il6st/Pfkm/Gbe1/Adgrf5/Pygm/Ppp1r3c/Sorbs1/Tcf7l2/Pygb/Ppp1r3d/Agl/Acadm/Lepr/Khk/Pgm2/Phkg1/Hmgb1/Gfpt1/Akt2/Gys1/Igf2/Insr/Ppp1r3b/Phkb/Mt3/Phka1 | 34 |
| BP | GO:0031529 | ruffle organization | 24/3915 | 57/17474 | 0,00069041 | 0,00673524 | 0,00501573 | Plek/Lpin1/Carmil1/Lima1/Aif1/Fam98a/Ston1/Pdgfrb/Csf1r/Tcirg1/Coro1b/Kank1/Aif1l/P2ry12/Arhgef26/Arhgap24/Cav1/Eps8/Eps8l1/Cyfip1/Inppl1/Eps8l2/Icam1/Rdx | 24 |
| BP | GO:0032388 | positive regulation of intracellular transport | 61/3915 | 186/17474 | 0,0006905 | 0,00673524 | 0,00501573 | Pgap1/Map2/Rab29/Fyn/Sar1a/Rab21/Myo1c/Spag5/Brca1/Prkd1/Gli3/Pik3r1/Prkcd/Xpo4/Dab2/Ubr5/Mlc1/Rapgef3/Hcls1/Ezr/Crebrf/Kif5b/Jak2/Kif20b/Prkcq/Stom/Wipf1/Pla2g4e/Mavs/Edem2/Ptpn22/Sh3glb1/Asph/Anp32b/Tek/Scp2/Ldlrap1/Gper1/Smo/Edem1/Camk1/Tnfrsf1a/Ehd2/Tgfb1/Akt2/Cib1/Cemip/Chp2/Itgam/Cd81/Gas6/Pcm1/Cdh1/Rdx/Uaca/Anxa2/Nedd4/Zic1/Mecp2/Flna/Msn | 61 |
| BP | GO:0044042 | glucan metabolic process | 31/3915 | 80/17474 | 0,00070429 | 0,00682939 | 0,00508584 | Irs1/Enpp1/Igf1/Grb10/Ugp2/Pygl/Ppp1r3g/Il6st/Pfkm/Gbe1/Pygm/Ppp1r3c/Sorbs1/Tcf7l2/Pygb/Ppp1r3d/Agl/Acadm/Lepr/Khk/Pgm2/Phkg1/Hmgb1/Gfpt1/Akt2/Gys1/Igf2/Insr/Ppp1r3b/Phkb/Phka1 | 31 |
| BP | GO:0009948 | anterior/posterior axis specification | 23/3915 | 54/17474 | 0,00073042 | 0,0070414 | 0,00524373 | Pgap1/Epb41l5/Hey2/Frs2/Ddit3/Lhx1/Wnt3/Tifab/Tasor/Wnt5a/Bmp4/Smad4/Nrarp/Wls/Tbx3/Tcf7l1/Lrp6/Pcsk6/Tbx18/Ctnnb1/Kdm6a/Gpc3/Zic3 | 23 |
| BP | GO:0045730 | respiratory burst | 14/3915 | 27/17474 | 0,00077636 | 0,00744944 | 0,00554759 | Slc11a1/Ncf2/Dusp10/Grn/Cybc1/Ncf4/Rac2/Trem2/Lbp/Ncf1/Rps19/Insr/Cyba/Cybb | 14 |
| BP | GO:0033628 | regulation of cell adhesion mediated by integrin | 21/3915 | 48/17474 | 0,00080379 | 0,00769479 | 0,00573031 | Itgb3/Syk/Plau/Nckap1l/Crkl/Cyp1b1/Fermt3/Lpxn/Dpp4/P2ry12/Efna1/Lyn/Acer2/Ptpn11/Podxl/Ret/Wnk1/Ptpn6/Cib1/Swap70/Adam9 | 21 |
| BP | GO:0031345 | negative regulation of cell projection organization | 68/3915 | 213/17474 | 0,00081155 | 0,00776001 | 0,00577888 | Map4k4/Map2/Neu4/Rab29/Tnr/Katna1/Fyn/Zfp365/Kremen1/Efemp1/Pmp22/Rap1gap2/Abi3/Gfap/Wnt3/Adam17/Wnt5a/Prkcd/Sema3g/Dab2/Sema5a/Lgals1/Dip2b/Cep97/H2-K1/H2-D1/Nrxn1/Sema6a/Coro1b/Kank1/Il15ra/Vim/Tsc1/Lrp4/B2m/Sema6d/Id1/Ctsz/Efna1/Snapin/Cers2/Odf2l/Dab1/Sema3d/Sema3c/Cd38/Arhgap24/Ptprz1/Apoe/Cib1/Inppl1/Dennd5a/Xylt1/Ccp110/Mt3/Cdh1/Itgb1/Yap1/Dnm2/Ntm/Neo1/Ryk/Rhoa/Ccr5/Thoc2/Fgf13/Arhgap4/Flna | 68 |
| BP | GO:0051642 | centrosome localization | 16/3915 | 33/17474 | 0,00086237 | 0,00815793 | 0,00607521 | Pard3b/Aspm/Syne2/Dlgap5/Sun2/Nde1/Dlg1/Ezr/Kif5b/Sema6a/Spout1/Ccdc141/Gpsm2/Akap9/Fhod1/Pard3 | 16 |
| BP | GO:0002011 | morphogenesis of an epithelial sheet | 25/3915 | 61/17474 | 0,00086917 | 0,00818769 | 0,00609737 | Vangl2/Map3k1/Wnt5a/Fermt2/Acvrl1/Phldb2/Lama1/Hbegf/Chuk/Notch1/Cd44/Lin7c/Jag1/Bmp7/Notch2/Ccn1/Phactr4/Pdpn/Arhgap24/Lrp6/Cd151/Pard3/Bmp5/Clasp2/Flna | 25 |
| BP | GO:0035051 | cardiocyte differentiation | 60/3915 | 184/17474 | 0,00089186 | 0,00834423 | 0,00621395 | Sox17/Vangl2/Prox1/Hey2/Igf1/Frs2/Egfr/Meis1/Sgcd/Dicer1/Edn1/Mef2c/Isl1/Bmpr1a/Bmp4/Myh6/Col14a1/Ppara/Myh11/Abi3bp/Cxadr/Vegfa/Trip10/Nr3c1/Pdgfrb/Smad4/Nrap/Nebl/Notch1/Tsc1/Ttn/Jag1/Bmp7/Vcam1/Camk2d/Pdlim5/Acadm/Zmpste24/Sema3c/Rbpj/Sgcb/Pdgfra/Rest/Tgfbr3/Myo18b/Tbx3/Gper1/Atg7/Ccnd2/Lrp6/Tgfb1/Sox6/Mapk3/Nrg1/Ednra/Agt/Itgb1/Tbx18/Ctnnb1/G6pdx | 60 |
| BP | GO:0051651 | maintenance of location in cell | 72/3915 | 229/17474 | 0,00093468 | 0,0086468 | 0,00643927 | Sp100/Dbi/Ptprc/Aspm/Pln/Gja1/Srgn/Ddit3/Ccdc88a/Gm2a/Gp1ba/Itgb3/Prkd1/F2r/Hexb/Dzip1/Sun2/Tspo/Twf1/Morc3/Slc25a23/Plcb3/Plce1/Spout1/Itgav/F2/Plcb2/Capn3/Plcb4/Fgf2/Casq2/Gstm7/Gpsm2/Slc30a7/Camk2d/Lyn/Chd7/Asph/Tmem38b/Akap9/Sri/Insig1/Pkd2/Slc8b1/P2rx7/Gper1/Kdelr2/Cav1/Hk2/Gp9/Ptpn6/Itpr2/Apoe/Tgfb1/Ftl1/Cemip/P2ry6/Tnrc6a/Atp7b/Ednra/Arl2bp/Plcg2/Cyba/Ubash3b/Hexa/Polr2m/Myzap/Ibtk/Plcd1/Ccr5/Flna/Dmd | 72 |
| BP | GO:0045807 | positive regulation of endocytosis | 45/3915 | 130/17474 | 0,0009586 | 0,00880888 | 0,00655997 | Appl2/Rab21/Cd63/Pld2/Ankfy1/Vtn/Sfrp4/Hfe/Syk/Wnt5a/Clu/Dab2/Angpt1/Ano6/Nckap1l/Itsn1/Vegfa/Trem2/Rab31/Cd14/Ckap5/B2m/Lbp/Fcgr1/Ldlrap1/Hip1r/Hip1/Cav1/Alms1/Gata2/Apoe/Axl/Picalm/Arrb1/Eef2k/Cln3/Cd151/Insr/Plcg2/Pard3/Itgb1/Dnm2/Anxa2/Apln/Gpc3 | 45 |
| BP | GO:0051208 | sequestering of calcium ion | 45/3915 | 130/17474 | 0,0009586 | 0,00880888 | 0,00655997 | Dbi/Ptprc/Pln/Ddit3/Gp1ba/Itgb3/Prkd1/F2r/Slc25a23/Plcb3/Plce1/Itgav/F2/Plcb2/Capn3/Plcb4/Fgf2/Casq2/Gstm7/Camk2d/Lyn/Chd7/Asph/Tmem38b/Sri/Pkd2/Slc8b1/P2rx7/Gper1/Gp9/Ptpn6/Itpr2/Tgfb1/Cemip/P2ry6/Atp7b/Ednra/Plcg2/Cyba/Ubash3b/Ibtk/Plcd1/Ccr5/Flna/Dmd | 45 |
| BP | GO:0042445 | hormone metabolic process | 66/3915 | 207/17474 | 0,00099573 | 0,00905656 | 0,00674442 | Chst10/Hsd11b1/Iyd/Enpp1/Ddo/Sgpl1/Hsp90b1/Igf1/Stat5b/Ace/P4hb/Dhrs7/Rdh11/Dio2/Hfe/Bmp6/Ctsl/Pde8b/Gcnt4/Ednrb/Ghr/Dab2/Cpq/Cyp2d22/Cyp1b1/Nr3c1/Hsd17b4/Aldh1a1/Papss2/Ide/Tcf7l2/Slco4a1/Myt1/Tiparp/Crabp2/Arnt/Ctsk/Nfkb1/Bmpr1b/Scp2/Zmpste24/Dhrs3/Reln/Plb1/Ppargc1a/Pdgfra/Rest/Hsd17b11/Spp1/Ptpn11/Retsat/Slco1c1/Pcsk6/Igf1r/Sult1a1/Igf2/Dhcr7/Cpe/Agt/Bco2/Adam10/Aldh1a2/Bmp5/Rbp1/Slc16a2/Hsd17b10 | 66 |
| BP | GO:0070997 | neuron death | 129/3915 | 450/17474 | 0,00099653 | 0,00905656 | 0,00674442 | Tfap2b/Casp8/Bok/Bcl2/Pm20d1/Rab29/Btg2/Ncf2/Ddr2/Fcgr2b/Trp53bp2/Capn2/Slc30a10/Enpp1/Fyn/Adora2a/Fbxo7/Igf1/Tmbim4/Ddit3/Erbb3/Rel/Rack1/Nlrp1b/Serpinf1/Stat3/Grn/Egln3/Snx6/Lgmn/Nqo2/Ntrk2/Mef2c/F2r/Il6st/Itga1/Isl1/Plau/Wnt5a/Sncg/Clu/Rb1/Ghr/Angpt1/Nrbp2/Ppara/Hdac7/Tmbim6/Cd200r1/Gbe1/Itsn1/Pcp4/Vegfa/Trem2/Rock1/Pcdhgc3/Nr3c1/Vegfb/Jak2/Fas/Casp7/Tsc1/Dlx1/Mdk/Spg11/Kcnip3/Bcl2l11/Ctsz/Pik3ca/Slc7a11/Nes/Csf1/Gclm/Usp53/Casp6/Tlr4/C1qa/Tnfrsf1b/Trp73/Xrcc2/Fgfr3/Ppargc1a/Tlr6/Kdr/Rest/Hrk/Diablo/Ptprz1/Smo/Hipk2/Casp2/Gpnmb/Npy/Grid2/Ndnf/Atg7/Tnfrsf1a/Zfp110/C5ar1/Apoe/Tgfb1/Axl/Akt2/Tyrobp/St8sia2/Picalm/Arrb1/Ilk/Cln3/Nupr1/Itgam/Ikbkb/Casp3/Mt3/Mt1/Nqo1/Agt/Il18/Neo1/Ephb1/Rhoa/Cx3cr1/Ctnnb1/Ccr5/Ndp/Xiap/Mecp2/G6pdx/Atp7a | 129 |
| BP | GO:0071407 | cellular response to organic cyclic compound | 132/3915 | 462/17474 | 0,00100958 | 0,00916511 | 0,00682525 | Stat1/Igfbp5/Gli2/Ralb/Atp1a2/Smyd3/Lats1/Akap7/Gna15/Hsp90b1/Igf1/Egfr/Ncor1/Nos2/Ppp1r1b/Stat3/Brca1/Pik3cg/Nfkbia/Zfp36l1/Gpld1/Smad5/Ntrk2/Ptch1/Mef2c/Isl1/Fam107a/Bmp4/Dab2/Slc1a3/Ubr5/Fbxo32/Ddx17/Mlc1/Rapgef3/Tfap4/Spidr/Ube2l3/Pak2/Gabpa/Ezr/Crebrf/Itpr3/Lbh/Cyp1b1/Zfp36l2/Riok3/Npc1/Ctnna1/Sting1/Nr3c1/Jak2/Lcor/Casp7/Ifih1/Slc1a2/Ddrgk1/Mavs/Tgm2/Bmp7/Cyp7b1/P2ry13/P2ry12/P2ry1/Tiparp/Rorc/Casq2/Slc16a1/Rap1a/Gstm7/Gnai3/Casp6/Nfkb1/Tmem38b/Heyl/Hdac1/Id3/Alpl/Padi2/Srarp/Akap9/Insig1/Gabrb1/Rest/Spp1/Pkd2/P2rx7/P2rx4/Gper1/Cav2/Cav1/Smo/Zc3hav1/Ezh2/Kdm3a/Hrh1/Lrp6/Cdkn1b/Pde3a/Itpr2/Tgfb1/Akt2/Zfp36/Pak4/Sirt2/Abhd2/Fes/P2ry6/Trim68/Mapk3/Eif4ebp1/Tlr3/Casp3/Smad1/Mmp2/Cdh1/Yap1/Dnm2/Kank2/Htr3a/Il18/Trip4/Rora/Nedd4/Rhoa/Slc26a6/Ctnnb1/Kdm6a/Mecp2/Flna/Msn/Hdac8 | 132 |
| BP | GO:0008202 | steroid metabolic process | 91/3915 | 302/17474 | 0,00102258 | 0,00927288 | 0,00690551 | Chst10/Acadl/Cyp27a1/Soat1/Pbx1/Lbr/Prox1/Hsd11b1/Enpp1/Sgpl1/Lss/Igf1/Rdh5/Adora2b/Pmp22/Acadvl/Dhrs11/Pctp/Stat5b/Fdxr/Npc2/Bmp6/Pde8b/Hmgcs1/Ephx2/Ednrb/Dab2/Cyp2d22/Tspo/Lima1/Lmf1/Cyp1b1/Npc1/Stard4/Fgf1/Nr3c1/Hsd17b4/Atp8b1/Lrp5/Hsd17b12/Cat/Pex2/Cyp7b1/Tiparp/Pmvk/Rorc/Fmo5/Hmgcs2/Abcd3/Nfkb1/Mttp/Bmpr1b/Abca1/Lepr/Scp2/Ldlrap1/Cyp51/Insig1/Ppargc1a/Pdgfra/Rest/Igfbp7/Hsd17b11/Spp1/Mvk/Hrh1/Lpcat3/Apoc1/Apoe/Lipe/Igf1r/Sult1a1/Hsd3b7/Igf2/Dhcr7/Erlin2/Asah1/Msmo1/Mt3/Lcat/Sdr42e1/Ldlr/Aplp2/Sc5d/Rora/Bmp5/Ebp/Nsdhl/Mecp2/G6pdx/Hsd17b10 | 91 |
| BP | GO:0007252 | I-kappaB phosphorylation | 12/3915 | 22/17474 | 0,00103226 | 0,00931972 | 0,00694039 | Ikbke/Chuk/Ddrgk1/Tlr2/Map3k7/Tlr4/Erc1/Ikbkb/Tlr3/Plcg2/Cx3cr1/Tlr7 | 12 |
| BP | GO:0010232 | vascular transport | 12/3915 | 22/17474 | 0,00103226 | 0,00931972 | 0,00694039 | Gja1/Slc22a5/Slc2a1/Mfsd2a/Abcb1a/Abcb1b/Slc5a6/Abcg2/Slc1a5/Apoe/Slc27a1/Slc16a2 | 12 |

| | | | | | | | | | |
|---|---|---|---|---|---|---|---|---|---|
| BP | GO:0006869 | lipid transport | 122/3915 | 423/17474 | 0,00105421 | 0,00949709 | 0,00707248 | Ecrg4/Gulp1/Acsl3/Tnfrsf11a/Dbi/Pla2g4a/Soat1/Eprs/Fabp7/Psap/Esyt1/Acsl6/Gm2a/Nos2/Pctp/Abcc3/Itgb3/Ace/Pitpnc1/Abca8a/Abca9/Abca6/C1qtnf1/Hbp1/Nfkbia/Abcd4/Npc2/Bmp6/Edn1/Syk/Ptch1/Prkcd/Xkr6/Abcc4/Dab2/Tspo/Abcd2/Ano6/Lima1/Apod/Atp5j/Runx1/Trem2/Npc1/Stard4/Atp8b1/Prelid3a/Stard6/Slc66a2/Slc22a8/Slc22a6/Pip4k2a/Ptges/Pla2r1/Itgav/Slc43a3/Nr1h3/Thbs1/Spg11/Lbp/Pltp/Fabp5/Atp11b/Crabp2/Abcd3/Abca4/Mttp/Ttpa/Abca1/Plin2/Scp2/Osbpl9/Slc2a1/Mfsd2a/Xkr8/Ldlrap1/Pla2g5/Abcb1a/Abcb1b/Crot/Atp8a1/Atp10d/Scarb2/Spp1/Gltp/Pla2g1b/Ptpn11/P2rx7/Cav1/Osbpl3/Abcg2/Lpcat3/Lrp6/Slco1c1/Slco1a4/Apoc1/Apoe/Ceacam1/Akt2/Atp10a/Stard5/Fzd4/Slco2b1/P2ry2/Stard10/Cln3/Atp11a/Nrg1/Slc27a1/Lcat/Agt/Ldlr/Anxa2/Aqp9/Plscr1/Plscr2/Plscr4/Rbp1/Slco2a1/Slc25a20/Atp11c/Irak1 | 122 |
| BP | GO:0043087 | regulation of GTPase activity | 103/3915 | 349/17474 | 0,00109148 | 0,0097901 | 0,00729069 | Map4k4/Dock10/Srgap2/Dennd1b/F11r/Lims1/Arhgap45/Fgd6/Arhgap9/Tns3/Rack1/Arhgef15/Rap1gap2/Poldip2/Adap2/Ccl9/Ccl6/Pecam1/Tbc1d16/Rgs6/Ttc8/Eif5/Net1/Gpr137b/Nedd9/Ntrk2/Rasgrf2/F2r/Wnt5a/Arhgap22/Fermt2/Lrch1/Tbc1d4/Rictor/Tbc1d22a/Plxnb2/Rapgef3/Crkl/Snx9/Rrp1b/Rasal3/Mmut/Pot1b/Vav1/Ralbp1/Rasgrp3/Arap3/Sipa1/Rasgrp2/Dock8/Usp6nl/Rsu1/Tsc1/Garnl3/Gapvd1/Itga6/Rapgef4/Cd40/Arhgef26/Rap1a/Vav3/S1pr1/Arhgap29/Bcar3/Dock7/Asap3/Arhgef19/Arap2/Arhgap24/Evi5/Rasa4/Cav2/Arhgef5/Ezh2/Chn2/Arhgap25/Wnk1/Akt2/Iqgap1/Picalm/Arrb1/Arap1/Sbf2/Pycard/Rgs10/Grtp1/Rasa3/Arhgef10/Gmip/Elmod2/Itgb1/Arhgap42/Icam1/Dock6/Rdx/Tbc1d2b/Rasa2/Plxnb1/Als2cl/Dock11/Tbc1d8b/Amot/Arhgap6 | 103 |
| BP | GO:0008360 | regulation of cell shape | 47/3915 | 138/17474 | 0,00111232 | 0,00995967 | 0,00741696 | Fn1/Vil1/F11r/Epb41l2/Arhgap18/Fyn/S100b/Gna13/Cdc42ep4/Rhoj/Hexb/Fermt2/Myo10/Plxnb2/Fmnl3/Fgd4/Dlg1/Brwd1/Ezr/Vegfa/Myl12a/Cdc42ep3/Csf1r/Fmnl2/Prpf40a/F2/Hck/P2ry1/S100a13/Zmpste24/Epb41/Pdpn/Kdr/Wasf3/Ptn/Eps8/Cyfip1/Fes/Arap1/Parva/Dlc1/Icam1/Rdx/Plxnb1/Limd1/Msn/Sh3kbp1 | 47 |
| MF | GO:0005178 | integrin binding | 79/3915 | 150/17474 | 5,1343E-16 | 4,8247E-13 | 3,5929E-13 | Dst/Col3a1/Fn1/Tnr/F11r/Utrn/Ccn2/Ccn6/Itgb2/Igf1/Egfr/Vtn/Itga2b/Gfap/Itgb3/Icam2/P4hb/Adam17/Fbln5/Itgb8/Lgals8/Syk/Tgfbi/Thbs4/Itga1/Fermt2/Mmp14/Lcp1/Dab2/Fbln1/Itga5/Emp2/Itgb5/Casr/Cxadr/Jam2/Lama3/Fgf1/Fermt3/Gfra1/Itga6/Itgav/Thbs1/Lama5/Fgf2/Vcam1/Npnt/Ccn1/Lyn/Tln1/Plpp3/Col16a1/Isg15/Kdr/Spp1/Hmgb1/Ptprz1/Ptn/Pdia4/Gpnmb/Cxcl12/Cd9/Vwf/Mfge8/Ilk/Itgal/Itgam/Cd151/Igf2/Cd81/Adam9/Nrg1/Itgb1/S1pr2/Icam1/Lamb2/Itga9/Dmd/Egfl6 | 79 |
| MF | GO:0050839 | cell adhesion molecule binding | 127/3915 | 298/17474 | 3,636E-15 | 1,6855E-12 | 1,2552E-12 | Dst/Col3a1/Fn1/Cdh20/Cdh19/Tnr/F11r/Utrn/Ccn2/Ccn6/Ctnna3/Itgb2/Igf1/Ptprb/Egfr/Vtn/Nos2/Itga2b/Gfap/Itgb3/Icam2/P4hb/Adam17/Numb/Fbln5/Itgb8/Lgals8/Dsp/Ninj1/Syk/Tgfbi/Thbs4/Itga1/Fermt2/Mmp14/Lcp1/Dab2/Cdh9/Fbln1/Itga5/Emp2/Itgb5/Casr/Cd200r1/Nectin3/Cxadr/Jam2/Ezr/Cd2ap/Ptprm/Epcam/Nrxn1/Lama3/Ctnna1/Fgf1/Fermt3/Tjp2/Gfra1/Itga6/Itgav/Ctnnd1/Thbs1/Ptprt/Lama5/Fgf2/Cd1d1/Kirrel/Vcam1/Npnt/Ccn1/Nexn/Lyn/Tln1/Ctnnal1/Ptprd/Plpp3/Col16a1/Isg15/Nos3/Prom1/Kdr/Spp1/Ptpn11/P2rx4/Hmgb1/Ptprz1/Ptn/Pdia4/Gpnmb/Cntn6/Cxcl12/Ptpn6/Cd9/Vwf/Ptprh/Nectin2/Bcam/Kirrel2/Nphs1/Fxyd5/Tjp1/Mfge8/Ilk/Itgal/Itgam/Cd151/Igf2/Cd81/Adam9/Nrg1/Cpe/Cdh5/Cdh1/Itgb1/Cntn5/S1pr2/Icam1/Esam/Rdx/Neo1/Lamb2/Itga9/Rpsa/Ctnnb1/Dmd/Msn/Egfl6 | 127 |
| MF | GO:0019838 | growth factor binding | 64/3915 | 139/17474 | 5,4979E-10 | 5,041E-08 | 3,754E-08 | Il1r1/Col3a1/Erbb4/Igfbp2/Igfbp5/Erbb3/Egfr/Flt4/Trim16/Col1a1/Itgb3/Ntrk2/Il6st/Ghr/Rpl37/Osmr/Lifr/Acvrl1/Vasn/Nrros/Rps2/Vegfa/Twsg1/Ltbp1/Pdgfrb/Csf1r/Fgfbp3/Kazald1/Eng/Itga6/Itgav/Thbs1/Shc1/Il6ra/S100a13/Ccn1/Il11ra1/Tgfbr1/Tek/Fgfr3/Fgfbp1/Pdgfra/Kdr/Igfbp7/Tgfbr3/Flt1/Col1a2/Ptprz1/Ptn/A2m/Rps19/Pcsk6/Igf1r/Lrrc32/Fgfr2/Htra1/Insr/Il10ra/Cd109/Nradd/Tgfbr2/Acvr2b/Il2rg/Chrdl1 | 64 |
| MF | GO:0005201 | extracellular matrix structural constituent | 65/3915 | 143/17474 | 7,858E-10 | 6,6168E-08 | 4,9275E-08 | Col9a1/Col19a1/Col3a1/Fn1/Prelp/Hmcn1/Lamc2/Lama2/Lama4/Col13a1/Col18a1/Lum/Emid1/Efemp1/Col23a1/Sparc/Mfap4/Vtn/Col1a1/Fbln5/Nid1/Aspn/Ogn/Tgfbi/Hapln1/Vcan/Thbs4/Nid2/Colq/Mmrn2/Scara3/Col14a1/Fbln1/Creld2/Abi3bp/Col11a2/Lama1/Ltbp1/Lama3/Efemp2/Thbs1/Matn4/Lama5/Npnt/Col27a1/Tnc/Col9a2/Col16a1/Tinagl1/Hspg2/Vwa1/Reln/Igfbp7/Eln/Col1a2/Vwf/Mfge8/Bmper/Vwa5a/Col12a1/Col6a6/Lamb2/Bgn/Col4a6/Col4a5 | 65 |
| MF | GO:0050840 | extracellular matrix binding | 34/3915 | 57/17474 | 1,4492E-09 | 1,1282E-07 | 8,4015E-08 | Adgrg6/Ntn4/Sparc/Vtn/Itga2b/Itgb3/Clec14a/Smoc1/Nid1/Tgfbi/Thbs4/Lgals3/Lgals1/Olfml2a/Itga6/Itgav/Thbs1/Ctss/Ccn1/Tinagl1/Sparcl1/Dmp1/Spp1/Eln/Bcam/Thsd1/Adam9/Plekha2/Adgrg1/Itgb1/Anxa2/Itga9/Rpsa/Bgn | 34 |
| MF | GO:0005539 | glycosaminoglycan binding | 85/3915 | 213/17474 | 6,0858E-09 | 3,8257E-07 | 2,849E-07 | Nrp2/Fn1/Serpine2/Prelp/Ptprc/Cfh/Lamc2/Ccn2/Ccn6/Col13a1/Stab2/Gns/Col23a1/Vtn/Smoc1/Ecm2/Ptch1/Hapln1/Vcan/Thbs4/Stab1/Colq/Bmp4/Ang/Egflam/Sema5a/Serpind1/Fstl1/Ccdc80/Abi3bp/Lipi/Adamts1/Eva1c/Thbs2/Vegfa/Trem2/Twsg1/Hbegf/Fgf1/Efemp2/Vegfb/Fgfbp3/Itih2/Eng/F2/Mdk/Hsd17b12/Cd44/Lgr4/Thbs1/Fbln7/Bmp7/Fgf2/Lxn/Tlr2/Bcan/Ccn1/Rspo1/Pla2g5/Rpl22/Tgfbr3/Pcolce/Hmgb1/Ptn/Gpnmb/Ndnf/Pglyrp1/Apoe/Pcsk6/Hapln3/Cemip/Lyve1/Itgam/Fgfr2/Spock3/Ncan/Adgre5/Nod2/Adgrg1/Crispld2/Aplp2/Rpl29/Tgfbr2/Clec3b/Bgn | 85 |
| MF | GO:1901681 | sulfur compound binding | 96/3915 | 258/17474 | 4,2515E-08 | 2,096E-06 | 1,5609E-06 | Nrp2/Acadl/Fn1/Serpine2/Dbi/Prelp/Ptprc/Cfh/Lamc2/Soat1/Ccn2/Ccn6/Col13a1/Ilvbl/Gns/Col23a1/Acadvl/Vtn/Acaca/Smoc1/Eci2/Tpmt/Ecm2/Ptch1/Thbs4/Fst/Tkt/Colq/Hacl1/Bmp4/Ang/Sema5a/Serpind1/Fstl1/Ccdc80/Abi3bp/Lipi/Adamts1/Eva1c/Hlcs/Thbs2/Cbs/Vegfa/Twsg1/Hbegf/Fgf1/Smad4/Kmt5b/Efemp2/Vegfb/Pank1/Fgfbp3/Dhtkd1/Acbd5/F2/Mdk/Hsd17b12/Lgr4/Thbs1/Fbln7/Bmp7/Fgf2/Lxn/Dbt/Ccn1/Acadm/Scp2/Rspo1/Hmgcl/Pla2g5/Rpl22/Hadha/Tgfbr3/Acacb/Acads/Pcolce/Hmgb1/Ptn/Tpk1/Gpnmb/Ndnf/Slc6a6/Apoe/Ftl1/Pcsk6/Sult1a1/Itgam/Fgfr2/Insr/Adgre5/Gcdh/Adgrg1/Crispld2/Aplp2/Rpl29/Clec3b | 96 |
| MF | GO:0019842 | vitamin binding | 58/3915 | 140/17474 | 3,4187E-07 | 1,2056E-05 | 8,9781E-06 | Lmbrd1/Mtarc2/P4ha1/Sgpl1/Ilvbl/Shmt2/Tcn2/Ddc/Shmt1/Slc46a1/Acaca/Ogfod3/Egln3/Pygl/Sptlc2/Mtr/Dhfr/Tkt/Hacl1/Gpt/Csad/Abat/Pdxdc1/Hlcs/Cbs/Mmut/Mocos/Pygm/Psat1/Gldc/Phyh/Dhtkd1/Gad2/Sardh/Gad1/Accs/Pygb/Crabp2/Abca4/Etnppl/Kyat3/Cth/Ttpa/Plod1/Acacb/Plod3/Tpk1/Thnsl2/Ggcx/Rlbp1/P4ha3/Oat/Gpt2/Aldh1a2/Plod2/Rbp1/Gadl1/Alas2 | 58 |
| MF | GO:0005518 | collagen binding | 35/3915 | 70/17474 | 3,7117E-07 | 1,2869E-05 | 9,5837E-06 | Ddr2/Srgn/Lum/Aebp1/Sparc/Vtn/P3h4/Mrc2/C1qtnf1/Nid1/Ecm2/Aspn/Tgfbi/Ctsl/Thbs4/Itga1/Abi3bp/Lox/Smad4/Dpp4/Hsd17b12/Thbs1/Ctsk/Ctss/Hspg2/Sparcl1/Pcolce/Antxr1/Vwf/Serpinh1/Adam9/Adgrg1/Itgb1/Crtap/Itga9 | 35 |
| MF | GO:0030170 | pyridoxal phosphate binding | 28/3915 | 52/17474 | 7,8058E-07 | 2,4585E-05 | 1,8309E-05 | Mtarc2/Sgpl1/Shmt2/Ddc/Shmt1/Pygl/Sptlc2/Gpt/Csad/Abat/Pdxdc1/Cbs/Mocos/Pygm/Psat1/Gldc/Gad2/Gad1/Accs/Pygb/Etnppl/Kyat3/Cth/Thnsl2/Oat/Gpt2/Gadl1/Alas2 | 28 |
| MF | GO:0003779 | actin binding | 139/3915 | 436/17474 | 2,4734E-06 | 6,6054E-05 | 4,919E-05 | Dst/Myo1b/Map2/Tns1/Vil1/Mlph/Lmod1/Tnni1/Rcsd1/Utrn/Phactr2/Epb41l2/Mical1/Ctnna3/Cnn2/Eef2/Smtn/Myo1g/Egfr/Ccdc88a/Pdlim4/Flii/Myo1c/Nos2/Myo19/Lasp1/Ace/P4hb/Cfl2/Daam1/Syne2/Clmn/Phactr1/Fam107a/Pxk/Vcl/Fermt2/Gmfb/Ang/Myh6/Gjb6/Lcp1/Rai14/Myo10/Ncald/Mtss1/Triobp/Parvg/Twf1/Fmnl3/Lima1/Myh11/Fgd4/Mylk/Hcls1/Myh15/Abi3bp/Ezr/Pacrg/Myo1f/Aif1/Ppp1r18/Daam2/Svil/Ctnna1/Spire1/Coro1b/Add3/Nrap/Ablim1/Nebl/Aif1l/Fmnl2/Cobll1/Wipf1/Ttn/Itprid2/Kif18a/Trpm7/Epb41l1/Phactr3/S100a4/Capza1/Cnn3/Synpo2/Pdlim5/Gbp2/Nexn/Tpm2/Tln1/Ctnnal1/Fkbp15/Macf1/Epb41/Phactr4/Wasf2/Nos3/Afap1/Limch1/Myo18b/Hip1r/Hip1/Arpc1b/Wasf3/Cald1/Capg/Antxr1/Eps8/Eps8l1/Vasp/Ceacam1/Fxyd5/Cyfip1/Iqgap1/Homer2/Myo7a/Inppl1/Parva/Eps8l2/Lsp1/Pdlim3/Palld/Tpm4/Nod2/Fhod1/Itgb1/Cnn1/Rdx/Pstpip1/Myo1e/Tmod3/Myo6/Clasp2/Ccr5/Shroom4/Was/Flna/Dmd/Msn | 139 |

| | | | | | | | | | |
|---|---|---|---|---|---|---|---|---|---|
| MF | GO:0005543 | phospholipid binding | 144/3915 | 458/17474 | 3,8376E-06 | 9,3415E-05 | 6,9566E-05 | Sgk3/Plekhb2/Cavin2/Myo1b/Pard3b/Vil1/Wdfy1/Dennd1b/Pla2g4a/Rcsd1/Aida/Psap/Appl2/Arhgap9/Esyt1/Dgka/Myo1g/Ccdc88a/Sh3pxd2b/Sap30l/Nlrp3/Flii/Pld2/Ankfy1/Myo1c/Adap2/Pctp/Tom1l1/Grb7/Pitpnc1/Wipi1/Kcnj2/Cd300a/Snx6/Zfyve26/Pxdc1/Zfyve16/Fcho2/Pxk/Fermt2/Amer2/Dab2/Myo10/Mtss1/Gsdmd/Ncf4/Baiap2l2/Pacsin2/Twf1/Bin2/Snx29/Pcyt1a/Snx4/Snx9/Itpr3/Fgd2/Pla2g7/Trem2/Dennd1c/Svil/Psd2/Arap3/Ticam2/Atp8b1/Myof/Pacsin3/Thbs1/Plcb2/Pla2g4e/Jag1/Kif16b/Snx5/Sdcbp2/Cpne1/Pltp/Pld1/Anxa5/Arfip1/Snx27/Golph3l/Syt6/F3/Dapp1/Plekhf2/Cpne3/Ttpa/Tln1/Abca1/Dab1/Epb41/Ptafr/Ldlrap1/Pla2g5/Plekhn1/Arap2/Scarb2/Anxa3/Gltp/Pla2g1b/Tpcn1/Hip1r/Ncf1/Hip1/Rasa4/Snx8/Hmgb1/Capg/Dysf/Anxa4/Tulp3/Itpr2/Apoe/Ceacam1/Axl/Plekhf1/Mfge8/Rlbp1/Fes/Iqgap1/Picalm/Fchsd2/Arap1/Inppl1/Tpp1/Sbf2/Pik3c2a/Gas6/Pard3/Tirap/Snx33/Snx22/Anxa2/Myo1e/Unc13c/Rasa2/Plcd1/Sytl5/Mtm1/Pcyt1b/Amer1/Ogt/Lpar4/Sytl4/Btk | 144 |
| MF | GO:0001968 | fibronectin binding | 20/3915 | 34/17474 | 4,8741E-06 | 0,00011426 | 8,5093E-05 | Igfbp5/Myoc/Ccn2/Itgb3/Ctsl/Thbs4/Fbln1/Ccdc80/Vegfa/Itgav/Hsd17b12/Thbs1/Sdc4/Ctsk/Ctss/Tnc/Loxl3/Plekha2/Mmp2/Itgb1 | 20 |
| MF | GO:0140375 | immune receptor activity | 46/3915 | 116/17474 | 2,1062E-05 | 0,00039379 | 0,00029326 | Il1r1/Il1rl2/Fcgr2b/Fcgr3/Fcer1g/Cr1l/Ifngr1/Il6st/Il17rd/Il17rb/Ghr/Osmr/Lifr/Il7r/Csf2rb2/Csf2rb/Il1rap/Ifnar2/Il10rb/Ifnar1/H2-Eb1/Cd74/Il15ra/Cd44/Il6ra/Fcgr1/Il11ra1/Lepr/Csf3r/Cmklr1/Il17rc/Il17ra/C3ar1/C5ar2/C5ar1/Fzd4/Il4ra/Il21r/Il10ra/Ctsh/Ccrl2/Cx3cr1/Ccr2/Ccr5/Il13ra1/Il2rg | 46 |
| MF | GO:0030246 | carbohydrate binding | 85/3915 | 256/17474 | 4,2945E-05 | 0,00071304 | 0,000531 | Col9a1/Prg4/Enpp1/P4ha1/Bsg/Galnt4/Ugp2/Canx/Galnt10/Asgr1/Clec10a/Vtn/Lgals9/Mrc2/Ogfod3/Egln3/Clec14a/Pygl/Lgals8/Ppp1r3g/Hk3/Lman2/Vcan/Hexb/Tkt/Galnt15/Lgals3/Cln5/Lgals1/Pfkm/Crybg3/Gbe1/Eva1c/H2-Eb1/C4b/Galnt14/Colec12/Galnt1/Lman1/Pygm/Ppp1r3c/Phyh/Mrc1/Eng/Cd302/Ly75/Pla2r1/Ganc/Gpcpd1/Cd93/Pygb/Ppp1r3d/Bcan/Agl/Alpk1/Manba/Frem1/Plod1/Fbxo2/Pgd/Slc2a5/Gusb/Plod3/Ptn/Clec5a/Hk2/Gfpt1/Clec4a3/Clec4a2/Clec2d/Clec1a/Clec7a/Lgals4/Siglecf/Cd33/Siglece/Gys1/P4ha3/Ppp1r3b/Ncan/Man2b1/Plod2/Clec3b/G6pdx/Prps2 | 85 |
| MF | GO:0050700 | CARD domain binding | 10/3915 | 13/17474 | 4,5762E-05 | 0,00074778 | 0,00055687 | Irgm1/Igtp/Irgm2/Card9/Mavs/Bcl10/Ripk2/Nod1/Nod2/Casp4 | 10 |
| MF | GO:0016829 | lyase activity | 67/3915 | 192/17474 | 4,913E-05 | 0,00079651 | 0,00059316 | Pm20d1/Npl/Echdc1/Amd1/Sgpl1/Ilvbl/Shmt2/Adcy1/Ddc/Ltc4s/Shmt1/Gucy2e/Aldoc/Car4/Dglucy/Pcbd2/Hacl1/Adcy4/Clybl/Ggact/Aco2/Csad/Pdxdc1/Ehhadh/Hacd2/Glo1/Cbs/Mocs1/Cyp1b1/Hsd17b4/Me2/Gldc/Gad2/Gad1/Ptgis/Car13/Car2/Gucy1a1/Car14/Etnppl/Kyat3/Cth/Car8/Aco1/Hacd4/Echdc2/Hmgcl/Hadha/Cd38/Paics/Sdsl/Hmgb1/Alox5ap/Tbxas1/Thnsl2/Ggcx/Dera/Echs1/Adcy7/Pts/Neil1/Ppcdc/Hacd3/Me1/Gadl1/Gucy2f/Car5b | 67 |
| MF | GO:0050660 | flavin adenine dinucleotide binding | 37/3915 | 90/17474 | 5,2731E-05 | 0,0008368 | 0,00062317 | Aox1/Aox3/Acadl/D2hgdh/Qsox1/Fmo1/Fmo2/Ddo/Mical1/Aifm2/Ilvbl/Acadvl/Nos2/Acox1/Nqo2/Kdm1b/Chdh/Aifm3/Prodh/Xdh/Sardh/Ivd/Sqor/Fmo5/Acadm/Nos3/Acox3/Acads/Txnrd3/Acadsb/Gcdh/Acad8/Etfa/Acad11/Cybb/Maoa/Maob | 37 |
| MF | GO:0008238 | exopeptidase activity | 38/3915 | 94/17474 | 6,5104E-05 | 0,00097858 | 0,00072875 | Lta4h/Metap2/Aebp1/Blmh/Scpep1/Scrn2/Ace/Adam17/Ctsl/Erap1/Mmp14/Cpq/Lnpep/Ermp1/Dpp7/Dpp4/Metap1d/Agbl2/Ctsa/Npepl1/Ctsz/Mindy1/Enpep/Pm20d2/Lap3/Agbl3/Anpep/Folh1/Ctsc/Prcp/Tpp1/Kdm8/Cpe/Naalad2/Dpp8/Mindy2/Adam10/Ctsh | 38 |
| MF | GO:0019199 | transmembrane receptor protein kinase activity | 31/3915 | 74/17474 | 0,00013718 | 0,00183168 | 0,00136405 | Nrp2/Erbb4/Ddr2/Erbb3/Egfr/Efemp1/Flt4/Ntrk2/Bmpr1a/Acvrl1/Pdgfrb/Csf1r/Mertk/Bmpr1b/Tgfbr1/Tek/Tie1/Fgfr3/Pdgfra/Kdr/Tgfbr3/Ephb4/Flt1/Ret/Axl/Igf1r/Fgfr2/Insr/Ephb1/Tgfbr2/Acvr2b | 31 |
| MF | GO:0008013 | beta-catenin binding | 38/3915 | 97/17474 | 0,00014347 | 0,00190644 | 0,00141972 | Sox17/Gja1/Ctnna3/Specc1l/Nos2/Axin2/Sox9/Numb/Gli3/Vcl/Amer2/Cxadr/Cd2ap/Ctnna1/Tcf4/Kank1/Tcf7l2/Ctnnd1/Csnk2a1/Ptprt/Foxo1/Med12l/Lef1/Klf4/Lzic/Ctnnbip1/Nos3/Tcf7l1/Tjp1/Sall1/Cdh5/Cdh1/Rora/Ctnnb1/Tbl1x/Amer1/Foxo4/Med12 | 38 |
| MF | GO:0003995 | acyl-CoA dehydrogenase activity | 9/3915 | 12/17474 | 0,00015863 | 0,00206897 | 0,00154076 | Acadl/Acadvl/Ivd/Acadm/Acads/Acadsb/Gcdh/Acad8/Acad11 | 9 |
| MF | GO:0052890 | oxidoreductase activity, acting on the CH-CH group of donors, with a flavin as acceptor | 9/3915 | 12/17474 | 0,00015863 | 0,00206897 | 0,00154076 | Acadl/Acadvl/Ivd/Acadm/Acads/Acadsb/Gcdh/Acad8/Acad11 | 9 |
| MF | GO:0004030 | aldehyde dehydrogenase [NAD(P)+] activity | 12/3915 | 19/17474 | 0,00015921 | 0,00206897 | 0,00154076 | Aldh9a1/Aldh1l2/Aldh3a2/Rdh11/Aldh7a1/Aldh3b1/Aldh1a1/Adh5/Aldh4a1/Aldh2/Aldh1l1/Aldh1a2 | 12 |
| MF | GO:0043028 | cysteine-type endopeptidase regulator activity involved in apoptotic process | 23/3915 | 50/17474 | 0,00018672 | 0,00233104 | 0,00173593 | Vil1/Rack1/Atp2a3/Serpinb9/Ctsl/Naip2/Naip5/Naip6/Tnfrsf10b/Tnfaip8/Ctsk/Ctss/Bcl10/Casp8ap2/Rps6ka1/Arrb1/Gas6/Mt3/Birc3/Rps27l/Ctsh/Xiap/Rps6ka3 | 23 |
| MF | GO:0043236 | laminin binding | 16/3915 | 30/17474 | 0,00021539 | 0,00265363 | 0,00197616 | Adgrg6/Ntn4/Nid1/Thbs4/Lgals3/Lgals1/Itga6/Thbs1/Ctss/Tinagl1/Bcam/Adam9/Plekha2/Itgb1/Itga9/Rpsa | 16 |
| MF | GO:0043177 | organic acid binding | 71/3915 | 216/17474 | 0,00024641 | 0,00295546 | 0,00220093 | Glul/Sesn1/Fabp7/P4ha1/Psap/Shmt2/Castor1/Ddc/Hba-a1/Shmt1/Slc46a1/Nos2/Acaca/Acox1/Ogfod3/Egln3/Pygl/Mtr/Aldh5a1/Dhfr/Hmgcs1/Glud1/Slc1a3/Rida/Prodh/Casr/Grik1/Hlcs/Cyp4f15/Cyp4f14/Ddah2/Gldc/Phyh/Gad2/Ptgds/Sardh/Ass1/Gad1/Mdk/Fabp5/Crabp2/Gstm7/Dbt/Adh5/Sh3glb1/Ddah1/Scp2/Hmgcl/Id3/Plod1/Pgd/Nos3/Acox3/Acacb/Plod3/Alox5ap/Ptn/Thnsl2/Gfpt1/Slc6a6/Apoc1/Siglecf/Cd33/Siglece/Stard5/P4ha3/Hbb-bt/Insr/Sesn3/Plod2/Rbp1 | 71 |
| MF | GO:0048029 | monosaccharide binding | 34/3915 | 86/17474 | 0,00025616 | 0,00304582 | 0,00226822 | P4ha1/Bsg/Ugp2/Asgr1/Clec10a/Ogfod3/Egln3/Pygl/Hk3/Lman2/Tkt/Lgals3/Cln5/Lgals1/Pfkm/Lman1/Phyh/Mrc1/Eng/Alpk1/Manba/Plod1/Slc2a5/Plod3/Hk2/Clec4a3/Clec4a2/Siglecf/Siglece/Gys1/P4ha3/Man2b1/Plod2/G6pdx | 34 |
| MF | GO:0016504 | peptidase activator activity | 23/3915 | 51/17474 | 0,00026854 | 0,0031359 | 0,0023353 | Cflar/Fn1/Vsir/Rack1/Nlrp1b/Atp2a3/Tifab/Ctsl/Tnfrsf10b/Fbln1/Tank/Ctsk/Ctss/Bcl10/Casp8ap2/Svbp/Pcolce/Cav1/Ctsc/Pycard/Clpx/Rps27l/Ctsh | 23 |
| MF | GO:0019209 | kinase activator activity | 46/3915 | 127/17474 | 0,00027045 | 0,00314024 | 0,00233854 | Ccnyl1/Mob3a/Igf1/Erbb3/Egfr/Irgm1/Igtp/Irgm2/Lgals9/Tom1l1/Fam20a/Gprc5c/Nek9/Pik3r1/Cab39l/Rgcc/Rictor/Dazap2/Nckap1l/Pak2/Itsn1/Cdkn1a/Trem2/Spdya/Dele1/Afap1l2/Map3k20/Pik3ca/Bcl10/Mob3b/Mob3c/P2rx7/Hmgb1/Mob1a/Wnk1/Tgfb1/Igf1r/Iqgap1/Gprc5b/Igf2/Insr/Gas6/Nrg1/Mt3/Rplp1/Fgf13 | 46 |
| MF | GO:0046943 | carboxylic acid transmembrane transporter activity | 55/3915 | 161/17474 | 0,00040871 | 0,00436877 | 0,00325342 | Slc16a9/Slc26a10/Slc1a4/Slc22a4/Slc47a1/Slc16a13/Slc13a5/Slc46a1/Abcc3/Slc38a6/Sfxn1/Slc38a9/Slc7a7/Abcc4/Slc1a3/Slc38a2/Grik1/Slc3a2/Slc16a12/Slc38a11/Slc43a3/Slc1a2/Slc13a3/Slc7a11/Slc16a1/Slc16a4/Slc26a7/Mfsd2a/Abcb1a/Abcb1b/Slc5a6/Abcg2/Sfxn5/Slc6a6/Slc6a11/Slc6a1/Slc25a18/Slc6a13/Slc6a12/Slco1c1/Slco1a4/Slc1a5/Ceacam1/Slc7a10/Slco2b1/Ucp2/Slc25a15/Slc7a2/Slc27a1/Slc10a7/Slc38a3/Slc26a6/Slc6a20a/Slc38a5/Slc16a2 | 55 |
| MF | GO:0035325 | Toll-like receptor binding | 9/3915 | 13/17474 | 0,00041258 | 0,0043874 | 0,0032673 | Ly96/Syk/Unc93b1/Tlr2/Tlr1/Tlr6/Ceacam1/Tirap/Myd88 | 9 |
| MF | GO:0140333 | glycerophospholipid flippase activity | 9/3915 | 13/17474 | 0,00041258 | 0,0043874 | 0,0032673 | Atp8b1/Abca4/Mfsd2a/Abcb1a/Abcb1b/Atp8a1/Atp10a/Atp11a/Atp11c | 9 |

| | | | | | | | | | |
|---|---|---|---|---|---|---|---|---|---|
| MF | GO:0008514 | organic anion transmembrane transporter activity | 73/3915 | 227/17474 | 0,00042228 | 0,00448475 | 0,00333979 | Gja1/Slc16a9/Slc26a10/Slc1a4/Slc22a4/Slc47a1/Slc16a13/Slc13a5/Slc46a1/Abcc3/Slc38a6/Sfxn1/Slc35d2/Slc38a9/Slc7a7/Abcc4/Slc1a3/Slc38a2/Abcc5/Grik1/Slc35b2/Slc25a23/Slc22a8/Slc22a6/Slc3a2/Slc16a12/Lrrc8a/Slc38a11/Slc43a3/Slc1a2/Slc52a3/Slc13a3/Slc2a10/Slco4a1/Slc7a11/Slc16a1/Slc16a4/Slc25a24/Slc39a8/Slc26a7/Slc35d1/Slc2a1/Mfsd2a/Abcb1a/Abcb1b/Slc5a6/Slc4a4/Abcg2/Sfxn5/Slc6a6/Slc6a11/Slc6a1/Slc25a18/Slc6a13/Slc6a12/Slco1c1/Slco1a4/Slc1a5/Ceacam1/Slc7a10/Slco2b1/Ucp2/Slc25a15/Slc7a2/Slc27a1/Slc10a7/Slc37a2/Slco2a1/Slc38a3/Slc26a6/Slc6a20a/Slc38a5/Slc16a2 | 73 |
| MF | GO:0031406 | carboxylic acid binding | 67/3915 | 206/17474 | 0,00050354 | 0,00524713 | 0,00390754 | Glul/Sesn1/Fabp7/P4ha1/Psap/Shmt2/Castor1/Ddc/Shmt1/Slc46a1/Nos2/Acaca/Acox1/Ogfod3/Egln3/Pygl/Mtr/Aldh5a1/Dhfr/Glud1/Slc1a3/Rida/Prodh/Casr/Grik1/Hlcs/Cyp4f15/Cyp4f14/Ddah2/Gldc/Phyh/Gad2/Ptgds/Sardh/Ass1/Gad1/Mdk/Fabp5/Crabp2/Gstm7/Dbt/Adh5/Sh3glb1/Ddah1/Scp2/Hmgcl/Id3/Plod1/Pgd/Nos3/Acox3/Acacb/Plod3/Alox5ap/Ptn/Thnsl2/Gfpt1/Apoc1/Siglecf/Cd33/Siglece/Stard5/P4ha3/Insr/Sesn3/Plod2/Rbp1 | 67 |
| MF | GO:0045028 | G protein-coupled purinergic nucleotide receptor activity | 8/3915 | 11/17474 | 0,00053697 | 0,00551128 | 0,00410425 | P2ry14/P2ry13/P2ry12/P2ry1/Ptafr/P2ry6/P2ry2/Gpr34 | 8 |
| MF | GO:0042887 | amide transmembrane transporter activity | 20/3915 | 44/17474 | 0,00058567 | 0,00589388 | 0,00438917 | Slc47a1/Slc46a1/Slc38a9/Slc1a3/Slc15a2/Grik1/Tap1/Tap2/Slc14a1/Slc15a3/Slc1a2/Slc13a3/Slc7a11/Slc5a6/Abcg2/Slc25a18/Slc27a1/Aqp9/Slc38a3/Slc38a5 | 20 |
| MF | GO:0030695 | GTPase regulator activity | 125/3915 | 429/17474 | 0,00060156 | 0,00600976 | 0,00447547 | Rgs20/Prex2/Arhgef4/Plcd4/Dock10/Srgap2/Dennd1b/Rgs5/Arhgap30/Arhgap18/Syde1/Arhgap45/Fgd6/Arhgap9/Tns3/Ccdc88a/Dock2/Arhgef15/Rap1gap2/Adap2/Tbc1d16/Arhgap5/Rgs6/Rin3/Eif5/Net1/Elmo1/Rasgrf2/Slc38a9/Arhgap22/Arhgef40/Spata13/Dock5/Tbc1d4/Farp1/Ranbp3l/Dennd3/Cyth4/Prr5/Tbc1d22a/Rapgef3/Nckap1l/Fgd4/Arhgap31/Itsn1/Fgd2/Rasal3/Dennd1c/Vav1/Ralbp1/Rasgrp3/Psd2/Arap3/Arhgef37/Sipa1/Rasgrp2/Rab3il1/Dock8/Plce1/Arhgap19/Usp6nl/Psd4/Garnl3/Gapvd1/Rapgef4/Rin2/Dennd2c/Rap1a/Gpsm2/Vav3/Arhgap29/Bcar3/Rgs3/Dennd4c/Dock7/Asap3/Arhgef19/Rgs12/Arap2/Arhgap24/Evi5/Git2/Rasa4/Stard13/Dennd11/Arhgef5/Chn2/Elmod3/Arhgap25/Fgd5/Arhgdib/Eps8l1/Plekhg2/Rinl/Ankrd27/Iqgap1/Arrb1/Arap1/Dennd5a/Sbf2/Arhgap17/Rgs10/Dock1/Eps8l2/Grtp1/Rasa3/Arhgef10/Dlc1/Fbxo8/Gmip/Elmod2/Arl2bp/Arhgap42/Dock6/Arhgef12/Dennd4a/Tbc1d2b/Rasa2/Als2cl/Rp2/Dock11/Arhgap4/Stard8/Tbc1d8b/Arhgap6 | 125 |
| MF | GO:0060589 | nucleoside-triphosphatase regulator activity | 125/3915 | 429/17474 | 0,00060156 | 0,00600976 | 0,00447547 | Rgs20/Prex2/Arhgef4/Plcd4/Dock10/Srgap2/Dennd1b/Rgs5/Arhgap30/Arhgap18/Syde1/Arhgap45/Fgd6/Arhgap9/Tns3/Ccdc88a/Dock2/Arhgef15/Rap1gap2/Adap2/Tbc1d16/Arhgap5/Rgs6/Rin3/Eif5/Net1/Elmo1/Rasgrf2/Slc38a9/Arhgap22/Arhgef40/Spata13/Dock5/Tbc1d4/Farp1/Ranbp3l/Dennd3/Cyth4/Prr5/Tbc1d22a/Rapgef3/Nckap1l/Fgd4/Arhgap31/Itsn1/Fgd2/Rasal3/Dennd1c/Vav1/Ralbp1/Rasgrp3/Psd2/Arap3/Arhgef37/Sipa1/Rasgrp2/Rab3il1/Dock8/Plce1/Arhgap19/Usp6nl/Psd4/Garnl3/Gapvd1/Rapgef4/Rin2/Dennd2c/Rap1a/Gpsm2/Vav3/Arhgap29/Bcar3/Rgs3/Dennd4c/Dock7/Asap3/Arhgef19/Rgs12/Arap2/Arhgap24/Evi5/Git2/Rasa4/Stard13/Dennd11/Arhgef5/Chn2/Elmod3/Arhgap25/Fgd5/Arhgdib/Eps8l1/Plekhg2/Rinl/Ankrd27/Iqgap1/Arrb1/Arap1/Dennd5a/Sbf2/Arhgap17/Rgs10/Dock1/Eps8l2/Grtp1/Rasa3/Arhgef10/Dlc1/Fbxo8/Gmip/Elmod2/Arl2bp/Arhgap42/Dock6/Arhgef12/Dennd4a/Tbc1d2b/Rasa2/Als2cl/Rp2/Dock11/Arhgap4/Stard8/Tbc1d8b/Arhgap6 | 125 |
| MF | GO:0051287 | NAD binding | 26/3915 | 63/17474 | 0,00060892 | 0,0060613 | 0,00451385 | Aox1/Aox3/Idh1/Aldh9a1/Aldh5a1/Nnt/Glud1/Ehhadh/Parp14/Me2/Aldh1a1/Phgdh/Hadh/Bdh2/Hadha/Ugdh/Aldh2/Hibadh/Ldhb/Sirt2/Idh2/Ctbp2/Sirt3/Hpgd/Me1/Hsd17b10 | 26 |
| MF | GO:0005126 | cytokine receptor binding | 75/3915 | 238/17474 | 0,00069006 | 0,00673524 | 0,00501573 | Stat1/Cflar/Casp8/Tmbim1/Cnih4/Tlr5/Kitl/Frs2/Ccdc88a/Tnfsf13/Tnfsf12/Cxcl16/Traf4/Ccl9/Ccl6/Stat3/Itgb3/Smurf2/Adam17/Snx6/Traf3/Siva1/Prl/Ripk1/Syk/Cxcl14/Erap1/Pik3r1/Il6st/Fermt2/Osmr/Lifr/Angpt1/Irak4/Il1rap/Cd2ap/Vegfa/Ticam2/Smad2/Vegfb/Jak2/Eng/Traf1/Traf6/Cd44/Spred1/Tnfsf10/Nes/Shc1/Il6ra/Csf1/Lyn/Nsmaf/Tgfbr1/Tgfbr3/Cxcl12/Ptpn6/Zfp110/Ceacam1/Tgfb1/Ctf1/Pycard/Angpt2/Casp3/Vegfc/Il15/Cdh5/Cklf/Il18/Nradd/Ccrl2/Tgfbr2/Myd88/Cx3cr1/Irak1 | 75 |
| MF | GO:0008483 | transaminase activity | 13/3915 | 24/17474 | 0,00069627 | 0,00676919 | 0,00504102 | Gfpt2/Gpt/Abat/Psat1/Accs/Tgm2/Etnppl/Kyat3/Gfpt1/Bcat2/Oat/Gpt2/Amt | 13 |
| MF | GO:0001784 | phosphotyrosine residue binding | 21/3915 | 48/17474 | 0,00080379 | 0,00769479 | 0,00573031 | Irs1/Grb10/Grap/Shc3/Syk/Pik3r1/Sla/Crkl/Vav1/Hck/Shc1/She/Bcar3/Pik3r3/Ldlrap1/Yes1/Sh3bp2/Ptpn11/Ptpn6/Mapk3/Plcg2 | 21 |
| MF | GO:0042910 | xenobiotic transmembrane transporter activity | 16/3915 | 33/17474 | 0,00086237 | 0,00815793 | 0,00607521 | Slc47a1/Slc46a1/Abcc3/Abca8a/Abcc4/Abcc5/Ralbp1/Slc43a3/Slc2a1/Abcb1a/Abcb1b/Abcg2/Slc6a6/Slc6a11/Slc6a13/Abcc6 | 16 |
| MF | GO:0038024 | cargo receptor activity | 31/3915 | 81/17474 | 0,00089924 | 0,00837586 | 0,0062375 | Prg4/Enpp1/Itgb2/Stab2/Asgr1/Cxcl16/Vtn/Lgals3bp/Sdc1/Mia2/Stab1/Scara5/Dab2/Tfrc/Colec12/Megf10/Mrc1/Cd44/Abca1/Lrp8/Scarb2/Loxl3/Tex261/Cd163/Lrp6/Lsr/Siglech/Lyve1/Itgam/Insr/Ldlr | 31 |
| MF | GO:0044548 | S100 protein binding | 9/3915 | 14/17474 | 0,00092515 | 0,0085876 | 0,00639518 | S100b/Ezr/Fgf1/Ahnak/S100a1/S100a6/S100a11/Iqgap1/Anxa2 | 9 |
| MF | GO:0005319 | lipid transporter activity | 55/3915 | 166/17474 | 0,00094051 | 0,00868127 | 0,00646494 | Esyt1/Gm2a/Pctp/Abcc3/Pitpnc1/Abca8a/Abca9/Abca6/Abcd4/Npc2/Abcc4/Abcd2/Ano6/Npc1/Stard4/Atp8b1/Prelid3a/Slc22a8/Slc22a6/Slc43a3/Pltp/Slco4a1/Fabp5/Atp11b/Abcd3/Abca4/Mttp/Ttpa/Abca1/Scp2/Osbpl9/Slc2a1/Mfsd2a/Xkr8/Abcb1a/Abcb1b/Atp8a1/Atp10d/Gltp/Osbpl3/Slco1c1/Slco1a4/Apoe/Ceacam1/Atp10a/Stard5/Slco2b1/Atp11a/Slc27a1/Slc10a7/Plscr1/Plscr2/Plscr4/Slco2a1/Atp11c | 55 |

GO_SO_protein

| ONTOLOGY | ID | Description | GeneRatio | BgRatio | pvalue | p.adjust | qvalue | geneID | Count |
|---|---|---|---|---|---|---|---|---|---|
| CC | GO:0043209 | myelin sheath | 60/1013 | 187/8928 | 1,47591E-14 | 8,55141E-11 | 7,89999E-11 | Plp1/Mobp/Cnp/Cldn11/Mog/Ermn/Nefm/Sirt2/Tspan2/Cryab/Mag/Myo1d/Nefl/Mbp/Pllp/Gjc3/Nfasc/Jam3/Ina/Gjc2/Ndrg1/Serinc5/Nefh/Llgl1/Septin4/Gsn/Cntn2/Phgdh/Gfap/Igsf8/Rdx/Pard3/Rap1a/Myh14/Gstm1/Tubb4a/Prdx1/Kcnj11/Cdc42/Glul/Ass1/Septin2/Scrib/Ckb/Mpdz/Tkt/Septin8/Omg/Cntnap1/Rala/Gdi2/Pals1/Fscn1/Msn/Gsto1/Rtn4/Actr1a/Cct3/Tcp1/Cct2 | 60 |
| CC | GO:0044304 | main axon | 32/1013 | 72/8928 | 1,13679E-12 | 3,29329E-09 | 3,04242E-09 | Ermn/Sirt2/Mag/Myo1d/Mbp/Nfasc/Hapln2/Gjc2/Cntn2/Sptbn4/Epb41l3/Ank3/Mapt/Pard3/Bin1/Kif13b/Kcnq3/Scn8a/Dlg2/Kcnq2/Tubb4a/Cntnap2/Kcna2/Kcna4/App/Trim46/Kcnj11/Kcna1/Cntnap1/Scn2a/Adam22/Map1b | 32 |
| CC | GO:0099513 | polymeric cytoskeletal fiber | 89/1013 | 432/8928 | 8,31741E-09 | 4,81911E-06 | 4,45201E-06 | Cnp/Cldn11/Nefm/Sirt2/Nefl/Jam3/Ina/Pls1/Ndrg1/Prph/Tppp3/Nefh/Synj2/Vim/Eml1/Dusp22/Rtn2/Sptbn4/Dst/Slain1/Gng12/Mapt/Gfap/Kif13b/Synm/Dcxr/Map4/Tuba8/Dync1li2/Mid1ip1/Reep3/Pbxip1/Shtn1/Tubb4a/Stmn1/Fez1/Dnm2/Dync1i2/Myo6/Rassf5/Kif5a/Dynlrb1/Gabarap/Map6/Jakmip1/Rusc1/Eml2/Slc8a3/Specc1/Pdlim4/Lmnb1/Septin2/Pdlim1/Dapk3/Yes1/Fer/Tpm4/Tubb3/Numa1/Clip2/Dync1h1/Kif5b/Rhoq/Zw10/Tcp11l1/Lmna/Apc2/Clasp2/Cep170b/Fsd1/Camsap3/Cep170/Eif6/Ckap5/Map1b/Kif21b/Fkbp4/Klc4/Dctn1/Cct3/Pafah1b1/Cdk5rap3/Dctn2/Tcp1/Eml6/Tsc1/Wdr44/Dync1i1/Cct2 | 89 |
| BP | GO:0007272 | ensheathment of neurons | 37/1013 | 106/8928 | 1,17856E-10 | 1,70715E-07 | 1,5771E-07 | Plp1/Cldn11/Sirt2/Tspan2/Mag/Bcas1/Mbp/Pllp/Gjc3/Nfasc/Jam3/Cd9/Ndrg1/Serinc5/Fa2h/Arhgef10/Enpp1/Cntn2/Qki/Epb41l3/Pard3/Abca2/Degs1/Cntnap2/Galc/Mtmr2/Lgi4/Kcnj10/Slc8a3/Mpdz/Omg/Cntnap1/Adam22/Pals1/Zpr1/Ckap5/Tsc1 | 37 |
| BP | GO:0008366 | axon ensheathment | 37/1013 | 106/8928 | 1,17856E-10 | 1,70715E-07 | 1,5771E-07 | Plp1/Cldn11/Sirt2/Tspan2/Mag/Bcas1/Mbp/Pllp/Gjc3/Nfasc/Jam3/Cd9/Ndrg1/Serinc5/Fa2h/Arhgef10/Enpp1/Cntn2/Qki/Epb41l3/Pard3/Abca2/Degs1/Cntnap2/Galc/Mtmr2/Lgi4/Kcnj10/Slc8a3/Mpdz/Omg/Cntnap1/Adam22/Pals1/Zpr1/Ckap5/Tsc1 | 37 |
| BP | GO:0042063 | gliogenesis | 57/1013 | 218/8928 | 6,66919E-10 | 6,44022E-07 | 5,94962E-07 | Plp1/Cnp/Opalin/Sirt2/Tspan2/Mag/Aspa/Dusp15/Cd9/Gjc2/Ndrg1/Fa2h/Il33/Arhgef10/Med12/Enpp2/Olig1/Enpp1/Vim/Cers2/Cntn2/Daam2/Qki/Phgdh/Gfap/Hdac11/Pard3/Rras2/Bin1/Abca2/Vcan/Ldlr/Rras/Cntnap2/Rela/Grin1/Hdac1/App/Abcc8/Lgi4/Kcnj10/Slc8a3/Scrib/Naglu/Omg/Cntnap1/Mapk3/Adam22/Pals1/Stat3/Rtn4/Ckap5/Cspg4/Ptpn11/Idh2/Pafah1b1/Atp1b2 | 57 |
| BP | GO:0000226 | microtubule cytoskeleton organization | 76/1013 | 390/8928 | 1,11014E-06 | 0,00049478 | 0,000457089 | Cnp/Nefm/Cryab/Nefl/Tppp3/Arhgef10/Nefh/Llgl1/Phldb1/Ccp110/Eml1/Cntn2/Map7d2/Dst/Slain1/Ank3/Stmn4/Dnaaf5/Mapt/Rock1/Pard3/Sbds/Trim36/Gpsm2/Map4/Rcc1/Tuba8/Dync1li2/Dixdc1/Cep97/Mid1ip1/Ttbk2/Tubb4a/Stmn1/Arl2/Trim46/Gabarap/Map6/Nubp1/Tacc1/Eml2/Tbcd/Mpdz/Chp1/Map7d1/Fer/Tubb3/Numa1/Kifbp/Clip2/Vps4b/Stag2/Dync1h1/Cdc42bpa/Bccip/Zpr1/Ift88/Zw10/Lmna/Apc2/Clasp2/Sik3/Fsd1/Chek1/Smc3/Camsap3/Smc1a/Usp33/Ckap5/Map1b/Fkbp4/Ccdc88a/Dctn1/Pafah1b1/Dctn2/Bicd2 | 76 |
| BP | GO:0006066 | alcohol metabolic process | 45/1013 | 197/8928 | 2,69009E-06 | 0,001039092 | 0,000959938 | Qdpr/Synj2/Enpp1/Nudt4/Asah2/Dhfr/Abhd4/Npc1/Abca2/Pecr/Hmgcs1/Dhcr24/Lipe/Cln8/Fgf1/Ldlr/Mvk/Galk1/Pmvk/Lss/Idh1/App/Ebp/Gpd1/Lbr/Sptlc1/Inppl1/Itpk1/Mvd/Adh5/G6pdx/Thtpa/Clcn2/Bpnt2/Asah1/Lmf1/Aplp2/Bpnt1/Plcd1/Fdps/Rdh11/Spr/Dpm1/Scp2/Idh2 | 45 |
| BP | GO:0044283 | small molecule biosynthetic process | 67/1013 | 340/8928 | 3,27413E-06 | 0,0011159 | 0,001030895 | Plp1/Fa2h/Qdpr/Gpt/Gamt/Psat1/Adi1/Qki/Ptgds/Carns1/Phgdh/Asah2/Gstp1/Dhfr/Scd2/Soga1/Ggt1/Fads6/Gstm7/Gstm3/Gatm/Abca2/Pecr/Degs1/Hmgcs1/Dhcr24/Nt5e/Stard3/Mid1ip1/Csad/Ilvbl/Gstm1/Fgf1/Mvk/Pmvk/Psph/Lss/Cbs/Ebp/Gpd1/Lbr/Sptlc1/Enoph1/Glul/Slc25a13/Ass1/Mvd/Acss2/Aldoc/G6pdx/Clcn2/Cad/Asah1/Mthfd1/Adal/Adk/Pgm2/Gsto1/Plcd1/Fdps/Acaca/Spr/Asns/Fasn/Scp2/Eif6/Mtap | 67 |
| BP | GO:0061564 | axon development | 71/1013 | 381/8928 | 1,29275E-05 | 0,003256593 | 0,003008517 | Plp1/Cnp/Nefm/Tspan2/Mag/Nefl/Mbp/Nfasc/Efnb3/Unc5b/Nefh/Llgl1/Enpp1/Vim/Cers2/Cntn2/Sptbn4/Ddr1/Epb41l3/Dst/Ank3/Mapt/Spart/Dhfr/Fgfr2/Kif13b/Dixdc1/Aplp1/Apbb2/Jak2/Shtn1/Stmn1/Plxnb3/Dnm2/Cntnap2/Grin1/App/Trim46/Spg11/Kif5a/Sema7a/Inpp5f/Map6/Rnd2/Zdhhc17/Rgma/Chl1/Tnr/Dip2b/Omg/Crtac1/Cntnap1/Tubb3/Kifbp/Vangl2/Zpr1/Aplp2/Kif5b/Rtn4/Clasp2/Rufy3/Ablim1/Usp33/Map1b/Ptpn11/Rab21/Mycbp2/Pafah1b1/Plxna1/Plxna4/Rtn4rl2 | 71 |
| BP | GO:0070507 | regulation of microtubule cytoskeleton organization | 31/1013 | 126/8928 | 2,05146E-05 | 0,004842731 | 0,004473827 | Phldb1/Slain1/Stmn4/Mapt/Rock1/Trim36/Gpsm2/Dixdc1/Cep97/Mid1ip1/Ttbk2/Tubb4a/Stmn1/Arl2/Map6/Eml2/Tbcd/Mpdz/Numa1/Vps4b/Dync1h1/Apc2/Clasp2/Fsd1/Camsap3/Ckap5/Map1b/Fkbp4/Dctn1/Pafah1b1/Bicd2 | 31 |
| BP | GO:0030705 | cytoskeleton-dependent intracellular transport | 38/1013 | 169/8928 | 2,32689E-05 | 0,004993323 | 0,004612948 | Nefm/Myo1d/Nefl/Nefh/Myo1e/Dst/Ank3/Mapt/Kif13b/Syt4/Dlg2/Fez1/Dync1i2/Myo6/Lamp1/App/Trim46/Spg11/Kif5a/Cdc42/Map6/Wdr35/Borcs6/Arhgap21/Dync1h1/Ift88/Kif5b/Ift22/Tmem201/Arl8a/Camsap3/Map1b/Rab21/Ap3b1/Ccdc88a/Pafah1b1/Bicd2/Dync1i1 | 38 |
| MF | GO:0015631 | tubulin binding | 52/1013 | 252/8928 | 1,09806E-05 | 0,002891883 | 0,002671588 | Sirt2/Cryab/Ndrg1/Tppp3/Eml1/Dst/Stmn4/Mapt/Sbds/Trim36/Kif13b/Gnas/Jakmip3/Map4/Dixdc1/Reep3/Syt11/Stmn1/Fez1/Dnm2/Pde4b/Trim46/Kif5a/Gabarap/Map6/Jakmip1/Fnta/Eml2/Cnn3/Tbcd/Dip2b/Chp1/Numa1/Clip2/Bccip/Kif5b/Apc2/Clasp2/Fsd1/Smc3/Camsap3/Map4k4/Vps41/Ckap5/Map1b/Kif21b/Ccdc88a/Dctn1/Pafah1b1/Vapb/Eml6/Dync1i1 | 52 |
| MF | GO:0016765 | transferase activity, transferring alkyl or aryl (other than methyl) groups | 15/1013 | 41/8928 | 2,18514E-05 | 0,004869508 | 0,004498565 | Gstp1/Gstm7/Gstm3/Gstm1/Cbs/Fnta/Gstm5/Fntb/Gstk1/Agps/Srm/Sms/Gsto1/Fdps/Mat2a | 15 |
| MF | GO:0005200 | structural constituent of cytoskeleton | 18/1013 | 57/8928 | 3,44496E-05 | 0,007128596 | 0,006585563 | Nefm/Nefl/Ina/Nefh/Vim/Epb41l3/Ank3/Gfap/Synm/Tuba8/Tubb4a/Tln1/Arpc1b/Lmnb1/Tubb3/Add3/Lmna/Ank2 | 18 |
| MF | GO:0016810 | hydrolase activity, acting on carbon-nitrogen (but not peptide) bonds | 21/1013 | 74/8928 | 4,76623E-05 | 0,008908245 | 0,008229644 | Sirt2/Aspa/Padi2/Acy1/Asah2/Hdac11/Ddah2/Rida/Hdac1/Asrgl1/Cad/Asah1/Amdhd2/Mthfd1/Adal/Atic/Ddah1/Pdf/Hint3/Dpysl4/Oplah | 21 |

GO_SP_protein

| ONTOLOGY | ID | Description | GeneRatio | BgRatio | pvalue | p.adjust | qvalue | geneID | Count |
|---|---|---|---|---|---|---|---|---|---|
| BP | GO:0006397 | mRNA processing | 261/1488 | 377/8928 | 1,1E-120 | 5,9E-117 | 5,3E-117 | Khdrbs3/Hnrnpl/Rbmx/Celf2/Srsf3/Rbfox1/Cpsf7/Rbfox3/Ddx5/Rnmt/Rbm14/Celf1/Hnrnpa0/Hnrnpa3/Hnrnpa2b1/Tra2a/Rbm42/Sugp2/Srsf7/U2af2/Hnrnpll/Snrpd1/Hnrnpa1/Dus3l/Dhx9/Ddx23/Khsrp/Ptbp2/U2af1/Prpf40a/Hnrnpc/Sfswap/Sf3b1/Snrnp70/Hnrnpk/Sfpq/Snrpd2/Npm1/Xrn2/Srsf12/Srsf1/Snrpa/Nudt21/Ccar2/Pabpn1/Cdk13/Snu13/Magoh/Snrnp40/Srsf6/Snrpa1/Zmat2/Psip1/Sart1/Ncl/Srsf10/Rbm25/Pnn/Srsf2/Srsf5/Usp39/Snrpb/Snrnp200/Ddx41/Sltm/Plrg1/Srrm2/Hnrnpu/Tra2b/Pqbp1/Cdc5l/Khdrbs1/Dhx15/Ddx39b/Rbm15b/Wbp11/Sf3a1/Rbm39/Prpf8/Hnrnpm/Prpf40b/Ramac/Rbm15/Nova2/Srrt/Tardbp/Prpf6/Son/Prpf19/Htatsf1/Thoc2/Scaf1/Rnps1/Rbm5/Zc3h14/Snw1/Rbm26/Ncbp1/Smu1/Rbm8a/Sf3a3/Sf3b2/Snrpe/Eftud2/Prpf3/Nono/Alyref/Dbr1/Thoc1/Snrpd3/Prpf31/Tcerg1/Eif4a3/Sugp1/Sart3/Rprd1a/Mbnl2/Snrpb2/Celf5/Snrpg/Safb/Ddx17/Zcchc8/Larp7/Raly/Rbm6/Ctnnbl1/Thoc6/Fip1l1/Sf3b3/Luc7l3/Prpf4/Luc7l/Rbm27/Thrap3/Rbmxl1/Sf3b4/Rbm10/Acin1/Wtap/Cpsf6/Ddx46/Cstf3/Rbm17/Thoc5/Nup98/Prpf4b/Mtrex/Rprd2/Isy1/Sympk/Ik/Sf1/Phf5a/Srsf4/Srrm1/Cdc40/Sf3a2/Hnrnph1/Dhx38/Rbm19/Sap18/Fmr1/Cdc73/Ppp1r8/Bcas2/Bud31/Cpsf1/Ythdc1/Cpsf4/Snrpf/Ptbp3/Syncrip/Habp4/Snrnp27/Cdk9/Celf4/Cstf1/Zranb2/Gpkow/Puf60/Gemin8/Xab2/Cmtr1/Cpsf2/Coil/Rbm22/Mbnl1/Smndc1/Tia1/Paf1/Scaf8/Khdrbs2/Cirbp/Rbfox2/Tfip11/Frg1/Akap8l/Wdr33/Dazap1/Aqr/Ncbp2/Iws1/Rprd1b/Rbm28/Fxr2/Ddx20/Zcrb1/Arl6ip4/Rbm3/Phrf1/Adar/Elavl4/Rbbp6/Alkbh5/Sf3b6/Gemin4/Gemin5/Cd2bp2/Mettl3/Hnrnpf/Fxr1/Cwc25/Safb2/Srek1/Crnkl1/Cpsf3/Lsm7/Gemin7/Sf3b5/Adarb1/Paxbp1/Lsm6/Cwc22/Ddx39a/Rngtt/Ddx1/Leo1/U2af1l4/Celf6/Srpk1/Thoc3/Zc3h13/Dhx8/Pabpc1/Fam172a/Cnot6l/Ddx47/Cwf19l1/Lsm8/Dhx16/Lsm2/Virma/Ppie/Dhx36/Rbm4b/Cstf2t/Cpeb3/Cstf2/Srpk2/Tdrd3 | 261 |
| BP | GO:0022613 | ribonucleoprotein complex biogenesis | 175/1488 | 318/8928 | 3,2E-57 | 2,23E-54 | 1,99E-54 | Fbll1/Rbmx/Celf2/Pelp1/Nol9/Cpsf7/Wdr18/Celf1/Snrpd1/Dhx9/Ptbp2/Sfswap/Sf3b1/Snrpd2/Npm1/Srsf12/Srsf1/Nop56/Nop58/Dkc1/Nudt21/Las1l/Snu13/Riox1/Srsf6/Rpl38/Psip1/Sart1/Nifk/Ncl/Srsf10/Usp39/Snrpb/Snrnp200/Ddx39b/Gar1/Wbp11/Sf3a1/Nat10/Prpf8/Ramac/Mrto4/Prpf6/Prpf19/Rps25/Rbm5/Ncbp1/Sf3a3/Snrpe/Prpf3/Krr1/Fbl/Snrpd3/Prpf31/Rps19/Sart3/Celf5/Rps14/Snrpg/Rpl14/Ddx17/Ythdf2/Luc7l3/Luc7l/Nhp2/Rps9/Rps7/Bysl/Rbm10/Cpsf6/Rpl7a/Rpl23a/Pes1/Rpl6/Mtrex/Isy1/Rps28/Rpl35/Rpl7/Rps8/Rpl13a/Rpl7l1/Sf3a2/Rps16/Rpl27/Rpl11/Fau/Cdc73/Ythdc1/Rps6/Rps27/Snrpf/Rpl10/Rpl24/Celf4/Rpf2/Ddx51/Rps5/Puf60/Nup88/Rpl35a/Gemin8/Xab2/Rpsa/Coil/Rpl5/Mbnl1/Rplp0/Utp14a/Frg1/Bop1/Wdr3/Ebna1bp2/Nop16/Mettl16/Rpl26/Nvl/Mybbp1a/Gtpbp4/Rps17/Ddx20/Nip7/Heatr1/Wdr43/Adar/Rps27l/Brix1/Rps15/Utp6/Nol11/Rrp12/Gemin4/Rps21/Gemin5/Mak16/Exosc10/Crnkl1/Nop2/Gnl3l/Gemin7/Pdcd11/Rpp30/Lsm6/Mdn1/Utp18/Celf6/Srpk1/Noc4l/Urb1/Ftsj3/Ddx47/Eif3a/Wdr36/Dis3/Dhx30/Ago2/Rrs1/Lsm2/Surf6/Nol6/Ddx3x/Exosc4/Rrp7a/Rrp9/Ago1/Ddx21/Wdr12/Exosc1/Eif3d/Exosc8/Eif2s3x/Eif6/Ruvbl1/Ruvbl2/Srpk2 | 175 |
| BP | GO:0022618 | ribonucleoprotein complex assembly | 97/1488 | 152/8928 | 2,13E-39 | 7,42E-37 | 6,63E-37 | Rbmx/Celf2/Cpsf7/Celf1/Snrpd1/Dhx9/Ptbp2/Sfswap/Sf3b1/Snrpd2/Srsf12/Srsf1/Nudt21/Snu13/Srsf6/Rpl38/Psip1/Srsf10/Usp39/Snrpb/Snrnp200/Ddx39b/Sf3a1/Prpf8/Ramac/Mrto4/Prpf6/Prpf19/Rps25/Rbm5/Ncbp1/Sf3a3/Snrpe/Prpf3/Snrpd3/Prpf31/Rps19/Sart3/Celf5/Rps14/Snrpg/Luc7l3/Luc7l/Cpsf6/Rpl23a/Rpl6/Isy1/Rps28/Rpl13a/Sf3a2/Rpl11/Fau/Cdc73/Ythdc1/Rps6/Rps27/Snrpf/Rpl10/Rpl24/Celf4/Rpf2/Rps5/Puf60/Gemin8/Xab2/Rpsa/Coil/Rpl5/Mbnl1/Rplp0/Bop1/Ddx20/Adar/Rps27l/Brix1/Rps15/Gemin4/Gemin5/Crnkl1/Nop2/Gemin7/Mdn1/Celf6/Srpk1/Eif3a/Dhx30/Ago2/Rrs1/Lsm2/Rrp7a/Ago1/Eif3d/Eif2s3x/Eif6/Ruvbl1/Ruvbl2/Srpk2 | 97 |
| BP | GO:0002181 | cytoplasmic translation | 85/1488 | 122/8928 | 4,26E-39 | 1,32E-36 | 1,18E-36 | Dhx9/Rpl22l1/Rps26/Rpl38/Hnrnpu/Hnrnpd/Rps10/Rps25/Ncbp1/Rps19/Rps14/Rpl14/Rps13/Rpl18/Rps15a/Ythdf2/Rps4x/Rps11/Rpl15/Rps9/Rps7/Rps23/Rpl7a/Rps3/Rpl23a/Rpl18a/Rps20/Rpl6/Rpl39/Rpl37a/Rps18/Rpl27a/Rps28/Rpl31/Rpl7/Rpl4/Rps8/Rpl13a/Rpl22/Rps16/Rpl12/Rpl27/Fmr1/Rpl13/Rpl11/Fau/Rps2/Rps6/Rpl17/Rps12/Syncrip/Rpl8/Rpl34/Rpl24/Rpl9/Rpl21/Rps5/Rpl35a/Rpl10a/Rpsa/Rpl5/Rpl3/Rpl19/Rplp0/Rpl23/Rplp2/Ncbp2/Rpl26/Rps17/Rpl36/Cpeb4/Rpl28/Rps15/Rps21/Mettl3/Drg1/Eif3a/Eif3i/Dhx36/Eif3d/Eif4a2/Zc3h15/Cpeb3/Eif2s3x/Csde1 | 85 |
| BP | GO:0071826 | ribonucleoprotein complex subunit organization | 99/1488 | 159/8928 | 8,06E-39 | 2,36E-36 | 2,11E-36 | Rbmx/Celf2/Cpsf7/Celf1/Snrpd1/Dhx9/Ptbp2/Sfswap/Sf3b1/Snrpd2/Srsf12/Srsf1/Nudt21/Snu13/Srsf6/Rpl38/Psip1/Srsf10/Usp39/Snrpb/Snrnp200/Ddx39b/Sf3a1/Prpf8/Ramac/Mrto4/Prpf6/Prpf19/Rps25/Rbm5/Ncbp1/Sf3a3/Snrpe/Prpf3/Snrpd3/Prpf31/Rps19/Sart3/Celf5/Rps14/Snrpg/Luc7l3/Luc7l/Cpsf6/Rpl23a/Rpl6/Isy1/Rps28/Rpl13a/Sf3a2/Rpl11/Fau/Cdc73/Ythdc1/Rps6/Rps27/Snrpf/Rpl10/Rpl24/Celf4/Rpf2/Rps5/Puf60/Gemin8/Xab2/Rpsa/Coil/Rpl5/Mbnl1/Rplp0/Tfip11/Bop1/Ddx20/Adar/Rps27l/Brix1/Rps15/Gemin4/Gemin5/Crnkl1/Nop2/Gemin7/Mdn1/Celf6/Srpk1/Dhx8/Eif3a/Dhx30/Ago2/Rrs1/Lsm2/Rrp7a/Ago1/Eif3d/Eif2s3x/Eif6/Ruvbl1/Ruvbl2/Srpk2 | 99 |
| BP | GO:0006325 | chromatin organization | 144/1488 | 330/8928 | 5,15E-32 | 1,25E-29 | 1,11E-29 | Cenpv/Gatad2b/Smarcb1/Hmg20a/Dpf1/Cbx5/Satb2/Trim28/Smarca4/Smarcc2/Smarcd3/Ddx23/Sf3b1/Npm1/Mta2/Smarce1/Arid1b/Smarcd1/Rnf2/Dpf2/Chd3/Hdac2/Riox1/Brd4/Tasor/Smchd1/Hnrnpu/Mta1/Cfdp1/Rbm15b/Bcl7a/Mecp2/Atrx/Rbm15/Ep400/Lmnb2/Kdm6a/Mbd3/Cbx3/Hira/Spin1/Actl6b/Set/Arid1a/Sart3/Sin3a/Tlk1/Macroh2a1/Chd5/Rbbp5/Tpr/Rbbp7/Cbx1/Mta3/Rbbp4/Hp1bp3/Smarca2/Cbx6/Kdm1a/Suz12/Actr8/Trrap/Wdr5/Top1/Rybp/Ythdc1/Chd4/Ssrp1/Kmt2a/Dcaf1/Brd9/Arid2/Brd1/Sgf29/Samd1/Dmap1/Smarca5/Ash2l/Meaf6/Chd6/Ubr5/Ncor1/Hmgn3/Hmgn5/Mybbp1a/Macroh2a2/Kdm5a/Hcfc1/Kdm5b/Msl2/Mbd2/Brd7/Rnf20/Epc1/Wac/Kat5/Tbl1xr1/Vps72/Rsf1/Kmt2d/Yeats2/Lmna/Ctr9/Epc2/Exosc10/Mettl3/Pbrm1/Nsd3/Ubn1/Rnf40/Yeats4/Nucks1/Kdm2a/Setd1a/Ing2/Chd2/Morc2a/Zmynd11/Baz1b/Yy1/Bptf/Morf4l1/Lmnb1/Fam172a/Brd3/Mcm3ap/Pwwp3a/Nr3c1/Hdac3/Dek/Ddx21/Usp22/Phf2/Selenof/Setdb1/Trip12/Ruvbl1/Nfat5/Ruvbl2/Tspyl4/Srpk2/Tdrd3/Usp7/Ogt | 144 |
| BP | GO:0016072 | rRNA metabolic process | 95/1488 | 170/8928 | 7,37E-32 | 1,71E-29 | 1,53E-29 | Fbll1/Pelp1/Nol9/Smarcb1/Wdr18/Smarca4/Npm1/Dkc1/Las1l/Snu13/Sart1/Nifk/Ncl/Gar1/Trir/Wbp11/Nat10/Mrto4/Rps25/Krr1/Spin1/Fbl/Rps19/Rps14/Rpl14/Macroh2a1/Ddx17/Ythdf2/Nhp2/Rps7/Bysl/Pes1/Mtrex/Tcof1/Rps28/Rpl35/Rpl7/Rps8/Rpl7l1/Rps16/Top1/Rpl27/Rpl11/Rps6/Rps27/Rpf2/Ddx51/Rpl35a/Ubtf/Rpl5/Utp14a/Frg1/Bop1/Wdr3/Ebna1bp2/Mettl16/Rpl26/Nvl/Gtpbp4/Rps17/Macroh2a2/Heatr1/Wdr43/Brix1/Rps15/Utp6/Nol11/Rrp12/Rps21/Mars1/Mak16/Exosc10/Nop2/Ercc3/Pdcd11/Rpp30/Lsm6/Utp18/Urb1/Ftsj3/Ddx47/Wdr36/Dis3/Rrs1/Nol6/Exosc4/Rrp7a/Rrp9/Ddx21/Wdr12/Exosc1/Exosc8/Eif6/Gtf3c1/Xrn1 | 95 |
| BP | GO:0006403 | RNA localization | 80/1488 | 134/8928 | 1,06E-29 | 1,9E-27 | 1,7E-27 | Srsf3/Poldip3/Hnrnpa3/Hnrnpa2b1/Srsf7/Hnrnpa1/Hnrnpab/Dhx9/Khsrp/Npm1/Srsf1/Nop58/Dkc1/Sarnp/Pabpn1/Magoh/Fyttd1/Nup210/Pom121/Chtop/Nup35/G3bp2/Hnrnpu/Khdrbs1/Zc3h11a/Ddx39b/Rbm15b/Ssb/Prpf6/Thoc2/Ncbp1/Rbm8a/Alyref/Nxf1/Fbl/Thoc1/Ahctf1/Eif4a3/Nup153/Nup188/Senp2/Thoc6/Tpr/Ranbp2/Cpsf6/Thoc5/Nup98/Nup43/Nup160/Fmr1/Ythdc1/Nup107/Nup133/Nup88/Nup155/Nup37/Akap8l/Nup93/Nup85/Dcp2/Ncbp2/Iws1/Nup214/Stau1/Alkbh5/Nup54/Phax/Exosc10/Nup62/Aaas/Ddx39a/Thoc3/Yy1/Mcm3ap/Seh1l/Rae1/Nol6/Dhx36/Sec13/Fubp3 | 80 |
| BP | GO:0051028 | mRNA transport | 61/1488 | 90/8928 | 2,68E-27 | 4,14E-25 | 3,7E-25 | Srsf3/Poldip3/Hnrnpa3/Hnrnpa2b1/Srsf7/Hnrnpa1/Dhx9/Khsrp/Srsf1/Sarnp/Pabpn1/Magoh/Fyttd1/Nup210/Pom121/Chtop/Nup35/G3bp2/Zc3h11a/Ddx39b/Rbm15b/Thoc2/Ncbp1/Rbm8a/Alyref/Nxf1/Thoc1/Ahctf1/Eif4a3/Nup188/Senp2/Thoc6/Tpr/Ranbp2/Thoc5/Nup98/Nup43/Nup160/Fmr1/Ythdc1/Nup107/Nup133/Nup88/Nup155/Nup37/Akap8l/Nup93/Nup85/Ncbp2/Iws1/Nup214/Alkbh5/Nup54/Nup62/Aaas/Ddx39a/Thoc3/Mcm3ap/Seh1l/Rae1/Sec13 | 61 |

| | | | | | | | | | |
|---|---|---|---|---|---|---|---|---|---|
| BP | GO:0016570 | histone modification | 107/1488 | 243/8928 | 2,53E-24 | 2,88E-22 | 2,57E-22 | Fbll1/Gatad2b/Smarcb1/Smarca4/Sf3b1/Sin3b/Sfpq/Mta2/Rnf2/Chd3/Phf20l1/Hdac2/Riox1/Brd4/Chtop/Atf2/Mta1/Mecp2/Ep400/Kdm6a/Mbd3/Snw1/Rcor3/Actl6b/Fbl/Sart3/Sin3a/Taf5l/Chd5/Sap30bp/Rbbp5/Sf3b3/Rbbp7/Mta3/Rbbp4/Kdm1a/Suz12/Trrap/Wdr5/Cdc73/Chd4/Cdk9/Kmt2a/Sgf29/Rtf1/Dmap1/Smarca5/Supt7l/Ash2l/Paf1/Meaf6/Akap8l/Ubr5/Ncor1/Iws1/Mybbp1a/Kdm5a/Hcfc1/Kdm5b/Msl2/Mbd2/Rnf20/Epc1/Wac/Kat5/Glyr1/Tbl1xr1/Vps72/Kmt2d/Yeats2/Ctr9/Epc2/Nsd3/Rnf40/Yeats4/Crebbp/Sf3b5/Kdm2a/Setd1a/Ncor2/Paxbp1/Ing2/Wdr82/Leo1/Baz1b/Trerf1/Supt3/Morf4l1/Akap8/Taf6l/Taf4/Nos1/Pbxip1/Hdac3/Dek/Ddx21/Usp22/Prmt1/Setdb1/Trip12/Ruvbl1/Naa50/Skic8/Ruvbl2/Ufl1/Usp7/Ogt | 107 |
| BP | GO:0010608 | post-transcriptional regulation of gene expression | 126/1488 | 313/8928 | 3,85E-24 | 4,29E-22 | 3,83E-22 | Celf2/Matr3/Nolc1/Nr2f1/Poldip3/Celf1/Elavl1/Hnrnpa0/Hnrnpr/Fus/Dus3l/Dhx9/Khsrp/Hnrnpc/Ilf3/Npm1/Srsf1/Dkc1/Nudt21/Sarnp/Magoh/Rpl38/Ncl/Hnrnpu/Hnrnpd/Khdrbs1/Ddx39b/Lsm14b/Nat10/Mecp2/Ssb/Tardbp/Zc3h14/Ncbp1/Zc3h18/Eif4a3/Pum2/Ythdf2/Tpr/Thrap3/Rbm10/Rps3/Tcof1/Pum1/Polr2g/Rpl13a/Rack1/Fmr1/Ythdc1/Syncrip/Larp4b/Rpl10/Tial1/Habp4/Caprin1/Helz/Rpl5/Tia1/Cirbp/Larp4/Dcp1a/Ythdf1/Dcp2/Ncbp2/Fxr2/Mettl16/Rpl26/Apex1/Taf15/Pum3/Tmed2/Rbm3/Ythdf3/Cpeb4/Adar/Elavl4/Rps27l/Alkbh5/Eif4g3/Gemin5/Phax/Mettl3/Fxr1/Upf1/Calr/Dnajc3/Pabpc4/Tnrc6b/Gigyf2/Pym1/Pabpc1/Cnot6l/Eif3k/Cnot11/Pcif1/Lsm14a/Ddx6/Ago2/Cnot2/Ddx3x/Eif3b/Secisbp2l/Eif3e/Ago1/Eif4g1/Tnrc6c/Dhx36/Rbm4b/Eif3d/Eif4g2/Exosc8/Cpeb3/Cnot10/Zc3h7b/Cnot7/Cnot1/Eif6/Cnot3/Prkca/Xrn1/Eif5b/Gcn1/Ptk2b/Eif2s1/Csde1/Cyfip1 | 126 |
| BP | GO:0006401 | RNA catabolic process | 85/1488 | 183/8928 | 2,17E-21 | 2,12E-19 | 1,9E-19 | Ddx5/Celf1/Elavl1/Hnrnpa0/Hnrnpr/Hnrnpab/Fus/Dhx9/Khsrp/Hnrnpc/Npm1/Xrn2/Srsf1/Dkc1/Magoh/Hnrnpu/Hnrnpd/Trir/Mrto4/Ssb/Tardbp/Zc3h4/Rnps1/Zc3h14/Ncbp1/Rbm8a/Zc3h18/Eif4a3/Pum2/Ythdf2/Thrap3/Rbm10/Snd1/Mtrex/Edc4/Pum1/Polr2g/Fmr1/Syncrip/Larp4b/Cirbp/Dcp1a/Ythdf1/Dcp2/Ncbp2/Fxr2/Mettl16/Apex1/Taf15/Ythdf3/Elavl4/Alkbh5/Phax/Exosc10/Mettl3/Fxr1/Upf1/Lsm7/Lsm6/Wdr82/Pabpc4/Tnrc6b/Gigyf2/Pym1/Pabpc1/Cnot6l/Dis3/Ago2/Cnot2/Lsm2/Exosc4/Eif3e/Ago1/Tnrc6c/Dhx36/Exosc1/Edc3/Exosc8/Cpeb3/Cnot10/Cnot7/Cnot1/Cnot3/Xrn1/Csde1 | 85 |
| BP | GO:0006406 | mRNA export from nucleus | 37/1488 | 49/8928 | 1,28E-19 | 1,13E-17 | 1,01E-17 | Srsf3/Poldip3/Hnrnpa2b1/Sarnp/Pabpn1/Magoh/Fyttd1/Chtop/Zc3h11a/Ddx39b/Rbm15b/Thoc2/Ncbp1/Rbm8a/Alyref/Nxf1/Thoc1/Eif4a3/Thoc6/Tpr/Thoc5/Nup160/Ythdc1/Nup107/Nup133/Nup88/Nup155/Akap8l/Nup93/Nup85/Iws1/Nup214/Alkbh5/Ddx39a/Thoc3/Mcm3ap/Rae1 | 37 |
| BP | GO:0051168 | nuclear export | 64/1488 | 125/8928 | 3,93E-19 | 3,15E-17 | 2,81E-17 | Srsf3/Poldip3/Atxn1/Hnrnpa2b1/Dhx9/Npm1/Sarnp/Pabpn1/Magoh/Fyttd1/Pom121/Chtop/Khdrbs1/Zc3h11a/Ddx39b/Rbm15b/Ssb/Thoc2/Ncbp1/Rbm8a/Alyref/Nxf1/Thoc1/Eif4a3/Nup153/Nup188/Thoc6/Tpr/Ranbp2/Rbm10/Cpsf6/Thoc5/Nup98/Nup160/Ythdc1/Nup107/Nup133/Nup88/Nup155/Rbm22/Akap8l/Nup93/Nup85/Ncbp2/Iws1/Nup214/Adar/Alkbh5/Rps15/Phax/Ranbp3/Nup62/Calr/Mdn1/Ddx39a/Thoc3/Mcm3ap/Rae1/Rrs1/Nol6/Hdac3/Rangap1/Nemf/Eif6 | 64 |
| BP | GO:0031047 | RNA-mediated gene silencing | 42/1488 | 66/8928 | 1,25E-17 | 8,93E-16 | 7,97E-16 | Srsf3/Cenpv/Nr2f1/Ddx5/Elavl1/Hnrnpa2b1/Dhx9/Mecp2/Srrt/Ncbp1/Pum2/Ddx17/Snd1/Pum1/Fmr1/Tial1/Nup155/Helz/Ncbp2/Rbm3/Adar/Mettl3/Fxr1/Prkra/Tnrc6b/Pabpc1/Fam172a/Cnot6l/Cnot11/Ddx6/Ago2/Cnot2/Ddx3x/Ago1/Eif4g1/Tnrc6c/Cnot10/Zc3h7b/Cnot7/Cnot1/Eif6/Cnot3 | 42 |
| BP | GO:2000112 | regulation of cellular macromolecule biosynthetic process | 107/1488 | 304/8928 | 1,19E-15 | 6,91E-14 | 6,17E-14 | Nolc1/Poldip3/Celf1/Elavl1/Hnrnpr/Dus3l/Dhx9/Khsrp/Ilf3/Npm1/Sarnp/Magoh/Rpl38/Ncl/Hnrnpu/Hnrnpd/Khdrbs1/Ddx39b/Lsm14b/Nat10/Golga2/Ssb/Tardbp/Tm9sf2/Ncbp1/Eif4a3/Pum2/Ythdf2/Tpr/Rps3/Tcof1/Pum1/Polr2g/Rpl13a/Rack1/Fmr1/Syncrip/Larp4b/Rpl10/Habp4/Caprin1/Rpl5/Tia1/Cirbp/Larp4/Dcp1a/Ythdf1/Dcp2/Ncbp2/Fxr2/Mettl16/Rpl26/Pum3/Tmed2/Rbm3/Ythdf3/Cpeb4/Elavl4/Rps27l/Eif4g3/Gemin5/Mettl3/Fxr1/Upf1/Calr/Dnajc3/Tnrc6b/Gigyf2/Dyrk2/Pym1/Pabpc1/Cnot6l/Eif3k/Cnot11/Pcif1/Lsm14a/Ddx6/Ago2/Cnot2/Ddx3x/Eif3b/Secisbp2l/Eif3e/Ago1/Eif4g1/Tnrc6c/Dhx36/Rbm4b/Eif3d/Eif4g2/Rabl3/Exosc8/Cpeb3/Cnot10/Cnot7/Cnot1/Eif6/Cnot3/Prkca/Xrn1/Eif5b/Gcn1/Ptk2b/Itm2c/Eif2s1/Csde1/Cyfip1 | 107 |
| BP | GO:0034248 | regulation of amide metabolic process | 106/1488 | 303/8928 | 2,63E-15 | 1,46E-13 | 1,31E-13 | Nolc1/Poldip3/Celf1/Elavl1/Hnrnpr/Dus3l/Dhx9/Khsrp/Ilf3/Npm1/Sarnp/Magoh/Rpl38/Ncl/Hnrnpu/Hnrnpd/Khdrbs1/Ddx39b/Lsm14b/Nat10/Ssb/Tardbp/Ncbp1/Eif4a3/Pum2/Ythdf2/Tpr/Rps3/Tcof1/Pum1/Sorl1/Polr2g/Rpl13a/Rack1/Fmr1/Syncrip/Larp4b/Rpl10/Habp4/Caprin1/Rpl5/Tmed10/Tia1/Cirbp/Larp4/Dcp1a/Ythdf1/Dcp2/Ncbp2/Fxr2/Mettl16/Rpl26/Pum3/Tmed2/Rbm3/Ythdf3/Cpeb4/Elavl4/Rps27l/Eif4g3/Gemin5/Mettl3/Fxr1/Upf1/Calr/Dnajc3/Samd8/Tnrc6b/Gigyf2/Pym1/Pabpc1/Cnot6l/Eif3k/Cnot11/Pcif1/Lsm14a/Ddx6/Ago2/Cnot2/Ddx3x/Eif3b/Secisbp2l/Eif3e/Ago1/Eif4g1/Tnrc6c/Dhx36/Rbm4b/Eif3d/Eif4g2/Exosc8/Cpeb3/Cnot10/Cnot7/Cnot1/Eif6/Cnot3/Prkca/Xrn1/Eif5b/Gcn1/Ptk2b/Eif2s1/Ntrk2/Csde1/Cyfip1 | 106 |
| BP | GO:0006974 | cellular response to DNA damage stimulus | 141/1488 | 458/8928 | 1,15E-14 | 6,1E-13 | 5,45E-13 | Smarcb1/Ddx5/Dpf1/Cbx5/Trim28/Smarca4/Fus/Smarcc2/Smarcd3/Dhx9/Gtf2h1/Wrnip1/Hnrnpk/Sfpq/Ppp1r10/Npm1/Xrn2/Bclaf1/Smarce1/Arid1b/Smarcd1/Dpf2/Ccar2/Pogz/Brd4/Atf2/Smchd1/Mta1/Cdc5l/Ddx39b/Bcl7a/Atrx/Ep400/Prpf19/Gtf2h3/Snw1/Cbx3/H2ax/Parp1/Nono/Actl6b/Thoc1/Arid1a/Cdkn2aip/Taf5l/Tlk1/Rbbp5/Sf3b3/Cbx1/Smarca2/Rps3/Kdm1a/Thoc5/Mtrex/Actr8/Ints7/Trrap/Ccnk/Fmr1/Chd4/Ssrp1/Hdgfl2/Cdk9/Zmynd8/Xab2/Arid2/Sgf29/Usp10/Ints3/Dmap1/Terf2/Smarcal1/Smarca5/Supt7l/Ash2l/Sfr1/Meaf6/Ubr5/Fxr2/Rpl26/Pds5b/Apex1/Ppp4r3a/Brd7/Rbbp6/Rps27l/Epc1/Wac/Kat5/Vps72/Epc2/Mettl3/Fxr1/Pbrm1/Upf1/Terf2ip/Ercc4/Msh2/Yeats4/Nucks1/Ercc3/Sf3b5/Kdm2a/Setd1a/Ppp4r3b/Pds5a/Chd2/Nabp2/Xrcc1/Ddx1/Hmga1/Clock/Morc2a/Baz1b/Yy1/Lig3/Usp45/Supt3/Morf4l1/Gigyf2/Dyrk2/Taf6l/Taf4/Msh6/Eef1e1/Pwwp3a/Polr2i/Dek/Usp22/Ctc1/Smc1a/Smc3/Nudt16l1/Stn1/Trip12/Ruvbl1/Nfat5/Ruvbl2/Ufl1/Ppp2r5c/Usp7 | 141 |
| BP | GO:2001020 | regulation of response to DNA damage stimulus | 74/1488 | 185/8928 | 1,97E-14 | 1,03E-12 | 9,18E-13 | Smarcb1/Ddx5/Dpf1/Trim28/Smarca4/Fus/Smarcc2/Smarcd3/Dhx9/Hnrnpk/Ppp1r10/Npm1/Bclaf1/Smarce1/Arid1b/Smarcd1/Dpf2/Ccar2/Pogz/Brd4/Smchd1/Ddx39b/Bcl7a/Ep400/Parp1/Actl6b/Thoc1/Arid1a/Taf5l/Sf3b3/Smarca2/Rps3/Kdm1a/Thoc5/Actr8/Trrap/Fmr1/Hdgfl2/Cdk9/Arid2/Sgf29/Dmap1/Smarca5/Supt7l/Meaf6/Ubr5/Fxr2/Rpl26/Brd7/Epc1/Kat5/Vps72/Epc2/Fxr1/Pbrm1/Terf2ip/Ercc4/Yeats4/Sf3b5/Ppp4r3b/Xrcc1/Baz1b/Yy1/Supt3/Morf4l1/Taf6l/Taf4/Eef1e1/Dek/Usp22/Nudt16l1/Trip12/Ruvbl1/Ruvbl2 | 74 |
| BP | GO:0000122 | negative regulation of transcription by RNA polymerase II | 120/1488 | 380/8928 | 1,78E-13 | 8,32E-12 | 7,43E-12 | Zeb2/Tcf4/Gatad2b/Hmg20a/Nr2f1/Ddx5/Atxn1/Cbx5/Satb2/Hnrnpa2b1/Trim28/Phf14/Smarca4/Hnrnpab/Smarcc2/Sin3b/Hnrnpk/Myef2/Sfpq/Mta2/Bcl11a/Rnf2/Foxg1/Mtdh/Chd3/Sarnp/Hdac2/Sap130/Atf2/Ylpm1/Hnrnpu/Mta1/Khdrbs1/Mecp2/Rbm15/Mbd3/Snw1/Cbx3/Parp1/Arid1a/Tcerg1/Sin3a/Rps14/Macroh2a1/Chd5/Larp7/Tpr/Rbbp7/Rbm10/Cbx1/Mta3/Rbbp4/Cbx6/Kdm1a/Suz12/Drap1/Pou3f1/Rybp/Cdc73/Chd4/Esrra/Rpl10/Irf2bpl/Zmynd8/Zbtb18/Dcaf1/Tle1/Mypop/Rtf1/Samd1/Dmap1/Maz/Pcbp3/Paf1/Scaf8/Pias1/Rpl23/Spen/Ncor1/Srebf2/Mepce/Ddx20/Macroh2a2/Pura/Kdm5a/Hcfc1/Hexim1/Naca/Mbd2/Epc1/Kat5/Tbl1xr1/Cic/Thap11/Yeats2/Ctr9/Nfib/Hdgf/Purb/Crebbp/Ncor2/Ing2/Calr/Mef2a/Nfix/Yy1/Bptf/Cry2/Zbtb20/Med25/Cnot2/Nr3c1/Hdac3/Mlip/Cpeb3/Setdb1/Cnot1/Tcf25/Sdcbp/Ogt | 120 |
| BP | GO:0033044 | regulation of chromosome organization | 61/1488 | 146/8928 | 4,14E-13 | 1,86E-11 | 1,66E-11 | Cenpv/Smarcb1/Dpf1/Hnrnpa2b1/Trim28/Smarca4/Smarcc2/Smarcd3/Hnrnpc/Sfpq/Ppp1r10/Smarce1/Arid1b/Smarcd1/Dkc1/Dpf2/Tasor/Ylpm1/Hnrnpu/Hnrnpd/Bcl7a/Nat10/Atrx/Bub3/Parp1/Actl6b/Arid1a/Sin3a/Macroh2a1/Tpr/Smarca2/Actr8/Ik/Arid2/Terf2/Sub1/Smarca5/Dcp2/Mad1l1/Brd7/Mad2l1/Kat5/Lmna/Exosc10/Pbrm1/Upf1/Terf2ip/Ercc4/Gnl3l/Nabp2/Xrcc1/Morc2a/Baz1b/Numa1/Yy1/Dhx36/Ctc1/Setdb1/Stn1/Ruvbl1/Ruvbl2 | 61 |
| BP | GO:0018205 | peptidyl-lysine modification | 75/1488 | 200/8928 | 6,75E-13 | 2,91E-11 | 2,6E-11 | Smarcb1/Hmg20a/Trim28/Smarca4/Sf3b1/Bcl11a/Senp3/Phf20l1/Hdac2/Brd4/Atf2/Mecp2/Ep400/Mbd3/Snw1/Actl6b/Senp7/Sin3a/Taf5l/Chd5/Rbbp5/Senp2/Sf3b3/Kdm1a/Suz12/Trrap/Wdr5/Kmt2a/Sgf29/Rtf1/Dmap1/Smarca5/Supt7l/Ash2l/Pias1/Meaf6/Iws1/Mybbp1a/Hcfc1/Msl2/Epc1/Kat5/Glyr1/Vps72/Sumo1/Kmt2d/Yeats2/Ctr9/Epc2/Gnl3l/Sae1/Yeats4/Crebbp/Sf3b5/Setd1a/Wdr82/Baz1b/Supt3/Morf4l1/Taf6l/Taf4/Nos1/Pbxip1/Zmiz1/Dek/Ddx21/Usp22/Setd3/Rangap1/Setdb1/Ruvbl1/Naa50/Skic8/Ruvbl2/Ogt | 75 |
| BP | GO:0071824 | protein-DNA complex subunit organization | 44/1488 | 89/8928 | 7,26E-13 | 3,07E-11 | 2,74E-11 | Tcf4/Cenpv/Smarcb1/Smarca4/Smarcc2/Smarcd3/Npm1/Smarce1/Smarcd1/Pogz/Atrx/Hira/Set/Arid1a/Sart3/Macroh2a1/Rbbp4/Hp1bp3/Smarca2/Creb1/Gtf2b/Ssrp1/Tbp/Med24/Arid2/Med17/Smarca5/Pias1/Rpl23/Macroh2a2/Rsf1/Gtf2f2/Med6/Ubn1/Med23/Chd2/Med14/Taf1/Taf6l/Taf4/Mcm3ap/Med25/Med16/Tspyl4 | 44 |

| | | | | | | | | | |
|---|---|---|---|---|---|---|---|---|---|
| BP | GO:0044270 | cellular nitrogen compound catabolic process | 91/1488 | 265/8928 | 9,16E-13 | 3,83E-11 | 3,42E-11 | Ddx5/Celf1/Elavl1/Hnrnpa0/Hnrnpr/Hnrnpab/Fus/Dhx9/Khsrp/Hnrnpc/Npm1/Xrn2/Srsf1/Dkc1/Magoh/Hnrnpu/Hnrnpd/Trir/Entpd6/Mrto4/Ssb/Tardbp/Zc3h4/Rnps1/Zc3h14/Ncbp1/Rbm8a/Zc3h18/Eif4a3/Pum2/Ythdf2/Thrap3/Rbm10/Snd1/Mtrex/Nmnat1/Edc4/Pum1/Polr2g/Fmr1/Syncrip/Larp4b/Cirbp/Dcp1a/Ythdf1/Dcp2/Ncbp2/Fxr2/Mettl16/Apex1/Taf15/Ythdf3/Elavl4/Alkbh5/Phax/Exosc10/Mettl3/Fxr1/Upf1/Lsm7/Lsm6/Wdr82/Pabpc4/Cyp3a13/Pde5a/Tnrc6b/Gigyf2/Pym1/Pabpc1/Cnot6l/Dis3/Ago2/Cnot2/Lsm2/Exosc4/Eif3e/Ago1/Tnrc6c/Dhx36/Exosc1/Edc3/Exosc8/Cpeb3/Cnot10/Cnot7/Entpd5/Cnot1/Cnot3/Fitm2/Xrn1/Csde1 | 91 |
| BP | GO:0019827 | stem cell population maintenance | 46/1488 | 97/8928 | 1,47E-12 | 6,08E-11 | 5,43E-11 | Smarcb1/Elavl1/Smarca4/Smarce1/Smarcd1/Dpf2/Hdac2/Sap130/Bcl7a/Srrt/Actl6b/Arid1a/Sin3a/Taf5l/Rbbp7/Rbbp4/Smarca2/Cdc73/Med24/Brd9/Rtf1/Med17/Paf1/Wdr43/Vps72/Ctr9/Med6/Mettl3/Nfib/Crebbp/Setd1a/Ing2/Med14/Leo1/Nfix/Zc3h13/Bcl9/Taf6l/Ddx6/Cnot2/Smc1a/Smc3/Stag2/Cnot1/Cnot3/Ogt | 46 |
| BP | GO:0042255 | ribosome assembly | 32/1488 | 54/8928 | 1,66E-12 | 6,75E-11 | 6,03E-11 | Rpl38/Mrto4/Rps25/Rps19/Rps14/Rpl23a/Rpl6/Rps28/Rpl11/Fau/Rps6/Rps27/Rpl10/Rpl24/Rpf2/Rps5/Rpsa/Rpl5/Rplp0/Bop1/Nip7/Rps27l/Brix1/Rps15/Nop2/Mdn1/Dhx30/Rrs1/Surf6/Ddx3x/Rrp7a/Eif6 | 32 |
| BP | GO:0046700 | heterocycle catabolic process | 90/1488 | 264/8928 | 1,96E-12 | 7,8E-11 | 6,97E-11 | Ddx5/Celf1/Elavl1/Hnrnpa0/Hnrnpr/Hnrnpab/Fus/Dhx9/Khsrp/Hnrnpc/Npm1/Xrn2/Srsf1/Dkc1/Magoh/Hnrnpu/Hnrnpd/Trir/Entpd6/Mrto4/Ssb/Tardbp/Zc3h4/Rnps1/Zc3h14/Ncbp1/Rbm8a/Zc3h18/Eif4a3/Pum2/Ythdf2/Thrap3/Rbm10/Snd1/Mtrex/Nmnat1/Edc4/Pum1/Polr2g/Fmr1/Syncrip/Larp4b/Cirbp/Dcp1a/Ythdf1/Dcp2/Ncbp2/Fxr2/Mettl16/Apex1/Taf15/Ythdf3/Elavl4/Alkbh5/Phax/Exosc10/Mettl3/Fxr1/Upf1/Lsm7/Lsm6/Wdr82/Pabpc4/Pde5a/Tnrc6b/Gigyf2/Pym1/Pabpc1/Cnot6l/Dis3/Ago2/Cnot2/Lsm2/Exosc4/Eif3e/Ago1/Tnrc6c/Dhx36/Exosc1/Edc3/Exosc8/Cpeb3/Cnot10/Cnot7/Entpd5/Cnot1/Cnot3/Fitm2/Xrn1/Csde1 | 90 |
| BP | GO:0098727 | maintenance of cell number | 46/1488 | 98/8928 | 2,34E-12 | 9,13E-11 | 8,15E-11 | Smarcb1/Elavl1/Smarca4/Smarce1/Smarcd1/Dpf2/Hdac2/Sap130/Bcl7a/Srrt/Actl6b/Arid1a/Sin3a/Taf5l/Rbbp7/Rbbp4/Smarca2/Cdc73/Med24/Brd9/Rtf1/Med17/Paf1/Wdr43/Vps72/Ctr9/Med6/Mettl3/Nfib/Crebbp/Setd1a/Ing2/Med14/Leo1/Nfix/Zc3h13/Bcl9/Taf6l/Ddx6/Cnot2/Smc1a/Smc3/Stag2/Cnot1/Cnot3/Ogt | 46 |
| BP | GO:0019439 | aromatic compound catabolic process | 92/1488 | 273/8928 | 2,34E-12 | 9,13E-11 | 8,15E-11 | Ddx5/Celf1/Elavl1/Hnrnpa0/Hnrnpr/Hnrnpab/Fus/Dhx9/Khsrp/Hnrnpc/Npm1/Xrn2/Srsf1/Dkc1/Magoh/Hnrnpu/Hnrnpd/Trir/Entpd6/Mrto4/Ssb/Tardbp/Zc3h4/Rnps1/Zc3h14/Ncbp1/Rbm8a/Zc3h18/Eif4a3/Pum2/Ythdf2/Thrap3/Rbm10/Snd1/Mtrex/Nmnat1/Edc4/Pum1/Polr2g/Fmr1/Syncrip/Larp4b/Cirbp/Dcp1a/Ythdf1/Dcp2/Ncbp2/Fxr2/Mettl16/Apex1/Taf15/Ythdf3/Elavl4/Alkbh5/Phax/Exosc10/Mettl3/Fxr1/Upf1/Lsm7/Lsm6/Wdr82/Pabpc4/Pde5a/Tnrc6b/Gigyf2/Pym1/Pabpc1/Cnot6l/Dis3/Moxd1/Ago2/Cnot2/Lsm2/Exosc4/Eif3e/Ago1/Tnrc6c/Dhx36/Exosc1/Edc3/Exosc8/Cpeb3/Cnot10/Cnot7/Entpd5/Cnot1/Cnot3/Fitm2/Xrn1/Ephx1/Csde1 | 92 |
| BP | GO:0140694 | non-membrane-bounded organelle assembly | 84/1488 | 250/8928 | 2,6E-11 | 8,99E-10 | 8,02E-10 | Ccsap/Rbm14/Npm1/Hdac2/Rpl38/Pogz/Tasor/G3bp2/Hnrnpu/Mdm1/Pqbp1/Golga2/Mrto4/Rps25/Map9/Rps19/Rps14/Pum2/Ythdf2/Tpr/Rps23/Rps3/Rpl23a/Rpl6/Rps28/Rpl11/Fau/Rps6/Gtf2b/Rps27/Rpl10/Rpl24/Rpf2/Rps5/Ubap2l/Rpsa/Rpl5/Tia1/Cirbp/Rplp0/Atxn2l/Ythdf1/Ncor1/Bop1/Nip7/Ythdf3/G3bp1/Rps27l/Brix1/Rps15/Nop2/Nup62/Mdn1/Aaas/Prrc2c/Mef2a/Numa1/Atxn2/Cnot6l/Drg1/Stag1/Lsm14a/Dhx30/Ddx6/Cnot2/Rrs1/Surf6/Ddx3x/Hdac3/Rrp7a/Edc3/Smc1a/Cds2/Smc3/Cnot7/Stag2/Cnot1/Eif6/Ripor2/Prkar1a/Fitm2/Rcc1/Eif2s1/Csde1 | 84 |
| BP | GO:2000036 | regulation of stem cell population maintenance | 25/1488 | 39/8928 | 4,05E-11 | 1,35E-09 | 1,2E-09 | Smarcb1/Elavl1/Smarca4/Smarce1/Smarcd1/Dpf2/Hdac2/Sap130/Bcl7a/Actl6b/Arid1a/Sin3a/Taf5l/Rbbp7/Rbbp4/Smarca2/Brd9/Wdr43/Ing2/Zc3h13/Taf6l/Cnot2/Cnot1/Cnot3/Ogt | 25 |
| BP | GO:1901361 | organic cyclic compound catabolic process | 91/1488 | 282/8928 | 4,66E-11 | 1,53E-09 | 1,36E-09 | Ddx5/Celf1/Elavl1/Hnrnpa0/Hnrnpr/Hnrnpab/Fus/Dhx9/Khsrp/Hnrnpc/Npm1/Xrn2/Srsf1/Dkc1/Magoh/Hnrnpu/Hnrnpd/Trir/Entpd6/Mrto4/Ssb/Tardbp/Zc3h4/Rnps1/Zc3h14/Ncbp1/Rbm8a/Zc3h18/Eif4a3/Pum2/Ythdf2/Thrap3/Rbm10/Snd1/Mtrex/Nmnat1/Edc4/Pum1/Polr2g/Fmr1/Syncrip/Larp4b/Cirbp/Dcp1a/Ythdf1/Dcp2/Ncbp2/Fxr2/Mettl16/Apex1/Taf15/Ythdf3/Elavl4/Alkbh5/Phax/Exosc10/Mettl3/Fxr1/Upf1/Lsm7/Lsm6/Wdr82/Pabpc4/Pde5a/Tnrc6b/Gigyf2/Pym1/Pabpc1/Cnot6l/Dis3/Moxd1/Ago2/Cnot2/Lsm2/Exosc4/Eif3e/Ago1/Tnrc6c/Dhx36/Exosc1/Edc3/Exosc8/Cpeb3/Cnot10/Cnot7/Entpd5/Cnot1/Cnot3/Fitm2/Xrn1/Csde1 | 91 |
| BP | GO:0034063 | stress granule assembly | 19/1488 | 25/8928 | 9,51E-11 | 3,04E-09 | 2,72E-09 | G3bp2/Pqbp1/Pum2/Ythdf2/Rps23/Ubap2l/Tia1/Cirbp/Atxn2l/Ythdf1/Ythdf3/G3bp1/Prrc2c/Atxn2/Lsm14a/Ddx6/Ddx3x/Eif2s1/Csde1 | 19 |
| BP | GO:0000819 | sister chromatid segregation | 52/1488 | 138/8928 | 1,97E-09 | 5,41E-08 | 4,83E-08 | Ccsap/Smarcb1/Dpf1/Smarca4/Smarcc2/Smarcd3/Smarce1/Arid1b/Smarcd1/Dpf2/Pogz/Tasor/Hnrnpu/Bcl7a/Golga2/Atrx/Bub3/Map9/Actl6b/Arid1a/Champ1/Macroh2a1/Tpr/Smarca2/Ik/Top2b/Gtf2b/Arid2/Smarca5/Akap8l/Mad1l1/Pds5b/Brd7/Mad2l1/Kat5/Pbrm1/Nup62/Pds5a/Aaas/Baz1b/Numa1/Akap8/Drg1/Stag1/Seh1l/Lsm14a/Rrs1/Smc1a/Smc3/Stag2/Ripor2/Naa50 | 52 |
| BP | GO:0065004 | protein-DNA complex assembly | 34/1488 | 73/8928 | 2,24E-09 | 6,07E-08 | 5,41E-08 | Tcf4/Cenpv/Smarca4/Npm1/Pogz/Atrx/Hira/Set/Sart3/Macroh2a1/Rbbp4/Hp1bp3/Smarca2/Creb1/Gtf2b/Ssrp1/Tbp/Med24/Med17/Smarca5/Pias1/Macroh2a2/Rsf1/Gtf2f2/Med6/Ubn1/Med23/Med14/Taf1/Taf6l/Taf4/Med25/Med16/Tspyl4 | 34 |
| BP | GO:0007059 | chromosome segregation | 70/1488 | 218/8928 | 1,06E-08 | 2,71E-07 | 2,42E-07 | Ccsap/Smarcb1/Dpf1/Smarca4/Smarcc2/Smarcd3/Smarce1/Arid1b/Smarcd1/Dpf2/Pogz/Brd4/Tasor/Hnrnpu/Bcl7a/Golga2/Atrx/Sun1/Bub3/Map9/Actl6b/Arid1a/Champ1/Tlk1/Macroh2a1/Pum2/Tpr/Smarca2/Rps3/Nup43/Pum1/Ik/Top1/Top2b/Rcc2/Gtf2b/Arid2/Nup37/Smarca5/Akap8l/Ncor1/Mad1l1/Pds5b/Brd7/Mad2l1/Kat5/Pbrm1/Ercc4/Nup62/Pds5a/Aaas/Baz1b/Srpk1/Numa1/Akap8/Drg1/Stag1/Seh1l/Lsm14a/Rrs1/Ddx3x/Hdac3/Smc1a/Smc3/Stag2/Ripor2/Naa50/Rgs14/Rcc1/Top3b | 70 |
| BP | GO:0043414 | macromolecule methylation | 57/1488 | 164/8928 | 1,08E-08 | 2,74E-07 | 2,44E-07 | Fbll1/Smarcb1/Trim28/Smarca4/Ilf3/Mta2/Hdac2/Snrpb/Chtop/Rbm15b/Mecp2/Ramac/Rbm15/Snw1/Parp1/Fbl/Snrpd3/Chd5/Larp7/Rbbp5/Wtap/Kdm1a/Suz12/Wdr5/Kmt2a/Trmt10b/Cmtr1/Rtf1/Smarca5/Ash2l/Paf1/Fam98b/Iws1/Mettl16/Mepce/Hcfc1/Trmt61a/Rnf20/Kmt2d/Ctr9/Mettl3/Nsd3/Nop2/Trmt6/Setd1a/Paxbp1/Wdr82/Trmt1l/Zc3h13/Ftsj3/Pcif1/Virma/Setd3/Prmt1/Setdb1/Skic8/Ogt | 57 |
| BP | GO:0006888 | endoplasmic reticulum to Golgi vesicle-mediated transport | 40/1488 | 100/8928 | 2,06E-08 | 5,06E-07 | 4,51E-07 | Tmed7/Golga2/Sorl1/Lman2l/Yipf4/Mia3/Tmed10/Ergic1/Ergic2/Tmed9/Tmed2/Lman2/Tmed4/Lman1/Ergic3/Ero1b/Gosr1/Hyou1/Golt1b/Sec16a/Erp29/Copa/Trip11/Preb/Arcn1/Copg2/Uso1/Copb1/Copb2/Cope/Sec24b/Sec13/Copg1/Sec23a/Sec31a/Sec24c/Vamp7/Scfd1/Stx18/Sec24a | 40 |
| BP | GO:0098732 | macromolecule deacylation | 35/1488 | 84/8928 | 4,63E-08 | 1,07E-06 | 9,52E-07 | Gatad2b/Sin3b/Sfpq/Mta2/Chd3/Ccar2/Hdac2/Mta1/Mbd3/Rcor3/Sin3a/Sap30bp/Abhd13/Rbbp7/Mta3/Rbbp4/Wdr5/Chd4/Sgf29/Smarca5/Akap8l/Ncor1/Ndst1/Mbd2/Tbl1xr1/Yeats2/Ncor2/Ing2/Trerf1/Morf4l1/Akap8/Ppt1/Hdac3/Abhd12/Fry | 35 |
| BP | GO:0006999 | nuclear pore organization | 11/1488 | 13/8928 | 1,5E-07 | 3,18E-06 | 2,84E-06 | Pom121/Nup35/Ahctf1/Nup153/Nup98/Nup107/Nup133/Nup205/Nup93/Nup54/Seh1l | 11 |
| BP | GO:0098813 | nuclear chromosome segregation | 55/1488 | 170/8928 | 3,05E-07 | 6,1E-06 | 5,44E-06 | Ccsap/Smarcb1/Dpf1/Smarca4/Smarcc2/Smarcd3/Smarce1/Arid1b/Smarcd1/Dpf2/Pogz/Tasor/Hnrnpu/Bcl7a/Golga2/Atrx/Sun1/Bub3/Map9/Actl6b/Arid1a/Champ1/Macroh2a1/Tpr/Smarca2/Ik/Top2b/Rcc2/Gtf2b/Arid2/Smarca5/Akap8l/Mad1l1/Pds5b/Brd7/Mad2l1/Kat5/Pbrm1/Ercc4/Nup62/Pds5a/Aaas/Baz1b/Numa1/Akap8/Drg1/Stag1/Seh1l/Lsm14a/Rrs1/Smc1a/Smc3/Stag2/Ripor2/Naa50 | 55 |
| BP | GO:0070316 | regulation of G0 to G1 transition | 16/1488 | 26/8928 | 3,26E-07 | 6,47E-06 | 5,78E-06 | Smarcb1/Dpf1/Smarca4/Smarcc2/Smarcd3/Smarce1/Arid1b/Smarcd1/Dpf2/Bcl7a/Actl6b/Arid1a/Smarca2/Arid2/Brd7/Pbrm1 | 16 |
| BP | GO:0031507 | heterochromatin formation | 26/1488 | 58/8928 | 4,46E-07 | 8,65E-06 | 7,72E-06 | Cenpv/Trim28/Smarca4/Tasor/Smchd1/Mecp2/Atrx/Lmnb2/Mbd3/Cbx3/Sin3a/Tpr/Kdm1a/Suz12/Samd1/Smarca5/Kdm5a/Mbd2/Kmt2d/Lmna/Ctr9/Morc2a/Lmnb1/Fam172a/Phf2/Setdb1 | 26 |

| | | | | | | | | | |
|---|---|---|---|---|---|---|---|---|---|
| BP | GO:0019080 | viral gene expression | 23/1488 | 48/8928 | 4,73E-07 | 9,09E-06 | 8,11E-06 | Smarcb1/Smarca4/Dhx9/Brd4/Ssb/Tardbp/Snw1/Larp7/Gtf2b/Cdk9/Pcbp2/Hexim1/Rsf1/Nucks1/Ccnt2/Eif3l/Eif3a/Eif3f/Spcs1/Spcs3/Eif3b/Eif3d/Csde1 | 23 |
| BP | GO:0032259 | methylation | 59/1488 | 190/8928 | 5,52E-07 | 1,04E-05 | 9,31E-06 | Fbll1/Smarcb1/Rnmt/Trim28/Smarca4/Ilf3/Mta2/Hdac2/Snrpb/Chtop/Rbm15b/Mecp2/Ramac/Rbm15/Snw1/Parp1/Fbl/Snrpd3/Chd5/Larp7/Rbbp5/Wtap/Kdm1a/Suz12/Wdr5/Kmt2a/Trmt10b/Cmtr1/Rtf1/Smarca5/Ash2l/Paf1/Fam98b/Prdm10/Iws1/Mettl16/Mepce/Hcfc1/Trmt61a/Rnf20/Kmt2d/Ctr9/Mettl3/Nsd3/Nop2/Trmt6/Setd1a/Paxbp1/Wdr82/Trmt1l/Zc3h13/Ftsj3/Pcif1/Virma/Setd3/Prmt1/Setdb1/Skic8/Ogt | 59 |
| BP | GO:0045023 | G0 to G1 transition | 16/1488 | 27/8928 | 6,77E-07 | 1,26E-05 | 1,13E-05 | Smarcb1/Dpf1/Smarca4/Smarcc2/Smarcd3/Smarce1/Arid1b/Smarcd1/Dpf2/Bcl7a/Actl6b/Arid1a/Smarca2/Arid2/Brd7/Pbrm1 | 16 |
| BP | GO:0140747 | regulation of ncRNA transcription | 12/1488 | 17/8928 | 1,2E-06 | 2,13E-05 | 1,9E-05 | Smarcb1/Smarca4/Ncl/Atrx/Zc3h4/Macroh2a1/Larp7/Mepce/Macroh2a2/Nol11/Mars1/Wdr82 | 12 |
| BP | GO:0006486 | protein glycosylation | 37/1488 | 103/8928 | 1,64E-06 | 2,88E-05 | 2,57E-05 | Galnt17/Galnt16/Golga2/Galnt9/B3galt6/Fut8/St8sia3/St6galnac5/Alg10b/Large1/Tusc3/Rpn2/Stt3a/Extl2/Tmem165/Dpy19l3/Ddost/Ube2j1/Rpn1/Tmtc3/Tmtc1/Mlec/Galnt2/Mogs/Stt3b/Pomgnt2/Trip11/Poglut1/B4gat1/Eogt/Fut11/Alg2/Pofut2/Entpd5/Uggt1/Pmm1/Ogt | 37 |
| BP | GO:0043413 | macromolecule glycosylation | 37/1488 | 103/8928 | 1,64E-06 | 2,88E-05 | 2,57E-05 | Galnt17/Galnt16/Golga2/Galnt9/B3galt6/Fut8/St8sia3/St6galnac5/Alg10b/Large1/Tusc3/Rpn2/Stt3a/Extl2/Tmem165/Dpy19l3/Ddost/Ube2j1/Rpn1/Tmtc3/Tmtc1/Mlec/Galnt2/Mogs/Stt3b/Pomgnt2/Trip11/Poglut1/B4gat1/Eogt/Fut11/Alg2/Pofut2/Entpd5/Uggt1/Pmm1/Ogt | 37 |
| BP | GO:0051983 | regulation of chromosome segregation | 27/1488 | 65/8928 | 1,7E-06 | 2,96E-05 | 2,64E-05 | Smarcb1/Dpf1/Smarca4/Smarcc2/Smarcd3/Smarce1/Arid1b/Smarcd1/Dpf2/Hnrnpu/Bcl7a/Bub3/Actl6b/Arid1a/Pum2/Tpr/Smarca2/Pum1/Ik/Rcc2/Arid2/Mad1l1/Brd7/Mad2l1/Kat5/Pbrm1/Numa1 | 27 |
| BP | GO:0032986 | protein-DNA complex disassembly | 10/1488 | 13/8928 | 2,83E-06 | 4,77E-05 | 4,26E-05 | Smarcb1/Smarca4/Smarcc2/Smarcd3/Smarce1/Smarcd1/Arid1a/Ssrp1/Arid2/Rpl23 | 10 |
| BP | GO:0070085 | glycosylation | 39/1488 | 116/8928 | 5,65E-06 | 8,99E-05 | 8,02E-05 | Galnt17/Galnt16/Golga2/Galnt9/B3galt6/Fut8/St8sia3/St6galnac5/Alg10b/Large1/Tusc3/Rpn2/Stt3a/Extl2/Tmem165/Dpy19l3/Ddost/B4galnt1/Ube2j1/Rpn1/Tmtc3/Tmtc1/Mlec/Galnt2/Mogs/Stt3b/Pomgnt2/Trip11/Poglut1/B4gat1/Eogt/Cds2/Fut11/Alg2/Pofut2/Entpd5/Uggt1/Pmm1/Ogt | 39 |
| BP | GO:2000736 | regulation of stem cell differentiation | 19/1488 | 41/8928 | 8,75E-06 | 0,000136 | 0,000121 | Gatad2b/Mta2/Nudt21/Chd3/Cdk13/Hdac2/Hnrnpu/Mta1/Mbd3/Ythdf2/Rbbp7/Mta3/Rbbp4/Ccnk/Chd4/Kat5/Setd1a/Dhx36/Ap2a2 | 19 |
| BP | GO:0033045 | regulation of sister chromatid segregation | 24/1488 | 59/8928 | 9,83E-06 | 0,000152 | 0,000136 | Smarcb1/Dpf1/Smarca4/Smarcc2/Smarcd3/Smarce1/Arid1b/Smarcd1/Dpf2/Hnrnpu/Bcl7a/Bub3/Actl6b/Arid1a/Tpr/Smarca2/Ik/Arid2/Mad1l1/Brd7/Mad2l1/Kat5/Pbrm1/Numa1 | 24 |
| BP | GO:0006337 | nucleosome disassembly | 9/1488 | 12/8928 | 1,32E-05 | 0,000194 | 0,000173 | Smarcb1/Smarca4/Smarcc2/Smarcd3/Smarce1/Smarcd1/Arid1a/Ssrp1/Arid2 | 9 |
| BP | GO:0045663 | positive regulation of myoblast differentiation | 15/1488 | 29/8928 | 1,49E-05 | 0,000217 | 0,000194 | Smarcb1/Smarca4/Smarcc2/Smarcd3/Smarce1/Arid1b/Smarcd1/Actl6b/Arid1a/Smarca2/Arid2/Brd7/Kat5/Pbrm1/Ripor2 | 15 |
| BP | GO:0010453 | regulation of cell fate commitment | 10/1488 | 15/8928 | 2,15E-05 | 0,000303 | 0,000271 | Gatad2b/Mta2/Chd3/Hdac2/Mta1/Mbd3/Rbbp7/Mta3/Rbbp4/Chd4 | 10 |
| BP | GO:0042659 | regulation of cell fate specification | 10/1488 | 15/8928 | 2,15E-05 | 0,000303 | 0,000271 | Gatad2b/Mta2/Chd3/Hdac2/Mta1/Mbd3/Rbbp7/Mta3/Rbbp4/Chd4 | 10 |
| BP | GO:0006334 | nucleosome assembly | 16/1488 | 33/8928 | 2,22E-05 | 0,000313 | 0,00028 | Smarca4/Npm1/Atrx/Hira/Set/Sart3/Macroh2a1/Rbbp4/Hp1bp3/Smarca2/Ssrp1/Smarca5/Macroh2a2/Rsf1/Ubn1/Tspyl4 | 16 |
| BP | GO:0001824 | blastocyst development | 27/1488 | 73/8928 | 2,24E-05 | 0,000315 | 0,000281 | Fbll1/Matr3/Smarcb1/Smarca4/Sf3b1/Brd4/Thoc2/Bysl/Thoc5/Ints1/Rpl7l1/Rpl13/Xab2/Rtf1/Hcfc1/Sf3b6/Tbl1xr1/Ctr9/Pbrm1/Nop2/Junb/Cdk11b/Cnot2/Rrp7a/Setdb1/Cnot1/Cnot3 | 27 |
| BP | GO:0034504 | protein localization to nucleus | 62/1488 | 227/8928 | 2,98E-05 | 0,000407 | 0,000363 | Nolc1/Ddx5/Elavl1/Trim28/Npm1/Srsf1/Pom121/Nup50/Nup35/Atf2/Plrg1/Hnrnpu/Hnrnpm/Sun1/Phip/Tardbp/Lmnb2/Parp1/Ints13/Sin3a/Nup153/Larp7/Nup188/Tpr/Ranbp2/Nup98/Rpl11/Tmem201/Nup107/Nup133/Rpf2/Nup88/Nup155/Rbm22/Dmap1/Supt7l/Paf1/Nup93/Ubr5/Nup85/Mepce/Nvl/Nup214/Adar/Sumo1/Nup54/Lmna/Hdgf/Nup62/Calr/Nup58/Lmnb1/Cry2/Dnajb6/Rrs1/Cdkl5/Hdac3/Cabp1/Rangap1/Sec13/Agap3/Ogt | 62 |
| BP | GO:0031330 | negative regulation of cellular catabolic process | 45/1488 | 152/8928 | 4,53E-05 | 0,000604 | 0,000539 | Elavl1/Hnrnpa0/Hnrnpab/Fus/Dhx9/Hnrnpc/Srsf1/Dkc1/Ccar2/Phf20l1/Hnrnpu/Hnrnpd/Golga2/Tardbp/Thrap3/Rps7/Rbm10/Nmnat1/Sorl1/Rybp/Fmr1/Rpl11/Foxk2/Syncrip/Larp4b/Rpl5/Cirbp/Rpl23/Mettl16/Taf15/Foxk1/Elavl4/Wac/Phax/Cnr1/Eif4g1/Dhx36/Eif3h/Eif4g2/Ctsa/Scfd1/Sdcbp/Csde1/Usp7/Ogt | 45 |
| BP | GO:0070988 | demethylation | 14/1488 | 28/8928 | 4,71E-05 | 0,000627 | 0,00056 | Trim28/Hnrnpab/Riox1/Kdm6a/Kdm1a/Syncrip/Apex1/Kdm5a/Alkbh5/Kdm2a/Cyp3a13/Otud4/Phf2/Usp7 | 14 |
| BP | GO:1905214 | regulation of RNA binding | 8/1488 | 11/8928 | 6E-05 | 0,000783 | 0,000699 | Ncbp1/Eif4a3/Fmr1/Cdk9/Nucks1/Eif3e/Eif4g1/Eif3d | 8 |
| BP | GO:0009247 | glycolipid biosynthetic process | 15/1488 | 32/8928 | 6,6E-05 | 0,000857 | 0,000765 | Ugcg/Slc30a5/Tm9sf2/St8sia3/B4galt6/Pigg/St6galnac5/Gal3st3/B4galnt1/Pgap4/Gpaa1/Pigt/Pigs/Pigu/Pigk | 15 |
| BP | GO:0045582 | positive regulation of T cell differentiation | 19/1488 | 48/8928 | 0,000127 | 0,001599 | 0,001427 | Smarcb1/Smarca4/Smarcc2/Smarcd3/Smarce1/Arid1b/Smarcd1/Sart1/Egr3/Actl6b/Arid1a/Smarca2/Arid2/Brd7/Kat5/Pbrm1/Zap70/Zmiz1/Rasgrp1 | 19 |
| BP | GO:0044770 | cell cycle phase transition | 73/1488 | 292/8928 | 0,000146 | 0,001811 | 0,001617 | Smarcb1/Dpf1/Smarca4/Smarcc2/Smarcd3/Ppp1r10/Npm1/Smarce1/Arid1b/Smarcd1/Dpf2/Ccar2/Brd4/Atf2/Plrg1/Cdc5l/Ddx39b/Bcl7a/Mecp2/Bub3/Prpf19/Actl6b/Thoc1/Arid1a/Sin3a/Macroh2a1/Larp7/Senp2/Tpr/Mta3/Smarca2/Thoc5/Ints7/Trrap/Ik/Cdc73/Rcc2/Rps6/Rpl17/Rpl24/Arid2/Ints3/Paf1/Pias1/Cirbp/Akap8l/Mad1l1/Mepce/Rpl26/Apex1/Brd7/Mad2l1/Rps27l/Wac/Nfib/Pbrm1/Upf1/Msh2/Cpsf3/Ercc3/Crebbp/Nabp2/Clock/Nfix/Gigyf2/Lmnb1/Akap8/Ankrd17/Ddx3x/Eif4g1/Ctc1/Rcc1/Ppp2r5c | 73 |
| BP | GO:0006260 | DNA replication | 37/1488 | 123/8928 | 0,000146 | 0,001811 | 0,001617 | Wdr18/Dhx9/Wrnip1/Npm1/Atrx/Parp1/Alyref/Thoc1/Sin3a/Senp2/Rbbp7/Rbbp4/Actr8/Ilkap/Top1/Ssrp1/Cdk9/Smarcal1/Smarca5/Meaf6/Gtpbp4/Pura/Rbbp6/Nfib/Upf1/Wiz/Nucks1/Pds5a/Nfix/Yy1/Lig3/Ankrd17/Ctc1/Smc3/Stn1/Ruvbl1/Ruvbl2 | 37 |
| BP | GO:1901798 | positive regulation of signal transduction by p53 class mediator | 10/1488 | 18/8928 | 0,000192 | 0,002321 | 0,002072 | Ddx5/Chd5/Rps7/Rps20/Rpl11/Rpl23/Rpl26/Hexim1/Rps15/Eef1e1 | 10 |
| BP | GO:0045445 | myoblast differentiation | 19/1488 | 50/8928 | 0,00024 | 0,002863 | 0,002556 | Smarcb1/Ddx5/Smarca4/Smarcc2/Smarcd3/Smarce1/Arid1b/Smarcd1/Actl6b/Arid1a/Ddx17/Smarca2/Arid2/Mbnl1/Brd7/Kat5/Pbrm1/Bcl9/Ripor2 | 19 |
| BP | GO:0030518 | intracellular steroid hormone receptor signaling pathway | 23/1488 | 66/8928 | 0,000252 | 0,003001 | 0,002679 | Ddx5/Smarca4/Parp1/Arid1a/Safb/Ddx17/Rbfox2/Ubr5/Ncor1/Kmt2d/Safb2/Ncor2/Calr/Clock/Trerf1/Cry2/Cnot2/Nr3c1/Zmiz1/Ufsp2/Cnot1/Ntrk2/Ufl1 | 23 |
| BP | GO:0045047 | protein targeting to ER | 14/1488 | 32/8928 | 0,000286 | 0,003355 | 0,002995 | Sec61a2/Hspa5/Ssr3/Sec61a1/Hyou1/Srp14/Srp72/Sec63/Srp68/Spcs1/Spcs2/Spcs3/Srprb/Get1 | 14 |

| | | | | | | | | | |
|---|---|---|---|---|---|---|---|---|---|
| BP | GO:0045070 | positive regulation of viral genome replication | 11/1488 | 22/8928 | 0,000312 | 0,003642 | 0,003252 | Top2b/Stau1/Adar/Adarb1/Srpk1/Ppih/Ddx3x/Ppie/Cnot7/Hacd3/Srpk2 | 11 |
| BP | GO:0140718 | facultative heterochromatin formation | 13/1488 | 29/8928 | 0,000352 | 0,004008 | 0,003579 | Trim28/Smarca4/Tasor/Mbd3/Kdm1a/Suz12/Samd1/Smarca5/Kdm5a/Mbd2/Morc2a/Phf2/Setdb1 | 13 |
| BP | GO:0006606 | protein import into nucleus | 36/1488 | 124/8928 | 0,000385 | 0,004345 | 0,003879 | Nolc1/Ddx5/Elavl1/Trim28/Pom121/Nup50/Nup35/Atf2/Phip/Tardbp/Nup153/Nup188/Tpr/Ranbp2/Nup98/Nup107/Nup133/Nup88/Nup155/Rbm22/Dmap1/Nup93/Ubr5/Nup85/Nup214/Adar/Sumo1/Nup54/Lmna/Nup62/Nup58/Cry2/Hdac3/Cabp1/Sec13/Agap3 | 36 |
| BP | GO:0007623 | circadian rhythm | 34/1488 | 116/8928 | 0,000456 | 0,005086 | 0,00454 | Ddx5/Dhx9/Pspc1/Sfpq/Ccar2/Hdac2/Hnrnpu/Hnrnpd/Mta1/Tardbp/Parp1/Nono/Sin3a/Thrap3/Creb1/Top1/Kmt2a/Ncor1/Mybbp1a/Kdm5a/Rai1/Kdm5b/Kat5/Npy2r/Mettl3/Kdm2a/Clock/Cry2/Hdac3/Rbm4b/Gabrb3/Ntrk2/Usp7/Ogt | 34 |
| BP | GO:0000075 | cell cycle checkpoint signaling | 27/1488 | 87/8928 | 0,000647 | 0,006933 | 0,00619 | Ppp1r10/Ccar2/Brd4/Atf2/Cdc5l/Ddx39b/Bub3/Prpf19/Thoc1/Tpr/Thoc5/Ints7/Trrap/Ik/Rps6/Rpl24/Ints3/Mad1l1/Rpl26/Mad2l1/Rps27l/Wac/Msh2/Nabp2/Clock/Gigyf2/Ppp2r5c | 27 |
| BP | GO:0035561 | regulation of chromatin binding | 8/1488 | 14/8928 | 0,000679 | 0,00721 | 0,006437 | Larp7/Senp2/Kdm1a/Mepce/Mbd2/Btaf1/Med25/Ctbp2 | 8 |
| BP | GO:0007346 | regulation of mitotic cell cycle | 66/1488 | 273/8928 | 0,000795 | 0,008353 | 0,007458 | Smarcb1/Dpf1/Smarca4/Smarcc2/Smarcd3/Ppp1r10/Smarce1/Arid1b/Smarcd1/Foxg1/Dpf2/Brd4/Atf2/Plrg1/Hnrnpu/Bcl7a/Mecp2/Bub3/Phip/Actl6b/Ints13/Arid1a/Sin3a/Larp7/Senp2/Tpr/Mta3/Smarca2/Trrap/Ik/Cdc73/Rcc2/Rps6/Rpl17/Trim35/Rpl24/Arid2/Ints3/Mad1l1/Mepce/Rpl26/Apex1/Brd7/Mad2l1/Rps27l/Wac/Pbrm1/Msh2/Cpsf3/Ercc3/Crebbp/Nup62/Nabp2/Cdk11b/Gigyf2/Lmnb1/Ankrd17/Ddx3x/Hdac3/Eif4g1/Usp22/Ctc1/Prkca/Rcc1/Ppp2r5c/Sdcbp | 66 |
| MF | GO:0003729 | mRNA binding | 161/1488 | 248/8928 | 5,85E-67 | 5,43E-64 | 4,85E-64 | Khdrbs3/Hnrnpl/Rbmx/Celf2/Srsf3/Rbfox1/Cpsf7/Rbfox3/Ddx5/Poldip3/Rbm14/Celf1/Elavl1/Hnrnpa0/Hnrnpa3/Hnrnpa2b1/Rbm42/Hnrnpr/Hnrnpll/Hnrnpa1/Fus/Dhx9/Khsrp/Ptbp2/Hnrnpc/Sf3b1/Snrnp70/Hnrnpk/Myef2/Ilf3/Srsf1/Nudt21/Rps26/Srsf6/Ncl/Fyttd1/Rbm25/Srsf5/Chtop/G3bp2/Srrm2/Elavl3/Hnrnpu/Hnrnpd/Tra2b/Khdrbs1/Rbm15b/Lsm14b/Hnrnpm/Mecp2/Rbm15/Ssb/Nova2/Tardbp/Thoc2/Rnps1/Rbm5/Ncbp1/Rbm8a/Alyref/Nxf1/Eif4a3/Celf5/Rps14/Pum2/Ddx17/Rsl1d1/Larp7/Rps13/Ythdf2/Luc7l3/Tpr/Luc7l/Nhp2/Rps7/Cpsf6/Rps3/Cstf3/Rpl6/Thoc5/Nup98/Pum1/Sf1/Rpl35/Rpl7/Srsf4/Rpl13a/Cdc40/Fmr1/Ppp1r8/Cpsf1/Ythdc1/Rps2/Rps6/Ptbp3/Syncrip/Larp4b/Tial1/Serbp1/Rpl24/Celf4/Rps5/Rpl5/Pcbp2/Pcbp3/Tia1/Scaf8/Zfp638/Khdrbs2/Fubp1/Cirbp/Spen/Rbfox2/Dazap1/Larp4/Dcp1a/Ythdf1/Aqr/Ncbp2/Fxr2/Rpl26/Taf15/Pum3/Mbd2/Ythdf3/Cpeb4/Elavl4/G3bp1/Rnf20/Sf3b6/Eif4g3/Gemin5/Mettl3/Fxr1/Rnf40/Purb/Calr/Pabpc4/Celf6/Pabpc1/Copa/Eif3a/Lsm14a/Ddx6/Ago2/Ddx3x/Secisbp2l/Exosc4/Eif4g1/Ppie/Dhx36/Rbm4b/Eif3d/Hdlbp/Edc3/Cstf2t/Eif4g2/Exosc8/Cpeb3/Cstf2/Fubp3 | 161 |
| MF | GO:0003682 | chromatin binding | 149/1488 | 327/8928 | 9,1E-36 | 2,41E-33 | 2,15E-33 | Rbmx/Tcf4/Pelp1/H1f10/Ddx5/Atxn1/Cbx5/Ncoa5/Satb2/Trim28/Smarca4/Fus/Smarcc2/Smarcd3/Dhx9/Sf3b1/Gtf2h1/Sin3b/Sfpq/Npm1/Mta2/Adnp/Arid1b/Smarcd1/Rnf2/Nudt21/Chd3/Sarnp/Hdac2/Lemd3/Psip1/Brd4/Tasor/Atf2/Trim33/Hnrnpu/Hnrnpd/Mta1/Mecp2/Atrx/Ep400/Zc3h4/Kdm6a/Mbd3/Cbx3/Parp1/Hira/Nono/Actl6b/Set/Arid1a/Sin3a/Nup153/Safb/Macroh2a1/Chd5/Ddx17/Tpr/Rbmxl1/Cbx1/Mta3/Hp1bp3/Smarca2/Cbx6/Kdm1a/Suz12/Nup98/Hdgfl3/Top1/Fmr1/Top2b/Chd4/Gtf2b/Ssrp1/Cdk9/Kmt2a/Ncoa1/Tle1/Gtf2f1/Ubtf/Samd1/Smarca5/Maz/Tox4/Paf1/Chd6/Ncor1/Hmgn3/Srebf2/Hmgn5/Polr2a/Apex1/Macroh2a2/Kdm5a/Hcfc1/Mbd2/Rnf20/Wac/Kat5/Polr2b/Glyr1/Cic/Polr3d/Polr3a/Pbrm1/Upf1/Ercc4/Msh2/Nucks1/Ercc3/Crebbp/Ncor2/Ing2/Chd2/Wdr82/Mef2a/Ddx1/Hmga1/Clock/Taf1/Morc2a/Nfix/Ccnt2/Yy1/Rxrb/Morf4l1/Sbno1/Akap8/Camta2/Brd3/Stag1/Mcm3ap/Msh6/Dhx30/Pwwp3a/Ctbp2/Nr3c1/Ankrd17/Hdac3/Smc1a/Smc3/Setdb1/Stag2/Nfat5/Rcc1/Ruvbl2/Tspyl4/Tdrd3/Ogt | 149 |
| MF | GO:0043565 | sequence-specific DNA binding | 152/1488 | 389/8928 | 2,29E-27 | 3,64E-25 | 3,25E-25 | Zeb2/Neurod2/Hnrnpl/Rbmx/Bcl11b/Tcf4/Mef2d/Gatad2b/Smarcb1/Nr2f1/Dpf1/Satb2/Hnrnpa2b1/Smarca4/Hnrnpa1/Hnrnpab/Dhx9/Hnrnpk/Erh/Myef2/Sfpq/Npm1/Xrn2/Mta2/Adnp/Bcl11a/Tcf20/Foxg1/Chd3/Neurod6/Hdac2/Brd4/Ncl/Chtop/Atf2/Sltm/Hnrnpu/Hnrnpd/Mta1/Cdc5l/Egr3/Mecp2/Tardbp/Kdm6a/Mbd3/Ccar1/Cbx3/Tef/Mbnl2/Purg/Safb/Macroh2a1/Nacc1/Rbbp5/Thrap3/Rbmxl1/Mta3/Phox2b/Rbbp4/Smarca2/Zfp512b/Rps3/Nr3c2/Kdm1a/Suz12/Drap1/Pou3f1/Creb1/Top1/Cpsf4/Esrra/Gtf2b/Foxk2/Gmeb1/Cdk9/Zbtb18/Tbp/Kmt2a/Ncoa1/Mypop/Ubtf/Terf2/Rfx3/Sub1/Zfp871/Maz/Ash2l/Fubp1/Zkscan16/Ncor1/Srebf2/Mybbp1a/Gmeb2/Polr2a/Apex1/Macroh2a2/Pura/Kdm5a/Hcfc1/Foxk1/Kdm5b/Mbd2/Brd7/Tbl1xr1/Cic/Thap11/Kmt2d/Gtf3c5/Camta1/Zbtb11/Nfib/Upf1/Wiz/Safb2/Terf2ip/Msh2/Hdgf/Purb/Crebbp/Kdm2a/Junb/Ncor2/Zscan26/Chd2/Nabp2/Mef2a/Hmga1/Clock/Taf1/Nfix/Yy1/Rxrb/Bptf/Lmnb1/Cry2/Camta2/Taf4/Nkrf/Zbtb20/Ago2/Preb/Arnt2/Nr3c1/Foxj3/Ago1/Hivep2/Dhx36/Ctc1/Prmt1/Stn1/Nfat5/Ruvbl2 | 152 |
| MF | GO:0140110 | transcription regulator activity | 165/1488 | 467/8928 | 5,96E-24 | 6,51E-22 | 5,81E-22 | Zeb2/Neurod2/Bcl11b/Tcf4/Mef2d/Smarcb1/Nr2f1/Ddx5/Rbm14/Satb2/Trim28/Smarca4/Fus/Smarcd3/Dhx9/Sin3b/Myef2/Npm1/Mta2/Adnp/Bclaf1/Bcl11a/Smarcd1/Tcf20/Foxg1/Mtdh/Neurod6/Ewsr1/Psip1/Brd4/Atf2/Ylpm1/Hnrnpu/Mta1/Cdc5l/Egr3/Mecp2/Prpf6/Ccar1/Snw1/Rcor3/Ahdc1/Tef/Tcerg1/Sin3a/Purg/Taf5l/Ddx17/Nacc1/Raly/Thrap3/Mta3/Phox2b/Smarca2/Zfp512b/Nr3c2/Kdm1a/Nup98/Drap1/Trrap/Hdgfl3/Gtf2i/Pou3f1/Creb1/Sap18/Rybp/Bud31/Esrra/Foxk2/Irf2bpl/Gmeb1/Kctd1/Hdgfl2/Zmynd8/Zbtb18/Med24/Ncoa1/Tle1/Mypop/Med17/Dmap1/Rfx3/Sub1/Maz/Supt7l/Pcbp3/Pias1/Sfr1/Spen/Rbfox2/Ncor1/Srebf2/Mybbp1a/Gmeb2/Apex1/Taf15/Pura/Kdm5a/Hcfc1/Naca/Foxk1/Kdm5b/Wdr43/Brd7/Rnf20/Kat5/Tbl1xr1/Cic/Thap11/Kmt2d/Camta1/Zbtb11/Med6/Nfib/Wiz/Hdgf/Purb/Tle3/Nucks1/Crebbp/Edf1/Kdm2a/Junb/Ncor2/Btaf1/Zscan26/Med14/Mef2a/Ddx1/Hmga1/Clock/Nfix/Zmynd11/Trerf1/Yy1/Rxrb/Supt3/Bcl9/Camta2/Taf6l/Nkrf/Zbtb20/Asah1/Preb/Pbxip1/Ctbp2/Arnt2/Nr3c1/Zmiz1/Hdac3/Foxj3/Hivep2/Usp22/Mlip/Phf2/Setd3/Med16/Cnot7/Tcf25/Ruvbl1/Nfat5/Ruvbl2/Scai/Ccdc124/Tdrd3 | 165 |
| MF | GO:0003712 | transcription coregulator activity | 109/1488 | 265/8928 | 7,26E-22 | 7,49E-20 | 6,68E-20 | Tcf4/Smarcb1/Ddx5/Rbm14/Trim28/Smarca4/Fus/Smarcd3/Dhx9/Sin3b/Npm1/Mta2/Bclaf1/Bcl11a/Smarcd1/Tcf20/Mtdh/Ewsr1/Psip1/Brd4/Hnrnpu/Mta1/Mecp2/Prpf6/Ccar1/Snw1/Rcor3/Tcerg1/Sin3a/Taf5l/Ddx17/Raly/Thrap3/Mta3/Smarca2/Kdm1a/Nup98/Drap1/Trrap/Hdgfl3/Sap18/Rybp/Bud31/Irf2bpl/Kctd1/Hdgfl2/Zmynd8/Med24/Ncoa1/Tle1/Med17/Dmap1/Sub1/Supt7l/Pias1/Sfr1/Spen/Rbfox2/Ncor1/Mybbp1a/Apex1/Taf15/Kdm5a/Hcfc1/Naca/Kdm5b/Brd7/Rnf20/Kat5/Tbl1xr1/Kmt2d/Camta1/Med6/Hdgf/Tle3/Nucks1/Crebbp/Edf1/Kdm2a/Ncor2/Btaf1/Med14/Ddx1/Hmga1/Zmynd11/Trerf1/Rxrb/Supt3/Bcl9/Camta2/Taf6l/Asah1/Pbxip1/Ctbp2/Nr3c1/Zmiz1/Hdac3/Usp22/Mlip/Phf2/Setd3/Med16/Cnot7/Tcf25/Ruvbl1/Ruvbl2/Scai/Ccdc124/Tdrd3 | 109 |
| MF | GO:0003735 | structural constituent of ribosome | 71/1488 | 142/8928 | 2,5E-20 | 2,29E-18 | 2,04E-18 | Rpl22l1/Rps26/Rpl38/Rps10/Rps25/Rps19/Rps14/Rpl14/Rps13/Rpl18/Rps15a/Rps4x/Rps11/Rpl15/Rps9/Rps7/Rps23/Rpl7a/Rps3/Rpl23a/Rpl18a/Rps20/Rpl6/Rpl39/Rpl37a/Rps18/Rpl27a/Rps28/Rpl31/Rpl35/Rpl7/Rpl4/Rps8/Rpl13a/Rpl7l1/Rpl22/Rps16/Rpl12/Rpl27/Rpl13/Rpl11/Fau/Rps2/Rps6/Rps27/Rpl17/Rps12/Rpl10/Rpl8/Rpl34/Rpl24/Rpl9/Rpl21/Rps5/Rpl35a/Rpl10a/Rpsa/Rpl5/Rpl3/Rpl19/Rplp0/Rpl23/Rplp2/Rpl26/Rps17/Rpl36/Rps27l/Rpl28/Rps15/Rps21/Mrpl57 | 71 |
| MF | GO:0001067 | transcription regulatory region nucleic acid binding | 117/1488 | 313/8928 | 2,27E-19 | 1,95E-17 | 1,74E-17 | Zeb2/Neurod2/Hnrnpl/Rbmx/Bcl11b/Tcf4/Mef2d/Smarcb1/Nr2f1/Satb2/Hnrnpa2b1/Smarca4/Dhx9/Hnrnpk/Myef2/Sfpq/Npm1/Xrn2/Adnp/Bcl11a/Tcf20/Chd3/Neurod6/Hdac2/Brd4/Atf2/Hnrnpu/Mta1/Cdc5l/Egr3/Tardbp/Kdm6a/Ccar1/Cbx3/Tef/Purg/Safb/Macroh2a1/Nacc1/Rbbp5/Thrap3/Rbmxl1/Phox2b/Rbbp4/Smarca2/Zfp512b/Rps3/Kdm1a/Suz12/Drap1/Pou3f1/Creb1/Top1/Esrra/Gtf2b/Foxk2/Gmeb1/Cdk9/Tbp/Ncoa1/Mypop/Ubtf/Rfx3/Sub1/Mbnl1/Zfp871/Maz/Ash2l/Fubp1/Zkscan16/Ncor1/Srebf2/Mybbp1a/Gmeb2/Polr2a/Macroh2a2/Pura/Kdm5a/Hcfc1/Foxk1/Brd7/Tbl1xr1/Cic/Thap11/Kmt2d/Gtf3c5/Zbtb11/Nfib/Wiz/Hdgf/Purb/Crebbp/Junb/Zscan26/Chd2/Mef2a/Hmga1/Clock/Taf1/Nfix/Yy1/Rxrb/Bptf/Cry2/Taf4/Nkrf/Zbtb20/Ago2/Preb/Arnt2/Nr3c1/Foxj3/Ago1/Hivep2/Dhx36/Nfat5/Ruvbl2 | 117 |

| | | | | | | | | | |
|---|---|---|---|---|---|---|---|---|---|
| MF | GO:0043021 | ribonucleoprotein complex binding | 68/1488 | 140/8928 | 1,2E-18 | 8,93E-17 | 7,97E-17 | Nolc1/Ddx5/Cbx5/Snrpd1/Dhx9/Hnrnpc/Snrnp70/Hnrnpk/Snrpd2/Npm1/Snrpa/Snrpb/Hnrnpu/Pqbp1/Rbm39/Prpf6/Snrpe/Snrpd3/Prpf31/Eif4a3/Snrpb2/Snrpg/Ddx17/Sec61a2/Cpsf6/Snd1/Stau2/Rpl35/Rack1/Fmr1/Nomo1/Ncln/Serbp1/Rpsa/Ythdf1/Rplp2/Bop1/Hspa5/Rpn2/Nvl/Gtpbp4/Rbm3/Ythdf3/Cpeb4/Gemin4/Gemin5/Cd2bp2/Sec61a1/Tmem147/Ccdc47/Tmco1/Pym1/Eif3k/Ppih/Srp72/Eif3c/Srp68/Spcs1/Ddx3x/Secisbp2l/Prmt1/Cpeb3/Nemf/Eif6/Derl1/Gcn1/Eif2s1/Csnk2a1 | 68 |
| MF | GO:0001098 | basal transcription machinery binding | 34/1488 | 47/8928 | 3,78E-17 | 2,51E-15 | 2,24E-15 | Tcf4/Nolc1/Dhx9/Nop58/Ccar2/Brd4/Hnrnpu/Scaf1/Fbl/Rprd1a/Drap1/Rprd2/Cdc73/Gtf2b/Tbp/Gtf2e2/Rtf1/Gtf2f1/Paf1/Scaf8/Rprd1b/Wdr43/Wac/Ctr9/Ercc4/Crebbp/Edf1/Leo1/Taf1/Pcif1/Ago2/Ago1/Ruvbl1/Ruvbl2 | 34 |
| MF | GO:0001099 | basal RNA polymerase II transcription machinery binding | 34/1488 | 47/8928 | 3,78E-17 | 2,51E-15 | 2,24E-15 | Tcf4/Nolc1/Dhx9/Nop58/Ccar2/Brd4/Hnrnpu/Scaf1/Fbl/Rprd1a/Drap1/Rprd2/Cdc73/Gtf2b/Tbp/Gtf2e2/Rtf1/Gtf2f1/Paf1/Scaf8/Rprd1b/Wdr43/Wac/Ctr9/Ercc4/Crebbp/Edf1/Leo1/Taf1/Pcif1/Ago2/Ago1/Ruvbl1/Ruvbl2 | 34 |
| MF | GO:0140640 | catalytic activity, acting on a nucleic acid | 117/1488 | 334/8928 | 7,66E-17 | 4,85E-15 | 4,33E-15 | Fbll1/Ddx5/Rnmt/Ddx50/Smarca4/Dus3l/Dhx9/Ddx23/Wrnip1/Xrn2/Dkc1/Chd3/Snrnp200/Ddx41/Dhx15/Ddx39b/Atrx/Ddx42/Ep400/Dbr1/Fbl/Arid1a/Eif4a3/Ddx59/Chd5/Ddx17/Rbbp4/Smarca2/Snd1/Rps3/Ddx46/Mtrex/Isy1/Dhx38/Top1/Top2b/Chd4/Trmt10b/Ddx51/Helz/Cmtr1/Terf2/Smarcal1/Smarca5/Chd6/Dcp1a/Dcp2/Dhx32/Aqr/Mettl16/Mepce/Farsa/Polr2a/Ddx20/Apex1/Trmt61a/G3bp1/Alkbh5/Polr2b/Ints11/Polr3d/Farsb/Rsf1/Toe1/Mars1/Exosc10/Mettl3/Polr3a/Upf1/Nop2/Ercc4/Msh2/Cpsf3/Ercc3/Polr2c/Polr2e/Btaf1/Rpp30/Chd2/Ddx39a/Xrcc1/Rngtt/Ddx1/Hmga1/Pld3/Dhx35/Trmt1l/Lig3/Dhx8/Bptf/Ftsj3/Cnot6l/Ddx47/Dis3/Pcif1/Msh6/Dhx30/Ddx6/Polr2i/Ago2/Cnot2/Dhx16/Rtcb/Ddx3x/Ddx21/Dhx36/Eif4a2/Polr3b/Polr1c/Smc3/Cnot7/Cnot1/Dhx57/Ruvbl1/Xrn1/Ruvbl2/Top3b | 117 |
| MF | GO:0003727 | single-stranded RNA binding | 41/1488 | 69/8928 | 1,03E-15 | 6,12E-14 | 5,47E-14 | Khdrbs3/Rbmx/Zfr/Hnrnpdl/Atxn1/U2af2/Hnrnpa1/Dhx9/Hnrnpc/Ilf3/Pabpn1/Strbp/Hnrnpu/Khdrbs1/Ssb/Zc3h14/Eif4a3/Rps7/Rbm10/Cbx6/Polr2g/Fmr1/Syncrip/Khdrbs2/Cirbp/Dazap1/Larp4/Aqr/Zfr2/Elavl4/Exosc10/Hnrnpf/Fxr1/Pabpc4/Ddx1/Pabpc1/Lsm14a/Ago2/Ddx3x/Ago1/Ppie | 41 |
| MF | GO:0008134 | transcription factor binding | 124/1488 | 377/8928 | 2,28E-15 | 1,29E-13 | 1,15E-13 | Tcf4/Mef2d/Nolc1/Smarcb1/Ddx5/Cbx5/Smarca4/Fus/Smarcd3/Dhx9/Gtf2h1/Npm1/Mta2/Smarce1/Bcl11a/Nop58/Mtdh/Hdac2/Psip1/Ncl/Atf2/Hnrnpu/Mta1/Cdc5l/Mecp2/Prpf6/Snw1/Cbx3/Parp1/Hira/Fbl/Arid1a/Tcerg1/Sin3a/Eef1d/Thrap3/Rps3/Nr3c2/Kdm1a/Suz12/Drap1/Gtf2i/Creb1/Sap18/Bud31/Chd4/Gtf2b/Kctd1/Cdk9/Tbp/Gtf2e2/Med24/Ncoa1/Dcaf1/Tle1/Arid2/Med17/Gtf2f1/Dmap1/Pias1/Rpl23/Spen/Rbfox2/Chd6/Dcp1a/Ncor1/Mybbp1a/Ddx20/Apex1/Pura/Hcfc1/Naca/Brd7/Kat5/Sumo1/Yeats2/Med6/Hnrnpf/Wiz/Ercc4/Ubn1/Hdgf/Purb/Nucks1/Crebbp/Edf1/Setd1a/Pdcd11/Junb/Ncor2/Paxbp1/Calr/Mef2a/Hmga1/Clock/Taf1/Ccnt2/Trerf1/Yy1/Rxrb/Bptf/Cry2/Akap8/Camta2/Taf4/Asah1/Med25/Cnot2/Pbxip1/Ctbp2/Arnt2/Ddx3x/Hdac3/Setd3/Keap1/Med16/Cnot7/Cnot1/Naaa/Trip12/Ruvbl1/Nfat5/Ruvbl2/Baiap2 | 124 |
| MF | GO:0042393 | histone binding | 65/1488 | 151/8928 | 1,24E-14 | 6,53E-13 | 5,83E-13 | Cbx5/Phf14/Smarca4/Smarcc2/Npm1/Dpf2/Chd3/Brd4/Ncl/Atrx/Phip/Cbx3/H2ax/Hira/Spin1/Set/Sart3/Chd5/Rbbp5/Rbbp7/Cbx1/Rbbp4/Smarca2/Cbx6/Suz12/Wdr5/Fmr1/Chd4/Ssrp1/Hdgfl2/Zmynd8/Kmt2a/Brd9/Brd1/Sgf29/Smarca5/Chd6/Kdm5a/Kdm5b/Brd7/Rnf20/Kat5/Glyr1/Tbl1xr1/Vps72/Rsf1/Kmt2d/Yeats2/Yeats4/Ing2/Chd2/Taf1/Morc2a/Zmynd11/Baz1b/Bptf/Sbno1/Brd3/Mcm3ap/Msh6/Dek/Phf2/Rcc1/Tspyl4/Tdrd3 | 65 |
| MF | GO:0004386 | helicase activity | 49/1488 | 99/8928 | 3,35E-14 | 1,71E-12 | 1,53E-12 | Ddx5/Ddx50/Smarca4/Dhx9/Ddx23/Wrnip1/Snrnp200/Ddx41/Dhx15/Ddx39b/Atrx/Ddx42/Ep400/Eif4a3/Ddx59/Chd5/Ddx17/Smarca2/Ddx46/Mtrex/Dhx38/Chd4/Ddx51/Helz/Smarcal1/Smarca5/Chd6/Dhx32/Aqr/Ddx20/G3bp1/Upf1/Ercc3/Chd2/Ddx39a/Ddx1/Dhx35/Dhx8/Ddx47/Dhx30/Ddx6/Dhx16/Ddx3x/Ddx21/Dhx36/Eif4a2/Dhx57/Ruvbl1/Ruvbl2 | 49 |
| MF | GO:0008186 | ATP-dependent activity, acting on RNA | 34/1488 | 57/8928 | 2,54E-13 | 1,17E-11 | 1,04E-11 | Ddx5/Ddx50/Dhx9/Ddx23/Snrnp200/Ddx41/Dhx15/Ddx39b/Ddx42/Eif4a3/Ddx59/Ddx17/Ddx46/Mtrex/Dhx38/Ddx51/Dhx32/Aqr/Ddx20/G3bp1/Upf1/Ddx39a/Ddx1/Dhx35/Dhx8/Ddx47/Dhx30/Ddx6/Dhx16/Ddx3x/Ddx21/Dhx36/Eif4a2/Dhx57 | 34 |
| MF | GO:0017069 | snRNA binding | 26/1488 | 38/8928 | 1,68E-12 | 6,77E-11 | 6,05E-11 | Snrnp70/Snrpa/Snu13/Snrpa1/Hnrnpu/Ddx39b/Prpf8/Eftud2/Snrpd3/Prpf31/Sart3/Snrpb2/Larp7/Sf3b3/Prpf4/Cdk9/Coil/Rbm22/Ncbp2/Mettl16/Mepce/Hexim1/Toe1/Gemin5/Ccnt2/Ddx21 | 26 |
| MF | GO:0140098 | catalytic activity, acting on RNA | 82/1488 | 238/8928 | 1,04E-11 | 3,78E-10 | 3,37E-10 | Fbll1/Ddx5/Rnmt/Ddx50/Dus3l/Dhx9/Ddx23/Xrn2/Snrnp200/Ddx41/Dhx15/Ddx39b/Ddx42/Dbr1/Fbl/Eif4a3/Ddx59/Ddx17/Snd1/Ddx46/Mtrex/Isy1/Dhx38/Trmt10b/Ddx51/Cmtr1/Dcp1a/Dcp2/Dhx32/Aqr/Mettl16/Mepce/Farsa/Polr2a/Ddx20/Apex1/Trmt61a/G3bp1/Alkbh5/Polr2b/Ints11/Polr3d/Farsb/Toe1/Mars1/Exosc10/Mettl3/Polr3a/Upf1/Nop2/Cpsf3/Polr2c/Polr2e/Rpp30/Ddx39a/Rngtt/Ddx1/Dhx35/Trmt1l/Dhx8/Ftsj3/Cnot6l/Ddx47/Dis3/Pcif1/Dhx30/Ddx6/Polr2i/Ago2/Cnot2/Dhx16/Rtcb/Ddx3x/Ddx21/Dhx36/Eif4a2/Polr3b/Polr1c/Cnot7/Cnot1/Dhx57/Xrn1 | 82 |
| MF | GO:0031490 | chromatin DNA binding | 27/1488 | 46/8928 | 1,27E-10 | 3,99E-09 | 3,56E-09 | H1f10/Dhx9/Hdac2/Lemd3/Hnrnpu/Mecp2/Atrx/Kdm6a/Macroh2a1/Ddx17/Suz12/Top1/Tox4/Hmgn3/Hmgn5/Apex1/Macroh2a2/Kdm5a/Hcfc1/Crebbp/Clock/Rxrb/Sbno1/Hdac3/Rcc1/Ruvbl2/Ogt | 27 |
| MF | GO:0000993 | RNA polymerase II complex binding | 20/1488 | 28/8928 | 1,95E-10 | 5,94E-09 | 5,31E-09 | Dhx9/Ccar2/Brd4/Hnrnpu/Scaf1/Rprd1a/Rprd2/Cdc73/Gtf2b/Rtf1/Paf1/Scaf8/Rprd1b/Wdr43/Wac/Ctr9/Leo1/Pcif1/Ago2/Ago1 | 20 |
| MF | GO:0036002 | pre-mRNA binding | 20/1488 | 30/8928 | 1,34E-09 | 3,74E-08 | 3,34E-08 | Hnrnpl/Celf2/Ddx5/Celf1/Hnrnpa2b1/U2af2/Hnrnpa1/U2af1/Hnrnpk/Srsf6/Srsf2/Hnrnpu/Tra2b/Prpf8/Tardbp/Sf1/Celf4/Rbm22/Elavl4/U2af1l4 | 20 |
| MF | GO:0019843 | rRNA binding | 29/1488 | 59/8928 | 7,28E-09 | 1,89E-07 | 1,68E-07 | Npm1/Ncl/Rps14/Rps13/Rps4x/Rps11/Rps9/Rps3/Rpl23a/Rpl6/Rps18/Rpl7/Rpl4/Rpl12/Rpl11/Rpl17/Rpl8/Rpl9/Rpf2/Rps5/Rpl5/Rpl3/Rpl19/Cirbp/Rplp0/Rpl23/Brix1/Rrs1/Ddx21 | 29 |
| MF | GO:0001091 | RNA polymerase II general transcription initiation factor binding | 15/1488 | 20/8928 | 1,34E-08 | 3,35E-07 | 2,99E-07 | Tcf4/Nolc1/Nop58/Hnrnpu/Fbl/Drap1/Tbp/Gtf2e2/Gtf2f1/Ercc4/Crebbp/Edf1/Taf1/Ruvbl1/Ruvbl2 | 15 |
| MF | GO:0061980 | regulatory RNA binding | 17/1488 | 25/8928 | 1,53E-08 | 3,8E-07 | 3,4E-07 | Matr3/Elavl1/Hnrnpa2b1/Hnrnpa1/Dhx9/Mecp2/Pum2/Rbm10/Pum1/Fmr1/Mbd2/Prkra/Fam172a/Ago2/Ago1/Ddx21/Zc3h7b | 17 |
| MF | GO:0008094 | ATP-dependent activity, acting on DNA | 31/1488 | 70/8928 | 4,99E-08 | 1,14E-06 | 1,02E-06 | Smarca4/Dhx9/Wrnip1/Chd3/Atrx/Ep400/Arid1a/Chd5/Rbbp4/Smarca2/Top2b/Chd4/Smarcal1/Smarca5/Chd6/G3bp1/Rsf1/Upf1/Msh2/Ercc3/Btaf1/Chd2/Ddx1/Bptf/Msh6/Dhx30/Ddx3x/Dhx36/Smc3/Ruvbl1/Ruvbl2 | 31 |
| MF | GO:0140034 | methylation-dependent protein binding | 24/1488 | 48/8928 | 9,55E-08 | 2,11E-06 | 1,88E-06 | Cbx5/Dpf2/Phf20l1/Atrx/Cbx3/Spin1/Chd5/Cbx1/Cbx6/Suz12/Wdr5/Fmr1/Hdgfl2/Zmynd8/Sgf29/Kdm5a/Glyr1/Ing2/Taf1/Zmynd11/Bptf/Msh6/Phf2/Tdrd3 | 24 |
| MF | GO:0140097 | catalytic activity, acting on DNA | 42/1488 | 115/8928 | 1,95E-07 | 4,06E-06 | 3,62E-06 | Smarca4/Dhx9/Wrnip1/Dkc1/Chd3/Atrx/Ep400/Arid1a/Chd5/Rbbp4/Smarca2/Rps3/Top1/Top2b/Chd4/Terf2/Smarcal1/Smarca5/Chd6/Apex1/G3bp1/Rsf1/Upf1/Ercc4/Msh2/Ercc3/Btaf1/Chd2/Xrcc1/Ddx1/Hmga1/Pld3/Lig3/Bptf/Msh6/Dhx30/Ddx3x/Dhx36/Smc3/Ruvbl1/Ruvbl2/Top3b | 42 |
| MF | GO:0106222 | lncRNA binding | 10/1488 | 12/8928 | 7,68E-07 | 1,42E-05 | 1,26E-05 | Celf2/Celf1/Elavl1/Smarca4/Hnrnpu/Pum2/Ddx17/Kdm1a/Suz12/Atp2a2 | 10 |
| MF | GO:0031491 | nucleosome binding | 14/1488 | 22/8928 | 1,02E-06 | 1,84E-05 | 1,64E-05 | H1f10/Arid1b/Parp1/Hira/Arid1a/Macroh2a1/Hp1bp3/Ssrp1/Smarca5/Hmgn3/Hmgn5/Glyr1/Pwwp3a/Rcc1 | 14 |
| MF | GO:0016758 | hexosyltransferase activity | 30/1488 | 76/8928 | 1,67E-06 | 2,91E-05 | 2,6E-05 | Galnt17/Ugcg/Galnt16/B3gat3/Galnt9/B3galt6/Fut8/B4galt6/Alg10b/Large1/Stt3a/Extl2/Dpy19l3/B4galnt1/Tmtc3/Tmtc1/Galnt2/Stt3b/Pomgnt2/Colgalt1/Poglut1/Hexa/B4gat1/Eogt/Fut11/Alg2/Pofut2/Chpf/Uggt1/Ogt | 30 |

| | | | | | | | | | |
|---|---|---|---|---|---|---|---|---|---|
| MF | GO:0004402 | histone acetyltransferase activity | 14/1488 | 24/8928 | 4,48E-06 | 7,31E-05 | 6,53E-05 | Atf2/Gtf2b/Med24/Ncoa1/Brd1/Meaf6/Kat5/Gtf3c4/Crebbp/Clock/Taf1/Mcm3ap/Usp22/Naa50 | 14 |
| MF | GO:0003725 | double-stranded RNA binding | 23/1488 | 54/8928 | 5,98E-06 | 9,49E-05 | 8,47E-05 | Zfr/Ilf2/Elavl1/Dhx9/Ilf3/Mtdh/Strbp/Hnrnpu/Dhx15/Stau2/Mbnl1/Zfr2/Stau1/Adar/Adarb1/Ddx1/Prkra/Lsm14a/Dhx30/Ago2/Ago1/Ddx21/Dhx36 | 23 |
| MF | GO:0140223 | general transcription initiation factor activity | 16/1488 | 31/8928 | 7,99E-06 | 0,000126 | 0,000112 | Gtf2h3/Drap1/Gtf2b/Tbp/Gtf2e2/Gtf2f1/Ubtf/Gtf3c2/Gtf3c5/Gtf3c4/Gtf2f2/Taf1/Gtf3c3/Taf6l/Taf4/Gtf3c1 | 16 |
| MF | GO:0030515 | snoRNA binding | 14/1488 | 25/8928 | 8,62E-06 | 0,000134 | 0,00012 | Nolc1/Nop56/Nop58/Dkc1/Snu13/Gar1/Nhp2/Bysl/Wdr3/Heatr1/Utp6/Rrp9/Ddx21/Nudt16l1 | 14 |
| MF | GO:0045182 | translation regulator activity | 37/1488 | 112/8928 | 1,53E-05 | 0,000223 | 0,000199 | Eif4a3/Eef1d/Rps14/Rps9/Eef1g/Fmr1/Rpl10/Cirbp/Fxr2/Pura/Cpeb4/Elavl4/Rps27l/Eif4g3/Fxr1/Purb/Eif3l/Eif3k/Eif3a/Eif3c/Eif3f/Ago2/Eif3i/Eif3b/Eif3e/Eif4g1/Eif3d/Eif3h/Eif4a2/Eif4g2/Cpeb3/Eif2s3x/Eif6/Eif5b/Gcn1/Eif2s1/Cyfip1 | 37 |
| MF | GO:0140658 | ATP-dependent chromatin remodeler activity | 13/1488 | 23/8928 | 1,59E-05 | 0,00023 | 0,000206 | Smarca4/Chd3/Atrx/Ep400/Arid1a/Chd5/Smarca2/Chd4/Smarcal1/Smarca5/Chd6/Rsf1/Chd2 | 13 |
| MF | GO:0042826 | histone deacetylase binding | 25/1488 | 65/8928 | 2,06E-05 | 0,000295 | 0,000263 | Mef2d/Cbx5/Satb2/Sfpq/Mta2/Nudt21/Hdac2/Hnrnpd/Mta1/Mecp2/Parp1/Chd5/Mta3/Rbbp4/Top2b/Chd4/Akap8l/Ncor1/Ddx20/Ncor2/Mef2a/Akap8/Camta2/Hdac3/Dhx36 | 25 |
| MF | GO:0070577 | lysine-acetylated histone binding | 11/1488 | 18/8928 | 2,7E-05 | 0,000375 | 0,000335 | Smarca4/Dpf2/Brd4/Phip/Zmynd8/Kmt2a/Brd9/Brd7/Yeats4/Taf1/Brd3 | 11 |
| MF | GO:0016922 | nuclear receptor binding | 32/1488 | 98/8928 | 7,29E-05 | 0,000945 | 0,000844 | Ddx5/Smarca4/Fus/Smarcd3/Gtf2h1/Smarce1/Prpf6/Snw1/Parp1/Arid1a/Thrap3/Kdm1a/Bud31/Gtf2b/Med24/Ncoa1/Dcaf1/Med17/Ncor1/Ncor2/Calr/Hmga1/Taf1/Trerf1/Rxrb/Cry2/Asah1/Med25/Ctbp2/Med16/Cnot1/Trip12 | 32 |
| MF | GO:0035613 | RNA stem-loop binding | 8/1488 | 13/8928 | 0,000341 | 0,00394 | 0,003518 | Dhx9/Eif4a3/Fmr1/Dazap1/Mettl16/Ddx3x/Cpeb3/Csde1 | 8 |
| MF | GO:0003743 | translation initiation factor activity | 18/1488 | 48/8928 | 0,00042 | 0,004725 | 0,004219 | Eif4g3/Eif3l/Eif3k/Eif3a/Eif3c/Eif3f/Eif3i/Eif3b/Eif3e/Eif4g1/Eif3d/Eif3h/Eif4a2/Eif4g2/Eif2s3x/Eif6/Eif5b/Eif2s1 | 18 |
| MF | GO:0070034 | telomerase RNA binding | 8/1488 | 14/8928 | 0,000679 | 0,00721 | 0,006437 | Hnrnpc/Dkc1/Snrpb/Hnrnpu/Gar1/Snrpd3/Nhp2/Dhx36 | 8 |
| CC | GO:0005681 | spliceosomal complex | 124/1488 | 165/8928 | 4,43E-63 | 3,52E-60 | 3,15E-60 | Rbmx/Ddx5/Hnrnpa3/Hnrnpa2b1/Tra2a/Hnrnpr/U2af2/Snrpd1/Hnrnpa1/Ddx23/Ptbp2/U2af1/Prpf40a/Hnrnpc/Sf3b1/Snrnp70/Hnrnpk/Snrpd2/Srsf1/Snrpa/Snu13/Magoh/Snrnp40/Snrpa1/Zmat2/Sart1/Api5/Ncl/Pnn/Srsf2/Usp39/Snrpb/Snrnp200/Ddx41/Plrg1/Srrm2/Hnrnpu/Tra2b/Cdc5l/Dhx15/Ddx39b/Sf3a1/Prpf8/Hnrnpm/Prpf40b/Prpf6/Prpf19/Htatsf1/Rbm5/Snw1/Smu1/Rbm8a/Sf3a3/Sf3b2/Snrpe/Eftud2/Prpf3/Alyref/Snrpd3/Prpf31/Eif4a3/Sugp1/Snrpb2/Snrpg/Zcchc8/Raly/Ctnnbl1/Sf3b3/Luc7l3/Prpf4/Luc7l/Rbmxl1/Sf3b4/Rbm17/Prpf4b/Mtrex/Isy1/Ik/Sf1/Phf5a/Srrm1/Cdc40/Sf3a2/Hnrnph1/Dhx38/Ppp1r8/Bcas2/Bud31/Snrpf/Syncrip/Gpkow/Xab2/Rbm22/Smndc1/Ppil2/Cirbp/Tfip11/Frg1/Dhx32/Aqr/Rbm28/Zcrb1/Rbm3/Adar/Sf3b6/Wac/Hnrnpf/Cwc25/Upf1/Srek1/Crnkl1/Lsm7/Sf3b5/Lsm6/Cwc22/U2af1l4/Dhx35/Dhx8/Pabpc1/Cwf19l1/Lsm8/Dhx16/Lsm2/Ppie | 124 |
| CC | GO:0000785 | chromatin | 183/1488 | 388/8928 | 1E-46 | 5,08E-44 | 4,54E-44 | Zeb2/Hnrnpl/Rbmx/Bcl11b/Tcf4/Pelp1/Gatad2b/Smarcb1/Dpf1/Cbx5/Satb2/Hnrnpa2b1/Trim28/Phf14/Smarca4/Smarcc2/Smarcd3/Dhx9/Ddx23/Sf3b1/Sin3b/Hnrnpk/Sfpq/Ppp1r10/Mta2/Smarce1/Arid1b/Bcl11a/Smarcd1/Rnf2/Dpf2/Chd3/Ccar2/Phf20l1/Hdac2/Psip1/Pogz/Brd4/Sap130/Srsf2/Tasor/Atf2/Mta1/Cfdp1/Bcl7a/Mecp2/Atrx/Ep400/Tardbp/Mbd3/Snw1/Cbx3/H2ax/Parp1/Hira/Actl6b/Set/Ahctf1/Arid1a/Sin3a/Taf5l/Macroh2a1/Chd5/Sf3b3/Rbbp7/Cbx1/Mta3/Phox2b/Cpsf6/Rbbp4/Hp1bp3/Smarca2/Cbx6/Kdm1a/Suz12/Actr8/Nmnat1/Trrap/Wdr5/Creb1/Top2b/Bud31/Chd4/Ssrp1/Zmynd8/Zbtb18/Tbp/Ncoa1/Brd9/H4c1/Arid2/Brd1/Sgf29/Dmap1/Rfx3/Smarca5/Supt7l/Ash2l/Tox4/Prdm10/Sfr1/Meaf6/Chd6/Akap8l/Ncor1/Aff4/Hmgn3/Hmgn5/Mybbp1a/Pds5b/Polr2a/Macroh2a2/Hcfc1/Msl2/Wdr43/Mbd2/Brd7/Rnf20/Epc1/Kat5/Glyr1/Vps72/Rsf1/Sumo1/Yeats2/Ctr9/Epc2/Exosc10/Pbrm1/Upf1/Nsd3/Rnf40/Yeats4/Nucks1/Crebbp/Sf3b5/Kdm2a/Setd1a/Ppp4r3b/Ncor2/Pds5a/Ing2/Chd2/Wdr82/Mef2a/Xrcc1/Hmga1/Clock/Taf1/Morc2a/Baz1b/Srpk1/Yy1/Bptf/Supt3/Morf4l1/Akap8/Camta2/Taf6l/Taf4/Brd3/Stag1/Msh6/Ddx6/Pbxip1/Ankrd17/Hdac3/Exosc4/Dek/Ddx21/Usp22/Setd3/Tmpo/Smc3/Stag2/Ruvbl1/Skic8/Rcc1/Ruvbl2/Tspyl4/Srpk2/Csnk2a1/Ogt | 183 |
| CC | GO:0016604 | nuclear body | 198/1488 | 444/8928 | 6,15E-46 | 2,64E-43 | 2,35E-43 | Fbll1/Srsf3/Nolc1/Ddx5/Poldip3/Rbm14/Cbx5/Hnrnpa2b1/Sugp2/Srsf7/U2af2/Dhx9/U2af1/Prpf40a/Sf3b1/Snrnp70/Pspc1/Sfpq/Ppp1r10/Npm1/Bclaf1/Srsf1/Bcl11a/Rnf2/Nop58/Tcf20/Mtdh/Dkc1/Nudt21/Chd3/Sarnp/Cdk13/Ewsr1/Snrnp40/Srsf6/Snrpa1/Sart1/Api5/Fyttd1/Srsf10/Rbm25/Sap130/Pnn/Srsf2/Srsf5/Chtop/Sltm/Plrg1/Ylpm1/Srrm2/Hnrnpu/Pqbp1/Cdc5l/Dhx15/Ddx39b/Rbm15b/Srsf11/Wbp11/Sf3a1/Rbm39/Hnrnpm/Atrx/Ddx42/Rbm15/Ep400/Pnisr/Srrt/Tardbp/Prpf6/Son/Prpf19/Zc3h14/Snw1/H2ax/Rbm8a/Sf3a3/Sf3b2/Parp1/Zc3h18/Eftud2/Prpf3/Nono/Nxf1/Ints13/Fbl/Thoc1/Snrpd3/Prpf31/Champ1/Sart3/Snrpb2/Taf5l/Chd5/Ddx17/Zcchc8/Senp2/Thoc6/Luc7l3/Prcc/Prpf4/Ppig/Thrap3/Nufip2/Rbm10/Acin1/Wtap/Cpsf6/Hp1bp3/Ddx46/Suz12/Nup98/Prpf4b/Nup43/Nmnat1/Ints7/Sympk/Ik/Sf1/Phf5a/Srsf4/Srrm1/Rnf112/Cdc40/Sf3a2/Rbm19/Sap18/Fmr1/Ppp1r8/Bcas2/Ythdc1/Gtf2b/Habp4/Cdk9/Zbtb18/Gtf2e2/Brd1/Coil/Terf2/Pias1/Atxn2l/Tfip11/Akap8l/Pcnp/Rpn2/Bpnt2/Apex1/Ppp4r3a/Adar/Rbbp6/Alkbh5/Vps72/Sumo1/Gemin4/Lmna/Toe1/Gemin5/Cd2bp2/Mettl3/Zmym2/Cwc25/Safb2/Terf2ip/Srek1/Ubn1/Gemin7/Crebbp/Setd1a/Ppp4r3b/Ncor2/Cwc22/Ddx39a/Ddx1/Srpk1/Zc3h13/Dhx8/Morf4l1/Cry2/Ppih/Drg1/Stag1/Trip11/Zbtb20/Fnbp4/Nr3c1/Virma/Eif3e/Ppie/Dhx36/Mlip/Rbm4b/Cnot7/Scaper/Cstf2/Trip12/Rgs14/Agap3/Srpk2/Csnk1a1 | 198 |
| CC | GO:0022626 | cytosolic ribosome | 71/1488 | 86/8928 | 2,26E-41 | 8,4E-39 | 7,5E-39 | Rps26/Rpl38/Rps10/Rps25/Rps19/Rps14/Rpl14/Rps13/Rpl18/Rps15a/Rps4x/Rps11/Rpl15/Rps9/Rps7/Rps23/Rpl7a/Rps3/Rpl23a/Rpl18a/Rps20/Rpl6/Rpl39/Rpl37a/Rps18/Rpl27a/Rps28/Rpl31/Rpl35/Rpl7/Rpl4/Rps8/Rpl13a/Rpl7l1/Rpl22/Rps16/Rpl12/Rpl27/Rpl13/Rpl11/Fau/Rps2/Rps6/Rps27/Rpl17/Rps12/Rpl10/Rpl8/Rpl34/Rpl24/Rpl9/Rpl21/Rps5/Rpl35a/Rpl10a/Rpsa/Rpl5/Rpl3/Rpl19/Rplp0/Rpl23/Larp4/Rplp2/Rpl26/Rps17/Rpl36/Rps27l/Rpl28/Rps15/Rps21/Ddx3x | 71 |
| CC | GO:0120114 | Sm-like protein family complex | 59/1488 | 79/8928 | 3,9E-30 | 7,24E-28 | 6,46E-28 | Nolc1/Rbm42/Snrpd1/Ddx23/Prpf40a/Sf3b1/Snrnp70/Snrpd2/Snrpa/Snu13/Snrnp40/Snrpa1/Zmat2/Sart1/Usp39/Snrpb/Snrnp200/Ddx39b/Sf3a1/Prpf8/Prpf40b/Prpf6/Htatsf1/Sf3a3/Sf3b2/Snrpe/Eftud2/Prpf3/Snrpd3/Prpf31/Sart3/Snrpb2/Snrpg/Larp7/Sf3b3/Luc7l3/Prpf4/Luc7l/Sf3b4/Phf5a/Sf3a2/Fmr1/Snrpf/Snrnp27/Gemin8/Mepce/Ddx20/Hexim1/Sf3b6/Gemin4/Gemin5/Cd2bp2/Lsm7/Gemin7/Sf3b5/Lsm6/Ppih/Lsm8/Lsm2 | 59 |
| CC | GO:0070603 | SWI/SNF superfamily-type complex | 50/1488 | 68/8928 | 3,49E-25 | 4,06E-23 | 3,62E-23 | Bcl11b/Gatad2b/Smarcb1/Dpf1/Smarca4/Smarcc2/Smarcd3/Sf3b1/Mta2/Smarce1/Arid1b/Bcl11a/Smarcd1/Dpf2/Chd3/Hdac2/Mta1/Cfdp1/Bcl7a/Ep400/Mbd3/Actl6b/Arid1a/Chd5/Rbbp7/Mta3/Rbbp4/Smarca2/Suz12/Actr8/Trrap/Chd4/Brd9/Arid2/Dmap1/Smarca5/Mybbp1a/Mbd2/Brd7/Kat5/Rsf1/Pbrm1/Baz1b/Yy1/Bptf/Dek/Ddx21/Ruvbl1/Ruvbl2/Csnk2a1 | 50 |
| CC | GO:1904949 | ATPase complex | 55/1488 | 96/8928 | 1,33E-19 | 1,16E-17 | 1,04E-17 | Bcl11b/Gatad2b/Smarcb1/Dpf1/Smarca4/Smarcc2/Smarcd3/Sf3b1/Mta2/Smarce1/Arid1b/Bcl11a/Smarcd1/Dpf2/Chd3/Hdac2/Mta1/Cfdp1/Bcl7a/Ep400/Mbd3/Actl6b/Arid1a/Chd5/Rbbp7/Mta3/Rbbp4/Smarca2/Suz12/Actr8/Trrap/Chd4/Brd9/Arid2/Dmap1/Smarca5/Mybbp1a/Mbd2/Brd7/Kat5/Rsf1/Pbrm1/Atp6ap2/Baz1b/Yy1/Bptf/Atp6ap1/Ccdc115/Dek/Ddx21/Atp6v0a2/Ruvbl1/Atp6v1g2/Ruvbl2/Csnk2a1 | 55 |

| | | | | | | | | | |
|---|---|---|---|---|---|---|---|---|---|
| CC | GO:0005844 | polysome | 44/1488 | 71/8928 | 9,27E-18 | 6,7E-16 | 5,99E-16 | Fus/Dhx9/Rps26/Rpl38/Rps25/Larp7/Rpl18/Rps4x/Nufip2/Rps23/Rpl7a/Rps3/Rpl18a/Rpl6/Rpl39/Rps28/Rpl31/Rpl7/Fmr1/Rpl11/Rps6/Rpl17/Larp4b/Rpl8/Rpl24/Rpl10a/Rpl19/Larp4/Fxr2/Rpl36/Elavl4/Rps21/Fxr1/Calr/Atxn2/Drg1/Ago2/Ago1/Eif4g1/Hdlbp/Eif3h/Btf3/Gcn1/Fubp3 | 44 |
| CC | GO:0005667 | transcription regulator complex | 93/1488 | 234/8928 | 1,29E-17 | 8,96E-16 | 8E-16 | Tcf4/Gatad2b/Rbm14/Cbx5/Satb2/Trim28/Hnrnpab/Gtf2h1/Sfpq/Mta2/Adnp/Chd3/Hdac2/Atf2/Hnrnpu/Mta1/Gtf2h3/Mbd3/Cbx3/Parp1/Rcor3/Nono/Sin3a/Taf5l/Chd5/Rbbp7/Mta3/Rbbp4/Drap1/Trrap/Pou3f1/Creb1/Sap18/Chd4/Gtf2b/Tbp/Gtf2e2/Med24/Ncoa1/Tle1/Arid2/Med17/Gtf2f1/Rfx3/Sub1/Prdm10/Spen/Dcp1a/Ncor1/Gtf3c2/Ddx20/Apex1/Mbd2/Kat5/Tbl1xr1/Thap11/Gtf3c5/Gtf3c4/Gemin5/Gtf2f2/Med6/Hdgf/Tle3/Ercc3/Crebbp/Junb/Ncor2/Med23/Ing2/Med14/Mef2a/Hmga1/Clock/Taf1/Trerf1/Yy1/Rxrb/Supt3/Bcl9/Gtf3c3/Taf6l/Taf4/Med25/Pbxip1/Ctbp2/Arnt2/Hdac3/Hivep2/Usp22/Med16/Gtf3c1/Nfat5/Csnk2a1 | 93 |
| CC | GO:0140534 | endoplasmic reticulum protein-containing complex | 47/1488 | 87/8928 | 1,42E-15 | 8,18E-14 | 7,3E-14 | Fam8a1/Sec61a2/Sel1l/Pdia6/Syvn1/Tusc3/Hspa5/Srebf2/Rpn2/Stt3a/Dnajb11/Ssr4/Ddost/Rpn1/Sec61a1/Mlec/Hsp90b1/Hyou1/Calr/Stt3b/Pdia4/Dnajc10/Pdia3/Gpaa1/Sec63/Sec11a/Spcs1/Spcs2/Spcs3/Pigt/Pigs/Sec11c/Ppib/Pigu/Pigk/Ganab/Sdf2l1/Srprb/Emc7/Prkcsh/Emc3/Emc10/Emc1/Emc2/Derl1/Emc4/Get1 | 47 |
| CC | GO:0034708 | methyltransferase complex | 43/1488 | 76/8928 | 2,47E-15 | 1,39E-13 | 1,24E-13 | Pelp1/Tex10/Rnmt/Cbx5/Snrpd1/Erh/Snrpd2/Senp3/Rnf2/Las1l/Hdac2/Snrpb/Rbm15b/Ramac/Rbm15/Kdm6a/Snrpe/Snrpd3/Prpf31/Snrpg/Rbbp5/Rbbp7/Wtap/Rbbp4/Suz12/Wdr5/Snrpf/Kmt2a/Ash2l/Hcfc1/Trmt61a/Kmt2d/Mettl3/Trmt6/Setd1a/Wdr82/Taf1/Zc3h13/Taf4/Virma/Prmt1/Ruvbl1/Ruvbl2 | 43 |
| CC | GO:0044815 | DNA packaging complex | 28/1488 | 42/8928 | 6,23E-13 | 2,71E-11 | 2,42E-11 | Ep400/H2ax/Macroh2a1/Hp1bp3/Trrap/H4c1/Dmap1/Terf2/Meaf6/Macroh2a2/Epc1/Kat5/Glyr1/Vps72/Epc2/Terf2ip/Yeats4/Morf4l1/Stag1/Ctc1/Smc1a/Smc3/Stag2/Stn1/Ruvbl1/Ruvbl2/Top3b/Tdrd3 | 28 |
| CC | GO:0032993 | protein-DNA complex | 40/1488 | 78/8928 | 1,78E-12 | 7,13E-11 | 6,37E-11 | Wdr18/Hnrnpk/Npm1/Plrg1/Cdc5l/Ep400/Prpf19/Gtf2h3/H2ax/Parp1/Macroh2a1/Hp1bp3/Suz12/Trrap/Top1/Bcas2/Chd4/Gtf2b/Tbp/H4c1/Dmap1/Terf2/Smarcal1/Meaf6/Macroh2a2/Kdm5a/Epc1/Kat5/Glyr1/Vps72/Gtf2f2/Epc2/Terf2ip/Yeats4/Ercc3/Morf4l1/Ctc1/Stn1/Ruvbl1/Ruvbl2 | 40 |
| CC | GO:0030684 | preribosome | 33/1488 | 58/8928 | 3,61E-12 | 1,38E-10 | 1,23E-10 | Fbll1/Nop56/Nop58/Las1l/Snu13/Riox1/Mrto4/Krr1/Fbl/Rsl1d1/Rps7/Bysl/Pes1/Utp14a/Bop1/Wdr3/Ebna1bp2/Nip7/Heatr1/Utp6/Mak16/Pdcd11/Mdn1/Utp18/Noc4l/Ftsj3/Wdr36/Rrs1/Nol6/Rrp7a/Rrp9/Wdr12/Eif6 | 33 |
| CC | GO:0005643 | nuclear pore | 35/1488 | 65/8928 | 7E-12 | 2,57E-10 | 2,29E-10 | Nup210/Pom121/Nup50/Nup35/Nxf1/Ahctf1/Nup153/Nup188/Senp2/Tpr/Ranbp2/Nup98/Nup43/Nup160/Nup107/Nup133/Nup205/Nup88/Nup155/Nup37/Nup93/Nup85/Mad1l1/Nup214/Mad2l1/Nup54/Ranbp3/Nup62/Aaas/Nup58/Mcm3ap/Seh1l/Rae1/Rangap1/Sec13 | 35 |
| CC | GO:0000428 | DNA-directed RNA polymerase complex | 34/1488 | 69/8928 | 3,38E-10 | 1,02E-08 | 9,08E-09 | Gtf2h1/Gtf2h3/Rprd1a/Taf5l/Rprd2/Trrap/Polr2g/Cdc73/Gtf2b/Tbp/Gtf2e2/Rtf1/Gtf2f1/Paf1/Rprd1b/Polr2a/Polr2b/Polr3d/Gtf2f2/Ctr9/Polr3a/Ercc3/Polr2c/Polr2e/Leo1/Taf1/Supt3/Taf6l/Taf4/Polr2i/Usp22/Polr3b/Polr1c/Skic8 | 34 |
| CC | GO:0031248 | protein acetyltransferase complex | 33/1488 | 67/8928 | 6,24E-10 | 1,84E-08 | 1,64E-08 | Phf20l1/Atf2/Ep400/Actl6b/Taf5l/Sf3b3/Trrap/Wdr5/Brd1/Sgf29/Dmap1/Supt7l/Meaf6/Hcfc1/Msl2/Epc1/Kat5/Vps72/Yeats2/Epc2/Naa12/Yeats4/Crebbp/Sf3b5/Supt3/Morf4l1/Taf6l/Taf4/Usp22/Ruvbl1/Naa50/Ruvbl2/Ogt | 33 |
| CC | GO:0034719 | SMN-Sm protein complex | 13/1488 | 17/8928 | 8,96E-08 | 1,99E-06 | 1,78E-06 | Snrpd1/Snrpd2/Snrpb/Snrpe/Snrpd3/Snrpg/Fmr1/Snrpf/Gemin8/Ddx20/Gemin4/Gemin5/Gemin7 | 13 |
| CC | GO:0042175 | nuclear outer membrane-endoplasmic reticulum membrane network | 107/1488 | 402/8928 | 1,69E-07 | 3,56E-06 | 3,17E-06 | Mtdh/Slc30a5/Fam8a1/Nucb2/Cherp/Dgat2/Nup153/Sec61a2/Ranbp2/Smpd4/Lman2l/Syne2/Sel1l/Nomo1/Ncln/Maco1/Pigg/Mia3/Ssr1/Jph4/Ergic1/Syvn1/Slc35b2/Tusc3/Hspa5/Srebf2/Rpn2/Stt3a/Ssr4/Tmed2/Ddost/Gorasp2/Lman2/Rpn1/Nos1ap/Ptdss1/Lman1/Ryr3/Ergic3/Ero1b/Sec61a1/Tmem147/Mlec/Ccdc47/Nup62/Hsp90b1/Calr/Samd8/Mogs/Atp6ap2/Tmco1/Stt3b/Pld3/Jph3/Sec16a/Cpt1c/Erlin1/Porcn/Erp44/Pdia3/Gpaa1/Sec63/Moxd1/Ccdc115/Sec11a/Erlin2/Nos1/Preb/Itpr1/Spcs1/Spcs2/Spcs3/Pigt/Rtcb/Pigs/Gramd1b/Scap/Sec11c/Pigu/Saraf/Abhd12/Pigk/Dnajb14/Srprb/Cds2/Emc7/Ero1a/Emc3/Gramd1a/Clptm1l/Dnajb12/Atp2a2/Emc10/Emc1/Emc2/Ptdss2/Derl1/Fitm2/Hacd3/Sec23a/Emc4/Rab2a/Atp13a1/Vamp7/Ufl1/Vps13c/Plpp6 | 107 |
| CC | GO:0005732 | sno(s)RNA-containing ribonucleoprotein complex | 12/1488 | 16/8928 | 4,15E-07 | 8,15E-06 | 7,28E-06 | Fbll1/Nolc1/Nop56/Nop58/Dkc1/Snu13/Gar1/Fbl/Nhp2/Rpp30/Lsm6/Rrp9 | 12 |
| CC | GO:0030134 | COPII-coated ER to Golgi transport vesicle | 22/1488 | 45/8928 | 5,41E-07 | 1,03E-05 | 9,15E-06 | Slc30a5/Tmed7/Golga2/Lman2l/Tmed10/Ergic1/Ergic2/Tmed9/Tmed2/Lman2/Tmed4/Lman1/Ergic3/Sec16a/Sec23ip/Uso1/Sec24b/Sec13/Sec23a/Sec31a/Sec24c/Sec24a | 22 |
| CC | GO:0005852 | eukaryotic translation initiation factor 3 complex | 11/1488 | 14/8928 | 5,94E-07 | 1,12E-05 | 9,98E-06 | Eif3l/Eif3k/Eif3a/Eif3c/Eif3f/Eif3i/Ddx3x/Eif3b/Eif3e/Eif3d/Eif3h | 11 |
| CC | GO:0005788 | endoplasmic reticulum lumen | 23/1488 | 49/8928 | 7,52E-07 | 1,39E-05 | 1,24E-05 | Os9/Pdia6/Erlec1/Manf/Hspa5/Lrpap1/Ero1b/Hsp90b1/Calr/Dnajc3/Pdia4/Erp29/Edem3/Erp44/Dnajc10/Pdia3/Colgalt1/Poglut1/Txndc16/Eogt/Selenof/Entpd5/Uggt1 | 23 |
| CC | GO:0000974 | Prp19 complex | 11/1488 | 17/8928 | 1,24E-05 | 0,000188 | 0,000167 | U2af2/Plrg1/Cdc5l/Prpf19/Ctnnbl1/Isy1/Bcas2/Xab2/Rbm22/Polr2a/Crnkl1 | 11 |
| CC | GO:0000139 | Golgi membrane | 46/1488 | 156/8928 | 4,18E-05 | 0,000561 | 0,000501 | Ugcg/Zdhhc17/B3gat3/Slc30a5/Golga2/Glg1/Qsox2/Rer1/Lman2l/Bet1l/Man2a2/Tmed10/Large1/Ergic1/Slc35b2/Man1a2/Casd1/Golga5/B4galnt1/Gorasp2/Lman2/Lman1/Ergic3/Gosr1/Samd8/Pgap4/Pld3/Acbd3/Numa1/Atp2c1/Chst2/Ndfip1/Scap/Copg2/Cabp1/Scyl3/Uso1/Copb2/Fut11/Copg1/Sec23a/Rab2a/Vps45/Scfd1/Pi4k2a/Scamp5 | 46 |
| CC | GO:0005793 | endoplasmic reticulum-Golgi intermediate compartment | 21/1488 | 53/8928 | 5,58E-05 | 0,000732 | 0,000653 | Tmed7/Nucb2/Rer1/Lman2l/Pdia6/Tmed10/Ergic1/Hspa5/Golgb1/Tmed9/Lrpap1/Tmed2/Lman2/Tmed4/Lman1/Erp44/Trip11/Ccdc115/Copg2/Uggt1/Copg1 | 21 |
| CC | GO:0030014 | CCR4-NOT complex | 8/1488 | 12/8928 | 0,000154 | 0,001889 | 0,001686 | Cnot6l/Cnot11/Cnot2/Cpeb3/Cnot10/Cnot7/Cnot1/Cnot3 | 8 |
| CC | GO:0005845 | mRNA cap binding complex | 7/1488 | 10/8928 | 0,000265 | 0,003113 | 0,00278 | Rnmt/Ramac/Ncbp1/Fmr1/Ncbp2/Ago2/Cyfip1 | 7 |
| CC | GO:0034518 | RNA cap binding complex | 7/1488 | 10/8928 | 0,000265 | 0,003113 | 0,00278 | Rnmt/Ramac/Ncbp1/Fmr1/Ncbp2/Ago2/Cyfip1 | 7 |
| CC | GO:0042405 | nuclear inclusion body | 7/1488 | 10/8928 | 0,000265 | 0,003113 | 0,00278 | Atxn1/Pabpn1/Nxf1/Nup153/Tpr/Ranbp2/Nup98 | 7 |
| CC | GO:0032806 | carboxy-terminal domain protein kinase complex | 10/1488 | 20/8928 | 0,000589 | 0,006471 | 0,005777 | Gtf2h1/Cdk13/Brd4/Gtf2h3/Snw1/Ccnk/Cdk9/Gtf2f2/Ercc3/Ccnt2 | 10 |
| CC | GO:0016442 | RISC complex | 7/1488 | 11/8928 | 0,000624 | 0,006706 | 0,005987 | Dhx9/Snd1/Dcp2/Prkra/Ddx6/Ago2/Ago1 | 7 |
| CC | GO:0031332 | RNAi effector complex | 7/1488 | 11/8928 | 0,000624 | 0,006706 | 0,005987 | Dhx9/Snd1/Dcp2/Prkra/Ddx6/Ago2/Ago1 | 7 |

GO_SR_protein

| ONTOLOGY | ID | Description | GeneRatio | BgRatio | pvalue | p.adjust | qvalue | geneID | Count |
|---|---|---|---|---|---|---|---|---|---|
| CC | GO:0098984 | neuron to neuron synapse | 28/110 | 452/8928 | 5,37E-13 | 1,44E-09 | 1,25E-09 | Fam81a/Arc/Slc30a3/Syt12/Dlgap3/Lrrtm2/Dcc/Psd/Rims1/Efnb2/Anks1b/Ptprs/Lrfn2/Gria2/Ephb2/Rogdi/Grm7/Homer2/Ogt/Homer1/Htt/Kcnd2/Stxbp5/Src/Kalrn/Erc1/Arhgap32/Iqsec3 | 28 |
| CC | GO:0008021 | synaptic vesicle | 17/110 | 218/8928 | 1,08E-09 | 9,67E-07 | 8,44E-07 | Slc30a3/Syt12/Rph3a/Atp8a1/Slc17a7/Slc6a17/Unc13a/Ptprs/Dmxl2/Wdr7/Gria2/Ap2a2/Stxbp5/Madd/Atp6v0a1/Erc1/Cttnbp2 | 17 |
| CC | GO:0014069 | postsynaptic density | 21/110 | 401/8928 | 1,4E-08 | 5,36E-06 | 4,67E-06 | Fam81a/Arc/Dlgap3/Lrrtm2/Dcc/Psd/Rims1/Efnb2/Anks1b/Ptprs/Lrfn2/Gria2/Homer2/Homer1/Htt/Kcnd2/Src/Kalrn/Erc1/Arhgap32/Iqsec3 | 21 |
| CC | GO:0097060 | synaptic membrane | 20/110 | 418/8928 | 1,45E-07 | 2,98E-05 | 2,6E-05 | Arc/Olfm1/Rph3a/Lrrtm2/Ntrk3/Dcc/Rims1/Unc13a/Efnb2/Ptprs/Lrfn2/Syt3/Gria2/Ephb2/Grm7/Htt/Kcnd2/Stxbp5/Src/Iqsec3 | 20 |
| CC | GO:0099522 | cytosolic region | 7/110 | 47/8928 | 1,49E-06 | 0,00021 | 0,000183 | Anks1b/Ogt/Homer1/Htt/Stxbp5/Pnkd/Erc1 | 7 |
| CC | GO:0019898 | extrinsic component of membrane | 13/110 | 227/8928 | 3,94E-06 | 0,000377 | 0,000329 | Olfm1/Rph3a/Psd/Rims1/Vps13c/Ryr2/Cadps/Stxbp5/Src/Arhgef25/Kalrn/Trio/Pik3r4 | 13 |
| CC | GO:0098685 | Schaffer collateral - CA1 synapse | 9/110 | 131/8928 | 3,18E-05 | 0,00213 | 0,001859 | Lrrtm2/Dcc/Efnb2/Anks1b/Ptprs/Lrfn2/Gria2/Stxbp5/Trio | 9 |
| CC | GO:0099568 | cytoplasmic region | 9/110 | 150/8928 | 9,2E-05 | 0,004836 | 0,00422 | Arc/Rims1/Unc13a/Gria2/Htt/Madd/Erc1/Pik3r4/Mark2 | 9 |
| CC | GO:0098982 | GABA-ergic synapse | 8/110 | 117/8928 | 9,21E-05 | 0,004836 | 0,00422 | Lrrtm2/Rims1/Slc6a17/Grm7/Ogt/Kcnd2/Erc1/Iqsec3 | 8 |
| BP | GO:0061564 | axon development | 21/110 | 381/8928 | 5,66E-09 | 3,02E-06 | 2,64E-06 | Nptx1/Islr2/Olfm1/Robo1/Ntrk3/Plxna1/Dscaml1/Epha6/Dcc/Efnb2/Ptprs/Ephb2/Cyfip2/Grm7/Stxbp5/Arhgef25/Kalrn/Arhgap32/Trio/Robo2/Mark2 | 21 |
| BP | GO:0007409 | axonogenesis | 20/110 | 348/8928 | 6,77E-09 | 3,02E-06 | 2,64E-06 | Nptx1/Islr2/Olfm1/Robo1/Ntrk3/Plxna1/Dscaml1/Epha6/Dcc/Efnb2/Ptprs/Ephb2/Cyfip2/Stxbp5/Arhgef25/Kalrn/Arhgap32/Trio/Robo2/Mark2 | 20 |
| BP | GO:0050808 | synapse organization | 21/110 | 459/8928 | 1,44E-07 | 2,98E-05 | 2,6E-05 | Nptx1/Arc/Rph3a/Dlgap3/Lrrtm2/Ntrk3/Flrt1/Sdk2/Psd/Unc13a/Efnb2/Ptprs/Lrfn2/Ephb2/Ogt/Homer1/Kalrn/Erc1/Cttnbp2/Iqsec3/Mark2 | 21 |
| BP | GO:0050804 | modulation of chemical synaptic transmission | 20/110 | 449/8928 | 4,59E-07 | 7,49E-05 | 6,53E-05 | Nptx1/Arc/Syt12/Dlgap3/Lrrtm2/Dcc/Rims1/Unc13a/Ptprs/Lrfn2/Gria2/Ephb2/Grm7/Homer1/Htt/Stxbp5/Src/Pnkd/Erc1/Trio | 20 |
| BP | GO:0099177 | regulation of trans-synaptic signaling | 20/110 | 450/8928 | 4,76E-07 | 7,49E-05 | 6,53E-05 | Nptx1/Arc/Syt12/Dlgap3/Lrrtm2/Dcc/Rims1/Unc13a/Ptprs/Lrfn2/Gria2/Ephb2/Grm7/Homer1/Htt/Stxbp5/Src/Pnkd/Erc1/Trio | 20 |
| BP | GO:0016082 | synaptic vesicle priming | 6/110 | 27/8928 | 7,31E-07 | 0,000109 | 9,49E-05 | Rph3a/Rims1/Unc13a/Cadps/Stxbp5/Erc1 | 6 |
| BP | GO:0050803 | regulation of synapse structure or activity | 14/110 | 251/8928 | 2,22E-06 | 0,000276 | 0,000241 | Arc/Lrrtm2/Ntrk3/Slc17a7/Flrt1/Psd/Ptprs/Lrfn2/Ephb2/Ogt/Homer1/Kalrn/Cttnbp2/Mark2 | 14 |
| BP | GO:1990138 | neuron projection extension | 11/110 | 153/8928 | 2,59E-06 | 0,000289 | 0,000252 | Islr2/Olfm1/Ntrk3/Flrt1/Plxna1/Rims1/Unc13a/Ptprs/Syt3/Cyfip2/Arhgap32 | 11 |
| BP | GO:0099601 | regulation of neurotransmitter receptor activity | 7/110 | 54/8928 | 3,91E-06 | 0,000377 | 0,000329 | Nptx1/Arc/Rph3a/Dlgap3/Ephb2/Homer1/Src | 7 |
| BP | GO:0060560 | developmental growth involved in morphogenesis | 12/110 | 193/8928 | 4,09E-06 | 0,000377 | 0,000329 | Islr2/Olfm1/Robo1/Ntrk3/Flrt1/Plxna1/Rims1/Unc13a/Ptprs/Syt3/Cyfip2/Arhgap32 | 12 |
| BP | GO:0046578 | regulation of Ras protein signal transduction | 10/110 | 139/8928 | 7,54E-06 | 0,000673 | 0,000587 | Robo1/Psd/Ephb2/Ogt/Sh3bp1/Madd/Src/Arhgef25/Iqsec3/Arfgef1 | 10 |
| BP | GO:0050807 | regulation of synapse organization | 13/110 | 245/8928 | 9,06E-06 | 0,000783 | 0,000683 | Arc/Lrrtm2/Ntrk3/Flrt1/Psd/Ptprs/Lrfn2/Ephb2/Ogt/Homer1/Kalrn/Cttnbp2/Mark2 | 13 |
| BP | GO:0007215 | glutamate receptor signaling pathway | 6/110 | 45/8928 | 1,68E-05 | 0,001323 | 0,001155 | Unc13a/Gria2/Grm7/Homer2/Homer1/Kalrn | 6 |
| BP | GO:0006935 | chemotaxis | 14/110 | 301/8928 | 1,8E-05 | 0,00134 | 0,001169 | Robo1/Ntrk3/Plxna1/Dscaml1/Camk1d/Epha6/Dcc/Efnb2/Ephb2/Cyfip2/Arhgef25/Kalrn/Trio/Robo2 | 14 |
| BP | GO:0042330 | taxis | 14/110 | 301/8928 | 1,8E-05 | 0,00134 | 0,001169 | Robo1/Ntrk3/Plxna1/Dscaml1/Camk1d/Epha6/Dcc/Efnb2/Ephb2/Cyfip2/Arhgef25/Kalrn/Trio/Robo2 | 14 |
| BP | GO:0099068 | postsynapse assembly | 6/110 | 47/8928 | 2,17E-05 | 0,001553 | 0,001355 | Nptx1/Lrrtm2/Ntrk3/Psd/Ptprs/Ephb2 | 6 |
| BP | GO:0048588 | developmental cell growth | 11/110 | 194/8928 | 2,51E-05 | 0,001723 | 0,001504 | Islr2/Olfm1/Ntrk3/Flrt1/Plxna1/Rims1/Unc13a/Ptprs/Syt3/Cyfip2/Arhgap32 | 11 |
| BP | GO:0031346 | positive regulation of cell projection organization | 14/110 | 321/8928 | 3,68E-05 | 0,002393 | 0,002088 | Islr2/Robo1/Ntrk3/Plxna1/Camk1d/Dcc/Ephb2/Htt/Nckipsd/Src/Kalrn/Arhgap32/Robo2/Mark2 | 14 |
| BP | GO:0099003 | vesicle-mediated transport in synapse | 12/110 | 242/8928 | 4,01E-05 | 0,002452 | 0,002139 | Arc/Syt12/Rph3a/Slc17a7/Rims1/Unc13a/Efnb2/Ap2a2/Cadps/Stxbp5/Atp6v0a1/Erc1 | 12 |

| | | | | | | | | | |
|---|---|---|---|---|---|---|---|---|---|
| BP | GO:0006836 | neurotransmitter transport | 10/110 | 176/8928 | 5,87E-05 | 0,003344 | 0,002918 | Syt12/Rph3a/Slc17a7/Rims1/Slc6a17/Unc13a/Cadps/Stxbp5/Pnkd/Erc1 | 10 |
| BP | GO:0017156 | calcium-ion regulated exocytosis | 6/110 | 63/8928 | 0,000117 | 0,005917 | 0,005163 | Syt12/Rph3a/Rims1/Unc13a/Syt3/Cadps | 6 |
| BP | GO:0098962 | regulation of postsynaptic neurotransmitter receptor activity | 4/110 | 22/8928 | 0,000134 | 0,006667 | 0,005817 | Nptx1/Dlgap3/Homer1/Src | 4 |
| BP | GO:0007265 | Ras protein signal transduction | 11/110 | 242/8928 | 0,000185 | 0,009005 | 0,007857 | Robo1/Psd/Ephb2/Ogt/Sh3bp1/Madd/Src/Arhgef25/Arhgap32/Iqsec3/Arfgef1 | 11 |
| MF | GO:0008046 | axon guidance receptor activity | 3/110 | 10/8928 | 0,000205 | 0,009308 | 0,008122 | Robo1/Ephb2/Robo2 | 3 |
| MF | GO:0005085 | guanyl-nucleotide exchange factor activity | 9/110 | 167/8928 | 0,000208 | 0,009308 | 0,008122 | Psd/Dock9/Madd/Arhgef25/Kalrn/Trio/Tbc1d10a/Iqsec3/Arfgef1 | 9 |

GO_SLM_protein

| ONTOLOGY | ID | Description | GeneRatio | BgRatio | pvalue | p.adjust | qvalue | geneID |
|---|---|---|---|---|---|---|---|---|
| CC | GO:0005743 | mitochondrial inner membrane | 189/1360 | 419/8928 | 6,75E-51 | 3,88E-47 | 3,29E-47 | Gcat/Slc25a24/Slc25a21/Tst/Prodh/Abcb10/Ndufa3/Tomm22/Idh2/Sfxn5/Fdxr/Mccc1/Glud1/Hadh/Eci1/Bdh1/Ndufv3/Mpst/Gstk1/Them4/Sqor/Mfn1/Sfxn1/Slc25a23/Ndufa1/Maob/Hmgcl/Uqcrb/Gcdh/Ppox/Bcs1l/Nipsnap1/Crat/Fahd1/Ndufa12/Abcb7/Micos13/Sfxn3/Atp23/Sfxn2/Tmem223/Tk2/Mtarc2/Aifm1/Letm1/Acad9/Exog/Ndufa10/Hsd17b10/Uqcrc1/Cox5b/Bcl2l1/Cox20/Apoo/Sdhb/Cpt1a/Timm21/Ptpmt1/Ckmt1/Hadha/Tamm41/Acat1/Timm22/Ndufs2/Atp5c1/Phb2/Stoml2/Tmem65/Tmem256/Afg3l2/Samm50/Sod2/Ndufv2/Spg7/Ndufb11/Hadhb/Maip1/Tufm/Ndufa13/Cox5a/Slc25a11/Tmem177/Slc25a10/Ndufb9/Immt/Etfdh/Ndufs3/Uqcrfs1/Gpd2/Ndufa2/Rdh13/Ndufa5/Shmt2/Ndufs7/Atp5pb/Agk/Ndufv1/Sirt5/Ndufa6/Ndufs1/Slc25a12/Guf1/Cox6c/Timm29/Fdx1/Dnajc19/Micu1/Apool/Slc25a40/Slc25a3/Mtx1/Ak2/Ndufs5/Timm44/Atp5o/Dnajc11/Uqcrc2/Timm50/Ndufaf5/Pmpca/Vdac1/Coq9/Ndufc2/Hspd1/Ccdc51/Cpt2/Coq6/Uqcrq/Mcu/Mdh2/Slc25a15/Tomm40/Abcb8/Slc25a44/Ndufb4/Slc25a19/Ndufa9/Chchd3/Coq3/Sdha/Ndufa8/Micu3/Acadvl/Cox4i1/Got2/Slc25a4/Slc25a42/Aldh18a1/Lyn/Fech/Acaa2/Slc25a1/Cyc1/Dmac2l/Ndufaf3/Mrpl53/Cox15/Timm17b/Vdac2/Mrpl24/Mrpl18/Ndufb6/Pmpcb/Opa1/Cox7a2l/Sirt3/Mrpl12/Adck1/Slc25a51/Tmem126a/Slc25a5/Cyb5b/Mrs2/Slc25a25/Dhrs1/Mtch2/Dhodh/Mrps22/Mrpl23/Mrpl9/Mrps34/Mrpl14/Pde2a/Slc25a20/Mrpl13/Aifm3/Trap1/Vdac3/Mrpl58 |
| CC | GO:0098798 | mitochondrial protein-containing complex | 107/1360 | 243/8928 | 1,15E-27 | 2,21E-24 | 1,88E-24 | Suclg2/Ndufa3/Tomm22/Mccc1/Mccc2/Ndufv3/Mfn1/Fxn/Ndufa1/Uqcrb/Ndufa12/Dbt/Etfa/Micos13/Ndufa10/Hsd17b10/Pdhx/Uqcrc1/Cox5b/Apoo/Sdhb/Timm21/Bax/Hadha/Timm22/Ndufs2/Atp5c1/Phb2/Etfb/Afg3l2/Samm50/Mtx2/Ndufv2/Spg7/Ndufb11/Hadhb/Ndufa13/Dld/Cox5a/Ndufb9/Immt/Etfdh/Ndufs3/Uqcrfs1/Ndufa2/Ndufa5/Pdha1/Trmt10c/Ndufs7/Atp5pb/Agk/Ndufv1/Ndufa6/Idh3g/Ndufs1/Tomm40l/Cox6c/Timm29/Dnajc19/Micu1/Pdhb/Apool/Mtx1/Ndufs5/Atp5o/Dnajc11/Uqcrc2/Nfs1/Timm50/Idh3a/Supv3l1/Vdac1/Mrps36/Ndufc2/Pnpt1/Uqcrq/Mcu/Tomm40/Pdk1/Ndufb4/Ndufa9/Chchd3/Sdha/Ndufa8/Micu3/Cox4i1/Slc25a4/Bckdha/Cyc1/Idh3b/Mrpl53/Timm17b/Mrpl24/Mrpl18/Ndufb6/Pmpcb/Cox7a2l/Mrpl12/Dlat/Slc25a5/Mrps22/Mrpl23/Mrpl9/Mrps34/Mrpl14/Mrpl13/Mrpl58 |
| CC | GO:1990204 | oxidoreductase complex | 55/1360 | 95/8928 | 8,52E-22 | 4,45E-19 | 3,78E-19 | Ndufa3/Gcsh/Ndufv3/Ndufa1/Uqcrb/Ndufa12/Dbt/Etfa/Gldc/Ndufa10/Pdhx/Uqcrc1/Sdhb/Dlst/Ndufs2/Etfb/Ogdhl/Ndufv2/Ndufb11/Cbr4/Ndufa13/Dld/Ndufb9/Hsd17b8/Etfdh/Ndufs3/Uqcrfs1/Gpd2/Ndufa2/Ndufa5/Pdha1/Ndufs7/Ndufv1/Ndufa6/Idh3g/Ndufs1/Pdhb/Ndufs5/Uqcrc2/Idh3a/Ogdh/Mrps36/Ndufc2/Uqcrq/Pdk1/Ndufb4/Ndufa9/Sdha/Ndufa8/Bckdha/Cyc1/Idh3b/Ndufb6/Dlat/Cyb5b |
| CC | GO:0097060 | synaptic membrane | 132/1360 | 418/8928 | 5,2E-18 | 1,15E-15 | 9,77E-16 | Grm2/Ntng1/Kcnj4/Grm8/Cplx3/Grm4/Il1rapl1/Lrrc4c/Igsf9b/Kcnj9/Slc16a3/Cntn6/Kcnj3/Adra2a/Atp2b4/Kcnj6/Kcna3/Lrp4/Ptprf/Tiam1/Hcn1/Ptprt/Kctd8/Lrrtm4/Ptprd/Dlg1/Itga3/Slc6a4/Nrxn2/Olfm2/Slc8a3/Kctd12/Ddn/Cspg5/Nrxn1/Itsn2/Gpc4/Adcy8/Hspb1/Gabrg1/Slc4a8/Slc1a2/Fxyd6/Entpd1/Shisa9/Nlgn1/Adgrl2/Ctnna2/Kcnc4/Dgki/Lrfn4/Nlgn3/Grik5/Cdh8/Grik2/Dbn1/Stx1a/Itgb1/Adgrb3/Pip5k1c/Nlgn4l/Lrfn5/L1cam/Cdh10/Lrfn3/Grm3/Gabra4/Nipsnap1/Magi2/Adcy1/Syt7/Pcdh17/Grid2/Nlgn2/Hip1r/Rims2/Fbxo2/Grid1/Apba1/Gabrd/Itgb5/Lrfn1/Neto2/Cacna2d1/Cdh2/Snta1/Slc6a2/Ptprz1/Slc6a1/Epb41l1/Dbnl/Adgrl1/Ndufs7/Flrt3/Slc2a3/Dnm3/Gphn/Grip2/Vdac1/Dag1/Gabbr2/Ephb2/Adgrl3/Cask/Palm/Nrcam/Adam22/Efnb1/Ctnnb1/Tenm2/Slitrk1/Pacsin1/Grip1/Pten/Iqsec2/Stxbp1/Lrrc4b/Dnaja3/Syne1/Ppp1r9b/Kctd16/Abhd17a/Atp2b2/Pde2a/Gabbr1/Septin3/Htr2a/Slitrk3/Dnm1l/Crkl/Gabrb1/Iqsec3 |
| CC | GO:0070161 | anchoring junction | 130/1360 | 453/8928 | 4,25E-14 | 6,11E-12 | 5,19E-12 | Calb2/Camk2d/Mylk/Lama1/Pdlim7/Cav1/Fermt1/Flna/Pacsin2/Cav2/Cdh13/Tiam1/Aqp4/Magi1/Des/Cxadr/Itga7/Dlg1/Actn1/Kirrel3/Slc6a4/Tgfb1i1/Gja1/Ptprj/Nrxn1/Ptprk/Pecam1/Vangl2/Plekha7/Tns2/Esam/Ahnak/Sh3kbp1/Gjb6/Ptprm/Trpc4/Rasip1/Lpp/Ctnna2/Cdh12/Tns1/Podxl/Lasp1/Anxa5/Afap1/Cdh8/Stard10/Dbn1/Pak1/Itgb1/Nectin2/Itga1/Itgb8/Iqgap1/Cdh10/Pxn/Magi2/Pdlim5/Atp1a1/Cdh20/Sorbs1/Cd2ap/Cdh6/Tln1/Fscn1/Ccdc85a/Pals2/Cdh5/Plec/Ctnnd2/Pcdhgc3/Pkp4/Abcb1a/Mlc1/Actn4/Vcl/Tln2/Afdn/Itgav/Ptpn6/Itgb5/Cgnl1/Jcad/Cdh2/Tmem65/Rexo2/Pcdh9/Flrt3/Evl/Tjp1/Itga6/Add3/Jam2/Ptprc/Pdzd11/Coro2b/Dag1/Synpo/Mapk3/Ilk/Arhgef2/Adgrl3/Cask/Niban2/Nck1/Lyn/Mapre1/Coro1a/Ajm1/Ctnnb1/Specc1l/Tenm2/Capn2/Hepacam/Wnk3/Sptbn2/Twf1/Atp1a2/Itgb2/Arhgef7/Pdxp/Wasf2/Csk/Lcp1/Ctnnd1/Pard6a/Pdcd6ip/Flrt2/Pak2/Sptan1 |
| CC | GO:0098803 | respiratory chain complex | 36/1360 | 65/8928 | 6,87E-14 | 9,18E-12 | 7,8E-12 | Ndufa3/Ndufv3/Ndufa1/Uqcrb/Ndufa12/Ndufa10/Uqcrc1/Cox5b/Sdhb/Ndufs2/Ndufv2/Ndufb11/Ndufa13/Cox5a/Ndufb9/Ndufs3/Uqcrfs1/Ndufa2/Ndufa5/Ndufs7/Ndufv1/Ndufa6/Ndufs1/Cox6c/Ndufs5/Uqcrc2/Ndufc2/Uqcrq/Ndufb4/Ndufa9/Sdha/Ndufa8/Cox4i1/Cyc1/Ndufb6/Cox7a2l |
| CC | GO:0070469 | respirasome | 36/1360 | 69/8928 | 7,76E-13 | 9,11E-11 | 7,73E-11 | Ndufa3/Ndufv3/Ndufa1/Uqcrb/Ndufa12/Ndufa10/Uqcrc1/Cox5b/Sdhb/Ndufs2/Ndufv2/Ndufb11/Ndufa13/Cox5a/Ndufb9/Ndufs3/Uqcrfs1/Ndufa2/Ndufa5/Ndufs7/Ndufv1/Ndufa6/Ndufs1/Cox6c/Ndufs5/Uqcrc2/Ndufc2/Uqcrq/Ndufb4/Ndufa9/Sdha/Ndufa8/Cox4i1/Cyc1/Ndufb6/Cox7a2l |
| CC | GO:0098636 | protein complex involved in cell adhesion | 20/1360 | 30/8928 | 2,58E-10 | 2,18E-08 | 1,85E-08 | Tnc/Lama1/Lamb2/Itga7/Itga3/Lgals1/Lamc1/Nid1/Nrxn1/Nlgn1/Itgb1/Itga1/Itgb8/Itgav/Itgb5/Itga6/Jam2/Lyn/Itgb2/Itgam |
| CC | GO:0045121 | membrane raft | 79/1360 | 260/8928 | 2,76E-10 | 2,3E-08 | 1,95E-08 | Cd44/Cavin1/Cav1/Ehd2/Atp2b4/Kcna3/Mme/Pacsin2/Cav2/Lrp4/Cdh13/Cavin2/Cxadr/Dlg1/Slc6a4/Gja1/Icam1/Trpc1/Trpc5/Pecam1/Adcy8/Nos3/Angpt1/Ahnak/Slc1a2/Trpc4/Slc1a1/H2-D1/Itgb1/Itga1/Prkar2b/Iqgap1/Serpinh1/Myo1c/L1cam/Il6st/Atp1a1/Sorbs1/Mlc1/Vcl/Insr/Cdh2/Slc6a2/Stoml2/Casp3/Igf1r/Slc2a3/Ptprc/Prkaca/Adcy6/Dag1/Hspd1/Mapk3/Cask/Slc25a4/Lyn/Adcy2/Efnb1/Ctnnb1/Capn2/Pld2/Vdac2/Grip1/Atp1a2/Itgb2/Slc25a5/Cr1l/Csk/Itgam/Rgs19/Atp2b2/Pde2a/Add2/Ctnnd1/Hk1/Gabbr1/Hdac6/Coro1c/Gnai2 |
| CC | GO:0045239 | tricarboxylic acid cycle enzyme complex | 15/1360 | 19/8928 | 1,09E-09 | 7,72E-08 | 6,56E-08 | Suclg2/Dbt/Pdhx/Dlst/Ogdhl/Dld/Idh3g/Pdhb/Idh3a/Ogdh/Mrps36/Sucla2/Bckdha/Idh3b/Dlat |
| CC | GO:0009986 | cell surface | 107/1360 | 414/8928 | 6,86E-09 | 4,38E-07 | 3,72E-07 | Cd44/Il1rapl1/Vwf/Aoc3/Kcnj3/Cav1/Kcnj6/Gfra2/Cav2/Lrp4/Cdh13/Aqp4/Hcn1/Ptprt/Itga7/Pdgfrb/Itga3/Lgals1/Bcam/Lrrc24/Ace/Acvr2a/Icam1/Ptprj/Nrxn1/Agrn/Gpc4/Ptprk/Pecam1/Cd93/H2-K1/Slc7a11/Slc1a2/Antxr1/Trpc4/Entpd1/Ace2/Nlgn1/Pam/Kcnc4/Lrfn4/Dcbld2/Slc1a1/Anxa5/C1qbp/Nlgn3/H2-D1/Vldlr/Gpc5/Cd200/Itgb1/Nectin2/Itga1/Itgb8/Fgb/Nlgn4l/Lrfn5/Gpc6/Robo2/L1cam/Lrfn3/F3/Anpep/Fgg/Il6st/Anxa4/Cdh5/Abcb1a/Nlgn2/Itgav/Insr/Itgb5/Lrfn1/Cdh2/Kcnk2/Enpep/Phb2/Slc6a2/Slc6a1/Neo1/Il1r1/Tjp1/Cntfr/Itga6/Jam2/Abcc4/Ptprc/Atp5o/Cdon/Dag1/Hspd1/Atp1b2/Ephb2/Got2/Fgfr3/Vcam1/Capn2/Map3k5/Alcam/Abcg1/Itgb2/Cr1l/Abca7/Itgam/Vcan/Slitrk3/Rnpep |
| CC | GO:0098982 | GABA-ergic synapse | 43/1360 | 117/8928 | 7,6E-09 | 4,8E-07 | 4,08E-07 | Cbln4/Grm8/Igsf9b/Adra2a/Cdh13/Gad2/Lrrtm4/Cspg5/Nrxn1/Gap43/Nlgn1/Arhgef9/Lrfn4/C1qbp/Nlgn3/Pak1/Lrfn5/Cdh10/Gabra4/Magi2/Mdga2/Syt7/Pcdh17/Septin11/Nlgn2/Rims2/Grid1/Gabrd/Phb2/Slc6a1/Calb1/Gphn/Dag1/Gabbr2/Slitrk1/Grip1/Arhgef7/Atp2b2/Gabbr1/Slitrk3/Hap1/Gabrb1/Iqsec3 |
| CC | GO:0098984 | neuron to neuron synapse | 112/1360 | 452/8928 | 3,72E-08 | 2,28E-06 | 1,93E-06 | Grm4/Il1rapl1/Lrrc4c/Igsf9b/Slc16a3/Adra2a/Lrp4/Ptprf/Tiam1/Ptprt/Sorbs2/Lrrtm4/C1ql3/Ptprd/Dlg1/Cald1/Slc8a3/Plxnc1/Nrxn1/Adcy8/Hspb1/Gap43/Myo6/Epb41/Slc4a8/Camk1/Entpd1/Akap7/Shisa9/Nlgn1/Arhgef9/Ctnna2/Dgki/Lrfn4/Slc1a1/Grik5/Camk2g/Grik2/Dbn1/Stx1a/Pak1/Itgb1/Adgrb3/Pip5k1c/Lrfn5/Pde4b/Lrfn3/Grm3/Map2/Magi2/Pdlim5/Atp1a1/Srcin1/Adcy1/Syt7/Snx1/Grid2/Ctnnd2/Pkp4/Septin11/Hip1r/Grid1/Lrfn1/Neto2/Cdh2/Phb2/Ptprz1/Calb1/Epb41l1/Dbnl/Il1r1/Adgrl1/Flrt3/Add3/Dnm3/Gphn/Grip2/Vdac1/Atp1b2/Ephb2/Arhgef2/Cask/Palm/Nrcam/Adam22/Lyn/Ctnnb1/Slitrk1/Nf1/Dapk1/Tnik/Pacsin1/Dclk1/Grip1/Iqsec2/Arhgef7/Lrrc4b/Ppp1r9b/Abhd17a/Plekha5/Inpp4a/Atp2b2/Add2/Ctnnd1/Slitrk3/Sh3gl1/Tanc2/Pak2/Abr/Sptbn1/Iqsec3/Kalrn |
| CC | GO:0098858 | actin-based cell projection | 48/1360 | 147/8928 | 8,11E-08 | 4,61E-06 | 3,92E-06 | Cd44/Calb2/Aoc3/Slc4a7/Whrn/Cxadr/Dlg1/Itga3/Icam2/Gap43/Myo6/Angpt1/Antxr1/Utrn/Nlgn1/Podxl/Rapgef3/Morn4/Stard10/Dbn1/Itgb1/Myo1c/Map2/Srcin1/Fscn1/Shtn1/Itgav/Ttyh1/Nherf2/Ptprz1/Calb1/Itga6/Adcy6/Dag1/Dync2i2/Palm/Tubb3/Myo5a/Myo1b/Coro1a/Ctnnb1/Vcam1/Wwox/Twf1/Pls3/Ppp1r9b/Lcp1/Rufy3 |

| | | | | | | | | |
|---|---|---|---|---|---|---|---|---|
| CC | GO:0099572 | postsynaptic specialization | 107/1360 | 433/8928 | 8,88E-08 | 4,95E-06 | 4,21E-06 | Il1rapl1/Lrrc4c/Igsf9b/Slc16a3/Adra2a/Lrp4/Ptprf/Tiam1/Ptprt/Sorbs2/Lrrtm4/Dlg1/Cald1/Slc8a3/Adcy8/Hspb1/Gap43/Myo6/Epb41/Camk1/Entpd1/Shisa9/Nlgn1/Arhgef9/Ctnna2/Dgki/Lrfn4/Nlgn3/Grik5/Camk2g/Grik2/Dbn1/Stx1a/Pak1/Adgrb3/Pip5k1c/Nlgn4l/Lrfn5/Pde4b/Cdh10/Lrfn3/Grm3/Gabra4/Map2/Magi2/Pdlim5/Atp1a1/Srcin1/Adcy1/Snx1/Grid2/Ctnnd2/Pkp4/Septin11/Nlgn2/Hip1r/Grid1/Lrfn1/Neto2/Cdh2/Phb2/Ptprz1/Epb41l1/Dbnl/Il1r1/Adgrl1/Flrt3/Add3/Dnm3/Gphn/Grip2/Vdac1/Arhgef2/Cask/Palm/Nrcam/Adam22/Lyn/Ctnnb1/Slitrk1/Yes1/Nf1/Dapk1/Tnik/Pacsin1/Dclk1/Grip1/Iqsec2/Arhgef7/Lrrc4b/Ppp1r9b/Abhd17a/Plekha5/Inpp4a/Dlgap4/Atp2b2/Add2/Ctnnd1/Slitrk3/Sh3gl1/Tanc2/Pak2/Gabrb1/Abr/Sptbn1/Iqsec3/Kalrn |
| CC | GO:0031253 | cell projection membrane | 62/1360 | 213/8928 | 1,37E-07 | 7,41E-06 | 6,29E-06 | Cd44/Tpm1/Tiam1/Aqp4/Hcn1/Lamp5/Dlg1/Itga3/Ace/Slc5a6/Ptprj/Gap43/Myo6/Gabrg1/Slc7a11/Slc1a2/Antxr1/Utrn/Hcn2/Shisa9/Ace2/Slc39a6/Kcnc4/Podxl/Rapgef3/Itgb1/Sgce/Robo2/Myo1c/Anpep/Gabra4/Map2/Psd2/Fscn1/Hip1r/Prph/Akt2/Itgav/Insr/Ttyh1/Jcad/Enpep/Ptprz1/Adcy6/Atp1b2/Bbs9/Arhgef2/Cask/Palm/Mapre1/Ctnnb1/Pld2/Pacsin1/Twf1/Appl2/Pdxp/Ppp1r9b/Abca7/Atp2b2/Bbs7/Gabbr1/Sptbn1 |
| CC | GO:0098797 | plasma membrane protein complex | 81/1360 | 305/8928 | 1,45E-07 | 7,75E-06 | 6,59E-06 | Dpp10/Kcnj3/Cav1/Hcn3/Flna/Kcna3/Cav2/Cdh13/Hcn1/Hcn4/Lrrtm4/Itga7/Dlg1/Sntb1/Itga3/Acvr2a/Olfm2/Gja1/Gnal/H2-K1/Ahnak/Olfm3/Gng4/Gjb6/Utrn/Hcn2/Shisa9/Ctnna2/Cdh12/Kcnc4/Grik5/H2-D1/Cdh8/Grik2/Itgb1/Sgce/Itga1/Itgb8/Pde4b/Cdh10/Il6st/Atp1a1/Cdh20/Sorbs1/Htra2/Cdh6/Grid2/Cdh5/Itgav/Insr/Ptpn6/Itgb5/Cacna2d1/Cacna2d3/Cdh2/Snta1/Kcnk2/Casp3/Igf1r/Neo1/Kcna6/Cntfr/Itga6/Dag1/Eps15l1/Atp1b2/Gabbr2/Lyn/Gnb5/Ctnnb1/Vcam1/Alcam/Atp1a2/Itgb2/Itgam/Ctnnd1/Dtna/Gabbr1/Chuk/Coro1c/Gnai2 |
| CC | GO:0015629 | actin cytoskeleton | 91/1360 | 355/8928 | 1,46E-07 | 7,75E-06 | 6,59E-06 | Myh11/Calb2/Tpm1/Mylk/Tpm2/Pdlim7/Flna/Arhgap6/Whrn/Actn1/Cald1/Gas7/Dgkh/Adcy8/Vangl2/Myo6/Epb41/Ahnak/Gjb6/Ablim3/Lpp/Lmod1/Ctnna2/Lasp1/Swap70/Afap1/Rapgef3/Asap1/Dbn1/Stx1a/Pak1/Iqgap1/Myo1c/Ablim2/Map2/Myl6b/Pxn/Pdlim5/Srcin1/Sorbs1/Arhgap33/Cd2ap/Tmod1/Fscn1/Rac3/Septin11/Actn4/Vcl/Cgnl1/Cdh2/Stoml2/Myl9/Tax1bp3/Calb1/Carmil1/Dbnl/Coro2b/Synpo/Bag3/Ilk/Epb41l2/Arhgef2/Mapk8/Myo5a/Myo1b/Coro1a/Vcam1/Specc1l/Yes1/Dapk1/Sptbn2/Twf1/Cfl2/Pdxp/Dnaja3/Pls3/Ppp1r9b/Septin5/Taok2/Mprip/Add2/Lcp1/Ptpn12/Pdcd6ip/Coro1c/Dpysl3/Cdc42bpb/Myh10/Sptbn1/Hax1/Sptan1 |
| CC | GO:0009897 | external side of plasma membrane | 45/1360 | 137/8928 | 1,69E-07 | 8,85E-06 | 7,51E-06 | Cd44/Vwf/Kcnj3/Gfra2/Cdh13/Aqp4/Itga7/Itga3/Ace/Icam1/Gpc4/Pecam1/H2-K1/Antxr1/Entpd1/Nlgn1/Anxa5/H2-D1/Itgb1/Itga1/Fgb/L1cam/Anpep/Fgg/Il6st/Cdh5/Itgav/Insr/Enpep/Il1r1/Cntfr/Itga6/Abcc4/Ptprc/Dag1/Atp1b2/Vcam1/Capn2/Map3k5/Alcam/Abcg1/Itgb2/Cr1l/Itgam/Rnpep |
| CC | GO:0031252 | cell leading edge | 87/1360 | 337/8928 | 1,97E-07 | 1E-05 | 8,52E-06 | Cd44/Plxnd1/Tpm1/Mylk/Pdlim7/Tiam1/Hcn1/Lamp5/Pdgfrb/Actn1/Gas7/Ptprj/Ptprk/Pecam1/Myo6/Gabrg1/Slc1a2/Ablim3/Antxr1/Ptprm/Hcn2/Shisa9/Slc39a6/Vim/Ctnna2/Kcnc4/Podxl/Rapgef3/Dbn1/Pak1/Itgb1/Sgce/Iqgap1/Robo2/Myo1c/Gabra4/Map2/Pxn/Psd2/Srcin1/Cd2ap/Tln1/Fscn1/Hip1r/Akt2/Tln2/Shtn1/Itgav/Insr/Itgb5/Jcad/Cdh2/Ptprz1/Carmil1/Dbnl/Evl/Cspg4/Dag1/Ilk/Arhgef2/Palm/Tubb3/Myo5a/Coro1a/Abi3/Ctnnb1/Pld2/Pacsin1/Twf1/Appl2/Arhgef7/Pdxp/Wasf2/Ppp1r9b/Abca7/Atp2b2/Lcp1/Ctnnd1/Gabbr1/Hdac6/Wasf3/Coro1c/Dpysl3/Myh10/Rufy3/Sptbn1/Hax1 |
| CC | GO:0043235 | receptor complex | 59/1360 | 203/8928 | 2,91E-07 | 1,45E-05 | 1,23E-05 | Cd44/Plxnd1/Adra2a/Gfra2/Flna/Kctd8/Lrrtm4/Itga7/Pdgfrb/Itga3/Acvr2a/Olfm2/Plxnc1/Kctd12/Trpc1/Gabrg1/Olfm3/Plxna3/Pex5l/Shisa9/Nlgn1/Grik5/Vldlr/Lrp1b/Grik2/Itgb1/Itpr3/Itga1/Itgb8/Ptprn2/Gabra4/Il6st/Htra2/Grid2/Itgav/Insr/Gabrd/Ptpn6/Itgb5/Ddr1/Igf1r/Cntfr/Itga6/Hspd1/Gabbr2/Plxnb2/Fgfr3/Lyn/Alcam/Itgb2/Cr1l/Kctd16/Itgam/Taok2/Itpr2/Gabbr1/Htr2a/Chuk/Gabrb1 |
| CC | GO:0009295 | nucleoid | 19/1360 | 39/8928 | 8,5E-07 | 3,41E-05 | 2,9E-05 | Lonp1/Dbt/Ssbp1/Hsd17b10/Hadha/Sod2/Hadhb/Tufm/Shmt2/Trmt10c/Supv3l1/Vdac1/Tfam/Acadvl/Fastkd5/Vdac2/Dnaja3/Slc25a5/Lrpprc |
| CC | GO:1902495 | transmembrane transporter complex | 68/1360 | 253/8928 | 9,23E-07 | 3,68E-05 | 3,13E-05 | Slc17a6/Dpp10/Kcnj3/Hcn3/Kcna3/Hcn1/Hcn4/Lrrtm4/Olfm2/Trpc1/Trpc5/Gabrg1/Olfm3/Pex5l/Ndufa3/Trpc4/Hcn2/Shisa9/Kcnc4/Ndufv3/Grik5/Grik2/Ndufa1/Uqcrb/Pde4b/Gabra4/Atp1a1/Ndufa12/Grid2/Ndufa10/Uqcrc1/Gabrd/Ttyh1/Cacna2d1/Cacna2d3/Sestd1/Kcnk2/Ndufs2/Ndufv2/Ndufb11/Ndufa13/Ndufb9/Ndufs3/Uqcrfs1/Ndufa2/Ndufa5/Ndufs7/Kcna6/Ndufv1/Ndufa6/Ndufs1/Micu1/Ndufs5/Uqcrc2/Ndufc2/Ccdc51/Uqcrq/Mcu/Abcb8/Atp1b2/Ndufb4/Ndufa9/Ndufa8/Micu3/Cyc1/Ndufb6/Atp1a2/Gabrb1 |
| CC | GO:1990351 | transporter complex | 71/1360 | 270/8928 | 1,29E-06 | 5E-05 | 4,24E-05 | Slc17a6/Dpp10/Kcnj3/Hcn3/Kcna3/Hcn1/Hcn4/Lrrtm4/Olfm2/Trpc1/Trpc5/Gabrg1/Olfm3/Pex5l/Ndufa3/Trpc4/Hcn2/Shisa9/Kcnc4/Ndufv3/Grik5/Grik2/Ndufa1/Uqcrb/Pde4b/Gabra4/Atp1a1/Ndufa12/Grid2/Atp11a/Ndufa10/Uqcrc1/Gabrd/Ttyh1/Cacna2d1/Cacna2d3/Sestd1/Kcnk2/Ndufs2/Ndufv2/Ndufb11/Atp8a1/Ndufa13/Ndufb9/Ndufs3/Uqcrfs1/Ndufa2/Ndufa5/Ndufs7/Kcna6/Ndufv1/Ndufa6/Ndufs1/Micu1/Ndufs5/Uqcrc2/Ndufc2/Ccdc51/Uqcrq/Mcu/Abcb8/Atp1b2/Ndufb4/Ndufa9/Ndufa8/Micu3/Cyc1/Ndufb6/Atp1a2/Atp11b/Gabrb1 |
| CC | GO:0019867 | outer membrane | 49/1360 | 165/8928 | 1,48E-06 | 5,59E-05 | 4,75E-05 | Ass1/Slc8a3/Gja1/Tomm22/Mfn1/Itpr3/Maob/Fis1/Mavs/Plec/Sfxn2/Mtarc2/Aifm1/Maoa/Bcl2l1/Nipsnap2/Cpt1a/Bax/Nlrx1/Phb2/Bad/Samm50/Mtx2/Hadhb/Tomm40l/Mtx1/Dnajc11/Rab11fip5/Rtn4ip1/Vdac1/Tomm40/Gk/Rhot1/Slc25a4/Vdac2/Qtrt1/Opa1/Pgam5/Rmdn3/Cyb5b/Mtch2/Armc1/Pde2a/Hk1/Acsl6/Vdac3/Dnm1l/Hax1/Clmn |
| CC | GO:0044853 | plasma membrane raft | 32/1360 | 95/8928 | 5,55E-06 | 0,000172 | 0,000146 | Cavin1/Cav1/Ehd2/Pacsin2/Cav2/Lrp4/Cdh13/Cavin2/Adcy8/Nos3/Trpc4/Atp1a1/Sorbs1/Mlc1/Insr/Cdh2/Igf1r/Slc2a3/Prkaca/Dag1/Mapk3/Ctnnb1/Pld2/Atp1a2/Itgb2/Cr1l/Itgam/Add2/Ctnnd1/Hk1/Hdac6/Coro1c |
| CC | GO:0098862 | cluster of actin-based cell projections | 34/1360 | 104/8928 | 6,04E-06 | 0,000186 | 0,000158 | Myh11/Calb2/Slc4a7/Flna/Mme/Tiam1/Whrn/Actn1/Ace/Slc5a6/Myo6/Slc7a11/Ace2/Rapgef3/Morn4/Itpr3/Myo1c/Anpep/Plec/Coro2a/Snx5/Ddr1/Nherf2/Enpep/Calb1/Add3/Gipc1/Myo1b/Coro1a/Pld2/Pls3/Rgs19/Dnm1l/Myh10 |
| CC | GO:0062023 | collagen-containing extracellular matrix | 50/1360 | 180/8928 | 9,52E-06 | 0,000284 | 0,000242 | Col1a2/Col6a3/Vwf/Col6a1/Tnc/Reln/Col4a2/Lama1/Prelp/Matn4/Col15a1/Hspg2/Tgm2/Lama5/Itih5/Timp3/Thbs4/Lamb2/Dlg1/Lgals1/Lamc1/F13a1/Nid1/Vwa1/Agrn/Gpc4/Angpt1/Entpd1/Lama4/Tspan9/Anxa5/Gpc5/Itgb1/Fgb/Gpc6/Serpinh1/F3/Fgg/Mmrn2/Anxa4/Lgals9/Ptprz1/Cspg4/Itga6/Dag1/Anxa3/Rell2/Plxnb2/Cask/Vcan |
| CC | GO:0097386 | glial cell projection | 16/1360 | 34/8928 | 1,11E-05 | 0,000324 | 0,000276 | Grm2/Aqp4/Slc4a8/Slc7a11/Slc1a2/Slc1a1/Itgb1/Grm3/Slc2a13/Mlc1/Adgrg1/Kcnk2/Atp1b2/Abca7/Lcp1/Wasf3 |
| CC | GO:0031256 | leading edge membrane | 39/1360 | 130/8928 | 1,32E-05 | 0,000382 | 0,000324 | Cd44/Tpm1/Tiam1/Hcn1/Lamp5/Ptprj/Ptprk/Myo6/Gabrg1/Slc1a2/Antxr1/Hcn2/Shisa9/Slc39a6/Kcnc4/Itgb1/Sgce/Robo2/Myo1c/Gabra4/Map2/Psd2/Hip1r/Akt2/Itgav/Insr/Jcad/Ptprz1/Arhgef2/Palm/Pacsin1/Twf1/Appl2/Pdxp/Ppp1r9b/Abca7/Atp2b2/Gabbr1/Sptbn1 |
| CC | GO:0043209 | myelin sheath | 51/1360 | 187/8928 | 1,36E-05 | 0,000391 | 0,000332 | Ass1/Dlg1/Itgb1/Itpr3/Atp1a1/Aco2/Fscn1/Prdx3/Plec/Ndufa10/Uqcrc1/Cox5b/Ckmt1/Dlst/Atp5c1/Sod2/Ndufv2/Tufm/Dld/Cox5a/Immt/Ndufs3/Uqcrfs1/Pdha1/Atp5pb/Ndufs1/Slc25a12/Prxl2b/Slc25a3/Atp5o/Uqcrc2/Idh3a/Vdac1/Hspd1/Mdh2/Sucla2/Tagln3/Sdha/Got2/Slc25a4/Gnb5/Vdac2/Pacsin1/Pten/Atp1a2/Stxbp1/Dlat/Slc25a5/Itpr2/Pdcd6ip/Sptan1 |
| CC | GO:0098552 | side of membrane | 70/1360 | 284/8928 | 1,74E-05 | 0,000489 | 0,000416 | Cd44/Vwf/Kcnj3/Gfra2/Cav2/Cdh13/Tiam1/Aqp4/Itga7/Dlg1/Itga3/Ace/Icam1/Gpc4/Pecam1/Gnal/H2-K1/Epb41/Gng4/Antxr1/Entpd1/Nlgn1/Anxa5/H2-D1/Th/Itgb1/Itga1/Fgb/Iqgap1/L1cam/Anpep/Fgg/Il6st/Htra2/Cdh5/Pkp4/Snx5/Itgav/Insr/Enpep/Il1r1/Cntfr/Itga6/Dnajc19/Abcc4/Gphn/Ptprc/Dag1/Atp1b2/Aspscr1/Fgfr3/Lyn/Gnb5/Vcam1/Capn2/Yes1/Map3k5/Alcam/Abcg1/Pten/Itgb2/Cr1l/Ppp1r9b/Cdk16/Itgam/Dtna/Rnpep/Chuk/Myh10/Gnai2 |
| CC | GO:0043083 | synaptic cleft | 12/1360 | 22/8928 | 2,17E-05 | 0,000591 | 0,000502 | Lama5/Lamb2/C1ql3/Lamc1/Agrn/Lama4/Slc1a1/Cdh8/Adgrb3/Dnm3/Lgi1/Grip1 |
| CC | GO:0009925 | basal plasma membrane | 45/1360 | 164/8928 | 3,66E-05 | 0,00096 | 0,000815 | Cd44/Kcnj4/Slc16a3/Cav1/Atp2b4/Slc4a7/Aqp4/Hcn1/Hcn4/Cxadr/Dlg1/Slc29a1/Itga3/Ace/Trpc1/Adcy8/Vangl2/Epb41/Slc4a8/Trpc4/Entpd1/Ctnna2/Rapgef3/Iqgap1/Myo1c/Atp1a1/Plec/Mlc1/Ddr1/Cdh2/Slc14a1/Oscp1/Tjp1/Itga6/Abcc4/Pdzd11/Dag1/Cask/Palm/Slc38a3/Ctnnb1/Slc23a2/Heph/Slc12a6/Cr1l |

| | | | | | | | | |
|---|---|---|---|---|---|---|---|---|
| CC | GO:0070069 | cytochrome complex | 12/1360 | 23/8928 | 3,91E-05 | 0,001007 | 0,000855 | Uqcrb/Uqcrc1/Cox5b/Cox5a/Uqcrfs1/Cox6c/Uqcrc2/Uqcrq/Cox4i1/Cyc1/Cox15/Cox7a2l |
| CC | GO:0048786 | presynaptic active zone | 32/1360 | 106/8928 | 6,71E-05 | 0,001641 | 0,001394 | Ntng1/Grm8/Cplx3/Grm4/Adra2a/Atp2b4/Hcn1/Itga3/Nrxn1/Adcy8/Ctnna2/Dgki/C1qbp/Stx1a/Cdh10/Lrfn3/Grm3/Rims2/Apba1/Lrfn1/Cacna2d1/Cdh2/Phb2/Vdac1/Ctnnb1/Gucy1b1/Grip1/Iqsec2/Stxbp1/Atp2b2/Ctnnd1/Gabrb1 |
| CC | GO:0001726 | ruffle | 37/1360 | 131/8928 | 9,13E-05 | 0,002091 | 0,001776 | Tpm1/Pdlim7/Tiam1/Pdgfrb/Actn1/Gas7/Ptprj/Pecam1/Myo6/Podxl/Pak1/Itgb1/Iqgap1/Myo1c/Map2/Psd2/Cd2ap/Tln1/Fscn1/Hip1r/Akt2/Tln2/Itgav/Jcad/Ptprz1/Dbnl/Cspg4/Arhgef2/Myo5a/Pacsin1/Twf1/Appl2/Pdxp/Wasf2/Ppp1r9b/Abca7/Lcp1 |
| CC | GO:0150034 | distal axon | 73/1360 | 314/8928 | 9,17E-05 | 0,002092 | 0,001777 | Cplx3/Grm4/Calb2/Adra2a/Pcsk1/Hcn3/Flna/Kcna3/Ptprf/Tiam1/Hcn1/Hcn4/Lamp5/Cxadr/Itga3/Slc8a3/Nrxn1/Agrn/Trpc5/Tpbg/Slc4a8/Slc18a2/Dbh/Kcnc4/Dgki/Slc1a1/Anxa5/Grik5/Th/Rapgef3/Cdh8/Grik2/Dbn1/Pak1/Iqgap1/L1cam/Ptprn2/Map2/Syt7/Sncb/Fscn1/Rac3/Slc2a13/Prph/Shtn1/Cad/Kcnk2/Ptprz1/Calb1/Neo1/Adgrl1/Kcna6/Pcdh9/Flrt3/Ilk/Tubb3/Pcdhgb1/Pacsin1/Septin6/Sri/Dclk1/Grip1/Arhgef7/Ppp1r9b/Septin5/Taok2/Ctnnd1/Fkbp4/Pard6a/Dpysl3/Hap1/Myh10/Rufy3 |
| CC | GO:0045178 | basal part of cell | 46/1360 | 176/8928 | 0,00011 | 0,002441 | 0,002073 | Cd44/Kcnj4/Slc16a3/Cav1/Atp2b4/Slc4a7/Aqp4/Hcn1/Hcn4/Cxadr/Dlg1/Slc29a1/Itga3/Ace/Trpc1/Adcy8/Vangl2/Epb41/Slc4a8/Trpc4/Entpd1/Ctnna2/Rapgef3/Itga1/Iqgap1/Myo1c/Atp1a1/Plec/Mlc1/Ddr1/Cdh2/Slc14a1/Oscp1/Tjp1/Itga6/Abcc4/Pdzd11/Dag1/Cask/Palm/Slc38a3/Ctnnb1/Slc23a2/Heph/Slc12a6/Cr1l |
| CC | GO:0031594 | neuromuscular junction | 27/1360 | 86/8928 | 0,000117 | 0,002573 | 0,002185 | Camk2d/Lama5/Lrp4/Thbs4/Des/Lamb2/Cxadr/Itga7/Dlg1/Itga3/Lamc1/Slc8a3/Nrxn1/Utrn/Nlgn1/Lama4/Kcnc4/Itgb1/Cd2ap/Snta1/Prkaca/Syngr3/Dnaja3/Pls3/Dlgap4/Crkl/Myh10 |
| CC | GO:0042383 | sarcolemma | 34/1360 | 121/8928 | 0,000193 | 0,00387 | 0,003287 | Col6a3/Col6a1/Camk2d/Kcnj3/Cav1/Atp2b4/Cav2/Aqp4/Des/Itga7/Dlg1/Slc8a3/Agrn/Nos3/Ahnak/Utrn/Akap7/Anxa5/Itgb1/Atp1a1/Plec/Vcl/Cacna2d1/Cdh2/Snta1/Igf1r/Adcy6/Dag1/Got2/Vcam1/Pld2/Sri/Atp1a2/Dtna |
| CC | GO:0060076 | excitatory synapse | 28/1360 | 93/8928 | 0,000197 | 0,003938 | 0,003345 | Slc17a6/Grm4/Calb2/Kcnj9/Slc16a3/Cntn6/Kcnj3/Kcnj6/Ptprf/Itga3/Plxnc1/Nrxn1/Adcy8/Nlgn1/Ctnna2/Dgki/Nlgn3/Dbn1/Itgb1/Adgrb3/Nlgn4l/Pde4b/Grid2/Nlgn2/Afdn/Sptbn2/Stxbp1/Atp2b2 |
| CC | GO:0043198 | dendritic shaft | 21/1360 | 62/8928 | 0,000203 | 0,004005 | 0,003402 | Grm4/Flna/Whrn/Hcn1/Kirrel3/Slc1a2/Hcn2/Nlgn1/Slc1a1/Dbn1/Prkar2b/Map2/Nlgn2/Grip2/Gipc1/Ilk/Arhgef2/Ctnnb1/Grip1/Gabbr1/Htr2a |
| CC | GO:0043679 | axon terminus | 40/1360 | 155/8928 | 0,000399 | 0,006858 | 0,005824 | Cplx3/Grm4/Calb2/Adra2a/Pcsk1/Hcn3/Kcna3/Hcn1/Hcn4/Slc8a3/Tpbg/Slc4a8/Slc18a2/Dbh/Kcnc4/Dgki/Slc1a1/Anxa5/Grik5/Th/Rapgef3/Cdh8/Grik2/L1cam/Ptprn2/Syt7/Sncb/Prph/Cad/Kcnk2/Calb1/Kcna6/Flrt3/Ilk/Pacsin1/Septin6/Sri/Grip1/Septin5/Hap1 |
| BP | GO:0009117 | nucleotide metabolic process | 138/1360 | 393/8928 | 1,34E-23 | 1,54E-20 | 1,31E-20 | Prps1l1/Slc4a7/Prps1/Prpsap2/Fmo1/Prps2/Galt/Nt5c1a/Prpsap1/Adcy8/Nos3/Suclg2/Ndufa3/Nudt13/Idh2/Fdxr/Mcee/Entpd1/Aldh1l2/Mccc2/Dtymk/Nadk2/Ndufv3/Dguok/Nme3/Mfn1/Gda/Ndufa1/Hmgcl/Gcdh/Nudt12/Fis1/Adcy1/Nqo1/Ndufa12/Ppat/Dut/Csl/Acss1/Nudt17/Tk2/Pde4a/Ndufa10/Pdhx/Bcl2l1/Acadsb/Sdhb/Insr/Cad/Galk1/Dlst/Acat1/Ndufs2/Atp5c1/Bad/Stoml2/Mmut/Ndufv2/Ndufb11/Nmnat2/Ndufa13/Fhit/Dld/Ndufb9/Ndufs3/Gpd2/Ndufa2/Nudt2/Ndufa5/Cs/Pdha1/Ndufs7/Atp5pb/Ndufv1/Ndufa6/Idh3g/Ndufs1/Slc25a12/Fdx1/Naxe/Pdhb/Acot9/Rpia/Ak2/Ndufs5/Atp5o/Prkaca/Pnp/Idh3a/Ogdh/Vdac1/Mlycd/Adcy6/Ndufc2/Ak3/Mdh2/Sucla2/Tkfc/Pdk1/Ndufb4/Ndufa9/Atic/Sdha/Cmpk1/Ndufa8/Cmpk2/Oxsm/Acaa2/Idh3b/Adcy2/Nt5m/Hsd11b1/Dmac2l/Gucy1b1/Ndufb6/Opa1/Pank4/Acot2/Ctps2/Pfkp/Atp1a2/Me2/Dlat/Slc25a25/Mtch2/Dhodh/Atp2b2/Pgd/Mpi/Pdk3/Pde2a/Hk1/Htr2a/Acsl6/Dnm1l/Oga/Flad1/Pfkm |
| BP | GO:0045333 | cellular respiration | 79/1360 | 173/8928 | 6,96E-22 | 4E-19 | 3,4E-19 | Suclg2/Ndufa3/Idh2/Lyrm7/Ndufv3/Dguok/Fxn/Slc25a23/Ndufa1/Uqcrb/Aco2/Ndufa12/Etfa/Csl/Plec/Ndufa10/Uqcrc1/Cox5b/Nipsnap2/Sdhb/Bax/Dlst/Ndufs2/Atp5c1/Stoml2/Etfb/Sod2/Ogdhl/Ndufv2/Ndufb11/Ndufa13/Dld/Cox5a/Ndufb9/Ndufs3/Uqcrfs1/Ndufa2/Ndufa5/Cs/Shmt2/Pdha1/Ndufs7/Atp5pb/Ndufv1/Ndufa6/Idh3g/Ndufs1/Slc25a12/Cox6c/Pdhb/Ndufs5/Atp5o/Uqcrc2/Prkaca/Idh3a/Ogdh/Coq9/Mrps36/Ndufc2/Pnpt1/Uqcrq/Mdh2/Sucla2/Ndufb4/Ndufa9/Sdha/Ndufa8/Cox4i1/Cyc1/Idh3b/Mtfr1l/Ndufb6/Cox7a2l/Sirt3/Dlat/Slc25a25/Mtch2/Pde2a/Trap1 |
| BP | GO:0006520 | amino acid metabolic process | 79/1360 | 181/8928 | 2,23E-20 | 6,76E-18 | 5,74E-18 | Atp2b4/Ass1/Tph2/Gcat/Gad2/Aldh1a1/Pm20d1/Tst/Prodh/Aass/Ddc/Nos3/Aldh6a1/Slc7a11/Ivd/Gcsh/Glud1/Mccc2/Wars2/Aldh4a1/Th/Mpst/Kyat3/Gls/Hmgcl/Gatc/Gstz1/Bcat2/Hmgcll1/Tha1/Dbt/Ppat/Etfa/Gldc/Acad8/Sardh/Pars2/Ddo/Vars2/Hsd17b10/Aldh5a1/Acadsb/Oat/Gatb/Cad/Cars2/Qrsl1/Iars2/Sars2/Hibch/Acat1/Nars2/Mmut/Etfb/Dld/Gls2/Mars2/Ears2/Shmt2/Cyp2d22/Slc25a12/Ddah1/Dars2/Lars2/Slc25a44/Hibadh/Got2/Aldh18a1/Bckdha/Rars2/Adhfe1/Tars2/Aarsd1/Ctps2/Yars2/Cbs/Blmh/Gclc/Gfpt1 |
| BP | GO:0016054 | organic acid catabolic process | 74/1360 | 172/8928 | 1,01E-18 | 2,53E-16 | 2,15E-16 | Slc16a3/Atp2b4/Gcat/Gad2/Prodh/Aass/Nos3/Aldh6a1/Acads/Ivd/Gcsh/Glud1/Aldh1l2/Mccc2/Hadh/Acad10/Aldh4a1/Eci1/Acadm/Acadl/Decr1/Gls/Hmgcl/Echs1/Gcdh/Gstz1/Bcat2/Crat/Hmgcll1/Tha1/Dbt/Ppat/Etfa/Gldc/Acad8/Sardh/Akt2/Ddo/Acsf3/Hsd17b10/Aldh5a1/Acadsb/Oat/Cpt1a/Hadha/Eci2/Hibch/Acat1/Etfb/Hadhb/Dld/Gls2/Etfdh/Shmt2/Cyp4f14/Ddah1/Npl/Pck2/Mlycd/Cpt2/Abhd3/Slc25a44/Hibadh/Acadvl/Got2/Bckdha/Acaa2/Adhfe1/Acot2/Cbs/Acat2/Blmh/Gnpda1/Pgd |
| BP | GO:0046390 | ribose phosphate biosynthetic process | 70/1360 | 158/8928 | 1,27E-18 | 2,93E-16 | 2,49E-16 | Prps1l1/Prps1/Prpsap2/Prps2/Prpsap1/Adcy8/Ndufa3/Entpd1/Ndufv3/Dguok/Nme3/Ndufa1/Gcdh/Adcy1/Ndufa12/Ppat/Acss1/Ndufa10/Pdhx/Bcl2l1/Sdhb/Cad/Acat1/Ndufs2/Atp5c1/Stoml2/Mmut/Ndufv2/Ndufb11/Ndufa13/Dld/Ndufb9/Ndufs3/Ndufa2/Nudt2/Ndufa5/Pdha1/Ndufs7/Atp5pb/Ndufv1/Ndufa6/Ndufs1/Slc25a12/Pdhb/Ak2/Ndufs5/Atp5o/Pnp/Vdac1/Mlycd/Adcy6/Ndufc2/Ak3/Pdk1/Ndufb4/Ndufa9/Atic/Sdha/Cmpk1/Ndufa8/Adcy2/Dmac2l/Gucy1b1/Ndufb6/Pank4/Ctps2/Dlat/Dhodh/Pdk3/Acsl6 |
| BP | GO:0044282 | small molecule catabolic process | 90/1360 | 243/8928 | 2,32E-17 | 4,94E-15 | 4,2E-15 | Slc16a3/Atp2b4/Galt/Gcat/Gad2/Aldh1a1/Prodh/Aass/Nos3/Aldh6a1/Acads/Ivd/Gcsh/Glud1/Aldh1l2/Mccc2/Hadh/Acad10/Aldh4a1/Eci1/Acadm/Acadl/Oxct1/Gda/Decr1/Gls/Hmgcl/Echs1/Gcdh/Gstz1/Bcat2/Crat/Hmgcll1/Tha1/Dbt/Ppat/Etfa/Gldc/Acad8/Sardh/Akt2/Ddo/Acsf3/Hsd17b10/Aldh5a1/Acadsb/Oat/Cpt1a/Hadha/Galk1/Eci2/Hibch/Acat1/Bad/Etfb/Hadhb/Dld/Gls2/Etfdh/Shmt2/Cyp4f14/Slc25a12/Ddah1/Npl/Pck2/Pnp/Mlycd/Cpt2/Abhd3/Slc25a44/Tkfc/Hibadh/Acadvl/Got2/Bckdha/Acaa2/Adhfe1/Hagh/Acot2/Pfkp/Pten/Pdxp/Cbs/Acat2/Blmh/Gnpda1/Pgd/Mpi/Hk1/Pfkm |
| BP | GO:0050808 | synapse organization | 134/1360 | 459/8928 | 3,76E-15 | 6,18E-13 | 5,25E-13 | Ntng1/Cbln4/Plxnd1/Il1rapl1/Lrrc4c/Tnc/Igsf9b/Reln/Lingo2/Flna/Lama5/Lrp4/Ptprf/Tiam1/Ptprt/Sorbs2/Lrrtm4/Lamb2/Dock10/C1ql3/Ptprd/Itga3/Lrtm2/Negr1/Kirrel3/Lrrc24/C1qa/Nrxn2/Slc8a3/Plxnc1/C1qb/Nrxn1/Agrn/Gpc4/Gap43/Myo6/Tpbg/Slc7a11/Camk1/Arhgap22/Nlgn1/Arhgef9/Adgrl2/Ctnna2/Lrfn4/Slc1a1/Nlgn3/Cdh8/Asap1/Dbn1/Pak1/Itgb1/Mfn1/Adgrb3/Nlgn4l/Lrfn5/Camkv/Lgi2/L1cam/Cdh10/Lrfn3/Magi2/Pdlim5/Mdga2/Srcin1/Sorbs1/Arhgap33/Cd2ap/Cdh6/Sncb/Pcdh17/Grid2/Rac3/Ctnnd2/Pcdhgc3/Adgrb2/Septin11/Nlgn2/Hip1r/Rims2/Grid1/Afdn/Insr/Lrfn1/Cdh2/Snta1/Afg3l2/Igf1r/Slc6a1/Dbnl/Adgrl1/Flrt3/Sema4d/Dnm3/Gphn/Grip2/Dag1/Synpo/Plxnb2/Ephb2/Adgrl3/Cask/Palm/Nrcam/Myo5a/Dab2ip/Efnb1/Ctnnb1/Slitrk1/Nf1/Sptbn2/Opa1/Pten/Arhgef7/Lrrc4b/Dnaja3/Wasf2/Slitrk4/Abhd17a/Itgam/Taok2/Atp2b2/Add2/Hdac6/Slitrk3/Ube2v2/Flrt2/Wasf3/Tanc2/Dnm1l/Crkl/Myh10/Iqsec3/Kalrn |
| BP | GO:1990542 | mitochondrial transmembrane transport | 39/1360 | 71/8928 | 8,92E-15 | 1,42E-12 | 1,21E-12 | Slc25a21/Slc8a3/Tst/Sfxn5/Sfxn1/Slc25a23/Abcb7/Sfxn3/Sfxn2/Aifm1/Letm1/Timm21/Afg3l2/Maip1/Slc25a12/Tomm40l/Dnajc19/Micu1/Slc25a40/Slc25a3/Timm44/Timm50/Vdac1/Hspd1/Pnpt1/Ccdc51/Mcu/Slc25a15/Tomm40/Abcb8/Micu3/Slc25a4/Slc25a1/Timm17b/Mrpl18/Opa1/Slc25a51/Slc25a5/Mrs2 |
| BP | GO:0006119 | oxidative phosphorylation | 49/1360 | 106/8928 | 2,68E-14 | 4,16E-12 | 3,53E-12 | Ndufa3/Ndufv3/Dguok/Fxn/Slc25a23/Ndufa1/Uqcrb/Ndufa12/Ndufa10/Uqcrc1/Cox5b/Nipsnap2/Sdhb/Ndufs2/Atp5c1/Stoml2/Ndufv2/Ndufb11/Ndufa13/Dld/Cox5a/Ndufb9/Ndufs3/Uqcrfs1/Ndufa2/Ndufa5/Shmt2/Ndufs7/Atp5pb/Ndufv1/Ndufa6/Ndufs1/Cox6c/Ndufs5/Atp5o/Uqcrc2/Coq9/Ndufc2/Uqcrq/Ndufb4/Ndufa9/Sdha/Ndufa8/Cox4i1/Cyc1/Ndufb6/Cox7a2l/Mtch2/Pde2a |

| | | | | | | | | |
|---|---|---|---|---|---|---|---|---|
| BP | GO:0034329 | cell junction assembly | 104/1360 | 331/8928 | 3,24E-14 | 4,9E-12 | 4,17E-12 | Cbln4/Plxnd1/Il1rapl1/Reln/Lingo2/Cav1/Arhgap6/Lrp4/Cdh13/Lrrtm4/C1ql3/Ptprd/Dlg1/Lrtm2/Actn1/Negr1/Kirrel3/Lamc1/Lrrc24/Ace/Nrxn2/Gja1/Ptprj/Nrxn1/Agrn/Gpc4/Ptprk/Pecam1/Gap43/Myo6/Tpbg/Esam/Gjb6/Ace2/Nlgn1/Arhgef9/Adgrl2/Cdh12/Tns1/Lrfn4/Nlgn3/Cdh8/Itgb1/Adgrb3/Nlgn4l/Iqgap1/Lrfn5/Lgi2/Myo1c/Cdh10/Lrfn3/Pxn/Magi2/Pdlim5/Cdh20/Mdga2/Srcin1/Sorbs1/Cdh6/Tln1/Pcdh17/Fscn1/Grid2/Cdh5/Plec/Ctnnd2/Pkp4/Adgrb2/Nlgn2/Actn4/Vcl/Afdn/Itgav/Lrfn1/Cdh2/Dbnl/Adgrl1/Flrt3/Tjp1/Sema4d/Dnm3/Coro2b/Prkaca/Plxnb2/Ephb2/Adgrl3/Efnb1/Ctnnb1/Slitrk1/Sptbn2/Pten/Lrrc4b/Slitrk4/Taok2/Add2/Ctnnd1/Slitrk3/Pdcd6ip/Ube2v2/Flrt2/Coro1c/Pak2/Crkl/Dusp3 |
| BP | GO:0007005 | mitochondrion organization | 122/1360 | 415/8928 | 4,13E-14 | 6,08E-12 | 5,17E-12 | Cav2/Cxadr/Ndufa3/Tomm22/Lyrm7/Tspan9/Them4/Mfn1/Nectin2/Fxn/Ndufa1/Hmgcl/Bcs1l/Lonp1/Fis1/Htra2/Ndufa12/Ssbp1/Micos13/Prdx3/Plec/Atp23/Hip1r/Tmem223/Tk2/Aifm1/Letm1/Acad9/Ndufa10/Hsd17b10/Bcl2l1/Nipsnap2/Cox20/Apoo/Timm21/Bax/Timm22/Ndufs2/Phb2/Bad/Stoml2/Nubpl/Afg3l2/Samm50/Sod2/Mtx2/Spg7/Ndufb11/Maip1/Hspa4/Ndufa13/Ndufb9/Immt/Ndufs3/Ndufa2/Ndufa5/Ndufs7/Agk/Ndufa6/Ndufs1/Tomm40l/Timm29/Dnajc19/Apool/Gpx1/Mtx1/Ndufs5/Timm44/Dnajc11/Rimoc1/Pnp/Timm50/Ndufaf5/Supv3l1/Pmpca/Vdac1/Ndufc2/Hspd1/Pnpt1/Mcu/Tfam/Taco1/Bag3/Tomm40/Ndufb4/Ndufa9/Chchd3/Ndufa8/Rhot1/Slc25a4/Mapk9/Acaa2/Fbxl4/Atg13/Cert1/Ndufaf3/Timm17b/Vdac2/Surf1/Mtfr1l/Ndufb6/Pmpcb/Ndufaf7/Opa1/Cox7a2l/Pgam5/Adck1/Cyrib/Tmem126a/Dnaja3/Slc25a5/Mtch2/Dhodh/Armc1/Pde2a/Gclc/Wdr45/Hdac6/Rap1gds1/Hap1/Dnm1l/Atg2b |
| BP | GO:0043648 | dicarboxylic acid metabolic process | 39/1360 | 74/8928 | 5,53E-14 | 7,75E-12 | 6,58E-12 | Ass1/Gad2/Prodh/Slc7a11/Suclg2/Glud1/Aldh1l2/Aldh4a1/Kyat3/Gls/Csl/Ddo/Acsf3/Aldh5a1/Sdhb/Oat/Dlst/Ogdhl/Dld/Gls2/Cs/Shmt2/Mthfd1l/Idh3g/Slc25a12/Pck2/Idh3a/Ogdh/Mrps36/Mdh2/Sucla2/Atic/Sdha/Got2/Aldh18a1/Idh3b/Adhfe1/Me2/Gclc |
| BP | GO:0098609 | cell-cell adhesion | 128/1360 | 462/8928 | 1,13E-12 | 1,27E-10 | 1,08E-10 | Ntng1/Cd44/Il1rapl1/Lrrc4c/Igsf9b/Cntn6/Cav1/Tenm1/Ass1/Lama5/Ptprf/Cdh13/Aqp4/Magi1/Ptprt/Thbs4/Ptprd/Cxadr/Itga7/Dlg1/Itga3/Lgals1/Negr1/Kirrel3/Icam2/Icam1/Nrxn1/Gpc4/Pecam1/Hspb1/Cd93/Plekha7/Esam/Igdcc4/Slc7a11/Ptprm/Lpp/Nlgn1/Ctnna2/Cdh12/Tspan9/Lrfn4/Podxl/Nlgn3/Swap70/Cd200/Cdh8/Itgb1/Nectin2/Itga1/Fgb/Pip5k1c/Nlgn4l/Lrfn5/Robo2/L1cam/Cdh10/Lrfn3/Fgg/Tenm3/Astn2/Il6st/Magi2/Cdh20/Mdga2/Cd2ap/Cdh6/Tln1/Pcdh17/Grid2/Cdh5/Ctnnd2/Pcdhgc3/Pkp4/Rap1gap/Nlgn2/Vcl/Tln2/Afdn/Itgav/Ptpn6/Itgb5/Ttyh1/Lgals9/Cdh2/Bad/Casp3/Neo1/Cadm2/Adgrl1/Flrt3/Tjp1/Sema4d/Efr3a/Itga6/Jam2/Ptprc/Pnp/Cdon/Hspd1/Atp1b2/Plxnb2/Fat3/Adgrl3/Nck1/Nrcam/Lyn/Coro1a/Efnb1/Ctnnb1/Vcam1/Specc1l/Tenm2/Slitrk1/Alcam/Pcdh10/Pten/Cyrib/Itgb2/Stxbp1/Lrrc4b/Dnaja3/Itgam/Taok2/Add2/Ctnnd1/Slitrk3/Dusp3 |
| BP | GO:0010257 | NADH dehydrogenase complex assembly | 29/1360 | 52/8928 | 1,51E-11 | 1,47E-09 | 1,25E-09 | Ndufa3/Ndufa1/Bcs1l/Ndufa12/Aifm1/Acad9/Ndufa10/Timm21/Ndufs2/Nubpl/Ndufb11/Ndufa13/Ndufb9/Ndufs3/Ndufa2/Ndufa5/Ndufs7/Ndufa6/Ndufs1/Ndufs5/Ndufaf5/Ndufc2/Ndufb4/Ndufa9/Ndufa8/Ndufaf3/Ndufb6/Ndufaf7/Tmem126a |
| BP | GO:0032981 | mitochondrial respiratory chain complex I assembly | 29/1360 | 52/8928 | 1,51E-11 | 1,47E-09 | 1,25E-09 | Ndufa3/Ndufa1/Bcs1l/Ndufa12/Aifm1/Acad9/Ndufa10/Timm21/Ndufs2/Nubpl/Ndufb11/Ndufa13/Ndufb9/Ndufs3/Ndufa2/Ndufa5/Ndufs7/Ndufa6/Ndufs1/Ndufs5/Ndufaf5/Ndufc2/Ndufb4/Ndufa9/Ndufa8/Ndufaf3/Ndufb6/Ndufaf7/Tmem126a |
| BP | GO:0099560 | synaptic membrane adhesion | 22/1360 | 34/8928 | 7,76E-11 | 7,08E-09 | 6,01E-09 | Ntng1/Lrrc4c/Igsf9b/Ptprf/Ptprd/Itga3/Nrxn1/Gpc4/Nlgn1/Lrfn4/Lrfn5/Cdh10/Lrfn3/Magi2/Mdga2/Cdh6/Pcdh17/Flrt3/Slitrk1/Lrrc4b/Taok2/Slitrk3 |
| BP | GO:0007416 | synapse assembly | 60/1360 | 176/8928 | 2,85E-10 | 2,34E-08 | 1,99E-08 | Cbln4/Plxnd1/Il1rapl1/Reln/Lingo2/Lrp4/Lrrtm4/C1ql3/Ptprd/Lrtm2/Negr1/Kirrel3/Lrrc24/Nrxn2/Nrxn1/Agrn/Gpc4/Gap43/Myo6/Tpbg/Nlgn1/Arhgef9/Adgrl2/Lrfn4/Nlgn3/Adgrb3/Nlgn4l/Lrfn5/Lgi2/Lrfn3/Magi2/Pdlim5/Mdga2/Srcin1/Pcdh17/Grid2/Adgrb2/Nlgn2/Lrfn1/Cdh2/Dbnl/Adgrl1/Flrt3/Sema4d/Dnm3/Plxnb2/Ephb2/Adgrl3/Efnb1/Ctnnb1/Slitrk1/Sptbn2/Pten/Lrrc4b/Slitrk4/Add2/Slitrk3/Ube2v2/Flrt2/Crkl |
| BP | GO:0050807 | regulation of synapse organization | 75/1360 | 245/8928 | 5,42E-10 | 4,15E-08 | 3,53E-08 | Il1rapl1/Reln/Lingo2/Lrp4/Ptprf/Tiam1/Ptprt/Lrrtm4/C1ql3/Ptprd/Lrtm2/Negr1/Lrrc24/Plxnc1/Nrxn1/Agrn/Gpc4/Tpbg/Slc7a11/Camk1/Arhgap22/Nlgn1/Adgrl2/Ctnna2/Lrfn4/Nlgn3/Cdh8/Asap1/Dbn1/Itgb1/Mfn1/Adgrb3/Lrfn5/Camkv/Lrfn3/Magi2/Pdlim5/Mdga2/Srcin1/Arhgap33/Grid2/Adgrb2/Septin11/Nlgn2/Grid1/Afdn/Lrfn1/Cdh2/Dbnl/Adgrl1/Flrt3/Sema4d/Dnm3/Dag1/Ephb2/Adgrl3/Cask/Nrcam/Dab2ip/Ctnnb1/Slitrk1/Nf1/Opa1/Pten/Lrrc4b/Slitrk4/Abhd17a/Taok2/Slitrk3/Ube2v2/Flrt2/Tanc2/Dnm1l/Myh10/Kalrn |
| BP | GO:0050803 | regulation of synapse structure or activity | 76/1360 | 251/8928 | 7,27E-10 | 5,5E-08 | 4,67E-08 | Slc17a6/Il1rapl1/Reln/Lingo2/Lrp4/Ptprf/Tiam1/Ptprt/Lrrtm4/C1ql3/Ptprd/Lrtm2/Negr1/Lrrc24/Plxnc1/Nrxn1/Agrn/Gpc4/Tpbg/Slc7a11/Camk1/Arhgap22/Nlgn1/Adgrl2/Ctnna2/Lrfn4/Nlgn3/Cdh8/Asap1/Dbn1/Itgb1/Mfn1/Adgrb3/Lrfn5/Camkv/Lrfn3/Magi2/Pdlim5/Mdga2/Srcin1/Arhgap33/Grid2/Adgrb2/Septin11/Nlgn2/Grid1/Afdn/Lrfn1/Cdh2/Dbnl/Adgrl1/Flrt3/Sema4d/Dnm3/Dag1/Ephb2/Adgrl3/Cask/Nrcam/Dab2ip/Ctnnb1/Slitrk1/Nf1/Opa1/Pten/Lrrc4b/Slitrk4/Abhd17a/Taok2/Slitrk3/Ube2v2/Flrt2/Tanc2/Dnm1l/Myh10/Kalrn |
| BP | GO:0046034 | ATP metabolic process | 54/1360 | 160/8928 | 3,37E-09 | 2,31E-07 | 1,96E-07 | Galt/Ndufa3/Entpd1/Ndufv3/Dguok/Ndufa1/Fis1/Ndufa12/Ndufa10/Bcl2l1/Sdhb/Insr/Galk1/Ndufs2/Atp5c1/Bad/Stoml2/Ndufv2/Ndufb11/Ndufa13/Ndufb9/Ndufs3/Ndufa2/Nudt2/Ndufa5/Ndufs7/Atp5pb/Ndufv1/Ndufa6/Ndufs1/Slc25a12/Ak2/Ndufs5/Atp5o/Prkaca/Ogdh/Ndufc2/Ak3/Tkfc/Ndufb4/Ndufa9/Sdha/Ndufa8/Dmac2l/Ndufb6/Pfkp/Atp1a2/Slc25a25/Mtch2/Mpi/Hk1/Htr2a/Dnm1l/Pfkm |
| BP | GO:0009081 | branched-chain amino acid metabolic process | 16/1360 | 23/8928 | 6,56E-09 | 4,23E-07 | 3,6E-07 | Aldh6a1/Ivd/Mccc2/Hmgcl/Bcat2/Hmgcll1/Dbt/Acad8/Hsd17b10/Acadsb/Hibch/Acat1/Dld/Slc25a44/Hibadh/Bckdha |
| BP | GO:0051962 | positive regulation of nervous system development | 70/1360 | 247/8928 | 6,89E-08 | 3,96E-06 | 3,37E-06 | Plxnd1/Il1rapl1/Reln/Tiam2/Lingo2/Mme/Tgm2/Ptprf/Tiam1/Lrrtm4/Ptprd/Lrtm2/Lrrc24/Ace/Plxnc1/Nrxn1/Agrn/Trpc5/Tpbg/Plxna3/Nlgn1/Vim/Adgrl2/Nlgn3/Dbn1/Pak1/Itgb1/Mfn1/Fxn/Adgrb3/Nr2e1/Robo2/L1cam/Il6st/Grid2/Adgrb2/Nlgn2/Shtn1/Sox8/Afdn/Ptprz1/Dbnl/Adgrl1/Flrt3/Sema4d/Cdon/Dag1/Ilk/Plxnb2/Ephb2/Adgrl3/Cask/Mapk8/Lyn/Nap1l1/Ctnnb1/Slitrk1/Tnik/Opa1/Grip1/Lrrc4b/Slitrk4/Slitrk3/Ube2v2/Tbc1d24/Flrt2/Hap1/Dnm1l/Rufy3/Kalrn |
| BP | GO:0006635 | fatty acid beta-oxidation | 27/1360 | 62/8928 | 8,35E-08 | 4,7E-06 | 3,99E-06 | Acads/Ivd/Aldh1l2/Hadh/Acad10/Eci1/Acadm/Acadl/Decr1/Echs1/Gcdh/Crat/Etfa/Akt2/Hsd17b10/Cpt1a/Hadha/Eci2/Acat1/Etfb/Hadhb/Etfdh/Mlycd/Cpt2/Acadvl/Acaa2/Acat2 |
| BP | GO:0031589 | cell-substrate adhesion | 64/1360 | 223/8928 | 1,52E-07 | 8,04E-06 | 6,83E-06 | Vwf/Fermt1/Flna/Lama5/Arhgap6/Cdh13/Tiam1/Lamb2/Itga7/Itga3/Actn1/Lgals1/Lamc1/Bcam/Nid1/Spry4/Ptprj/Cspg5/Ptprk/Pecam1/Angpt1/Antxr1/Utrn/C1qbp/Dbn1/Itgb1/Itga1/Itgb8/Fgb/Iqgap1/L1cam/Fgg/Pxn/Srcin1/Sorbs1/Rac3/Actn4/Vcl/Itgav/Itgb5/Ddr1/Ttyh1/Ptprz1/Carmil1/Tesk1/Itga6/Gfus/Coro2b/Dag1/Atp1b2/Rell2/Ilk/Cask/Coro1a/Ctnnb1/Vcam1/Nf1/Pten/Itgb2/Itgam/Taok2/Coro1c/Crkl/Dusp3 |
| BP | GO:1901137 | carbohydrate derivative biosynthetic process | 92/1360 | 364/8928 | 2,49E-07 | 1,26E-05 | 1,07E-05 | Prps1l1/Prps1/Prpsap2/Prps2/Pdgfrb/Prpsap1/Adcy8/Vangl2/Angpt1/Necab1/Ndufa3/Entpd1/Dtymk/Ndufv3/Dguok/Nme3/Ndufa1/Gcdh/Adcy1/Ndufa12/Ppat/Dut/Pgm2/Acss1/Ndufa10/Pdhx/Bcl2l1/Sdhb/Insr/Cad/Acat1/Ndufs2/Atp5c1/Stoml2/Mmut/Poglut3/Ndufv2/Ndufb11/Ndufa13/Dld/Ndufb9/Ndufs3/Ndufa2/Nudt2/Ndufa5/Pdha1/Ndufs7/Atp5pb/Ndufv1/Ndufa6/Ndufs1/Slc25a12/Pdhb/Gfus/Ak2/Ndufs5/Atp5o/Pnp/Vdac1/Mlycd/Adcy6/Ndufc2/Ak3/Pdk1/Ndufb4/Ndufa9/Atic/Sdha/Cmpk1/Ndufa8/Cmpk2/Gk/Adcy2/Ctnnb1/Dmac2l/Gucy1b1/Gykl1/Ndufb6/Pank4/Ctps2/Dlat/Uap1l1/Dhodh/Abca7/Gnpda1/Mpi/Pdk3/Nanp/Acsl6/Oga/Gfpt1/Gmppb |
| BP | GO:0006418 | tRNA aminoacylation for protein translation | 15/1360 | 25/8928 | 3,69E-07 | 1,68E-05 | 1,43E-05 | Wars2/Pars2/Vars2/Cars2/Iars2/Sars2/Nars2/Mars2/Ears2/Dars2/Lars2/Rars2/Tars2/Aarsd1/Yars2 |
| BP | GO:0043113 | receptor clustering | 24/1360 | 55/8928 | 4,17E-07 | 1,86E-05 | 1,58E-05 | Reln/Flna/Lrp4/Sorbs2/Lrrtm4/Nrxn2/Nrxn1/Agrn/Slc7a11/Nlgn1/Arhgef9/Grik5/Grik2/Pak1/Magi2/Sorbs1/Nlgn2/Cdh2/Gphn/Grip2/Itgb2/Dnaja3/Slitrk3/Crkl |
| BP | GO:0035383 | thioester metabolic process | 26/1360 | 63/8928 | 5,25E-07 | 2,27E-05 | 1,93E-05 | Suclg2/Mcee/Hmgcl/Gcdh/Csl/Acss1/Pdhx/Acadsb/Acat1/Mmut/Dld/Cs/Pdha1/Pdhb/Acot9/Ogdh/Vdac1/Mlycd/Sucla2/Pdk1/Oxsm/Acaa2/Acot2/Dlat/Pdk3/Acsl6 |

| | | | | | | | | |
|---|---|---|---|---|---|---|---|---|
| BP | GO:0010975 | regulation of neuron projection development | 100/1360 | 420/8928 | 1,53E-06 | 5,7E-05 | 4,84E-05 | Ntng1/Plxnd1/Il1rapl1/Lrrc4c/Reln/Tiam2/Pcp4/Rap1gap2/Csmd3/Flna/Lrp4/Ptprf/Tiam1/Ptprd/Itga3/Lgals1/Negr1/Plxnc1/Nrxn1/Agrn/Trpc5/Hspb1/H2-K1/Camk1/Plxna3/Sema6b/Nlgn1/Vim/Ctnna2/Dguok/Nlgn3/H2-D1/Camk2g/Vldlr/Dbn1/Pak1/Itgb1/Mfn1/Fxn/Adgrb3/Iqgap1/Nr2e1/Robo2/L1cam/Tenm3/Map2/Magi2/Pdlim5/Srcin1/Arhgap33/Grid2/Hecw2/Shtn1/Afdn/Ddr1/Mark1/Cdh2/Igf1r/Dip2b/Ptprz1/Neo1/Kidins220/Dbnl/Sema4d/Cspg4/Itga6/Dnm3/Dpysl5/Rtn4ip1/Adcy6/Atp1b2/Ilk/Plxnb2/Ephb2/Fat3/Cask/Nck1/Nrcam/Lyn/Dab2ip/Slitrk1/Nf1/Tnik/Pacsin1/Opa1/Twf1/Grip1/Pten/Ppp2r5d/Ube2v2/Tbc1d24/Acsl6/Dpysl3/Hap1/Tanc2/Pak2/Dnm1l/Crkl/Rufy3/Kalrn |
| BP | GO:0045216 | cell-cell junction organization | 40/1360 | 125/8928 | 1,73E-06 | 6,38E-05 | 5,42E-05 | Cav1/Flna/Cdh13/Whrn/Cxadr/Dlg1/Ace/Gja1/Pecam1/Plekha7/Esam/Gjb6/Ace2/Cdh12/Cdh8/Itgb1/Myo1c/Cdh10/Cdh20/Cdh6/Fscn1/Cdh5/Plec/Ctnnd2/Pkp4/Actn4/Vcl/Afdn/Cdh2/Tjp1/Prkaca/Ephb2/Ajm1/Ctnnb1/Specc1l/Csk/Ctnnd1/Pard6a/Pdcd6ip/Pak2 |
| BP | GO:0006839 | mitochondrial transport | 39/1360 | 121/8928 | 1,9E-06 | 6,92E-05 | 5,88E-05 | Slc25a24/Tomm22/Them4/Bcs1l/Fis1/Hip1r/Aifm1/Bcl2l1/Timm21/Bax/Timm22/Bad/Samm50/Mtx2/Spg7/Maip1/Hspa4/Ndufa13/Slc25a10/Agk/Tomm40l/Timm29/Dnajc19/Timm44/Timm50/Pmpca/Hspd1/Bag3/Tomm40/Rhot1/Slc25a4/Acaa2/Timm17b/Vdac2/Pmpcb/Slc25a5/Mtch2/Slc25a20/Gclc |
| BP | GO:0050731 | positive regulation of peptidyl-tyrosine phosphorylation | 31/1360 | 89/8928 | 3,51E-06 | 0,000117 | 9,9E-05 | Cd44/Reln/Adra2a/Ehd4/Cav2/Lrp4/Thbs4/Ace/Icam1/Ptprj/Agrn/Pecam1/Angpt1/Tspan9/Itgb1/Iqgap1/Il6st/Srcin1/Ptprz1/Sema4d/Cspg4/Ptprc/Arhgef2/Fgfr3/Lyn/Abi3/Yes1/Ctnnd1/Htr2a/Pak2/Hax1 |
| BP | GO:0033627 | cell adhesion mediated by integrin | 19/1360 | 43/8928 | 5,35E-06 | 0,000168 | 0,000143 | Fermt1/Itga7/Itga3/Icam1/Podxl/Swap70/Itgb1/Itga1/Itgb8/L1cam/Rac3/Itgav/Ptpn6/Itgb5/Itga6/Lyn/Itgb2/Itgam/Crkl |
| BP | GO:0035384 | thioester biosynthetic process | 14/1360 | 26/8928 | 5,38E-06 | 0,000168 | 0,000143 | Gcdh/Acss1/Pdhx/Acat1/Mmut/Dld/Pdha1/Pdhb/Vdac1/Mlycd/Pdk1/Dlat/Pdk3/Acsl6 |
| BP | GO:1901888 | regulation of cell junction assembly | 51/1360 | 182/8928 | 5,92E-06 | 0,000183 | 0,000155 | Il1rapl1/Lingo2/Cav1/Arhgap6/Lrrtm4/Ptprd/Lrtm2/Negr1/Lrrc24/Ace/Gja1/Ptprj/Nrxn1/Agrn/Gpc4/Tpbg/Ace2/Nlgn1/Adgrl2/Lrfn4/Nlgn3/Adgrb3/Iqgap1/Lrfn5/Myo1c/Lrfn3/Pdlim5/Mdga2/Srcin1/Grid2/Adgrb2/Nlgn2/Vcl/Lrfn1/Adgrl1/Flrt3/Tjp1/Sema4d/Prkaca/Ephb2/Adgrl3/Ctnnb1/Slitrk1/Pten/Lrrc4b/Slitrk4/Slitrk3/Ube2v2/Flrt2/Coro1c/Dusp3 |
| BP | GO:0006734 | NADH metabolic process | 10/1360 | 15/8928 | 9,39E-06 | 0,000282 | 0,00024 | Nudt13/Nudt12/Nudt17/Dlst/Gpd2/Idh3g/Idh3a/Ogdh/Mdh2/Idh3b |
| BP | GO:0006851 | mitochondrial calcium ion transmembrane transport | 10/1360 | 15/8928 | 9,39E-06 | 0,000282 | 0,00024 | Slc8a3/Slc25a23/Letm1/Afg3l2/Maip1/Micu1/Vdac1/Mcu/Micu3/Opa1 |
| BP | GO:2000807 | regulation of synaptic vesicle clustering | 8/1360 | 10/8928 | 9,6E-06 | 0,000284 | 0,000242 | Nlgn1/Nlgn3/Magi2/Pcdh17/Nlgn2/Bcl2l1/Cdh2/Pten |
| BP | GO:0140053 | mitochondrial gene expression | 42/1360 | 145/8928 | 1,62E-05 | 0,000457 | 0,000389 | Wars2/C1qbp/Gatc/Hsd17b10/Gatb/Qrsl1/Iars2/Sars2/Tufm/Mtif2/Ears2/Tsfm/Shmt2/Trmt10c/Mto1/Dars2/Lars2/Mtrf1l/Supv3l1/Pnpt1/Tfam/Taco1/Rmnd1/Rars2/Fastkd5/Rpusd3/Mrpl53/Mrpl24/Mrpl18/Sirt3/Gfm2/Mrpl12/Yars2/Mrps22/Mrpl23/Mrpl9/Mrps34/Mrpl14/Mrpl13/Tbrg4/Lrpprc/Mrpl58 |
| BP | GO:0051560 | mitochondrial calcium ion homeostasis | 13/1360 | 25/8928 | 1,94E-05 | 0,000535 | 0,000454 | Tgm2/Slc8a3/Slc25a27/Slc25a23/Fis1/Letm1/Afg3l2/Maip1/Immt/Micu1/Mcu/Micu3/Rap1gds1 |
| BP | GO:0090151 | establishment of protein localization to mitochondrial membrane | 12/1360 | 23/8928 | 3,91E-05 | 0,001007 | 0,000855 | Tomm22/Bcs1l/Bax/Timm22/Samm50/Maip1/Hspa4/Ndufa13/Agk/Timm29/Tomm40/Mtch2 |
| BP | GO:0007015 | actin filament organization | 77/1360 | 332/8928 | 6,45E-05 | 0,001591 | 0,001351 | Kank4/Tpm1/Tpm2/Flna/Tenm1/Arhgap6/Sorbs2/Dlg1/Actn1/Cald1/Gas7/Tmsb10/Gja1/Kank2/Icam1/Pecam1/Myo6/Esam/Sh3kbp1/Stmn1/Lmod1/Ctnna2/Swap70/Rapgef3/Dbn1/Pak1/Myo1c/Arhgap25/Pxn/Sorbs1/Cd2ap/Tmod1/Fscn1/Rac3/Arhgef10l/Plec/Actn4/Hip1r/Shtn1/Itgb5/Cgnl1/Carmil1/Dbnl/Tesk1/Evl/Tjp1/Add3/Coro2b/Synpo/Mcu/Arhgef2/Nck1/Myo5a/Myo1b/Coro1a/Specc1l/Pacsin1/Twf1/Msrb2/Coro7/Cyrib/Cfl2/Pdxp/Wasf2/Pls3/Ppp1r9b/Add2/Lcp1/Wasf3/Coro1c/Dpysl3/Pak2/Myh10/Rufy3/Sptbn1/Hax1/Sptan1 |
| BP | GO:0018958 | phenol-containing compound metabolic process | 21/1360 | 58/8928 | 6,78E-05 | 0,00165 | 0,001401 | Reln/Kcnj6/Tph2/Pde1b/Trpc1/Ddc/Slc7a11/Dbh/Slc1a1/Cpq/Gch1/Th/Maob/Sncb/Maoa/Cyp2d22/Aldh2/Gipc1/Myo5a/Itgam/Atp2b2 |
| BP | GO:1905475 | regulation of protein localization to membrane | 44/1360 | 163/8928 | 6,8E-05 | 0,00165 | 0,001401 | Dpp10/Camk2d/Atp2b4/Lrp4/Sorbs2/Dlg1/Itga3/Nrxn1/Gpc4/Slc7a11/Slc1a1/Camk2g/Gpc5/Dbn1/Pak1/Itgb1/Gpc6/Myo1c/Magi2/Fis1/Sorbs1/Nlgn2/Akt2/Bcl2l1/Neto2/Cdh2/Lypla1/Grip2/Dag1/Ephb2/Myo5a/Abi3/Wnk3/Tnik/Lyplal1/Grip1/Iqsec2/Ppp1r9b/Csk/Itgam/Ppp2r5a/Crkl/Sptbn1/Kalrn |
| BP | GO:0030155 | regulation of cell adhesion | 94/1360 | 425/8928 | 7,26E-05 | 0,001752 | 0,001488 | Cd44/Plxnd1/Tnc/Tpm1/Lama1/Cav1/Fermt1/Flna/Ass1/Hspg2/Tgm2/Lama5/Arhgap6/Cdh13/Magi1/Lamb2/Dlg1/Itga3/Lgals1/Lamc1/Plxnc1/Nid1/Icam1/Spry4/Ptprj/Cspg5/Hspb1/Angpt1/Plxna3/Utrn/Lama4/Podxl/C1qbp/Swap70/Dbn1/Itgb1/Fgb/Iqgap1/L1cam/Lrfn3/Fgg/Tenm3/Il6st/Magi2/Mdga2/Srcin1/Rac3/Rap1gap/Actn4/Adgrg1/Vcl/Afdn/Itgav/Ptpn6/Ddr1/Lgals9/Bad/Casp3/Ptprz1/Spg7/Carmil1/Tesk1/Tjp1/Sema4d/Itga6/Jam2/Gfus/Ptprc/Coro2b/Pnp/Dag1/Hspd1/Rell2/Ilk/Plxnb2/Ephb2/Cask/Nck1/Lyn/Coro1a/Efnb1/Vcam1/Specc1l/Pld2/Nf1/Pten/Cyrib/Itgb2/Dnaja3/Taok2/Coro1c/Faf1/Crkl/Dusp3 |
| BP | GO:0034220 | monoatomic ion transmembrane transport | 107/1360 | 498/8928 | 8,13E-05 | 0,001924 | 0,001634 | Kcnj4/Dpp10/Kcnh7/Kcnh3/Reln/Tmem163/Camk2d/Kcnj9/Kcnj3/Cav1/Atp2b4/Kcnj6/Hcn3/Flna/Kcna3/Hcn1/Hcn4/Dlg1/Slc6a4/Slc8a3/Pm20d1/Trpc1/Slc5a6/Agrn/Trpc5/Gabrg1/Slc4a8/Ahnak/Slc1a2/Fxyd6/Trpc4/Hcn2/Akap7/Slc39a6/Kcnc4/Slc1a1/Nlgn3/Grik5/Rapgef3/Itgb1/Itpr3/Slc25a23/Pde4b/Atp2a3/Gabra4/Slc39a10/Atp1a1/Abcb7/Hecw2/Abcb1a/Letm1/Nipsnap2/Itgav/Gabrd/Ptpn6/Bax/Neto2/Slc12a4/Cacna2d1/Cacna2d3/Nherf2/Sestd1/Snta1/Kcnk2/Phb2/Slc6a2/Afg3l2/Slc6a1/Maip1/Ndufs7/Kcna6/Micu1/Ptprc/Asph/Vdac1/Ccdc51/Mcu/Taco1/Abcb8/Atp1b2/Ephb2/Micu3/Slc25a4/Myo5a/Lyn/Coro1a/Gnb5/Cox15/Vdac2/Wnk3/Opa1/Slc23a2/Slc12a6/Sri/Grip1/Pten/Atp1a2/Mrs2/Slc25a25/Itpr2/Atp2b2/Hk1/Htr2a/Vdac3/Hap1/Oga/Gabrb1 |
| BP | GO:0006739 | NADP metabolic process | 16/1360 | 39/8928 | 8,9E-05 | 0,002078 | 0,001765 | Prps1/Fmo1/Prps2/Nudt13/Idh2/Fdxr/Aldh1l2/Nadk2/Nudt12/Nqo1/Nudt17/Fdx1/Rpia/Hsd11b1/Me2/Pgd |
| BP | GO:0070585 | protein localization to mitochondrion | 27/1360 | 85/8928 | 9,39E-05 | 0,002132 | 0,001811 | Tomm22/Bcs1l/Fis1/Mavs/Aifm1/Timm21/Bax/Timm22/Samm50/Maip1/Hspa4/Ndufa13/Agk/Tomm40l/Timm29/Dnajc19/Timm44/Timm50/Pmpca/Hspd1/Bag3/Tomm40/Timm17b/Pmpcb/Mtch2/Hk1/Dnm1l |
| BP | GO:1902414 | protein localization to cell junction | 34/1360 | 119/8928 | 0,000136 | 0,002943 | 0,0025 | Reln/Flna/C1ql3/Dlg1/Nrxn2/Nrxn1/Gpc4/Pecam1/Hspb1/Nlgn1/Nlgn3/Gpc6/Magi2/Cdh5/Nlgn2/Actn4/Vcl/Afdn/Neto2/Cgnl1/Tjp1/Gphn/Grip2/Dag1/Mapk8/Adam22/Mapk9/Tnik/Lgi1/Grip1/Iqsec2/Slitrk3/Pak2/Kalrn |
| BP | GO:0097119 | postsynaptic density protein 95 clustering | 7/1360 | 10/8928 | 0,000147 | 0,00313 | 0,002658 | Reln/Nrxn2/Nrxn1/Nlgn1/Dbn1/Nlgn2/Cdh2 |
| BP | GO:0003018 | vascular process in circulatory system | 36/1360 | 129/8928 | 0,000148 | 0,00313 | 0,002658 | Cav1/Adra2a/Slc6a4/Ace/Gja1/Icam1/Slc5a6/Ptprj/Plekha7/Nos3/Angpt1/Ptprm/Dbh/Gch1/Itga1/Fgb/Fgg/Cdh5/Plec/Abcb1a/Snta1/Sod2/Tjp1/Add3/Ddah1/Gpx1/Grip2/Adcy6/Mgll/Atp1a2/Cbs/Pde2a/Gclc/Htr2a/Rap1gds1/Abr |

| | | | | | | | | |
|---|---|---|---|---|---|---|---|---|
| BP | GO:0010770 | positive regulation of cell morphogenesis involved in differentiation | 19/1360 | 53/8928 | 0,000174 | 0,003613 | 0,003068 | Il1rapl1/Reln/Ptprf/Tiam1/Ptprd/Dbn1/Mfn1/Afdn/Dbnl/Ilk/Ephb2/Cask/Mapk9/Tnik/Opa1/Grip1/Tbc1d24/Dnm1l/Kalrn |
| BP | GO:0043605 | amide catabolic process | 8/1360 | 13/8928 | 0,000178 | 0,003645 | 0,003096 | Nt5c1a/Pm20d1/Prodh/Aldh4a1/Gda/Nit1/Oat/Pnp |
| BP | GO:0019674 | NAD metabolic process | 12/1360 | 26/8928 | 0,000179 | 0,003645 | 0,003096 | Nudt13/Nadk2/Nudt12/Nudt17/Dlst/Nmnat2/Gpd2/Idh3g/Idh3a/Ogdh/Mdh2/Idh3b |
| BP | GO:0000959 | mitochondrial RNA metabolic process | 16/1360 | 41/8928 | 0,00018 | 0,003653 | 0,003102 | Wars2/Hsd17b10/Sars2/Ears2/Trmt10c/Mto1/Dars2/Supv3l1/Pnpt1/Tfam/Fastkd5/Sirt3/Mrpl12/Yars2/Tbrg4/Lrpprc |
| BP | GO:0071709 | membrane assembly | 17/1360 | 45/8928 | 0,000182 | 0,003662 | 0,00311 | Il1rapl1/Cav1/Pacsin2/Cav2/Lrp4/Ptprd/Nrxn2/Nrxn1/Nlgn1/Nlgn3/Nlgn4l/Magi2/Nlgn2/Cdh2/Ephb2/Pten/Sptbn1 |
| BP | GO:0015711 | organic anion transport | 53/1360 | 216/8928 | 0,000202 | 0,003995 | 0,003393 | Grm2/Slc17a6/Slc16a3/Slc4a7/Slc25a24/Slc25a21/Ace/Gja1/Slc5a6/Myo6/Slc4a8/Slc7a11/Slc1a2/Trpc4/Slc7a2/Sfxn5/Ace2/Slc1a1/Stard10/Pak1/Itgb1/Sfxn1/Slc25a23/Fis1/Sfxn3/Sfxn2/Abcb1a/Akt2/Apba1/Rbp1/Slc6a1/Slc25a11/Slc25a10/Slc25a12/Slc2a3/Abcc4/Gipc1/Slc25a15/Slc25a44/Slc25a19/Slc38a3/Slc25a4/Slc25a42/Mapk9/Slc25a1/Pnpla8/Nf1/Slc23a2/Stxbp1/Slc25a5/Slc25a25/Gabbr1/Acsl6 |
| BP | GO:0006935 | chemotaxis | 69/1360 | 301/8928 | 0,000221 | 0,004323 | 0,003671 | Plxnd1/Reln/Lama1/Cntn6/Lama5/Cdh13/Tiam1/Cntn4/Thbs4/Lamb2/Cxadr/Pdgfrb/Lrtm2/Plxnc1/Ptprj/Agrn/Hspb1/Gap43/Vangl2/Lsp1/Tpbg/Angpt1/Plxna3/Ptprm/Sema6b/Padi2/C1qbp/Swap70/Itga1/Pip5k1c/Robo2/L1cam/Pde4b/Hmgb2/Rac3/Plec/Akt2/Itgav/Lgals9/Dpysl4/Neo1/Flrt3/Evl/Sema4d/Dpysl5/Dag1/Mcu/Mapk3/Plxnb2/Ephb2/Tubb3/Nrcam/Lyn/Coro1a/Efnb1/Vcam1/Tenm2/Alcam/Lgi1/Cyrib/Itgb2/Itgam/Flrt2/Dnm1l/Pla2g7/Crkl/Myh10/Kalrn/Dusp3 |
| BP | GO:0042330 | taxis | 69/1360 | 301/8928 | 0,000221 | 0,004323 | 0,003671 | Plxnd1/Reln/Lama1/Cntn6/Lama5/Cdh13/Tiam1/Cntn4/Thbs4/Lamb2/Cxadr/Pdgfrb/Lrtm2/Plxnc1/Ptprj/Agrn/Hspb1/Gap43/Vangl2/Lsp1/Tpbg/Angpt1/Plxna3/Ptprm/Sema6b/Padi2/C1qbp/Swap70/Itga1/Pip5k1c/Robo2/L1cam/Pde4b/Hmgb2/Rac3/Plec/Akt2/Itgav/Lgals9/Dpysl4/Neo1/Flrt3/Evl/Sema4d/Dpysl5/Dag1/Mcu/Mapk3/Plxnb2/Ephb2/Tubb3/Nrcam/Lyn/Coro1a/Efnb1/Vcam1/Tenm2/Alcam/Lgi1/Cyrib/Itgb2/Itgam/Flrt2/Dnm1l/Pla2g7/Crkl/Myh10/Kalrn/Dusp3 |
| BP | GO:0051345 | positive regulation of hydrolase activity | 75/1360 | 335/8928 | 0,000261 | 0,004946 | 0,004201 | Rgs6/Tiam2/Rasgrp2/Rap1gap2/Cav2/Arhgap6/Tiam1/Dock10/Pdgfrb/Chn2/Icam1/Agrn/Antxr1/Sipa1l2/Arhgap22/Rasgrf2/Slc1a1/Garnl3/Rapgef3/Cd200/Asap1/Itgb1/Itga1/Arhgap25/Atp2a3/F3/Magi2/Slc39a10/Fis1/Hmgb2/Htra2/Agap1/Pkp4/Rap1gap/Hip1r/Akt2/Aifm1/Afdn/Gramd4/Bax/Phb2/Bad/Mmut/Tax1bp3/Ndufa13/Sema4d/Itga6/Ptprc/Asph/Hspd1/Rabgap1/Fgfr3/Mapk8/Mapk9/Lyn/Gnb5/Nf1/Dapk1/Map3k5/Ralgapb/Rabgap1l/Arhgef7/Dnaja3/Mtch2/Tbc1d10a/Sgsm1/Htr2a/Evi5l/Ralgapa1/Coro1c/Rap1gds1/Crkl/Abr/Iqsec3/Kalrn |
| BP | GO:0070584 | mitochondrion morphogenesis | 12/1360 | 27/8928 | 0,000278 | 0,005232 | 0,004443 | Mfn1/Fis1/Ssbp1/Plec/Bcl2l1/Bax/Nubpl/Supv3l1/Pnpt1/Cert1/Opa1/Dnm1l |
| BP | GO:2001224 | positive regulation of neuron migration | 9/1360 | 17/8928 | 0,000327 | 0,006047 | 0,005136 | Reln/Flna/Shtn1/Ptprz1/Il1r1/Arhgef2/Mapk8/Dab2ip/Tbc1d24 |
| BP | GO:0015867 | ATP transport | 7/1360 | 11/8928 | 0,000351 | 0,006372 | 0,005412 | Slc25a24/Gja1/Slc25a23/Slc25a4/Slc25a42/Slc25a5/Slc25a25 |
| BP | GO:0042886 | amide transport | 58/1360 | 247/8928 | 0,000359 | 0,006377 | 0,005417 | Grm2/Cplx3/Slc17a6/Adra2a/Kcnj6/Tiam1/Rbp4/Gja1/Trpc1/Slc5a6/Adcy8/Myo6/Slc7a11/Slc1a2/Pex5l/Glud1/Slc18a2/Hadh/Slc1a1/Anxa5/Stx1a/Pak1/Oxct1/Itgb1/Fgb/Ptprn2/Fgg/Syt7/Abcb1a/Nlgn2/Rims2/Apba1/Cpt1a/Ptpmt1/Bad/Slc14a1/Pde8b/Slc25a12/Pck2/Prkaca/Rab11fip5/Mcu/Gipc1/Cask/Slc38a3/Myo5a/Cert1/Nf1/Sirt3/Abcg1/Sri/Stxbp1/Arhgef7/Cdk16/Gabbr1/Oga/Kalrn/Pfkm |
| BP | GO:0016226 | iron-sulfur cluster assembly | 11/1360 | 24/8928 | 0,00036 | 0,006377 | 0,005417 | Fdx2/Fxn/Glrx5/Iba57/Abcb7/Nubpl/Isca2/Nfs1/Hscb/Nfu1/Ciapin1 |
| BP | GO:0031163 | metallo-sulfur cluster assembly | 11/1360 | 24/8928 | 0,00036 | 0,006377 | 0,005417 | Fdx2/Fxn/Glrx5/Iba57/Abcb7/Nubpl/Isca2/Nfs1/Hscb/Nfu1/Ciapin1 |
| BP | GO:0046189 | phenol-containing compound biosynthetic process | 11/1360 | 24/8928 | 0,00036 | 0,006377 | 0,005417 | Tph2/Trpc1/Ddc/Slc7a11/Dbh/Gch1/Th/Cyp2d22/Aldh2/Gipc1/Myo5a |
| BP | GO:0007229 | integrin-mediated signaling pathway | 20/1360 | 60/8928 | 0,000363 | 0,006408 | 0,005442 | Lama1/Fermt1/Flna/Lama5/Lamb2/Itga7/Itga3/Lamc1/Nid1/Itgb1/Itga1/Itgb8/Pxn/Tln1/Itgav/Itgb5/Itga6/Ilk/Itgb2/Itgam |
| BP | GO:0051046 | regulation of secretion | 92/1360 | 433/8928 | 0,000383 | 0,006728 | 0,005715 | Grm2/Grm8/Cplx3/Il1rapl1/Pcp4/Adra2a/Kcnj6/Pcsk1/Rab3b/Tmem132a/Tiam1/Doc2a/Rbp4/Slc6a4/Ace/Acvr2a/Gja1/Trpc1/Cspg5/Adcy8/Myo6/Slc4a8/Pex5l/Idh2/Entpd1/Glud1/Nlgn1/Hadh/Pam/Kcnc4/Anxa5/Sv2c/Cd200/Stx1a/Oxct1/Fgb/Maob/Rab3c/Ptprn2/Anpep/Fgg/Srcin1/Cd2ap/Syt7/Nlgn2/Hip1r/Rims2/Bcl2l1/Snx5/Cpt1a/Ptpmt1/Lgals9/Bad/Pde8b/Slc6a1/Neo1/Sergef/Pck2/Prkaca/Rab11fip5/Mcu/Cask/Rhot1/Myo5a/Mapk9/Lyn/Pld2/Nf1/Wnk3/Sirt3/Abcg1/Sri/Itgb2/Stxbp1/Arhgef7/Septin5/Cdk16/Itgam/Gabbr1/Pard6a/Htr2a/Pdcd6ip/Hgs/Rap1gds1/Hap1/Dnm1l/Oga/Myh10/Abr/Kalrn/Gnai2/Pfkm |
| BP | GO:0051494 | negative regulation of cytoskeleton organization | 37/1360 | 140/8928 | 0,000395 | 0,006811 | 0,005785 | Kank4/Tpm1/Arhgap6/Tmsb10/Kank2/Pecam1/Stmn1/Lmod1/Ctnna2/Map6d1/Swap70/Dbn1/Map2/Tmod1/Cdh5/Hip1r/Cgnl1/Carmil1/Evl/Tjp1/Add3/Coro2b/Arhgef2/Mapre1/Coro1a/Specc1l/Twf1/Cyrib/Arhgef7/Wasf2/Add2/Fkbp4/Hdac6/Eml4/Pak2/Sptbn1/Sptan1 |
| BP | GO:1902473 | regulation of protein localization to synapse | 12/1360 | 28/8928 | 0,000419 | 0,007123 | 0,00605 | Dlg1/Gpc4/Nlgn1/Nlgn3/Gpc6/Magi2/Nlgn2/Neto2/Dag1/Tnik/Iqsec2/Kalrn |
| BP | GO:0019725 | cellular homeostasis | 103/1360 | 496/8928 | 0,00042 | 0,007123 | 0,00605 | Grm2/Calb2/Prkg2/Camk2d/Cav1/Adra2a/Atp2b4/Slc4a7/Flna/Tgm2/Cav2/Tiam1/Aqp4/Hcrtr2/Slc8a3/Inpp4b/Icam1/Trpc1/Trpc5/Adcy8/Tns2/Slc4a8/Slc7a11/Trpc4/Slc39a6/Slc1a1/Dmxl2/Grik2/Oxct1/Slc25a27/Fxn/Slc25a23/Gls/Txn2/Ptprn2/Atp2a3/Tpp2/Slc39a10/Atp1a1/Fis1/Abcb7/Prdx3/Cdh5/Letm1/Txnrd2/Itgav/Ptpn6/Bax/Ptpmt1/Slc12a4/Fggy/Ftl1/Bad/Stoml2/Afg3l2/Pde8b/Igf1r/Sod2/Maip1/Dld/Calb1/Neo1/Immt/Micu1/Gpx1/Ptprc/Pck2/Asph/Prkaca/Rab11fip5/Synpo/Ccdc51/Mcu/Bag3/Abcb8/Atp1b2/Mapk3/Micu3/Rhot1/Slc38a3/Myo5a/Lyn/Slc30a9/Coro1a/Vcam1/Wnk3/Heph/Rmdn3/Slc12a6/Abcg1/Sri/Atp1a2/Cfl2/Cdk16/Itpr2/Atp2b2/Pdk3/Hk1/Gclc/Htr2a/Rap1gds1/Hap1/Pfkm |
| BP | GO:0070373 | negative regulation of ERK1 and ERK2 cascade | 17/1360 | 48/8928 | 0,000444 | 0,007511 | 0,006379 | Cav1/Timp3/Dlg1/Spred3/Spred1/Spry4/Ace2/Spred2/Ptprr/Ptprc/Ephb2/Lyn/Dab2ip/Sirt3/Pten/Csk/Dusp3 |
| BP | GO:0042417 | dopamine metabolic process | 13/1360 | 32/8928 | 0,000455 | 0,007677 | 0,00652 | Pde1b/Ddc/Dbh/Slc1a1/Gch1/Th/Maob/Sncb/Maoa/Cyp2d22/Aldh2/Myo5a/Itgam |

| | | | | | | | | |
|---|---|---|---|---|---|---|---|---|
| BP | GO:0098771 | inorganic ion homeostasis | 69/1360 | 309/8928 | 0,000491 | 0,008171 | 0,00694 | Slc17a6/Calb2/Camk2d/Cav1/Atp2b4/Flna/Tgm2/Cav2/Cp/Hcrtr2/Ace/Slc8a3/Inpp4b/Trpc1/Trpc5/Adcy8/Trpc4/Slc39a6/Slc1a1/Grik2/Slc25a27/Fxn/Slc25a23/Atp2a3/Anpep/Slc39a10/Atp1a1/Fis1/Abcb7/Cdh5/Letm1/Snx5/Itgav/Ptpn6/Bax/Slc12a4/Ftl1/Stoml2/Afg3l2/Sod2/Maip1/Calb1/Neo1/Immt/Micu1/Ptprc/Asph/Synpo/Mcu/Atp1b2/Micu3/Fgfr3/Myo5a/Lyn/Slc30a9/Fech/Coro1a/Heph/Rmdn3/Slc12a6/Sri/Atp1a2/Itpr2/Atp2b2/Htr2a/Cnnm3/Rap1gds1/Hap1/Gnai2 |
| BP | GO:0031346 | positive regulation of cell projection organization | 71/1360 | 321/8928 | 0,000545 | 0,009034 | 0,007673 | Plxnd1/Il1rapl1/Reln/Tiam2/Pcp4/Flna/Tenm1/Ptprf/Tiam1/Ptprd/Fnbp1l/Itga3/Negr1/Plxnc1/Nrxn1/Agrn/Trpc5/Hspb1/Camk1/Plxna3/Nlgn1/Vldlr/Dbn1/Pak1/Itgb1/Mfn1/Fxn/Iqgap1/Robo2/L1cam/Tenm3/Magi2/Fscn1/Shtn1/Afdn/Ddr1/Igf1r/Ptprz1/Carmil1/Kidins220/Dbnl/Sema4d/Itga6/Dnm3/Atp1b2/Ilk/Plxnb2/Ephb2/Cask/Palm/Nck1/Lyn/Dab2ip/Slitrk1/Nf1/Tnik/Pacsin1/Opa1/Twf1/Grip1/Wasf2/Ppp2r5d/Ube2v2/Tbc1d24/Coro1c/Acsl6/Dpysl3/Hap1/Dnm1l/Rufy3/Kalrn |
| BP | GO:0007189 | adenylate cyclase-activating G protein-coupled receptor signaling pathway | 18/1360 | 53/8928 | 0,000547 | 0,009034 | 0,007673 | Adra2a/Atp2b4/Gpr26/Adcy8/Gnal/Adgrl2/Adgrb3/Pde4b/Adcy1/Adgrb2/Adgrg1/Pde4a/Adgrl1/Prkaca/Adcy6/Adgrl3/Adcy2/Gnai2 |
| BP | GO:2000177 | regulation of neural precursor cell proliferation | 18/1360 | 53/8928 | 0,000547 | 0,009034 | 0,007673 | Flna/Slc6a4/Itgb1/Nr2e1/Adgrg1/Afdn/Cdh2/Ptprz1/Cdon/Ilk/Fgfr3/Mapk8/Lyn/Nap1l1/Ctnnb1/Nf1/Appl2/Gnai2 |
| BP | GO:0001505 | regulation of neurotransmitter levels | 46/1360 | 188/8928 | 0,000553 | 0,009108 | 0,007736 | Grm8/Cplx3/Slc17a6/Rab3b/Pde1b/Gad2/Doc2a/Slc29a1/Slc6a4/Nrxn2/Cspg5/Nrxn1/Slc4a8/Slc1a2/Slc18a2/Dbh/Nlgn1/Kcnc4/Dgki/Sv2c/Grik5/Th/Stx1a/Pak1/Itgb1/Maob/Ptprn2/Syt7/Rims2/Maoa/Aldh5a1/Bcl2l1/Slc6a2/Slc6a1/Pdzd11/Prkaca/Syngr3/Cask/Aldh9a1/Rhot1/Nf1/Sptbn2/Atp1a2/Stxbp1/Septin5/Dnm1l |
| BP | GO:0006816 | calcium ion transport | 57/1360 | 246/8928 | 0,000573 | 0,009327 | 0,007922 | Camk2d/Mylk/Cav1/Atp2b4/Flna/Pdgfrb/Ace/Slc8a3/Gja1/Icam1/Trpc1/Trpc5/Nos3/Epb41/Ahnak/Trpc4/Camk2g/Rapgef3/Itpr3/Slc25a23/Pde4b/Atp2a3/Letm1/Nipsnap2/Itgav/Ptpn6/Bax/Cacna2d1/Cacna2d3/Sestd1/Afg3l2/Pacsin3/Maip1/Micu1/Ptprc/Asph/Vdac1/Mcu/Micu3/Cask/Myo5a/Lyn/Coro1a/Gnb5/Ctnnb1/Wnk3/Opa1/Sri/Atp1a2/Slc25a25/Itpr2/Atp2b2/Htr2a/Hap1/Dnm1l/Oga/Gnai2 |
| BP | GO:0050769 | positive regulation of neurogenesis | 48/1360 | 199/8928 | 0,000598 | 0,009658 | 0,008203 | Plxnd1/Il1rapl1/Reln/Tiam2/Mme/Tgm2/Ptprf/Tiam1/Ptprd/Ace/Plxnc1/Trpc5/Plxna3/Vim/Dbn1/Pak1/Itgb1/Mfn1/Fxn/Nr2e1/Robo2/L1cam/Il6st/Shtn1/Sox8/Afdn/Ptprz1/Dbnl/Sema4d/Cdon/Dag1/Ilk/Plxnb2/Ephb2/Cask/Mapk8/Lyn/Nap1l1/Ctnnb1/Slitrk1/Tnik/Opa1/Grip1/Tbc1d24/Hap1/Dnm1l/Rufy3/Kalrn |
| BP | GO:0032272 | negative regulation of protein polymerization | 22/1360 | 71/8928 | 0,000598 | 0,009658 | 0,008203 | Kank4/Tpm1/Tmsb10/Pecam1/Stmn1/Lmod1/Map2/Tmod1/Cdh5/Hip1r/Carmil1/Evl/Add3/Mapre1/Vdac2/Twf1/Cyrib/Arhgef7/Add2/Fkbp4/Sptbn1/Sptan1 |
| BP | GO:1902904 | negative regulation of supramolecular fiber organization | 38/1360 | 148/8928 | 0,000612 | 0,009848 | 0,008364 | Kank4/Tpm1/Hspg2/Arhgap6/Tmsb10/Kank2/Pecam1/Stmn1/Lmod1/Ctnna2/Map6d1/Swap70/Dbn1/Map2/Tmod1/Cdh5/Hip1r/Cgnl1/Carmil1/Evl/Tjp1/Add3/Coro2b/Arhgef2/Mapre1/Coro1a/Specc1l/Twf1/Cyrib/Arhgef7/Wasf2/Add2/Fkbp4/Hdac6/Eml4/Pak2/Sptbn1/Sptan1 |
| MF | GO:0016491 | oxidoreductase activity | 145/1360 | 437/8928 | 5,15E-22 | 3,29E-19 | 2,79E-19 | Aoc3/Oxr1/Tph2/Vat1l/Fmo1/Cp/Ptgr3/Aldh1a1/Prodh/Mgst3/Aass/Nos3/Aldh6a1/Suclg2/Acads/Ivd/Idh2/Fdxr/Glud1/Aldh1l2/Cryzl2/Dbh/Bco2/Hadh/Pcbd2/Acad10/Pam/Aldh4a1/Bdh1/Acadm/Fdx2/Th/Acadl/Gstk1/Sqor/Fxn/Maob/Decr1/Uqcrb/Gcdh/Txn2/Glrx5/Suox/Ppox/Crat/Dhrs4/Nqo1/Etfa/Gldc/Prdx3/Acad8/Sardh/Sdr39u1/Mecr/Ddo/Mtarc2/Aifm1/Acad9/Maoa/Ndufa10/Txnrd2/Hsd17b10/Aldh5a1/Pdhx/Cox5b/Acadsb/L2hgdh/Sdhb/Hadha/Aldh7a1/Ndufs2/Hsdl2/Etfb/Sod2/Ogdhl/Ndufv2/Cbr4/Blvra/Dld/Cox5a/Hsd17b8/Etfdh/Ndufs3/Uqcrfs1/Gpd2/Ndufa2/Rdh13/Pdha1/Ndufs7/Mthfd1l/Cyp4f14/Ndufv1/Cyp2d22/Idh3g/Ndufs1/Aldh1b1/Fdx1/Pdhb/Aldh2/Gphn/Prxl2b/Gfus/Gpx1/Pdpr/Asph/Ndufaf5/Pir/Idh3a/Ogdh/Rtn4ip1/Mrps36/Coq6/Mdh2/Hibadh/Ndufa9/Ethe1/Sdha/Gfer/Cryz/Acadvl/Cox4i1/Aldh9a1/Aldh18a1/Bckdha/Cyc1/Idh3b/Hsd11b1/Adhfe1/Vcam1/Wwox/Cox15/Surf1/Heph/Msrb2/Gfod1/Me2/Dlat/Cyb5b/Dhrs1/Cbs/Dhodh/Pgd/Aifm3/Dhrs7b/Cyp2j9 |
| MF | GO:0050839 | cell adhesion molecule binding | 77/1360 | 201/8928 | 5,83E-16 | 1,08E-13 | 9,17E-14 | Ntng1/Vwf/Lrrc4c/Cntn6/Fermt1/Tenm1/Lama5/Ptprf/Cdh13/Ptprt/Thbs4/Lamb2/Ptprd/Cxadr/Itga7/Itga3/Actn1/Kirrel3/Icam2/Bcam/Nrxn2/Ctnnal1/Icam1/Ptprj/Nrxn1/Nos3/Esam/Ptprm/Trpc4/Utrn/Nlgn1/Ctnna2/Cdh12/Nlgn3/Cd200/Cdh8/Itgb1/Nectin2/Itga1/Itgb8/Fgb/L1cam/Cdh10/Fgg/Tenm3/Pxn/Cdh20/Cd2ap/Cdh6/Tln1/Cdh5/Ctnnd2/Pkp4/Nlgn2/Tln2/Shtn1/Afdn/Itgav/Ptpn6/Itgb5/Cdh2/Ptprz1/Neo1/Adgrl1/Tjp1/Itga6/Jam2/Ilk/Nrcam/Lyn/Ctnnb1/Vcam1/Tenm2/Itgb2/Itgam/Lcp1/Ctnnd1 |
| MF | GO:0016874 | ligase activity | 39/1360 | 107/8928 | 4,91E-08 | 2,94E-06 | 2,5E-06 | Ass1/Suclg2/Mccc1/Acsf2/Mccc2/Wars2/Lipt2/Gatc/Pcca/Pccb/Acss1/Pars2/Acsf3/Vars2/Gatb/Cad/Cars2/Qrsl1/Iars2/Sars2/Atp5c1/Nars2/Hlcs/Mars2/Ears2/Atp5pb/Mthfd1l/Dars2/Lars2/Atp5o/Sucla2/Rars2/Tars2/Aarsd1/Ctps2/Yars2/Gclc/Acsbg1/Acsl6 |
| MF | GO:0003779 | actin binding | 86/1360 | 324/8928 | 6,1E-08 | 3,58E-06 | 3,04E-06 | Myh11/Tagln/Tpm1/Mylk/Tpm2/Pdlim7/Fermt1/Flna/Sntb1/Actn1/Slc6a4/Cald1/Gas7/Ace/Ctnnal1/Tmsb10/Trpc5/Adcy8/Myo6/Nos3/Lsp1/Epb41/Cobll1/Gjb6/Ablim3/Antxr1/Utrn/Lmod1/Ctnna2/Tns1/Lasp1/Afap1/Dbn1/Itgb1/Iqgap1/Myo1c/Ablim2/Map2/Pdlim5/Tln1/Tmod1/Fscn1/Plec/Actn4/Hip1r/Phactr3/Vcl/Tln2/Shtn1/Coro2a/Afdn/Snta1/Slc6a2/Epb41l1/Dbnl/Evl/Add3/Coro2b/Dag1/Synpo/Gipc1/Epb41l2/Myo5a/Myo1b/Coro1a/Sptbn2/Twf1/Msrb2/Coro7/Cfl2/Wasf2/Syne1/Pls3/Ppp1r9b/Mprip/Ncald/Add2/Lcp1/Hdac6/Impact/Wasf3/Coro1c/Myh10/Sptbn1/Clmn/Sptan1 |
| MF | GO:0015453 | oxidoreduction-driven active transmembrane transporter activity | 15/1360 | 24/8928 | 1,72E-07 | 8,89E-06 | 7,55E-06 | Uqcrb/Ndufa10/Cox5b/Ndufs2/Ndufv2/Cox5a/Ndufs3/Uqcrfs1/Ndufa2/Ndufs7/Ndufv1/Ndufs1/Cox4i1/Cyc1/Surf1 |
| MF | GO:0004812 | aminoacyl-tRNA ligase activity | 15/1360 | 25/8928 | 3,69E-07 | 1,68E-05 | 1,43E-05 | Wars2/Pars2/Vars2/Cars2/Iars2/Sars2/Nars2/Mars2/Ears2/Dars2/Lars2/Rars2/Tars2/Aarsd1/Yars2 |
| MF | GO:0050660 | flavin adenine dinucleotide binding | 27/1360 | 66/8928 | 3,93E-07 | 1,77E-05 | 1,5E-05 | Fmo1/Prodh/Nos3/Acads/Ivd/Acad10/Acadm/Acadl/Sqor/Maob/Gcdh/Etfa/Acad8/Sardh/Ddo/Aifm1/Acad9/Maoa/Txnrd2/Acadsb/Dld/Mto1/Coq6/Sdha/Gfer/Acadvl/Aifm3 |
| MF | GO:0051536 | iron-sulfur cluster binding | 23/1360 | 53/8928 | 8,18E-07 | 3,38E-05 | 2,87E-05 | Fdx2/Fxn/Glrx5/Aco2/Ppat/Sdhb/Ndufs2/Nubpl/Ndufv2/Etfdh/Uqcrfs1/Ndufs7/Ndufv1/Ndufs1/Fdx1/Isca2/Nfs1/Cisd3/Nfu1/Fech/Dph1/Ciapin1/Aifm3 |
| MF | GO:0051540 | metal cluster binding | 23/1360 | 53/8928 | 8,18E-07 | 3,38E-05 | 2,87E-05 | Fdx2/Fxn/Glrx5/Aco2/Ppat/Sdhb/Ndufs2/Nubpl/Ndufv2/Etfdh/Uqcrfs1/Ndufs7/Ndufv1/Ndufs1/Fdx1/Isca2/Nfs1/Cisd3/Nfu1/Fech/Dph1/Ciapin1/Aifm3 |
| MF | GO:0050136 | NADH dehydrogenase (quinone) activity | 9/1360 | 11/8928 | 1,77E-06 | 6,49E-05 | 5,51E-05 | Nqo1/Ndufa10/Ndufs2/Ndufv2/Ndufs3/Ndufa2/Ndufs7/Ndufv1/Ndufs1 |
| MF | GO:0005178 | integrin binding | 30/1360 | 92/8928 | 2,22E-05 | 0,000602 | 0,000512 | Vwf/Fermt1/Lama5/Thbs4/Lamb2/Cxadr/Itga7/Itga3/Actn1/Icam2/Icam1/Utrn/Itgb1/Itga1/Itgb8/L1cam/Pxn/Tln1/Tln2/Itgav/Itgb5/Ptprz1/Itga6/Jam2/Ilk/Lyn/Vcam1/Itgb2/Itgam/Lcp1 |
| MF | GO:0140101 | catalytic activity, acting on a tRNA | 26/1360 | 78/8928 | 5,05E-05 | 0,001279 | 0,001087 | Wars2/Nsun6/Gatc/Dtd2/Mtfmt/Pars2/Vars2/Dtd1/Gatb/Cars2/Qrsl1/Iars2/Sars2/Nars2/Mars2/Ears2/Trmt10c/Dars2/Lars2/Trmu/Rars2/Tars2/Qtrt1/Aarsd1/Yars2/Mrpl58 |
| MF | GO:0051287 | NAD binding | 19/1360 | 51/8928 | 9,57E-05 | 0,002157 | 0,001832 | Aldh1a1/Idh2/Glud1/Hadh/Hsd17b10/Aldh5a1/Hadha/Ndufs2/Dld/Hsd17b8/Ndufv1/Sirt5/Idh3g/Aldh2/Idh3a/Hibadh/Aldh9a1/Sirt3/Me2 |
| MF | GO:0005509 | calcium ion binding | 75/1360 | 328/8928 | 0,00013 | 0,002829 | 0,002403 | Calb2/Aoc3/Rasgrp2/Pcp4/Ehd4/Ehd2/Matn4/Hpcal1/Hspg2/Tgm2/Lrp4/Cdh13/Slc25a24/Thbs4/Doc2a/Actn1/Nid1/Nrxn1/Itsn2/Agrn/Cd93/Necab1/Pam/Cdh12/Padi2/Anxa5/Gch1/Vldlr/Lrp1b/Cdh8/Itpr3/Tpd52/Slc25a23/Iqgap1/Cdh10/Astn2/Myl6b/Cdh20/Cdh6/Syt7/Sncb/Anxa4/Cdh5/Actn4/Esyt1/Ttyh1/Pitpnm2/Cdh2/Pitpnm3/Myl9/Calb1/Gpd2/Slc25a12/Micu1/Pdp1/Asph/Gca/Dag1/Anxa3/Eps15l1/Fat3/Micu3/Adgrl3/Rhot1/Myo5a/Capn2/Sri/Pls3/Slc25a25/Itpr2/Ncald/Atp2b2/Lcp1/Vcan/Sptan1 |

| | | | | | | | | |
|---|---|---|---|---|---|---|---|---|
| MF | GO:1901702 | salt transmembrane transporter activity | 75/1360 | 331/8928 | 0,000176 | 0,003645 | 0,003096 | Kcnj4/Slc17a6/Kcnh7/Kcnh3/Kcnj9/Slc16a3/Kcnj3/Atp2b4/Kcnj6/Hcn3/Slc4a7/Kcna3/Aqp4/Hcn1/Slc25a24/Hcn4/Slc6a4/Slc8a3/Trpc1/Slc5a6/Trpc5/Gabrg1/Slc4a8/Slc7a11/Slc1a2/Trpc4/Slc7a2/Sfxn5/Slc18a2/Hcn2/Slc39a6/Kcnc4/Slc1a1/Anxa5/Grik5/Grik2/Itpr3/Slc25a23/Atp2a3/Gabra4/Slc39a10/Atp1a1/Letm1/Itgav/Gabrd/Ttyh1/Slc12a4/Cacna2d1/Cacna2d3/Kcnk2/Slc6a2/Slc14a1/Slc6a1/Slc25a11/Slc25a10/Kcna6/Slc25a12/Abcc4/Ccdc51/Mcu/Slc25a15/Atp1b2/Slc25a19/Slc38a3/Slc25a4/Slc25a42/Slc25a1/Slc23a2/Slc12a6/Atp1a2/Slc25a5/Slc25a25/Itpr2/Atp2b2/Gabrb1 |
| MF | GO:0000062 | fatty-acyl-CoA binding | 9/1360 | 16/8928 | 0,000178 | 0,003645 | 0,003096 | Acads/Acadm/Acadl/Hmgcl/Gcdh/Hadha/Eci2/Acadvl/Acbd6 |
| MF | GO:0008514 | organic anion transmembrane transporter activity | 34/1360 | 121/8928 | 0,000193 | 0,00387 | 0,003287 | Slc17a6/Slc16a3/Slc4a7/Slc25a24/Slc25a21/Gja1/Slc5a6/Slc4a8/Slc7a11/Slc1a2/Slc7a2/Sfxn5/Slc1a1/Sfxn1/Slc25a23/Sfxn3/Sfxn2/Abcb1a/Slc6a1/Slc25a11/Slc25a10/Slc25a12/Slc2a3/Abcc4/Slc25a15/Slc25a44/Slc25a19/Slc38a3/Slc25a4/Slc25a42/Slc25a1/Slc23a2/Slc25a5/Slc25a25 |
| MF | GO:0005201 | extracellular matrix structural constituent | 24/1360 | 75/8928 | 0,000199 | 0,003962 | 0,003365 | Col1a2/Col6a3/Vwf/Col6a1/Tnc/Reln/Col4a2/Lama1/Prelp/Matn4/Col15a1/Hspg2/Lama5/Thbs4/Lamb2/Lamc1/Nid1/Vwa1/Agrn/Lama4/Fgb/Fgg/Mmrn2/Vcan |
| MF | GO:0030165 | PDZ domain binding | 27/1360 | 89/8928 | 0,000222 | 0,004325 | 0,003674 | Kcnj4/Kcnj9/Atp2b4/Cxadr/Dlg1/Sntb1/Kirrel3/Acvr2a/Gja1/Hcn2/Shisa9/Nlgn1/Adgrl2/Grik5/Grik2/L1cam/Grid2/Apba1/Snta1/Kidins220/Gipc1/Cask/Grip1/Pten/Tbc1d10a/Atp2b2/Dtna |
| MF | GO:0015075 | monoatomic ion transmembrane transporter activity | 86/1360 | 395/8928 | 0,000257 | 0,004885 | 0,004149 | Kcnj4/Slc17a6/Kcnh7/Kcnh3/Kcnj9/Slc16a3/Kcnj3/Atp2b4/Kcnj6/Hcn3/Slc4a7/Kcna3/Hcn1/Hcn4/Slc6a4/Slc8a3/Gja1/Trpc1/Slc5a6/Trpc5/Gabrg1/Slc4a8/Slc1a2/Pex5l/Trpc4/Sfxn5/Slc18a2/Hcn2/Slc39a6/Kcnc4/Slc1a1/Lasp1/Anxa5/Grik5/Grik2/Itpr3/Sfxn1/Uqcrb/Atp2a3/Gabra4/Slc39a10/Atp1a1/Grid2/Sfxn3/Slc2a13/Sfxn2/Letm1/Cox5b/Grid1/Itgav/Gabrd/Ttyh1/Slc12a4/Cacna2d1/Cacna2d3/Kcnk2/Atp5c1/Slc6a2/Slc6a1/Cox5a/Uqcrfs1/Atp5pb/Kcna6/Slc25a12/Atp5o/Vdac1/Ccdc51/Mcu/Tomm40/Atp1b2/Cox4i1/Slc38a3/Slc25a4/Slc30a9/Cyc1/Vdac2/Surf1/Slc23a2/Slc12a6/Atp1a2/Slc25a5/Mrs2/Itpr2/Atp2b2/Vdac3/Gabrb1 |
| MF | GO:0030552 | cAMP binding | 9/1360 | 17/8928 | 0,000327 | 0,006047 | 0,005136 | Hcn3/Hcn1/Hcn4/Hcn2/Rapgef3/Prkar2b/Pde4b/Pde4a/Pde2a |
| MF | GO:0050998 | nitric-oxide synthase binding | 9/1360 | 17/8928 | 0,000327 | 0,006047 | 0,005136 | Camk2d/Cav1/Atp2b4/Slc6a4/Nos3/Cdh2/Snta1/Dnm3/Ctnnb1 |
| MF | GO:1901567 | fatty acid derivative binding | 9/1360 | 17/8928 | 0,000327 | 0,006047 | 0,005136 | Acads/Acadm/Acadl/Hmgcl/Gcdh/Hadha/Eci2/Acadvl/Acbd6 |
| MF | GO:0005518 | collagen binding | 15/1360 | 39/8928 | 0,000344 | 0,006312 | 0,005361 | Vwf/Col6a1/Hspg2/Thbs4/Itga3/Nid1/Antxr1/Pak1/Itgb1/Itga1/Serpinh1/Adgrg1/Ddr1/Cspg4/Rell2 |
| MF | GO:0048038 | quinone binding | 7/1360 | 11/8928 | 0,000351 | 0,006372 | 0,005412 | Aoc3/Sqor/Ndufs2/Cbr4/Etfdh/Ndufs7/Dhodh |
| MF | GO:0008013 | beta-catenin binding | 22/1360 | 69/8928 | 0,000384 | 0,006728 | 0,005715 | Ptprt/Cxadr/Gja1/Ptprj/Ptprk/Nos3/Trpc4/Ctnna2/Pxn/Cd2ap/Cdh5/Ctnnd2/Vcl/Nherf2/Cdh2/Tax1bp3/Tjp1/Ctnnb1/Specc1l/Calcoco1/Ctnnd1/Hdac6 |
| MF | GO:0038023 | signaling receptor activity | 70/1360 | 312/8928 | 0,000388 | 0,006761 | 0,005742 | Grm2/Cd44/Grm8/Plxnd1/Grm4/Adra2a/Gfra2/Ptprf/Gpr26/Ptprt/Pdgfrb/Hcrtr2/Acvr2a/Nrxn2/Plxnc1/Nid1/Nrxn1/Pecam1/Gabrg1/Plxna3/Antxr1/Ptprm/Nlgn1/Adgrl2/Dcbld2/Or11h6/Nlgn3/Grik5/Vldlr/Grik2/Cd180/Itgb1/Itgb8/Adgrb3/Nlgn4l/Nr2e1/Robo2/L1cam/Grm3/Gabra4/Il6st/Grid2/Adgrb2/Nlgn2/Adgrg1/Grid1/Insr/Gabrd/Ptpn6/Itgb5/Ddr1/Igf1r/Neo1/Il1r1/Adgrl1/Sema4d/Cspg4/Cntfr/Gabbr2/Plxnb2/Ephb2/Adgrl3/Fgfr3/Itgb2/Cr1l/Abca7/Gabbr1/Htr2a/Ogfr/Gabrb1 |
| MF | GO:0060089 | molecular transducer activity | 70/1360 | 312/8928 | 0,000388 | 0,006761 | 0,005742 | Grm2/Cd44/Grm8/Plxnd1/Grm4/Adra2a/Gfra2/Ptprf/Gpr26/Ptprt/Pdgfrb/Hcrtr2/Acvr2a/Nrxn2/Plxnc1/Nid1/Nrxn1/Pecam1/Gabrg1/Plxna3/Antxr1/Ptprm/Nlgn1/Adgrl2/Dcbld2/Or11h6/Nlgn3/Grik5/Vldlr/Grik2/Cd180/Itgb1/Itgb8/Adgrb3/Nlgn4l/Nr2e1/Robo2/L1cam/Grm3/Gabra4/Il6st/Grid2/Adgrb2/Nlgn2/Adgrg1/Grid1/Insr/Gabrd/Ptpn6/Itgb5/Ddr1/Igf1r/Neo1/Il1r1/Adgrl1/Sema4d/Cspg4/Cntfr/Gabbr2/Plxnb2/Ephb2/Adgrl3/Fgfr3/Itgb2/Cr1l/Abca7/Gabbr1/Htr2a/Ogfr/Gabrb1 |
| MF | GO:0016597 | amino acid binding | 16/1360 | 44/8928 | 0,000463 | 0,007781 | 0,006609 | Ass1/Gad2/Prodh/Ddc/Nos3/Glud1/Slc1a1/Th/Gldc/Cad/Shmt2/Ddah1/Got2/Yars2/Gclc/Gfpt1 |
| MF | GO:0005516 | calmodulin binding | 37/1360 | 143/8928 | 0,000613 | 0,009848 | 0,008364 | Myh11/Grm4/Camk2d/Mylk/Pcp4/Atp2b4/Pde1b/Sntb1/Ace/Slc8a3/Adcy8/Gap43/Myo6/Nos3/Epb41/Camk1/Rasgrf2/Map6d1/Camk2g/Iqgap1/Camkv/Myo1c/Map2/Adcy1/Syt7/Snta1/Tjp1/Add3/Cask/Myo5a/Myo1b/Dapk1/Atp2b2/Add2/Myh10/Sptbn1/Sptan1 |
