## Supplementary material for "The molecular diversity of hippocampal regions and strata at synaptic resolution revealed by integrated transcriptomic and proteomic profiling": Table 2

| ONTOLOGY | ID | Description | GeneRatio | BgRatio | pvalue | p.adjust | qvalue | geneID | Count |
|---|---|---|---|---|---|---|---|---|---|
| MF | GO:0003735 | structural constituent of ribosome | 39/539 | 157/17474 | 1,31163E-24 | 6,75753E-21 | 6,22266E-21 | Rpsa/Rplp0/Rpl13a/Rps2/Rpl13/Rpl36/Rpl9/Rps20/Rpl26/Rps14/Rps18/Rps11/Rps4x/Rpl18a/Rps6/Rps15/Rps24/Rps25/Rps3/Rpl32/Rps17/Rps12/Rpl19/Rps5/Rplp2/Mrpl10/Srbd1/Rps9/Rpl10/Rpl35/Mrps2/Mrps18a/Mrpl49/Mrpl11/Mrpl44/Rpl34/Mrpl19/Mrps9/Rpl4 | 39 |
| MF | GO:0019843 | rRNA binding | 14/539 | 74/17474 | 4,96782E-08 | 1,50554E-05 | 1,38638E-05 | Rplp0/Rpl9/Rps14/Rps18/Rps11/Rps4x/Rps3/Rpl19/Rps5/Rps9/Mrps18a/Mrpl11/Rpl4/Ddx21 | 14 |
| MF | GO:0051020 | GTPase binding | 29/539 | 328/17474 | 3,91253E-07 | 0,000106091 | 9,76941E-05 | Rock2/Cyrib/Acap2/Arhgap1/Dennd5b/Rabgap1l/Clta/Dock9/Pak1/Becn1/Prkaca/Kif3a/Rabggtb/Nutf2/Pak3/Tsc2/Radil/Wdr44/Mff/Sgsm2/Mfn2/Dock3/Mical3/Dnm1l/Sgsm3/Xpo6/Fmnl1/Golga5/Coro1c | 29 |
| MF | GO:0043531 | ADP binding | 8/539 | 42/17474 | 3,64075E-05 | 0,004362131 | 0,004016861 | Pgk1/Hspa8/Vcp/Ppp5c/Myh10/Atp1a1/Chordc1/Prps1 | 8 |
| MF | GO:0044389 | ubiquitin-like protein ligase binding | 24/539 | 323/17474 | 6,92467E-05 | 0,007379027 | 0,006794965 | Pcbp2/Becn1/Prkaca/Tpi1/Srprb/Ube2k/Ube2n/Ywhae/Elob/Hspa8/Hspd1/Aup1/Bid/Cct2/Vcp/Tsg101/Lrpprc/Trim37/Mfn2/Yod1/Dnm1l/Nae1/Cdk5rap3/Cul5 | 24 |
| CC | GO:0022626 | cytosolic ribosome | 30/539 | 100/17474 | 8,43427E-22 | 1,74771E-18 | 1,60937E-18 | Rpsa/Rplp0/Rpl13a/Rps2/Rpl13/Rpl36/Rpl9/Rps20/Rpl26/Rps14/Rps18/Rps11/Rps4x/Rpl18a/Rps6/Rps15/Rps24/Rps25/Rps3/Rpl32/Rps17/Rps12/Rpl19/Rps5/Rplp2/Rps9/Rpl10/Rpl35/Rpl34/Rpl4 | 30 |
| CC | GO:0044391 | ribosomal subunit | 39/539 | 186/17474 | 1,01769E-21 | 1,74771E-18 | 1,60937E-18 | Rpsa/Rplp0/Rpl13a/Rps2/Rpl13/Rpl36/Rpl9/Rps20/Rpl26/Rps14/Rps18/Rps11/Rps4x/Rpl18a/Rps6/Rps15/Rps24/Rps25/Rps3/Rpl32/Rps17/Rps12/Rack1/Rpl19/Rps5/Rplp2/Mrpl10/Rps9/Rpl10/Rpl35/Mrps2/Mrps18a/Mrpl49/Mrpl11/Mrpl44/Rpl34/Mrpl19/Mrps9/Rpl4 | 39 |
| CC | GO:0099572 | postsynaptic specialization | 47/539 | 480/17474 | 2,49283E-12 | 1,16755E-09 | 1,07514E-09 | Hnrnpa2b1/Pcbp2/Kcnab2/Tanc2/Dlgap4/Dnajc6/Vdac1/Inpp4a/Pak1/Dclk1/Prrt1/Bmpr2/Strn/Gap43/Lrrc4b/Rplp0/Pcbp1/Rps14/Rps18/Rpl18a/Rps25/Rps3/Hspa8/Lrfn1/Camk2b/Lhfpl4/Kpna1/Pak3/Tsc2/Grik5/Eif4g2/Nlgn2/Bcr/Lrfn4/Lrrtm2/Rps6kc1/Lrfn3/Rgs7bp/Lrfn5/Rtn1/Atp1a1/Rheb/Rpl4/Arfgap1/Add2/Mpp2/Dmtn | 47 |
| CC | GO:0032279 | asymmetric synapse | 45/539 | 457/17474 | 6,16571E-12 | 2,44352E-09 | 2,25011E-09 | Hnrnpa2b1/Pcbp2/Kcnab2/Tanc2/Dnajc6/Vdac1/Inpp4a/Pak1/Dclk1/Prrt1/Bmpr2/Strn/Gap43/Lrrc4b/Rplp0/Pcbp1/Rps14/Rps18/Rpl18a/Rps25/Rps3/Hspa8/Lrfn1/Camk2b/Kpna1/Pak3/Tsc2/Grik5/Eif4g2/Bcr/Lrfn4/Lrrtm2/Rps6kc1/Lrfn3/Rgs7bp/Lrfn5/Slc4a8/Rtn1/Atp1a1/Rheb/Rpl4/Arfgap1/Add2/Mpp2/Dmtn | 45 |
| CC | GO:0043209 | myelin sheath | 23/539 | 216/17474 | 2,48512E-07 | 7,11296E-05 | 6,54996E-05 | Plec/Gnb5/Vdac1/Got2/Mdh2/Atp1b1/Sptan1/Ppia/Napb/Cycs/Hspa8/Hspd1/Pebp1/Cct2/Vcp/Prkci/Dlat/Atp1a1/Ndufa10/Napg/Idh3a/Plcb1/Sucla2 | 23 |
| CC | GO:0005759 | mitochondrial matrix | 26/539 | 295/17474 | 1,65941E-06 | 0,00040711 | 0,000374886 | Mipep/Vdac1/Timm44/Got2/Mdh2/Rps3/Hspd1/Mrpl10/Pptc7/Lrpprc/Pdhx/Dlat/Iba57/Fahd1/Mtg1/Pde2a/Mrps2/Hagh/Mrps18a/Mrpl49/Mrpl11/Acot13/Mrpl44/Mrpl19/Idh3a/Mrps9 | 26 |
| CC | GO:0150034 | distal axon | 31/539 | 394/17474 | 1,93247E-06 | 0,000452549 | 0,000416729 | Kcnab2/Slc9a6/L1cam/Agrn/Pak1/Dclk1/Slc18a3/Cdk5r2/Ap3d1/Got1/Calb1/Rpl26/Ywhae/Hspa8/Pcsk1/Pebp1/Ptprn2/Crmp1/Tsc2/Grik5/Myh10/Katnb1/Myo9a/Wdr47/Slc4a8/Sncb/Rtn4r/Ppp1r2/Tmod2/Septin6/Ror1 | 31 |
| CC | GO:0000502 | proteasome complex | 11/539 | 63/17474 | 3,1278E-06 | 0,000700628 | 0,000645172 | Psmb5/Psmc3/Psmd7/Rad23b/Psmc4/Psmc6/Psmb4/Vcp/Psmd6/Usp14/Psma1 | 11 |
| CC | GO:0008021 | synaptic vesicle | 23/539 | 256/17474 | 4,77893E-06 | 0,000984336 | 0,000906424 | Trim9/Clta/Vdac1/Atp6v0a1/Slc18a3/Prrt1/Cops5/Ppt1/Hspa8/Pebp1/Ptprn2/Ica1/Mff/Svop/Slc4a8/Dnm1l/Syngr3/Rab40c/Rab14/Rab27b/Sv2a/Scamp5/Septin6 | 23 |
| CC | GO:0098982 | GABA-ergic synapse | 15/539 | 134/17474 | 1,67366E-05 | 0,002612931 | 0,002406113 | Pak1/Gap43/Camk4/Calb1/Gabrg3/Rps6/Lhfpl4/Nlgn2/Gabbr2/Lrfn4/Lrrtm2/Cbln4/Lrfn5/Sv2a/Plcb1 | 15 |
| CC | GO:0005741 | mitochondrial outer membrane | 17/539 | 185/17474 | 6,16537E-05 | 0,00675829 | 0,00622336 | Plec/Vdac1/Gdap1/Hk1/Opa1/Pebp1/Bid/Vps13a/Dnajc11/Mff/Mfn2/Rps6kb1/Dnm1l/Pde2a/Tomm6/Slc25a46/Armc1 | 17 |
| CC | GO:0019867 | outer membrane | 18/539 | 211/17474 | 9,94587E-05 | 0,008820017 | 0,008121898 | Plec/Vdac1/Gdap1/Hk1/Opa1/Nutf2/Pebp1/Bid/Vps13a/Dnajc11/Mff/Mfn2/Rps6kb1/Dnm1l/Pde2a/Tomm6/Slc25a46/Armc1 | 18 |
| CC | GO:0099522 | cytosolic region | 8/539 | 49/17474 | 0,000115104 | 0,009883585 | 0,009101282 | Hnrnpa2b1/Trim9/Calb1/Prkce/Hspa8/Sncb/Pin1/Plcb1 | 8 |
| BP | GO:0002181 | cytoplasmic translation | 30/539 | 140/17474 | 2,91088E-17 | 2,99937E-14 | 2,76196E-14 | Rpsa/Rplp0/Rpl13a/Rps2/Rpl13/Rpl36/Rpl9/Rps20/Rpl26/Rps14/Rps18/Rps11/Rps4x/Rpl18a/Rps6/Rps15/Rps24/Rps25/Rps3/Rpl32/Rps17/Rps12/Rpl19/Rps5/Rplp2/Dph1/Rps9/Csde1/Rpl34/Rpl4 | 30 |
| BP | GO:0006163 | purine nucleotide metabolic process | 37/539 | 493/17474 | 6,21029E-07 | 0,000159977 | 0,000147315 | Kcnab2/Slc25a25/Slc4a7/Vdac1/Prkaca/Ldha/Mdh2/Pgk1/Atp1b1/Hk1/Tpi1/Opa1/Hspa8/Pfkfb2/Hint1/Vcp/Gpd1l/Pdhx/Dlat/Ppat/Dnm1l/Rptor/Cacnb4/Prpsap2/Hkdc1/Ndufv1/Pdk3/Pde2a/Ndufa10/Dmac2l/Ola1/Mpc2/Idh3a/Prps1/Itpa/Gda/Sucla2 | 37 |
| BP | GO:0019693 | ribose phosphate metabolic process | 32/539 | 433/17474 | 4,96754E-06 | 0,000984336 | 0,000906424 | Slc25a25/Vdac1/Prkaca/Pgk1/Atp1b1/Hk1/Tpi1/Opa1/Hspa8/Pfkfb2/Hint1/Vcp/Pdhx/Dlat/Ppat/Dnm1l/Rptor/Cacnb4/Prpsap2/Hkdc1/Ndufv1/Pdk3/Pde2a/Ndufa10/Dmac2l/Ola1/Mpc2/Nudt14/Prps1/Itpa/Gda/Sucla2 | 32 |
| BP | GO:0006090 | pyruvate metabolic process | 15/539 | 124/17474 | 6,4785E-06 | 0,001236193 | 0,001138346 | Vdac1/Prkaca/Ldha/Pgk1/Hk1/Tpi1/Pfkfb2/Pdhx/Dlat/Rptor/Fahd1/Hkdc1/Pdk3/Mpc2/Srr | 15 |
| BP | GO:0050807 | regulation of synapse organization | 25/539 | 302/17474 | 7,95737E-06 | 0,001464156 | 0,001348266 | Rock2/Tanc2/Agrn/Slc18a3/Lrrc4b/Ptk2/Gpc4/Opa1/Hspa8/Vcp/Lrfn1/Camk2b/Lhfpl4/Pak3/Tsc2/Myh10/Nlgn2/Mfn2/Lrfn4/Lrrtm2/Lrfn3/Lrfn5/Dnm1l/Rheb/Ube2v2 | 25 |
| BP | GO:0006605 | protein targeting | 24/539 | 284/17474 | 8,52192E-06 | 0,001513964 | 0,001394131 | C2cd5/Mipep/Pex10/Pak1/Timm44/Ap3d1/Becn1/Gdap1/Srprb/Ywhae/Hspa8/Hspd1/Bid/Vps13a/Hspa4/Mff/Trim37/Mfn2/Immp2l/Sgtb/Sec63/Ywhab/Pex14/Dmtn | 24 |
| BP | GO:0050803 | regulation of synapse structure or activity | 25/539 | 310/17474 | 1,24955E-05 | 0,002145892 | 0,001976041 | Rock2/Tanc2/Agrn/Slc18a3/Lrrc4b/Ptk2/Gpc4/Opa1/Hspa8/Vcp/Lrfn1/Camk2b/Lhfpl4/Pak3/Tsc2/Myh10/Nlgn2/Mfn2/Lrfn4/Lrrtm2/Lrfn3/Lrfn5/Dnm1l/Rheb/Ube2v2 | 25 |
| BP | GO:0022411 | cellular component disassembly | 31/539 | 437/17474 | 1,57257E-05 | 0,00253184 | 0,002331441 | Lrba/Atg2b/Dnajc6/Tpm1/Vdac1/Smarce1/Becn1/Sptan1/Napb/Mtpn/Hspa8/Vcp/Gbf1/Smarcc2/Epg5/Mfn2/Pelo/Zfand1/Katnb1/Wdr47/Mical3/Dnm1l/Uba5/Cdk5rap3/Slc25a46/Chmp5/Rb1cc1/Tmod2/Add2/Pex14/Dmtn | 31 |
| BP | GO:0006086 | acetyl-CoA biosynthetic process from pyruvate | 5/539 | 12/17474 | 1,81384E-05 | 0,002683355 | 0,002470963 | Vdac1/Pdhx/Dlat/Pdk3/Mpc2 | 5 |
| BP | GO:0042255 | ribosome assembly | 10/539 | 62/17474 | 1,82293E-05 | 0,002683355 | 0,002470963 | Rpsa/Rplp0/Rps14/Rps6/Rps15/Rps25/Rps5/Rpl10/Mrps2/Mrto4 | 10 |
| BP | GO:0000028 | ribosomal small subunit assembly | 6/539 | 20/17474 | 2,24505E-05 | 0,003116957 | 0,002870245 | Rpsa/Rps14/Rps6/Rps15/Rps25/Rps5 | 6 |
| BP | GO:0048488 | synaptic vesicle endocytosis | 11/539 | 77/17474 | 2,299E-05 | 0,003116957 | 0,002870245 | Dnajc6/Ap3d1/Napb/Ap2m1/Tbc1d24/Nlgn2/Mff/Dnm1l/Sncb/Rab27b/Scamp5 | 11 |
| BP | GO:0032271 | regulation of protein polymerization | 18/539 | 201/17474 | 5,30586E-05 | 0,005942565 | 0,005472201 | Cyrib/Clip1/Mapre3/Tpm1/Pak1/Prkce/Sptan1/Mtpn/Rps3/Pak3/Gba2/Mapk8/Arpc5l/Tmod2/Add2/Gda/Tenm1/Dmtn | 18 |
| BP | GO:0010763 | positive regulation of fibroblast migration | 6/539 | 24/17474 | 7,0181E-05 | 0,007379027 | 0,006794965 | Pak1/Prkce/Ptk2/Pak3/Tsc2/Dmtn | 6 |
| BP | GO:0006914 | autophagy | 31/539 | 475/17474 | 7,64681E-05 | 0,007724776 | 0,007113347 | Lrba/Atg2b/Vdac1/Atp6v0a1/Rnf5/Sptlc1/Becn1/Ptk2/Hspa8/Aup1/Bid/Vcp/Vps13a/Tsc2/Eif4g2/Soga3/Epg5/Mfn2/Pik3cb/Yod1/Wdr47/Dnm1l/Rptor/Uba5/Vps11/Slc25a46/Nsfl1c/Chmp5/Rb1cc1/Ubqln2/Tbk1 | 31 |
| BP | GO:0061919 | process utilizing autophagic mechanism | 31/539 | 475/17474 | 7,64681E-05 | 0,007724776 | 0,007113347 | Lrba/Atg2b/Vdac1/Atp6v0a1/Rnf5/Sptlc1/Becn1/Ptk2/Hspa8/Aup1/Bid/Vcp/Vps13a/Tsc2/Eif4g2/Soga3/Epg5/Mfn2/Pik3cb/Yod1/Wdr47/Dnm1l/Rptor/Uba5/Vps11/Slc25a46/Nsfl1c/Chmp5/Rb1cc1/Ubqln2/Tbk1 | 31 |
| BP | GO:0000266 | mitochondrial fission | 8/539 | 47/17474 | 8,47186E-05 | 0,008235283 | 0,007583446 | Cyrib/Gdap1/Mff/Mfn2/Ddhd2/Dnm1l/Slc25a46/Tmem135 | 8 |
| BP | GO:0099173 | postsynapse organization | 20/539 | 249/17474 | 9,62307E-05 | 0,008820017 | 0,008121898 | Rock2/Tanc2/Agrn/Gap43/Lrrc4b/Opa1/Hspa8/Lrfn1/Camk2b/Lhfpl4/Pak3/Tsc2/Myh10/Nlgn2/Mfn2/Lrfn4/Lrrtm2/Myo9a/Dnm1l/Rheb | 20 |
