## Supplementary material for "The molecular diversity of hippocampal regions and strata at synaptic resolution revealed by integrated transcriptomic and proteomic profiling": Table 3

| ONTOLOGY | ID | Description | GeneRatio | BgRatio | pvalue | p.adjust | qvalue | geneID | Count |
|---|---|---|---|---|---|---|---|---|---|
| BP | GO:0016054 | organic acid catabolic process | 47/946 | 236/17474 | 5,1266E-15 | 1,6925E-11 | 1,5048E-11 | Slc27a2/Bcat2/Gcat/Glud1/Oat/Eci1/Acadm/Bckdha/Arsb/Acadvl/Acadl/Cpt2/Dbt/Lipe/Slc16a3/Nos3/Renbp/Cpt1a/Acads/Mccc2/Abcd2/Hadh/Echs1/Hadha/Aldh5a1/Acaa2/Aldh4a1/Aldh1l2/Gldc/Gcsh/Abhd3/Etfdh/Ddo/Hadhb/Hibadh/Sardh/Nudt7/Decr1/Shmt2/Pgd/Npl/Etfb/Cyp4f14/Aldh6a1/Prodh/Eci2/Gstz1 | 47 |
| BP | GO:0046395 | carboxylic acid catabolic process | 47/946 | 236/17474 | 5,1266E-15 | 1,6925E-11 | 1,5048E-11 | Slc27a2/Bcat2/Gcat/Glud1/Oat/Eci1/Acadm/Bckdha/Arsb/Acadvl/Acadl/Cpt2/Dbt/Lipe/Slc16a3/Nos3/Renbp/Cpt1a/Acads/Mccc2/Abcd2/Hadh/Echs1/Hadha/Aldh5a1/Acaa2/Aldh4a1/Aldh1l2/Gldc/Gcsh/Abhd3/Etfdh/Ddo/Hadhb/Hibadh/Sardh/Nudt7/Decr1/Shmt2/Pgd/Npl/Etfb/Cyp4f14/Aldh6a1/Prodh/Eci2/Gstz1 | 47 |
| BP | GO:0044282 | small molecule catabolic process | 54/946 | 337/17474 | 6,045E-13 | 1,3305E-09 | 1,1829E-09 | Khk/Slc27a2/Bcat2/Gcat/Aldh1a1/Glud1/Oat/Eci1/Acadm/Pfkm/Inpp1/Bckdha/Arsb/Acadvl/Acadl/Cpt2/Dbt/Lipe/Slc16a3/Nos3/Renbp/Cpt1a/Acads/Mccc2/Abcd2/Hadh/Inpp5a/Echs1/Hadha/Aldh5a1/Acaa2/Aldh4a1/Aldh1l2/Gldc/Gcsh/Abhd3/Etfdh/Ddo/Hadhb/Hibadh/Sardh/Nudt7/Decr1/Shmt2/Dhdh/Pgd/Npl/Etfb/Cyp4f14/Aldh6a1/Nudt5/Prodh/Eci2/Gstz1 | 54 |
| BP | GO:0007229 | integrin-mediated signaling pathway | 22/946 | 99/17474 | 1,1461E-08 | 1,21E-05 | 1,0757E-05 | Lamc1/Itgam/Itgb1/Nid1/Itgb2/Lama1/Tln1/Ptpn11/Itgav/Dab2/Apoa1/Itgb8/Itga1/Lama2/Lama5/Lamb2/Itga6/Itga3/Fermt3/Plpp3/Plek/Rhoa | 22 |
| BP | GO:0006635 | fatty acid beta-oxidation | 19/946 | 76/17474 | 1,466E-08 | 1,21E-05 | 1,0757E-05 | Slc27a2/Eci1/Acadm/Acadvl/Acadl/Cpt2/Cpt1a/Acads/Abcd2/Hadh/Echs1/Hadha/Acaa2/Aldh1l2/Etfdh/Hadhb/Decr1/Etfb/Eci2 | 19 |
| BP | GO:0031589 | cell-substrate adhesion | 47/946 | 366/17474 | 3,2794E-08 | 2,3404E-05 | 2,0808E-05 | Sorbs1/Map4k4/Ptprz1/Utrn/Lamc1/Tiam1/Sdc4/Ninj1/Hsd17b12/Nid2/Itgam/Itgb1/Nid1/Col1a1/Itgb2/Atp1b2/Marcks/Ptpn11/Dnm2/Itgav/Tyro3/Dab2/Apoa1/Rsu1/Ctnnb1/Jup/Ddr1/Rac2/Pecam1/Cd36/Itgb8/Itga1/Mertk/Lama5/Lamb2/Cntn2/Bcan/Itga6/Itga3/Carmil1/Cspg5/Vwf/Fermt3/Poldip2/Rhoa/Bcam/Sorbs3 | 47 |
| BP | GO:1902904 | negative regulation of supramolecular fiber organization | 29/946 | 174/17474 | 5,8727E-08 | 3,2315E-05 | 2,873E-05 | Eml4/Gsn/Hspg2/S1pr1/Snca/F11r/Tmsb4x/Stmn1/Inpp5j/Evl/Clu/Pecam1/Specc1l/Capza1/Mapre1/Swap70/Cgnl1/Carmil1/Kank4/Wasf2/Kank2/Pak2/Arpin/Tubb4a/Lima1/Dysf/Flii/Hip1r/Add3 | 29 |
| BP | GO:0051494 | negative regulation of cytoskeleton organization | 28/946 | 171/17474 | 1,4543E-07 | 6,4019E-05 | 5,6917E-05 | Eml4/Gsn/S1pr1/Snca/F11r/Tmsb4x/Stmn1/Inpp5j/Evl/Pecam1/Specc1l/Capza1/Mapre1/Swap70/Cgnl1/Carmil1/Kank4/Wasf2/Tmem67/Kank2/Pak2/Arpin/Tubb4a/Lima1/Dysf/Flii/Hip1r/Add3 | 28 |
| BP | GO:0007264 | small GTPase mediated signal transduction | 49/946 | 425/17474 | 4,7985E-07 | 0,00015842 | 0,00014085 | Syde2/Map4k4/Mcf2l/Itsn1/Camk2d/Adcyap1r1/Arhgef12/Tiam1/F11r/Nras/Itgb1/Dnm2/Itgav/Stmn1/Rasa2/Nucb2/G3bp2/Rab33a/Apoa1/Rsu1/Sos2/Rac2/Pecam1/Iqsec3/Rapgef1/Rgl3/Garnl3/Rap1gap2/Chuk/Arhgdib/Itga3/Cgnl1/Psd2/Wasf2/Rab9b/Kank2/Arhgap25/Dock10/Sipa1l1/Dock2/Rtkn/Shtn1/Plcd4/Vangl2/Cd2ap/Git2/Cdc42ep4/Rhoa/Cyth1 | 49 |
| BP | GO:0007409 | axonogenesis | 53/946 | 482/17474 | 7,3055E-07 | 0,00021804 | 0,00019386 | Ank3/Aplp2/Dip2b/Plxnb2/Ptprz1/Nfasc/Tiam1/Trak2/Efna3/Septin7/Itgb1/Bsg/Lama1/Prkca/Mag/Ptprm/Ptpn11/Dnm2/Stmn1/Evl/Neo1/Wdr36/Prickle1/Cdkl5/Lama2/Foxg1/Lama5/Lamb2/Cntn2/Kif5b/Fgfr3/Nr2e1/Sema6d/Picalm/Trim46/Slit1/Dixdc1/Slitrk2/B4gat1/Sipa1l1/Pak2/Plxnb1/Rtn4rl1/Shtn1/Lrp4/Vangl2/Bcl11b/Tubb2b/Atg7/Bcl11a/Stk11/Slit3/B4galt6 | 53 |
| BP | GO:0044242 | cellular lipid catabolic process | 31/946 | 222/17474 | 1,2021E-06 | 0,00030569 | 0,00027178 | Pnpla7/Slc27a2/Sorl1/Eci1/Acadm/Acadvl/Acadl/Cpt2/Lipe/Cpt1a/Pld2/Acads/Abcd2/Hadh/Echs1/Hadha/Acaa2/Gpcpd1/Aldh1l2/Abhd6/Abhd3/Etfdh/Hadhb/Abhd12/Nudt7/Decr1/Gdpd1/Etfb/Cyp4f14/Eci2/Abhd16a | 31 |
| BP | GO:0015980 | energy derivation by oxidation of organic compounds | 39/946 | 318/17474 | 1,5566E-06 | 0,00036007 | 0,00032013 | Khk/Aco2/Sorbs1/Agl/Snca/Nipsnap2/Phkg1/Phka1/Cox4i1/Acadm/Pfkm/Adsl/Slc1a3/Gaa/Uqcrh/Il6st/Ndufv3/Afg1l/Grb10/Cox7a2l/Slc25a23/Phkb/Pygb/Coq9/Sdha/Uqcr10/Ugp2/Ndufa3/Mtfr1l/Sdhc/Shmt2/Cs/Ndufb8/Gbe1/Slc25a18/Etfb/Ide/Rhoa/Suclg2 | 39 |
| BP | GO:0032535 | regulation of cellular component size | 45/946 | 392/17474 | 1,5814E-06 | 0,00036007 | 0,00032013 | Inf2/Aqp4/Hp1bp3/Gsn/Dip2b/Dlg1/Bin1/Tmsb4x/Mag/Dnm2/Hcls1/Slc12a2/Evl/Pecam1/Specc1l/Wdr36/Ccdc51/Cdkl5/Pls1/Rilp/Capza1/Rap1gap2/Cntn2/Swap70/Carmil1/Kank4/Sema6d/Picalm/Trim46/Slit1/Lrrc8a/Lats1/Bag4/Shtn1/Esam/Slc12a9/Borcs6/Lima1/Plek/Flii/Hip1r/Rhoa/Add3/Bcl11a/Snapin | 45 |

| | | | | | | | | | |
|---|---|---|---|---|---|---|---|---|---|
| BP | GO:0110053 | regulation of actin filament organization | 34/946 | 273/17474 | 5,1296E-06 | 0,00084676 | 0,00075283 | Arhgef10l/Gsn/Dlg1/S1pr1/Bin1/Sdc4/F11r/Tmsb4x/Gja1/Hcls1/Stmn1/Evl/Apoa1/Pecam1/Specc1l/Capza1/Swap70/Cgnl1/Carmil1/Kank4/Wasf2/Kank2/Lats1/Bag4/Pak2/Esam/Arpin/Lima1/Plek/Flii/Hip1r/Rhoa/Add3/Sorbs3 | 34 |
| BP | GO:0007015 | actin filament organization | 47/946 | 443/17474 | 8,1298E-06 | 0,00129476 | 0,00115113 | Sorbs1/Arhgef10l/Gsn/Dlg1/Myo6/Cald1/S1pr1/Bin1/Sdc4/F11r/Tmsb4x/Gja1/Marcks/Hcls1/Stmn1/Evl/Apoa1/Rac2/Pecam1/Specc1l/Pls1/Capza1/Shroom1/Swap70/Cgnl1/Carmil1/Kank4/Wasf2/Kank2/Lats1/Arhgap25/Bag4/Pak2/Samd14/Shtn1/Sh3kbp1/Esam/Arpin/Nebl/Lima1/Plek/Flii/Hip1r/Cd2ap/Rhoa/Add3/Sorbs3 | 47 |
| BP | GO:0043547 | positive regulation of GTPase activity | 32/946 | 255/17474 | 8,2356E-06 | 0,00129476 | 0,00115113 | Map4k4/Agap1/Tiam1/S1pr1/Bin1/F11r/Itgb1/Mmut/Tbc1d10a/Eif5/Rsu1/Sos2/Iqsec3/Zc3h15/Rapgef1/Cdkl5/Rgl3/Garnl3/Rap1gap2/Itga6/Tbc1d12/Picalm/Scrib/Chn2/Tbc1d5/Tbc1d17/Arhgap25/Dock10/Sipa1l1/Snx18/Plxnb1/Cav2 | 32 |
| BP | GO:0043648 | dicarboxylic acid metabolic process | 17/946 | 94/17474 | 1,0424E-05 | 0,00149632 | 0,00133034 | Glud1/Oat/Kyat3/Dglucy/Aldh5a1/Aldh4a1/Aldh1l2/Sdha/Asl/Ddo/Me2/Shmt2/Cs/Nit2/Slc7a11/Prodh/Suclg2 | 17 |
| BP | GO:0071711 | basement membrane organization | 10/946 | 36/17474 | 1,4435E-05 | 0,00190623 | 0,00169477 | Lamc1/Itgb1/Nid1/Lama1/Cav1/Prickle1/Pxdn/Lama2/Lamb2/Cav2 | 10 |
| BP | GO:0032970 | regulation of actin filament-based process | 43/946 | 401/17474 | 1,5096E-05 | 0,00195452 | 0,0017377 | Arhgef10l/Gsn/Dlg1/S1pr1/Bin1/Sdc4/F11r/Hck/Tmsb4x/Gja1/Hcls1/Cav1/Stmn1/Evl/Apoa1/Jup/Pecam1/Specc1l/Capza1/Arhgdib/Swap70/Cgnl1/Carmil1/Sri/Kank4/Atp1a2/Dixdc1/Wasf2/Kank2/Lats1/Bag4/Pak2/Vangl2/Esam/Arpin/Lima1/Plek/Flii/Hip1r/Cd2ap/Rhoa/Add3/Sorbs3 | 43 |
| BP | GO:0008089 | anterograde axonal transport | 12/946 | 53/17474 | 2,0093E-05 | 0,00234312 | 0,00208319 | Ank3/Fmr1/Trak2/Cnih2/Hspb1/Kif5b/Trim46/Ap3s2/Arl8a/Dlg2/Caly/Snapin | 12 |
| BP | GO:0032272 | negative regulation of protein polymerization | 15/946 | 80/17474 | 2,1847E-05 | 0,00244499 | 0,00217376 | Gsn/Snca/Tmsb4x/Stmn1/Inpp5j/Evl/Pecam1/Capza1/Mapre1/Carmil1/Kank4/Tubb4a/Flii/Hip1r/Add3 | 15 |
| BP | GO:0043087 | regulation of GTPase activity | 38/946 | 349/17474 | 3,3381E-05 | 0,00335396 | 0,0029819 | Map4k4/Plxnb2/Agap1/Tiam1/S1pr1/Bin1/F11r/Itgb1/Mmut/Stmn1/Rasa2/Tbc1d10a/Eif5/Rsu1/Sos2/Pecam1/Iqsec3/Zc3h15/Rapgef1/Cdkl5/Rgl3/Garnl3/Rap1gap2/Itga6/Tbc1d12/Picalm/Scrib/Chn2/Tbc1d5/Tbc1d17/Arhgap25/Dock10/Sipa1l1/Snx18/Plxnb1/Poldip2/Tmed2/Cav2 | 38 |
| BP | GO:0006898 | receptor-mediated endocytosis | 31/946 | 265/17474 | 4,5279E-05 | 0,00439673 | 0,003909 | Myo6/Hspg2/Itsn1/Fmr1/Snca/Sorl1/Itgam/Itgb1/Itgb2/Prkca/Dnm2/Itgav/Cav1/Hpca/Pld2/Dab2/Clu/Cd36/Cntn2/Dlg4/Picalm/Scrib/Tbc1d5/Micall1/Grk2/Caly/Hip1r/Cd2ap/Syt11/Cav2/Unc119 | 31 |
| BP | GO:0030100 | regulation of endocytosis | 29/946 | 242/17474 | 5,0885E-05 | 0,0048695 | 0,00432932 | Ank3/Pacsin2/Nedd4l/Itsn1/Fmr1/Bin1/Snca/Itgb1/Prkca/Dnm2/Itgav/Cav1/Hpca/Pld2/Dab2/Clu/Cd36/Dlg4/Sh3gl1/Picalm/Snph/Scrib/Tbc1d5/Caly/Ehd4/Hip1r/Cd2ap/Syt11/Unc119 | 29 |
| BP | GO:0006163 | purine nucleotide metabolic process | 48/946 | 493/17474 | 6,0454E-05 | 0,00562224 | 0,00499856 | Khk/Nadk2/Dip2a/Snca/Slc4a4/Parg/Ldhb/Mmut/Fdx1/Pfkm/Adsl/Hmgcs2/Entpd1/Nudt2/Nos3/Hpca/Adcy5/Hmgcr/Ndufv3/Slc2a6/Mccc2/Fdxr/Nqo1/Nudt16/Atp1a2/Acaa2/Aldh1l2/Sdha/Pank1/Nudt4/Acsl6/Me2/Acss1/Nudt7/Ndufa3/Fis1/Sdhc/Cs/Mcee/Ndufb8/Pgd/Nudt12/Nudt5/Rhoa/Nme7/Ak3/Nme3/Suclg2 | 48 |
| BP | GO:1905039 | carboxylic acid transmembrane transport | 20/946 | 140/17474 | 6,5728E-05 | 0,00591973 | 0,00526305 | Slc1a2/Slc16a2/Itgb1/Slc3a2/Slc2a1/Slc7a2/Abcb1a/Slc1a3/Slc16a3/Cd36/Slc7a1/Slc5a6/Abcd2/Lrrc8a/Acsl6/Sfxn1/Slc25a18/Slc7a11/Slc25a15/Slc7a5 | 20 |
| BP | GO:0005977 | glycogen metabolic process | 14/946 | 79/17474 | 7,7922E-05 | 0,00625041 | 0,00555705 | Khk/Sorbs1/Agl/Phkg1/Phka1/Acadm/Pfkm/Gaa/Il6st/Grb10/Phkb/Pygb/Ugp2/Gbe1 | 14 |

| | | | | | | | | | |
|---|---|---|---|---|---|---|---|---|---|
| BP | GO:0006073 | cellular glucan metabolic process | 14/946 | 79/17474 | 7,7922E-05 | 0,00625041 | 0,00555705 | Khk/Sorbs1/Agl/Phkg1/Phka1/Acadm/Pfkm/Gaa/Il6st/Grb10/Phkb/Pygb/Ugp2/Gbe1 | 14 |
| BP | GO:2001046 | positive regulation of integrin-mediated signaling pathway | 6/946 | 15/17474 | 8,1289E-05 | 0,00638991 | 0,00568108 | Lamc1/Nid1/Lama1/Dab2/Lama2/Lamb2 | 6 |
| BP | GO:0044042 | glucan metabolic process | 14/946 | 80/17474 | 8,9817E-05 | 0,00691787 | 0,00615047 | Khk/Sorbs1/Agl/Phkg1/Phka1/Acadm/Pfkm/Gaa/Il6st/Grb10/Phkb/Pygb/Ugp2/Gbe1 | 14 |
| BP | GO:1905475 | regulation of protein localization to membrane | 25/946 | 201/17474 | 9,0101E-05 | 0,00691787 | 0,00615047 | Ank3/Camk2g/Sorbs1/Zmynd8/Gsn/Dlg1/Camk2d/Itgam/Itgb1/Csk/Hpca/Dab2/Pls1/Kif5b/Dlg4/Itga3/Ppp2r5a/Picalm/Atp2c1/Tcaf1/Lrp4/Fis1/Gpc6/Tmed2/Slc7a11 | 25 |
| BP | GO:0042428 | serotonin metabolic process | 6/946 | 16/17474 | 0,00012412 | 0,00871858 | 0,00775143 | Ddc/Aldh2/Rnf180/Maoa/Tph2/Cyp2d22 | 6 |
| BP | GO:0016358 | dendrite development | 34/946 | 320/17474 | 0,0001373 | 0,00948364 | 0,00843161 | Nedd4l/Zmynd8/Nsmf/Ptprz1/Dip2a/Myo6/Itsn1/Arid1b/Fmr1/Farp1/Tiam1/Trak2/Septin7/Itgb1/Marcks/Prickle1/Cdkl5/Mef2a/Dlg4/Nr2e1/Wls/Picalm/Pak4/Dock10/Sipa1l1/Pdlim5/Pak2/Lrp4/Camk1/Actl6b/Rhoa/Bcl11a/Stk11/Palm | 34 |
| CC | GO:0005759 | mitochondrial matrix | 42/946 | 295/17474 | 9,0088E-09 | 1,21E-05 | 1,0757E-05 | Aco2/Snca/Mmut/Glud1/Oat/Acadm/Fdx1/Aldh2/Bckdha/Acadvl/Acadl/Dbt/Hmgcs2/Acads/Mccc2/Pdpr/Dglucy/Hadha/Acaa2/Lonp1/Pitrm1/Ccar2/Poldip2/Pcca/Etfdh/Hadhb/Me2/Sardh/Pccb/Mccc1/Mrpl9/Acss1/Dhx30/Mrpl24/Pmpcb/Mrps22/Shmt2/Cs/Gstk1/Etfb/Ak3/Suclg2 | 42 |
| CC | GO:0042383 | sarcolemma | 28/946 | 166/17474 | 7,6215E-08 | 3,8712E-05 | 3,4417E-05 | Ank3/Aqp4/Ryr3/Dlg1/Camk2d/Utrn/Bin1/Akap7/Itgb1/Slc2a1/Bsg/Krt19/Des/Cav1/Nos3/Adcy5/Pld2/Cd36/Lama2/Slc30a1/Snta1/Sri/Atp1a2/Cacnb2/Dtna/Nos1ap/Dysf/Cav2 | 28 |
| CC | GO:0044853 | plasma membrane raft | 24/946 | 133/17474 | 1,7587E-07 | 7,2581E-05 | 6,4529E-05 | Pacsin2/Sorbs1/Adcyap1r1/Cavin1/Itgam/Hck/Itgb2/Slc2a1/Ctnnd1/Ptpn11/Bmpr1a/Cav1/Lipe/Nos3/Pld2/Ctnnb1/Cd36/Cavin2/Atp1a2/Ehd2/Lrp4/Grk2/Nos1ap/Cav2 | 24 |
| CC | GO:0031253 | cell projection membrane | 41/946 | 320/17474 | 2,6272E-07 | 9,6374E-05 | 8,5683E-05 | Aqp4/Ptprz1/Dlg1/Myo6/Utrn/Slc1a2/Tiam1/Ninj1/Itgb1/Ace/Atp1b2/Dnm2/Itgav/Inpp5j/Hpca/Anpep/Pld2/Ctnnb1/Cd36/Tmem231/Cdkl5/Jcad/Slc5a6/Mapre1/Itga3/Kcnc3/Wls/Psd2/Bbs9/Tmem67/Pex19/Snap29/Lima1/Plek/Hip1r/Rhoa/Podxl/Gabrg1/Slc7a11/Palm/Slc7a5 | 41 |
| CC | GO:0014069 | postsynaptic density | 50/946 | 437/17474 | 4,6233E-07 | 0,00015842 | 0,00014085 | Ank3/Camk2g/Nsmf/Ptprz1/Dlg1/Epb41l1/Myo6/Cald1/Fmr1/Tiam1/Cnih2/Igsf11/Itpr1/Hspb1/Map4/Ctnnd1/Dnm2/Entpd1/Slc16a3/Ctnnb1/Iqsec3/Insyn2a/Cdkl5/Fam81a/Slc30a1/Dlg4/Sh3gl1/Dgkb/Camk2n1/Picalm/Abhd17b/Scrib/Lrrtm3/Sipa1l1/Pdlim5/Pak2/Samd14/Lrp4/Dlg2/Camk1/Rnf112/Grk2/Tacc3/Hip1r/Homer2/Add3/Bcl11a/Syt11/Palm/Erc1 | 50 |
| CC | GO:0032279 | asymmetric synapse | 51/946 | 457/17474 | 7,595E-07 | 0,00021804 | 0,00019386 | Ank3/Camk2g/Nsmf/Ptprz1/Dlg1/Epb41l1/Myo6/Cald1/Fmr1/Tiam1/Cnih2/Itgb1/Igsf11/Itpr1/Hspb1/Map4/Ctnnd1/Dnm2/Entpd1/Slc16a3/Ctnnb1/Iqsec3/Insyn2a/Cdkl5/Fam81a/Slc30a1/Dlg4/Sh3gl1/Dgkb/Camk2n1/Picalm/Abhd17b/Scrib/Lrrtm3/Sipa1l1/Pdlim5/Pak2/Samd14/Lrp4/Dlg2/Camk1/Rnf112/Grk2/Tacc3/Hip1r/Homer2/Add3/Bcl11a/Syt11/Palm/Erc1 | 51 |
| CC | GO:0031594 | neuromuscular junction | 20/946 | 108/17474 | 1,2037E-06 | 0,00030569 | 0,00027178 | Ank3/Dlg1/Camk2d/Utrn/Lamc1/Itgb1/Des/Lama2/Lama5/Snta1/Lamb2/Dlg4/Itga3/Kcnc3/Erbin/Lrp4/Dlg2/Cd2ap/Cyth1/Thbs4 | 20 |
| CC | GO:0098636 | protein complex involved in cell adhesion | 14/946 | 57/17474 | 1,463E-06 | 0,00035778 | 0,00031809 | Lamc1/Itgam/Itgb1/Nid1/Itgb2/Lama1/Itgav/Itgb8/Itga1/Lama2/Lamb2/Itga6/Itga3/Jam2 | 14 |
| CC | GO:0043197 | dendritic spine | 29/946 | 206/17474 | 2,2323E-06 | 0,00044666 | 0,00039711 | Aplp2/Zmynd8/Nsmf/Ptprz1/Dip2a/Itsn1/Cald1/Fmr1/Slc1a2/Tiam1/Cnih2/Comt/Itgb1/Marcks/Ctnnd1/Slc1a3/Hpca/Lama2/Dlg4/Sri/Atp1a2/Fbxo2/Dock10/Sipa1l1/Grk2/Hip1r/Rhoa/Syt11/Palm | 29 |

| | | | | | | | | | |
|---|---|---|---|---|---|---|---|---|---|
| CC | GO:0098857 | membrane microdomain | 44/946 | 385/17474 | 2,3222E-06 | 0,00045099 | 0,00040096 | Pacsin2/Sorbs1/Dlg1/Adcyap1r1/Slc1a2/Sdc4/Cavin1/Itgam/Hck/Itgb1/Itgb2/Itpr1/Ldhb/Slc2a1/Bsg/Serpinh1/Prkca/Gja1/Ctnnd1/Ptpn11/Bmpr1a/Csk/Cav1/Lipe/Nos3/Adcy5/Pld2/Il6st/Ctnnb1/Pecam1/Cd36/Pi4k2a/Pag1/Itga1/Cavin2/Atp1a2/Ehd2/Rtn4rl1/Lrp4/Plpp3/Grk2/Nos1ap/Dysf/Cav2 | 44 |
| CC | GO:0044304 | main axon | 17/946 | 86/17474 | 2,9634E-06 | 0,00054354 | 0,00048324 | Ank3/Dlg1/Nfasc/Camk2d/Slc1a2/Tiam1/Bin1/Cldn5/Mag/Cntn2/Bcan/Dlg4/Kcnc3/Trim46/Gjc2/Dlg2/Tubb4a | 17 |
| CC | GO:0044309 | neuron spine | 29/946 | 212/17474 | 3,9982E-06 | 0,00069475 | 0,00061768 | Aplp2/Zmynd8/Nsmf/Ptprz1/Dip2a/Itsn1/Cald1/Fmr1/Slc1a2/Tiam1/Cnih2/Comt/Itgb1/Marcks/Ctnnd1/Slc1a3/Hpca/Lama2/Dlg4/Sri/Atp1a2/Fbxo2/Dock10/Sipa1l1/Grk2/Hip1r/Rhoa/Syt11/Palm | 29 |
| CC | GO:0005777 | peroxisome | 22/946 | 145/17474 | 1,1013E-05 | 0,00151493 | 0,00134687 | Akap11/Ech1/Slc27a2/Pxmp2/Cav1/Hmgcr/Hsdl2/Abcd2/Vwa8/Mavs/Pex19/Acsl6/Mtarc2/Ddo/Dhrs4/Pecr/Nudt7/Fis1/Gstk1/Nudt12/Ide/Eci2 | 22 |
| CC | GO:0042579 | microbody | 22/946 | 145/17474 | 1,1013E-05 | 0,00151493 | 0,00134687 | Akap11/Ech1/Slc27a2/Pxmp2/Cav1/Hmgcr/Hsdl2/Abcd2/Vwa8/Mavs/Pex19/Acsl6/Mtarc2/Ddo/Dhrs4/Pecr/Nudt7/Fis1/Gstk1/Nudt12/Ide/Eci2 | 22 |
| CC | GO:0032432 | actin filament bundle | 17/946 | 97/17474 | 1,6055E-05 | 0,0020387 | 0,00181254 | Myh11/Septin5/Sorbs1/Septin9/Septin7/Marcks/Ptpn11/Pls1/Mylk/Myl12a/Lpp/Pdlim5/Vangl2/Myl9/Nebl/Lima1/Bag3 | 17 |
| CC | GO:0030018 | Z disc | 20/946 | 128/17474 | 1,7577E-05 | 0,00218979 | 0,00194688 | Ank3/Ryr3/S100a1/Bin1/Hspb1/Slc2a1/Krt19/Des/Ctnnb1/Jup/Sphkap/Sri/Ppp2r5a/Myl12a/Pdlim5/Myl9/Nos1ap/Nebl/Bag3/Stk11 | 20 |
| CC | GO:0030055 | cell-substrate junction | 24/946 | 186/17474 | 6,9066E-05 | 0,00591973 | 0,00526305 | Sorbs1/Focad/Gsn/Map4k4/Tns1/Sdc4/Hck/Itgb1/Itgb2/Tln1/Itgav/Cav1/Evl/Rsu1/Itgb8/Itga1/Mapre1/Itga6/Erbin/Lpp/Fermt3/Lima1/Sorbs3/Cav2 | 24 |
| CC | GO:0005604 | basement membrane | 17/946 | 109/17474 | 7,531E-05 | 0,00621591 | 0,00552638 | Dlg1/Hspg2/Lamc1/Nid2/Itgb1/Nid1/Lama1/Col18a1/Timp3/Entpd1/Pxdn/Lama2/Lama5/Lamb2/Itga6/Vwa1/Thbs4 | 17 |
| CC | GO:0031252 | cell leading edge | 41/946 | 408/17474 | 0,00010328 | 0,00774921 | 0,00688959 | Gsn/Ptprz1/Myo6/Mcf2l/Itsn1/Slc1a2/Tiam1/Itgb1/Tln1/Ptprm/Ctnnd1/Dnm2/Itgav/Inpp5j/Evl/Hpca/Pld2/Ctnnb1/Rac2/Pecam1/Cdkl5/Jcad/Kcnc3/Wls/Carmil1/Psd2/Mylk/Scrib/Wasf2/Shtn1/Arpin/Lima1/Dysf/Plek/Hip1r/Cd2ap/Rhoa/Podxl/Gabrg1/Adgre5/Palm | 41 |
| CC | GO:0005938 | cell cortex | 32/946 | 295/17474 | 0,00014589 | 0,00982959 | 0,00873919 | Septin3/Septin5/Nsmf/Gsn/Septin9/Fry/Cald1/Cldn5/Snca/Septin7/Epb41l2/Slc2a1/Krt19/Marcks/Ctnnd1/Hcls1/Cav1/Ctnnb1/Pls1/Capza1/Shroom1/Mapre1/Dlg4/Myl12a/Septin10/Rtkn/Hip1r/Cd2ap/Rhoa/Add3/Itpr2/Erc1 | 32 |
| MF | GO:0050839 | cell adhesion molecule binding | 37/946 | 298/17474 | 2,1411E-06 | 0,00044181 | 0,0003928 | Ank3/Ptprz1/Nfasc/Utrn/Ninj1/F11r/Itgam/Itgb1/Itgb2/Prkca/Tln1/Ptprm/Ctnnd1/Ptpn11/Itgav/Nos3/Neo1/Dab2/Ctnnb1/Jup/Itgb8/Itga1/Lama5/Lamb2/Cntn2/Itga6/Itga3/Frmd5/Vwf/Fermt3/Shtn1/Esam/Plpp3/Jam2/Cd2ap/Bcam/Thbs4 | 37 |
| MF | GO:0005178 | integrin binding | 22/946 | 150/17474 | 1,9042E-05 | 0,00228607 | 0,00203248 | Ptprz1/Utrn/F11r/Itgam/Itgb1/Itgb2/Prkca/Tln1/Itgav/Dab2/Itgb8/Itga1/Lama5/Lamb2/Itga6/Itga3/Frmd5/Vwf/Fermt3/Plpp3/Jam2/Thbs4 | 22 |
| MF | GO:0050660 | flavin adenine dinucleotide binding | 16/946 | 90/17474 | 2,3548E-05 | 0,00259144 | 0,00230397 | Dus3l/Acadm/Acadvl/Acadl/Nos3/Acads/Aifm3/Maoa/Maob/Sdha/Ddo/Sardh/Pcyox1/Cyb5r3/Sqor/Prodh | 16 |
| MF | GO:0003729 | mRNA binding | 34/946 | 314/17474 | 9,4987E-05 | 0,00720917 | 0,00640945 | Eif4g3/Hnrnpc/Fmr1/Eif3d/Celf1/Cryz/Hnrnpa1/Cnbp/Tial1/G3bp2/Ddx5/Sf1/Nudt16/Khnyn/Pum1/Samd4b/Rbfox3/Thoc5/Lsm14b/Rbm14/Mettl3/Shfl/Casc3/Fyttd1/Hnrnpll/Rnps1/Ncbp2/Nhp2/Luc7l/Shmt2/Cdc40/Rbfox1/Khdrbs3/Celf2 | 34 |
| MF | GO:0003779 | actin binding | 43/946 | 436/17474 | 0,00010816 | 0,00802458 | 0,00713441 | Inf2/Myh11/Gsn/Crocc/Klhl3/Epb41l1/Myo6/Tns1/Utrn/Cald1/Bin1/Snca/Epb41l2/Itgb1/Ace/Tmsb4x/Tln1/Marcks/Hcls1/Nos3/Evl/Hpca/Cobll1/Pls1/Capza1/Shroom1/Snta1/Mylk/Dixdc1/Wasf2/Phactr3/Sipa1l1/Cacnb2/Pdlim5/Samd14/Shtn1/Ncald/Nebl/Lima1/Flii/Hip1r/Homer2/Add3 | 43 |
| MF | GO:0050998 | nitric-oxide synthase binding | 7/946 | 22/17474 | 0,00011093 | 0,00805101 | 0,00715791 | Camk2d/Dnm2/Cav1/Nos3/Ctnnb1/Snta1/Nos1ap | 7 |
| MF | GO:0051287 | NAD binding | 12/946 | 63/17474 | 0,00012201 | 0,00871858 | 0,00775143 | Ugdh/Ldhb/Aldh1a1/Glud1/Aldh2/Hadh/Hadha/Aldh5a1/Uxs1/Me2/Hibadh/Cyb5r3 | 12 |
| MF | GO:0043236 | laminin binding | 8/946 | 30/17474 | 0,00014418 | 0,00981439 | 0,00872568 | Itgb1/Nid1/Pxdn/Itga6/Itga3/Tinagl1/Bcam/Thbs4 | 8 |
