## Supplementary material for "The molecular diversity of hippocampal regions and strata at synaptic resolution revealed by integrated transcriptomic and proteomic profiling": Table 4

| iBAQ | rna | C..Type | N..Size | N..P.value | N..Benj..Hoch..FDR | T..Names |
|---|---|---|---|---|---|---|
| **0.527322** | 0.428839 | GOCC name | 152 | 0 | 0 | cell junction |
| **0.604497** | 0.477006 | GOCC name | 126 | 0 | 0 | synapse |
| **0.613918** | 0.572433 | GOCC name | 86 | 2.22045e-16 | 6.4837e-14 | glutamatergic synapse |
| **0.660375** | 0.427231 | GOCC name | 56 | 2.37775e-10 | 5.20728e-8 | postsynapse |
| **0.502977** | 0.245831 | GOCC name | 95 | 3.40425e-9 | 5.96424e-7 | cytoplasmic vesicle |
| **0.502977** | 0.245831 | GOCC name | 95 | 3.40425e-9 | 4.9702e-7 | intracellular vesicle |
| **0.318965** | 0.089795 | GOBP name | 296 | 7.06229e-9 | 4.15687e-5 | localization |
| **0.743845** | 0.454222 | GOCC name | 37 | 8.00014e-9 | 1.00116e-6 | presynapse |
| **0.479313** | 0.229501 | GOCC name | 100 | 9.08997e-9 | 9.95351e-7 | vesicle |
| **0.274799** | 0.29569 | GOBP name | 197 | 4.82242e-8 | 1.41924e-4 | regulation of biological quality |
| **0.481719** | 0.530051 | GOCC name | 53 | 4.8577e-8 | 4.72817e-6 | synaptic membrane |
| **0.323784** | 0.318006 | GOBP name | 145 | 5.74602e-8 | 1.12737e-4 | regulation of localization |
| **0.633385** | 0.5539 | GOCC name | 37 | 6.48096e-8 | 5.67732e-6 | postsynaptic density |
| **0.633385** | 0.5539 | GOCC name | 37 | 6.48096e-8 | 5.1612e-6 | postsynaptic specialization |
| **0.639176** | 0.557712 | GOBP name | 36 | 7.57214e-8 | 1.11424e-4 | regulation of vesicle-mediated transport |
| **0.285864** | 0.306823 | GOBP name | 172 | 7.62408e-8 | 8.97506e-5 | regulation of cellular component organization |
| **0.313814** | 0.0784224 | GOBP name | 243 | 8.83695e-8 | 8.66905e-5 | establishment of localization |
| **0.319223** | 0.0997892 | GOBP name | 228 | 1.20517e-7 | 1.01338e-4 | transport |
| **0.365524** | 0.332412 | GOBP name | 109 | 1.72569e-7 | 1.26968e-4 | regulation of transport |
| **0.382534** | 0.370736 | GOCC name | 88 | 3.17471e-7 | 2.31754e-5 | plasma membrane region |
| **0.469138** | 0.41501 | GOBP name | 61 | 3.63315e-7 | 2.37608e-4 | modulation of chemical synaptic transmission |
| **0.469138** | 0.41501 | GOBP name | 61 | 3.63315e-7 | 2.13847e-4 | regulation of trans-synaptic signaling |

| | | | | | | |
|---|---|---|---|---|---|---|
| **0.810692** | 0.69501 | GOCC name | 20 | 3.80062e-7 | 2.56103e-5 | Schaffer collateral - CA1 synapse |
| **0.316229** | 0.303507 | GOCC name | 135 | 3.90047e-7 | 2.44058e-5 | cell projection |
| **0.255745** | 0.124458 | GOCC name | 404 | 5.89007e-7 | 3.4398e-5 | membrane |
| **0.227946** | 0.276739 | GOBP name | 205 | 8.57895e-7 | 4.59052e-4 | regulation of signaling |
| **0.315747** | 0.292504 | GOCC name | 131 | 8.81368e-7 | 4.82549e-5 | plasma membrane bounded cell projection |
| **0.54251** | -0.658545 | GOBP name | 14 | 9.72639e-7 | 4.7708e-4 | translation |
| **0.333062** | 0.315797 | GOCC name | 110 | 1.18773e-6 | 6.12029e-5 | neuron projection |
| **0.378751** | -0.514473 | GOBP name | 25 | 1.21054e-6 | 5.48097e-4 | peptide metabolic process |
| **0.662686** | 0.523408 | GOCC name | 29 | 1.39653e-6 | 6.79643e-5 | transport vesicle membrane |
| **0.567353** | 0.624765 | GOBP name | 29 | 1.65234e-6 | 6.94692e-4 | trans-synaptic signaling |
| **0.262969** | 0.253328 | GOCC name | 183 | 1.81975e-6 | 8.39001e-5 | plasma membrane |
| **0.222277** | 0.269171 | GOBP name | 202 | 1.9383e-6 | 7.60589e-4 | regulation of cell communication |
| **0.50922** | -0.615198 | GOBP name | 15 | 2.16574e-6 | 7.96723e-4 | peptide biosynthetic process |
| **-0.292429** | -0.531032 | GOBP name | 52 | 2.17806e-6 | 7.5412e-4 | ncRNA processing |
| **0.174524** | 0.215783 | GOMF name | 430 | 3.00175e-6 | 0.00400734 | protein binding |
| **0.289819** | 0.321171 | GOBP name | 115 | 3.0554e-6 | 9.99116e-4 | positive regulation of signaling |
| **0.267612** | 0.263395 | GOMF name | 161 | 3.15577e-6 | 0.00210648 | enzyme binding |
| **0.514568** | 0.601674 | GOBP name | 31 | 3.35442e-6 | 0.00103916 | synaptic signaling |
| **0.766547** | -0.840057 | GOMF name | 7 | 3.62396e-6 | 0.00161266 | structural constituent of ribosome |
| **-0.378479** | -0.64143 | GOBP name | 32 | 4.83752e-6 | 0.00142368 | rRNA processing |
| **0.546211** | 0.6215 | GOBP name | 27 | 5.77649e-6 | 0.00161907 | anterograde trans-synaptic signaling |
| **0.546211** | 0.6215 | GOBP name | 27 | 5.77649e-6 | 0.00154547 | chemical synaptic transmission |
| **0.543307** | -0.0371169 | GOMF name | 39 | 5.87648e-6 | 0.00196128 | structural molecule activity |

| | | | | | | |
|---|---|---|---|---|---|---|
| **0.287257** | 0.310531 | GOBP name | 113 | 6.13305e-6 | 0.00156953 | positive regulation of cell communication |
| **0.554321** | 0.469907 | GOBP name | 35 | 6.13667e-6 | 0.00150502 | cell junction organization |
| **0.517215** | 0.542484 | GOBP name | 33 | 6.24792e-6 | 0.00147101 | cell-cell signaling |
| **0.565021** | 0.535871 | GOBP name | 30 | 7.41237e-6 | 0.00167805 | response to metal ion |
| **0.66141** | 0.616651 | GOCC name | 22 | 7.60803e-6 | 3.33232e-4 | excitatory synapse |
| **-0.0925834** | -0.250341 | GOBP name | 293 | 7.69754e-6 | 0.00167806 | cellular nitrogen compound metabolic process |
| **-0.258551** | -0.473624 | GOBP name | 59 | 7.80984e-6 | 0.00164174 | ncRNA metabolic process |
| **0.83776** | -0.703948 | GOBP name | 7 | 8.7887e-6 | 0.0017838 | cytoplasmic translation |
| **0.594163** | 0.528051 | GOBP name | 28 | 9.45822e-6 | 0.0018557 | synapse organization |
| **0.737014** | 0.653963 | GOCC name | 18 | 9.66149e-6 | 4.03022e-4 | synaptic vesicle |
| **0.37759** | 0.369739 | GOBP name | 66 | 9.70809e-6 | 0.00184328 | positive regulation of transport |
| **0.144198** | 0.222301 | GOBP name | 631 | 9.92031e-6 | 0.00182472 | regulation of biological process |
| **0.712247** | 0.547983 | GOCC name | 21 | 1.17553e-5 | 4.68075e-4 | exocytic vesicle membrane |
| **0.712247** | 0.547983 | GOCC name | 21 | 1.17553e-5 | 4.47724e-4 | synaptic vesicle membrane |
| **-0.125605** | -0.248094 | GOBP name | 268 | 1.25895e-5 | 0.00224551 | nucleobase-containing compound metabolic process |
| **0.464086** | 0.524509 | GOBP name | 35 | 1.38933e-5 | 0.00240517 | signaling |
| **0.373203** | 0.435732 | GOBP name | 53 | 1.39401e-5 | 0.00234433 | regulation of cell projection organization |
| **-0.352369** | -0.570451 | GOBP name | 36 | 1.50116e-5 | 0.00245439 | rRNA metabolic process |
| **0.360673** | 0.390803 | GOMF name | 62 | 1.5291e-5 | 0.00408271 | kinase binding |
| **0.529269** | 0.587426 | GOCC name | 27 | 1.53302e-5 | 5.59553e-4 | postsynaptic membrane |
| **-0.14705** | -0.239428 | GOBP name | 257 | 2.07415e-5 | 0.00329957 | nucleic acid metabolic process |
| **0.698492** | -0.78033 | GOCC name | 7 | 2.09992e-5 | 7.35813e-4 | ribosomal subunit |
| **0.556212** | 0.381386 | GOCC name | 35 | 2.13991e-5 | 7.20984e-4 | secretory vesicle |

| | | | | | | |
|---|---|---|---|---|---|---|
| **0.61273** | 0.54631 | GOBP name | 24 | 2.25589e-5 | 0.00349425 | regulation of endocytosis |
| **0.71446** | 0.520316 | GOCC name | 20 | 2.43634e-5 | 7.90456e-4 | exocytic vesicle |
| **0.360972** | 0.430547 | GOBP name | 52 | 2.4709e-5 | 0.00372915 | regulation of plasma membrane bounded cell projectio |
| **0.327361** | 0.143094 | GOMF name | 121 | 2.68116e-5 | 0.00596558 | identical protein binding |
| **0.635831** | 0.524715 | GOBP name | 23 | 2.91381e-5 | 0.00428767 | endocytosis |
| **0.61261** | 0.49826 | GOCC name | 25 | 3.00355e-5 | 9.39682e-4 | transport vesicle |
| **-0.119188** | -0.235596 | GOBP name | 272 | 3.16946e-5 | 0.00455011 | heterocycle metabolic process |
| **0.245265** | -0.4026 | GOBP name | 35 | 3.18061e-5 | 0.0044574 | amide metabolic process |
| **0.349492** | 0.324851 | GOMF name | 72 | 3.26783e-5 | 0.00623222 | protein domain specific binding |
| **0.419521** | 0.494749 | GOBP name | 37 | 3.54353e-5 | 0.00485052 | regulation of neuron projection development |
| **0.57478** | 0.535967 | GOCC name | 25 | 3.60611e-5 | 0.00108929 | dendritic spine |
| **0.57779** | 0.506542 | GOCC name | 26 | 3.72369e-5 | 0.00108732 | neuron spine |
| **0.392896** | 0.314001 | GOCC name | 62 | 3.98612e-5 | 0.0011264 | cell body |
| **-0.0902718** | -0.252687 | GOBP name | 221 | 4.02969e-5 | 0.00539063 | RNA metabolic process |
| **0.184024** | -0.166625 | GOCC name | 133 | 4.05046e-5 | 0.00110881 | ribonucleoprotein complex |
| **0.699253** | 0.631711 | GOBP name | 17 | 4.05976e-5 | 0.00531017 | actin cytoskeleton organization |
| **0.146259** | 0.203882 | GOBP name | 646 | 4,094E-02 | 0.00523854 | biological regulation |
| **0.301027** | 0.295139 | GOBP name | 92 | 4.13737e-5 | 0.00518139 | positive regulation of cellular component organization |
| **0.250098** | 0.0245405 | GOBP name | 204 | 4.28771e-5 | 0.00525781 | cellular localization |
| **-0.0415974** | -0.277769 | GOBP name | 160 | 4.48624e-5 | 0.00538899 | RNA processing |
| **0.382394** | 0.179039 | GOBP name | 79 | 4.75364e-5 | 0.00559599 | vesicle-mediated transport |
| **-0.111867** | -0.229701 | GOBP name | 275 | 4.99281e-5 | 0.00576229 | cellular aromatic compound metabolic process |
| **0.274427** | 0.0434597 | GOBP name | 162 | 5.26434e-5 | 0.00595882 | macromolecule localization |

| | | | | | | |
|---|---|---|---|---|---|---|
| **0.40841** | 0.299768 | GOCC name | 57 | 6.36956e-5 | 0.00169083 | neuronal cell body |
| **-0.125622** | -0.252902 | GOCC name | 198 | 6.6288e-5 | 0.00170789 | nuclear protein-containing complex |
| **0.196627** | 0.235066 | GOBP name | 181 | 6.7574e-5 | 0.00750454 | signal transduction |
| **0.400087** | 0.46989 | GOBP name | 38 | 6.86951e-5 | 0.00748777 | positive regulation of cell projection organization |
| **0.4067** | 0.250205 | GOCC name | 61 | 7.7189e-5 | 0.00193193 | vesicle membrane |
| **-0.0836259** | -0.204483 | GOBP name | 416 | 7.75832e-5 | 0.00830282 | nitrogen compound metabolic process |
| **-0.0749398** | -0.203293 | GOBP name | 437 | 7.89772e-5 | 0.00830107 | primary metabolic process |
| **0.66737** | 0.886432 | GOMF name | 11 | 8.46486e-5 | 0.0141257 | ligand-gated channel activity |
| **0.66737** | 0.886432 | GOMF name | 11 | 8.46486e-5 | 0.0125562 | ligand-gated monoatomic ion channel activity |
| **0.280201** | 0.0578589 | GOBP name | 147 | 8.52646e-5 | 0.00880469 | organic substance transport |
| **-0.105431** | -0.22175 | GOBP name | 280 | 8.56717e-5 | 0.0086942 | organic cyclic compound metabolic process |
| **0.41183** | 0.247964 | GOCC name | 59 | 8.6274e-5 | 0.00209933 | cytoplasmic vesicle membrane |
| **0.374343** | 0.292138 | GOCC name | 63 | 9.2753e-5 | 0.00219599 | dendrite |
| **0.226268** | 0.224026 | GOBP name | 156 | 1.03337e-4 | 0.0103091 | response to chemical |
| **0.233337** | 0.0158581 | GOCC name | 208 | 1.07456e-4 | 0.00247714 | organelle membrane |
| **0.484407** | 0.439594 | GOBP name | 32 | 1.07502e-4 | 0.010546 | regulation of synaptic plasticity |
| **0.267322** | 0.0563203 | GOBP name | 159 | 1.10165e-4 | 0.01063 | cellular macromolecule localization |
| **0.267322** | 0.0563203 | GOBP name | 159 | 1.10165e-4 | 0.0104586 | protein localization |
| **0.192266** | -0.0758567 | GOBP name | 801 | 1.1365e-4 | 0.0106182 | cellular process |
| **0.328737** | -0.30741 | GOBP name | 32 | 1.17104e-4 | 0.0107699 | organonitrogen compound biosynthetic process |
| **0.345992** | 0.363771 | GOMF name | 55 | 1.19949e-4 | 0.0160132 | protein kinase binding |
| **-0.113522** | -0.194792 | GOBP name | 401 | 1.20928e-4 | 0.0109505 | macromolecule metabolic process |
| **0.357971** | 0.0972775 | GOMF name | 81 | 1.26787e-4 | 0.0153873 | mRNA binding |

| | | | | | | |
|---|---|---|---|---|---|---|
| **0.235626** | 0.230616 | GOBP name | 139 | 1.27122e-4 | 0.0113369 | response to organic substance |
| **0.607301** | 0.508034 | GOBP name | 21 | 1.28295e-4 | 0.0112708 | cellular response to metal ion |
| **0.440477** | 0.387784 | GOBP name | 39 | 1.31038e-4 | 0.0113425 | response to inorganic substance |
| **0.664482** | 0.379787 | GOCC name | 21 | 1.31721e-4 | 0.00295867 | neuron to neuron synapse |
| **0.231616** | 0.332446 | GOBP name | 83 | 1.34631e-4 | 0.0114846 | response to oxygen-containing compound |
| **0.134141** | 0.186022 | GOBP name | 609 | 1.35078e-4 | 0.0113582 | regulation of cellular process |
| **0.298057** | 0.372378 | GOBP name | 59 | 1.44976e-4 | 0.0120187 | cellular response to oxygen-containing compound |
| **0.514059** | 0.525349 | GOCC name | 24 | 1.58279e-4 | 0.00346632 | postsynaptic specialization membrane |
| **0.268063** | 0.242916 | GOBP name | 108 | 1.59593e-4 | 0.0130467 | cellular response to chemical stimulus |
| **0.662963** | 0.78468 | GOMF name | 12 | 1.63144e-4 | 0.0181497 | neurotransmitter receptor activity |
| **0.259344** | 0.00933758 | GOBP name | 143 | 1.78803e-4 | 0.0144169 | establishment of localization in cell |
| **0.244541** | 0.310196 | GOBP name | 86 | 1.7943e-4 | 0.014272 | positive regulation of signal transduction |
| **0.657143** | 0.635056 | GOMF name | 15 | 1.91772e-4 | 0.0196935 | calmodulin binding |
| **0.581946** | 0.567489 | GOBP name | 19 | 1.94839e-4 | 0.0152909 | actin filament-based process |
| **0.260616** | 0.302385 | GOBP name | 83 | 2.08876e-4 | 0.0161769 | system process |
| **0.494489** | 0.43084 | GOBP name | 29 | 2.12512e-4 | 0.0162448 | import into cell |
| **0.741748** | 0.895043 | GOMF name | 9 | 2.13599e-4 | 0.0203682 | extracellular ligand-gated monoatomic ion channel acti |
| **0.741748** | 0.895043 | GOMF name | 9 | 2.13599e-4 | 0.0190103 | transmitter-gated channel activity |
| **0.741748** | 0.895043 | GOMF name | 9 | 2.13599e-4 | 0.0178222 | transmitter-gated monoatomic ion channel activity |
| **0.37495** | 0.40067 | GOBP name | 42 | 2.18312e-4 | 0.0164742 | regulation of monoatomic ion transport |
| **0.92485** | -0.775551 | GOBP name | 4 | 2.19782e-4 | 0.0163752 | translation at postsynapse |
| **0.92485** | -0.775551 | GOBP name | 4 | 2.19782e-4 | 0.0161705 | translation at presynapse |
| **0.92485** | -0.775551 | GOBP name | 4 | 2.19782e-4 | 0.0159708 | translation at synapse |

| | | | | | | |
|---|---|---|---|---|---|---|
| **0.789368** | -0.725577 | GOCC name | 5 | 2.39517e-4 | 0.00511749 | cytosolic small ribosomal subunit |
| **0.789368** | -0.725577 | GOCC name | 5 | 2.39517e-4 | 0.00499564 | small ribosomal subunit |
| **0.4712** | 0.53289 | GOMF name | 24 | 2.51274e-4 | 0.0197324 | transmembrane signaling receptor activity |
| **-0.309994** | -0.0462426 | GOBP name | 96 | 2,613E-01 | 0.0187563 | protein-DNA complex organization |
| **0.436226** | 0.478329 | GOBP name | 29 | 2.62725e-4 | 0.0186313 | cell projection morphogenesis |
| **0.436226** | 0.478329 | GOBP name | 29 | 2.62725e-4 | 0.0184095 | cellular anatomical entity morphogenesis |
| **0.547922** | 0.411218 | GOBP name | 25 | 2.86118e-4 | 0.0198128 | positive regulation of synaptic transmission |
| **-0.071722** | -0.185963 | GOBP name | 451 | 2.97348e-4 | 0.0203511 | organic substance metabolic process |
| **0.544101** | -0.340226 | GOCC name | 14 | 3.01529e-4 | 0.00614278 | ribosome |
| **0.46721** | 0.576817 | GOMF name | 21 | 3.28623e-4 | 0.0243728 | actin binding |
| **0.69853** | 0.633429 | GOBP name | 13 | 3.33666e-4 | 0.0225742 | receptor-mediated endocytosis |
| **0.686027** | 0.699495 | GOMF name | 12 | 3.45488e-4 | 0.0242751 | PDZ domain binding |
| **0.327448** | 0.493976 | GOCC name | 32 | 3.52055e-4 | 0.00700909 | transporter complex |
| **0.731453** | 0.841893 | GOMF name | 9 | 3.97553e-4 | 0.0265367 | postsynaptic neurotransmitter receptor activity |
| **0.238036** | -0.0545671 | GOBP name | 118 | 4.11765e-4 | 0.0275414 | nitrogen compound transport |
| **0.458859** | 0.58722 | GOBP name | 20 | 4.2022e-4 | 0.0277911 | calcium ion transport |
| **0.682352** | 0.624251 | GOBP name | 13 | 4.37219e-4 | 0.0285941 | long-term synaptic potentiation |
| **0.419412** | 0.385995 | GOBP name | 35 | 4.38129e-4 | 0.0283388 | regulation of neurogenesis |
| **0.714983** | 0.641077 | GOBP name | 12 | 4.45111e-4 | 0.0284774 | regulation of actin filament length |
| **0.714983** | 0.641077 | GOBP name | 12 | 4.45111e-4 | 0.0281712 | regulation of actin filament polymerization |
| **0.714983** | 0.641077 | GOBP name | 12 | 4.45111e-4 | 0.0278715 | regulation of actin polymerization or depolymerization |
| **0.576943** | 0.294223 | GOMF name | 24 | 4,783E-01 | 0.0304062 | phospholipid binding |
| **0.519864** | 0.291668 | GOBP name | 29 | 4.90644e-4 | 0.0303993 | regulation of synapse organization |

| | | | | | | |
|---|---|---|---|---|---|---|
| **0.265969** | 0.258251 | GOBP name | 86 | 4.9155e-4 | 0.0301382 | regulation of cellular localization |
| **0.548269** | 0.498727 | GOCC name | 20 | 4.95723e-4 | 0.00965008 | cytoplasmic region |
| **0.795549** | 0.951185 | GOMF name | 7 | 4.99679e-4 | 0.0303214 | glutamate receptor activity |
| **0.359405** | 0.285763 | GOBP name | 53 | 5.14727e-4 | 0.0312338 | cell communication |
| **0.294822** | 0.516214 | GOCC name | 29 | 5.22228e-4 | 0.00994503 | monoatomic ion channel complex |
| **0.294822** | 0.516214 | GOCC name | 29 | 5.22228e-4 | 0.00973344 | transmembrane transporter complex |
| **0.358857** | 0.412071 | GOBP name | 37 | 5.26696e-4 | 0.031634 | regulation of transmembrane transport |
| **0.275541** | 0.279626 | GOBP name | 75 | 5.26985e-4 | 0.0313316 | regulation of protein localization |
| **0.492435** | 0.59694 | GOCC name | 18 | 5.27907e-4 | 0.0096343 | GABA-ergic synapse |
| **0.286051** | -0.383977 | GOBP name | 23 | 5.30192e-4 | 0.0312071 | amide biosynthetic process |
| **0.371751** | 0.40574 | GOBP name | 36 | 5.82116e-4 | 0.0339241 | regulation of monoatomic ion transmembrane transpor |
| **0.2203** | 0.264726 | GOBP name | 97 | 5.98632e-4 | 0.0345446 | positive regulation of developmental process |
| **0.428985** | 0.350598 | GOCC name | 35 | 6.00356e-4 | 0.0107329 | cell surface |
| **0.415958** | 0.458859 | GOBP name | 28 | 6.05663e-4 | 0.034611 | plasma membrane bounded cell projection morphoger |
| **0.510668** | 0.468831 | GOCC name | 22 | 6.46415e-4 | 0.0113252 | presynaptic membrane |
| **-0.0727055** | -0.174576 | GOBP name | 472 | 6.5121e-4 | 0.036856 | metabolic process |
| **0.490122** | 0.512499 | GOCC name | 21 | 6.53071e-4 | 0.0112174 | receptor complex |
| **0.623732** | 0.509888 | GOBP name | 16 | 6.58782e-4 | 0.0369294 | regulation of actin filament organization |
| **-0.0205581** | -0.175064 | GOBP name | 372 | 6.59485e-4 | 0.0366201 | cellular metabolic process |
| **0.157377** | 0.225943 | GOBP name | 156 | 6.60745e-4 | 0.0363472 | regulation of signal transduction |
| **0.682891** | 0.823654 | GOCC name | 9 | 6.68334e-4 | 0.0112589 | neuromuscular junction |
| **0.697811** | 0.615152 | GOBP name | 12 | 6.69489e-4 | 0.0364871 | response to calcium ion |
| **0.487389** | 0.447981 | GOBP name | 24 | 6.77193e-4 | 0.0365684 | cellular response to inorganic substance |

| | | | | | | |
|---|---|---|---|---|---|---|
| **0.401718** | 0.48936 | GOBP name | 26 | 6.88655e-4 | 0.0368493 | regulation of protein-containing complex assembly |
| **0.657201** | 0.440035 | GOBP name | 16 | 6.8963e-4 | 0.036569 | regulation of postsynapse organization |
| **0.572684** | 0.680641 | GOBP name | 13 | 6.98756e-4 | 0.0367221 | regulation of postsynaptic membrane potential |
| **0.257252** | 0.313018 | GOBP name | 66 | 7.02497e-4 | 0.036592 | response to nitrogen compound |
| **0.820244** | 0.891601 | GOCC name | 7 | 7.04258e-4 | 0.0116402 | ionotropic glutamate receptor complex |
| **0.820244** | 0.891601 | GOCC name | 7 | 7.04258e-4 | 0.0114246 | neurotransmitter receptor complex |
| **0.602307** | 0.53068 | GOBP name | 16 | 7.18213e-4 | 0.0370825 | calcium ion homeostasis |
| **0.602307** | 0.53068 | GOBP name | 16 | 7.18213e-4 | 0.03676 | intracellular calcium ion homeostasis |
| **0.412693** | 0.397713 | GOCC name | 32 | 7.22709e-4 | 0.0115108 | plasma membrane protein complex |
| **0.702413** | 0.663884 | GOBP name | 11 | 7.26446e-4 | 0.0368609 | vesicle-mediated transport in synapse |
| **0.631001** | 0.568248 | GOBP name | 14 | 8.15334e-4 | 0.0410176 | regulation of receptor-mediated endocytosis |
| **0.511292** | 0.549345 | GOBP name | 18 | 8.2248e-4 | 0.0410264 | positive regulation of monoatomic ion transport |
| **0.425644** | 0.372809 | GOBP name | 32 | 8.24591e-4 | 0.0407861 | intracellular chemical homeostasis |
| **0.597611** | 0.478113 | GOBP name | 17 | 8.48255e-4 | 0.0416069 | regulation of cellular component size |
| **0.424714** | 0.263413 | GOCC name | 39 | 8.60208e-4 | 0.0134561 | endosome |
| **0.390489** | 0.319179 | GOBP name | 40 | 8.7162e-4 | 0.0423996 | regulation of nervous system development |
| **0.257447** | 0.317347 | GOBP name | 62 | 8.97802e-4 | 0.0433153 | nervous system process |
| **0.167453** | 0.129969 | GOCC name | 310 | 9.21572e-4 | 0.0141631 | cytoplasm |
| **0.567918** | 0.438882 | GOBP name | 19 | 9.28531e-4 | 0.0444336 | regulation of actin cytoskeleton organization |
| **0.264682** | -0.126844 | GOCC name | 941 | 9.41166e-4 | 0.0142149 | cellular anatomical entity |
| **0.303896** | 0.0783276 | GOCC name | 84 | 9.55064e-4 | 0.0141803 | endoplasmic reticulum |
| **0.423377** | 0.517208 | GOBP name | 22 | 9.60965e-4 | 0.0456148 | positive regulation of neuron projection development |
| **0.441914** | 0.433441 | GOBP name | 26 | 9.68087e-4 | 0.0455853 | regulation of monoatomic cation transmembrane trans |

| | | | | | | |
|---|---|---|---|---|---|---|
| **-0.18692** | -0.576345 | GOCC name | 22 | 9.77752e-4 | 0.0142752 | preribosome |
| **0.589757** | 0.5116 | GOBP name | 16 | 0.00100135 | 0.0467773 | regulation of protein localization to membrane |
| **0.378616** | 0.295245 | GOBP name | 43 | 0.00100501 | 0.0465788 | negative regulation of cellular component organization |
| **0.592953** | 0.46175 | GOCC name | 17 | 0.0010347 | 0.014859 | neuron projection cytoplasm |
| **0.592953** | 0.46175 | GOCC name | 17 | 0.0010347 | 0.0146193 | plasma membrane bounded cell projection cytoplasm |
| **0.276387** | 0.348403 | GOBP name | 50 | 0.00107775 | 0.0495596 | regulation of anatomical structure morphogenesis |
| **0.252052** | 0.128582 | GOCC name | 121 | 0.00116274 | 0.0161676 | bounding membrane of organelle |
| **0.425507** | 0.451955 | GOCC name | 25 | 0.00116647 | 0.0159661 | perikaryon |
| **0.347334** | 0.324078 | GOCC name | 43 | 0.00124257 | 0.016746 | axon |
| **0.57238** | 0.425587 | GOCC name | 18 | 0.00131975 | 0.0175167 | neuron projection terminus |
| **0.681574** | 0.473905 | GOCC name | 13 | 0.00136601 | 0.0178601 | axon terminus |
| **0.255308** | 0.389807 | GOCC name | 42 | 0.00137403 | 0.0177007 | cytoskeleton |
| **0.61904** | 0.568562 | GOCC name | 13 | 0.00142014 | 0.0180295 | dendritic shaft |
| **0.187394** | 0.063117 | GOCC name | 250 | 0.00156539 | 0.0195898 | cytosol |
| **0.478824** | 0.49826 | GOCC name | 19 | 0.00158503 | 0.0195562 | postsynaptic density membrane |
| **0.311343** | 0.15679 | GOCC name | 70 | 0.00172843 | 0.0210293 | membrane protein complex |
| **-0.228714** | 0.105391 | GOCC name | 76 | 0.00174071 | 0.0208885 | chromatin |
| **0.35522** | 0.326357 | GOCC name | 39 | 0.0017564 | 0.020792 | anchoring junction |
| **0.607926** | 0.65572 | GOCC name | 11 | 0.00178573 | 0.0208573 | terminal bouton |
| **0.718558** | 0.509128 | GOCC name | 11 | 0.00180029 | 0.0207507 | dendrite cytoplasm |
| **0.492895** | 0.816493 | GOCC name | 9 | 0.00188437 | 0.0214378 | presynaptic active zone membrane |
| **0.619239** | 0.90122 | GOCC name | 7 | 0.0018971 | 0.0213059 | parallel fiber to Purkinje cell synapse |
| **-0.116035** | -0.598925 | GOCC name | 17 | 0.00222512 | 0.0246735 | small-subunit processome |

| | | | | | | |
|---|---|---|---|---|---|---|
| **0.58706** | 0.26722 | GOCC name | 18 | 0.00223983 | 0.0245262 | side of membrane |
| **0.584586** | 0.508386 | GOCC name | 14 | 0.00226291 | 0.0244729 | lamellipodium |
| **0.476662** | 0.356797 | GOCC name | 23 | 0.0025389 | 0.0271228 | site of polarized growth |
| **0.490833** | 0.493282 | GOCC name | 17 | 0.00262065 | 0.0276589 | leading edge membrane |
| **0.643644** | -0.933267 | GOCC name | 3 | 0.00291193 | 0.0303673 | large ribosomal subunit |
| **0.830869** | 0.626561 | GOCC name | 7 | 0.0033721 | 0.0347524 | cytosolic region |
| **-0.208894** | 0.0751706 | GOCC name | 90 | 0.00342394 | 0.0348764 | protein-DNA complex |
| **0.681549** | 0.618328 | GOCC name | 9 | 0.00361987 | 0.0364483 | asymmetric synapse |
| **0.639666** | 0.523071 | GOCC name | 11 | 0.00376571 | 0.0374859 | plasma membrane signaling receptor complex |
| **933** | -958 | GOCC name | 2 | 0.00382702 | 0.0376682 | cytosolic large ribosomal subunit |
| **0.69489** | 0.368774 | GOCC name | 11 | 0.00411384 | 0.0400414 | cytoplasmic side of membrane |
| **0.250015** | -0.110444 | GOCC name | 53 | 0.00422006 | 0.0406238 | spliceosomal complex |
| **-0.133682** | -0.118189 | GOCC name | 343 | 0.00453161 | 0.0431488 | nucleoplasm |
| **0.39799** | -0.15208 | GOCC name | 21 | 0.00473039 | 0.0445572 | coated vesicle |
| **0.493043** | 0.19731 | GOCC name | 22 | 0.00485992 | 0.0452903 | mitochondrial membrane |
| **844** | -986 | GOCC name | 2 | 0.00508478 | 0.046887 | cytoplasmic side of endoplasmic reticulum membrane |
| **844** | -986 | GOCC name | 2 | 0.00508478 | 0.0463986 | cytoplasmic side of rough endoplasmic reticulum mem |
| **0.558844** | 0.237245 | GOCC name | 42 | 3.78474e-5 | 0.01529040000000 | synapse |
| **0.64966** | 0.555697 | GOCC name | 14 | 0.00100067 | 0.04491909999999 | neuron spine |
| **0.461395** | -0.1018 | GOCC name | 310 | 0 | 0 | mitochondrion |
| **0.517454** | -0.065476 | GOCC name | 113 | 1.35891e-13 | 5.95204e-11 | mitochondrial inner membrane |
| **0.509911** | -0.0636819 | GOCC name | 114 | 2.65343e-13 | 7.74802e-11 | organelle inner membrane |
| **0.449616** | -0.0394475 | GOCC name | 149 | 9.54348e-13 | 2.09002e-10 | mitochondrial membrane |

| | | | | | | |
|---|---|---|---|---|---|---|
| **0.292638** | -0.0941108 | GOBP name | 253 | 1.76898e-10 | 1.31913e-6 | small molecule metabolic process |
| **0.623602** | 0.00512974 | GOCC name | 61 | 2.81052e-10 | 4.92402e-8 | mitochondrial protein-containing complex |
| **0.220313** | -0.0601957 | GOBP name | 454 | 2.18382e-8 | 8.14236e-5 | cellular metabolic process |
| **0.122011** | 0.309129 | GOCC name | 264 | 2.46143e-8 | 3.59369e-6 | cell junction |
| **0.163882** | -0.130463 | GOBP name | 488 | 2.86518e-8 | 7.12187e-5 | organic substance metabolic process |
| **0.182615** | -0.0996053 | GOBP name | 549 | 3.50509e-8 | 6.53437e-5 | metabolic process |
| **0.330249** | -0.0710271 | GOBP name | 149 | 4.35275e-8 | 6.49169e-5 | oxoacid metabolic process |
| **0.326904** | -0.0728898 | GOBP name | 148 | 5.86632e-8 | 7.29086e-5 | carboxylic acid metabolic process |
| **0.317514** | -0.0752586 | GOBP name | 151 | 8.99931e-8 | 9.58683e-5 | organic acid metabolic process |
| **0.372263** | -0.0581499 | GOBP name | 113 | 1.08466e-7 | 1.01104e-4 | organonitrogen compound biosynthetic process |
| **0.843987** | 0.274984 | GOCC name | 25 | 2.0819e-7 | 2.60535e-5 | respiratory chain complex |
| **0.149304** | -0.126458 | GOBP name | 441 | 4.05161e-7 | 3.35698e-4 | primary metabolic process |
| **0.240023** | 0.326203 | GOCC name | 147 | 5.54602e-7 | 6.07289e-5 | synapse |
| **0.486353** | 0.0782246 | GOBP name | 69 | 6.93795e-7 | 5.17363e-4 | generation of precursor metabolites and energy |
| **0.7005** | 0.193366 | GOBP name | 33 | 7.86425e-7 | 5.33125e-4 | cellular respiration |
| **0.686259** | 0.153275 | GOCC name | 34 | 7.99667e-7 | 7.78343e-5 | inner mitochondrial membrane protein complex |
| **0.228949** | -0.111218 | GOBP name | 201 | 8.71716e-7 | 5.41699e-4 | cellular nitrogen compound metabolic process |
| **0.707761** | 0.202127 | GOBP name | 32 | 9.02567e-7 | 5.17726e-4 | aerobic respiration |
| **0.378024** | -0.0562225 | GOBP name | 92 | 1.04581e-6 | 5.57041e-4 | small molecule catabolic process |
| **0.370053** | -0.0594907 | GOBP name | 95 | 1.07999e-6 | 5.36898e-4 | organic cyclic compound biosynthetic process |
| **0.571867** | 0.217632 | GOBP name | 48 | 1.41643e-6 | 6.60143e-4 | energy derivation by oxidation of organic compounds |
| **0.30825** | -0.0950018 | GOMF name | 119 | 1.41738e-6 | 0.00279791 | oxidoreductase activity |
| **0.514751** | 0.0538364 | GOCC name | 56 | 1.47817e-6 | 1.29488e-4 | mitochondrial matrix |

| | | | | | | |
|---|---|---|---|---|---|---|
| **0.143229** | -0.126468 | GOBP name | 386 | 1.78919e-6 | 7.84822e-4 | nitrogen compound metabolic process |
| **0.548076** | 0.0758782 | GOBP name | 49 | 1,849E-03 | 7.65999e-4 | nucleoside phosphate biosynthetic process |
| **0.548076** | 0.0758782 | GOBP name | 49 | 1,849E-03 | 7.25683e-4 | nucleotide biosynthetic process |
| **0.201877** | -0.00206365 | GOCC name | 696 | 2.20213e-6 | 1.75369e-4 | intracellular membrane-bounded organelle |
| **0.156952** | -0.117032 | GOBP name | 346 | 2.41926e-6 | 9.02021e-4 | organonitrogen compound metabolic process |
| **0.336635** | -0.0316854 | GOBP name | 113 | 3.13783e-6 | 0.00111423 | cellular nitrogen compound biosynthetic process |
| **0.250429** | -0.0418615 | GOBP name | 209 | 3.1818e-6 | 0.00107848 | biosynthetic process |
| **0.677283** | -0.0199499 | GOBP name | 27 | 3.5779e-6 | 0.00116002 | nucleoside triphosphate biosynthetic process |
| **0.577898** | 0.077302 | GOBP name | 41 | 3.91273e-6 | 0.00121572 | ribonucleotide biosynthetic process |
| **0.577898** | 0.077302 | GOBP name | 41 | 3.91273e-6 | 0.00116709 | ribose phosphate biosynthetic process |
| **0.233303** | -0.00322717 | GOCC name | 289 | 3.96548e-6 | 2.8948e-4 | organelle membrane |
| **0.182152** | -0.148235 | GOBP name | 196 | 4.2487e-6 | 0.00121856 | organic cyclic compound metabolic process |
| **0.264354** | -0.0242936 | GOBP name | 191 | 4,433E-03 | 0.00122433 | cellular biosynthetic process |
| **0.209754** | -0.131139 | GOBP name | 176 | 4.54148e-6 | 0.00120949 | cellular aromatic compound metabolic process |
| **0.666684** | 0.142373 | GOCC name | 31 | 4.6279e-6 | 3.11849e-4 | oxidoreductase complex |
| **0.377975** | 0.0940232 | GOCC name | 103 | 4.77378e-6 | 2.98702e-4 | catalytic complex |
| **-0.109005** | 0.143694 | GOCC name | 412 | 4.96913e-6 | 2.90197e-4 | plasma membrane |
| **0.542038** | 0.080593 | GOBP name | 46 | 5.10267e-6 | 0.00131209 | purine-containing compound biosynthetic process |
| **0.382677** | -0.0281903 | GOBP name | 83 | 5.10902e-6 | 0.00126993 | aromatic compound biosynthetic process |
| **0.381116** | -0.01208 | GOBP name | 86 | 5.52074e-6 | 0.001328 | heterocycle biosynthetic process |
| **-0.0759074** | 0.154832 | GOBP name | 685 | 5.56812e-6 | 0.00129754 | regulation of cellular process |
| **-0.06737** | 0.161926 | GOBP name | 720 | 5.57783e-6 | 0.00126042 | regulation of biological process |
| **0.592372** | 0.0899141 | GOBP name | 38 | 5.58948e-6 | 0.0012259 | purine ribonucleotide biosynthetic process |

| | | | | | | |
|---|---|---|---|---|---|---|
| **0.359607** | -0.0790885 | GOBP name | 82 | 5.63841e-6 | 0.0012013 | carboxylic acid catabolic process |
| **0.359607** | -0.0790885 | GOBP name | 82 | 5.63841e-6 | 0.00116793 | organic acid catabolic process |
| **0.193092** | -0.0043002 | GOCC name | 708 | 5.81205e-6 | 3.1821e-4 | membrane-bounded organelle |
| **0.730477** | 0.0223672 | GOBP name | 23 | 6.04693e-6 | 0.0012187 | purine nucleoside triphosphate biosynthetic process |
| **0.730477** | 0.0223672 | GOBP name | 23 | 6.04693e-6 | 0.00118663 | purine ribonucleoside triphosphate biosynthetic proces |
| **0.553506** | 0.104045 | GOBP name | 44 | 6.07327e-6 | 0.00116124 | purine nucleotide biosynthetic process |
| **0.540684** | -0.0264646 | GOBP name | 40 | 6.51376e-6 | 0.00121433 | nucleoside triphosphate metabolic process |
| **0.473864** | -0.0226598 | GOBP name | 52 | 7.42639e-6 | 0.0013507 | carbohydrate derivative biosynthetic process |
| **0.249429** | -0.0382276 | GOBP name | 195 | 7.47553e-6 | 0.00132726 | organic substance biosynthetic process |
| **0.680643** | 0.00286174 | GOBP name | 25 | 9.51513e-6 | 0.0016501 | ribonucleoside triphosphate biosynthetic process |
| **0.90848** | 0.599772 | GOCC name | 14 | 1.00196e-5 | 5.16302e-4 | myelin sheath |
| **0.191807** | -0.132828 | GOBP name | 180 | 1.17857e-5 | 0.0019974 | heterocycle metabolic process |
| **0.558755** | 2.02757e-4 | GOBP name | 36 | 1.38335e-5 | 0.00229236 | purine nucleoside triphosphate metabolic process |
| **0.335858** | -0.0517897 | GOCC name | 92 | 1.45181e-5 | 7.06548e-4 | intracellular organelle lumen |
| **0.335858** | -0.0517897 | GOCC name | 92 | 1.45181e-5 | 6.69361e-4 | membrane-enclosed lumen |
| **0.335858** | -0.0517897 | GOCC name | 92 | 1.45181e-5 | 6.35893e-4 | organelle lumen |
| **0.579376** | 0.0254584 | GOBP name | 34 | 1.54891e-5 | 0.00251092 | purine ribonucleoside triphosphate metabolic process |
| **0.835839** | 0.141562 | GOBP name | 17 | 1.69159e-5 | 0.00268386 | proton motive force-driven ATP synthesis |
| **0.835839** | 0.141562 | GOBP name | 17 | 1.69159e-5 | 0.00262795 | proton motive force-driven mitochondrial ATP synthesi |
| **0.384058** | -0.0513768 | GOBP name | 69 | 1.73459e-5 | 0.00263976 | amino acid metabolic process |
| **-0.0745206** | 0.14693 | GOBP name | 743 | 1.76135e-5 | 0.00262687 | biological regulation |
| **0.553348** | 0.0116473 | GOBP name | 36 | 1.92546e-5 | 0.00281533 | ribonucleoside triphosphate metabolic process |
| **0.430632** | -0.0221116 | GOBP name | 56 | 2.45316e-5 | 0.00351793 | alpha-amino acid metabolic process |

| | | | | | | |
|---|---|---|---|---|---|---|
| **-0.168513** | 0.0864822 | GOBP name | 292 | 2.45738e-5 | 0.00345748 | signal transduction |
| **0.709895** | -0.0753684 | GOBP name | 19 | 2.64126e-5 | 0.00364738 | mitochondrial respiratory chain complex assembly |
| **0.304094** | -0.0532849 | GOBP name | 104 | 2.80783e-5 | 0.0038069 | nucleobase-containing small molecule metabolic proce |
| **0.73763** | 0.083707 | GOBP name | 20 | 3.1814e-5 | 0.00423638 | ATP biosynthetic process |
| **0.828312** | 0.126397 | GOCC name | 16 | 3.35019e-5 | 0.00139751 | mitochondrial respiratory chain complex I |
| **0.828312** | 0.126397 | GOCC name | 16 | 3.35019e-5 | 0.00133398 | NADH dehydrogenase complex |
| **0.828312** | 0.126397 | GOCC name | 16 | 3.35019e-5 | 0.00127598 | respiratory chain complex I |
| **0.254313** | 0.364078 | GOCC name | 79 | 3.43985e-5 | 0.00125554 | glutamatergic synapse |
| **0.788208** | -0.0031923 | GOBP name | 16 | 3.75138e-5 | 0.00490772 | mitochondrial respiratory chain complex I assembly |
| **0.788208** | -0.0031923 | GOBP name | 16 | 3.75138e-5 | 0.00482311 | NADH dehydrogenase complex assembly |
| **0.2412** | -0.0718651 | GOBP name | 147 | 4.01813e-5 | 0.00507851 | cellular catabolic process |
| **0.185872** | 0.0171243 | GOCC name | 790 | 4.11435e-5 | 0.00144167 | intracellular organelle |
| **0.119486** | -0.0978441 | GOMF name | 509 | 4.28201e-5 | 0.0422634 | catalytic activity |
| **0.191619** | 0.0404838 | GOCC name | 814 | 4.47737e-5 | 0.00150853 | organelle |
| **0.191534** | 0.272805 | GOCC name | 142 | 5.05695e-5 | 0.0016407 | neuron projection |
| **0.304419** | -0.0552181 | GOBP name | 96 | 5.10291e-5 | 0.00634207 | nucleoside phosphate metabolic process |
| **0.304419** | -0.0552181 | GOBP name | 96 | 5.10291e-5 | 0.0062381 | nucleotide metabolic process |
| **0.392761** | 0.0273302 | GOBP name | 68 | 5.13042e-5 | 0.00617057 | ribonucleotide metabolic process |
| **0.59327** | 0.0502503 | GOBP name | 29 | 5.3011e-5 | 0.00627466 | ATP metabolic process |
| **0.392327** | 0.0634457 | GOBP name | 70 | 6.06693e-5 | 0.00706892 | nucleobase-containing compound biosynthetic proces |
| **0.34266** | 0.51978 | GOCC name | 36 | 6.14231e-5 | 0.00192167 | postsynaptic specialization |
| **0.29858** | -0.0468435 | GOBP name | 100 | 6.24622e-5 | 0.00716585 | carbohydrate derivative metabolic process |
| **0.692134** | 0.158234 | GOBP name | 22 | 6.29849e-5 | 0.00711634 | electron transport chain |

| | | | | | | |
|---|---|---|---|---|---|---|
| **0.289595** | 0.448478 | GOCC name | 49 | 6.46693e-5 | 0.00195346 | postsynapse |
| **-0.385055** | 0.0936104 | GOMF name | 53 | 6.66604e-5 | 0.0438625 | GTPase regulator activity |
| **-0.385055** | 0.0936104 | GOMF name | 53 | 6.66604e-5 | 0.0328969 | nucleoside-triphosphatase regulator activity |
| **0.0906405** | 0.230686 | GOCC name | 241 | 6.73959e-5 | 0.00196796 | cell projection |
| **0.7253** | 0.209808 | GOBP name | 20 | 6,935E-02 | 0.00771855 | respiratory electron transport chain |
| **0.378623** | 0.0150725 | GOBP name | 69 | 7.36927e-5 | 0.00808127 | ribose phosphate metabolic process |
| **0.392154** | 0.0325326 | GOBP name | 65 | 8.09907e-5 | 0.00875286 | purine ribonucleotide metabolic process |
| **-0.549132** | 0.108424 | GOMF name | 25 | 1.11088e-4 | 0.0438574 | guanyl-nucleotide exchange factor activity |
| **0.341605** | 0.507978 | GOCC name | 34 | 1.32155e-4 | 0.00373444 | postsynaptic density |
| **0.288348** | -0.059597 | GOBP name | 93 | 1.3272e-4 | 0.0141385 | purine-containing compound metabolic process |
| **-0.155872** | 0.0843698 | GOBP name | 264 | 1.3764e-4 | 0.0144561 | regulation of signal transduction |
| **-0.149808** | 0.14876 | GOBP name | 161 | 1.43016e-4 | 0.0148121 | regulation of intracellular signal transduction |
| **-0.100477** | 0.130679 | GOBP name | 304 | 1.49878e-4 | 0.0153102 | regulation of signaling |
| **-0.137344** | 0.0847185 | GOBP name | 326 | 1.67711e-4 | 0.0169003 | regulation of response to stimulus |
| **-0.101135** | 0.127346 | GOBP name | 303 | 1.8495e-4 | 0.0183889 | regulation of cell communication |
| **0.80831** | 0.371542 | GOBP name | 14 | 1.85157e-4 | 0.0181673 | aerobic electron transport chain |
| **0.152628** | 0.34364 | GOBP name | 80 | 2.24139e-4 | 0.0217065 | regulation of neuron projection development |
| **0.289489** | -0.0508236 | GOBP name | 88 | 2.2873e-4 | 0.0218672 | purine nucleotide metabolic process |
| **0.0483092** | 0.224824 | GOCC name | 207 | 2.32609e-4 | 0.00636766 | plasma membrane region |
| **0.249511** | 0.36892 | GOCC name | 61 | 2.33299e-4 | 0.00619302 | synaptic membrane |
| **0.093421** | 0.218524 | GOCC name | 224 | 2.38424e-4 | 0.00614292 | plasma membrane bounded cell projection |
| **-0.0495519** | 0.245051 | GOBP name | 122 | 2.48943e-4 | 0.0234983 | anatomical structure morphogenesis |
| **0.203485** | -0.076621 | GOBP name | 151 | 2.73774e-4 | 0.0255191 | nucleobase-containing compound metabolic process |

| | | | | | | |
|---|---|---|---|---|---|---|
| **0.456823** | -0.0530603 | GOBP name | 34 | 3.17202e-4 | 0.0292022 | fatty acid beta-oxidation |
| **0.456823** | -0.0530603 | GOBP name | 34 | 3.17202e-4 | 0.0288461 | fatty acid oxidation |
| **0.163591** | 0.121274 | GOCC name | 388 | 3.22308e-4 | 0.0080669 | protein-containing complex |
| **0.34829** | 0.0400334 | GOBP name | 70 | 3.30683e-4 | 0.0297097 | organophosphate biosynthetic process |
| **0.143225** | 0.297937 | GOBP name | 101 | 3.4501e-4 | 0.0306278 | regulation of cell projection organization |
| **0.143225** | 0.297937 | GOBP name | 101 | 3.4501e-4 | 0.0302675 | regulation of plasma membrane bounded cell projectio |
| **0.163973** | -0.100536 | GOBP name | 180 | 3.4656e-4 | 0.03005 | organic substance catabolic process |
| **0.491671** | 0.771374 | GOCC name | 13 | 3.49775e-4 | 0.0085112 | Schaffer collateral - CA1 synapse |
| **0.429425** | 0.0366234 | GOBP name | 44 | 3.51759e-4 | 0.0301502 | amino acid catabolic process |
| **0.550465** | 0.739343 | GOCC name | 13 | 3.7594e-4 | 0.00890063 | glial cell projection |
| **0.16866** | -0.0824502 | GOBP name | 194 | 3.80553e-4 | 0.0322475 | catabolic process |
| **0.109395** | 0.503849 | GOBP name | 33 | 3.85212e-4 | 0.0322756 | neuron projection morphogenesis |
| **0.00862941** | 0.332126 | GOBP name | 71 | 4.10838e-4 | 0.0340402 | regulation of nervous system development |
| **0.241385** | 0.180252 | GOCC name | 132 | 4.27011e-4 | 0.00984374 | membrane protein complex |
| **0.290799** | -0.0268283 | GOBP name | 85 | 4.43691e-4 | 0.0363583 | monocarboxylic acid metabolic process |
| **-0.168003** | 0.113039 | GOBP name | 151 | 4.47246e-4 | 0.0362512 | cell surface receptor signaling pathway |
| **0.111931** | 0.491379 | GOBP name | 34 | 4.52786e-4 | 0.0363057 | plasma membrane bounded cell projection morphoger |
| **0.171107** | 0.348202 | GOBP name | 68 | 4.92475e-4 | 0.0390679 | cell junction organization |
| **0.446492** | -0.0249132 | GOBP name | 35 | 4.96809e-4 | 0.0389969 | lipid oxidation |
| **0.336095** | 0.466224 | GOCC name | 32 | 5.4134e-4 | 0.0121593 | neuron spine |
| **0.119165** | 0.46275 | GOBP name | 37 | 6.05715e-4 | 0.0470502 | cell projection morphogenesis |
| **0.119165** | 0.46275 | GOBP name | 37 | 6.05715e-4 | 0.0465651 | cellular anatomical entity morphogenesis |
| **-0.110054** | 0.15405 | GOBP name | 164 | 6.54687e-4 | 0.0498164 | positive regulation of cell communication |

| | | | | | | |
|---|---|---|---|---|---|---|
| **0.0369631** | 0.154444 | GOCC name | 722 | 7.40224e-4 | 0.0162109 | membrane |
| **0.314972** | 0.453072 | GOCC name | 31 | 0.00104785 | 0.0223882 | dendritic spine |
| **0.412012** | 0.49897 | GOCC name | 21 | 0.00169683 | 0.0353909 | presynaptic membrane |
| **0.371487** | 0.0847756 | GOCC name | 47 | 0.00196199 | 0.0399699 | transmembrane transporter complex |
| **0.857454** | 0.522799 | GOCC name | 8 | 0.00198266 | 0.039473 | cytochrome complex |
| **0.505115** | 0.799118 | GOCC name | 9 | 0.00209929 | 0.0408661 | astrocyte projection |
| **0.0689594** | 0.250141 | GOCC name | 107 | 0.00227134 | 0.0432543 | cell-cell junction |
| **0.260989** | 0.279857 | GOCC name | 59 | 0.00228683 | 0.0426227 | axon |
| **0.018177** | 0.209969 | GOCC name | 147 | 0.00229201 | 0.0418292 | anchoring junction |
| **0.118314** | 0.283841 | GOCC name | 80 | 0.00243123 | 0.0434645 | cell body |

| Strata |
| --- |
| SP |
| SP |
| SP |
| SP |
| SP |
| SP |
| SP |
| SP |
| SP |
| SP |
| SP |
| SP |
| SP |
| SP |
| SP |
| SP |
| SP |
| SP |
| SP |
| SP |
| SP |
| SP |

| |
|---|
| SP |
| SP |
| SP |
| SP |
| SP |
| SP |
| SP |
| SP |
| SP |
| SP |
| SP |
| SP |
| SP |
| SP |
| SP |
| SP |
| SP |
| SP |
| SP |
| SP |
| SP |
| SP |
| SP |

| |
|---|
| SP |
| SP |
| SP |
| SP |
| SP |
| SP |
| SP |
| SP |
| SP |
| SP |
| SP |
| SP |
| SP |
| SP |
| SP |
| SP |
| SP |
| SP |
| SP |
| SP |
| SP |
| SP |
| SP |

| |
|---|
| SP |
| SP |
| SP |
| SP |
| SP |
| SP |
| SP |
| SP |
| SP |
| SP |
| SP |
| SP |
| SP |
| SP |
| SP |
| SP |
| SP |
| SP |
| SP |
| SP |
| SP |
| SP |
| SP |

| |
|---|
| SP |
| SP |
| SP |
| SP |
| SP |
| SP |
| SP |
| SP |
| SP |
| SP |
| SP |
| SP |
| SP |
| SP |
| SP |
| SP |
| SP |
| SP |
| SP |
| SP |
| SP |
| SP |
| SP |

| |
|---|
| SP |
| SP |
| SP |
| SP |
| SP |
| SP |
| SP |
| SP |
| SP |
| SP |
| SP |
| SP |
| SP |
| SP |
| SP |
| SP |
| SP |
| SP |
| SP |
| SP |
| SP |
| SP |
| SP |

| |
|---|
| SP |
| SP |
| SP |
| SP |
| SP |
| SP |
| SP |
| SP |
| SP |
| SP |
| SP |
| SP |
| SP |
| SP |
| SP |
| SP |
| SP |
| SP |
| SP |
| SP |
| SP |
| SP |
| SP |

| |
|---|
| SP |
| SP |
| SP |
| SP |
| SP |
| SP |
| SP |
| SP |
| SP |
| SP |
| SP |
| SP |
| SP |
| SP |
| SP |
| SP |
| SP |
| SP |
| SP |
| SP |
| SP |
| SP |
| SP |

| |
|---|
| SP |
| SP |
| SP |
| SP |
| SP |
| SP |
| SP |
| SP |
| SP |
| SP |
| SP |
| SP |
| SP |
| SP |
| SP |
| SP |
| SP |
| SP |
| SP |
| SP |
| SP |
| SP |
| SP |

| |
|---|
| SP |
| SP |
| SP |
| SP |
| SP |
| SP |
| SP |
| SP |
| SP |
| SP |
| SP |
| SP |
| SP |
| SP |
| SP |
| SP |
| SP |
| SP |
| SP |
| SP |
| SP |
| SP |
| SP |

| |
|---|
| SP |
| SP |
| SP |
| SP |
| SP |
| SP |
| SP |
| SP |
| SP |
| SP |
| SP |
| SP |
| SP |
| SP |
| SP |
| SP |
| SP |
| SR |
| SR |
| SLM |
| SLM |
| SLM |
| SLM |

| |
|---|
| SLM |
| SLM |
| SLM |
| SLM |
| SLM |
| SLM |
| SLM |
| SLM |
| SLM |
| SLM |
| SLM |
| SLM |
| SLM |
| SLM |
| SLM |
| SLM |
| SLM |
| SLM |
| SLM |
| SLM |
| SLM |
| SLM |
| SLM |

| |
|---|
| SLM |
| SLM |
| SLM |
| SLM |
| SLM |
| SLM |
| SLM |
| SLM |
| SLM |
| SLM |
| SLM |
| SLM |
| SLM |
| SLM |
| SLM |
| SLM |
| SLM |
| SLM |
| SLM |
| SLM |
| SLM |
| SLM |
| SLM |

| |
|---|
| SLM |
| SLM |
| SLM |
| SLM |
| SLM |
| SLM |
| SLM |
| SLM |
| SLM |
| SLM |
| SLM |
| SLM |
| SLM |
| SLM |
| SLM |
| SLM |
| SLM |
| SLM |
| SLM |
| SLM |
| SLM |
| SLM |
| SLM |

| |
|---|
| SLM |
| SLM |
| SLM |
| SLM |
| SLM |
| SLM |
| SLM |
| SLM |
| SLM |
| SLM |
| SLM |
| SLM |
| SLM |
| SLM |
| SLM |
| SLM |
| SLM |
| SLM |
| SLM |
| SLM |
| SLM |
| SLM |
| SLM |

| |
|---|
| SLM |
| SLM |
| SLM |
| SLM |
| SLM |
| SLM |
| SLM |
| SLM |
| SLM |
| SLM |
| SLM |
| SLM |
| SLM |
| SLM |
| SLM |
| SLM |
| SLM |
| SLM |
| SLM |
| SLM |
| SLM |
| SLM |
| SLM |

| |
|---|
| SLM |
| SLM |
| SLM |
| SLM |
| SLM |
| SLM |
| SLM |
| SLM |
| SLM |
| SLM |
| SLM |
| SLM |
| SLM |
| SLM |
| SLM |
| SLM |
| SLM |
| SLM |
| SLM |
| SLM |
| SLM |
| SLM |
| SLM |

| |
|---|
| SLM |
| SLM |
| SLM |
| SLM |
| SLM |
| SLM |
| SLM |
| SLM |
| SLM |
| SLM |
